# Population genomics of postglacial western eurasia

**DOI:** 10.1101/2022.05.04.490594

**Authors:** Morten E. Allentoft, Martin Sikora, Alba Refoyo-Martínez, Evan K. Irving-Pease, Anders Fischer, William Barrie, Andrés Ingason, Jesper Stenderup, Karl-Göran Sjögren, Alice Pearson, Bárbara Sousa da Mota, Bettina Schulz Paulsson, Alma Halgren, Ruairidh Macleod, Marie Louise Schjellerup Jørkov, Fabrice Demeter, Lasse Sørensen, Poul Otto Nielsen, Rasmus A. Henriksen, Tharsika Vimala, Hugh McColl, Ashot Margaryan, Melissa Ilardo, Andrew Vaughn, Morten Fischer Mortensen, Anne Birgitte Nielsen, Mikkel Ulfeldt Hede, Niels Nørkjær Johannsen, Peter Rasmussen, Lasse Vinner, Gabriel Renaud, Aaron Stern, Theis Zetner Trolle Jensen, Gabriele Scorrano, Hannes Schroeder, Per Lysdahl, Abigail Daisy Ramsøe, Andrei Skorobogatov, Andrew Joseph Schork, Anders Rosengren, Anthony Ruter, Alan Outram, Aleksey A. Timoshenko, Alexandra Buzhilova, Alfredo Coppa, Alisa Zubova, Ana Maria Silva, Anders J. Hansen, Andrey Gromov, Andrey Logvin, Anne Birgitte Gotfredsen, Bjarne Henning Nielsen, Borja González-Rabanal, Carles Lalueza-Fox, Catriona J. McKenzie, Charleen Gaunitz, Concepción Blasco, Corina Liesau, Cristina Martinez-Labarga, Dmitri V. Pozdnyakov, David Cuenca-Solana, David O. Lordkipanidze, Dmitri En’shin, Domingo C. Salazar-García, T. Douglas Price, Dušan Borić, Elena Kostyleva, Elizaveta V. Veselovskaya, Emma R. Usmanova, Enrico Cappellini, Erik Brinch Petersen, Esben Kannegaard, Francesca Radina, Fulya Eylem Yediay, Henri Duday, Igor Gutiérrez-Zugasti, Ilya Merts, Inna Potekhina, Irina Shevnina, Isin Altinkaya, Jean Guilaine, Jesper Hansen, Joan Emili Aura Tortosa, João Zilhão, Jorge Vega, Kristoffer Buck Pedersen, Krzysztof Tunia, Lei Zhao, Liudmila N. Mylnikova, Lars Larsson, Laure Metz, Levon Yepiskoposyan, Lisbeth Pedersen, Lucia Sarti, Ludovic Orlando, Ludovic Slimak, Lutz Klassen, Malou Blank, Manuel González-Morales, Mara Silvestrini, Maria Vretemark, Marina S. Nesterova, Marina Rykun, Mario Federico Rolfo, Marzena Szmyt, Marcin Przybyła, Mauro Calattini, Mikhail Sablin, Miluše Dobisíková, Morten Meldgaard, Morten Johansen, Natalia Berezina, Nick Card, Nikolai A. Saveliev, Olga Poshekhonova, Olga Rickards, Olga V. Lozovskaya, Olivér Gábor, Otto Christian Uldum, Paola Aurino, Pavel Kosintsev, Patrice Courtaud, Patricia Ríos, Peder Mortensen, Per Lotz, Per Persson, Pernille Bangsgaard, Peter de Barros Damgaard, Peter Vang Petersen, Pilar Prieto Martinez, Piotr Włodarczak, Roman V. Smolyaninov, Rikke Maring, Roberto Menduiña, Ruben Badalyan, Rune Iversen, Ruslan Turin, Sergey Vasilyev, Sidsel Wåhlin, Svetlana Borutskaya, Svetlana Skochina, Søren Anker Sørensen, Søren H. Andersen, Thomas Jørgensen, Yuri B. Serikov, Vyacheslav I. Molodin, Vaclav Smrcka, Victor Merz, Vivek Appadurai, Vyacheslav Moiseyev, Yvonne Magnusson, Kurt H. Kjær, Niels Lynnerup, Daniel J. Lawson, Peter H. Sudmant, Simon Rasmussen, Thorfinn Korneliussen, Richard Durbin, Rasmus Nielsen, Olivier Delaneau, Thomas Werge, Fernando Racimo, Kristian Kristiansen, Eske Willerslev

## Abstract

Western Eurasia witnessed several large-scale human migrations during the Holocene^1–5^. To investigate the cross-continental impacts we shotgun-sequenced 317 primarily Mesolithic and Neolithic genomes from across Northern and Western Eurasia. These were imputed alongside published data to obtain diploid genotypes from >1,600 ancient humans. Our analyses revealed a ‘Great Divide’ genomic boundary extending from the Black Sea to the Baltic. Mesolithic hunter-gatherers (HGs) were highly genetically differentiated east and west of this zone, and the impact of the neolithisation was equally disparate. Large-scale ancestry shifts occurred in the west as farming was introduced, including near-total replacements of HGs in many areas, whereas no substantial ancestry shifts happened east of the zone during the same period. Similarly, relatedness decreased in the west from the Neolithic transition onwards, while east of the Urals relatedness remained high until ∼4,000 BP, consistent with persistence of localised HG groups. The boundary dissolved when Yamnaya-related ancestry spread across western Eurasia around 5,000 BP resulting in a second major turnover that reached most parts of Europe within a 1,000-year span. The genetic origin and fate of the Yamnaya have remained elusive but we demonstrate that HGs from the Middle Don region contributed ancestry to them. Yamnaya-groups later admixed with individuals associated with the Globular Amphora Culture before expanding into Europe. Similar turnovers occurred in western Siberia, where we report new genomic data from a ‘Neolithic steppe’ cline spanning the Siberian forest steppe to Lake Baikal. These prehistoric migrations had profound and lasting effects on the genetic diversity of Eurasian populations.

## Introduction

Genetic diversity in West Eurasian human populations was largely shaped by three major prehistoric migrations: anatomically modern hunter-gatherers (HGs) occupying the area from c. 45,000 BP^4,6^; Neolithic farmers expanding from the Middle East from c. 11,000 BP^4^; and steppe pastoralists coming out of the Pontic Steppe c. 5,000 BP^1,2^. Palaeogenomic analyses have uncovered the early post-glacial colonisation routes^7^ resulting in a basal ancestral dichotomy between HGs in central/western Europe and HG groups represented further east^8^. Western HG (WHG) ancestry appears to be derived directly from ancestry sources related to Epigravettian, Azilian and Epipalaeolithic cultures (the ‘Villabruna Cluster’)^9^, while Eastern HG (EHG) ancestry shows additional admixture with an Upper Palaeolithic (UP) Siberian source (‘Ancient North Eurasian’, ANE)^10^. The WHG ancestry composition was regionally variable in the Mesolithic populations. There is evidence for continuous local admixture in Iberian HGs^11^, contrasting with a more homogenous WHG ancestry profile in Britain and northwestern continental Europe, suggesting ancestry formation prior to expansion^12^. The timing of the ancestry admixture that formed EHG has been estimated at 13-15,000 BP and the composition seem to follow a cline broadly correlated with geography: Baltic and Ukrainian HGs showing more affinity to the Villabruna UP cluster ancestry, compared to HGs in Russia who displayed more ANE^5,7,13,14^. Genomic analyses of Mesolithic skeletal material from the Scandinavian Peninsula has revealed varied mixes of WHG and EHG ancestry among the later Mesolithic populations^3,15,16^. Beyond these broad scale characterizations our knowledge on Mesolithic population structure and demographic admixture processes is limited and with significant chronological and geographic information gaps. This is partly owing to a relative paucity of well-preserved Mesolithic human skeletons older than 8,000 years and partly because most ancient DNA (aDNA) studies on the Mesolithic and Neolithic periods have been restricted to individuals from Europe. The archaeological record indicates a boundary from the eastern Baltic to the Black Sea, east of which HG societies persisted for much longer than in western Europe despite similar distance to the distribution centre for early agriculture in the Middle East^17^. Components of eastern and western HG ancestry appear highly variable in this boundary region^5,18,19^ but the wider spatiotemporal genetic implications of the east-west division is unclear. The spatiotemporal mapping of population dynamics east of Europe, including Northern and Central Asia during the same time period is limited. In these regions the term ‘Neolithic’ is characterised by cultural and economic changes including societal network differences, changes in lithic technology and use of pottery. For instance, the Neolithic cultures of the Central Asian steppe and the Russian taiga belt possessed pottery, but retained a HG economy alongside stone blade technology similar to the preceding Mesolithic cultures^20^. A fundamental lack of data from some key regions and periods has prohibited a deeper understanding of how the neolithisation differed in timing, mechanisms, and impact across Northern and Western Eurasia.

The transition from hunting and gathering to farming was based on domesticated plants and animals of Middle Eastern origin, and represents one of the most fundamental shifts in demography, health, lifestyle and culture in human prehistory. The neolithisation process in large parts of Europe was accompanied by the arrival of immigrants of Anatolian descent^21^. For example, in Iberia the Neolithic began with the abrupt spread of immigrant farmers of Anatolian-Aegean ancestry along the Mediterranean and Atlantic coasts, after which admixture with local HGs gradually took place^11^. Similarly, in Southeast and Central Europe farming rapidly spread with Anatolian Neolithic farmers, who were to some extent subsequently admixed with local HGs^22–27^. Conversely, in Britain data suggest a complete replacement of the HG population when agriculture was introduced by incoming continental farmers and without a subsequent resurgence of local HG ancestry^12,28^. In the East Baltic region a markedly different neolithisation trajectory occurred, with introduction of domesticates only at the emergence of the Corded Ware Complex around 4,800 cal. BP^18,19^. Similarly, in eastern Ukraine, HGs of Mesolithic ancestry co-existed for millennia with farming groups further west^5,29^. These recent studies have all provided important regional contributions towards understanding West Eurasian population history, but from a broader cross-continental perspective, our knowledge is still patchy. A fundamental lack of data from some key regions and periods has prohibited a deeper understanding of how neolithisation differed in timing, mechanisms, and impact across Northern and Western Eurasia.

From approximately 5,000 BP, an ancestry component related to Early Bronze Age steppe pastoralists such as the Yamnaya Culture rapidly spread across Europe through the expansion of the Corded Ware Complex (CWC) and related cultures^1,2^. Although previous studies have identified these large-scale migrations into Europe and Central Asia, central aspects concerning the demographic processes are not resolved. Yamnaya ancestry (i.e. ‘steppe’ ancestry) has been characterised broadly as a mix between EHG ancestry and Caucasus Hunter-Gatherer (CHG), formed in a hypothetical admixture between a ‘Northern’ steppe source and a ‘Southern’ Caucasus source^30^. However, the exact origins of these ancestry sources have not been identified.

Furthermore, with a few exceptions^31–33^ published Yamnaya Y-chromosomal haplogroups do not match those found in post 5,000 BP Europeans and the origin of this patrilineal lineage is also unresolved. Finally, in Europe ‘steppe’ ancestry has hitherto only been identified in admixed form, but the origin of this admixture event and the mechanism by which the ancestry subsequently spread with the CWC have remained elusive.

To investigate these formative processes at a cross-continental scale, we sequenced the genomes of 317 radiocarbon-dated (AMS) individuals of primarily Mesolithic and Neolithic origin, covering major parts of Eurasia. We combined these with published shotgun-sequenced data to impute a dataset of >1600 diploid ancient genomes. Of the 317 sampled ancient skeletons (Fig. 1, Extended Data Fig. 1, Supplementary Data I) 272 were radiocarbon-dated within the project, while 30 dates were derived from published literature, and 15 were dated by archaeological context. Dates were corrected for marine and freshwater reservoir effects (Supplementary Note 4) and ranged from the UP c. 25,700 calibrated years before present (cal. BP) to the mediaeval period (c. 1200 cal. BP). However, 97% of the individuals (N=309) date to between 11,000 and 3,000 cal. BP, with a heavy focus on individuals associated with various Mesolithic and Neolithic cultures. Geographically, the 317 sampled skeletons cover a vast territory across Eurasia, from Lake Baikal to the Atlantic coast and from Scandinavia to the Middle East, deriving from contexts that include burial mounds, caves, bogs and the seafloor (Supplementary Notes 6-7). Broadly, we can divide our research area into three large regions: 1) central, western and northern Europe, 2) eastern Europe, including western Russia, Belarus and Ukraine, and 3) the Urals and western Siberia (Supplementary Notes 6-7). Samples cover many of the key Mesolithic and Neolithic cultures in Western Eurasia, such as the Maglemose, Ertebølle, Funnel Beaker (TRB) and Corded Ware/Single Grave Cultures in Scandinavia, the Cardial in the Mediterranean, the Körös and Linear Pottery (LBK) in SE and Central Europe, and many archaeological cultures in the Ukraine, western Russia, and the trans-Ural (e.g. Veretye, Lyalovo, Volosovo, Kitoi). Our sampling was particularly dense in Denmark, from where we present a detailed and continuous sequence of 100 genomes spanning the Early Mesolithic to the Bronze Age (*Allentoft, Sikora, Fischer et al. submitted*\*^1^). Dense sampling was also obtained from Ukraine, Western Russia, and the trans-Ural, spanning from the Early Mesolithic through the Neolithic, up to c. 5,000 BP.

**Fig. 1:**
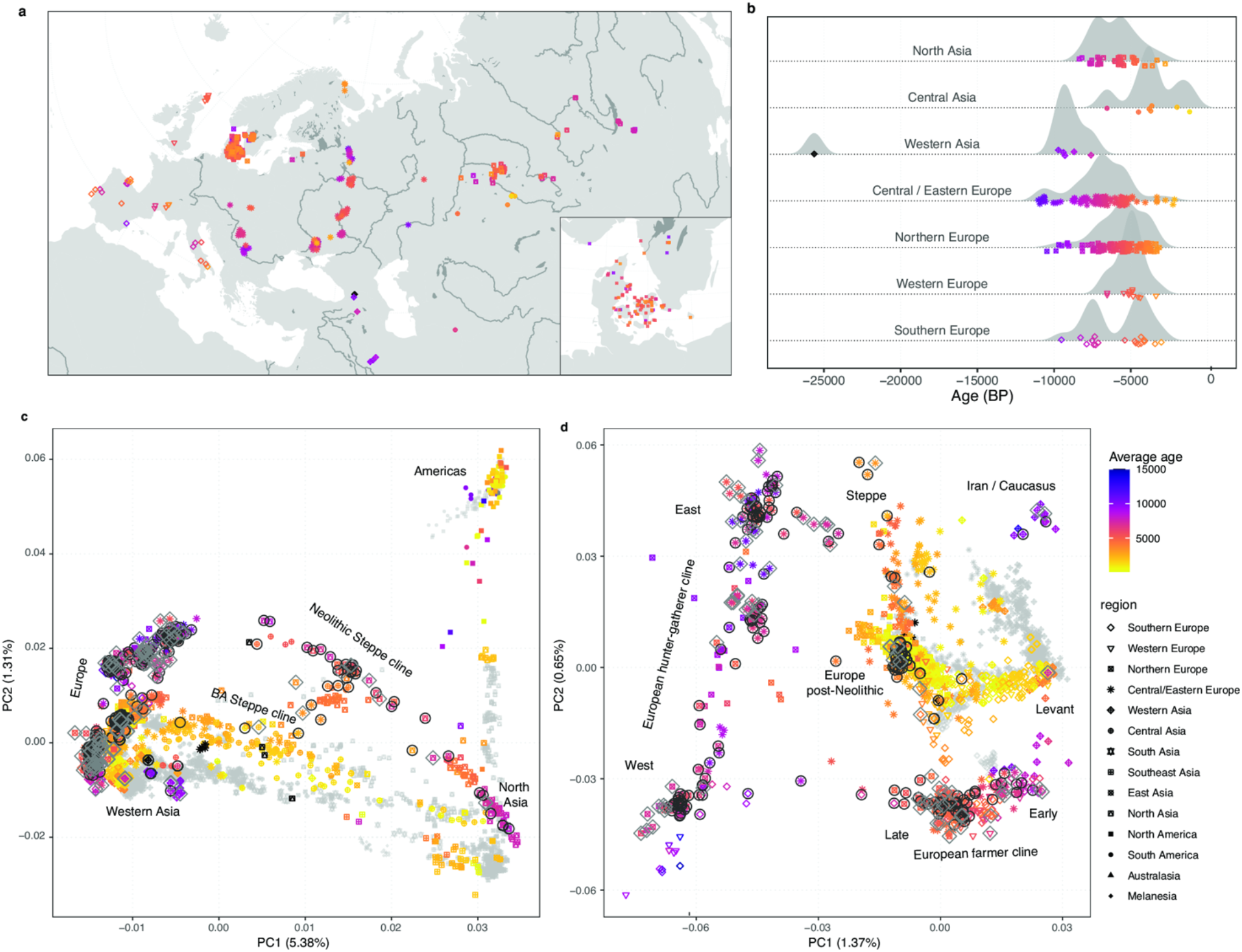
Sample overview and broad scale genetic structure. (A), (B) Geographic and temporal distribution of the 317 ancient genomes sequenced and reported in this study. Insert shows dense sampling in Denmark (see *Allentoft, Sikora, Fischer et al*., *submitted*). Age and geographic region of ancient individuals are indicated by plot symbol colour and shape, respectively. Colour scale for age is capped at 15,000 years, older individuals are indicated with black colour. Random jitter was added to geographic coordinates to avoid overplotting. (C), (D) Principal component analysis of 3,316 modern and ancient individuals from Eurasia, Oceania, and the Americas (C), and restricted to 2,126 individuals from western Eurasia (west of the Urals) (D). Principal components were defined using both modern and imputed ancient (n=1492) genomes passing all filters, with the remaining low-coverage ancient genomes projected. Ancient genomes sequenced in this study are indicated with black circles (imputed genomes passing all filters, n=213) or grey diamonds (pseudo-haploid projected genomes, n=104). Genomes of modern individuals are shown in grey, with population labels corresponding to their median coordinates.

**Fig. 2:**
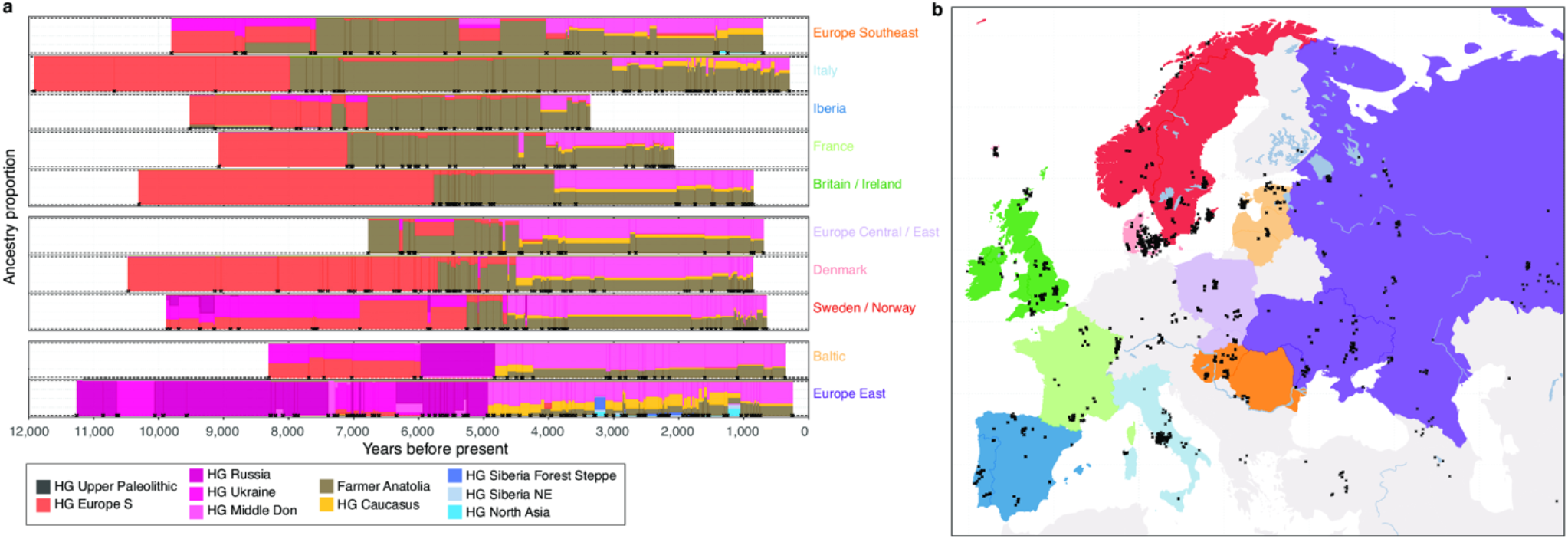
Genetic ancestry transects of Western Eurasia. (a) Regional timelines of genetic ancestry compositions within the past 12,000 years in western Eurasia. Ancestry proportions in 1,012 imputed ancient genomes (representing populations west of the Urals) inferred using supervised ancestry modelling with the “deep” HG ancestry source groups. Coloured bars within the timelines represent ancestry proportions for temporally consecutive individuals, with the width corresponding to their age difference. Individuals with identical age were offset along the time axis by adding random jitter. (b) Map highlighting geographic areas (coloured areas) for samples included in the individual regional timelines, and excavation locations (black crosses). Only shotgun sequenced genomes were used in our study, why the exact timing of ancestry shifts may differ slightly from previous studies if based on different types of data from different individuals.

## Results and Discussion

### Broad scale genetic structure

Ancient DNA was extracted from either dental cementum or petrous bones and the 317 genomes were shotgun sequenced to a depth of coverage ranging between 0.01X and 7.1X (mean = 0.75X, median = 0.26X), with >1X coverage for 81 genomes (Supplementary Note 1). We utilised a new computational method optimised for low-coverage data^34^, to impute genotypes using the 1000 Genomes phased data^35^ as a reference panel. This was jointly applied to >1300 previously published shotgun-sequenced genomes (Supplementary Data VII), resulting in a dataset of 8.5 million common SNPs (>1% Minor Allele Frequency (MAF) and imputation info score > 0.5) for 1,664 imputed diploid ancient genomes (Extended Data Fig. 2). For most downstream analyses n=71 individuals were excluded as close relatives or with a contamination estimate >5%, resulting in 1,593 genomes of which 1,492 were analysed as imputed (213 sequenced in this study) while n=101 were analysed as pseudo-haploids due to low coverage (<0.1X), and/or low imputation quality (average genotype probability < 0.98).

We conducted a broad-scale characterization of this dataset using principal component analysis (PCA) and model-based clustering (ADMIXTURE), which recapitulated previously described ancestry clines in ancient Eurasian populations at increased resolution (Fig. 1; Extended Data Fig. 1; Supplementary Note 3d). Our imputed whole genomes allowed us to perform principal component analysis using ancient genomes as input, instead of projecting onto a space defined by modern variation. Strikingly, this resulted in much higher differentiation among the ancient individuals than observed previously (Extended Data Fig. 1). This is particularly notable in a PCA of West Eurasian individuals, where the variance explained by the first two PCs increases ∼1.5 fold, and present-day populations are confined within a small central area of the PCA space (Fig. 1d; Extended Data Fig. 1c,d). These results are consistent with higher genetic differentiation between ancient Europeans than present-day populations, reflecting more genetic isolation and lower effective population sizes among ancient groups.

To obtain a finer-scale characterization of genetic ancestries across space and time, we employed an approach akin to the widely used CHROMOPAINTER/FINESTRUCTURE workflow^36–38^. We first performed community detection on a network constructed from pairwise identity-by-descent (IBD)-sharing similarities between ancient individuals to group them into hierarchically related clusters of similar genetic ancestry (Extended Data Fig. 3; Supplementary Note 3c). At higher levels of the hierarchy, the resulting clusters represented previously described ancestry groups reflecting broad genetic structure, such as eastern and western European hunter-gatherers (“HG_EuropeE”, “HG_EuropeW”; Extended Data Fig. 3). Clusters at the lowest level resolved fine-scale genetic structure, grouping individuals within restricted spatiotemporal ranges and/or archaeological contexts but also revealing previously unknown connections across broader geographical areas (Extended Data Fig. 3; Supplementary Note 3f). These resulting clusters were subsequently used in supervised ancestry modelling where sets of ‘target’ individuals were modelled as mixtures of ‘source’ groups (Methods).

### Post-LGM Population structure of HGs

Our study comprises 113 shotgun sequenced and imputed HG genomes of which 79 were sequenced in this study. Among them, we report a 0.83X genome of an UP skeleton from Kotias Klde Cave in Georgia, Caucasus (NEO283), directly dated to 26,052 - 25,323 cal. BP (95%). In the PCA of all non-African individuals, it occupied a position distinct from other previously sequenced UP individuals, shifted towards west Eurasians along PC1 (Supplementary Note 3d). Using admixture graph modelling, we find that a well-fitting graph for this Caucasus UP lineage derives it as a mixture of predominantly West Eurasian UP HG ancestry (76%) with ∼24% contribution from a ‘basal Eurasian’ ghost population, first observed in West Asian Neolithic individuals^4^ (Supplementary Note 3d, Fig. S3d.16). To further explore the fine-scale structure of later European HGs, we then performed supervised ancestry modelling using sets of increasingly proximate source clusters (Extended Data Fig. 4). We replicate previous results of broad-scale genetic differentiation between HGs in eastern and western Europe after the Last Glacial Maximum (LGM)^5,7^. We show that the deep ancestry divisions in the Eurasian human gene pool that were established during early post-LGM dispersals^7^ persisted throughout the Mesolithic (Extended Data Fig. 4). Using distal sets of pre-LGM HGs as sources, western HGs were modelled as predominantly derived from a source related to the herein reported Caucasus UP individual from Kotias Klde cave (Caucasus_25000BP), whereas eastern HGs showed varying amounts of ancestry related to a Siberian HG from Mal’ta (Malta_24000BP; Extended Data Fig. 4a; Supplementary Data XII). Using post-LGM sources, this divide is best represented by ancestry related to southern European (Italy_15000BP_9000 BP) and Russian (RussiaNW_11000BP_8000BP) HGs, respectively, corresponding to the ‘WHG’ and ‘EHG’ labels commonly used in previous studies.

Adding additional proximate sources allowed us to further refine the ancestry composition of Northern European HGs. In Denmark, our 28 sequenced and imputed HG genomes derived almost exclusively from a southern European source (Italy_15000BP_9000), with remarkable homogeneity across a 5,000 year transect (Extended Data Fig. 4a; Supplementary Data XII) (*Allentoft, Sikora, Fischer et al. submitted*). In contrast, we observed marked geographic variation in ancestry composition of HGs from other parts of Scandinavia. Mesolithic individuals from Scandinavia were broadly modelled as mixtures with varying proportions of eastern and western HGs using distal post-LGM sources (“hgEur1”, Extended Data Fig. 4a), as previously reported^15^. In Mesolithic individuals from southern Sweden, the eastern HG ancestry component was largely replaced by a southeast European source (Romania_8800BP) in more proximate models, making up between 60%-70% of the ancestry (Extended Data Fig. 4a; Supplementary Data XII). Ancestry related to Russian HGs increased in a cline towards the far north, peaking at ∼75% in a late HG from Tromso (VK531; ∼4,350BP) (Extended Data Fig. 4a,c; Supplementary Data XII), also reflected in highest IBD sharing of those individuals with Northern Russian HGs (Extended Data Fig. 4d). During the late Mesolithic, we observed higher southern European HG ancestry in coastal individuals (NEO260 from Evensås; NEO679 from Skateholm) than in earlier individuals from further inland. Adding Danish HGs as proximate source substantially improved the fit for those two individuals (“hgEur3”, Extended Data Fig. 4b), with estimated 58%-76% ancestry derived from Danish HGs (“hgEur3”, Extended Data Fig.7a; Supplementary Data XII), suggesting a population genetic link with Denmark where this ancestry prevailed (Extended Data Fig. 4c). These results indicate at least three distinct waves of northwards HG ancestry into Scandinavia: (i) a predominantly southern European source into Denmark and coastal southwestern Sweden; (ii) a source related to southeastern European HGs into the Baltic and southeastern Sweden; and (iii) a northwest Russian source into the far north, which then spread south along the Atlantic coast of Norway^15^ (Extended Data Fig. 4c). These movements likely represent post glacial expansions from refugia areas shared with many plant and animal species^39^.

On the Iberian Peninsula, the earliest individuals, including a ∼9,200-year-old HG (NEO694) from Santa Maira (eastern Spain), sequenced in this study, showed predominantly southern European HG ancestry with a minor contribution from UP HG sources (Extended Data Fig. 4a). This observed UP HG ancestry source mix likely reflects the pre-LGM Magdalenian-related ancestry component previously reported in Iberian HGs^11^, for which a good source population proxy is lacking in our dataset. In contrast, later individuals from Northern Iberia were more similar to HGs from southeastern Europe, deriving ∼30-40% of their ancestry from a source related to HGs from the Balkans in more proximate models^11,40^ (Extended Data Fig. 4a; Supplementary Data XII). The earliest evidence for this gene flow was observed in a Mesolithic individual from El Mazo, Spain (NEO646) that was dated, calibrated and reservoir-corrected to c. 8,200 BP (8365-8182 cal. BP, 95%) but dated slightly earlier by context (8550-8330 BP^41^). The directly dated age coincides with some of the oldest Mesolithic geometric microliths in northern Iberia, appearing around 8,200 BP at this site^41^. An influx of southeastern European HG-related ancestry in Ukrainian individuals after the Mesolithic (Extended Data Fig. 4a; Supplementary Data XII) suggests a similar eastwards expansion in south-eastern Europe^5^. Interestingly, two newly reported ∼7,300-year-old genomes from the Middle Don River region in the Pontic-Caspian steppe (Golubaya Krinitsa, NEO113 & NEO212) were found to be predominantly derived from earlier Ukrainian HGs, but with ∼18-24% of their ancestry contributed from a source related to HGs from the Caucasus (Caucasus_13000BP_10000BP) (Extended Data Fig. 4a; Supplementary Data XII). Additional lower coverage (non-imputed) genomes from the same site project in the same PCA space (Fig. 1d), shifted away from the European HG cline towards Iran and the Caucasus. Using the linkage-disequilibrium-based method DATES^42^ we dated this admixture to ∼8,300 BP (Supplementary Data XIV). These results document genetic contact between populations from the Caucasus and the steppe region much earlier than previously known, evidencing admixture prior to the advent of later nomadic steppe cultures, in contrast to recent hypotheses, and further to the west than previously reported^5,43^.

### Major genetic transitions in Europe

Previous ancient genomics studies have documented multiple episodes of large-scale population turnover in Europe within the last 10,000 years^e.g. 1,2,5,44^ but the 317 novel genomes reported here fill important knowledge gaps. Our analyses reveal profound differences in the spatiotemporal neolithisation dynamics across Europe. Supervised admixture modelling (using the ‘deep’ ancestry set; Supplementary Data IX) and spatiotemporal kriging^45^ document a broad east-west distinction along a boundary zone running from the Black Sea to the Baltic. On the western side of this ‘Great Divide’, the Neolithic transition is accompanied by large-scale shifts in genetic ancestry from local HGs to farmers with Anatolian-related ancestry (Boncuklu_10000BP, Fig. 3a, Fig. 4; Extended Data Figs. 5-7). The arrival of Anatolian-related ancestry in different regions spans an extensive time period of over 3,000 years, from its earliest evidence in the Balkans (Lepenski Vir) at ∼8,700 BP^5^ to c. 5,900 BP in Denmark.

**Fig. 3:**
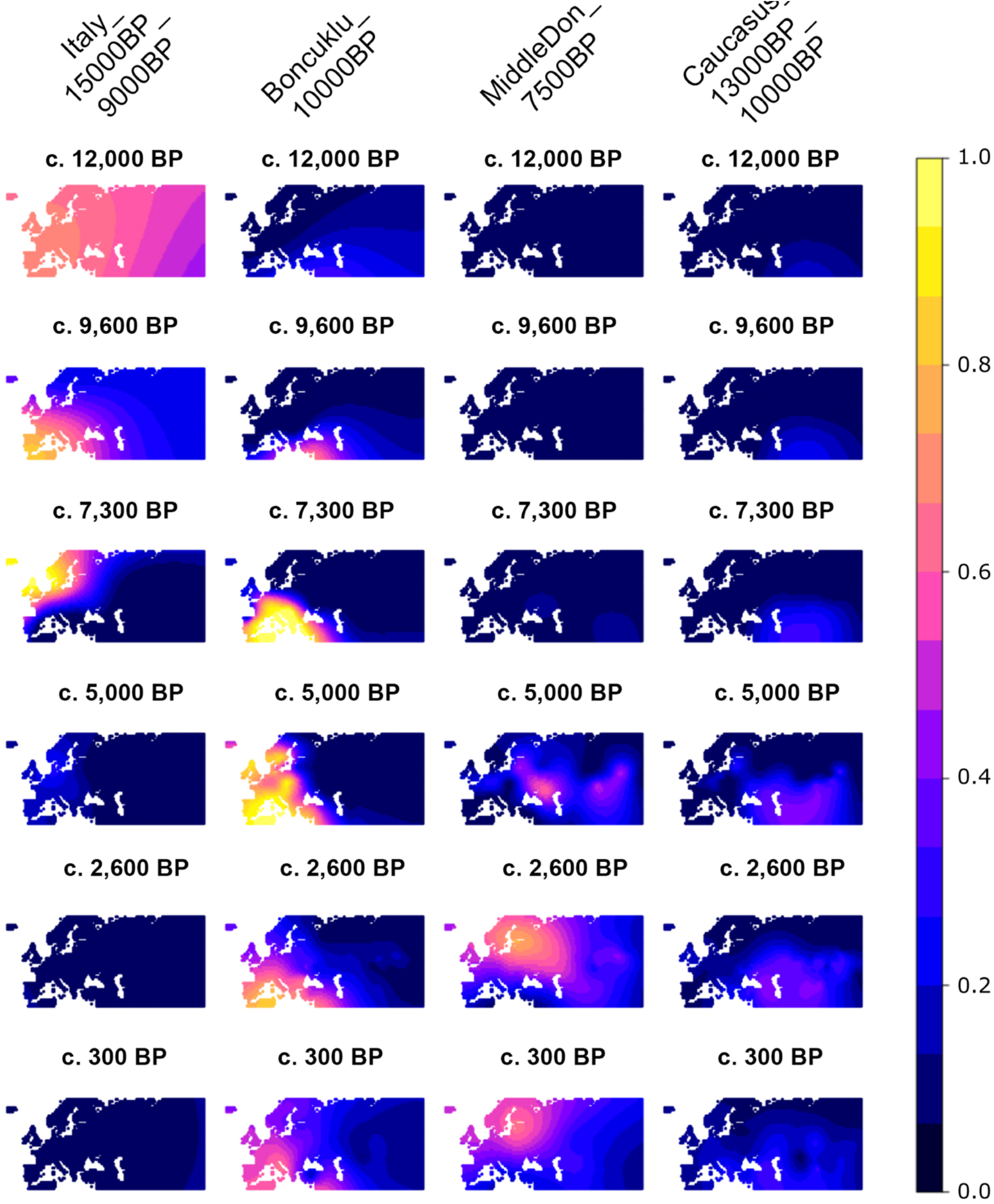
Spatiotemporal kriging analysis of major ancestries. The temporal transects demonstrate how WHG ancestry (Italy_15000BP_9000BP), was replaced by Neolithic farmer ancestry (Boncuklu_10000BP) during the Neolithic transition in Europe. Later, the steppe migrations around 5,000 cal. BP introduce both EHG (MiddleDon_7500BP) and CHG (Caucasus_13000BP_10000BP) ancestry into Europe, thereby reducing Neolithic farmer ancestry.

Further, we corroborate previous reports^e.g. 2,5,44,46^ of widespread, low-level admixture between early European farmers and local HGs resulting in a resurgence of HG ancestry in many regions of Europe during subsequent centuries (Extended Data Fig. 8b,c; Supplementary Data XIII). The resulting estimated HG ancestry proportions rarely exceeded 10%, with notable exceptions observed in individuals from south-eastern Europe (Iron Gates), Sweden (Pitted Ware Culture) as well as herein reported Early Neolithic genomes from Portugal (western Cardial), which are estimated to harbour 27% – 43% Iberian HG ancestry (Iberia_9000BP_7000BP). The latter result, together with an estimated admixture date of just ∼200 years earlier (Supplementary Data XIV) suggests extensive first-contact admixture, and is in agreement with archaeological inferences derived from modelling the spread of farming along west Mediterranean Europe^47^. Neolithic individuals from Denmark showed some of the highest overall HG ancestry proportions (up to ∼25%), but mostly derived from non-local Western European-related HGs (EuropeW_13500BP_8000BP), with only a small contribution from local Danish HG groups in some individuals (Extended Data Fig. 8b; Supplementary Note 3f).

We find evidence for regional stratification in early Neolithic farmer ancestries in subsequent Neolithic groups. Specifically, southern European early farmers were found to have provided major genetic ancestry to Neolithic groups of later dates in Western Europe, while central European early farmer ancestry was mainly observed in subsequent Neolithic groups in eastern Europe and Scandinavia (Extended Data Fig. 8e). These results are consistent with distinct migratory routes of expanding farmer populations as previously suggested^48^.

On the eastern side of the ‘Great Divide’ no ancestry shifts can be observed during this period. In the East Baltic region (see also^49^), the Ukraine and Western Russia, local HG ancestry prevailed until ∼5,000 BP without noticeable input of Anatolian-related farmer ancestry (Fig. 3-4; Extended Data Figs. 5-7). This Eastern genetic continuity is in congruence with the archaeological record, which shows persistence of pottery-using forager groups in this wide region, and a delayed introduction of cultivation and animal husbandry by several thousand years (Supplementary Note 5). Around 5,000 BP major demographic events unfolded on the Eurasian Steppe resulting in steppe-related ancestry spreading rapidly both eastwards and westwards^1,2^, marking the end of the great population genomic divide (Fig. 4, Fig. 8). We find that this second transition happened at a faster pace than during the neolithisation, reaching most parts of Europe within a ∼1,000-year time period after first appearing in the eastern Baltic region ∼4,800 cal. BP (Fig. 3). In line with previous reports we observe that by c. 4,200 cal. BP, steppe-related ancestry was already dominant in individuals from Britain, France and the Iberian Peninsula^12,50^. Strikingly, because of the delayed neolithisation in southern Scandinavia these dynamics resulted in two episodes of large-scale genetic turnover in Denmark and southern Sweden within roughly a 1,000-year period (Fig. 3) (*Allentoft, Sikora, Fischer et al. submitted*).

**Fig. 4:**
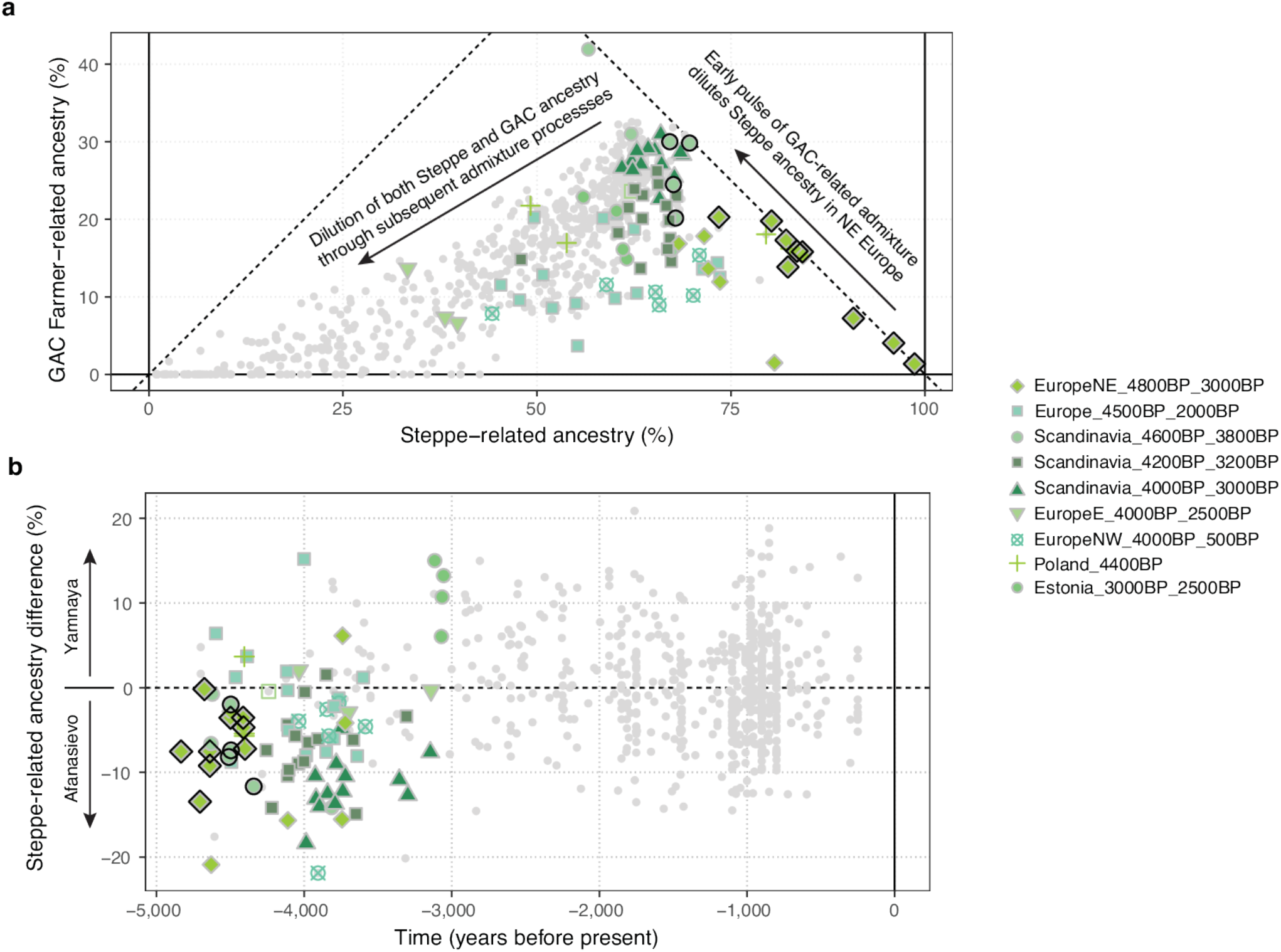
Fine-scale structure and temporal dynamics of steppe-related ancestry during the second transition in Europe. (a) Correlation between estimated proportions of steppe-related and GAC farmer-related ancestries (“postNeol” source set), across west Eurasian target individuals. (b) Timeline of difference in estimated steppe-related ancestry proportions, using individuals from genetic cluster “Steppe_5000BP_4300BP” associated with either Yamnaya or Afansievo cultural contexts as separate sources. Individuals from European post-Neolithic genetic clusters before 3,000 cal. BP are indicated with coloured symbols, while other west Eurasian target individuals are indicated with grey symbols. Symbols with black outlines highlight early steppe-related individuals associated with either Corded Ware or related (e.g., Battle Axe) cultural contexts.

While the broader impacts of the steppe migrations around 5,000 cal. BP are well known, the origin of this ancestry has remained a mystery. Here we demonstrate that the steppe ancestry composition (Steppe_5000BP_4300BP) can be modelled as a mixture of ∼65% ancestry related to herein reported HG genomes from the Middle Don River region (MiddleDon_7500BP) and ∼35% ancestry related to HGs from Caucasus (Caucasus_13000BP_10000BP) (Extended Data Fig. 6; Supplementary Data IX). Thus, Middle Don HGs, who already carried ancestry related to Caucasus HGs (Extended Data Fig. 4a), serve as a hitherto unknown proximal source for the majority ancestry contribution into Yamnaya-related genomes. The individuals in question derive from the burial ground Golubaya Krinitsa (Supplementary Note 3). Material culture and burial practices at this site are similar to the Mariupol-type graves, which are widely found in neighbouring regions of the Ukraine, for instance along the Dnepr River. They belong to the group of complex pottery-using HGs mentioned above, but the genetic composition at Golubaya Krinitsa is different from the remaining Ukrainian sites (Fig 2A, Extended Data Fig. 5). Lazaridis et al.^30^ suggested a model for the formation of Yamnaya ancestry that includes a ‘Northern’ steppe source (EHG+CHG ancestry) and a ‘Southern’ Caucasus Chalcolithic source (CHG ancestry) but without identifying the exact origin of these sources. The Middle Don genomes analysed here display the appropriate balance of EHG/CHG ancestry, suggesting them as likely candidates for the missing Northern proximate source for Yamnaya ancestry.

The dynamics of the continent-wide transition from Neolithic farmer ancestry to Steppe-related ancestry also differs markedly between geographic regions. The contribution of local Neolithic ancestry to the incoming groups was high in eastern, western and southern Europe, reaching >50% on the Iberian Peninsula (“postNeol” set; Extended Data Fig. 6; Supplementary Data X)^40^. Scandinavia, however, portrays a dramatically different picture, with much lower contributions (<15%), including near-complete replacement of the local population in some regions (Extended Data Fig. 9b). Steppe-related ancestry accompanies and spreads with the formation of the CWC across Europe and our results provide new evidence on the foundational admixture event. Individuals associated with the CWC carry a mix of steppe-related and Neolithic farmer-related ancestry and we show that the latter can be modelled as deriving exclusively from a genetic cluster associated with the Late Neolithic Globular Amphora Culture (GAC) (Poland_5000BP_4700BP), and this ancestry co-occurred with steppe-related ancestry across all sampled European regions (Fig. 5a; Extended Data Fig. 6). This suggests that the spread of steppe-related ancestry was predominantly mediated through groups already admixed with GAC-related farmer groups of the eastern European plains — an observation that has major implications for understanding the emergence of the CWC.

**Fig. 5:**
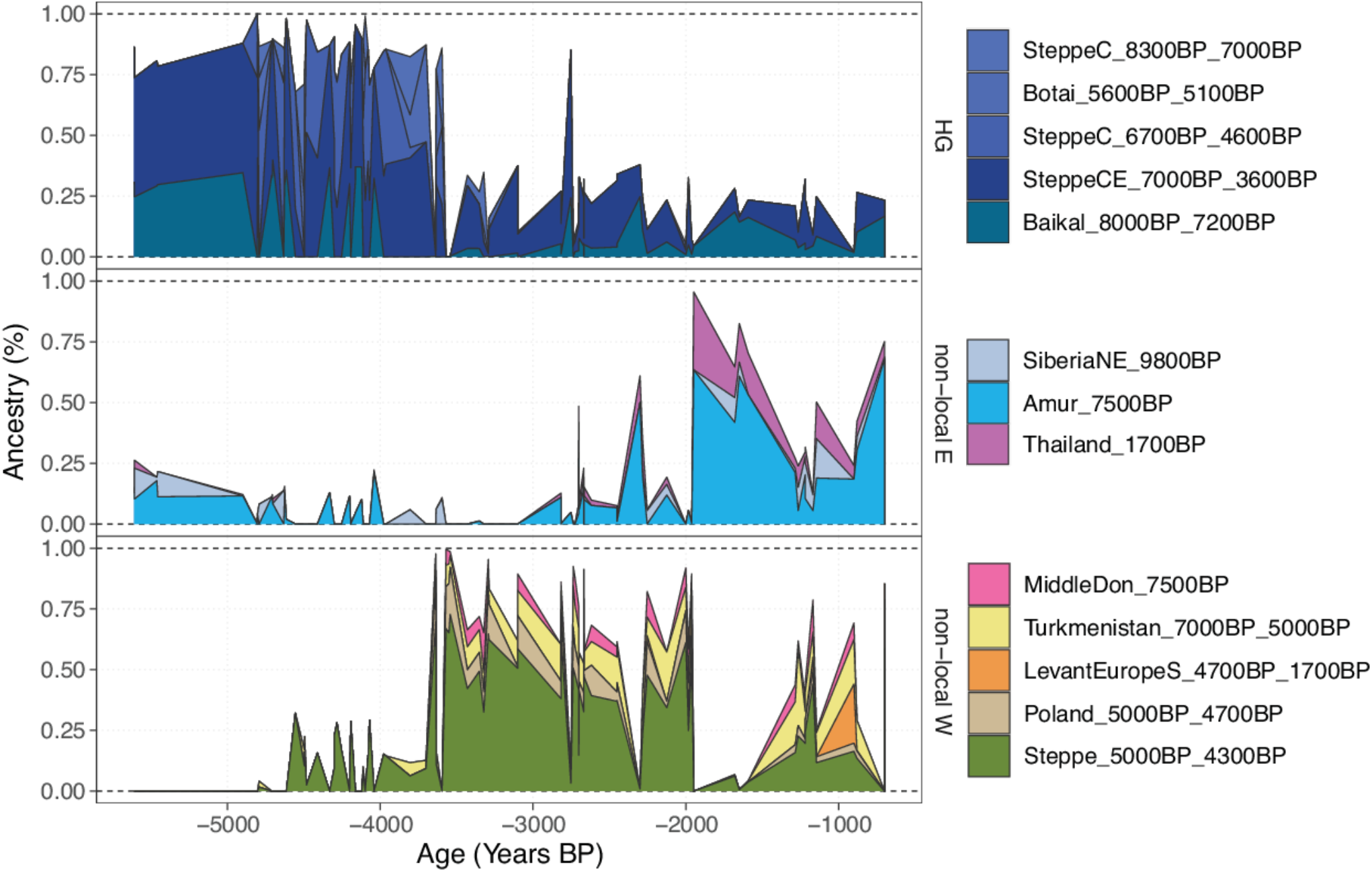
Genetic transects east of the Urals. Timelines of genetic ancestry compositions within the past 6,000 years east of the Urals. Shown are ancestry proportions in 148 imputed ancient genomes from this region, inferred using supervised ancestry modelling (“postNeol” source set). Panels separate ancestry proportions from local forest steppe HGs (HG) and sources representing ancestries originating further east or west.

A stylistic connection between GAC and CWC ceramics has long been suggested, including the use of amphora-shaped vessels and the development of cord decoration patterns^51^. Moreover, shortly before the emergence of the earliest CWC groups, eastern GAC and western Yamnaya groups exchanged cultural elements in the forest-steppe transition zone northwest of the Black Sea, where GAC ceramic amphorae and flint axes were included in Yamnaya burials, and the typical Yamnaya use of ochre was included in GAC burials^52^, indicating close interaction between these groups. Previous ancient genomic data from a few individuals suggested that this was limited to cultural influences and not population admixture^53^. However, in the light of our new genetic evidence it appears that this zone, and possibly other similar zones of contact between GAC and groups from the steppe (e.g. Yamnaya), were key in the formation of the CWC through which steppe-related ancestry and GAC-related ancestry co-dispersed far towards the west and the north^cf. 54^. This resulted in regionally diverse situations of interaction and admixture^14,32^ but a significant part of the CWC dispersal happened through corridors of cultural and demic transmission which had been established by the GAC during the preceding period^33,55^. Differences in Y-chromosomal haplogroups between CWC and Yamnaya suggests that the currently published Yamnaya-associated genomes do not represent the most direct source for the steppe ancestry component in CWC^32,33^. This notion was here supported by proximate ancestry modelling using published genomes^1^ associated with Yamnaya or Afanasievo cultural contexts as separate sources, which revealed a subtle increase in affinity for an Afanasievo-related source over a Yamnaya-related source in early European steppe-ancestry carrying individuals before 3,000 cal. BP (Fig. 5b; Extended Data Fig. 9d). The result confirms subtle population genomic structure in the population associated with Yamnaya/Afanasievo, showing that more dense sampling across the steppe horizon will be required to find the direct source(s) for steppe ancestry in early CWC.

### HG resilience east of the Urals

In contrast to the significant number of ancient HG genomes from western Eurasia studied to date, genomic data from HGs east of the Urals have remained sparse. As noted above, these regions are characterised by an early introduction of pottery from areas further east and were inhabited by complex forager societies with permanent and sometimes fortified settlements^20,56^. Here, we substantially expand knowledge on ancient populations of this region by reporting new genomic data from 38 individuals, 28 of which date to pottery-associated HG contexts between 8,300-5,000 cal. BP (Supplementary Data II). The majority of these genomes form a previously only sparsely sampled^13,42^ ‘Neolithic steppe’ cline spanning the Siberian forest steppe zones of the Irtysh, Ishim, Ob, and Yenisei River basins to the Lake Baikal region (Fig. 1c; Extended Data Fig. 1A, 3E). Supervised admixture modelling (using the “deep” set of ancestry sources; Supplementary Data IX) revealed contributions from three major sources in these HGs from east of the Urals: early West Siberian HG ancestry (SteppeC_8300BP_7000BP) dominated in the western Forest Steppe; Northeast Asian HG ancestry (Amur_7500BP) was highest at Lake Baikal; and Paleosiberian ancestry (SiberiaNE_9800BP) was observed in a cline of decreasing proportions from northern Lake Baikal westwards across the forest steppe^13^ (Extended Data Figs. 7, 10a). We used these Neolithic HG clusters (“postNeol” ancestry source set, Extended Data Fig. 7) as putative source groups in more proximal admixture modelling to investigate the spatiotemporal dynamics of ancestry compositions across the steppe and the Lake Baikal region after the Neolithic period. We replicate previously reported evidence for a genetic shift towards higher forest steppe HG ancestry (source SteppeCE_7000BP_3600BP) in Late Neolithic and Early Bronze Age individuals (LNBA) at Lake Baikal (clusters Baikal_5600BP_5400BP and Baikal_4800BP_4200BP) ^13,57^. However, ancestry related to this cluster is also already observed at ∼7,000 BP in herein-reported Neolithic HG individuals both at Lake Baikal (NEO199, NEO200), and along the Angara river to the north (NEO843) (Extended Data Fig. 7). Both male individuals at Lake Baikal belonged to Y-chromosome haplogroup Q1b1, characteristic of the later LNBA groups in the same region (Supplementary Note 3b, Figure S3b.5). Together with an early estimated admixture time (∼7,300 cal. BP upper bound) for the LNBA groups (Supplementary Data XIV), these results suggest that gene flow between HGs of Lake Baikal and the south Siberian forest steppe regions already occurred during the Eastern Early Neolithic, consistent with archaeological interpretations of contact. In this region, bifacially flaked tools first appeared near Baikal^58^ from where the technique spread far to the west. We find echoes of such bifacial flaking in archaeological complexes (Shiderty 3, Borly, Sharbakty 1, Ust-Narym, etc.) in Northern and Eastern Kazakhstan, around 6,500-6,000 cal. BP^59,60^. Here, Mesolithic cultural networks with Southwest Asia have also been recorded, as evidenced by pebble and flint lithics known from Southwest Asia cultures^61^.

Genomes reported here also shed light on the genetic origins of the Early Bronze Age Okunevo Culture in the Minusinsk Basin in Southern Siberia. In contrast to previous results, we find no evidence for Lake Baikal HG-related ancestry in the Okunevo^13,57^ when using our newly reported Siberian forest steppe HG genomes jointly with Lake Baikal LNBA genomes as putative proximate sources. Instead, we found that they originate from admixture of a forest steppe HG source (best modelled as mixture of clusters Steppe_6700BP_4600BP and SteppeCE_7000BP_3600BP) and steppe-related ancestry (Steppe_5300BP_4000BP; Extended Data Fig. 7, set “postBA”; Supplementary Data XI). We date the admixture with steppe-related ancestry to ∼4,600 BP (Supplementary Data XIV), and found it to be modelled exclusively from an Afanasievo-related source in proximate modelling separating the Yamnaya and Afanasievo steppe-ancestries (Extended Data Figs. 9d, 10c,e). This is direct evidence for gene flow from peoples of the Afanasievo Culture that were closely related to Yamnaya and existed near Altai and Minusinsk Basin during the era of the steppe migrations^1,57^.

From around 3,700 cal. BP, individuals across the steppe and Lake Baikal regions display markedly different ancestry profiles (Fig. 6; Extended Data Fig. 7, 9b). We document a sharp increase in nonlocal ancestries, with only limited ancestry contributions from local HGs. The early stages of this transition are characterised by influx of steppe-related ancestry, which decays over time from its peak of ∼70% in the earliest individuals. Similar to the dynamics in western Eurasia, steppe-related ancestry is here correlated with GAC-related farmer ancestry (Poland_5000BP_4700BP; Fig. 6; Extended Data Fig. 10b), recapitulating previously documented gene flow from GAC groups into steppe/forest steppe neighbouring groups and the eastward expansion of admixed Western steppe pastoralists from the Sintashta and Andronovo complexes during the Bronze Age^42,62^. However, GAC-related ancestry is notably absent in individuals of the Okunevo Culture, and individuals with steppe ancestry after 3,700BP show slight excess in affinity to Yamnaya over Afanasievo in proximate modelling (Extended Data Fig. 10d), providing further support for two distinct eastward migrations of Western steppe pastoralists during the early (Yamnaya-related) and later (Sintashta, Andronovo) Bronze Age. The later stages of the transition are characterised by increasing Central Asian (Turkmenistan_7000 BP_5000BP) and Northeast Asian-related (Amur_7500BP) ancestry components (Fig. 6; Extended Data Fig. 10b). Together, these results show that deeply structured HG ancestry dominated the eastern Eurasian steppe substantially longer than in West Eurasia, before successive waves of population expansions swept across the steppe within the last 4,000 years. These included a large-scale introduction of domesticated horse lineages concomitant with new equestrian equipment and spoke-wheeled chariotry^62,63^, as well as the adoption of millet as robust subsistence crop^64^.

**Fig. 6:**
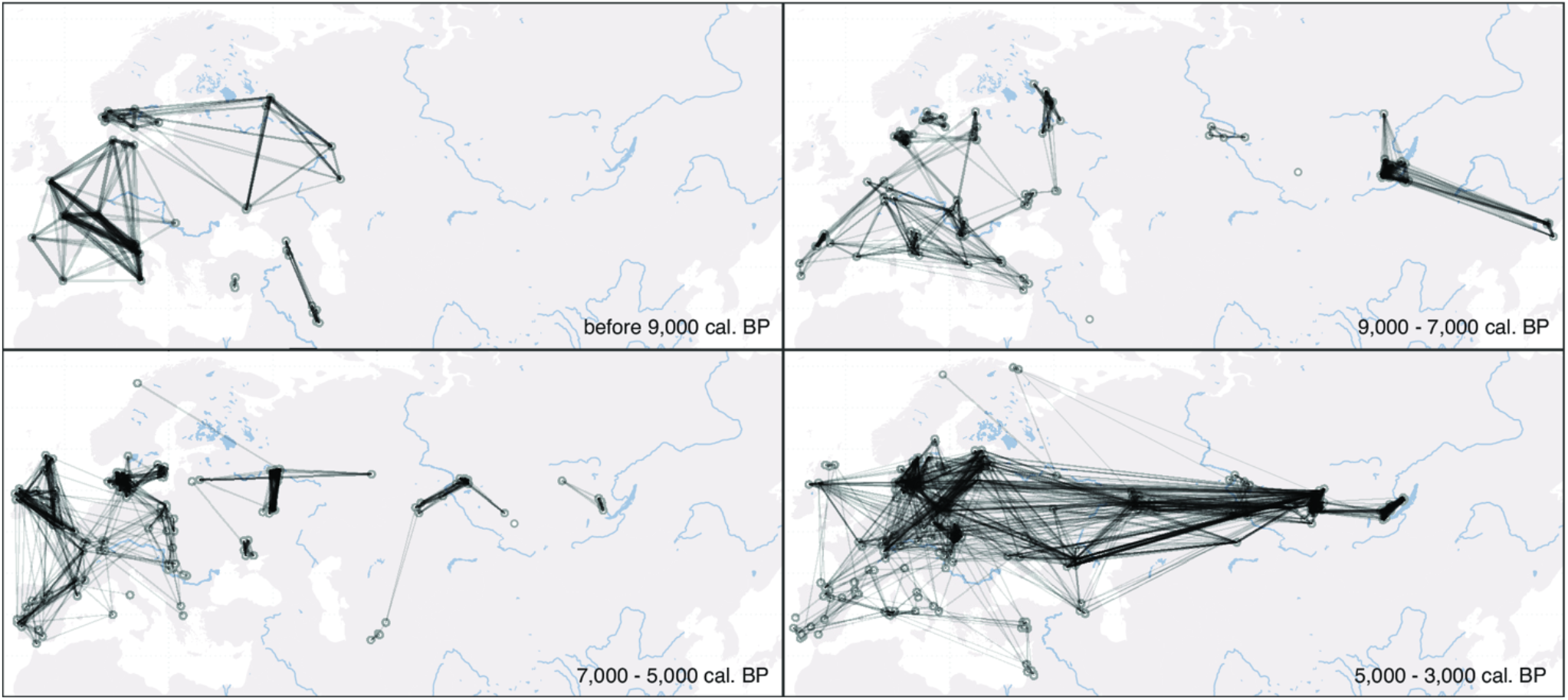
Genetic relatedness across Western Eurasia. Maps showing networks of highest IBD sharing (top 10 highest sharing per individual) during different time periods for 579 imputed genomes predating 3,000 cal. BP and located in the geographic region shown. Shading and thickness of lines is scaled to represent the amount of IBD shared between two individuals. In the earliest periods, sharing networks exhibit strong links within relatively narrow geographic regions, representing predominantly close genetic ties between small HG communities, and rarely crossing the East-West divide extending from the Baltic to the Black Sea. From ∼9,000 cal. BP onwards, a more extensive network with weaker individual ties appears in the south, linking Anatolia to the rest of Europe, as early Neolithic farmer communities spread across the continent. The period 7,000 - 5,000 cal. BP shows more connected subnetworks of western European and eastern/northern European Neolithic farmers, while locally connected networks of HG communities prevail on the eastern side of the divide. From c. 5,000 BP onwards the divide finally collapses, and continental-wide genetic relatedness unifies large parts of Western Eurasia.

### Sociocultural insights

We used patterns of pairwise IBD sharing between individuals to examine our data for temporal shifts in relatedness within genetic clusters. We found clear trends of a reduction of within-cluster relatedness over time, in both western and eastern Eurasia (Extended Data Fig. 11a). This pattern is consistent with a scenario of increasing effective population sizes during this period^65^. Nevertheless, we observe notable differences in temporal relatedness patterns between western and eastern Eurasia, mirroring the wider difference in population dynamics discussed above. In the west, within-group relatedness changed substantially during the Neolithic transition (∼9,000 to ∼6,000 BP), where clusters of individuals with Anatolian farmer-related ancestry show overall reduced IBD sharing compared to clusters of individuals with HG-associated ancestry (Extended Data Fig. 11a). In the east, genetic relatedness remained high until ∼4,000 BP, consistent with a much longer persistence of smaller localised HG groups (Fig. 6; Extended Data Fig. 11a). Next, we examined the data for evidence of recent parental relatedness, by identifying individuals harbouring > 50cM of their genomes in long (>20cM) ROH segments^66^. We only detect 29 such individuals out of a total sample of 1,396 imputed ancient genomes from across Eurasia (Extended Data Fig. 11b). This suggests that close kin mating was not common in the regions and periods covered by our data. No obviously discernible spatiotemporal or cultural clustering was observed among the individuals with recent parental relatedness. Interestingly, a ∼1,700-year-old Sarmatian individual from Temyaysovo (tem003)^67^ was found homozygous for almost the entirety of chromosome 2, but without evidence of ROHs elsewhere in the genome, suggesting the first documented case of uniparental disomy in an ancient individual (Extended Data Fig. 11c). Among several noteworthy familial relationships (see Supplementary Fig. S3c.2), we report a Mesolithic father/son burial at Ertebølle (NEO568/NEO569), as well as a Mesolithic mother/daughter burial at Dragsholm (NEO732/NEO733), Denmark (see also *Allentoft, Sikora, Fischer et al. submitted*).

### Formation and dissolution of the divide

We have demonstrated the existence of a clear east-west genetic division extending from the Black Sea to the Baltic, mirroring archaeological observations, and persisting over several millennia. We show that this deep ancestry division in the Eurasian human gene pool that was established during early post-LGM dispersals^7^ was maintained throughout the Mesolithic and Neolithic ages (Fig. 8). Accordingly, we show that the genetic impact of the Neolithic transition was highly distinct east and west of this boundary. These observations raise a series of questions related to understanding the underlying drivers.

In eastern Europe, the expansion of Neolithic farming was halted for around 3,000 years and this delay may be linked to environmental factors, with regions east of the division having more continental climates and harsher winters, possibly less suited for Middle Eastern agricultural practices^68^. Here, highly developed HG societies persisted with stable, complex and sometimes fortified settlements, long distance exchange and large cemeteries^69,70^. A diet including freshwater fish is clear both from our isotopic data (Supplementary Data II) and from analyses of lipids in pottery^70^. In the northern forested regions of this boundary zone, HG societies persisted until the emergence of the CWC around 5,000 cal. BP, whereas in the southern and eastern steppe regions, hunting and gathering was eventually complemented with some animal husbandry (cattle and sheep), and possibly horse herding in central Asia^71^. Some of these groups, such as Khvalynsk at the Volga saw the emergence of male sodalities involved in wide-ranging trade connections of copper objects from east central Europe and the Caucasus^29^. Settlements were confined mainly to the flat floodplains and river valleys, while the steppe belt remained largely unexploited. The eventual dissolution of this genetic, economic, and social border was driven by events unfolding in the steppe region. Here, two temporal phases of technological innovations can be observed archaeologically: the widespread dispersal of ox-drawn wheeled vehicles around 5,500 cal. BP and the later development of horse riding, and possible changing environmental conditions^72^. This opened up the steppe as an economic zone, allowing Yamnaya groups to exploit the steppe as pastoral nomads around 5,000 cal. BP^73^and all Eneolithic settlements along river valleys were replaced by this new mobile economy^74^ finally dissolved the great genomic boundary that had persisted in the preceding millenia (Fig. 8).

By 4,000 cal. BP the invention of chariot warfare and the adoption of millet as a food crop allowed the final eastward expansion into central Asia and beyond by Andronovo and related groups, with global legacies for the expansion of Indo-European languages^75^. Our study has provided new genetic knowledge on these steppe migrations on two levels: we have identified a hitherto unknown source of ancestry in HGs from the Middle Don region contributing ancestry to the steppe pastoralists, and we have documented how the later spread of steppe-related ancestry into Europe via the CWC was first mediated through peoples associated with the GAC. In a contact zone that included forested northern regions, the CWC was rapidly formed from a cultural and genetic amalgamation of steppe-groups related to Yamnaya and the GAC groups in eastern Europe. In accordance with their mixed cultural and genetic background, the CWC practised a mixed economy, employing various subsistence strategies in different environments. This flexibility would have contributed substantially to their success in settling and adapting to very different ecological and climatic settings over a very short period of time^33^.

## Supporting information

Supplementary Data VII

Supplementary Data VIII-XIII

Supplementary Data XIV

## Methods

### aDNA data generation and authentication

Sampling of ancient human remains was undertaken in collaboration with co-authors responsible for the curation and contextual analyses of these, and with approval of the relevant institutions responsible for the archaeological remains (detailed in the Reporting Summary). Laboratory work was undertaken in dedicated aDNA cleanlab facilities (Globe Institute, University of Copenhagen) following optimised aDNA protocols^1,76^ (Supplementary Note 1). Double-stranded blunt-end libraries were constructed from the extracted DNA using NEBNext DNA Prep Master Mix Set E6070 (New England Biolabs Inc.) and sequenced (80bp and 100bp single read) on Illumina HiSeq 2500 and 4000 platforms. Initial shallow shotgun-screening identified 317 of 962 ancient samples with sufficient DNA preservation for deeper sequencing. Of these, 211 were teeth, 91 were petrous bones, and 15 were sampled from long bones, ribs and cranial bones (Supplementary Data II). Reads were mapped to the human reference genome build 37 and also to the mitochondrial genome (rCRS) alone. Mapped reads were filtered for mapping quality 30 and sorted using *Picard (v*.*1*.*127)* (http://picard.sourceforge.net) and *samtools*^*77*^. Data was merged to library level and duplicates removed using *Picard MarkDuplicates (v*.*1*.*127)* and merged to sample level. Sample-level BAMs were re-aligned using *GATK (v*.*3*.*3*.*0)* and hereafter had the md-tag updated and extended BAQs calculated using *samtools calmd (v*.*1*.*10)*^*77*^. Read depth and coverage were determined using *pysam* (https://github.com/pysam-developers/pysam) and *BEDtools (v*.*2*.*23*.*0)*^*78*^. Post-mortem DNA damage patterns were determined using *mapDamage2*.*0*^*79*^. For the 317 samples we observed C-to-T deamination fractions ranging from 10.4% to 67.8%, with an average of 38.3% across all samples (Supplementary Data I). These numbers indicate DNA molecule degradation consistent with a millennia-scale depositional age. Three different methods were used to estimate DNA contamination: two based on mitochondrial sequences^80,81^ and one method investigating X-chromosomal data in males (ANGSD, Supplementary Note 1). All contamination estimates are reported in Supplementary Data V (summary values in Supplementary Data I). Based on this approach we have a total of 15 samples flagged as ‘possibly contaminated’ in our downstream analyses (Supplementary Note 1).

### Imputation of ancient genomes

We imputed the ancient genomes in this study using the imputation and phasing tool GLIMPSE v1.0.0^34^ and 1000 Genomes phase3^35^ as a reference panel. We first generated genotype likelihoods at the biallelic 1000 Genomes variant sites from the bam files with bcftools v1.10 and the command *bcftools mpileup with parameters -I -E -a ‘FORMAT/DP’ --ignore-RG*, followed by *bcftools call - Aim -C alleles*. Using GLIMPSE_chunk, the genotype likelihood data were first split into chunks of sizes between 1 and 2 Mb with a buffer region of 200 kb at each side. We then imputed each chunk with GLIMPSE_phase with parameters *--burn 10, --main 15 and --pbwt-depth 2*. Finally, the imputed chunks were ligated with *GLIMPSE_ligate*. To validate the accuracy of the imputation, 42 high coverage (5X to 39X) genomes, including a Neolithic trio, were downsampled for testing^82^ (Supplementary Note 2). We evaluated imputation accuracy on the basis of depth of coverage; minor allele frequency; and ancestry and timeframe of ancient genomes, using high coverage ancient genomes^82^. >1X genomes provided remarkably high imputation accuracy (closely matching that obtained for modern samples, Extended Data Fig. 2), except for African genomes that had lower accuracy due to poor representation of this ancestry in the reference panel. Imputation accuracy was influenced by both MAF and coverage (Supplementary Fig. S2.3). We found that coverage as low as 0.1X and 0.4X was sufficient to obtain r^2^ imputation accuracy of 0.8 and 0.9 at common variants (MAF≥10%), respectively. We conclude that ancient genomes can be imputed confidently from coverages above 0.4X, and genome-wide aggregate analyses relying on common SNPs (e.g. PCA and admixture modelling) can be carried out with a low amount of bias for genome coverage from as low as 0.1X when using specific QC on the imputed data (although at very low coverage a bias arises towards the major allele, see Supplementary Note 2). We additionally tested for possible effects of bias affecting inferred ancestry components^82^ propagating biases in individual-level pairwise analyses, using D-statistics, which indicated that imputed ancient genomes down to 0.1x coverage are not significantly affected (Supplementary Note 2).

### Demographic inference

We determined the genetic sex of the study individuals using the ratio of reads aligning to either of the sex chromosomes (R_Y_ statistic)^83^. Y chromosomes of inferred male individuals were further analysed using phylogenetic placement^84^. We built a reference phylogenetic tree of 1,244 male individuals from the 1000 Genomes project with RAxML-NG^85^, using the general time-reversible model including among-site rate heterogeneity and ascertainment correction (model GTR+G+ASC_LEWIS). For each ancient sample, haploid genotypes given the positions and alleles in the reference panel were called using ‘bcftools call’ (options *-C alleles –ploidy 1 -i*). The resulting genotypes were converted to fasta format and placed onto the reference tree using EPA-ng^84^. Phylogenetic placements were processed and visualised using gappa^86^. To convert phylogenetic placements into haplogroup calls, we assigned each branch of the reference phylogeny to its representing haplogroup, using SNP annotations from ISOGG (version 15.73). For each ancient sample, haplogroups were then called using the most basal branch accumulating 99% of the placement weights, obtained using *accumulate* in gappa. Phylogenetic analyses of reconstructed mitochondrial genomes were also undertaken using RAxML-ng^84^ (see Supplementary Note 3a). To infer genetic relatedness between the study individuals we used the allele-frequency free inference method introduced by^87^. For each pair of individuals, three relatedness estimators were calculated, R0, R1 and KING-robust^88^ using the site-frequency-spectrum (SFS)-based approach. We used the realSFS^89^ method implemented in the ANGSD^90^ package to infer the 2D-SFS, selecting the SFS with the highest likelihood across ten replicates. We used a set of 1,191,529 autosomal transversion SNPs with minor allele frequency ≥ 0.05 from the 1000 Genomes Project^35^ for the analysis. Previously established cut-offs^88^ for the KING-robust estimator were applied to assign individual pairs to first-, second-or third-degree relationships. Parent-offspring relationships were distinguished from sibling relationships using R0 and R1 ratios, by requiring that R0 ≤ 0.02 and 0.4 ≤ R1 ≤ 0.6 to infer a parent-offspring relative pair. Individual pairs with less than 20,000 sites contributing to the estimators were excluded.

We generated a dataset for population genetic analysis by combining the 317 newly sequenced individuals with 1,347 previously published ancient genomes with genomic coverage >0.1X generated using shotgun sequencing (Supplementary Data VII). Imputed genotype data (Supplementary Note 2) for this set of 1,664 ancient genomes were merged with genotypes of 2,504 modern individuals from the 1,000 Genomes project^35^ used as a reference panel in the imputation. We retained only SNPs passing the 1000 Genomes strict mask, resulting in a final dataset of 4,168 individuals genotyped at 7,321,965 autosomal SNPs (“1000G” dataset). In addition to imputed genotypes, we also generated pseudo-haploid genotypes for each ancient individual by randomly sampling an allele from sequencing reads covering those SNPs. For population structure analyses in the context of global genetic diversity, we generated a second dataset by intersecting the ancient genotype data with SNP array data of 2,180 modern individuals from 213 world-wide populations^3,4,91,92^ (“HO” dataset).

To facilitate filtering for downstream analyses, we flagged individuals to potentially exclude based on the following criteria: i) Contamination estimate >5% (“contMT5pct”, “contNuc5pct”; Supplementary Note 1); ii) Autosomal coverage <0.1X (“lowcov”), III) Genome-wide average imputation genotype probability <0.98 (“lowGpAvg”), IV) Individual is the lower quality sample in a close relative pair (“1d_rel”, “2d_rel”; Supplementary Note 3c). A total of 1,492 individuals (213 newly reported) passed all filters, which were used in the majority of downstream analyses unless otherwise noted.

We investigated overall population structure among the dataset individuals using principal component analyses (PCA) and model-based clustering (ADMIXTURE^93^). We carried out PCA using different subsets of individuals in the “HO’’ dataset. For the PCA including only imputed diploid samples, we used GCTA^94^, excluding SNPs with minor allele frequency (MAF) < 0.05 in the respective panel. For PCA projecting low coverage or flagged individuals, we used smartpca^95,96^ with options ‘*lsqproject: YES*’ and ‘*autoshrink: YES*’ on a fixed set of 400,186 SNPs with MAF ≥ 0.05 in non-African individuals passing all filters. We ran ADMIXTURE on a set of 1,593 ancient individuals from the “1000G” dataset, excluding individuals flagged as close relatives or a contamination estimate >5%. For the 1,492 individuals passing all filters we used imputed genotypes, the remaining 101 lower coverage samples were represented by pseudo-haploid genotypes. We restricted the analysis to transversion SNPs with imputation INFO score ≥ 0.8 and MAF ≥ 0.05. We further performed linkage disequilibrium (LD) pruning and filtering for missingness using plink^97^ (options *--indep-pairwise 500 50 0*.*4 –geno 0*.*8*), for a final analysis set of 142,550 SNPs.

We performed admixture graph fitting (qpGraph) to investigate deep Eurasian population structure using ADMIXTOOLS2^98^. For these analyses, pairwise *f*_2_-statistics were pre-computed from pseudo-haploid genotypes in the “1000G” dataset using the ‘extract_f2’ function with ‘afProd=TRUE’. We grouped individuals into populations using their membership in the genetic clusters inferred from IBD sharing (Supplementary Note 3f), with the exception of the UP European individual Kostenki 14, which was treated as a separate population (new cluster label “Europe_37000BP_33000BP_Kostenki”). We carried out admixture graph fitting using a semi-automatic iterative approach (Supplementary Note 3d).

We used IBDseq^99^ to detect genomic segments shared identical-by-descent (IBD) between all individuals in the “1000G” dataset, restricting to transversion SNPs with imputation INFO score ≥ 0.8 and MAF ≥ 0.01. We filtered the resulting IBD segments for LOD score ≥ 3 and a minimum length of 2 centimorgans (cM), and further removed regions of excess long IBD following^100^. First, we used the GenomicRanges^101^ package in R to calculate the total number of long IBD segments (>10cm) overlapping each position along the genome, and calculated their 3% trimmed mean and standard deviation (SD). We then called regions of excess IBD if they were > 10 trimmed SD from the trimmed mean, and removed any segments overlapping the excess IBD regions. For analyses of runs of homozygosity (ROH) we used a shorter length cutoff of 1cM.

We carried out genetic clustering of the ancient individuals using hierarchical community detection on a network of pairwise identity-by-descent (IBD)-sharing similarities^102^. To facilitate detection of clusters at a finer scale, we ran IBDseq (version r1206) on a dataset restricting to ancient samples only, and applied more lenient filters of imputation INFO score > 0.5, and minimum IBD segment length of 1 cM. We constructed a weighted network of the individuals using the igraph^103^ package in R, with the fraction of the genome shared IBD between pairs of individuals as weights. We then performed iterative community detection on this network using the Leiden algorithm^104^ implemented in the leidenAlg R package (v1.01, https://github.com/kharchenkolab/leidenAlg). We used a resolution parameter of r=0.5 as the starting value for each level of community detection. If more than one community was detected, we split the network into the respective communities, and repeated the community detection step. If no communities were detected, we incremented the resolution parameter in steps of 0.5 until a maximum value of r=3. The initial clustering was completed when no more communities were detected at the highest resolution parameter, across all subcommunities. To convert the resulting hierarchy into a final clustering, we simplified the initial clustering by collapsing nodes into single clusters based on observed spatiotemporal annotations of the samples. We note that the obtained clusters should not be interpreted as ‘populations’ in the sense of a local community of individuals, but rather as sets of individuals with shared ancestry. While this approach is an oversimplification of the complex spatiotemporally structured populations investigated here, the obtained clusters nevertheless captured real effects, grouping individuals within restricted spatiotemporal ranges and/or archaeological contexts and recapitulating known relationships between clusters.

To circumvent some of the pitfalls of grouping individuals into discrete clusters, we used supervised ancestry modelling where sets of ‘target’ individuals were modelled as mixtures of ‘source’ groups, selected to represent particular ancestry components. As an illustrative case, an individual of European HG ancestry with a minor contribution of Neolithic farmer admixture might be inferred to be a member of a HG genetic cluster, but will be modelled as a mixture of a HG and Neolithic farmer sources in the ancestry modelling. To estimate ancestry proportions from patterns of pairwise IBD sharing, we applied an approach akin to “chromosome painting”^105^. We first inferred an IBD-based “painting profile” for each target individual, by summing up the total amount of IBD shared with each “donor” group (using population labels for modern donors or IBD-based genetic clusters for ancient donors), and normalising them to the interval [0,1]. We used a leave-one-out approach following^37^ to account for the fact that recipient individuals cannot be included as donors from their own group. We then used these painting profiles in supervised modelling of target individuals as mixtures from different sets of putative source groups^37,106^, using non-negative least squares implemented in the R package limSolve^107^. We estimated standard errors of ancestry proportions using a weighted block jacknife, leaving out each chromosome in turns. A comparison of results obtained using this approach to other commonly used methods (supervised ADMIXTURE, qpAdm) is shown in Supplementary Note 3f). We focussed our analyses on three panels of putative source clusters reflecting different temporal depths: ‘deep’, using a set of deep ancestry source groups reflecting major ancestry poles; ‘postNeol’, using diverse Neolithic and earlier source groups; and ‘postBA’, using Late Neolithic and Bronze Age source groups (Extended Data Figs. 5-7). We also used additional source sets in follow-up analyses of more restricted spatiotemporal contexts (Supplementary Datas VII-XIII).

Finally, we aimed to infer the geographic and temporal spread of major ancestries (Supplementary Note 3e). We used a method^45^ applying spatiotemporal ordinary kriging on latent ancestry proportion estimates from ancient and present-day genomes. This way, we obtained spatiotemporal maps reflecting the dynamics of the spread of ancestry during the transition from the Mesolithic to the Neolithic, Bronze Age, Iron Age and more recent periods. We obtained ancestry proportions estimated using ADMIXTURE^108^ with K=9 latent ancestry clusters (Supplementary Note 3d) on a sequence dataset including both whole-genome shotgun-sequenced genomes and genomic sequences obtained via SNP capture (Supplementary Note 2, intersection with “HO” dataset). We performed spatiotemporal kriging^109^ of these proportions over the last 12,900 years, in intervals of 300 years, with a 5,000-point spatial grid spanning Western and Central Eurasia. We used the R package *gstat* to fit a spatiotemporal variogram via a metric covariance model, and perform ordinary kriging^110^. We focused on the ancestry clusters for which we could fit variogram models that were not static over time.

### ^14^C chronology and reservoir effects

Of the 317 individuals sequenced in this study 272 were ^14^C-dated in the project, while 30 ^14^C-dates were obtained from literature, and 15 were dated by archaeological context (Supplementary Note 4, Supplementary Data II). Some individuals were dated twice. Most of the dates (n=242) were performed at the ^14^CHRONO Centre laboratory at Queen’s University, Belfast, following published sample pretreatment and laboratory protocols^111^. Additional samples were analysed by the Oxford Radiocarbon Accelerator Unit (ORAU) laboratory (n=24) and by the Keck-CCAMS Group, Irvine, California, USA (n=6) (see ^112,113^ for laboratory procedures). Only datings with a C/N ratio of 2.9-3.6 were accepted; both δ13C and δ15N collagen measurements were also performed, and were used in estimates of marine (MRE) and freshwater (FRE) reservoir effects (see Supplementary Note 4, Supplementary Data IV). Published values of MRE and FRE were used where available, but for some regions, such as sites in western Russia, a standard FRE value of 500 years was applied. A diet-weighted reservoir offset was then applied to the ^14^C central value before calibration. Calibrations were made in Oxcal 4.4 using the Intcal20 calibration curve^114^. For display and calculation purposes a midpoint of the reservoir corrected and calibrated 95% interval was calculated. Full details of the reservoir correction and calibration procedure are given in Supplementary Note 4 and the calculations are found in Table S4.1.

## Extended Data Figures

**Extended Data Fig. 1:**
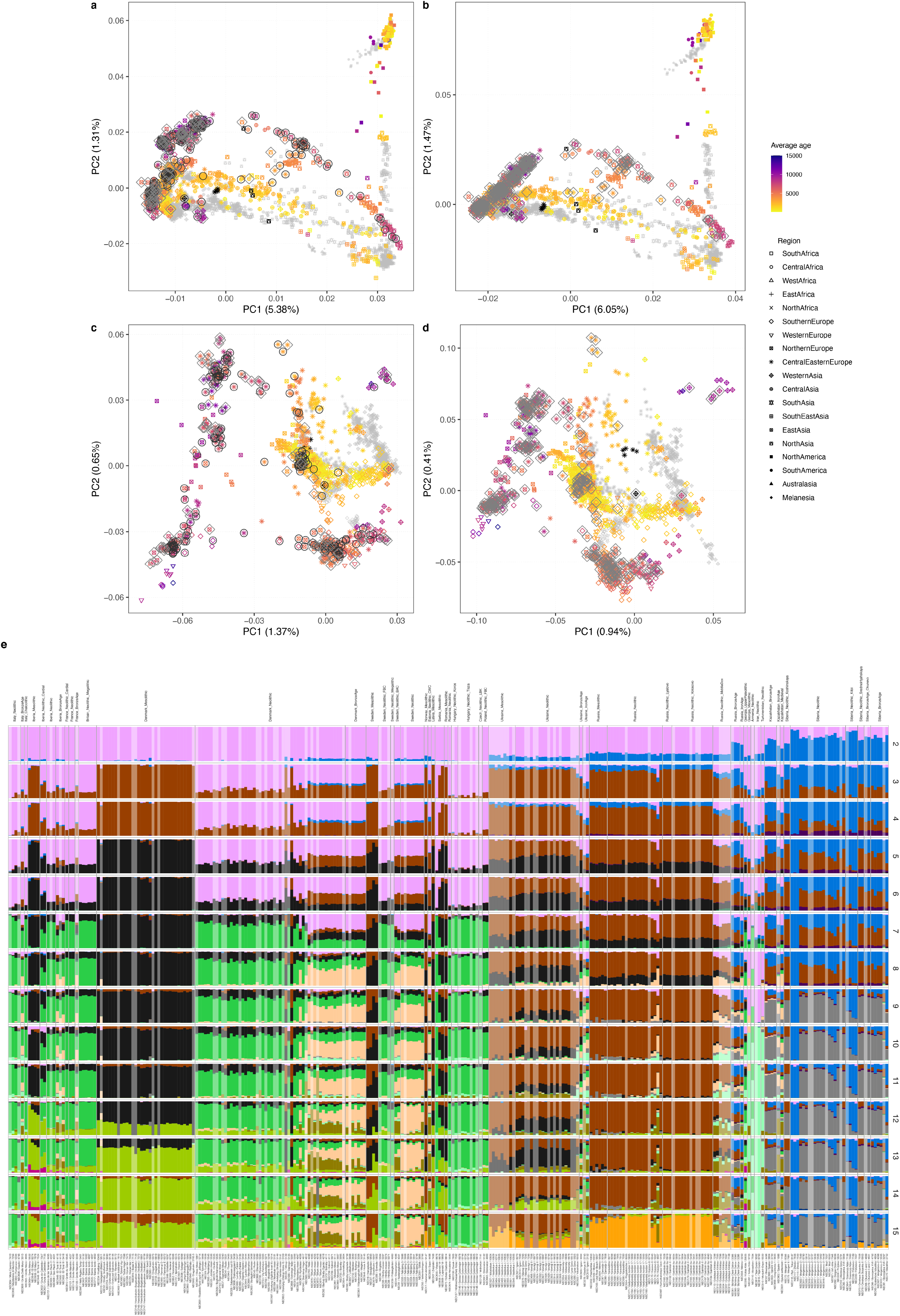
Genetic structure of the 317 herein-reported ancient genomes. (a-d) Principal component analysis of 3,316 modern and ancient individuals from Eurasia, Oceania and the Americas (a, b), as well as restricted to 2,126 individuals from western Eurasia (west of the Urals) (c, d). Shown are analyses with principal components inferred either using both modern and imputed ancient genomes passing all filters, and projecting low coverage ancient genomes (a, c); or only modern genomes and projecting all ancient genomes (b, d). Ancient genomes sequenced in this study are indicated either with black circles (imputed genomes) or grey diamonds (projected genomes). (e) Model-based clustering results using ADMIXTURE for 284 newly reported genomes (excluding close relatives and individuals flagged for possible contamination). Results shown are based on ADMIXTURE runs from K=2 to K=15 on 1,593 ancient individuals, corresponding to the full set of 1,492 imputed genomes passing filters as well as 101 low coverage genomes represented by pseudo-haploid genotypes (flags “lowcov” or “lowGpAvg”, Supplementary Data VII; indicated with alpha transparency in plot).

**Extended Data Fig. 2.**
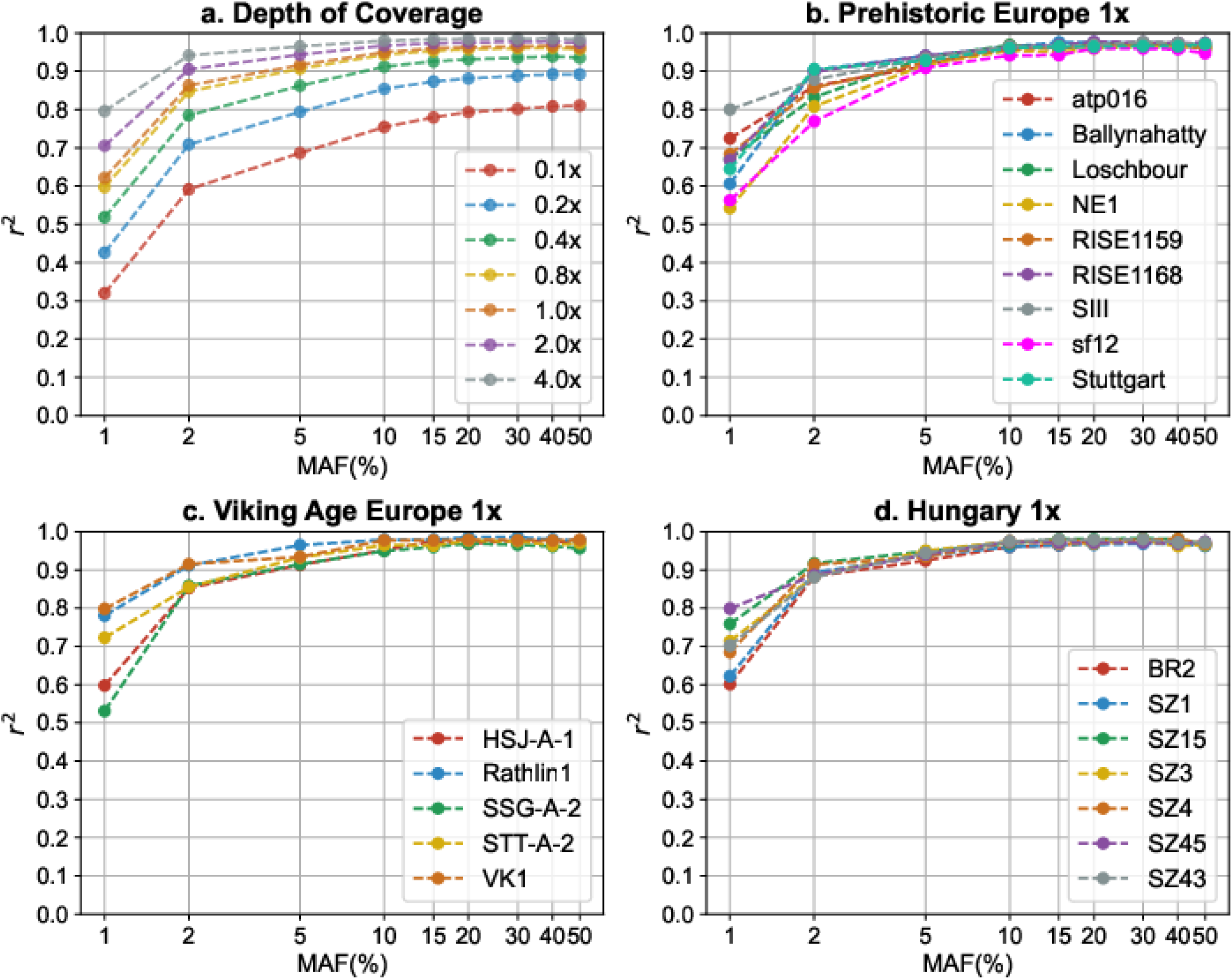
Imputation accuracy of aDNA. (a) imputation accuracy across 42 high-coverage ancient genomes when downsampled to lower depth of coverage values (see Supplementary Note 2, Table 2.1). (b-d) imputation accuracy for 1X depth of coverage across 21 ancient European genomes. In all panels, imputation accuracy is shown as the squared Pearson correlation between imputed and true genotype dosages as a function of minor allele frequency of the target variant sites.

**Extended Data Fig. 3.**
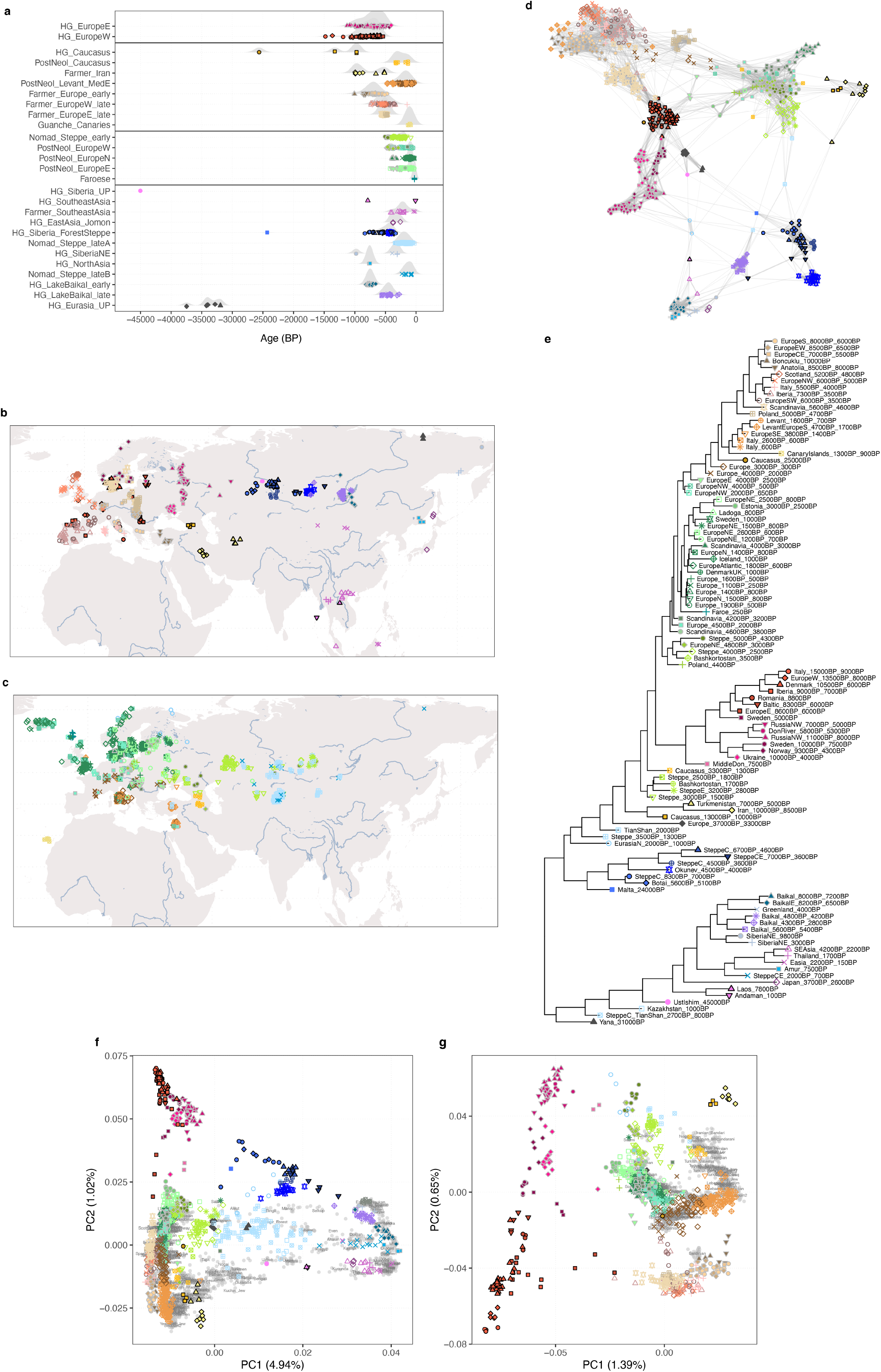
Genetic clustering of ancient individuals. Characterization of genetic clusters for 1,401 imputed ancient individuals from Eurasia (i.e. excluding 91 individuals from Africa and Americas), inferred from pairwise identity-by-descent (IBD) sharing (indicated using coloured symbols throughout) (a) Temporal distribution of clustered individuals, grouped by broad ancestry cluster. (b), (c) Geographical distribution of clustered individuals, shown for individuals predating 3,000 BP (b) and after 3,000 BP (c). (d) Network graph of pairwise IBD sharing between 596 ancient Eurasians predating 3,000 BP, highlighting within- and between-cluster relationships. Each node represents an individual, and the width of edges connecting nodes indicates the fraction of the genome shared IBD between the respective pair of individuals. Network edges were restricted to the 10 highest sharing connections for each individual, and the layout was computed using the force-directed Fruchterman-Reingold algorithm. (e) Neighbour-joining tree showing relationships between genetic clusters, inferred using total variation distance (TVD) of IBD painting palettes. (f), (g) PCA of 3,119 Eurasian (f) or 2,126 west Eurasian (g) ancient and modern individuals (“HO” dataset).

**Extended Data Fig. 4:**
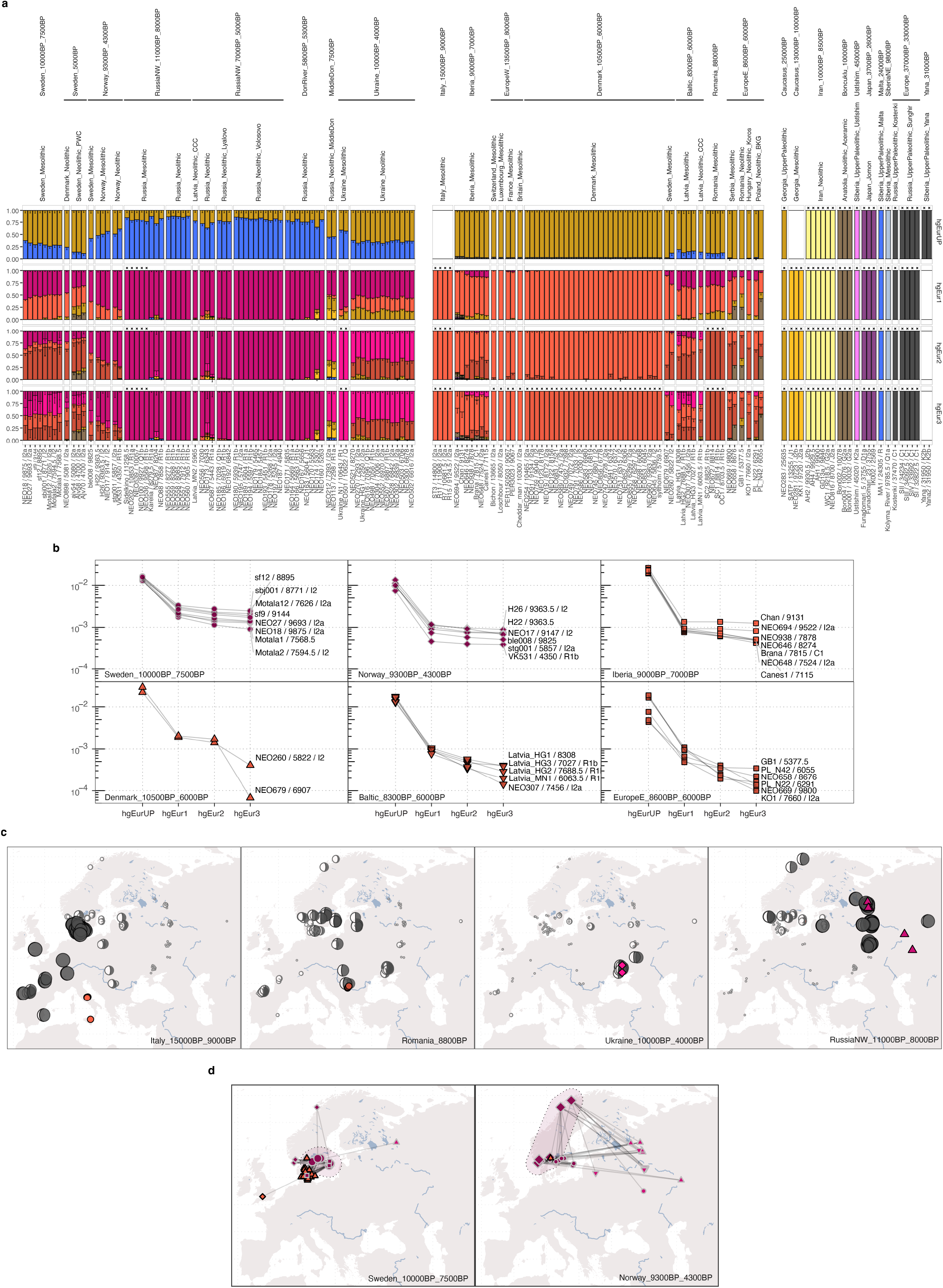
Genetic structure of European hunter-gatherers after the LGM. (a) Supervised ancestry modelling using non-negative least squares on IBD sharing profiles. Panels show estimated ancestry proportions for target individuals from genetic clusters representing European HGs, using different sets of increasingly proximal source groups. Individuals used as sources in a particular set are indicated with black crosses and coloured bars with 100% ancestry proportion. Black lines indicate 1 standard error for the respective ancestry component. (b) Residuals for model fit of target individuals from selected genetic clusters across different source sets. (c) Moon charts showing spatial distribution of ancestry proportions in European HGs deriving from four European source groups (set “hgEur2”; source origins shown with coloured symbol). Estimated ancestry proportions are indicated by both size and amount of fill of moon symbols. Note that ‘Italy_15000BP_9000 BP’ and ‘RussiaNW_11000BP_8000BP’ correspond to ‘WHG’ and ‘EHG’ labels used in previous studies. (d) Maps showing networks of highest between-cluster IBD sharing (top 10 highest sharing per individual) for individuals from two genetic clusters representing Scandinavian HGs. See Supplementary Datas I and VII for details of individual sample IDs presented here.

**Extended Data Fig. 5:**
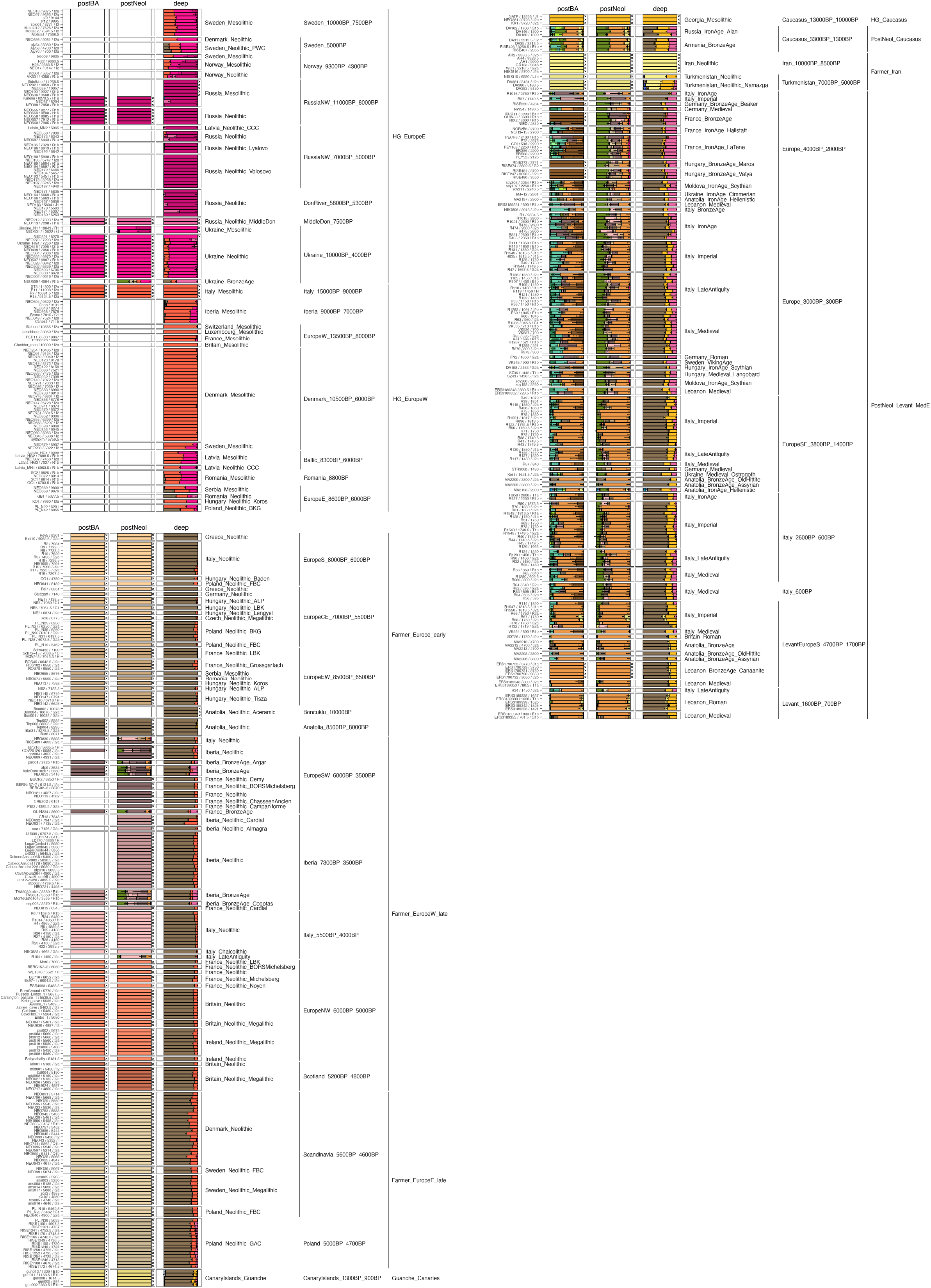
Ancestry modelling for HG and Neolithic farmer-associated genetic clusters. Supervised ancestry modelling using non-negative least squares on IBD sharing profiles. Panels show estimated ancestry proportions of two global Eurasian clusters, corresponding to European HGs before 4,000 BP and individuals from Europe and Western Asia from around 10,000 BP until historical times, including Anatolian-associated (Neolithic) farmers, Caucasus HGs and recent individuals with genetic affinity to the Levant. Columns show results of modelling target individuals using three panels of increasingly distal source groups: “postBA”: Bronze Age and Neolithic source groups; “postNeol”, Bronze Age and later targets using Late Neolithic/early Bronze Age and earlier source groups; “deep”, Mesolithic and later targets using deep ancestry source groups. Individuals used as sources in a particular set are indicated with black crosses and coloured bars with 100% ancestry proportion. Black lines indicate 1 standard error for the respective ancestry component.

**Extended Data Fig. 6:**
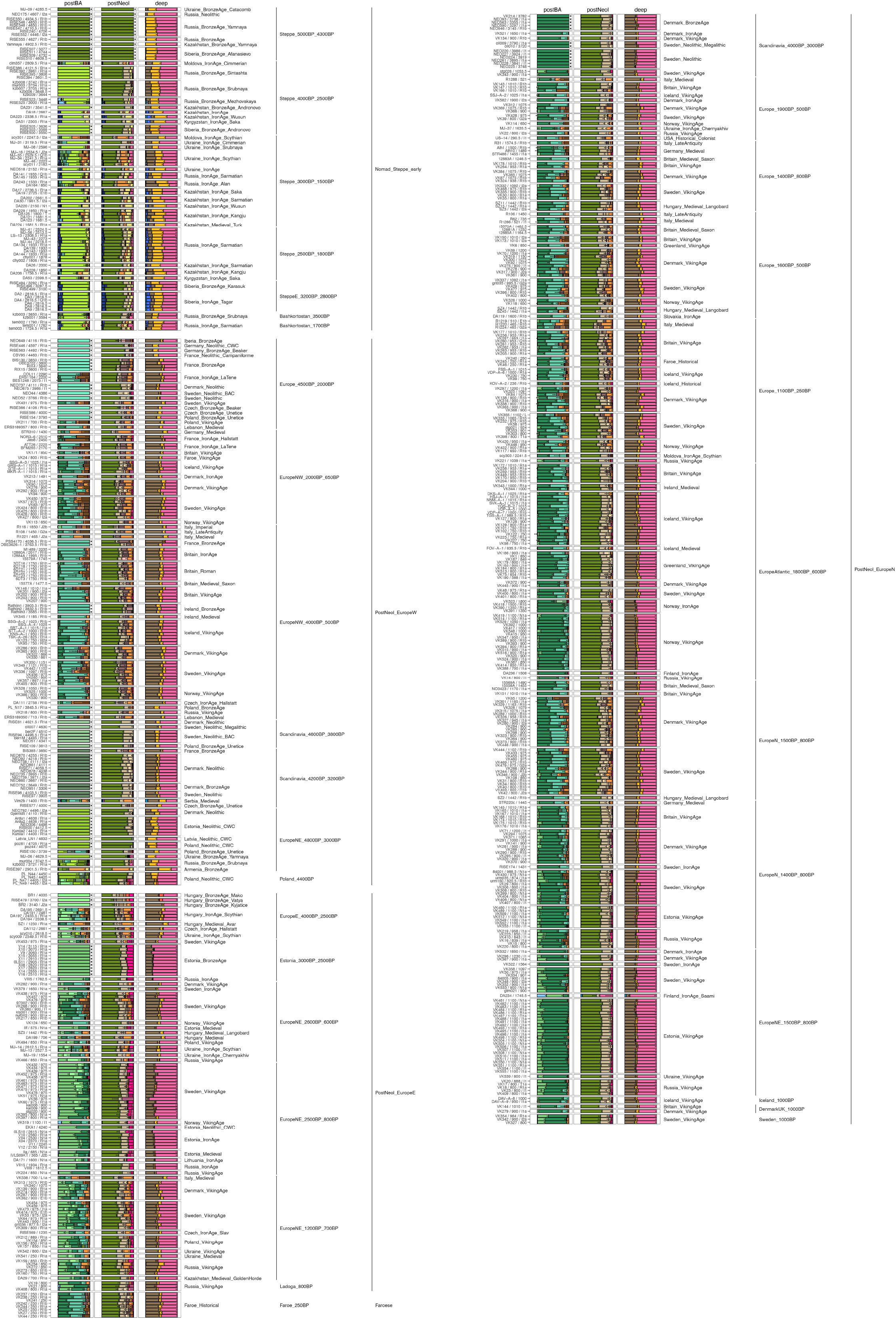
Ancestry modelling for post-Neolithic genetic clusters. Supervised ancestry modelling using non-negative least squares on IBD sharing profiles. Panels show estimated ancestry proportions of a global Eurasian cluster corresponding to European individuals after 5,000 BP, as well as pastoralist groups from the Eurasian steppe. Columns show results of modelling target individuals using three panels of increasingly distal source groups: “postBA”: Bronze Age and Neolithic source groups; “postNeol”, Bronze Age and later targets using Late Neolithic/early Bronze Age and earlier source groups; “deep”, Mesolithic and later targets using deep ancestry source groups. Individuals used as sources in a particular set are indicated with black crosses and coloured bars with 100% ancestry proportion. Black lines indicate 1 standard error for the respective ancestry component.

**Extended Data Fig. 7:**
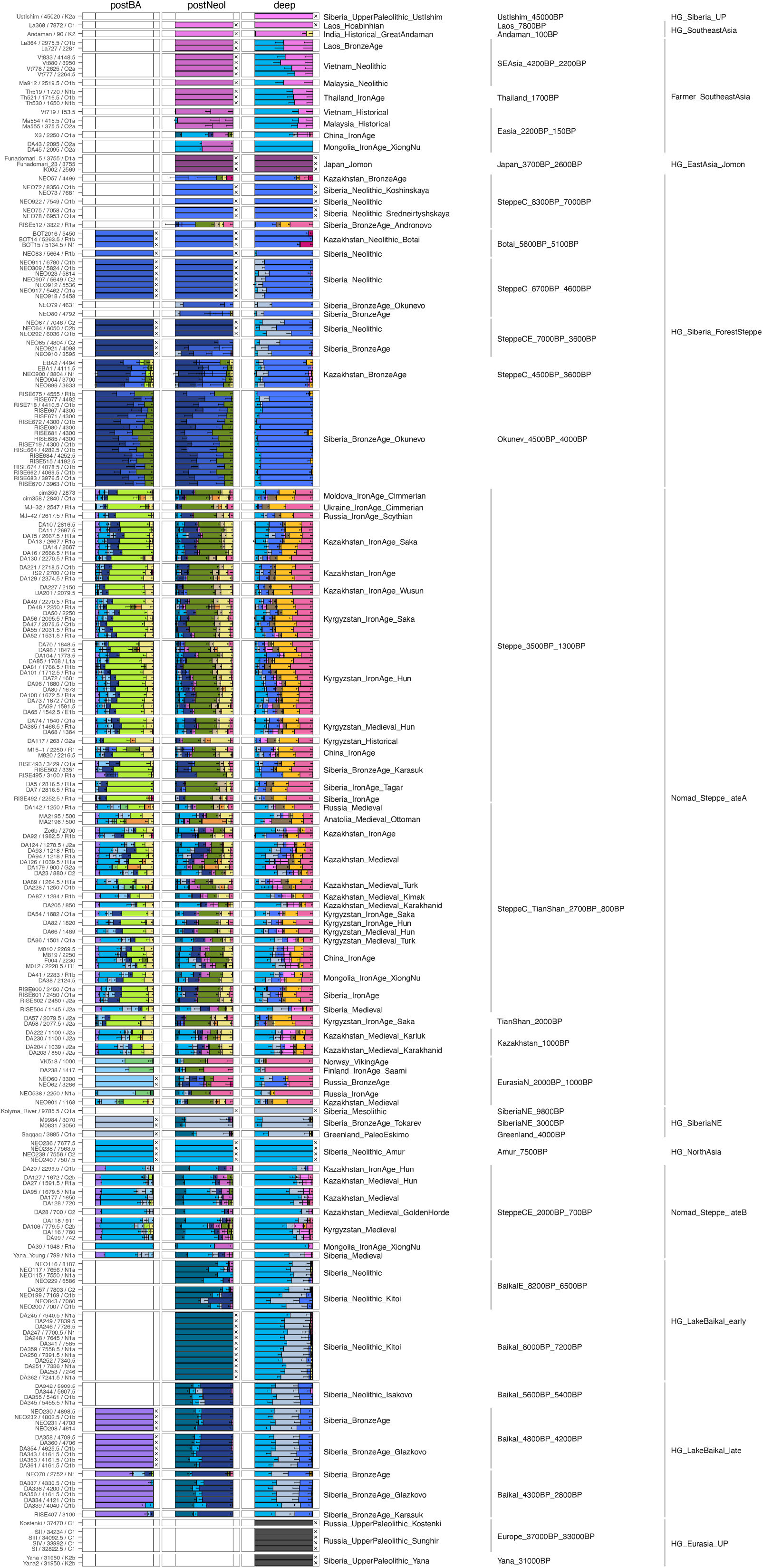
Ancestry modelling for genetic clusters east of the Urals. Supervised ancestry modelling using non-negative least squares on IBDaring profiles. Panels show estimated ancestry proportions of a global Eurasian cluster corresponding to Central, East and North Asian individuals with east Eurasian genetic affinities. Columns show results of modelling target individuals using three panels of increasingly distal source groups: “postBA”: Bronze Age and Neolithic source groups; “postNeol”, Bronze Age and later targets using Late Neolithic/early Bronze Age and earlier source groups; “deep”, Mesolithic and later targets using deep ancestry source groups. Individuals used as sources in a particular set are indicated with black crosses and coloured bars with 100% ancestry proportion. Black lines indicate 1 standard error for the respective ancestry component.

**Extended Data Fig. 8:**
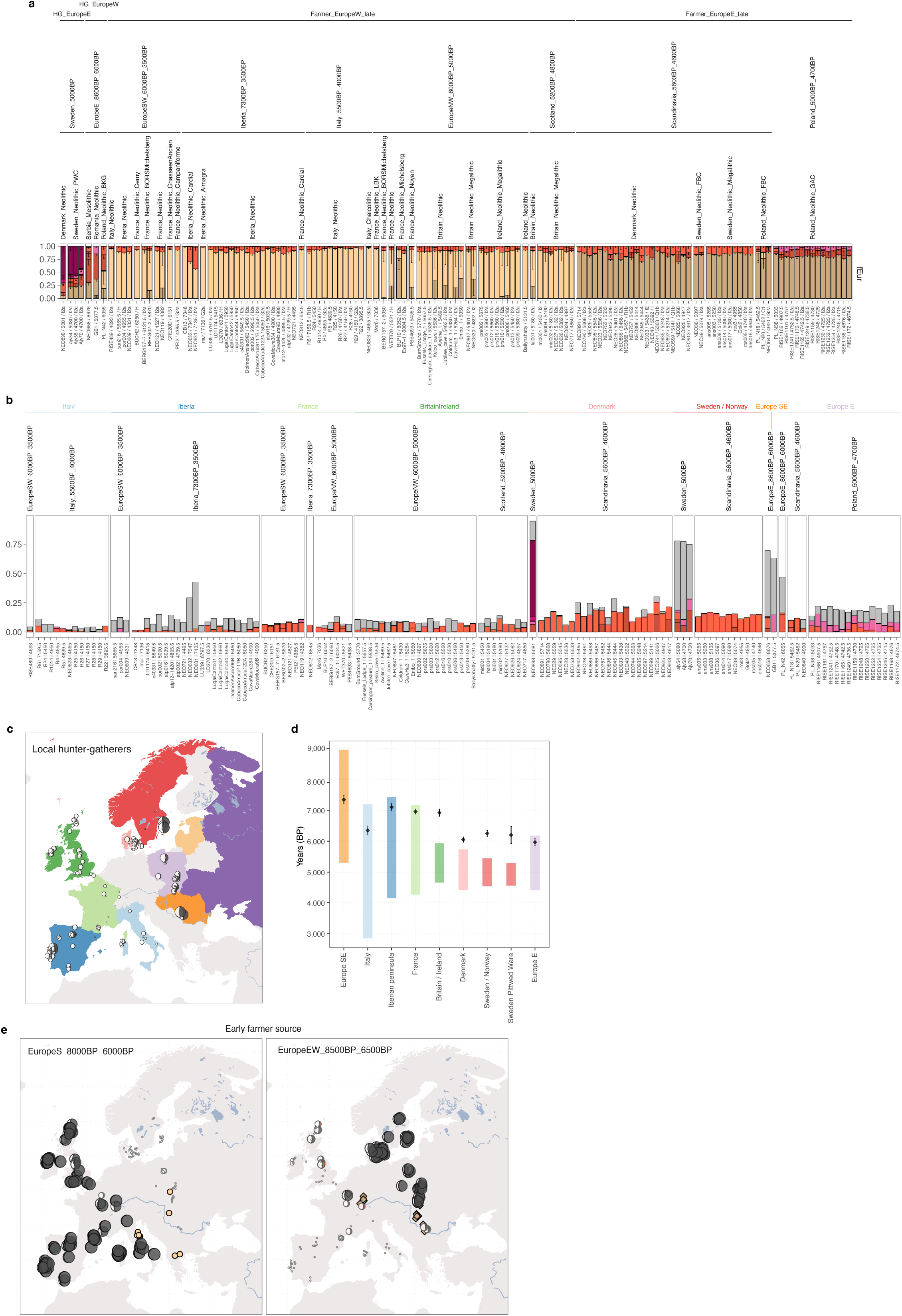
Dynamics of the Neolithic transition in Europe. (a) Supervised ancestry modelling using non-negative least squares on IBD sharing profiles. Panels show estimated ancestry proportions for target individuals from genetic clusters representing European Neolithic farmer individuals (“fEur” source set). Individuals used as sources in a particular set are indicated with black crosses and coloured bars with 100% ancestry proportion. Black lines indicate 1 standard error for the respective ancestry component. (b) Composition of HG ancestry proportions from different source groups in individuals with Neolithic farmer ancestry, shown as barplots. Grey bars represent contributions from a source with ancestry related to local HGs. (c) Moon charts showing spatial distribution of estimated ancestry proportions related to local HGs across Europe. Estimated ancestry proportions are indicated by size and amount of fill of moon symbols. Coloured areas indicate the geographic extent of individuals included as local sources in the respective regions. (d) Estimated time of admixture between local HG groups and Neolithic farmers. Black diamonds and error bars represent point estimate and standard errors of admixture time, coloured bars show temporal range of included target individuals. The time to admixture was adjusted backwards by the average age of individuals for each region. (e) Moon charts showing spatial distribution of estimated ancestry proportions derived from genetic clusters of early Neolithic European farmers (locations indicated with coloured symbols). Estimated ancestry proportions are indicated by size and amount of fill of moon symbols. Red symbols indicate individuals where standard errors exceed the point estimates for the respective ancestry source.

**Extended Data Fig. 9:**
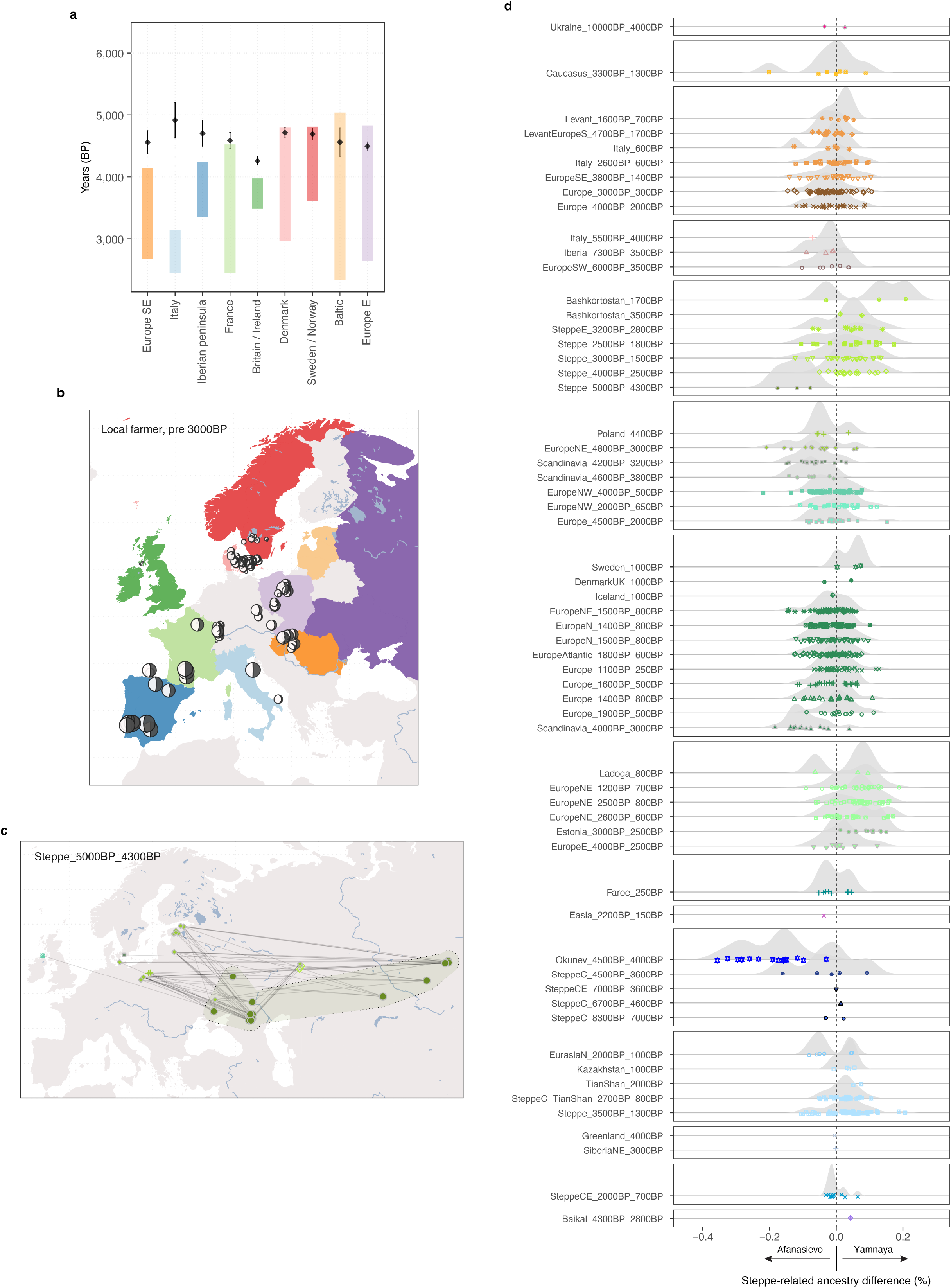
Dynamics of the steppe transition in Europe. (a) Estimated time of admixture between local hunter-gatherer groups and Neolithic farmers. Black diamonds and error bars represent point estimate and standard errors of admixture time, coloured bars show temporal range of included target individuals. The time to admixture was adjusted backwards by the average age of individuals for each region. (b) Moon charts showing spatial distribution of estimated ancestry proportions related to local Neolithic farmers across Europe. Estimated ancestry proportions are indicated by size and amount of fill of moon symbols. Coloured areas indicate the geographic extent of individuals included as local sources in the respective regions. (c) Maps showing networks of highest between-cluster IBD sharing (top 10 highest sharing per individual) for individuals from genetic cluster “Steppe_5000BP_4300BP” representing the major steppe ancestry source for Europeans. (d) Distributions of difference in estimated steppe-related ancestry proportions, using individuals from the genetic cluster “Steppe_5000BP_4300BP”, associated with either Yamnaya or Afansievo cultural contexts as separate sources.

**Extended Data Fig. 10:**
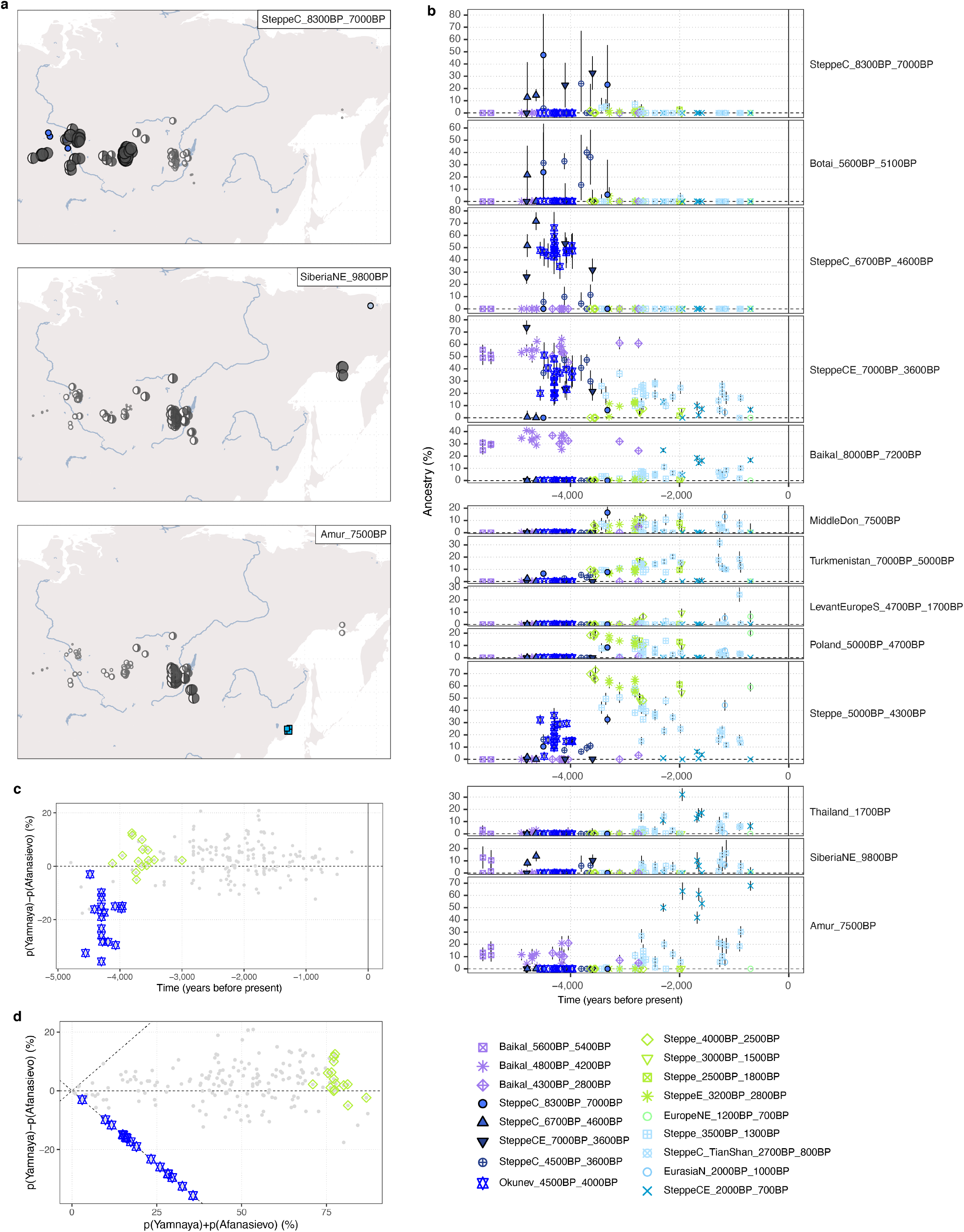
Genetic transformations across the Eurasian steppe. (a) Moon charts showing spatial distribution of estimated ancestry proportions of Siberian HGs from the “deep” Siberian ancestry sources (names and locations indicated with coloured symbols). Estimated ancestry proportions are indicated by size and amount of fill of moon symbols. (b) Timelines of ancestry proportions from “postNeol” sources in Central and North Asian ancient individuals after 5,000 BP. Symbol shape and colour indicate the genetic cluster of each individual. Black lines indicate 1 standard error. (c), (d). Difference in estimated steppe-related ancestry proportions, using individuals from genetic cluster “Steppe_5000BP_4300BP” associated with either Yamnaya or Afansievo cultural contexts as separate sources, as a function of time (c) or total estimated steppe-ancestry proportion (d). Individuals from genetic clusters of individuals associated with Okunevo (blue stars) or Sintashta/Andronovo (green diamonds) contexts are indicated with coloured symbols.

**Extended Data Fig. 11:**
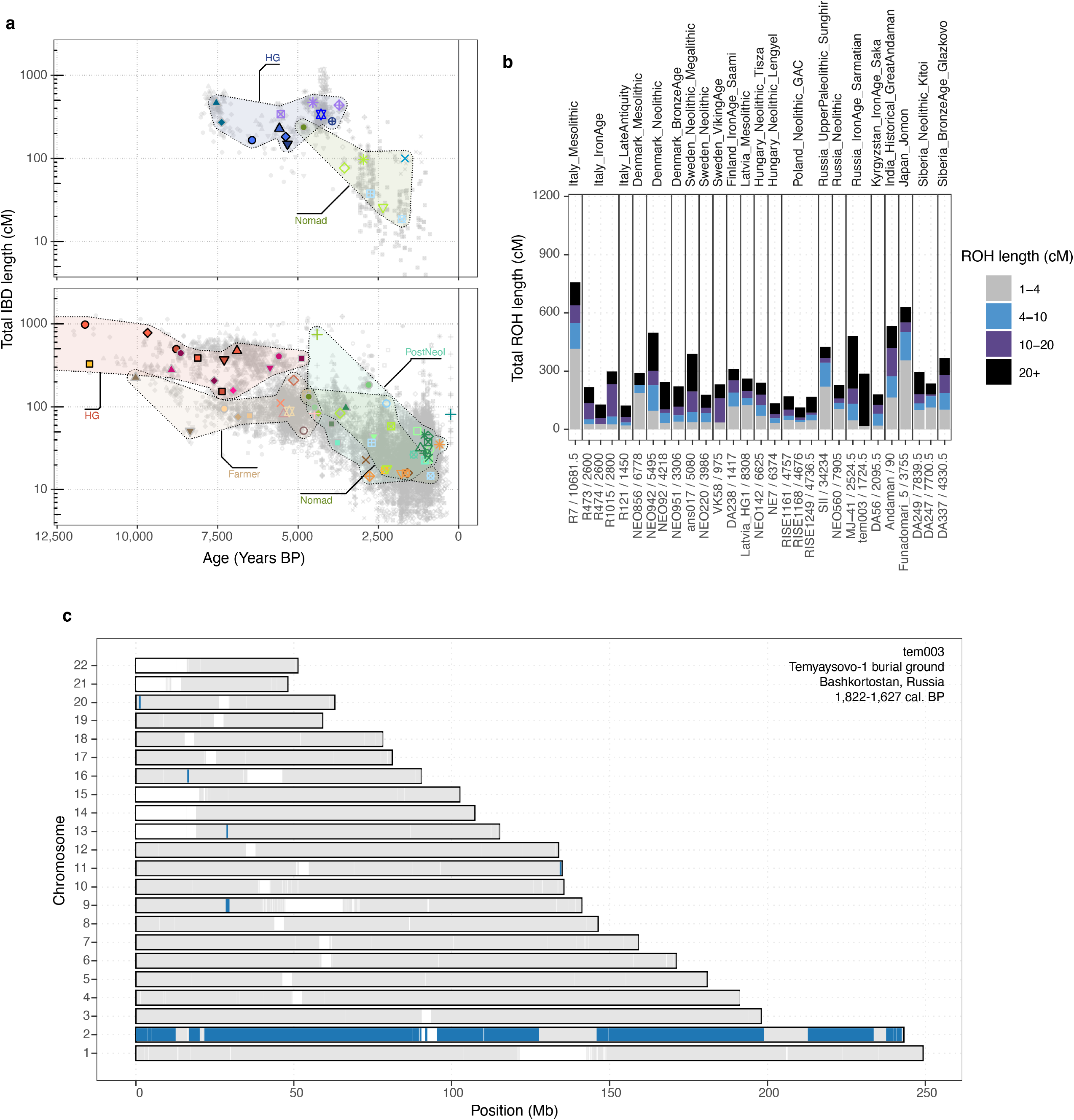
Patterns of co-ancestry. (a) Panels show within-cluster genetic relatedness over time, measured as the total length of genomic segments shared IBD between individuals. Results for both measures are shown separately for individuals from western versus eastern Eurasia. Small grey dots indicate estimates for individual pairs, with larger coloured symbols indicating median values within genetic clusters. Ranges of median values for major ancestry groups are indicated with labelled convex hulls. (b) Distribution of ROH lengths for 29 individuals with evidence for recent parental relatedness (>50 cM total in ROHs > 20 cM). (c) Karyogram showing genomic distribution of ROH in individual tem003, an ancient case of uniparental disomy for chromosome 2. Regions within ROH are indicated with blue colour.

## Data availability

All adapter-trimmed sequence data (fastq) for the samples sequenced in this study are publicly available on the European Nucleotide Archive under accession PRJEB64656, together with sequence alignment map files, aligned using human build GRCh37. The full analysis dataset including both imputed and pseudohaploid genotypes for all ancient individuals used in this study is available at https://doi.org/10.17894/ucph.d71a6a5a-8107-4fd9-9440-bdafdfe81455. Aggregated IBD-sharing data as well as hi-resolution versions of supplementary figures are available at Zenodo under accession 10.5281/zenodo.8196989. Previously published ancient genomic data used in this study is detailed in Supplementary Data VII, and are all already publicly available. Bioarchaeological data (including Accelerator Mass Spectrometry results) are included in the online supplementary materials of this submission. Map figures were created using Natural Earth Data (in Figs 1,2,3,6 and Extended Data Figs 1,3,4,8,9,10,11.).

## Code availability

All analyses relied upon available software which has been fully referenced in the manuscript and detailed in the relevant supplementary notes. A collection of R functions for IBD-based mixture model inference is available at https://github.com/martinsikora/mixmodel_ibd.

### Acknowledgements

We acknowledge Pia Bennike, involved in initiating this project, for her substantial contributions to its conception and to prehistoric research more broadly; she passed away in 2017. We thank Line Olsen and Pernille Selmer Olsen for administrative and technical assistance, respectively. We thank UK Biobank Ltd. for access to the UK Biobank genomic resource; thankfully acknowledge Illumina Inc. for collaboration. We thank Sturla Ellingvåg for assistance in sample access. E.W. thanks St. John’s College, Cambridge, for providing a stimulating environment of discussion and learning. The Lundbeck Foundation GeoGenetics Centre is supported by grants from the Lundbeck Foundation (R302-2018-2155, R155-2013-16338), the Novo Nordisk Foundation (NNF18SA0035006), the Wellcome Trust (214300), Carlsberg Foundation (CF18-0024), the Danish National Research Foundation (DNRF94, DNRF174), the University of Copenhagen (KU2016 programme), and Ferring Pharmaceuticals A/S to E.W.. This research has been conducted using the UK Biobank Resource and the iPSYCH Initiative, funded by the Lundbeck Foundation (R102-A9118 and R155-2014-1724). This work was further supported by the Swedish Foundation for Humanities and Social Sciences grant (Riksbankens Jubileumsfond M16-0455:1) to K.K. M.E.A. was supported by Marie Skłodowska-Curie Actions of the EU (grant no. 300554), The Villum Foundation (grant no. 10120) and Independent Research Fund Denmark (grant no. 7027-00147B). W.B. is supported by the Hanne and Torkel Weis-Fogh Fund (Department of Zoology, University of Cambridge); AP is funded by Wellcome grant WT214300, B.S.d.M and O.D. by the Swiss National Science Foundation (SFNS PP00P3_176977) and European Research Council (ERC 679330); R. Macleod by an SSHRC doctoral studentship grant (G101449: ‘Individual Life Histories in Long-Term Cultural Change’); G.R. by a Novo Nordisk Foundation Fellowship (gNNF20OC0062491); N.N.J. by Aarhus University Research Foundation; B.S.P. by an ERC-Starter Grant ‘NEOSEA’ (grant no. 949424); H.S. by a Carlsberg Foundation Fellowship (CF19-0601); G.S. by Marie Skłodowska-Curie Individual Fellowship ‘PALAEO-ENEO’ (grant agreement number 751349); A.J. Schork by a Lundbeckfonden Fellowship (R335-2019-2318) and the National Institute on Aging (NIH award numbers U19AG023122, U24AG051129, and UH2AG064706); A.V.L. and I.V.S. by the Science Committee, Ministry of Education and Science of the Republic of Kazakhstan (AP08856317); B.G.R. and MGM by the Spanish Ministry of Science and Innovation (Project HAR2016-75605-R); C.M.-L. and O.R. by the Italian Ministry for the Universities (grants ‘2010-11 prot.2010EL8TXP_001 Biological and cultural heritage of the central-southern Italian population through 30 thousand Years’ and ‘2008 prot. 2008B4J2HS_001 Origin and diffusion of farming in central-southern Italy: a molecular approach’); D.C.-S. and I.G.Z. by the Spanish Ministry of Science and Innovation (Project HAR2017-86262-P). D.C.S.G. acknowledges funding from the Generalitat Valenciana (CIDEGENT/2019/061) and the Spanish Government (EUR2020-112213); D.B. was supported by the NOMIS Foundation and Marie Skłodowska-Curie Global Fellowship ‘CUSP’ (grant no. 846856); E.R.U. by the Science Committee, Ministry of Education and Science of the Republic of Kazakhstan (AP09261083: “Transcultural Communications in the Late Bronze Age (Western Siberia - Kazakhstan - Central Asia)”); E.C. by Villum Fonden (17649); J.E.A.T. by the Spanish Ministry of Economy and Competitiveness, (HAR2013-46861-R) and Generalitat Valenciana (Aico/ 2018/125 and Aico 2020/97); P.K. by the Russian Ministry of Science and Higher Education (Ural Federal University Program of Development within the Priority-2030 Program) and acknowledges the Museum of the Institute of Plant and Animal Ecology (UB RAS, Ekaterinburg). L.Y. acknowledges funding by the Science Committee of the Armenian Ministry of Education and Science (Project 21AG-1F025), L.O. by ERC Consolidator Grant ‘PEGASUS’ (agreement no. 681605); M. Sablin by the Russian Ministry of Science and Higher Education (075-15-2021-1069); N.C. by Historic Environment Scotland; S.V. and E.V. by the Russian Ministry of Science and Higher Education (075-15-2022-328); V.M. by the Science Committee, Ministry of Education and Science of the Republic of Kazakhstan (AR08856925). V.A. is supported by a Lundbeckfonden Fellowship (R335-2019-2318); P.H.S. by the National Institute of General Medical Sciences (R35GM142916); S.R. by the Novo Nordisk Foundation (NNF14CC0001); R.D. by the Wellcome Trust (WT214300); R.N. by the National Institute of General Medical Sciences (NIH grant R01GM138634); F. Racimo by a Villum Fonden Young Investigator Grant (no. 00025300). T.W. and V.A. are supported by the Lundbeck Foundation iPSYCH initiative (R248-2017-2003).

## Author Information

These authors contributed equally: Morten E. Allentoft, Martin Sikora, Alba Refoyo-Martínez, Evan K. Irving-Pease, Anders Fischer, William Barrie & Andrés Ingason

These authors equally supervised research: Morten E. Allentoft, Martin Sikora, Thorfinn Korneliussen, Richard Durbin, Rasmus Nielsen, Olivier Delaneau, Thomas Werge, Fernando Racimo, Kristian Kristiansen & Eske Willerslev

### Contributions

M.E.A., M.S., A.R.-M., E.K.I.-P., A.F., W.B., and A.I. contributed equally to this work. M.E.A., M.S., T.S.K., R.D., R.N., O.D., T.W., F. Racimo, K.K. and E.W. led the study. M.E.A., M.S., A.F., C.L.-F., R.N., T.W., K.K. and E.W. conceptualised the study. M.E.A., M.S., H.S., L.O., T.S.K., R.D., R.N., O.D., T.W., F. Racimo, K.K. and E.W. supervised the research. M.E.A., L.O., R.D., R.N., T.W., K.K. and E.W. acquired funding for research. A.F., J.S., K.G.S., M.L.S.J., M.U.H., A.A.T., A.C., A.Z., A.M.S., A.J.H, A.G., A.V.L., B.H.N., B.G.R, C.B., C.L., C.M-L., D.V., D.C.-S., D.L., D.N., D.C.S.-G., D.B., E.K., E.V.V., E.R.U., E. Kannegaard, F. Radina, H.D., I.G.Z., I.P., I.V.S., J.G., J.H., J.E.A.T., J.Z., J.V., K.B.P., K.T., L.N., L.L., L.M., L.Y., L.P., L. Sarti, L. Slimak, L.K., M.G.M., M. Silvestrini, M.V., M.S.N., M.P.R., M.H.S., M.P., M.C., M. Sablin, N.C., O.P., O.R., O.V.L., P.A., P.K., P.C., P. Ríos, P. Lotz, P. Lysdahl, P.P., P.B., P.d.B.D., P.V.P., P.P.M., P.W., R.V.S., R. Maring, R. Menduiña, R.B., R.T., S.V., S.W., S.B., S.N.S., S.A.S., S.H.A., T.D.P., T.J., Y.B.S., V.I.M., V.S., V.M, Y.M., I.M., O.G. and N.L. were involved in sample collection. M.E.A., M.S., A.R.-M., E.K.I.-P., W.B., A.I., J.S., A.P., B.S.d.M., M.I., L.V., A.J. Stern, C.G., F.E.Y, D.J.L., T.S.K., R.D., R.N., O.D., F. Racimo, K.K. and E.W. were involved in developing and applying methodology. M.E.A., J.S., C.G. and L.V. led the DNA laboratory work research component. K.G.S., A.F., M.E.A. led bioarchaeological data curation. M.E.A., M.S., A.R.-M., E.K.I.-P., W.B., A.I., A.P., B.S.d.M., B.S.P., A.S.H., R.A.H., T.V., H.M., A.M., A.V., A.B.N., P. Rasmussen, G.R., A. Ramsøe, A.S., A.J. Schork, A. Rosengren, C.J.M., I.A., L.Z., R.Maring, V.S., V.A., P.H.S, S.R., T.S.K., O.D. and F. Racimo undertook formal analyses of data. M.E.A., M.S., A.R.-M., E.K.I.-P., A.F., W.B., A.I., K.G.S., R. Macleod, D.J.L., P.H.S., T.S.K., F. Racimo and E.W. drafted the main text (M.E.A. and M.S. led this). M.E.A., M.S., A.R.-M., E.K.I.-P., A.F., W.B., A.I., K.G.S., A.P., B.S.d.M., B.S.P, A.S.H., R. Macleod, R.A.H., T.V., M.F.M., A.B.N., M.U.H., P. Rasmussen, A.J. Stern, N.N.J., H.S., G.S., A. Ramsøe, A.S., A. Rosengren, A.K.O., A.B., A.C., A.G., A.V.L., A.B.G., C.J.M., D.C.S.-G., E. Kostyleva, E.R.U., E. Kannegaard, I.G.Z., I.P., I.V.S., J.G., J.H., J.E.A.T., L.Z, L.Y., L.P., L.K., M.B., M.G.M., M.V., M.P.R., M.J., N.B., O.V.L., O.C.U., P.K., P. Lysdahl, P.B., P.W., R.V.S., R. Maring, R.B., R.I., S.V., S.W., S.B., S.H.A., T.J., V.S., D.J.L., P.H.S., S.R., T.S.K., O.D. and F. Racimo drafted supplementary notes and materials. M.E.A., M.S., A.R.-M., E.K.I.-P., A.F., W.B., A.I., G.G.S., A.S.H., M.L.S.J., F.D., R. Macleod, L. Sørensen, P.O.N., R.A.H., T.V., H.M., A.M., N.N.J., H.S., A. Ramsøe, A.S., A.J. Schork, A. Ruter, A.K.O., B.H.N., B.G.R., D.C.-S., D.C.S.-G., I.G.Z., I.P., J.G., J.E.A.T., L.Z., L.O., L.K., M.G.M., P.d.B.D., R.I., S.A.S., D.J.L., I.M., O.G., P.H.S., T.S.K., R.D., R.N., O.D., T.W., F. Racimo, K.K. and E.W. were involved in reviewing drafts and editing (M.E.A., M.S., A.F., K.G.S., R. Macleod, and E.W. led this, and subsequent finalisation of the study). All co-authors read, commented on, and agreed upon the submitted manuscript.

## Footnotes

1* M.E.A., M. Sikora, A.F., K.-G.S., A.I., R. Macleod, A. Rosengren, B.S.P., M.L.S.J, Maria Novosolov, J.S., T.D.P., M.F.M., A.B.N., M.U.H., L.S., P.O.N., P.R., T.Z.T.J., A.R.-M., E.K.I.-P., W.B., W.B., A.P., B.S.d.M., F.D., R.A.H, T.V., H.M., A.V., L.V., G.R., A.J. Stern, N.N.J., A. Ramsøe, A.J. Schork, A. Ruter, A.B.G, B.H.N., E.B.P., E.K., J.H., K.B.P., L.P., L.K., M.M., M.J., O.C.U., P.L., P.B., P.V.P., R. Maring, R.I., S.W., S.A.S., S.H.A., T.J., N.L., D.J.L., S.R., T.S.K., K.H.K., R.D., F.R., R.N., O.D., T.W., K.K., E.W., 100 Ancient Genomes Show Two Rapid Population Turnovers in Neolithic Denmark. *(submitted)*

