## Supplementary Information for "Population genomics of postglacial western eurasia"

### **Supplementary Notes 1 to 7**

**Note**: High resolution versions of supplementary figures are available at Zenodo under accession 10.5281/zenodo.8196989, alongside aggregated IBD-sharing data.

#### Contents


Contents 1

**1) Data Generation and Authentication 8**

Sampling, lab work and sequencing 8

Basic bioinformatics 9

DNA authentication 10

DNA contamination - results and implications 11

References 12

**2) Imputation of ancient DNA 15**

Introduction 15

Methods 15

Results 17

Effects of Low coverage 19

References 31

**3) Demographic inference 34**

3a) Phylogenetic analysis of mtDNA sequences 34

Methods 34

Results 34

Conclusion 35

References 39

3b) Y chromosome / Sex determination 40

Methods 40

Results 40

Sex determination 40

Phylogenetic placement 41

Phylogenetics of selected haplogroups including newly reported samples 42

References 51

3c) Relatedness 52

Results 52

References 54

3d) Overall Population Structure 56

Results 56
Comparison of imputed and pseudo-haploid genotypes in PCA space 56
PCA position of samples flagged as contaminated 61
Genetic ancestry of newly reported samples 62
Admixture graph fitting of deep Eurasian genomes 69
References 71
3e) Inferring the spatiotemporal spread of population movements in the past 13 millennia 73
Introduction 73
Methods 73
Results - ADMIXTURE ancestry proportions 74
Results - IBD cluster proportions 74
Figures 75
References 86
3f) HBD/ IBD sharing/ROH/clustering 88
Results 88
IBD-based hierarchical graph clustering 88
Runs of homozygosity and IBD sharing within clusters 115
Mixture models 118
References 124
**4) 14C chronology and estimates of reservoir effects 126**
Reservoir effects 127
References 129
**5) From forager to farmer in western Eurasia: an archaeological overview 136**
Introduction 136
The cradle of European agriculture - southwestern Asia 137
The later Stone Age of western and central Europe 138
The later Stone Age of eastern Europe and western Siberia 141
References 143
**6) Catalogue of Danish archaeological sites 150**
Introduction 150
The structure of the catalogue 153
Acknowledgements 153
Avlebjerg, NEO961; Strøby 05.06.12-8A, Zealand. Stone cist 155
Barhøj, NEO92; Strøby 05.06.12-22, Zealand. Stone cist 155
Bjørnsholm, NEO751; Ranum 12.07.10-20, North Jutland. Shell midden with inhumation 156
Bodal K (Knoglebo), NEO814; Stenlille 03.03.09-176, Zealand. Settlement and unassociated human remains 156
Borreby, NEO735+737; Magleby 04.04.11-45, Zealand. Passage grave 156
Bybjerg, NEO563; Orø 03.07.09-34, off Zealand. Mesolithic kitchen midden with Bronze Age burials 158
Bygholm Nørremark, NEO564; Hatting 17.04.03-128, East Jutland. Earthen long barrow 158
Dalmosegaard, NEO886; Borre 05.05.02-95, Møen. Bog skeleton 160
Dragsholm, NEO732+733+822+962b; Fårevejle 03.04.03-503, Zealand. Dwelling site with a Mesolithic and a Neolithic burial, and a loose human bone 160
Døjringe, NEO566; Munke-Bjergby 04.01.08-109, Zealand. Wetland with two bog skeletons 161

Elkenøre, NEO888; Idestrup 07.02.05-91, Falster. Bog skeleton 161
Ertebølle, NEO568+569; Strandby 12.02.12-63, North Jutland. Kitchen midden with inhumations and loose human bones 162
Fannerup D, NEO855; Ginnerup 14.01.05-71, East Jutland. Kitchen midden with inhumation 163
Fannerup E, NEO570; Ginnerup 14.01.05-72, East Jutland. Kitchen midden with inhumation 164
Fannerup F, NEO930; Ginnerup 14.01.05-107, East Jutland. Kitchen midden with inhumation 164
Gammellung (Troldebjerg), NEO934; Lindelse 09.03.04-98, Langeland. Ceremonial bog deposition 164
Grøfte, NEO571; Kindertofte 04.03.06-3, Zealand. Megalithic monument with two dolmen chambers 165
Havnø, NEO941; Visborg 12.04.13-(45?), North Jutland. Human mandible from a kitchen midden 166
Hedegaard, NEO13; Bislev 12.05.01-113, North Jutland. Wetland with human skull 166
Henriksholm-Bøgebakken, NEO745+746+747+748+749; Søllerød 02.03.10-157, Zealand. Mesolithic dwelling site with burials 166
Holmegaard-Djursland, NEO1; Hyllested 14.02.07-24, East Jutland. Kitchen midden with inhumation 167
Hove Å (Gundsømagle Mose), NEO946; Hvedstrup 02.04.06-29, Zealand. Wetland with bog skeletons 168
Jorløse Mose (Jordløse Mose), NEO23; Jorløse 03.06.06-192, Zealand. Wetland with bog skeleton 168
Jørlundegård, NEO702; Jørlunde 01.03.06-85, Zealand. Wetland with bog skeleton 168
Kainsbakke, NEO25; Ginnerup 14.01.05-118, East Jutland. Ritual pit at coastal settlement 169
Klokkehøj, NEO580; Horne 09.04.12-2, Funen. Dolmen 170
Klæstrupholm Mose, NEO951; Jerslev 10.01.06-269, Vendsyssel. Wetland with bog skeleton 171
Koed, NEO583+586; Koed 14.10.07-37, East Jutland. Kitchen midden and inhumations 172
Koelbjerg, NEO254; Vissenbjerg 08.04.15-78, Funen. Wetland with bog skeleton 172
Kolind, NEO738+739; Kolind 14.02.08-13, East Jutland. Settlement deposits with stray human bones 173
Kongemose, NEO587; Stenmagle 04.01.12-232, Zealand. Settlement and stray human skeletal remains 173
Korsør Nor (Korsør Glasværk), NEO589+791; off Korsør 401433-17, Zealand. Habitation site with inhumations 174
Kyndeløse (Møllehøj), NEO878; Kirke Hyllinge 02.06.05-6, Zealand. Passage grave 174
Køge Sønakke, NEO759; 401379-38, east of Zealand. Human skeleton from the sea floor 175
Langø Skaldynge, NEO853; Stubberup 08.01.12-(51?), Funen. Human skull from shell midden 175
Lohals, NEO29; Hou 09.02.02-17, Langeland. Shell midden with inhumation 176
Lollikhuse, NEO857; Selsø 01.02.06-77, Zealand. Kitchen midden with stray human remains 176
Lundby-Falster, NEO865+866; Brarup 07.01.01-28, Falster. Wetland with bog skeletons 176
Læsten Mose (Volstrup Mose), NEO945; Ålum 13.12.15-(71?) North Jutland. Wetland with bog skeleton 177
Madesø, NEO752; Jorløse 03.06.06-53, Zealand. Wetland with bog skeleton 177

Magleø, NEO590; off Korsør 401433-6, Zealand. Stray Bronze Age skeleton on Mesolithic habitation site 178
Mandemarke, NEO896; Magleby 05.05.07-375, Møen. Wetland with bog skeleton 179
Mosede Mose (Karlslunde Mose), NEO860+861; Karlslunde 02.05.04-9, Zealand. Wetland with bog skeletons 179
Myrebjerg Mose, NEO925; Magleby 09.03.06-78, Langeland. Ceremonial wetland deposition 179
Nederst, NEO856; Albøge 14.02.01-49, East Jutland. Settlement with shell middens and inhumations 180
Neverkær Mose, NEO594; Vissenbjerg 08.04.15-24, Funen. Wetland with bog skeleton 180
Norsminde, NEO852; Malling 15.04.05-71, East Jutland. Kitchen midden with burials and loose human bones 180
Næs, NEO792; Aastrup 07.02.14-(45?), Falster. Megalithic tomb 181
Orehoved Sejlrende, NEO122+123; 401651-33, south of Zealand. Submerged settlement with stray human remains 182
Pandebjerg, NEO595; Føllenslev 03.06.04-76, west of Zealand. Kitchen midden with stray human bones 182
Porsmose, NEO795; Toksværd 05.04.10-15, Zealand. Wetland with bog bog skeleton 183
Ravnsbjerggård II (Rygård), NEO960; Undløse 03.03.18-54, Zealand. Settlement site with bog deposition 184
Roskilde Fjord (south of Jyllinge), NEO891; 401256-78 off Zealand. Stray human bone from shell bank 184
Rude, NEO41+043; Saksild 15.02.12-6, East Jutland. Long barrow with two stone chambers 184
Rødhals (Sejerø), NEO8; Sejerø 03.06.07-49, off Zealand. Kitchen midden with a burial and a stray human skull 185
Rønsten, NEO19; 401275-1, off East Jutland. Settlement with stray human bone 186
Salpetermose, NEO28; Hillerød 01.03.01-180A, Zealand. Wetland with bog skeleton 186
Sejerby, NEO757; Sejerø 03.06.07-36B, off Zealand. Burial 187
Sigersdal, NEO7; Stenløse 01.06.05-110, Zealand. Wetland with bog skeletons 187
Sigersdal Mose, NEO753; Stenløse 01.06.05-99, Zealand. Wetland with bog skeleton 188
Sludegaard Sømose, NEO933; Frørup 09.06.06-34A, Funen. Wetland sacrificial site with bog skeletons 188
Stenderup Hage, NEO943; South Jutland 401513c-1. Submerged settlement and stray human bones 189
Storelyng Eel Picker (Øgaarde boat III), NEO597; Undløse 03.03.18-331, Zealand. Wetland with bog skeleton 189
Storelyng Fire Lighter (Østrup homo II), NEO602; Undløse 03.03.18-26B, Zealand. Wetland with bog skeleton 190
Strøby Grøftemark, NEO91; Strøby 05.06.12-54, Zealand. Wetland with bog skeleton 190
Strøby Ladeplads, NEO93; Strøby 050612-31, Zealand. Stone cist 190
Svinninge Vejle, NEO898; Svinninge 03.07.12-91, Zealand. Reclaimed fjord with stray human bone 191
Sølager, NEO598; Torup 01.05.09-7, Zealand. Kitchen midden with stray human bones 191
Tissøe, NEO942; Buerup 03.02.02-219, Zealand. Lake deposition (?) 192
Toftum Mose, NEO870+872+875+876; Søvind 16.05.08-17, East Jutland. Wetland with bog skeletons 192
Tudse Hage, NEO932; 401441-11, west of Zealand. Submarine settlement site with stray human remains 193

Tybrind Vig, NEO683; 401521-6, west of Funen. Submerged settlement with burials 193
Tysmose, NEO790; Ledøje 02.02.09-65, Zealand. Wetland with bog skeletons 194
Vanløse Mose, NEO599; Stenmagle 04.01.12-418, Zealand. Wetland with bog skeletons 194
Vasagård, NEO815; Åker 06.02.05-58, Bornholm. Passage grave 194
Vedbæk Boldbaner, NEO600; Søllerød 02.03.10-135, Zealand. Settlement with burials and stray human bones 195
Vibygårds Mose, NEO935; Syv 02.01.10-17, Zealand. Wetland with bog skeleton 196
Vig Femhøve, NEO744; Vig 03.04.12-150, Zealand. Megalithic monument with a dolmen chamber 196
Viksø Mose (Rolandsgårdens Mose), NEO601; Veksø 01.06.06-42, Zealand. Wetland with bog skeleton 197
Vittrup, NEO33; Børglum 10.01.02-56, Vendsyssel. Wetland with bog skeleton 198
Vængesø II, NEO3; Helgenæs 14.05.06-39, East Jutland. Kitchen midden with burials and loose human bones 198
References 199
**7) Catalogue of non-Danish archaeological sites 209**
Armenia 211
Aknashen, Armavir, Armenia. Settlement and burials. 211
Czech Republic 211
Vedrovice, Moravia, Czech Republic. Cemeteries and settlement. 211
England 212
Hetty Pegler’s Tump (Uley Barrow), Gloucestershire, England. Long barrow. 212
Estonia 213
Sope 213
France 214
Grotte Gazel, Aude, France. Double burial. 214
Grotte Mandrin, Drôme, Rhône-Alpes, France. Rock shelter. 215
Georgia 216
Kotias Klde, Chiatura, Georgia. Cave site. 216
Hungary 217
Deszk – Olajkút 217
Hödmezövasarhely Kotac 217
Gorzsa Cukormajor, region Csongrád, Hungary 218
Iran 220
Tepe Guran, Hulailan Valley, Central Zagros, Iran, grave/pit find 220
Italy 221
Fontenoce, Recanati, Italy 221
Gaudo, Paestum, Italy 221
Grotta delle Mura, Monopoli, Bari, Italy 223
Grotta Nisco, Cassano Murge, Bari, Italy, Cave 224
Maddalena di Muccia A, Province of Macerata, Marche Region, Italy. Settlement. 224
Mora Cavorso cave, Jenne, Rome, Italy. Cave. 225
Kazakhstan 226
Gregorievka 1, Pavlodar, Kazakhstan. Cemetery. 226
Shauke settlement, Pavlodarsk, Kazakhstan. Settlement and burial 228
Shauke 1, Pavlodarsk, Kazakhstan. Cemetery. 228
Sjiderty 10, Pavlodarsk, Kazakhstan. Barrows. 230

Bestamak, Kazakhstan. Cemetery. 231
Latvia 232
Zvejnieki, Latvia. Cemetery and settlement. 232
Norway 233
Hummervikholmen, Søgne, Agder, Norway. Submarine find. 233
Poland 234
Słonowice, Kazimierza Wielka district, site 5, Poland. Cemetery (long barrows) 234
Portugal 235
Gruta do Caldeirão, Tomar, Portugal. Cave site. 235
São Paulo II, Almada, Lisbon, Portugal. Hypogeum. 235
Romania 236
Schela Cladovei, Danube Gorges, Romania. Settlement with burials. 236
Băile Herculane – Peştera Hoţilar, Banat, Romania. Cave. 236
Russia 237
Afontova Gora, Krasnoyarsk, Southern Siberia, Russia. Flat grave. 237
Bazaiha village, Krasnoyarsk district, Southern Siberia, Russia. Cemetery. 238
Borovyanka XVII, Omsk region, Russia. Cemetery. 238
Dolgoe Ozero (Long lake), Krasnoyarsk, Russia. Settlement and graves. 239
Fofonovo, Baikal region, Russia. Cemetery. 240
Golubaya Krinitsa, Middle Don, Russia. Cemetery. 241
Itkul (Bol’shoy Mys), Altai region, Russia. Cemetery. 242
Karavaikha, Vologda region, Russia. Cemetery. 243
Kostenkova Izbushka, Altai region, Russia. Cemetery. 244
Kumyshanskaya cave, Ural, Russia 245
Ksizovo 6, Lipetsk, Russia. Settlement and burial ground. 246
Mergen’ 6, Tumen region, Russia. Settlement 247
Minino I and II, Vologda region, Russia. Cemetery. 247
Okunevo 5 and 7, Omsk region, Russia. Cemetery. 249
Omskaya Stoyanka (Omsk settlement), Russia. Settlements and cemeteries. 250
Ostrov 2, Tumen region, Russia. Cemetery. 251
Pad’ Tokui, Krasnochikoysky region, Russia. Cemetery. 251
Peschanitsa, Archangelsk region, Russia. Burial(s). 252
Pogostishche I, Vologda region, Russia. Cemetery. 252
Protoka, Novosibirsk region, Russia. Cemetery. 253
Sakhtysh, Ivanovo region, Russia. Settlements and cemeteries. 254
Sosnovyi Mys, Angara valley, Russia. Settlement and cemetery. 258
Ural River, Orenburg region, Cis-Urals, Russia. River beach find. 259
Ust’-Isha, Altai region, Russia. Cemetery. 260
Vasilyevsky Cordon 17, Lipetsk, Russia. Settlement and burial ground 260
Vengerovo-2A, Novosibirsk Region, Russia 261
Zamostje 2, Sergiev-Posad district, Moscow region, Russia. Settlement. 262
Zjindo, Krasnochikoysky region, Russia. Cemetery. 263
Scotland 264
Banks Chambered Tomb, South Ronaldsay, Orkney, Scotland 264
Serbia 265
Vlasac, Danube Gorges, Serbia. Settlement with burials. 265
Lepenski Vir, Danube Gorges, Serbia. Settlement with burials. 266

Spain 267
Camino de las Yeseras, San Fernando de Henares, Madrid, Spain. Chalcolithic Tomb, Área 15, El-08-I 267
El Toral III, Andrín, Asturias, Spain. Rock shelter. 268
El Mazo, Andrín, Asturias, Spain. Rock shelter. 269
Coves de Santa Maira, Castell de Castells, Alacant, Spain. Cave. 270
Sweden 270
Bredgården, RAÄ 113 Marbäck sn, Västergötland, Sweden. Wetland find. SHM 33241. 270
Dösemarken, Scania, Sweden. Grave and settlement. 271
Evensås, Skaftö 85, Bohuslän, Sweden. Flat graves. 272
Fredriksberg, Falköping stad 5, Västergötland, Sweden. Gallery grave. 273
Frälsegården, Gökhem 94, Västergötland, Sweden. Passage grave. 274
Hanaskede, N Ving sn, Västergötland, Sweden. Wetland find. VGM 106100. 275
Hindby mosse, MHM 1505, Scania, Sweden. Central place. 275
Kastanjegården, Scania, Sweden. Cemetery. 276
Sillvik, Torslanda 43, Bohuslän, Sweden. Flat grave. 277
Skateholm I, Scania, Sweden. Cemetery. 278
Vattenledningen, Scania, Sweden. Cemetery and settlement. 278
Ängdala, Scania, Sweden. Grave and flint mines. 279
Turkmenistan 280
Monjukly Depe. Settlement. 280
Ukraine 281
Igren 8. Cemetery. 281
Kleshnya III, the Seversky Donets River, Ukraine. Burial in a settlement. 282
Lysa Gora, Lower Dnepr, Ukraine. Burial ground. 283
Mamai Gora, Lower Dnieper, Ukraine. Burial ground. 284
Vasilevka I, Dnepr river valley, Ukraine. Cemetery. 285
Vovnigi I and Vovnigi II, Dnepr river valley, Ukraine. Cemeteries. 286
Voloshskoe, Dnepr river valley, Ukraine. Cemetery. 287
Volnensky, Dnepr river valley, Ukraine. Cemetery. 288
References 289

### 1) Data Generation and Authentication

Morten E. Allentoft[1,2], Simon Rasmussen[3], Gabriel Renaud[4], Abigail D. Ramsøe[2], Thorfinn Korneliussen[2], Martin Sikora[2]

[1]Trace and Environmental DNA (TrEnD) Lab, Curtin University, Perth, Australia
[2]Lundbeck Foundation GeoGenetics Centre, Globe Institute, University of Copenhagen, Copenhagen, Denmark
[3]Novo Nordisk Foundation Center for Protein Research, University of Copenhagen, Copenhagen, Denmark
[4]Department of Health Technology, Section of Bioinformatics, Technical University of Denmark, Kongens Lyngby, Denmark

#### Sampling, lab work and sequencing

The lab work component of this project followed the same procedures outlined in Allentoft et al.[1] and Damgaard et al.[2], but also included sampling of the petrous part of the temporal bone following the discovery of exceptional DNA preservation in these bones[3,4]. While new ancient DNA (aDNA) extraction and library methods are continually optimised and presented in the literature, we prioritised method consistency throughout the project period to avoid the risk of introducing batch effects in the data.

A total of 962 later Stone Age and early Bronze Age human skeletons from across Eurasia were sampled for this project. An initial molecular 'screening' to assess the endogenous DNA content (proportion of DNA sequences identified as human) was performed by shallow shotgun sequencing resulting in 317 samples (Supplementary Data I) being selected for deeper sequencing. We applied a threshold of <1% endogenous DNA for rejecting samples in the project, except for a few Danish skeletons that were prioritised despite displaying even lower contents. Of the 317 samples, 211 were teeth, 91 were petrous bones, and 15 were pieces of other types of bones (long bones, ribs, cranial bones).

All the pre-PCR-amplification lab work was conducted in dedicated clean laboratories at the Lundbeck Foundation GeoGenetics Centre (Globe Institute, University of Copenhagen), according to strict aDNA guidelines[5–7]. To reduce the amount of non-target DNA in the extracts, the outermost surfaces of the samples were first removed using a sterile cutting disc. Teeth were processed by separating the crown from the root by a cutting disc and the inner dentine was then removed from the root with a pointy drilling bit. By this procedure each root sample was proportionally enriched for the outer cementum layer which is known for its high endogenous DNA content[2,8,9]. Petrous bones were sampled by cutting off slices (with a cutting disc) until reaching the dense otic capsule which was used for DNA extraction. The samples were crushed into smaller pieces before the lysis step.

To further increase the endogenous DNA yield, we performed a brief ‘pre-digestion’ step prior to the extraction protocol following Damgaard et al.[2]. After this pre-digestion, we added 3.5mL of fresh digestion buffer to each sample and incubated them for 24h before the DNA was purified with silica-in-solution similar to Rohland & Hofreiter[10] but using the optimised binding buffer from Allentoft et al.[1]. Double-stranded blunt-end libraries were constructed from the extracted DNA using NEBNext DNA Prep Master Mix Set E6070 (New England Biolabs Inc.) with protocol modifications[11] and then amplified with indexed Illumina-specific adapters prepared as in[12]. The DNA concentration of each amplified library was quantified on an Agilent 2200 Tapestation and sequencing (80bp and 100bp single read) was performed on Illumina HiSeq 2500 and Illumina HiSeq 4000 platforms at the Danish National High-throughput DNA Sequencing Centre.

#### Basic bioinformatics

The Illumina data were base-called using Illumina software *CASAVA (v.1.8.2)*[13] and sequences were de-multiplexed with the requirement of full matching of the six nucleotide index which was used for library preparation. Adapter sequences and leading/trailing stretches of Ns were trimmed from the reads and bases with quality 2 or less were removed using *AdapterRemoval (v.2.1.3)*. Trimmed reads of at least 30bp were mapped using *bwa (v.0.7.10)*[14] with the seed disabled to allow for higher sensitivity[15]. Reads were mapped to the human reference genome build 37 including mitochondrial DNA (rCRS) and to mitochondrial DNA alone. Mapped reads were filtered for mapping quality 30 and sorted using *Picard (v.1.127)* (http://picard.sourceforge.net) and *samtools*[16]. Data was merged to library level and duplicates removed using *Picard MarkDuplicates (v.1.127)* and hereafter merged to sample level. Sample level BAMs were re-aligned using *GATK (v.3.3.0)* and hereafter had the md-tag updated and extended BAQs calculated using *samtools calmd (v.1.10)*[16]. Read depth and coverage were determined using *pysam* (https://github.com/pysam-developers/pysam) and *BEDtools (v.2.23.0)*[17].

#### DNA authentication

To investigate the authenticity of the ancient DNA molecules, post-mortem DNA damage patterns were determined using *mapDamage2.0*[18]. Cytosine deamination was recorded for each sample as the fraction of C-to-T transitions at the first 5'position of the DNA reads when compared to the reference genome. For the 317 samples included here, we observed C-to-T deamination fractions ranging from 10.4% to 67.8%, with an average of 38.3% across all 317 samples (Supplementary Data I). These numbers generally reflect highly damaged molecules as expected for DNA that is thousands of years old.

Next, we applied three different methods to estimate levels of DNA contamination; two based on mitochondrial genome data and one method investigating X-chromosomal data in males. All contamination estimates are reported in Supplementary Data V with summary values provided in Supplementary Data I.

First, estimates of present-day human contamination for the mitochondrial genome were performed using the iterative Bayesian framework implemented in *Schmutzi*[19]. Briefly, ancient DNA sequences were realigned to the mitochondrial genome of the revised Cambridge Reference Sequence (rCRS, NCBI Reference Sequence: NC_012920.1) using *BWA*[14] with parameters for increased sensitivity (*-n 0.01 -o 2 -l 16500*). The mapping was performed exclusively to the mitochondrial genome to mitigate the impacts of nuclear NUMTs on the mitochondrial alignments. The resulting BAM file was used as input for *Schmutzi* using five iterations and by subsampling samples with coverage to 500X, should they exceed that[19]. The iterative approach was run using a database of Eurasian mitogenomes. The point estimate for the final contamination rate with the maximum *a posteriori* probability is reported in the sample summary Supplementary Data I, whereas the 95% confidence interval (lower and upper bound as well as the point estimate) are reported in Supplementary Data V.

Next, we applied *ContamMix* in order to estimate the fraction of non-endogenous reads in the mitochondrial genome by comparing the reconstructed mtDNA consensus sequence to 311 possible contaminant genomes[20]. For each sample, an in-house perl script was used to construct two different versions of the endogenous mitochondrial genome. The first approach (CONTAMIX_APPROX_1Xdif05) used sites with at least 1x coverage, and at each position a base was only called if it was observed in at least 50% of reads covering the site. The second approach (CONTAMIX_PRECISE_5Xdif07) only considered sites with at least 5x coverage and 70% of reads agreeing. Both approaches used reads with a base quality of ≥20 and mapping quality of ≥30.

Lastly, we applied *ANGSD* [21] on X-chromosomal data in males. This approach quantifies heterozygosity on the X chromosome. As males only have one copy of the X chromosome, any heterozygosity is expected to arise from either contamination or sequencing error. As heterozygosity due to contamination is expected to be restricted to mainly known diagnostic polymorphic sites, *ANGSD* quantifies the heterozygosity in these sites. It then compares it to adjacent sites in order to ascertain the level of background sequencing error, and thus estimates the extent of contamination. For each sample, we removed the pseudoautosomal regions on the X chromosome and filtered out reads with a base quality <20 and mapping quality <30.

#### DNA contamination - results and implications

Supplementary Data V lists all the contamination estimation results for the 317 samples across the three applied methods. The vast majority of the samples show very low levels of contamination (≤ 5%) across all methods. A total of 33 samples, however, display contamination estimates >5% in one or more of the methods, but there are considerable inconsistencies between the methods (Supplementary Data V). It is well established that contamination estimates are not reliable for very low coverage genomic data[5–7] and this is further complicated by DNA damage in the sequences. Indeed, we observe that the 33 potentially problematic samples have an average coverage of 0.11X and a median of 0.06X which is considerably lower than the full dataset with its average coverage of 0.75X and median of 0.26X. So, instead of simply excluding data from these precious samples based on estimates that are likely imprecise, we “flagged” samples as potentially contaminated in downstream analyses and took a more analytical approach in the evaluation. Flagging was applied as follows:

**Samples with nuclear coverage <0.1X and MT coverage <10X:** Not flagged, since both estimates are likely unreliable

**Samples with nuclear coverage <0.1X and MT coverage ≥10X:** Flagged as possibly contaminated if any MT estimate is >5%; ignore nuclear estimate as likely unreliable

**Male samples with nuclear coverage ≥0.1X:** Flagged as possibly contaminated if any nuclear (X-chromosomal) estimate is > 5%; ignore MT as only nuclear data are relevant for genome-wide analyses

**Female samples with nuclear coverage ≥0.1X:** Flagged as possibly contaminated if any MT estimate is >5% as no nuclear estimate is available

Based on this approach we have a total of 15 samples (NEO1, NEO3, NEO76, NEO77, NEO158, NEO162, NEO168, NEO221, NEO226, NEO537, NEO657, NEO671, NEO677, NEO746, NEO815) that we have flagged as "possibly contaminated" (See Table S2) in our downstream analyses. The further analytical evaluation of these samples is described in Supplement Note S3d.

### 2) Imputation of ancient DNA

Bárbara Sousa da Mota[1,2], Andrew Vaughn[3] & Olivier Delaneau[1,2]

[1] Department of Computational Biology, University of Lausanne, Switzerland
[2] Swiss Institute of Bioinformatics, University of Lausanne, Switzerland
[3] Center for Computational Biology, University of California, Berkeley, USA

#### Introduction

GLIMPSE is a statistical method developed to impute low-coverage human genomes. It has been shown that GLIMPSE efficiently produces accurate results when employed to impute low-coverage present-day genomes. Here we seek to demonstrate that GLIMPSE is also a suitable imputation tool for low-coverage ancient genomes. To benchmark it, we used a subset of 42 previously published ancient genomes with a mean depth of coverage above 10x (sample list is given in Table S2.1) that we downsampled to lower coverages in order to match the coverage we observed for the 317 genomes sequenced in the present study. Then, we imputed the resulting genomes and assessed the accuracy of the imputed calls by comparing with the original high-coverage genomes. Specifically, we examined the imputation performance regarding (i) depth of coverage, (ii) minor allele frequency, (iii) ancestry and living period of target samples. We also imputed the samples with the present 'gold standard' method, Beagle4.1[1], to show how it compares with GLIMPSE v1.0.1[2] (https://odelaneau.github.io/GLIMPSE/). Finally, we obtained D-statistics to ascertain the effect of imputation upon pairwise comparisons. Here we specifically tested for bias introduced by the reference panel used for imputation affecting signals of WHG ancestry, using the Loschbour[3] genome to represent WHG ancestry.

#### Methods

We first prepared all necessary files for the imputation step. We used samtools 1.10 to downsample the 42 high-coverage genomes to coverages 0.1x, 0.2x, 0.4x, 0.8x,1.0x, 2.0x and 4.0x. Then, we prepared a list of candidate variant sites at which imputation was performed by retaining all sites in 1000 Genomes version 3 that were (i) bi-allelic SNPs and (ii) non-singleton (i.e. informative for imputation). For each of the seven tested coverages, we computed genotype likelihoods (VCF/PL field) at all candidate variant sites across all target samples using bcftools 1.10. To minimise computation time, we restricted this data generation procedure to chromosome 20.

Then, we performed imputation of all the resulting VCF files. We first divided chromosome 20 into 35 chunks with size between 1Mb and 2Mb. To prevent edge effects, we also added additional buffer regions of 200kb on each side. Splitting chromosome 20 according to these parameters was done using GLIMPSE_chunk v1.0.1. We then performed the imputation with GLIMPSE_phase using the reference panel 1000 Genomes version 3, a cosmopolitan collection of whole genome sequenced modern samples that we feel well adapted to the various ancestries included in our data set. GLIMPSE_phase was run using the following parameters: 10 burn-in iterations (*--burn 10*), 15 main iterations (*--main 15*) and a depth of 2 for the conditional state selection based on Positional Burrows-Wheeler transform (*--pbwt-depth 2*). Finally, we ligated all imputed chunks back together into chromosome-wide VCF files using GLIMPSE_ligate v1.0.1.

In addition to this, we also performed imputation with Beagle 4.1 with exactly the same input data and reference panel and set its parameters *modelscale* to 2 and *niterations* to 0. These parameter settings allow Beagle v4.1 to run with tractable running times on the data while retaining good accuracy. The chunks of data imputed with Beagle v4.1 were ligated together with bcftools concat 1.10.

To evaluate imputation performance, we employed GLIMPSE_concordance v1.0.1 using as a validation set all genotypes in high-coverage data that were covered by at least eight sequencing reads and at which the most likely genotype was at least 1,000 times more likely than the second best given the genotype likelihoods reported in the VCF/PL fields. Specifically, we computed (i) the squared correlation and (ii) the concordance between imputed and validation genotypes. For (i), we compared minor allele dosages (VCF/DS field) within multiple minor allele frequency (MAF) bins. For (ii), we compared best guess genotypes (VCF/GT field) and stratified the results depending on the type of validation genotype: homozygous reference allele, heterozygous and homozygous alternative allele. As further validation, we increased genomic coverage to 27.5X, 18.9X and 5.4X by deep sequencing a previously published family trio (mother, father, son) from the Late Neolithic mass burial at Koszyce in Poland[4]. This presented an opportunity to validate imputed genotypes and haplotypes on the basis of Mendel's rules of inheritance[5].

To ascertain the potential effects of imputation bias in pairwise analyses, we computed population or individual allele frequencies using FrAnTK [6] (https://github.com/morenomayar/FrAnTK). We removed SNPs with MAF<1% that are imputed less accurately and we kept only SNPs also present in the “1240K” dataset[7]. To minimise the impact of postmortem deamination for the high-coverage ancient genomes, we kept only transversion sites (208,220 SNPs). Most of the ancient genomes were treated as part of a cluster with one individual. The exception was the cluster of 6th Century Hungarian genomes (SZ#)[8] that we pooled together by coverage, (i.e., all un-downsampled high-coverage SZ#s form one cluster, all imputed 1x SZ#s form another cluster and so on). In the end, we obtained 285 clusters. As before, we used GLIMPSE1[9] to impute the genomes and 1000 Genomes phase 3[10] as the imputation reference panel.

#### Results

In Fig. S2.1 and S2.2, we present the imputation accuracy per downsampled genome to 1.0x for chromosome 20. We divided samples into eight classes based on expected genetic proximity and plotted each class separately. In Fig. S2.1, the minor allele frequency (MAF) for each of these groups was estimated from the 1000G reference panel, using European, African, South East Asian, East Asian or American allele frequencies according to the place of origin of samples. As expected, imputation accuracy decreases as minor allele frequency decreases too. For common variants (MAF≥5%), imputation accuracy is remarkably high (>0.9) and closely matches what is usually obtained for modern samples. An exception to this are African samples which exhibit lower accuracy in some cases (especially for baa01). This likely results from the reference panel we used that does not represent well the underlying ancestries of these samples. In Fig. S2.2, we present imputation accuracy per genome as genotype discordance: the fraction of validation genotypes incorrectly imputed stratified by homozygous and heterozygous genotypes. As expected, homozygous reference alleles exhibit lower error rates than heterozygous and homozygous alternative alleles. Error rates are remarkably low: less than 1% overall and less than 5% for the most challenging genotypes to impute (RA and AA). Again, African samples exhibit much higher error rates.

In Fig. S2.3, we present how imputation accuracy varies for all 42 samples depending on multiple factors expected to affect accuracy. First, we look at coverage and find that 0.4x coverage is enough to get 0.9 imputation accuracy at common variants (MAF≥10%). Of note, even 0.1x allows reaching 0.8 at common variants. Second, we considered whether imputing the 42 jointly with the remaining 1,622 low coverage samples could decrease imputation accuracy and did not find evidence of this happening: we get very similar accuracy results. Finally, we check how GLIMPSE imputation does compare to Beagle 4.1 in case of ancient low coverage samples and find that GLIMPSE brings a substantial accuracy boost across the entire frequency range. Altogether, our validation analysis showed that ancient European genomes can be imputed confidently from coverages above 0.4x and highly valuable data can still be obtained with coverages as low as 0.1X when using specific QC on the imputed data.

*Imputation of the full dataset*

Given the outcome of the benchmarking described above, we then proceeded with the imputation of the full dataset. In total, we retained 1,664 samples with at least 0.1x mean coverage. Similarly as before, we extracted all variable positions in 1000 Genomes version 3 that correspond to non-singleton bi-allelic SNPs and call genotype likelihoods at all these variants for all samples using bcftools v1.10, thereby resulting in a VCF containing data for 1,664 samples across 43,285,119 SNPs. Of note, the reference genome used in this analysis was hg19, b37. We then used GLIMPSE_chunk to split all the data into 1,841 chunks of 1Mb to 2Mb with overlapping 200kb buffers on each side. All these chunks of data were imputed using GLIMPSE_phase v1.0.1 with 1000 Genomes version 3 as a reference panel of haplotypes. Imputed chunks were ligated back together using GLIMPSE_ligate v1.0.1, resulting in chromosome-wide VCF files containing the following information: (i) best genotype guesses (VCF/GT field), (ii) expected non-reference allele dosage (VCF/DS field), (iii) genotype posterior probabilities /VCF/GP field) and (iv) haplotype pairs sampled during imputation (VCF/HS field). Finally, we used GLIMPSE_sample v1.0.1 to produce consensus haplotype calls at all variants for all samples from the VCF/HS field.

*Pairwise testing using D-statistics*

We first calculated D-statistics of the form D(CHB, Loshbour; H3, YRI) to test for WHG admixture in H3, that include 36 ancient groups and all the 1000 Genomes populations, using the Han Chinese population in the 1000 Genomes as a related population to H3, and Yoruba as an outgroup. Given that in our previous genetic clustering analyses, the WHG-like component increased in imputed genomes at the expense of the Anatolian farmer-like component, we tested WHG and Anatolian farmer admixture in individuals of European ancestry by computing D(Stuttgart, Loshbour; H3, YRI), where the Stuttgart[3] genome represents this second source. Fig. S2.10 contains the D-stats results we obtained when restricting to high-coverage ancient genomes and the abovementioned present-day populations, which we use here as our ground truth.

We obtained similar D-statistics values for high-coverage and imputed data (Figure S2.11). For the first test, D(CHB, Loshbour; H3, YRI), we find only four genomes falling outside the y=x line at 0.1x (where x and y are the D-stats values for imputed and high-coverage genomes, respectively) in Figure Y, and none of them at higher coverages. In the case of the second test, all 95% confidence intervals included the y=x line for all tests and coverages. Furthermore, when defining significance by a |Z-score| > 3, we did not find any change but for two individual samples (Table S2.2). In the first test, D-statistics calculated with imputed 0.1x, 0.25x and 0.5x BOT2016 [11] (Z-scores: -2.75, -2.84 and -2.97, respectively) were not significant contrary to the corresponding high-coverage data (-3.49), whereas the imputed 0.1x SSG-A-2 genome [12] yielded a significant D-statistics value (-3.06 vs. -2.98).These significance changes are mostly due to small variations around the significance threshold and not due to major differences in the Z-scores.

In conclusion, these analyses show that imputation has very little impact on D-statistics when considering samples with at least 0.1x sequencing coverage. Even though differences between D-statistics calculated with high-coverage and imputed data tend to be smaller for coverages above 0.1x, we arrived at the same results in regard to their significance (|Z|>3) for 46 out of the 50 D-tests.

#### Effects of Low coverage

It should be noted, however, that certain issues may arise when using this imputation on very low coverage data. Specifically, the imputation errors GLIMPSE makes with low coverage data tend to predominantly occur by incorrectly filling in the major allele at a given SNP. This is not an issue specific to GLIMPSE itself but is instead inherent to any kind of Bayesian approach to imputation. In the absence of informative data about the allele at a particular SNP, imputation methods will fall back on the reference panel, or a set of confidently imputed genomes, for imputation, which will tend to fill in missing data with the major allele at each SNP.

To illustrate this phenomenon, we took the 1,492 samples that passed all filters (described in detail in Supplementary Note 3d) and retained only SNPs passing the 1000 Genomes strict mask. For computational considerations, we considered only chromosome 8, which gave 1,139,150 total SNPs. In Fig. S2.4, we plotted, for each of the 2,984 haplotypes, the total number of allele differences between that genome and the reference sequence against the coverage of that sample. Computing Spearman's ρ showed a substantial correlation, which persisted after filtering for SNPs on INFO > 0.5, leaving 584,280 SNPs, and INFO > 0.8, leaving 336,842 SNPs (Fig. S2.5 and S2.6 respectively). Visually, it appeared that this correlation was driven by a reduced number of differences to the reference sequence among very low coverage genomes, and we confirmed this by considering SNPs of all INFO scores and noticing that the correlation decreases sharply when samples below 0.3x are dropped (Fig. S2.8) and can be decreased even further by dropping samples below 2x coverage (Fig. S2.9). We also confirmed that the correlation is very small when only retaining very high INFO score SNPs, specifically INFO > 0.97, which retained 56,925 SNPs (Fig. S2.7). This is to be expected, as the imputation for high-confidence SNPs should show no significant biases.

This phenomenon, where coverage can be predictive of sequence, is important to keep in mind when running certain types of analyses on these data. Analyses such as PCA and admixture modelling, which mainly rely on common SNPs that are shared among many individuals, are not expected to be significantly affected, as imputation is quite accurate for SNPs with high MAF (see Supplementary Note 3d for a thorough analysis of how coverage affects PCA). However, this observation has important implications for genealogy reconstruction and other analyses that rely on overall sequence similarity or otherwise utilise rare SNPs. We recommend that researchers using these imputed data carefully consider what effect the inclusion of low coverage samples might have on their analyses and then utilise appropriate MAF/INFO filters on SNPs and/or coverage filters on samples as necessary.

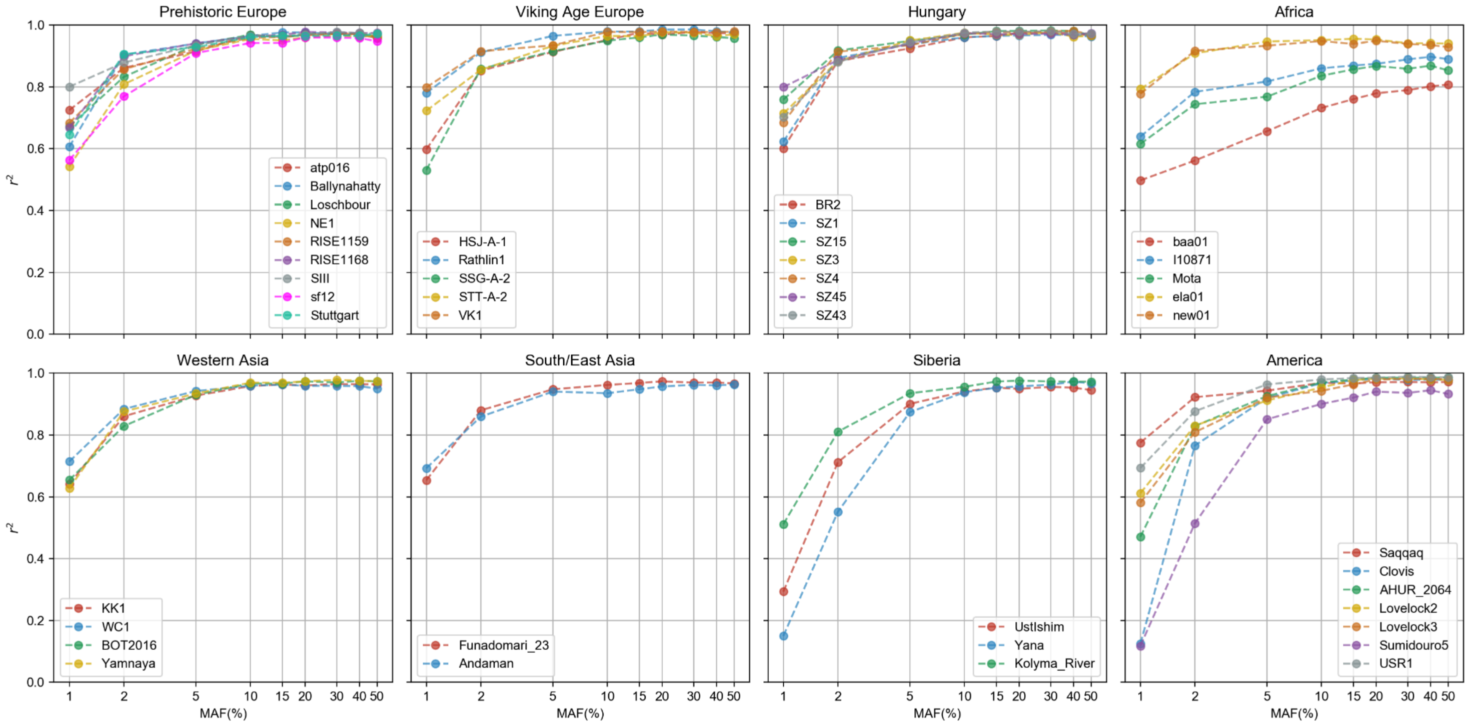


***Figure S2.1: Per sample imputation accuracy (1).*** *Imputation accuracy as squared correlation between imputed and validation genotypes (y-axis). Samples were grouped into eight broad categories based on genetic proximity. Each of the plots corresponds to one of such categories. Minor allele frequencies (MAF; x-axis) were estimated from 1000 Genomes version 3 for matched continental groups (see Table S2.1 for details).*

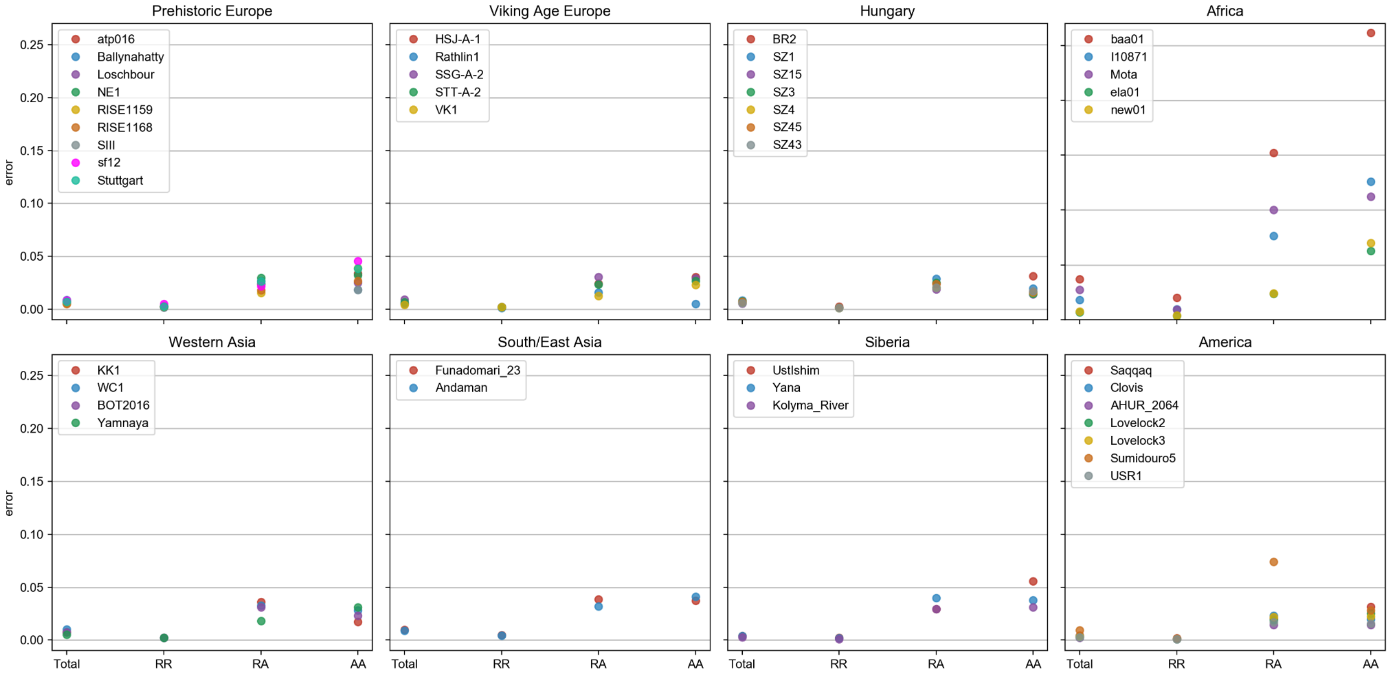


***Figure S2.2: Per-sample imputation accuracy (2).*** *Imputation accuracy as discordance between imputed and validation genotypes (y-axis). Samples were grouped into eight broad categories based on genetic proximity. Each of the plots corresponds to one of such categories. We report results across four types of genotypes: (i) Total; all genotypes together, (ii) RR; validation genotypes with two copies of the reference allele, (iii) RA; heterozygous genotypes (iv) AA; validation genotypes with two copies of the alternative allele.*

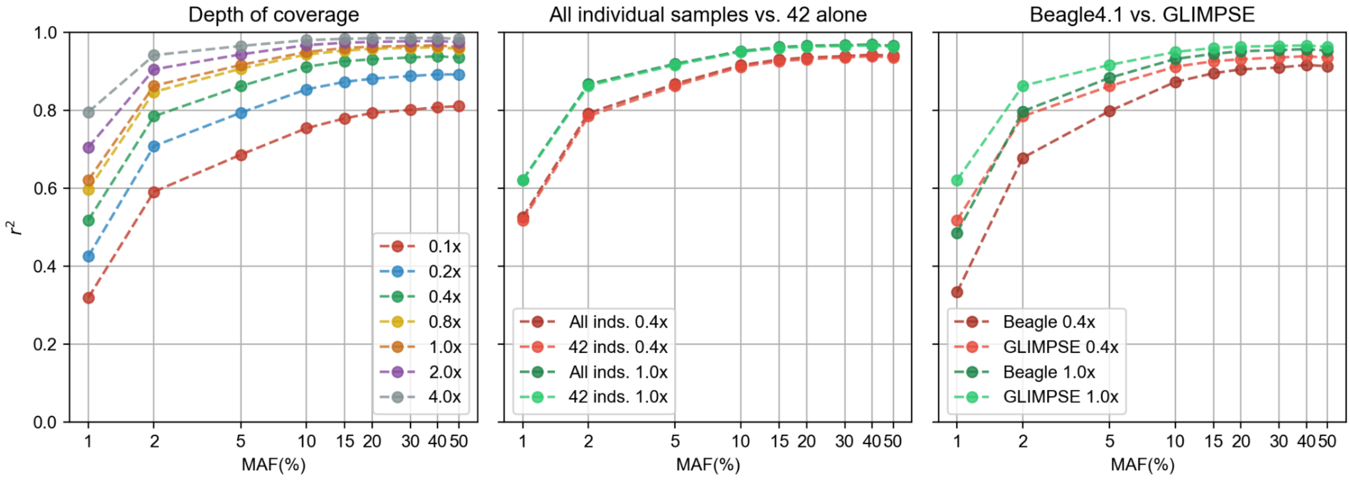


***Figure S2.3: Main parameters affecting imputation accuracy.*** *Imputation accuracy of GLIMPSE across the 42 samples regarding, from left to right: (i) sequencing coverage; (ii) effect of jointly imputing all 1.6K individual samples compared to imputing only the 42 downsampled genomes, (iii) imputation done with Beagle4.1.*

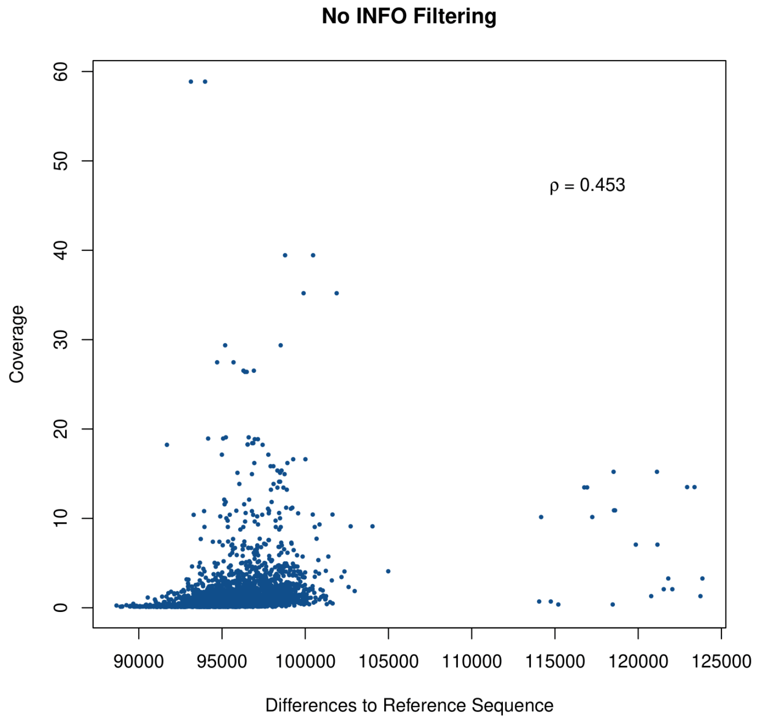


***Figure S2.4.*** *Total number of allele differences to the reference sequence for each of the 2,984 haplotypes against coverage, without INFO filtering.*

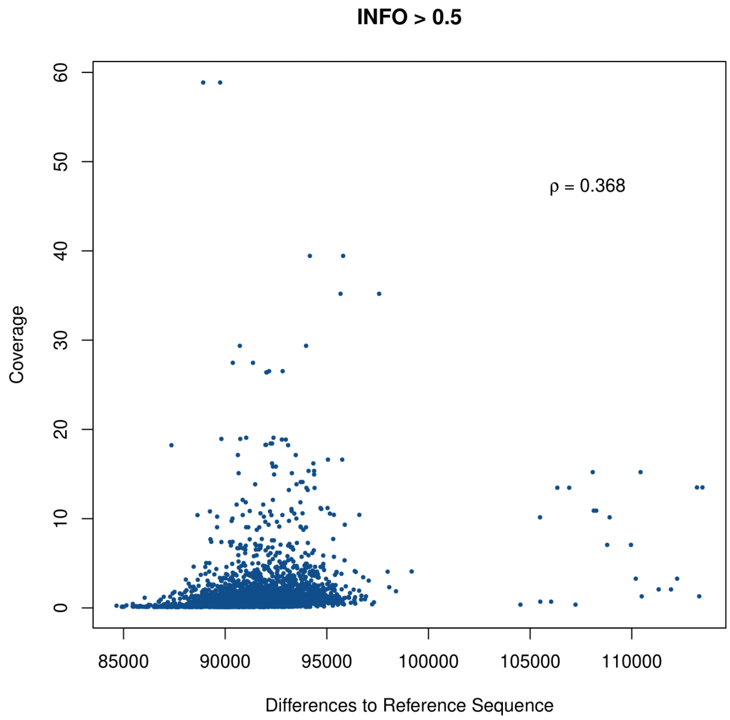


***Figure S2.5.*** *Total number of allele differences to the reference sequence for each of the 2,984 haplotypes against coverage, with filtering for SNPs on INFO >0.5.*

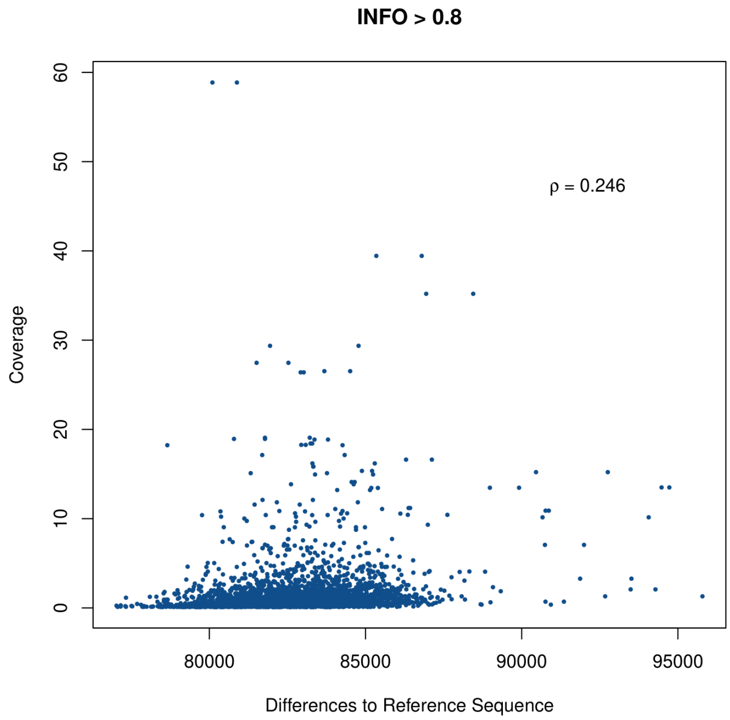


***Figure S2.6.*** *Total number of allele differences to the reference sequence for each of the 2,984 haplotypes against coverage, with filtering for SNPs on INFO >0.8.*

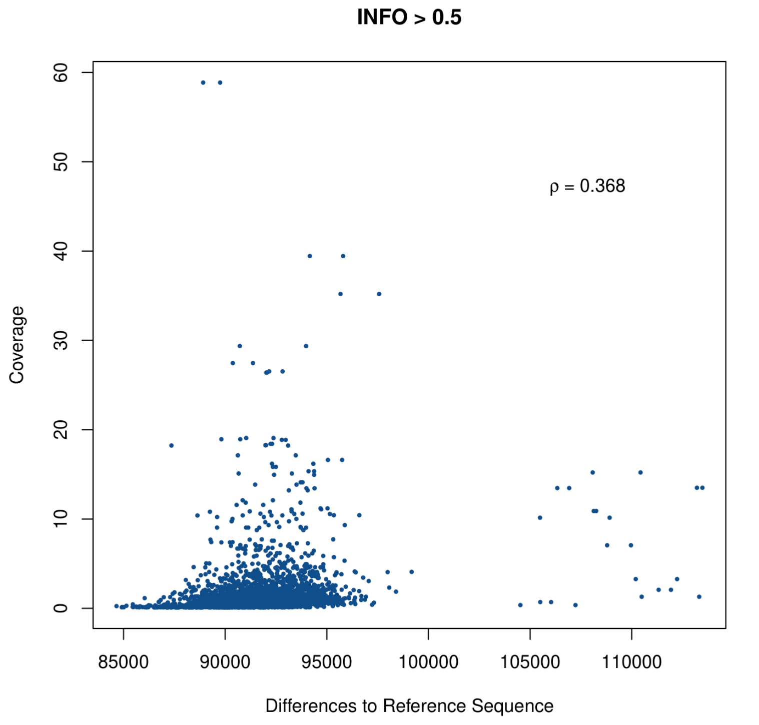


***Figure S2.7.*** *Total number of allele differences to the reference sequence for each of the 2,984 haplotypes against coverage, with filtering for SNPs on INFO >0.97.*

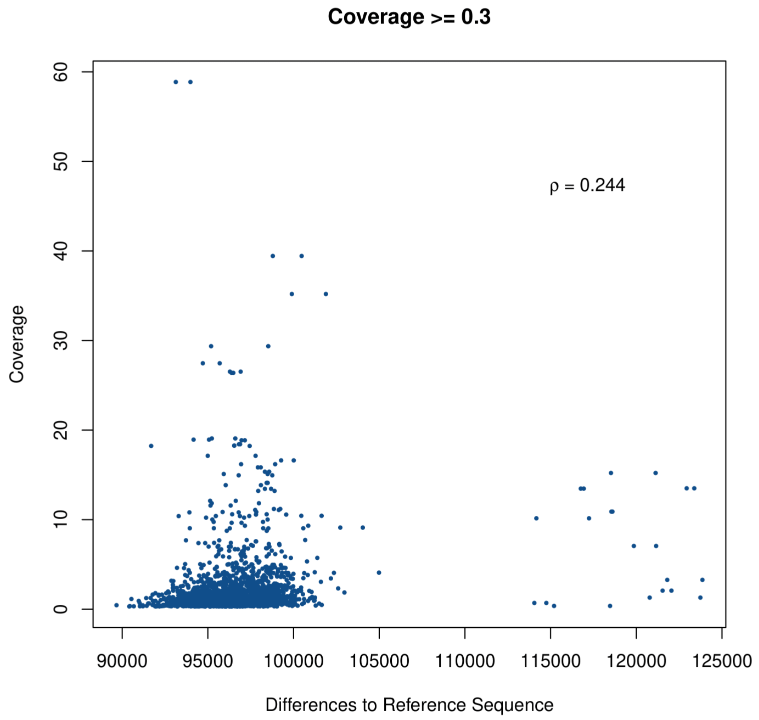


***Figure S2.8.*** *Total number of allele differences to the reference sequence for each of the 2,984 haplotypes against coverage, with filtering for samples on coverage ≥0.3.*

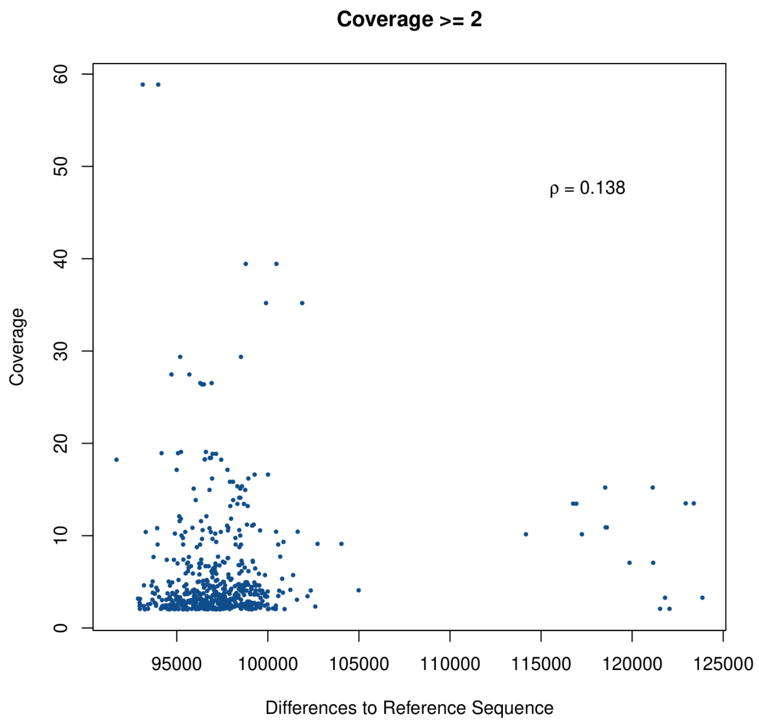


***Figure S2.9.*** *Total number of allele differences to the reference sequence for each of the 2,984 haplotypes against coverage, with filtering for samples on coverage ≥2.*

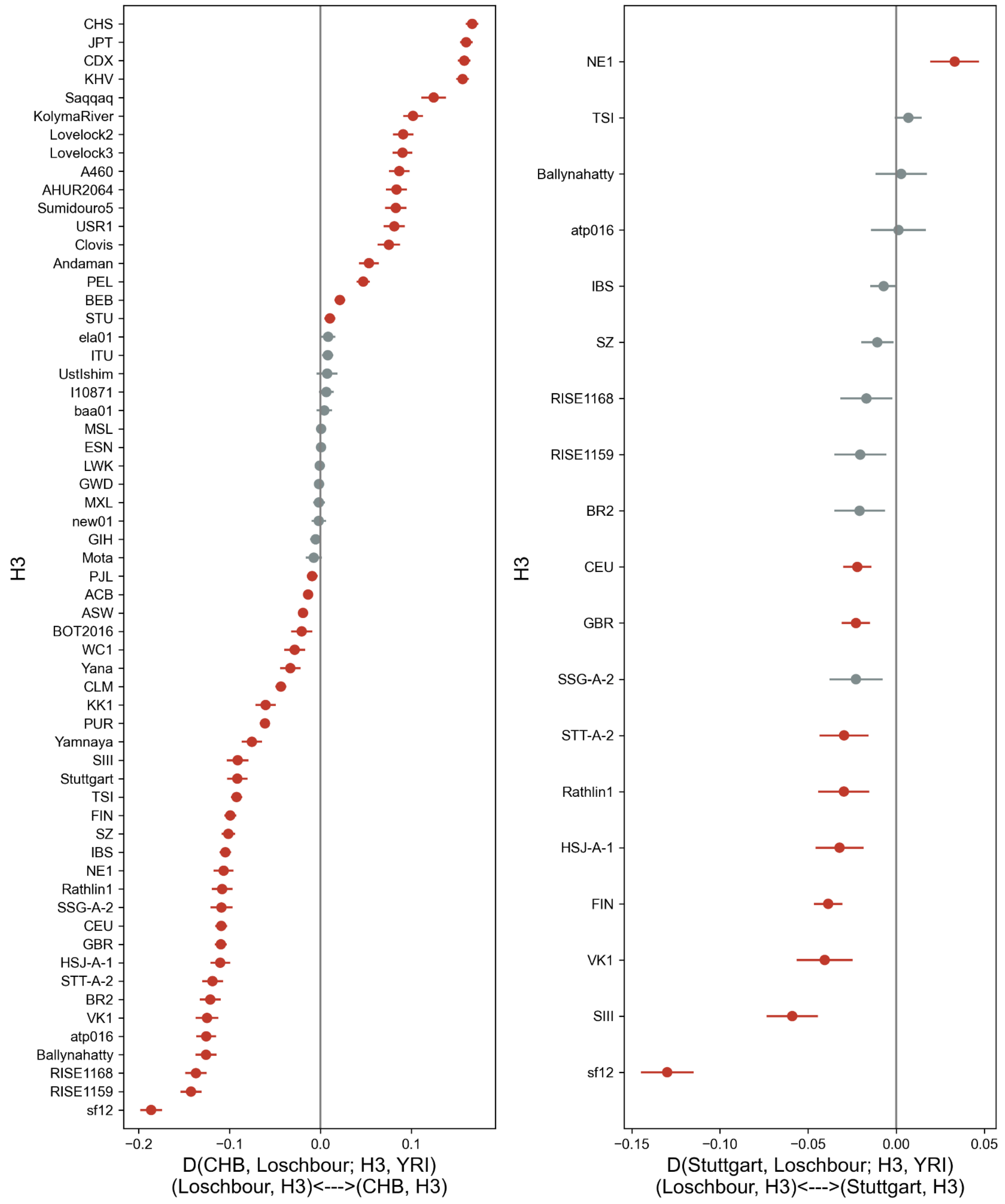


**Figure S2.10: D-statistics testing for WHG admixture.** WHG ancestry is represented by the Loschbour genome and outgroup is the Yoruba (YRI) population in the 1000 Genomes reference panel. Red points indicate significant D-statistics values (|Z|>3). Error bars represent 1.96 times the standard error. Left: D-statistics of the form D(CHB, Loshbour; H3, YRI), where CHB are Han Chinese in the 1000 Genomes panel and H3 is either a high-coverage ancient individual(s) or another 1000 Genomes population. Right: D-statistics of the form D(Stuttgart, Loshbour; H3, YRI), where Stuttgart is an ancient gen

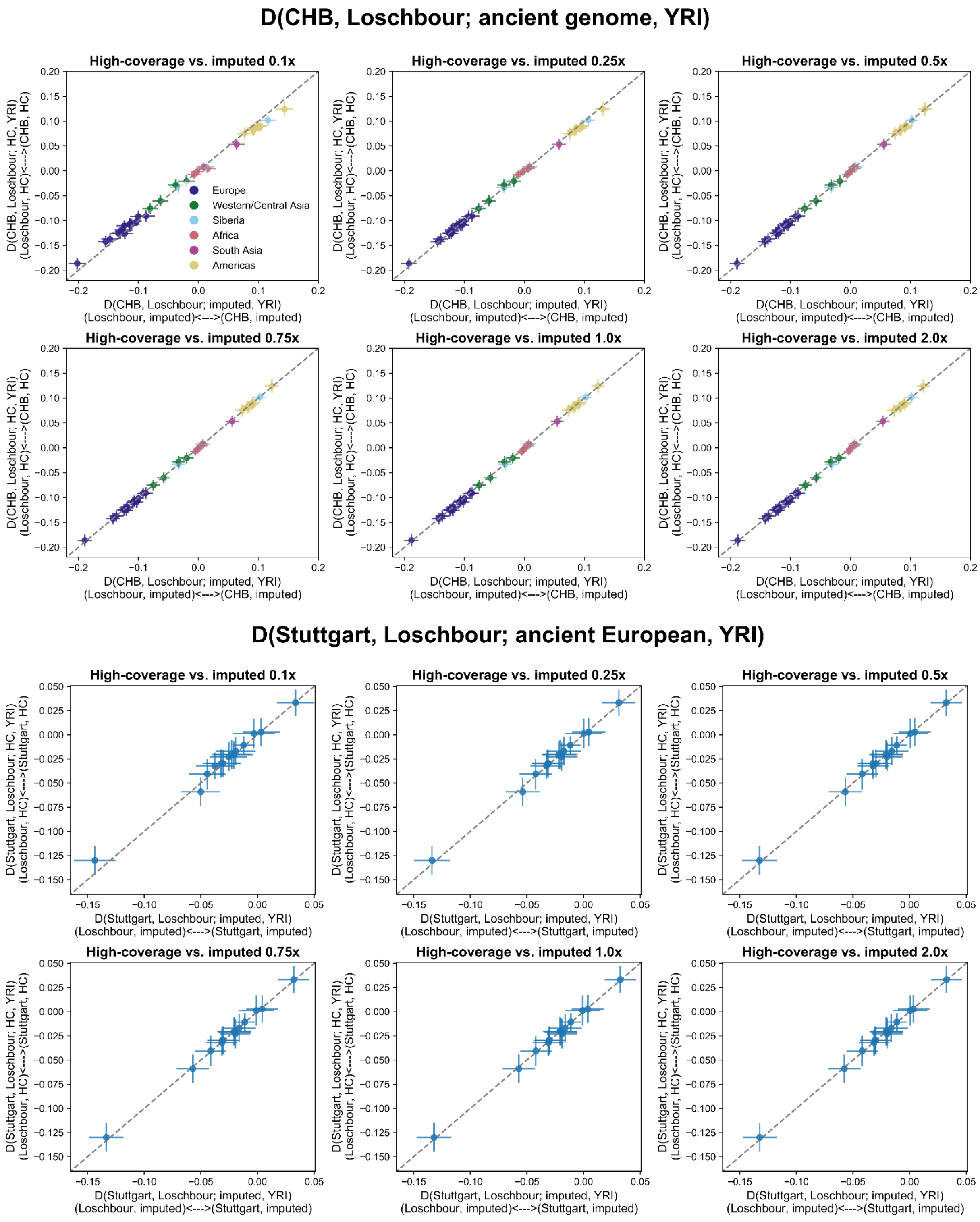


ome representing Anatolian farmer ancestry and H3 is either a high-coverage ancient European individual(s) or populations of European ancestry in the 1000 Genomes panel.

**Figure S2.11: Comparison of D-statistics calculated with imputed (x-axis) and high-coverage ancient genomes (y-axis).** Top: D-statistics tests of the form D(CHB, Loshbour; H3, YRI), where H3 corresponds to ancient genomes coloured by region. Bottom:

D-statistics tests of the form D(Stuttgart, Loshbour; H3, YRI), where H3 corresponds to ancient European genomes. Each subplot corresponds to the imputed genomes' depth of coverage and error bars are 1.96 times the standard error estimated for the D-statistics.

| SampleID | Country | Age (yBP) | MAF 1000G group | Coverage | Reference |
|---|---|---|---|---|---|
| atp16 | Spain | 4867 – 5212 | EUR | 13X | Günther et al. 2015[13] |
| Stuttgart | Germany | 7020 – 7260 | EUR | 16X | Lazaridis et al. 2014[3] |
| Loschbour | Luxembourg | 7940 – 8160 | EUR | 18X | Lazaridis et al. 2014[3] |
| Ballynahatty | Ireland | 4970 – 5293 | EUR | 10X | Cassidy et al. 2016[14] |
| sf12 | Sweden | 8757 – 9033 | EUR | 59X | Günther et al. 2018[15] |
| NE1 | Hungary | 7021 – 7256 | EUR | 18X | Gamba et al. 2014[16] |
| Sunghir III | Russia | 33031 – 35154 | EUR | 11X | Sikora et al. 2017[17] |
| Rathlin1 | Ireland | 3835 – 3976 | EUR | 11X | Cassidy et al. 2016[14] |
| SSG-A2 | Iceland | 950 – 1100 | EUR | 10X | Ebenesersdóttir et al. 2018[12] |
| HSJ-A1 | Iceland | 950 – 1080 | EUR | 29X | Ebenesersdóttir et al. 2018[12] |
| STT-A2 | Iceland | 950 – 1050 | EUR | 14X | Ebenesersdóttir et al. 2018[12] |
| VK1 | Greenland | 750 – 950 | EUR | 12X | Margaryan et al. 2020[18] |
| BR2 | Hungary | 3060 – 3220 | EUR | 18X | Gamba et al. 2014[16] |
| SZ15 | Hungary | 1346 – 1538 | EUR | 11X | Amorim et al. 2018[8] |
| SZ3 | Hungary | 1346 – 1538 | EUR | 11X | Amorim et al. 2018[8] |
| SZ4 | Hungary | 1347 – 1538 | EUR | 10X | Amorim et al. 2018[8] |
| SZ45 | Hungary | 1348 – 1538 | EUR | 10X | Amorim et al. 2018[8] |
| SZ43 | Hungary | 1349 – 1538 | EUR | 12X | Amorim et al. 2018[8] |
| SZ1 | Hungary | 1150 – 1350 | EUR | 11X | Amorim et al. 2018[8] |
| baa01 | South Africa | 1831 – 1986 | AFR | 14X | Schlebusch et al. 2017[19] |
| ela01 | South Africa | 453 – 533 | AFR | 13X | Schlebusch et al. 2017[19] |
| new01 | South Africa | 327 – 508 | AFR | 11X | Schlebusch et al. 2017[19] |

| | | | | | |
|---|---|---|---|---|---|
| I10871 | Cameroon | 7800 – 7970 | AFR | 15X | Lipson et al. 2020[20] |
| Mota | Ethiopia | 4419 – 4525 | AFR | 10X | Gallego Llorente et al. 2015[21] |
| KK1 | Georgia | 9550 – 9890 | EUR | 12X | Broushaki et al. 2016[22] |
| WC1 | Iran | 9032 – 9405 | EUR | 10X | Broushaki et al. 2016[22] |
| BOT2016 | Kazakhstan | 5318 – 5582 | EUR | 14X | de Barros Damgaard et al. 2018[11] |
| Yamnaya Karagash | Kazakhstan | 4837 – 4968 | EUR | 26X | de Barros Damgaard et al. 2018[11] |
| Andaman | India | 30 – 150 | SAS | 17X | Moreno-Mayar et al. 2018[23] |
| Funadomari 23 | Japan | 3550 – 3960 | EAS | 39X | Kanzawa-Kiriyama et al. 2019[24] |
| Ust'-Ishim | Russia | 42560 – 47480 | ALL | 35X | Fu et al. 2014[25] |
| Yana 1 | Russia | 30950 – 32950 | ALL | 27X | Sikora et al. 2019[26] |
| Kolyma 1 | Russia | 9665 – 9906 | ALL | 15X | Sikora et al. 2019[26] |
| USR1 | USA | 11270 – 11600 | ALL | 17X | Moreno-Mayar et al. 2018[27] |
| AHUR_2064 | USA | 10770 – 11170 | AMR | 19X | Moreno-Mayar et al. 2018[23] |
| Lovelock2 | USA | 1818 – 1942 | AMR | 15X | Moreno-Mayar et al. 2018[23] |
| Lovelock3 | USA | 567 – 687 | AMR | 19X | Moreno-Mayar et al. 2018[23] |
| Saqqaq | Greenland | 3600 – 4170 | AMR | 13X | Rasmussen et al. 2010[28] |
| Clovis | USA | 12572 – 12726 | AMR | 15X | Rasmussen et al. 2014[29] |
| Sumidouro5 | Brazil | 10258 – 10552 | AMR | 16X | Moreno-Mayar et al. 2018[23] |
| RISE1159 * | Poland | 4840 – 4709 | EUR | 27X | Schroeder et al. 2019[4] |
| RISE1168 * | Poland | 4840 – 4709 | EUR | 19X | Schroeder et al. 2019[4] |
| RISE1160 * | Poland | 4840 – 4709 | EUR | 5X | Schroeder et al. 2019[4] |

***Table S2.1: Detailed list of high coverage ancient validation genomes.*** *From left to right: (1) original sample ID, (2) country of origin, (3) estimated age, (4) best matching continental group in 1000 Genomes used to stratify imputation accuracy results and (5) original coverage from which down-sampling has been performed. * indicates the Neolithic Koszyce trio that was first published in Schroeder et al.[4] but now sequenced to higher depth in this* *study and used for imputation validation purposes. RISE1160 is not counted among the 42 high coverage genomes.*

**Table S2.2: Genomes for which we observed a difference in significance (|Z|>3).** Test 1 corresponds to D-tests of the form D(CHB, Loshbour; H3, YRI) and test 2 to D(Stuttgart, Loshbour; H3, YRI).

| Test | Sample Id | D-stat | Z-score |
|---|---|---|---|
| 1 | BOT2016 high coverage | -0.021 | -3.49 |
| 1 | BOT2016 0.1x | -0.020 | -2.75 |
| 1 | BOT2016 0.25x | -0.018 | -2.84 |
| 1 | BOT2016 0.5x | -0.018 | -2.97 |
| 1 | BOT2016 0.75x | -0.019 | -3.28 |
| 1 | BOT2016 1.0x | -0.019 | -3.29 |
| 1 | BOT2016 2.0x | -0.019 | -3.31 |
| 2 | SSG-A-2 high coverage | -0.023 | -2.98 |
| 2 | SSG-A-2 0.1x | -0.025 | -3.06 |
| 2 | SSG-A-2 0.5x | -0.020 | -2.85 |
| 2 | SSG-A-2 0.25x | -0.019 | -2.68 |
| 2 | SSG-A-2 0.75x | -0.020 | -2.85 |
| 2 | SSG-A-2 1.0x | -0.019 | -2.77 |
| 2 | SSG-A-2 2.0x | -0.020 | -2.91 |

### 3) Demographic inference

#### 3a) Phylogenetic analysis of mtDNA sequences

Tharsika Vimala[1], Martin Sikora[1]

[1]Lundbeck Foundation GeoGenetics Centre, GLOBE Institute, University of Copenhagen, Copenhagen, Denmark

##### Methods

We carried out phylogenetic analysis of the reconstructed mitochondrial genomes from the human remains presented in this study. Only sequences with less than 2000 missing sites were included in the analysis to avoid biases caused by missing data. Sequences were aligned using mafft[1] and used to build a phylogenetic tree using RAxML-ng[2]. The analysis was carried out by using the ‘all-in-one’ option, performing both an ML search and bootstrapping (--all --bs-trees 100) along with the substitution model GTR+I+G4.

##### Results

From the resulting phylogenetic tree, we obtain an overview of how the remains are distributed across the haplogroups in our dataset. We find haplogroup U to be the most common haplogroup in the analysed set of individuals. In particular, subclade U5 is commonly observed among European hunter-gatherers. Focusing on the subclade U5a, we find the remains distributed into two main sub-haplogroups classified as U5a1 and U5a2. U5a1 is mainly influenced by Eastern European hunter-gatherers from Russia and Ukraine, while we also find the Scandinavian remains, NEO752 and NEO18, represented in this clade. In particular, the 9.8 kya old remains of NEO18 are interesting as the genetic structure analysis of the autosomes of NEO18 shows evidence of Ukrainian hunter-gatherer ancestry (Figure 2, main). Haplogroup U5a2 shows a higher representation of Danish hunter-gatherers, specifically in subclade U5a2a, in which we also identify two Mesolithic Iberian individuals, NEO648 and NEO938. We likewise observe a Mesolithic Latvian individual (NEO307) within the U5a2d subclade, which is primarily dominated by Ukrainian and Russian hunter-gatherers (Figure S3a.1). These observations are congruent with the autosomal structure analysis displaying Ukrainian hunter-gatherer ancestry in these individuals. The clade representing haplogroup U5b is mainly influenced by Danish hunter-gatherers along with a few Western European hunter-gatherers from Britain, France, and Iberia. We do, however, also observe a few Ukrainian hunter-gatherers clustering closely to the Danish hunter-gatherer individuals, which could indicate a continuous level of gene flow from Eastern Europe into Scandinavia. Additionally, we identify a single farmer individual, NEO597, carrying U5b, which is a rare example of genetic continuity of a hunter-gatherer associated haplogroup (Figure S3a.1). This contrasts the genetic transition otherwise observed with the arrival of the early farmers. We find a similar overall pattern within the genetic variation of U4, which is mainly influenced by Eastern European hunter-gatherers. Furthermore, we identify individuals with steppe-ancestry as well as two Danish hunter-gatherer remains clustering within the same clade. In the clade of haplogroup U2 we find a single Mesolithic Iberian carrying haplogroup U2’3’4’7’8’9, while the rest of the remains in the U2-clade belong to the sub-haplogroup U2e. U2e is carried by Eastern European hunter-gatherers, although we also identify a significant number of remains from the forest steppe clustering in U2e as well. The Danish Neolithic remains of NEO792 are interesting as this individual carried the highest proportion of steppe-ancestry among the Danish individuals. Haplogroup K, a descending haplogroup of U8, includes a combination of farmers and hunter-gatherers. Specifically, we find the haplogroup K1e to be carried by Danish hunter-gatherers, while K1a and K1b are mainly influenced by Neolithic individuals from Scandinavia (Figure S3a.2). The highest frequency of early farmers is found within the genetic variation of haplogroup H, a descending haplogroup of HV. Both HV and JT are mainly influenced by Western European farmers (Figure S3a.3).

#### Conclusion

In overall we find most of the Scandinavian hunter-gatherers clustering within the variation of haplogroup U. Given the high number of human remains represented, we were able to obtain a phylogeny of a relatively high resolution of this particular haplogroup. Our results show evidence of a continuous migration of especially Eastern hunter-gatherers into Scandinavia. Most of the early farmers carried haplogroup H or fell within haplogroup JT, while a few farmers carried haplogroup K. We find individuals with steppe ancestry mainly clustering together under macro haplogroup M, although we also identify a few steppe individuals in U2 closely related to a Danish Neolithic individual, NEO792, who also carried a high proportion of steppe-ancestry in the nuclear genome.

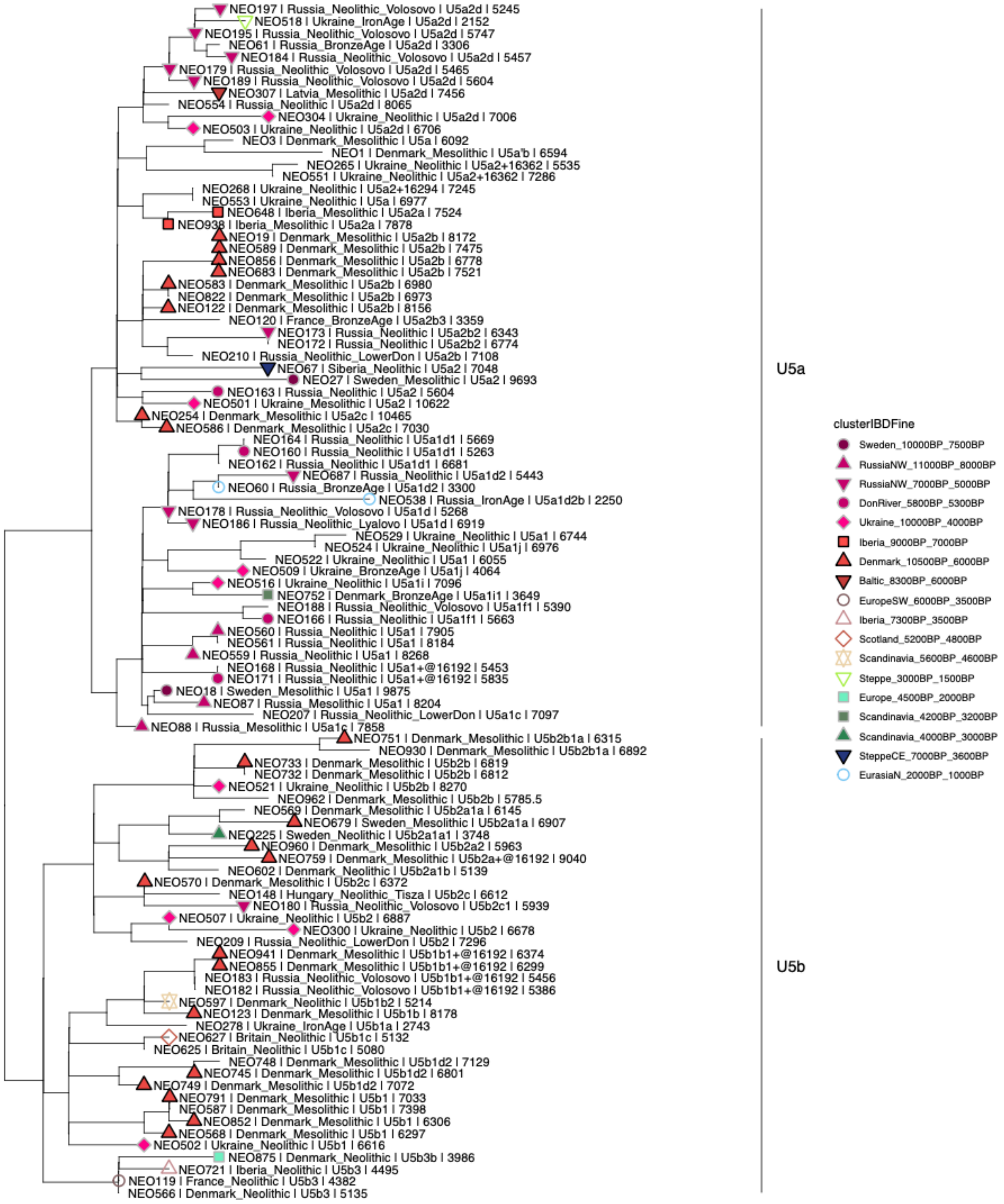


***Figure S3a.1. Maximum likelihood tree of haplogroup U5***

*Maximum likelihood tree displaying the phylogenetic relationship between the human remains carrying haplogroup U5 and more specifically the two subclades U5a and U5b. Labels include information on sample ID, group ID, haplogroup, and age of the respective remains. Symbols indicate the specific fine scale IBD cluster listed in the legend.*

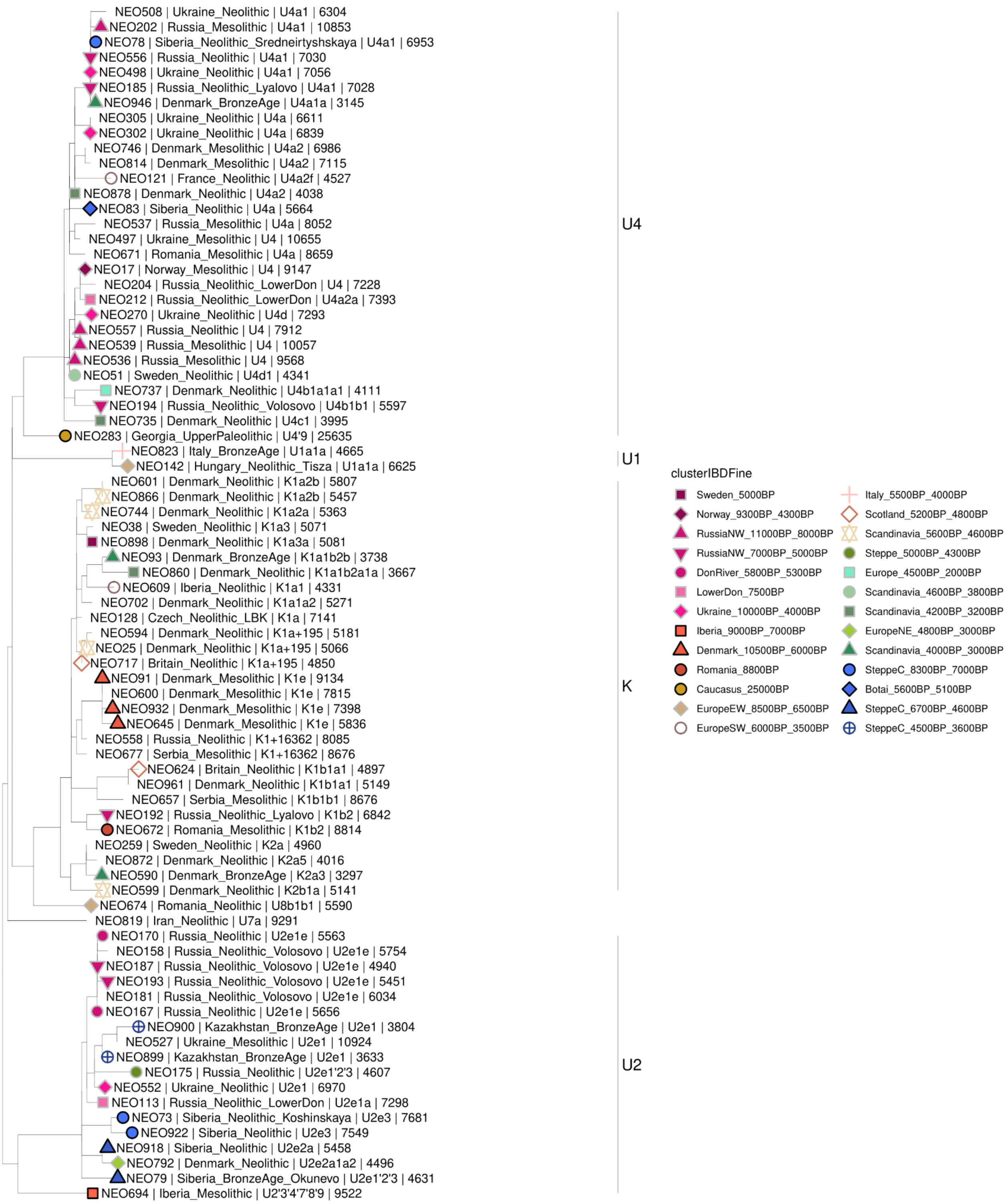


***Figure S3a.2. Maximum likelihood tree of haplogroup U excluding U5***

*Maximum likelihood tree displaying the phylogenetic relationship between the human remains carrying a descending haplogroup of U with the exception of U5. Labels include information on sample ID, group ID, haplogroup, and age of the respective remains. Symbols indicate the specific fine scale IBD cluster listed in the legend.*

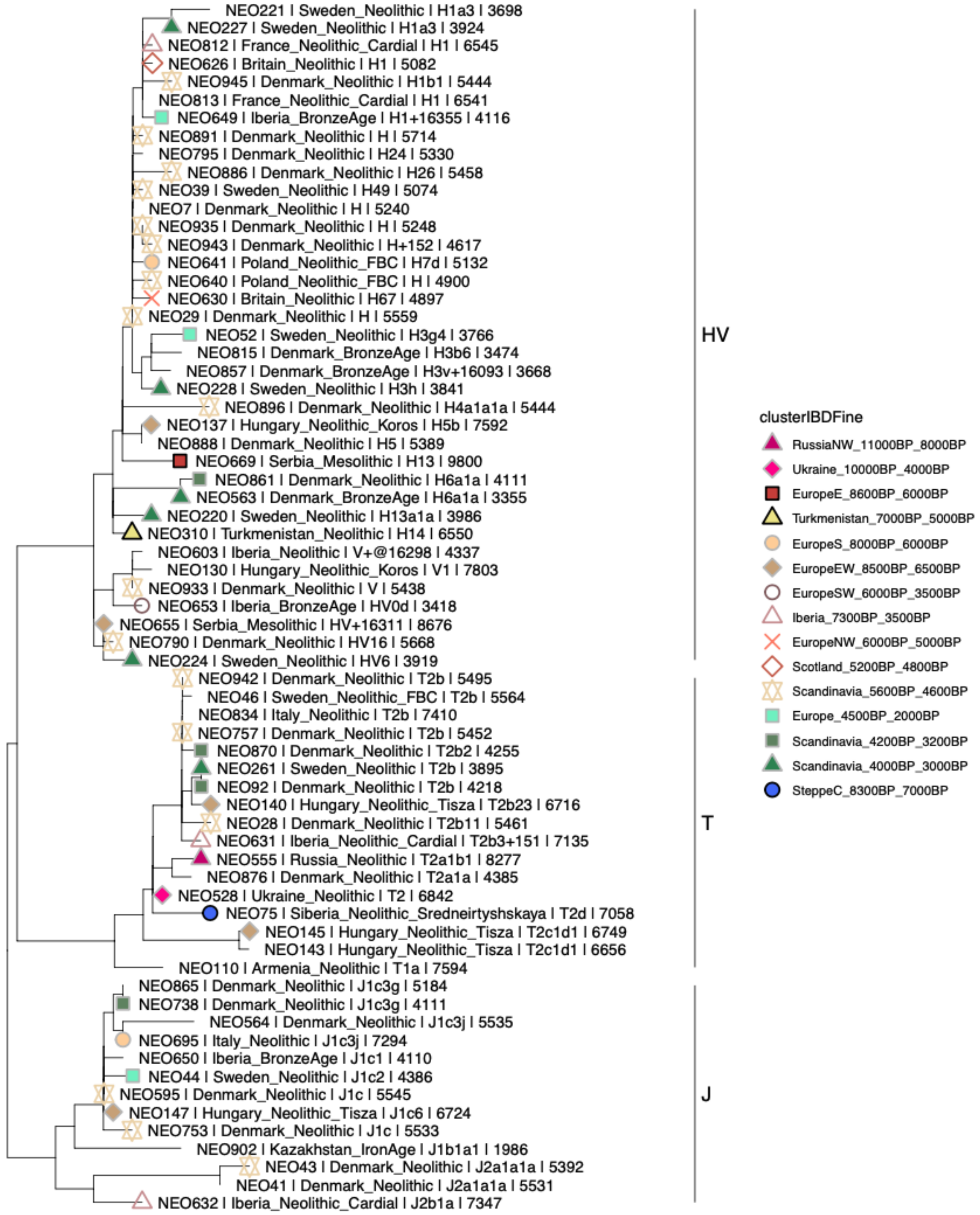


***Figure S3a.3. Maximum likelihood tree of haplogroup farmer-associated haplogroups***

*Maximum likelihood tree displaying the phylogenetic relationship between the human remains carrying an HV or JT descending haplogroup. Labels include information on sample ID, group ID, haplogroup, and age of the respective remains. Symbols indicate the specific fine scale IBD cluster listed in the legend.*

### 3b) Y chromosome / Sex determination

Martin Sikora[1]

[1]Lundbeck Foundation GeoGenetics Centre, GLOBE Institute, University of Copenhagen, Copenhagen, Denmark

#### Methods

Chromosomal sex was inferred through the ratio of reads aligned to each sex chromosome ($R_Y$ statistic)[1]. Reconstructed Y-chromosomal sequences were analysed through phylogenetic placement, based on a reference phylogenetic tree generated from the 1,244 males from the 1000 genomes project[2] (see Methods section of main text).

For in-depth phylogenetic analyses of haplogroups I, R1, and Q, we compiled extended reference panels of high-coverage modern individuals belonging to those haplogroups from publicly available sources[3–6]. To increase resolution for the placement of ancient samples, we also included ancient individuals with Y chromosome coverage ≥1.5X in the reference panels. For each haplogroup panel, we called haploid genotypes individually per sample using bcftools call, setting genotypes with read depth <2 or quality score <30 to missing. Individual VCF files were then merged and filtered to retain only biallelic SNPs polymorphic in the reference panel. For each haplogroup reference panel, we built phylogenetic trees using RAxML-NG and performed phylogenetic placement as described above, restricting to target samples with >0.1X coverage.

#### Results

##### Sex determination

We unambiguously determined genetic sex for all 317 study individuals (118 female, 199 males; Supplementary Data V) based on $R_Y$ statistics (ratio of reads aligned to each sex chromosome)[1]. In a plot of normalised sequencing depth across the X and Y chromosomes, the final dataset individuals form two clearly separated clusters corresponding to XX and XY karyotypes (Fig. S3b.1). The exception is individual YGS-B-2, an Icelandic Viking Age individual previously found to carry an XXY karyotype[7].

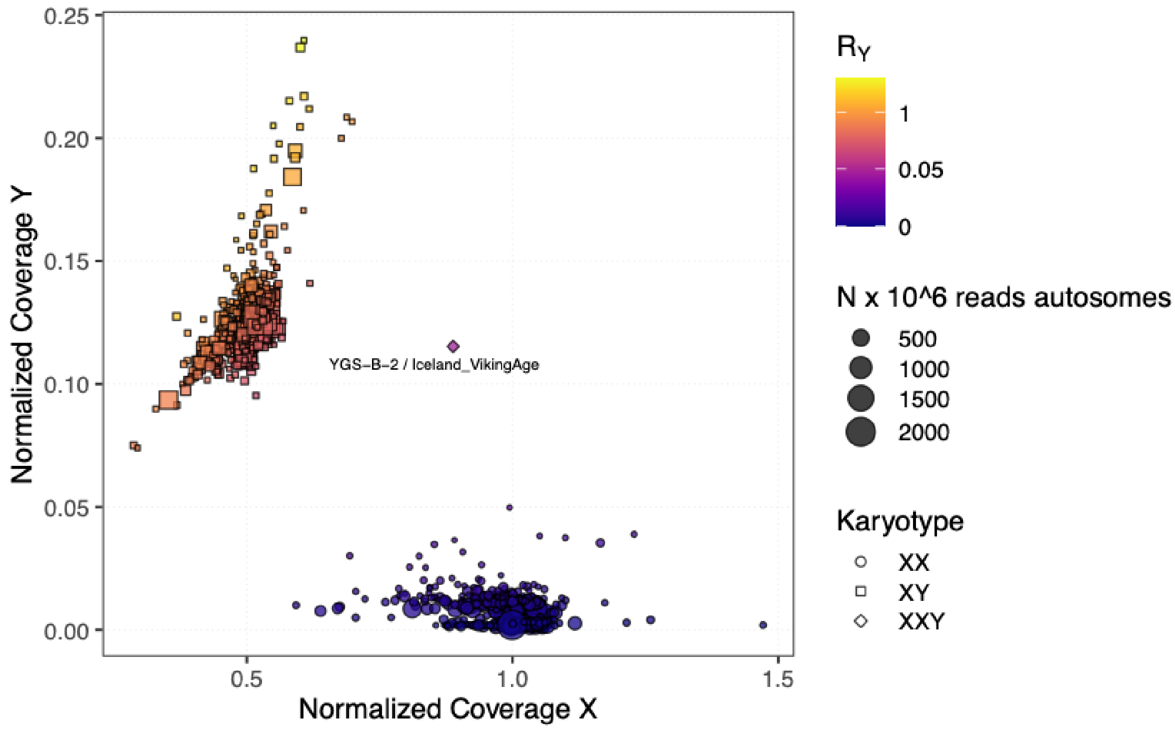


***Fig. S3b.1. Sex determination.*** *Plot shows coverage on X and Y chromosomes normalised by autosomal coverage, for each individual. Symbol colour indicates $R_Y$ ration values, and shape the inferred sex chromosome karyotype. Total number of autosomal reads are indicated by symbol size.*

(hi-res https://www.dropbox.com/s/snhujlmcvl82wgr/sfig_chrY_01_XY.png?dl=0)

#### Phylogenetic placement

We used phylogenetic placement to analyse Y chromosome diversity in our dataset, using a reference . For each ancient sample we obtain a distribution of placement weights across the reference phylogeny, hereby incorporating uncertainty in the placement due to low coverage. The placements can be subjected to analyses such as grafting as a pendant edge to the most likely placement, or clustering of multiple samples. As Y chromosome haplogroups are labels for clades of the phylogeny descending from specific ancestral branches, we can convert the placements into haplogroup calls for a specific sample by assigning haplogroup labels to each branch in the reference phylogeny, and finding the most basal branch that accumulates placement weights up to a specified threshold for the sample. We chose a conservative threshold of 0.99 for the weight accumulation; lower thresholds result in more derived haplogroup calls but with potentially higher uncertainty. Fig. S3b.2 gives an example of this approach for NEO962, a Mesolithic individual from Denmark with low coverage of 0.036X.

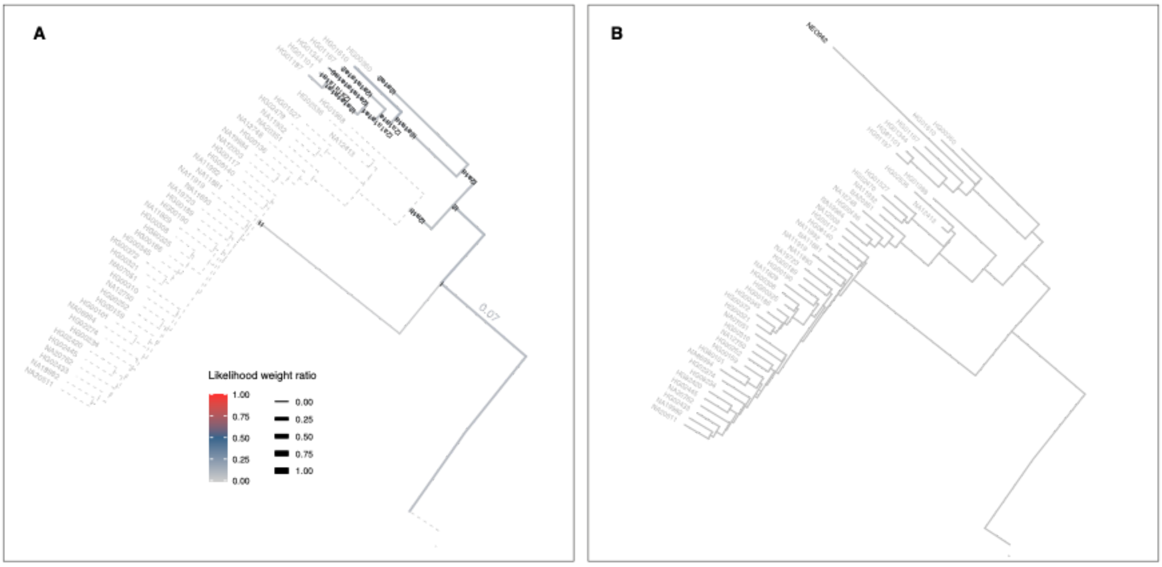


***Fig. S3b.2. Phylogenetic placement.*** *Plot showing phylogenetic placement weights (A) and graft tree with most likely placement (B) for individual NEO962 on a subtree representing reference individuals with haplogroup I. (A) Weights for individual branches are indicated with edge colour and width, edges without placements are indicated with dashed line. While the majority of the placement weight mass is distributed among branches of haplogroup I2, non-zero weights are also found on branches ancestral to I1 (0.02) and I (0.07). The individual is hence conservatively assigned to haplogroup I. (B) The most likely branching of NEO962 was found within the subclade I2a1a1a, albeit this placement is associated with considerable uncertainty.*

(hi-res https://www.dropbox.com/s/np81jfdm6owt71a/sfig_chrY_02_placement.png?dl=0)

#### Phylogenetics of selected haplogroups including newly reported samples

Haplogroup I

Haplogroup I2 was common among newly reported samples from western Eurasian hunter-gatherer contexts, as well as later Neolithic farmers. In particular, the 25 Danish Mesolithic male individuals were exclusively carriers of haplogroup I2, albeit with considerable diversity across different sub-haplogroups (Fig. S3b.3). Neolithic farmer individuals from Scandinavia were predominantly placed within an ancient-only subclade of haplogroup I2a1a2, containing other individuals from Neolithic farmer contexts across Europe.

The earliest presence of haplogroup I1, which is the most common haplogroup among present-day Scandinavians, was found ~4,000BP among late Neolithic and early Bronze Age Scandinavians newly reported in this study (Fig. S3b.3). A single Swedish Mesolithic individual (sf11) was placed at the base of the I1 clade; however, its low coverage (0.1X) precludes to conclude with certainty whether early I1-related lineages were indeed present among Scandinavian hunter-gatherers.

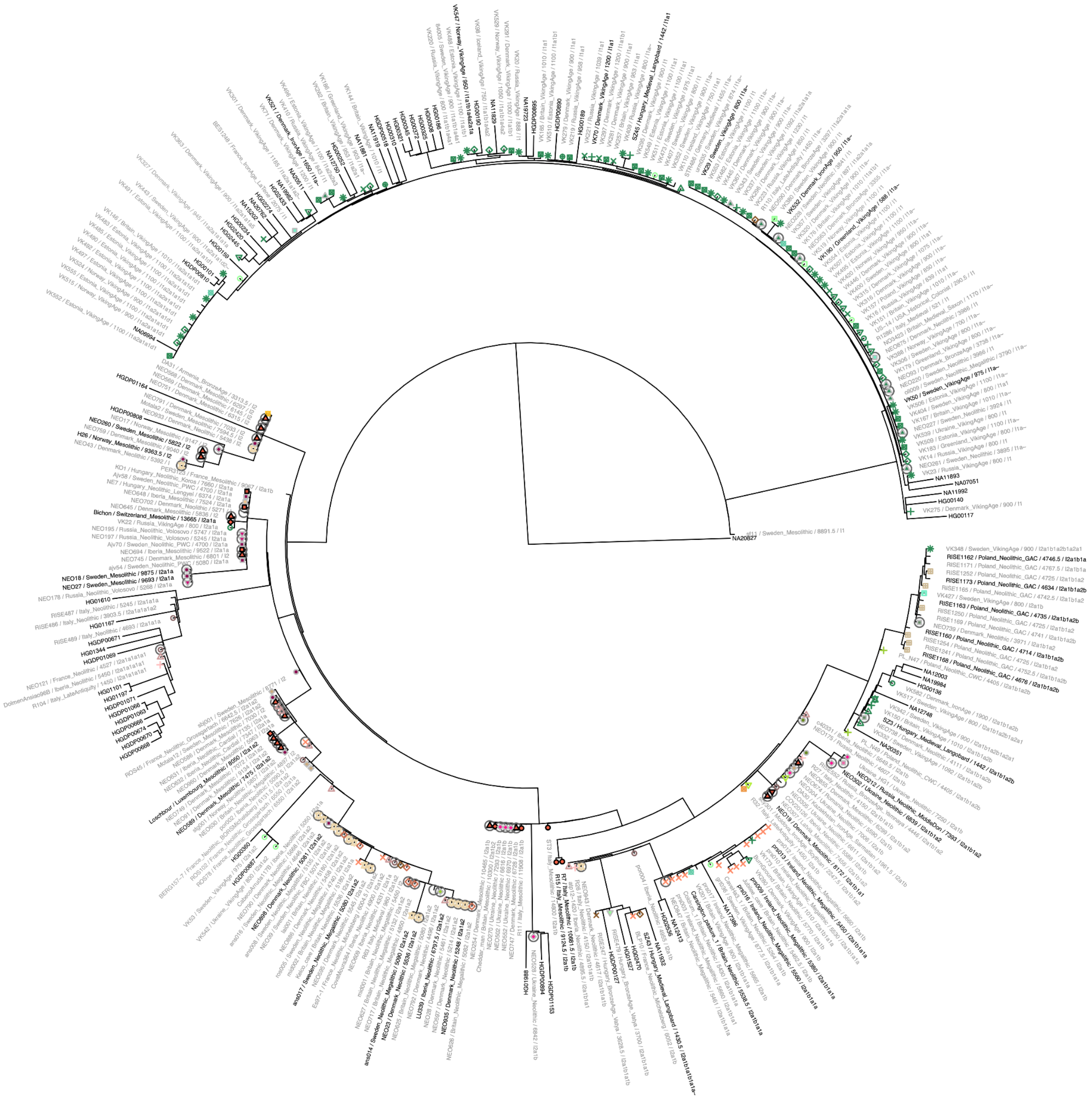

***Fig. S3b.3. Phylogeny of haplogroup I.*** *Phylogenetic tree with most likely placements of ancient samples. Samples labelled with black colour were used to infer the reference tree, whereas samples with grey labels were grafted from phylogenetic placement. Terminal branches for ancient samples were shortened to aid visualisation. Symbol colours and*

*shapes indicate genetic clusters from IBD-based clustering. Newly reported individuals are highlighted with circled symbols.*

(hi-res https://www.dropbox.com/s/j6dslmpqptn4hlw/sfig_chrY_03_hg_I.png?dl=0)

#### Haplogroup J

Haplogroup J was found in four newly reported samples. Among those, a Mesolithic HG individual from Kotias Klde Cave in Georgia, Caucasus (NEO281) belonged to haplogroup J2b. In contrast, two previously reported Caucasus HG males from the same site belonged to haplogroups J1 (SATP) and J2a (KK1) (Fig. S3b.4), demonstrating high diversity of paternal lineages of haplogroup J in the Caucasus region already during the Mesolithic period.

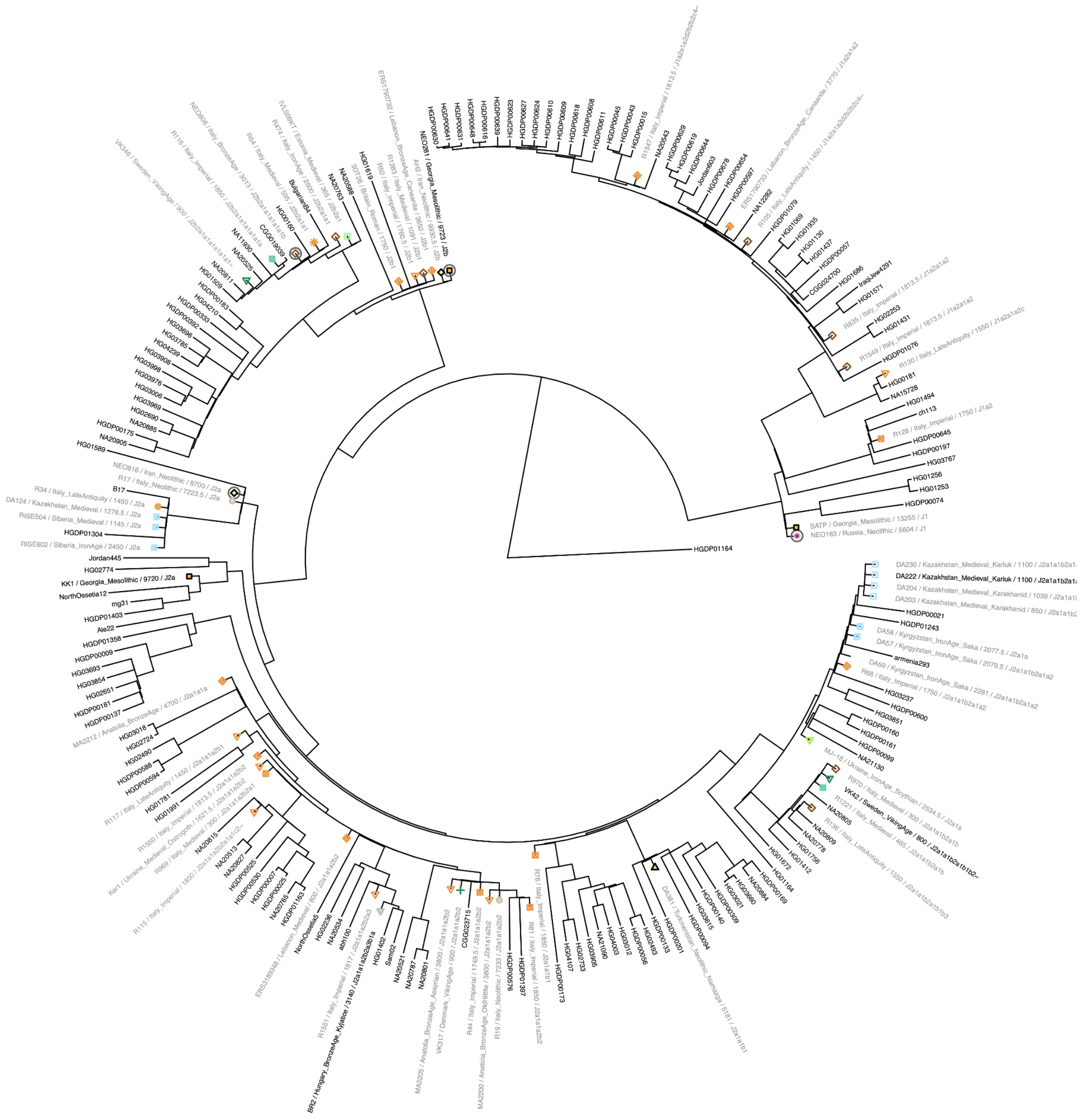

***Fig. S3b.4. Phylogeny of haplogroup J.*** *Phylogenetic tree with most likely placements of ancient samples. Samples labelled with black colour were used to infer the reference tree, whereas samples with grey labels were grafted from phylogenetic placement. Terminal branches for ancient samples were shortened to aid visualisation. Symbol colours and shapes indicate genetic clusters from IBD-based clustering. Newly reported individuals are highlighted with circled symbols.*

(hi-res https://www.dropbox.com/s/xp9b2jow43qctia/sfig_chrY_04_hg_J.png?dl=0)

Haplogroup Q

Haplogroup Q1 was common among newly reported Neolithic hunter-gatherer individuals from the Siberian Forest steppe and the Lake Baikal region (Fig. S3b. 5). We observed haplogroup Q1b2, rare among ancient West Eurasians, in two Ukrainian hunter-gatherers (NEO501, NEO516) as well as two Danish Neolithic farmer individuals (NEO599, NEO744).

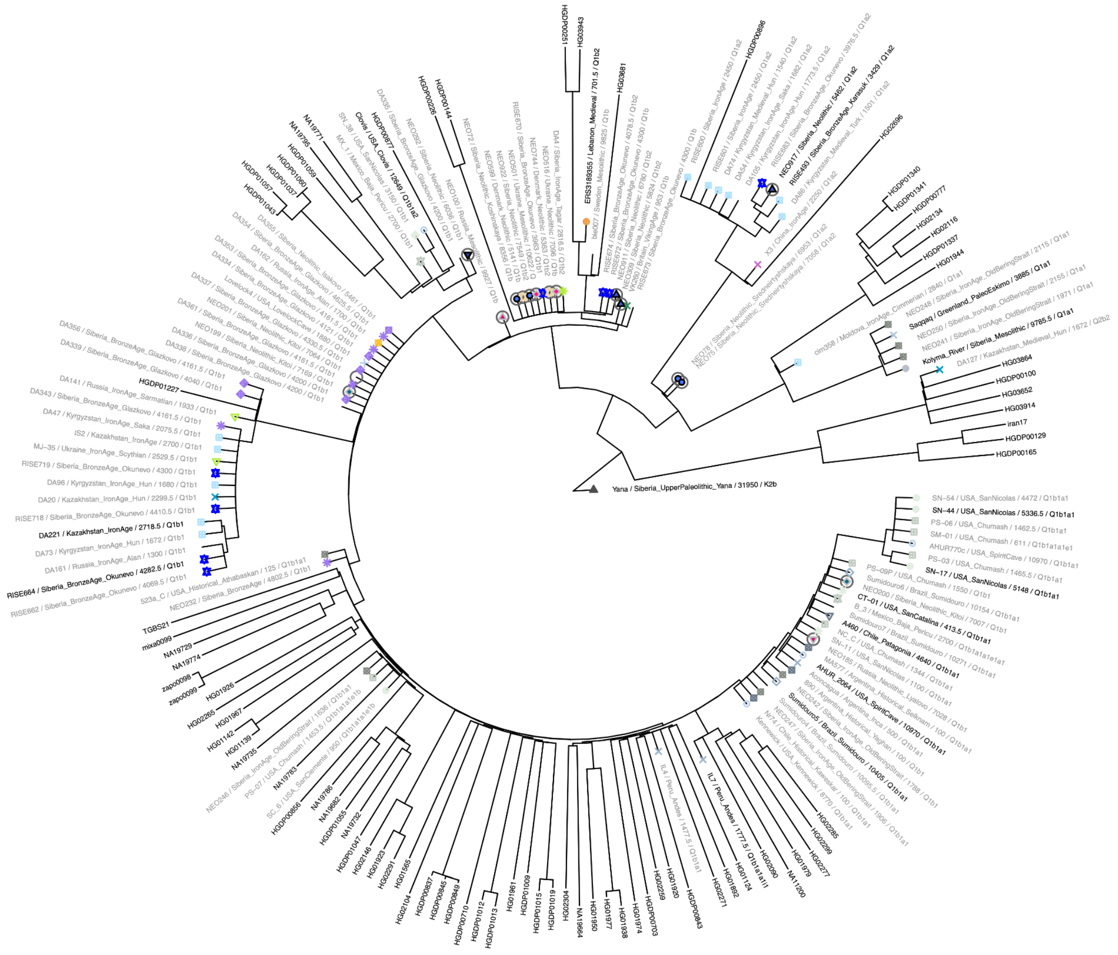

***Fig. S3b.5.*** *Phylogeny of haplogroup Q. Phylogenetic tree with most likely placements of ancient samples. Samples labelled with black colour were used to infer the reference tree, whereas samples with grey labels were grafted from phylogenetic placement. Terminal branches for ancient samples were shortened to aid visualisation. Symbol colours and shapes indicate genetic clusters from IBD-based clustering. Newly reported individuals are highlighted with circled symbols.*

(hi-res [https://www.dropbox.com/s/68ef06mvcfz370s/sfig_chrY_05_hg_Q.png?dl=0](https://www.dropbox.com/s/68ef06mvcfz370s/sfig_chrY_05_hg_Q.png?dl=0))

<u>Haplogroup R1a</u>

Haplogroup R1a was found in the newly reported samples mainly among Eastern European hunter-gatherer individuals. Phylogenetic placement suggests that the oldest individuals from Mesolithic and Neolithic Russia represent early diverging lineages (Fig. S3b.6). Notably, a

~7,300-year-old Neolithic individual from the Middle Don region (NEO113) was placed in a basal R1a clade together with early individuals associated with the Corded Ware complex (poz81, RISE446), which would make it the earliest observation of this lineage reported to date.

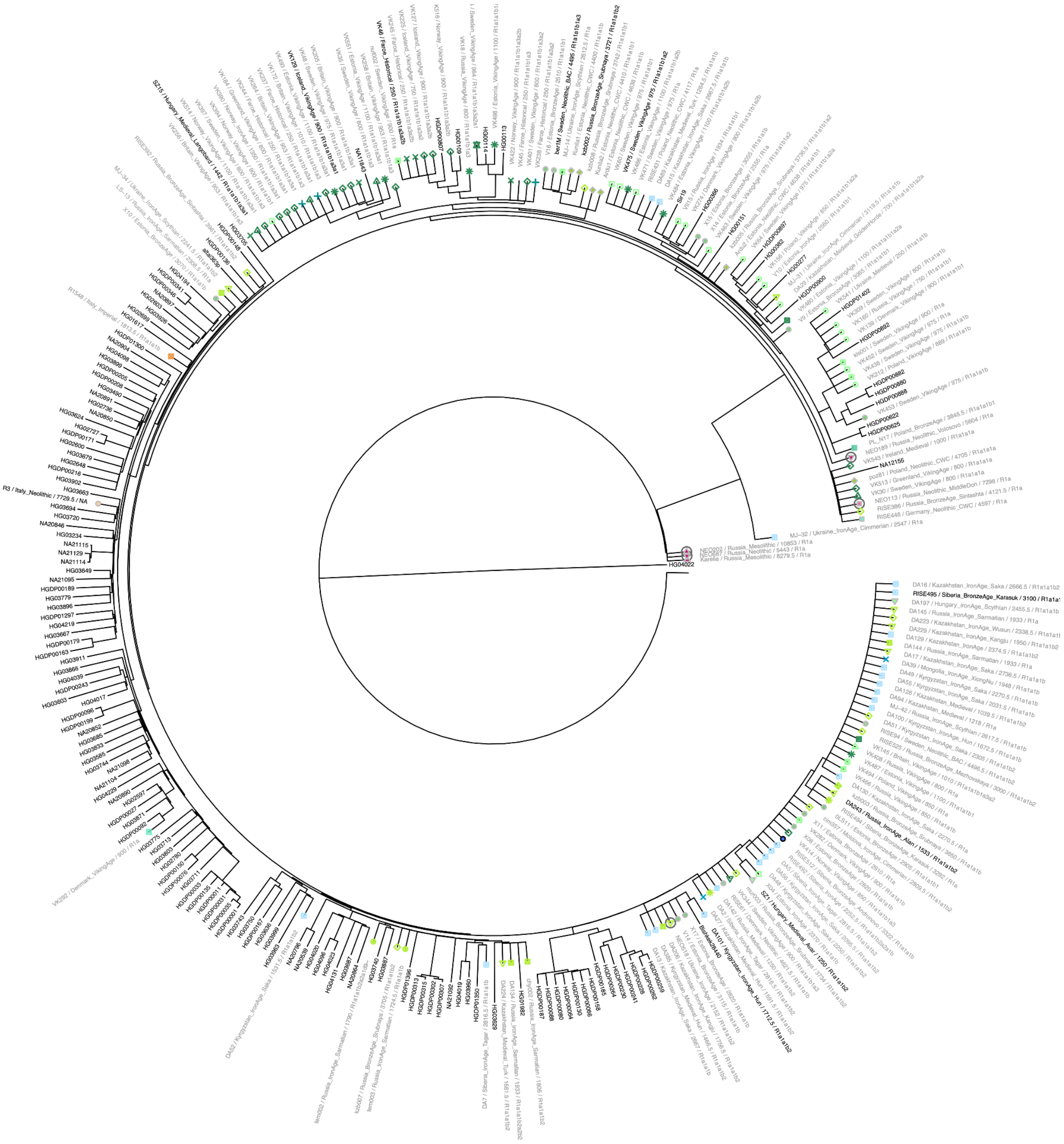

***Fig. S3b.6. Phylogeny of haplogroup R1a.*** *Phylogenetic tree with most likely placements of ancient samples. Samples labelled with black colour were used to infer the reference tree, whereas samples with grey labels were grafted from phylogenetic placement. Terminal branches for ancient samples were shortened to aid visualisation. Symbol colours and shapes indicate genetic clusters from IBD-based clustering. Newly reported individuals are highlighted with circled symbols.*

(hi-res [https://www.dropbox.com/s/oguqdevp92qakyz/sfig_chrY_06_hg_R1a.png?dl=0](https://www.dropbox.com/s/oguqdevp92qakyz/sfig_chrY_06_hg_R1a.png?dl=0))

Haplogroup R1b

Newly reported samples belonging to haplogroup R1b were distributed between two distinct groups depending on whether they formed part of the major European subclade R1b1a1b (R1b-M269). Individuals placed outside this subclade were predominantly from Eastern European Mesolithic and Neolithic contexts, and formed part of rare early diverging R1b lineages (Fig. S3b.7). Two Ukrainian individuals belonged to a subclade of R1b1b (R1b-V88) found among present-day Central and North Africans, lending further support[5,10] to an ancient Eastern European origin for this clade. Haplogroup R1b1a1a (R1b-M73) was frequent among Russian Neolithic individuals.

Individuals placed within the R1b-M269 clade on the other hand were from Scandinavian Late Neolithic and early Bronze Age contexts (Fig. S3b.7). Interestingly, more fine-scale sub-haplogroup placements of those individuals revealed that Y chromosome lineages distinguished samples from distinct genetic clusters inferred from autosomal IBD sharing. In particular, individuals associated with the Scandinavian cluster *Scandinavia_4200BP_3200BP* were all placed within the sub-haplogroup R1b1a1b1a1a1 (R1b-U106), whereas the two Scandinavian males associated with the Western European cluster *Europe_4500BP_2000BP* were placed within R1b1a1b1a1a2 (R1b-P312) (Fig. S3b.8).

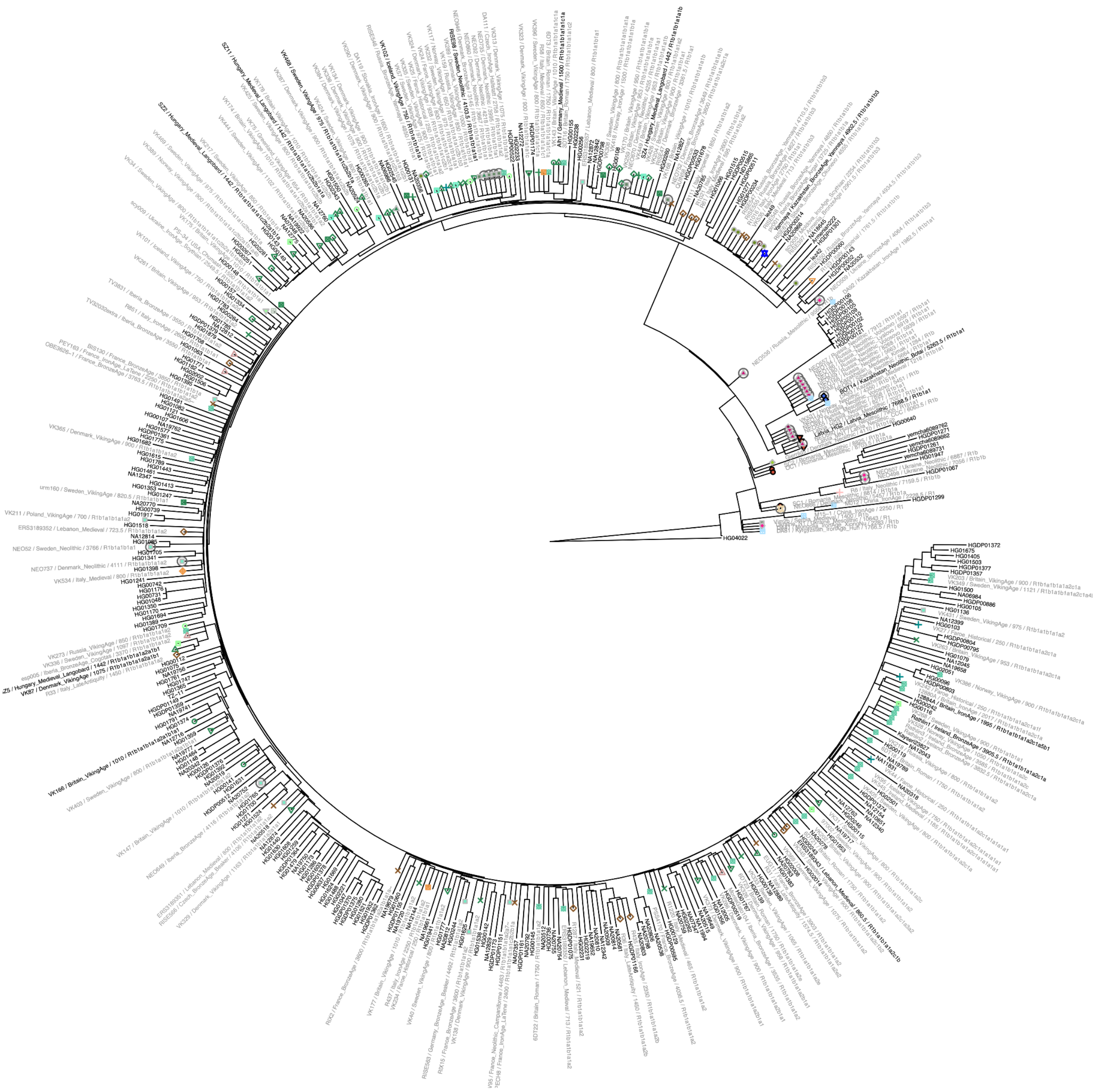

***Fig. S3b.7. Phylogeny of haplogroup R1b.*** *Phylogenetic tree with most likely placements of ancient samples. Samples labelled with black colour were used to infer the reference tree, whereas samples with grey labels were grafted from phylogenetic placement. Terminal branches for ancient samples were shortened to aid visualisation. Symbol colours and shapes indicate genetic clusters from IBD-based clustering. Newly reported individuals are highlighted with circled symbols.*

(hi-res https://www.dropbox.com/s/9dmi1a49d0chamm/sfig_chrY_07_hg_R1b.png?dl=0)

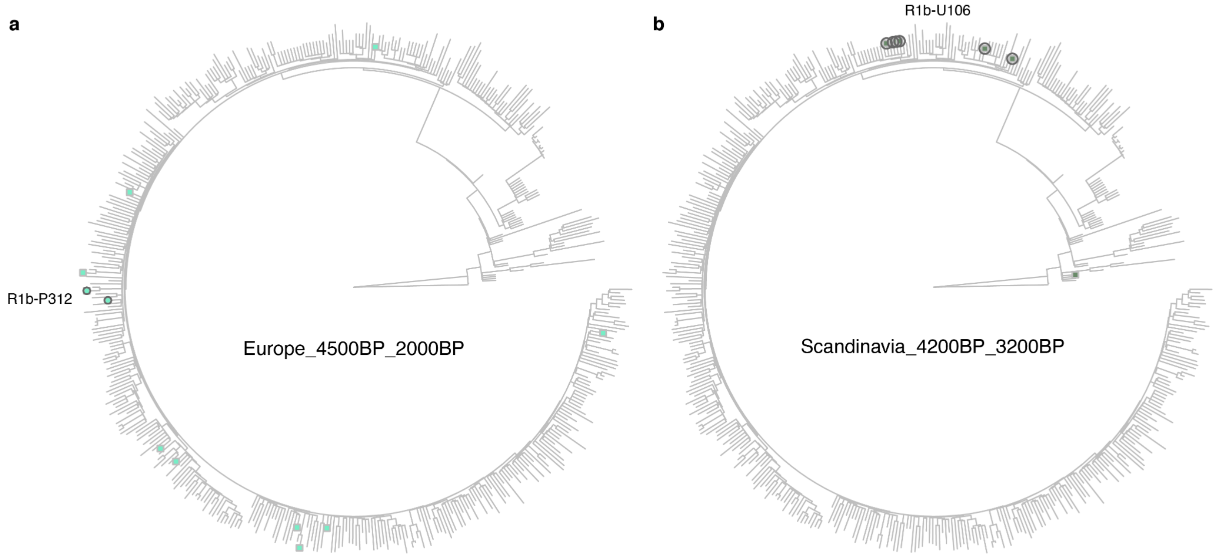


***Fig. S3b.8. Sub-haplogroups of R1b for genetic clusters.*** *Phylogenetic trees showing most likely placements of ancient samples from genetic clusters Scandinavia_4200BP_3200BP (a) and Europe_4500BP_2000BP (b). Terminal branches for ancient samples were shortened to aid visualisation. Symbol colours and shapes indicate samples belonging to the respective genetic clusters. Individuals from Scandinavia within each cluster are highlighted with circled symbols, and annotated with the respective sub-haplogroup labels.*

(hi-res https://www.dropbox.com/s/yrhg95lr2fhq751/sfig_chrY_08_R1_contrast.png?dl=0)

### 3c) Relatedness

Martin Sikora[1]

[1]Lundbeck Foundation GeoGenetics Centre, GLOBE Institute, University of Copenhagen, Copenhagen, Denmark

#### Results

We detected a total of 92 close relative pairs among the 1,664 dataset individuals, including 24 parent-offspring pairs, 36 siblings and 30 2nd degree pairs (Fig S3c.1, Supplementary Data VI), based on allele frequency-free inference of relatedness indices(Manichaikul *et al.*, 2010; Waples, Albrechtsen and Moltke, 2019). We further found evidence of two duplicate / monozygotic twin relationships. Sample NEO70 presented in this study was inferred to be from the same individual as RISE554 previously reported in Allentoft *et al*[6]. Additionally, two male individuals MJ-15 and MJ-32 reported in Järve *et al*[7] were also inferred as duplicate/twin pairs. In both cases genetic sex, mitochondrial as well as Y chromosome haplogroups were all consistent with their inferred relatedness status.

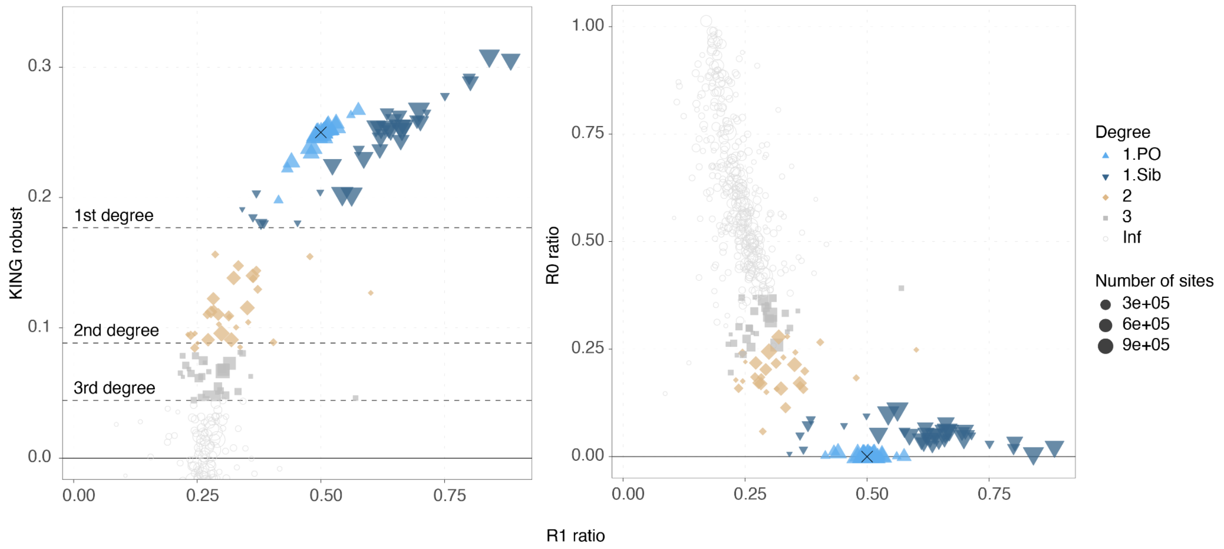


***Fig. S3c.1. Relatedness inference.*** *Plots showing relatedness estimators R1/KING-robust (left) and R1/R2 (right) for pairs of individuals. Inferred relatedness status for each pair is indicated by plot symbol and colour, with symbol size scaled by the total number of informative sites. Black crosses indicate expected values for parent-offspring relationships.* (hi-res https://www.dropbox.com/s/f4hhqh6iome54ze/sfig_rel_01_rel.png?dl=0)

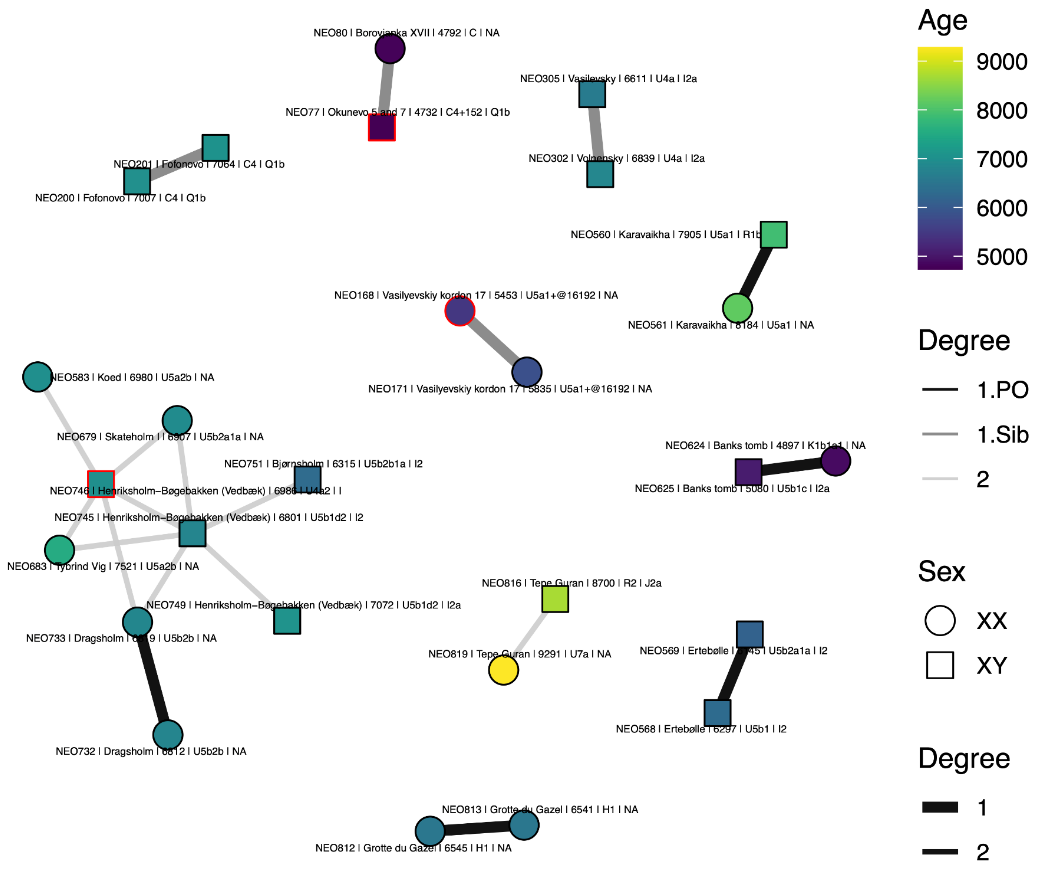


***Fig. S3c.2. Relatedness among newly reported individuals.*** *Network showing first and second-degree relationships, indicated by edge width and colour. Age of individuals is indicated by fill colour, and individuals are labelled with site name, age, mitochondrial and Y chromosome haplogroups. Individuals flagged for possible contamination are indicated in red.*

(hi-res https://www.dropbox.com/s/pj0fe8saa8c29hd/sfig_rel_02_network.png?dl=0)

We identified a total of 10 first- and 12 second-degree relative pairs among the newly reported individuals. However, inspection of the relatedness network revealed that the majority of 2$^{nd}$ degree connections are between Danish Mesolithic individuals from different sites and two individuals from Henriksholm (NEO745, NEO746), one of which was flagged as contaminated (Fig. S3c.2). As excess heterozygosity due to contamination can lead to artificially increased relatedness estimates, we excluded any pair involving those two individuals, as well as two other pairs involving contaminated individuals from the final list of close relatives (Table S3c.1). Finally, three individuals reported here were inferred to be either the same individual (2) or close relatives (1) of samples previously published using targeted SNP capture (Table S3c.2).

| Individual 1 | Individual 2 | Site | Country | Degree | Notes |
|---|---|---|---|---|---|
| NEO568 | NEO569 | Ertebølle | Denmark | 1.PO | NEO568 ("Ertebølle man") is the father of infant NEO569 |
| NEO732 | NEO733 | Dragsholm | Denmark | 1.PO | Mother-daughter relationship, direction unknown |
| NEO624 | NEO625 | Banks tomb | UK | 1.PO | NEO625 is the father of juvenile NEO624 |
| NEO813 | NEO812 | Grotte du Gazel | France | 1.PO | NEO813 is the mother of infant NEO812 |
| NEO560 | NEO561 | Karavaikha | Russia | 1.PO | NEO561 likely the mother of NEO560 (age and MT haplogroup) |
| NEO201 | NEO200 | Fofonovo | Russia | 1.Sib | |
| NEO302 | NEO305 | Volnensky / Vasilevsky | Ukraine | 1.Sib | |
| NEO816 | NEO819 | Tepe Guran | Iran | 2 | |

***Table S3c.1. Close relatives among newly reported individuals.***

| Study Individual | Related individual | Publication | Relationship |
|---|---|---|---|
| NEO306 | SOP002 | Saag et al 2021 *Science Advances* | same individual |
| NEO202 | PES001 | Saag et al 2021 *Science Advances* | same individual |
| NEO73 | I1960 | Narasimhan et al 2019 *Science* | same individual |
| NEO669 | I5407 | Mathieson et al 2018 *Nature* | same individual |
| NEO60 | BOO005 | Lamnidis et al 2018 *Nature Communications* | first degree, infant NEO60 likely daughter of BOO005 |

***Table S3c.2. Study individuals with related published individuals.***

### 3d) Overall Population Structure

Martin Sikora[1]

[1]Lundbeck Foundation GeoGenetics Centre, GLOBE Institute, University of Copenhagen, Copenhagen, Denmark

#### Results

##### Comparison of imputed and pseudo-haploid genotypes in PCA space

Methods for the determination of population structure including principal components analysis[1,2] and model-based clustering (ADMIXTURE[3]) are detailed in the methods section of the main text. To determine the consistency of imputed and pseudo-haploid genotypes when used in PCA, we followed the approach of Antonio *et al.*[4] comparing the coordinates of both sets of genotypes for each individual when projected onto principal components inferred from modern individuals (Fig. S3d.1, S3d.3). We did not use the "*autoshrink*" option of *smartpca* for this analysis to avoid possible systematic differences in the projection correction between the two sets of genotypes. Projected PCA positions for samples passing all filters were consistent between imputed and pseudo-haploid genotypes, with no evidence for systematic shifts (Fig. S3d.2, S3d.4) and only a very subtle relationship of PCA distance between genotypes with genomic coverage (Fig. S3d.3, S3d.6). More substantial shifts were only observed with low coverage (<0.1X) flagged samples.

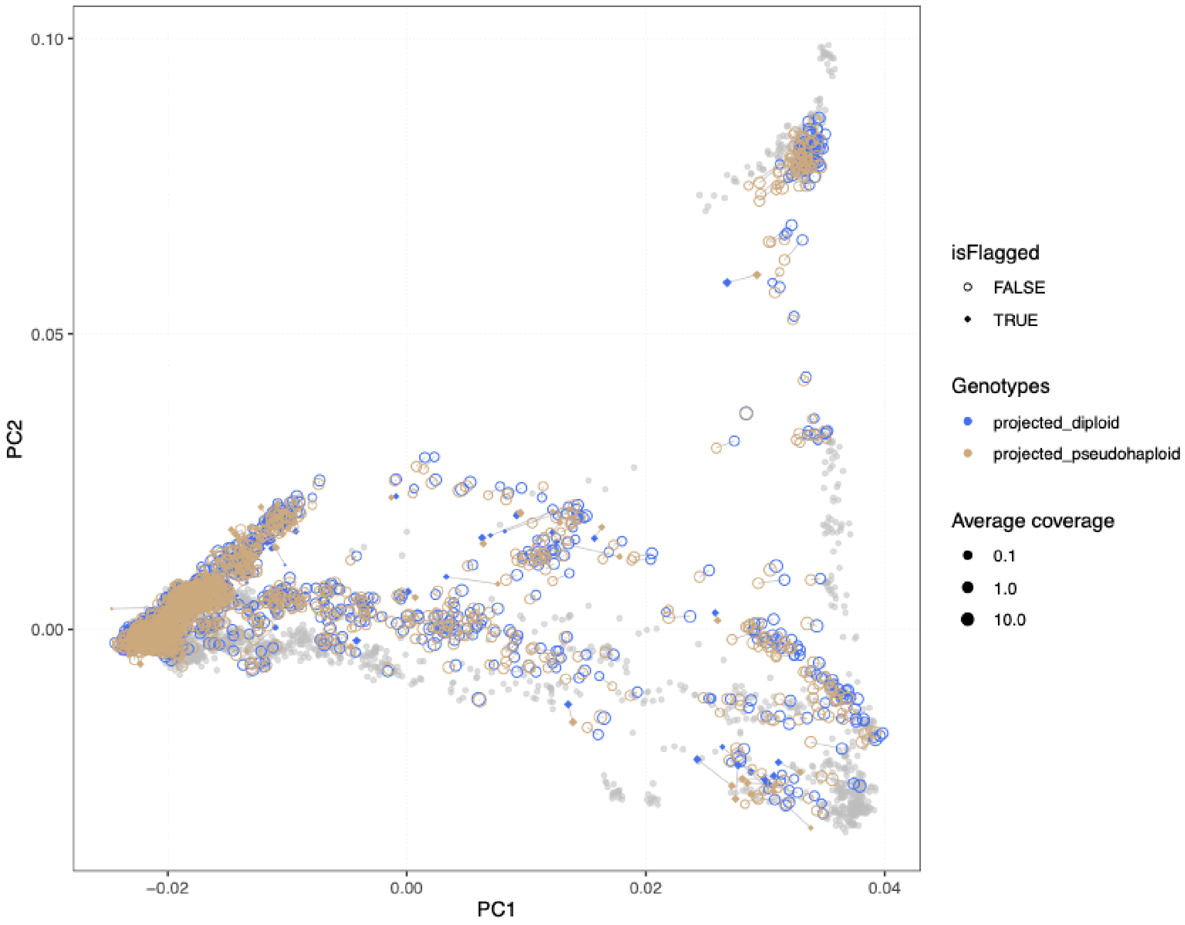


***Fig. S3d.1. PCA projection of imputed and pseudo-haploid genotypes.*** *Coloured symbols show the position of imputed diploid (blue) and pseudo-haploid (beige) genotypes for each ancient individual, projected onto principal components inferred from modern individuals from Eurasia, Oceania, and the Americas. Genotype pairs from the same individual are connected by grey lines.*

(hi-res https://www.dropbox.com/s/fsok1ekf6fp1tax/sfig_structure_01_pca_ooa.png?dl=0)

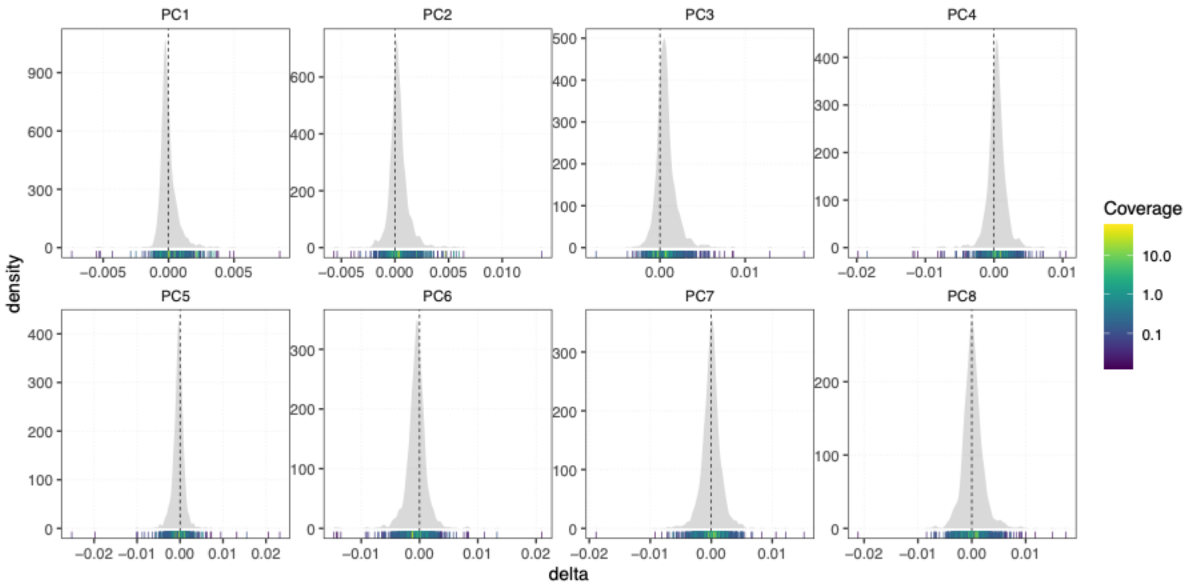


***Fig. S3d.2. Distribution of differences between genotypes in PC space.*** *Density plots show differences along individual PCs between imputed and pseudo-haploid genotypes for each individual in Fig. S3d.1, as a function of their average read depth. Marginal rug plots show individual observations, coloured by the average read depth of the respective individual.*

(hi-res [https://www.dropbox.com/s/zijj2o9u5ia2b3a/sfig_structure_02_pca_ooa_delta_dist.png?dl=0](https://www.dropbox.com/s/zijj2o9u5ia2b3a/sfig_structure_02_pca_ooa_delta_dist.png?dl=0))

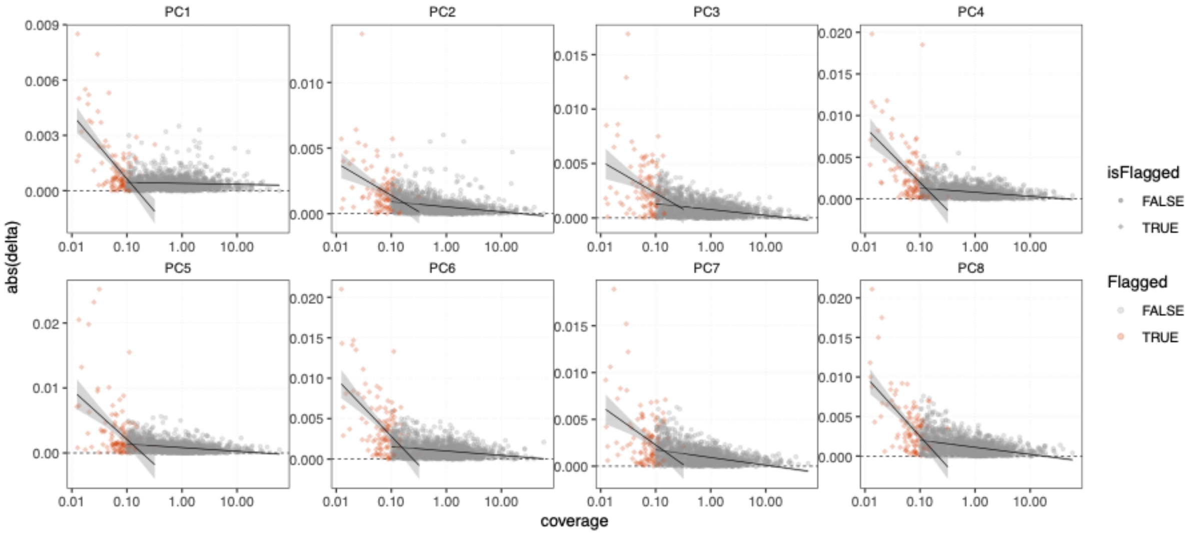


***Fig. S3d.3. Relationship of read depth and PCA position.*** *Plot shows absolute value of differences along individual PCs between imputed and pseudo-haploid genotypes for each individual in Fig. S3d.1, as a function of their average read depth. Individuals flagged for low coverage or low GP average are indicated with red symbols. Linear regression lines for flagged and unflagged individuals are shown with black lines.*

(hi-res [https://www.dropbox.com/s/hnz1665nqpdjven/sfig_structure_03_pca_ooa_delta_dp.png?dl=0](https://www.dropbox.com/s/hnz1665nqpdjven/sfig_structure_03_pca_ooa_delta_dp.png?dl=0))

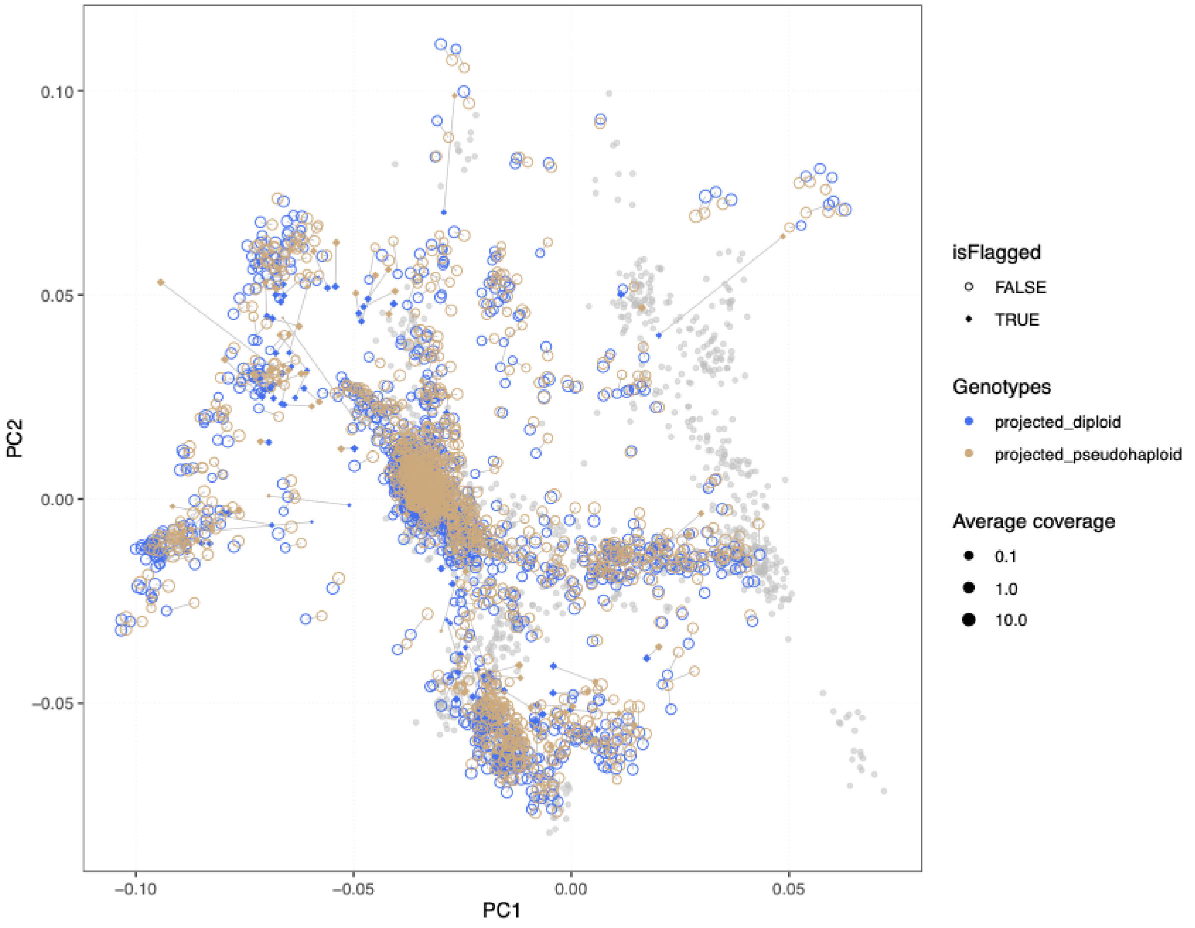


***Fig. S3d.4. PCA projection of imputed and pseudo-haploid genotypes.*** *Coloured symbols show the position of imputed diploid (blue) and pseudo-haploid (beige) genotypes for each ancient individual, projected onto principal components inferred from modern individuals from Western Eurasia. Genotype pairs from the same individual are connected by grey lines.*

(hi-res [https://www.dropbox.com/s/7kh9oqbbrv4h8du/sfig_structure_04_pca_eurWAsia.png?dl=0](https://www.dropbox.com/s/7kh9oqbbrv4h8du/sfig_structure_04_pca_eurWAsia.png?dl=0))

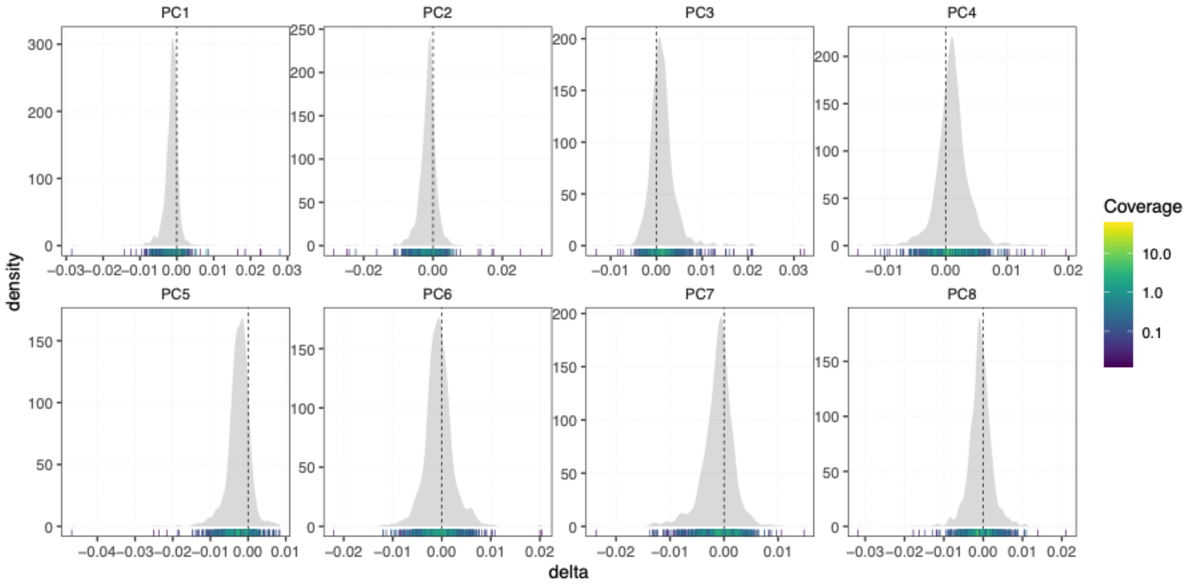


***Fig. S3d.5. Distribution of differences between genotypes in PC space.*** *Density plots show differences along individual PCs between imputed and pseudo-haploid genotypes for each individual in Fig. S3d.4, as a function of their average read depth. Marginal rug plots show individual observations, coloured by average read depth of the respective individual.*

(hi-res https://www.dropbox.com/s/v8wrjxqqpxkduz6/sfig_structure_05_pca_eurWAsia_delta_dist.png?dl=0)

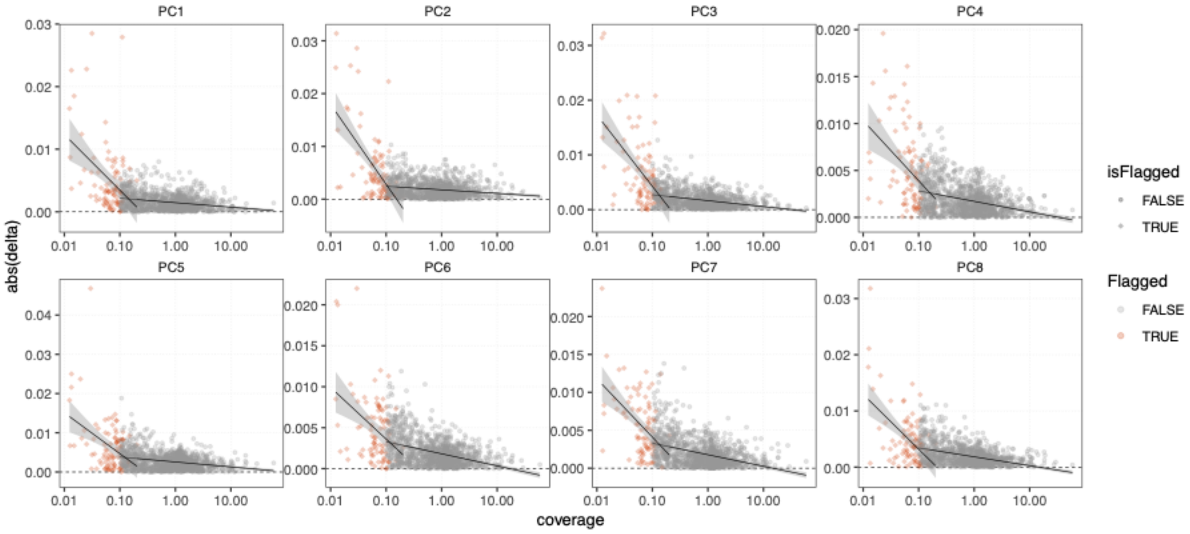


***Fig. S3d.6. Relationship of read depth and PCA position.*** *Plot shows absolute difference along individual PCs between imputed and pseudo-haploid genotypes for each individual in Fig. S3d.4, as a function of their average read depth. Individuals flagged for low coverage or low GP average are indicated with red symbols. Linear regression lines for flagged and unflagged individuals are shown with black lines.*

(hi-res https://www.dropbox.com/s/6bhlweqbosi6os8/sfig_structure_06_pca_eurWAsia_delta_dp.png?dl=0)

#### PCA position of samples flagged as contaminated

To investigate the effect of elevated contamination estimates on the position of individuals flagged as possibly contaminated, we projected them onto the principal components inferred from modern and ancient individuals passing all filters. We found that the majority of those individuals projected consistently with ancient samples of related age and regional contexts (Fig. S3d.7). An exception to this is seen in the Mesolithic Danish individual NEO1, which shows a clear shift towards present-day Europeans along PC1 and PC2. Overall, our results suggest that inferences about broad patterns of deep Eurasian population structure are likely not affected in the majority of the flagged individuals. We nevertheless opted for a conservative approach and excluded those individuals from in-depth analyses further downstream.

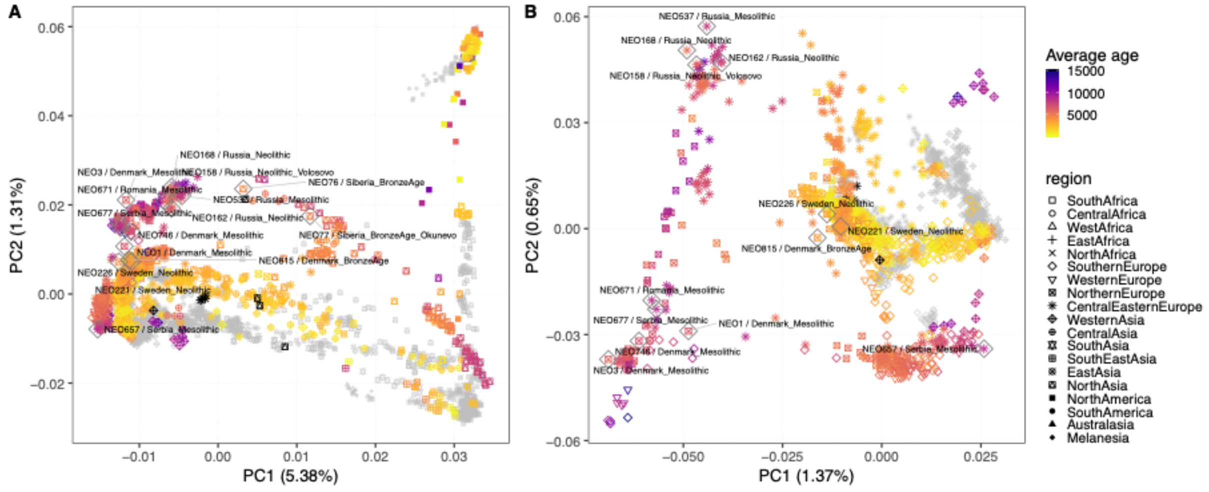


**Fig. S3d.7. PCA positions of individuals flagged as contaminated.** Flagged individuals are labelled and outlined with grey diamonds. Principal components were inferred using ancient and modern individuals from (A) Eurasia, Oceania, and the Americas or (B) Western Eurasia. Plot symbols indicate geographic region, coloured by age of the respective individual. Present-day individuals are indicated in grey.

(hi-res https://www.dropbox.com/s/29a8k7qef6nrrkl/sfig_structure_07_pca_contam.png?dl=0)

#### Genetic ancestry of newly reported samples

##### Overview

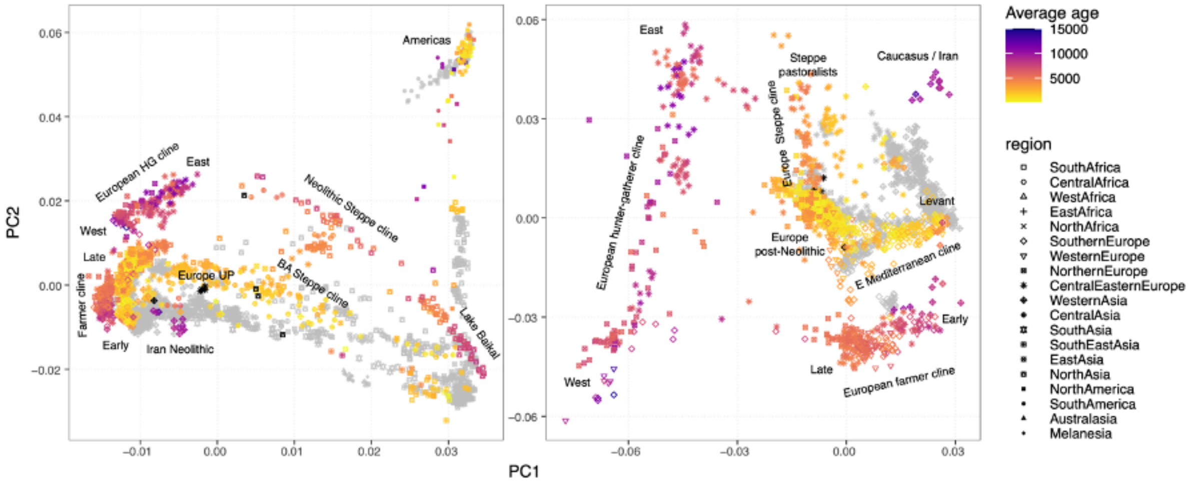


***Fig. S3d.8. Overview of genetic structure****. PCA of ancient and modern individuals from Eurasia, Oceania, and the Americas (left), or Western Eurasia (right). Plot symbols indicate geographic regions, coloured by the age of the respective individual. Present-day individuals are indicated in grey. Terms for spatiotemporal ancestry clusters and clines are indicated.*

(hi-res https://www.dropbox.com/s/mkgel7s93elvvnv/sfig_structure_08_pca_overview.png?dl=0)

Genetic structure in a PCA using 3,316 individuals from regions outside Africa is dominated by continental-scale differentiation among western Eurasia (defined here as west of the Urals), east Asia and the Americas (Fig S3d.8). Two west-east clines of ancient individuals connect western and eastern Eurasia: A “Neolithic Steppe cline” between hunter-gatherers of the West Siberian Forest Steppe and Lake Baikal; as well as a later “BA Steppe cline” linking Western Steppe pastoralists with the Altai mountain region (Fig S3d.8).

Focusing the PCA on 2,126 modern and ancient individuals from Western Eurasia, the extremes of the PCA space are defined by clines and clusters related to previously described “deep” ancestry sources, including: A “European hunter-gatherer cline” between western and eastern European Mesolithic individuals; A “European farmer cline” ranging from early Neolithic individuals from Anatolia and Southern Europe to mid- and late Neolithic European individuals; and hunter-gatherers and early farmers from Iran and the Caucasus. European individuals from the late Neolithic and early Bronze Age onwards form an extended “European post-Neolithic” cluster in the centre of the PCA, differentiated along either a “European Steppe cline” between Steppe pastoralists and late European farmers, or an

“Eastern Mediterranean cline” anchored in the east by Anatolian and Levantine Bronze Age individuals (Fig S3d.8). The newly reported genomes from western Eurasia cluster across the entire range of the PCA, resulting in increased fine-scale resolution along the major ancestry clines, particularly the European hunter-gather and farmer clines. The following sections provide regional descriptions for the patterns of ancestry observed in the newly reported samples.

#### Southern Europe

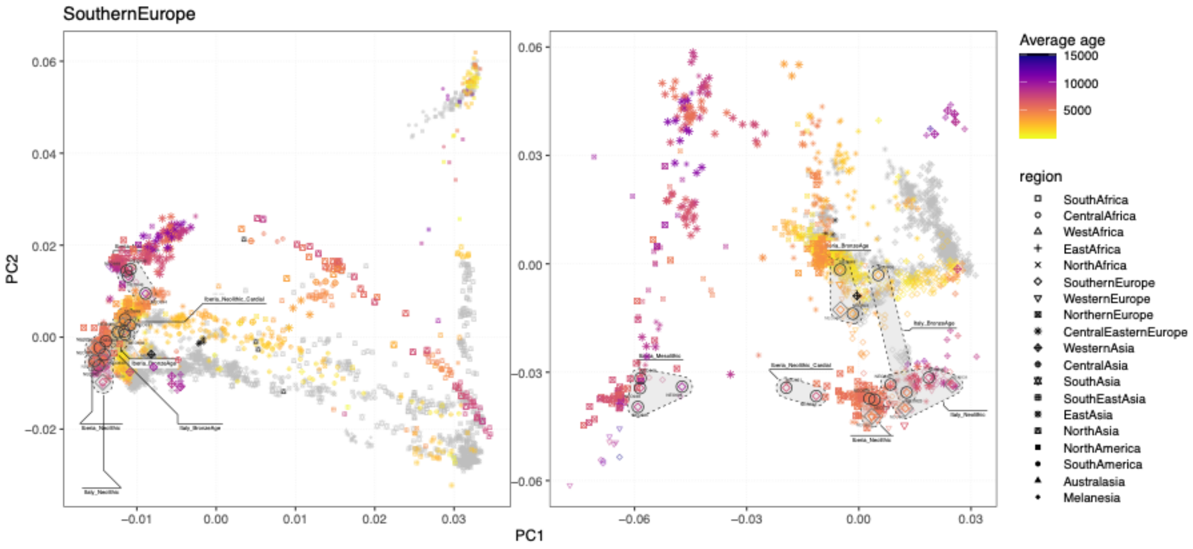


***Fig. S3d.9. Newly reported individuals from Southern Europe.*** *PCA positions of newly reported individuals are highlighted with black circles (imputed) or grey diamonds (pseudo-haploid, projected). Individuals from the same spatiotemporal group are connected with shaded hulls.*

(hi-res https://www.dropbox.com/s/hdgnvp3hb1dfyhe/sfig_structure_09_pca_regions_europeS.png?dl=0)

We report 18 new individuals from Italy (6) and the Iberian Peninsula (12), distributed across European hunter-gatherer (HG), farmer, and post-Neolithic ancestry clusters (Fig S3d.9). Four Iberian Mesolithic individuals cluster with other Southern European Mesolithic individuals at the “western” end of the European HG cline. Among the individuals falling within the European farmer cline, two early Neolithic individuals from Portugal (NEO631, NOE632; ~ 7,300 BP) are shifted towards the European HG cline suggestive of increased HG ancestry. The four most recent individuals (from ~4,100 BP) form part of the extended European post-Neolithic cluster.

Western Europe

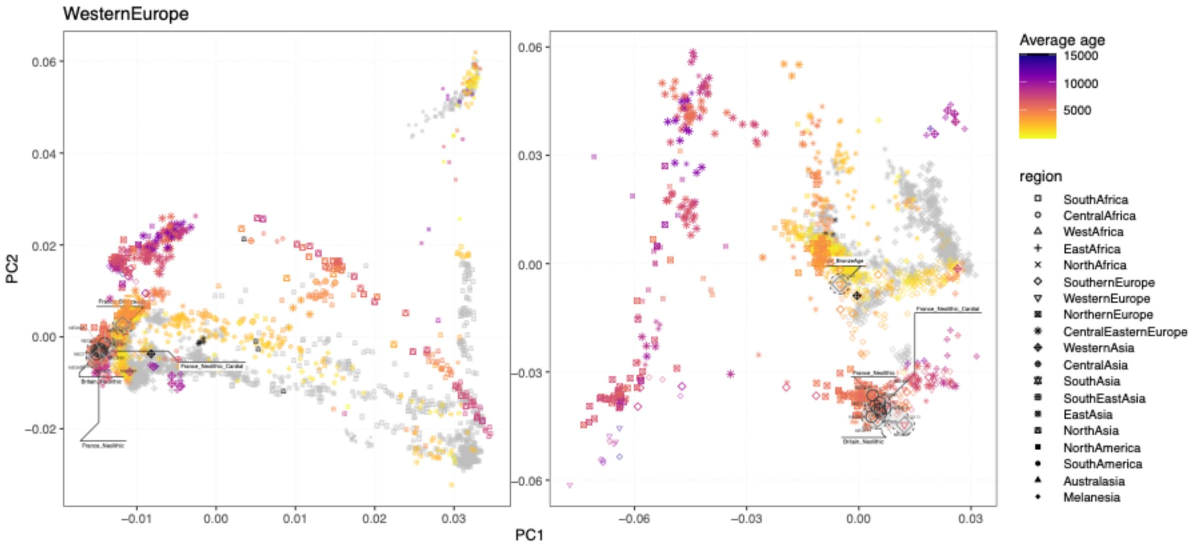


***Fig. S3d.10. Newly reported individuals from Western Europe.*** *PCA positions of newly reported individuals are highlighted with black circles (imputed) or grey diamonds (pseudo-haploid, projected). Individuals from the same spatiotemporal group are connected with shaded hulls.*

(hi-res https://www.dropbox.com/s/1rngjx21530ri8i/sfig_structure_10_pca_regions_europeW.png?dl=0)

We report 12 new individuals from France (5) and the UK (7), from the early Neolithic to the Bronze Age. All 11 Neolithic individuals fall within the European farmer cline, whereas a single Bronze Age individual from Grotte Mandrin (NEO120, ~3,400BP) clustered with post-Neolithic Europeans (Fig. S3d.10).

Northern Europe

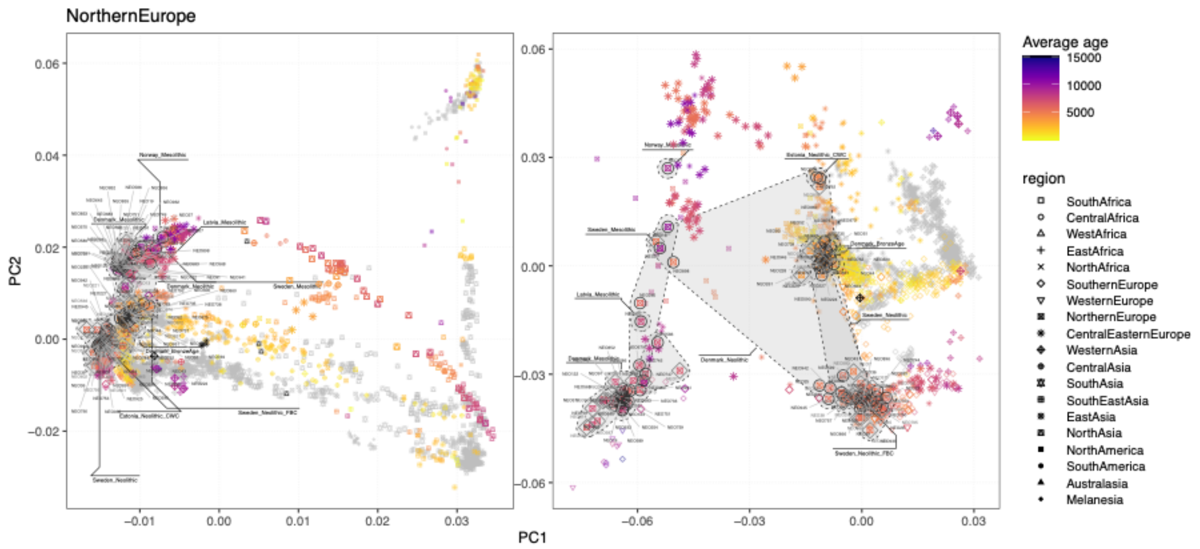


***Fig. S3d.11. Newly reported individuals from Northern Europe.*** *PCA positions of newly reported individuals are highlighted with black circles (imputed) or grey diamonds (pseudo-haploid, projected). Individuals from the same spatiotemporal group are connected with shaded hulls.*

(hi-res https://www.dropbox.com/s/e83qh8i5rrvbclj/sfig_structure_11_pca_regions_europeN.png?dl=0)

We report 124 new individuals from Denmark (100), Sweden (21), Norway (1) and the Baltic (2), spanning a period from ~10,500BP to 3,100BP. This transect includes 46 Mesolithic individuals, all of which cluster within the European HG cline (Fig. S3d.10). The 40 HG individuals from Denmark fall towards the “western” end of the cline, whereas the other Scandinavian and Baltic individuals occupy varied positions shifted towards the “eastern” end of the cline. Neolithic Scandinavian individuals generally fall towards the “late” end of the European farmer cline, with later Neolithic individuals also found among the extended post-Neolithic Europe cluster. Three late Neolithic (~4,500BP) individuals from Denmark (NEO876, NEO792) and Estonia (NEO306) are shifted further up along PC2 towards the Steppe pastoralist cluster, suggesting higher amounts of Steppe-related ancestry (Fig. S3d.11).

Central and Eastern Europe

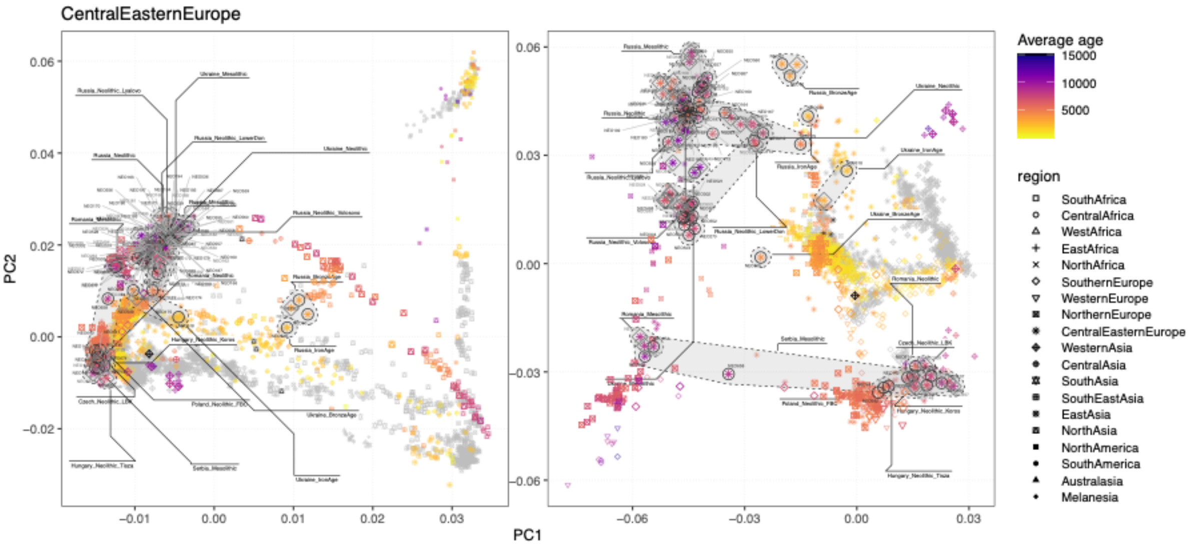


***Fig. S3d.12. Newly reported individuals from Southern Europe.*** *PCA positions of newly reported individuals are highlighted with black circles (imputed) or grey diamonds (pseudo-haploid, projected). Individuals from the same spatiotemporal group are connected with shaded hulls.*

(hi-res https://www.dropbox.com/s/4y3md5tx85udu4n/sfig_structure_12_pca_regions_europeCE.png?dl=0)

We report 112 new individuals from Central and Eastern Europe, falling into two distinct groups. The 92 individuals from Russia (57) and Ukraine (35) predominantly occupy a broad area between the centre and "eastern" end of the European HG cline, roughly corresponding to a geographic cline from the south (Ukraine) to the north (Russia) (Fig. S3d.11). Among the southern Russian samples, six individuals from Golubaya Krinitsa in the Middle Don region are shifted on a cline along PC1 towards Iranian and Caucasus Mesolithic and Neolithic at the other extreme, falling close to later Steppe pastoralists from the region.  Three Bronze Age individuals from Northwestern Russia (Bol'shoy Oleni Ostrov; NEO60, NEO61, NEO62) are positioned between the Neolithic and BA Steppe clines, centrally between the West and East Eurasian poles in the extended PCA of all non-Africans (Fig. S3d.12).

The remaining 20 samples from Central and Southeastern Europe include Mesolithic individuals at the "western" end of the European HG cline, as well as early Neolithic individuals on the farmer cline. An early Neolithic individual from Iron Gates, Serbia (NEO658) is found intermediate between the HG and farmer clines, suggestive of recent farmer/HG admixture (Fig. S3d.12).

#### Western Asia

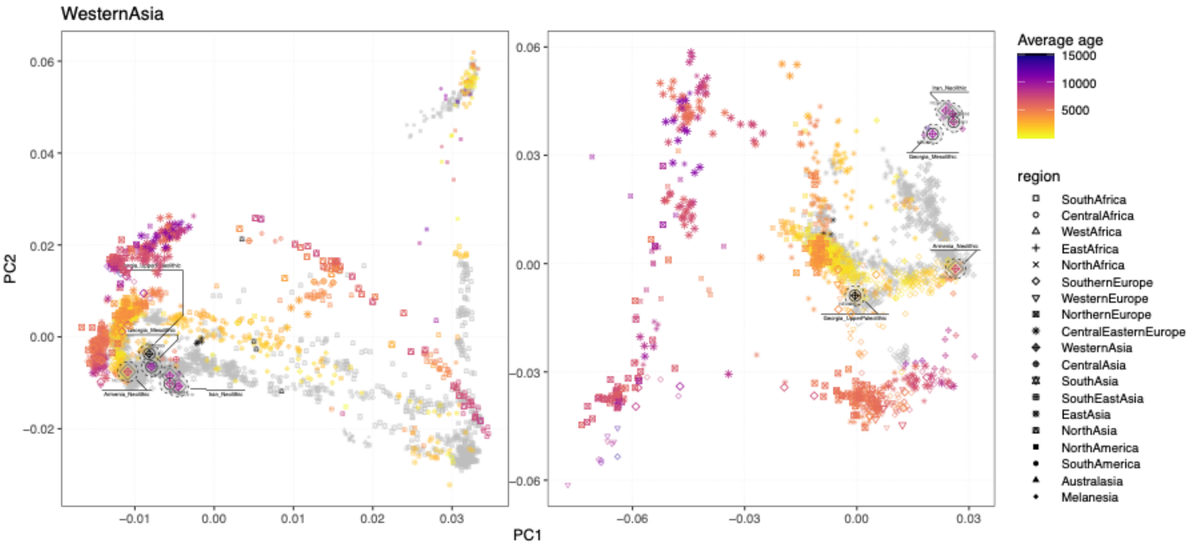


***Fig. S3d.13. Newly reported individuals from Western Asia.*** *PCA positions of newly reported individuals are highlighted with black circles (imputed) or grey diamonds (pseudo-haploid, projected). Individuals from the same spatiotemporal group are connected with shaded hulls.*

(hi-res https://www.dropbox.com/s/8ge5xxsh43jhnav/sfig_structure_13_pca_regions_asiaW.png?dl=0)

We report 6 new individuals from Iran (3) and the South Caucasus region (3). The oldest sample in the dataset, a ~25,000-year-old individual from Georgia is positioned intermediate between Upper Paleolithic Europeans and early Neolithic farmers in both the west Eurasian and extended non-African PCA (Fig. S3d.13). Three Iranian Neolithic individuals (~9,200 BP) as well as one Mesolithic Georgian individual (NEO281; ~9,700 BP) fall with other previously published samples of similar provenance, defining one of the extremes of PC1/PC2 space. One Neolithic individual from Armenia (NEO110, ~7,600 BP) is found at the “eastern” extreme of an eastern Mediterranean cline between ancient Levantine individuals and Southern post-Neolithic Europeans.

#### Central Asia

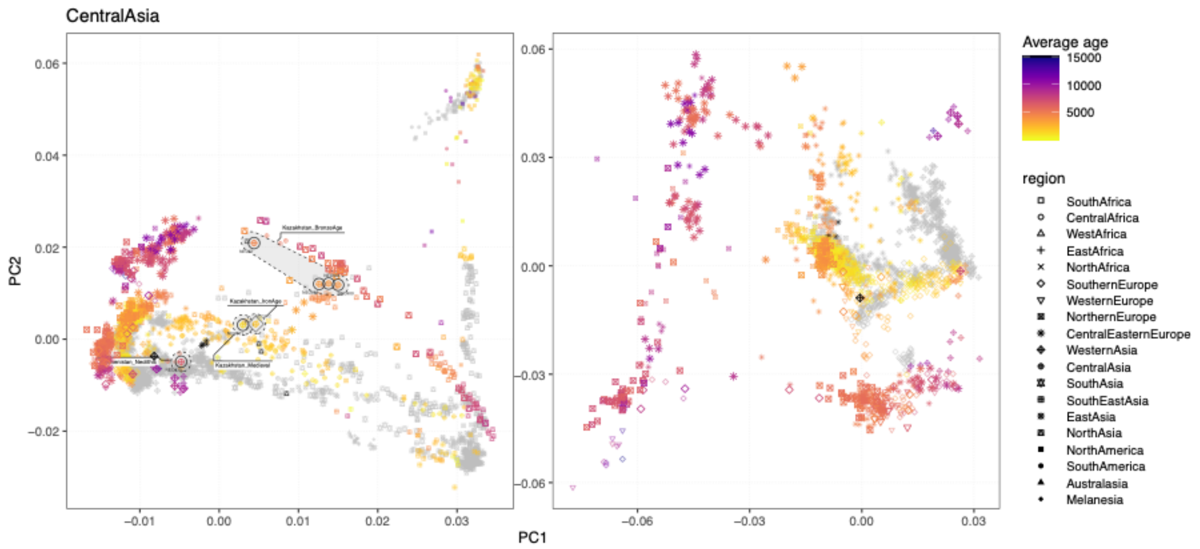


***Fig. S3d.14. Newly reported individuals from Central Asia.*** *PCA positions of newly reported individuals are highlighted with black circles (imputed) or grey diamonds (pseudo-haploid, projected). Individuals from the same spatiotemporal group are connected with shaded hulls.*

(hi-res https://www.dropbox.com/s/ia9a2hik7qw27wl/sfig_structure_14_pca_regions_asiaC.png?dl=0)

We report 7 new individuals from Kazakhstan (6) and Turkmenistan (1). The Neolithic individual from Turkmenistan (~6,500 BP) clusters close to Neolithic Iranians. The individuals from Kazakhstan are more recent (~4,500BP – 2,000BP), with the older individuals forming part of the Neolithic Steppe cline, and the younger individuals along the BA Steppe cline (Fig. S3d.14).

<u>North Asia</u>

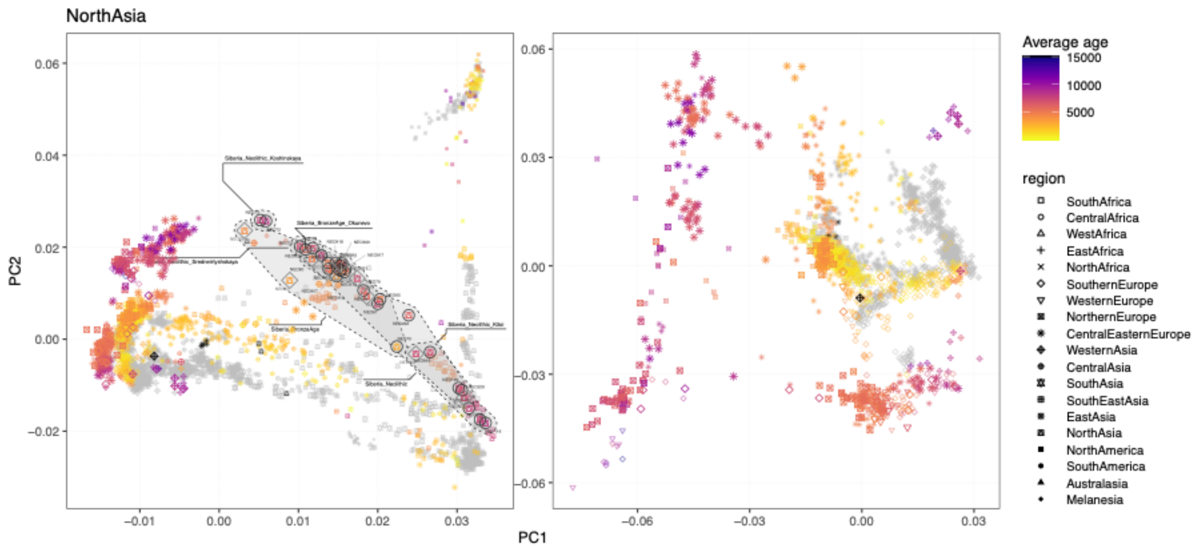


***Fig. S3d.15. Newly reported individuals from North Asia.*** *PCA positions of newly reported individuals are highlighted with black circles (imputed) or grey diamonds (pseudo-haploid, projected). Individuals from the same spatiotemporal group are connected with shaded hulls.*

(hi-res https://www.dropbox.com/s/g53v2ml0r0ojz4w/sfig_structure_15_pca_regions_asiaN.png?dl=0)

We report 38 new individuals from Western Siberia and Lake Baikal, spanning a period from ~8,300BP to 2,800 BP. The individuals fall along the entire range of the Neolithic Steppe cline, spanning from early Forest Steppe hunter-gatherers at the “western” end (NEO72, NEO73) to Lake Baikal hunter-gatherers at the “eastern” end (Fig. S3d.15)

#### Admixture graph fitting of deep Eurasian genomes

We carried out admixture graph fitting using a semi-automatic iterative approach. First, we inferred a well-fitting base graph without admixture edges including seven clusters representing deep world-wide ancestries:

*SouthAfrica_2000BP_1000BP*
*Ethiopia_4500BP*
*UstIshim_45000BP*
*Europe_37000BP_33000BP_Kostenki*
*Europe_37000BP_33000BP*
*Japan_3700BP_2600BP*
*Laos_7800BP*

Onto this base graph, we iteratively added the following additional genetic clusters:

*Yana_31000BP*
*Caucasus_25000BP*
*Malta_24000BP*
*Italy_15000BP_9000BP*
*Iran_10000BP_8500BP*
*Caucasus_13000BP_10000BP*
*RussiaNW_11000BP_8000BP*
*Boncuklu_10000BP*

For each new cluster X, we performed the following fitting procedure:

- Calculate fit for all possible graph topologies adding X as unadmixed leaf
- If no good fit (Z<3) is found, take the 10 best unadmixed graph topologies from the previous step, and generate all possible new topologies adding one admixture edge where X is either source or the target of the new edge.
- If no good fit is found, take the 40 best graph topologies from the previous step and repeat the same procedure, up to a maximum of three admixture edges for each X

The resulting final best-fitting graph is shown in Fig. S3d.16. The newly reported Caucasus UP lineage (NEO283) was best modelled as a mixture of predominantly West Eurasian UP hunter-gatherer ancestry (76%) with ~24% contribution from a “basal Eurasian” ghost population, similar to but with lower amount than West Asian Neolithic individuals. All post-LGM individuals included in the modelling derived some part of their ancestry from the Caucasus UP lineage, suggesting an important role of the descendants related to this lineage in the formation of later West Eurasian populations[5].

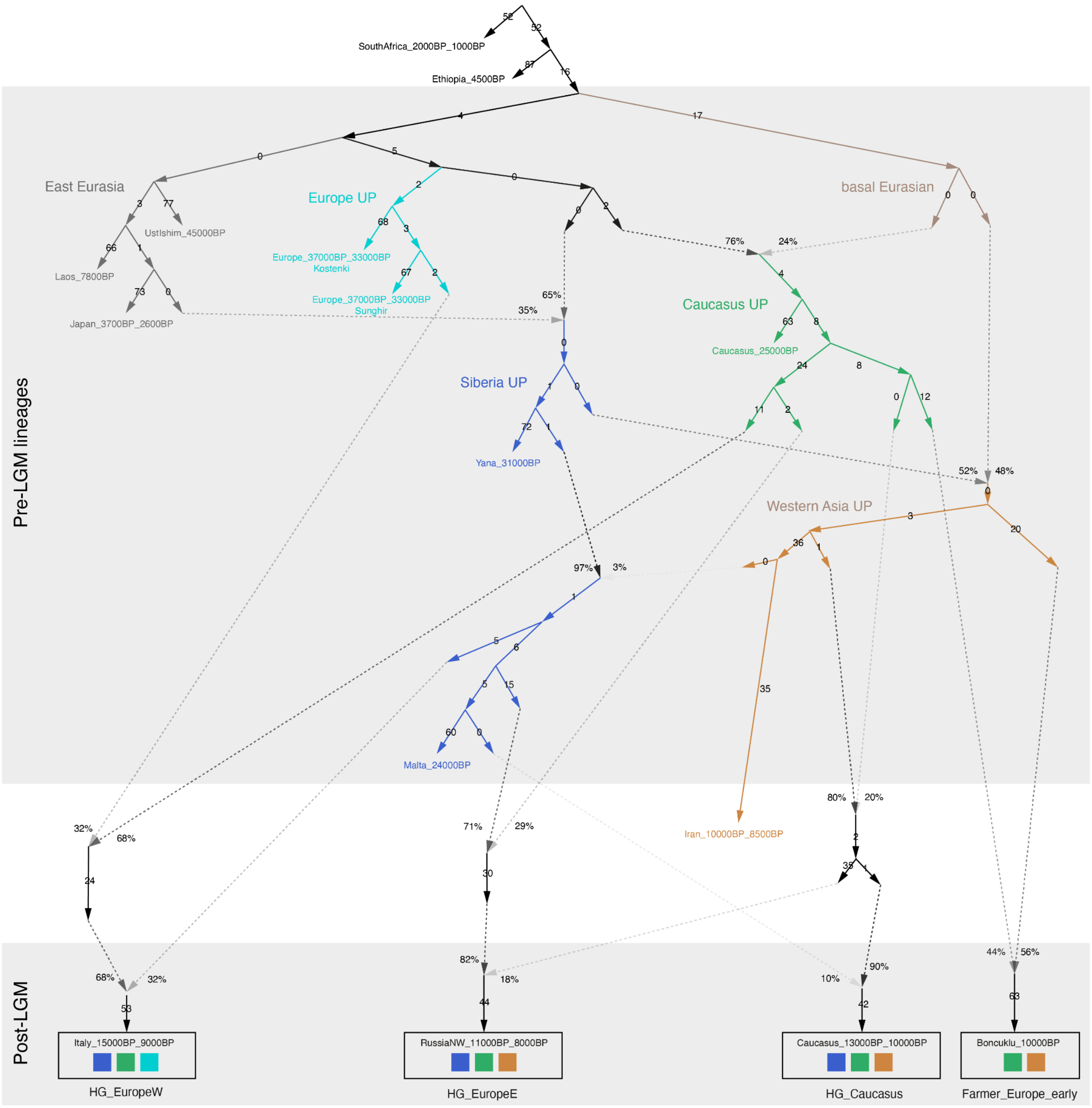


**Fig. S3d.16. Admixture graph of deep Eurasian lineages.** Admixture graph fit relating deep Eurasian lineages predating the Last Glacial Maximum (LGM) to later West Eurasian ancestry clusters (worst |Z| = 3.65).

(hi-res https://www.dropbox.com/s/o0sxqiu0pf9ebrd/sfig_structure_16_admixgraph_deep.png?dl=0)

### 3e) Inferring the spatiotemporal spread of population movements in the past 13 millennia

Fernando Racimo[1]

[1]Lundbeck Foundation GeoGenetics Centre, GLOBE Institute, University of Copenhagen, Copenhagen, Denmark

#### Introduction

We aimed to infer the geographic and temporal spread of major population movements in the past 13 millennia of Western Eurasian history. We used a method developed in Racimo et al.[1], which uses spatiotemporal ordinary kriging on latent ancestry proportion estimates from ancient and present-day genomes. This way, we obtained detailed spatiotemporal maps reflecting the dynamics of the spread of ancestry during the transition from the Mesolithic to the Neolithic, Bronze Age, Iron Age and more recent periods, finally resulting in the complex ancestry make-up of present-day populations in the region.

#### Methods

We first obtained ancestry proportions estimated using ADMIXTURE[2] with K=9 latent ancestry clusters (Supplementary Note S3d) on a sequence dataset including both whole-genome shotgun-sequenced genomes and genomic sequences obtained via SNP capture (Supplementary Note S2, intersection with “HO” dataset). We performed spatiotemporal kriging[3] of these proportions over the last 12,900 years, in intervals of 300 years, with a 5,000-point spatial grid spanning Western and Central Eurasia. We used the R package *gstat* to fit a spatiotemporal variogram via a metric covariance model, and perform ordinary kriging[4]. We focused on the ancestry clusters for which we could fit variogram models that were not static over time.

We then repeated the procedure above, but using the ancestry proportions estimated through IBD clustering, to look at finer-scale genetic structure (Supplementary Note 3f). We focused on cluster assignments for which we had large data across time and space in Western Eurasia, from 12,900 years ago to the present.

#### Results - ADMIXTURE ancestry proportions

We were able to fit spatiotemporal variogram functions to six of the nine ancestries. We label these as WHG, EHG, IRN, LVN, SIB and EAS. The first four are roughly maximised in Mesolithic western European hunter-gatherers, Mesolithic eastern European hunter-gatherers, Iranian Neolithic populations, Levant Neolithic populations and ancient Siberian populations, respectively (see[5] for a model positing the first four of these populations as the major sources of ancestry in present-day Europeans). We depict the spatiotemporal spread of the first four of these ancestries in Fig. S3e.1 and Supplementary Animations 1-4. The fifth ancestry (SIB) occurs at much lower rates in western Eurasia, and rises in frequency in northeastern Europe during the Iron Age (Fig. S3e.2, Supplementary Animation 5)[6–8]. A sixth ancestry (EAS) has affinities to East Asians, and expands into the Caucasus in recent times (Fig. S3e.2, Supplementary Animation 6).

These spatiotemporal maps evince interesting patterns of ancestry change across the landscape. For example, the advancement of Neolithic Levant (LVN) ancestry appears staggered: we observe different periods of advancement followed by stasis. In addition to the Bronze Age movement of EHG ancestry, there is a southern incursion of IRN ancestry via South Europe[9–11]. This is particularly obvious in Bronze Age Greek and Iron Age Roman samples, and may be due to contacts with Anatolia and Northern Africa (where this ancestry is also present). Additionally, we observe small incursions of very late SIB ancestry into Eastern Europe (Fig. S3e.2 Supplementary Animation 5). This signal is driven by the presence of SIB ancestry in Iron Age Cimmerian nomads[12] and in a medieval Serbian[13], and could perhaps be linked to the introduction of languages from the Finno-Ugric family into the Hungarian Plain. An incursion of this ancestry into Western Eurasia can also be seen in Medieval Ottoman Anatolians[14].

We can focus on local timelines of kriged ancestry changes in different points of the map (Fig. S3e.3). Here, we observe that the timing and duration of the rise in LVN ancestry was different in different points in Europe (Fig. S3e.3). We also observe that, in certain regions of Europe, the rise in IRN and EHG ancestry are largely decoupled from each other (see e.g. “Rome” in Fig. S3e.2)[9–11,15].

#### Results - IBD cluster proportions

We were able to fit spatiotemporal variogram functions to seven IBD-based ancestry assignment:

- *Italy_15000BP_9000BP*

- *Boncuklu_10000BP*
- *MiddleDon_7500BP*
- *Caucasus_13000BP_10000BP*
- *SiberiaNE_9800BP*
- *SteppeC_8300BP_7000BP*
- *Ukraine_10000BP_4000BP*

These are depicted in Figure S3e.4 and Figure S3e.5. We also include animations for each of them (Animations S3e.5-11).

#### Figures

High resolution versions of figures are available here:
https://www.dropbox.com/scl/fo/2yp6227x06f761l6udppm/h?rlkey=29n5m7nrblh6sml2ggisvk2cd&dl=0

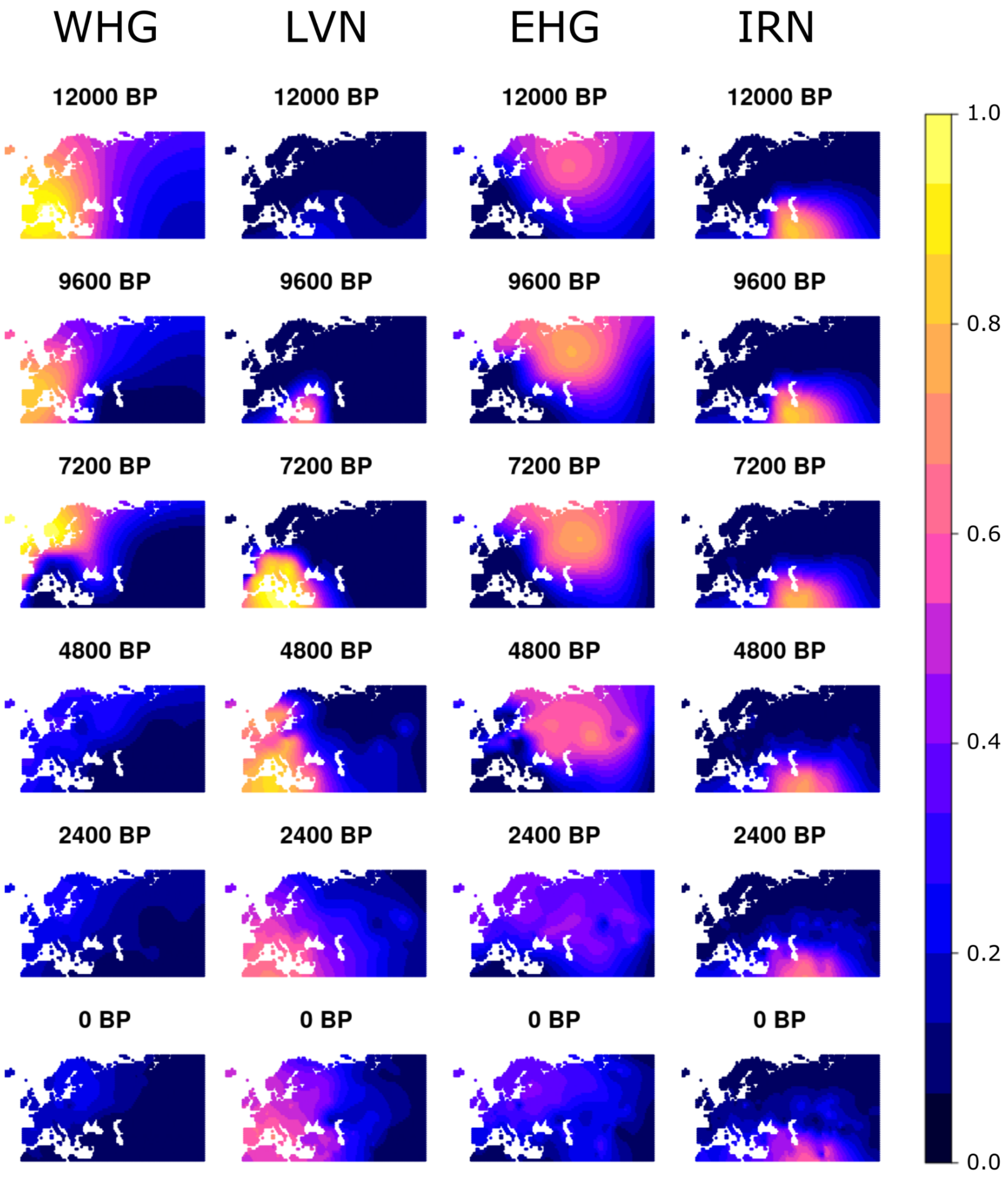


**Figure S3e.1.** Spatiotemporal kriging of four major ADMIXTURE ancestry clusters over the last 12,000 years. WHG = ancestry maximised in western European hunter-gatherers. LVN = ancestry maximised in Anatolian farmer populations. EHG = ancestry maximised in eastern European hunter-gatherers. IRN = ancestry maximised in Iranian Neolithic individuals.

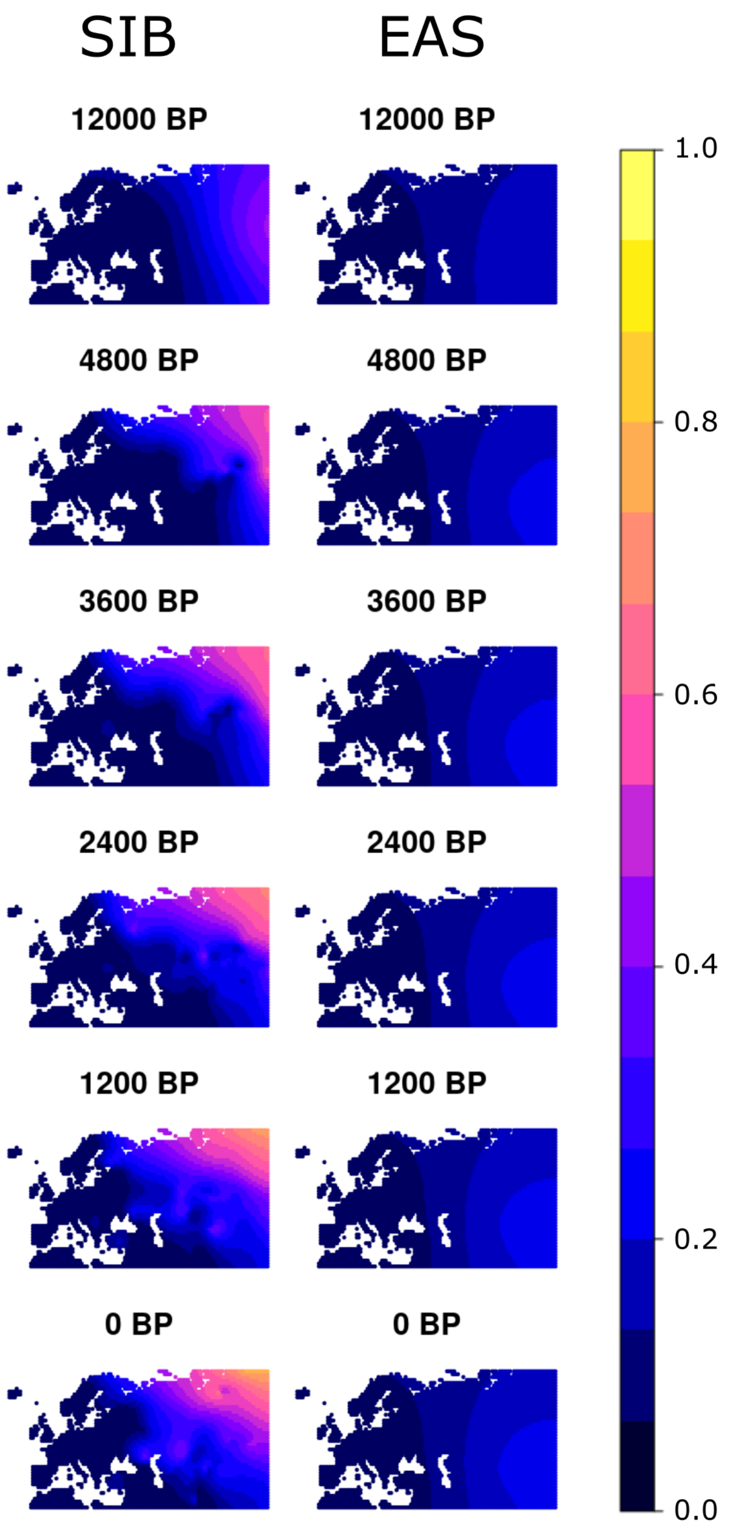


**Figure S3e.2.** Spatiotemporal kriging of two additional ADMIXTURE ancestry clusters with later incursions into Western Eurasia over the last 12,000 years, particularly focusing on the last 5,000 years. SIB = ancestry maximised in ancient Siberian individuals. EAS = ancestry maximised in East Asian individuals.

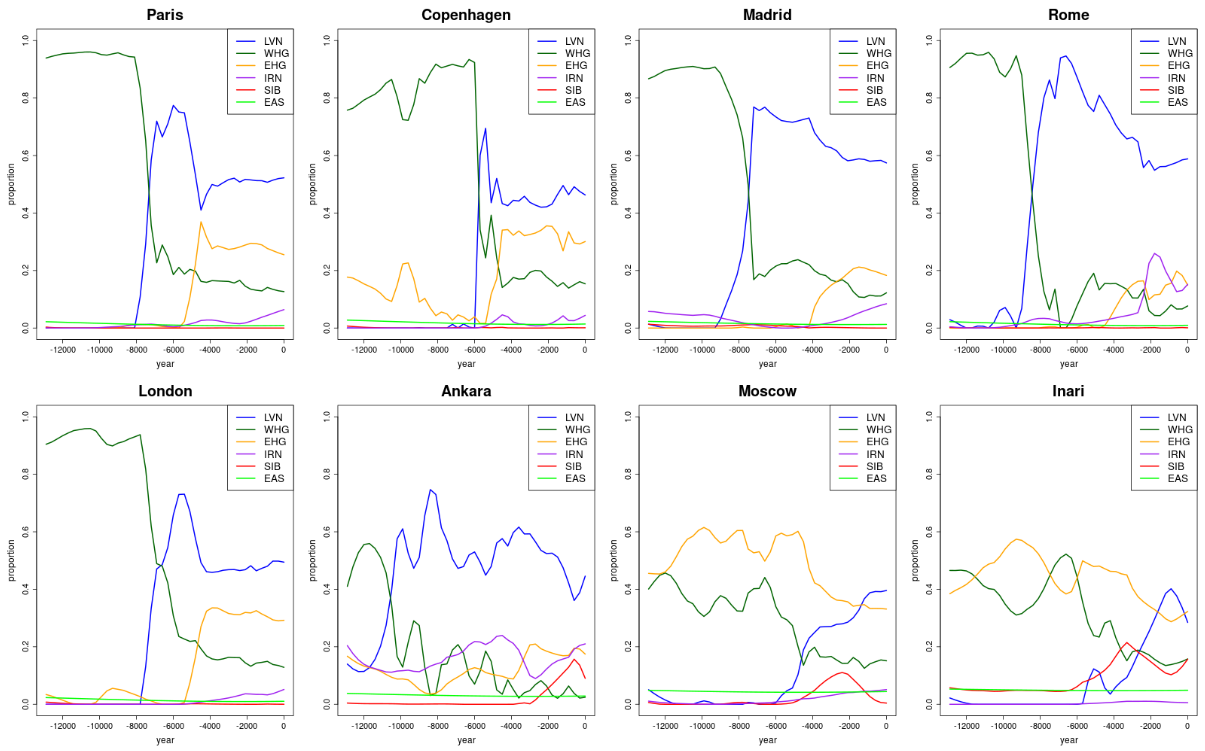


**Figure S3e.3.** Local timelines in what are now 8 urban centres across western Eurasia, reflecting local differences in the tempo and mode of ADMIXTURE ancestry changes over time. LVN = ancestry maximised in Anatolian farmer populations. WHG = ancestry maximised in western European hunter-gatherers. EHG = ancestry maximised in eastern European hunter-gatherers. IRN = ancestry maximised in Iranian Neolithic individuals / Caucasus hunter-gatherers. SIB = ancestry maximised in ancient Siberian individuals. EAS = ancestry maximised in East Asian individuals.

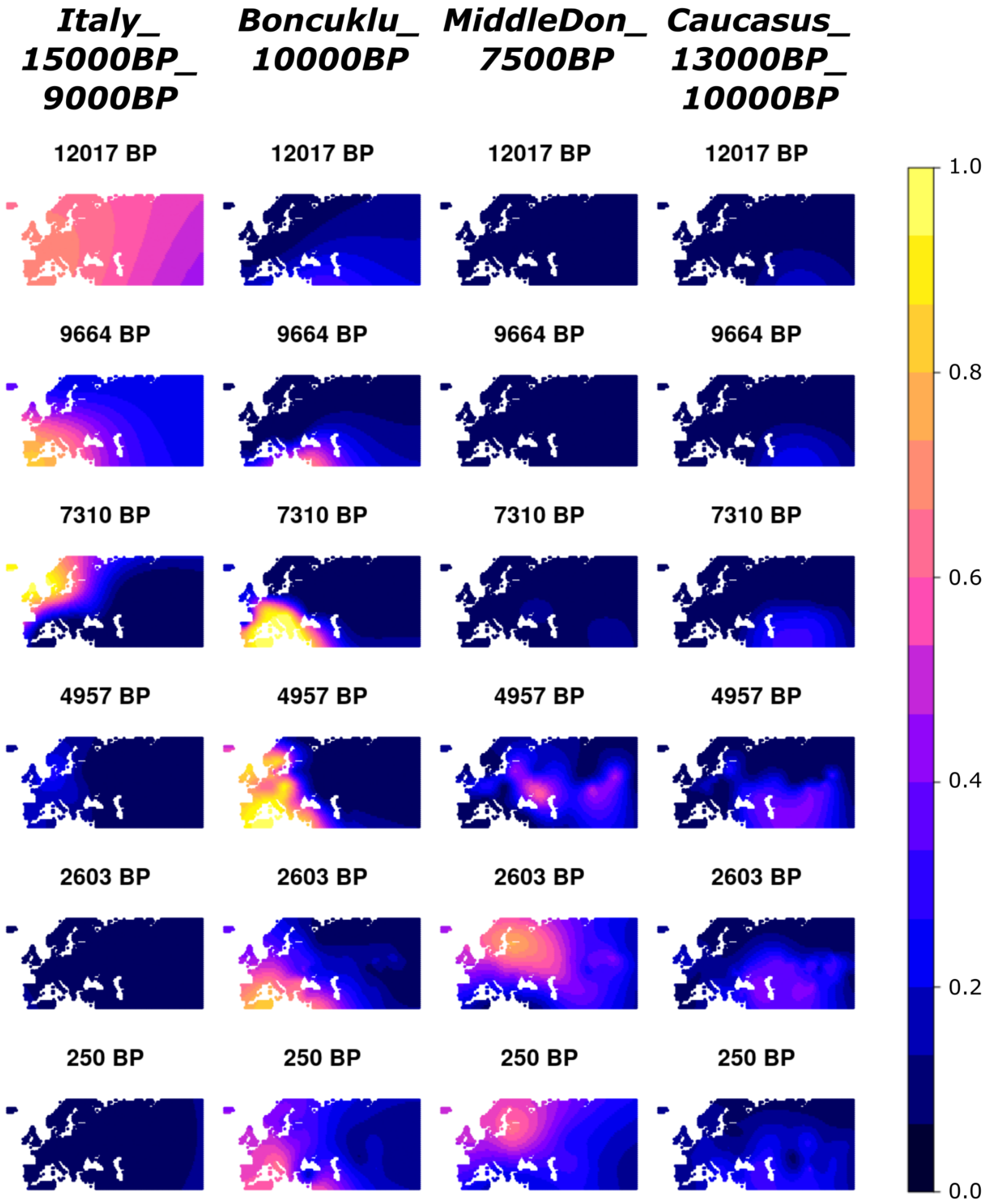


**Figure S3e.4.** Spatiotemporal kriging of four ancestry IBD-based clusters over the last 12,000 years: *Italy_15000BP_9000BP, Boncuklu_10000BP, MiddleDon_7500BP and Caucasus_13000BP_10000BP.*

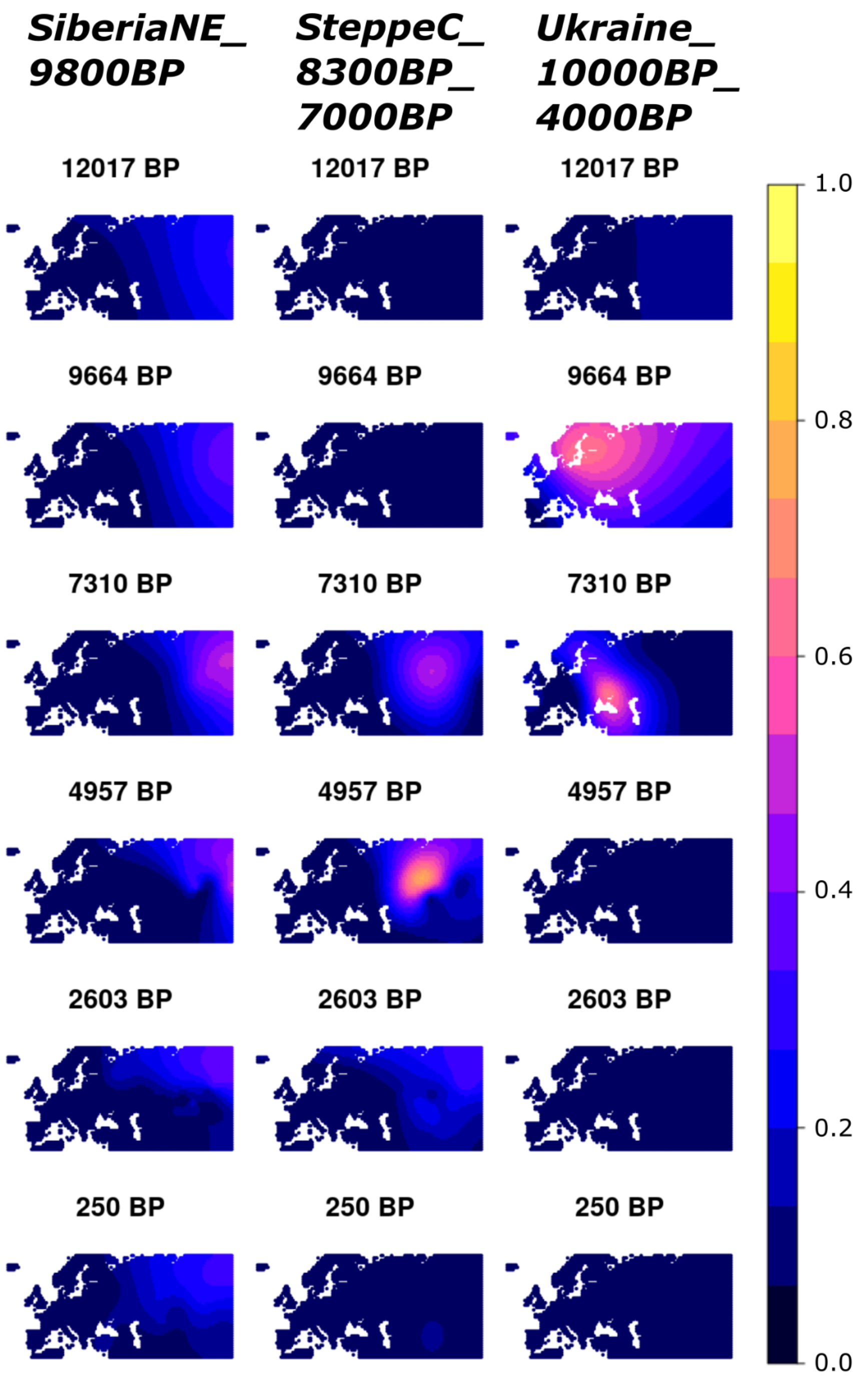


**Figure S3e.5.** Spatiotemporal kriging of three ancestry IBD-based clusters over the last 12,000 year: *SiberiaNE_9800BP, SteppeC_8300BP_7000BP* and *Ukraine_10000BP_4000BP*

**Animation S3e.1. IRN kriged ancestry from ADMIXTURE analysis.**

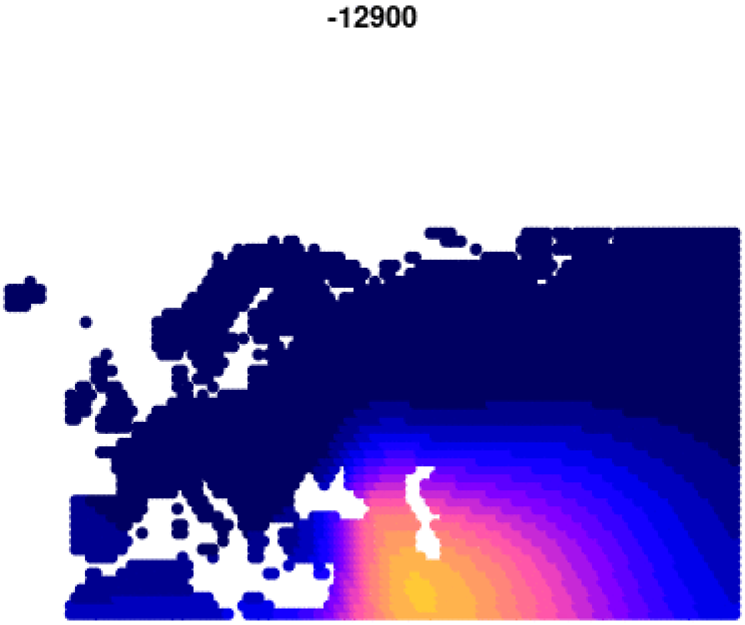


**Animation S3e.2. EHG kriged ancestry from ADMIXTURE analysis.**

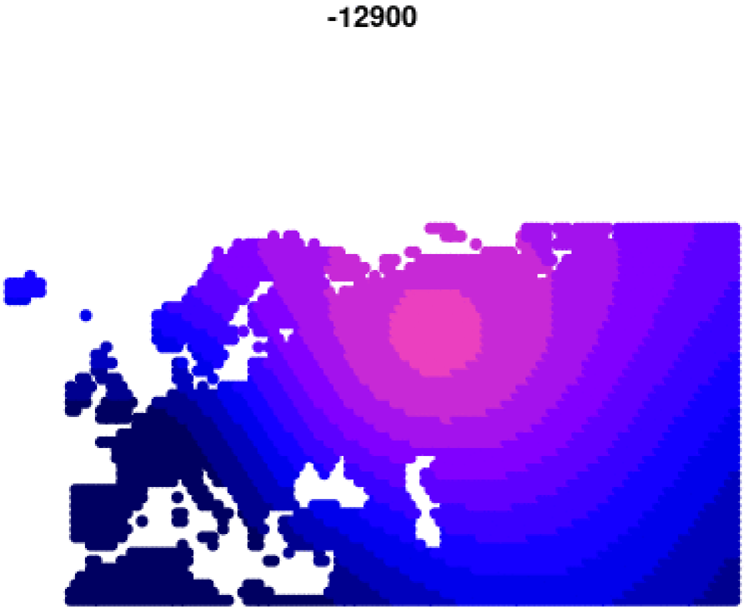

**Animation S3e.3. LVN kriged ancestry from ADMIXTURE analysis.**

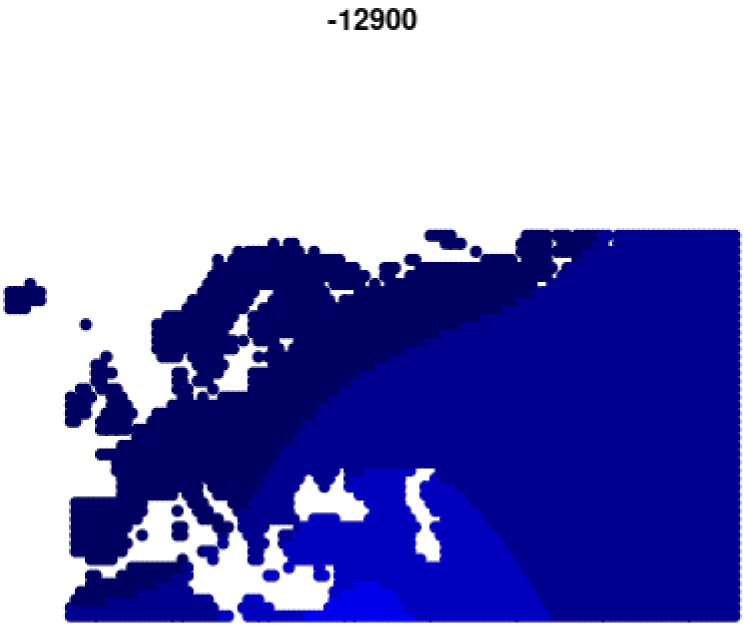


**Animation S3e.4. WHG kriged ancestry from ADMIXTURE analysis.**

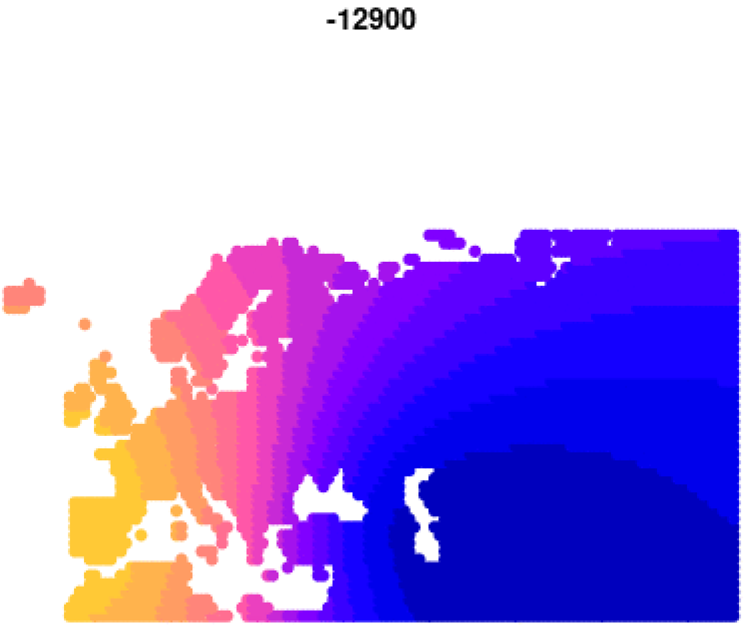

**Animation S3e.5. “Boncuklu_10000BP” kriged ancestry from IBD analysis.**

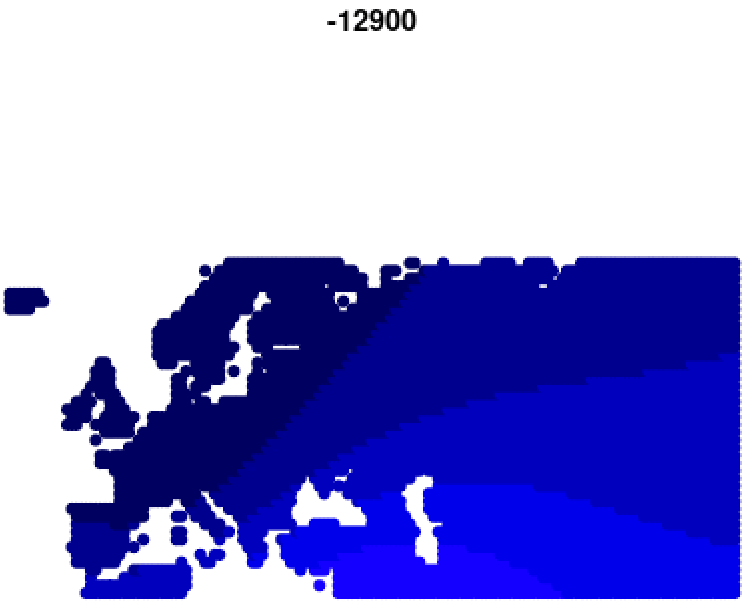


**Animation S3e.6. “Italy_15000BP_9000BP” kriged ancestry from IBD analysis.**

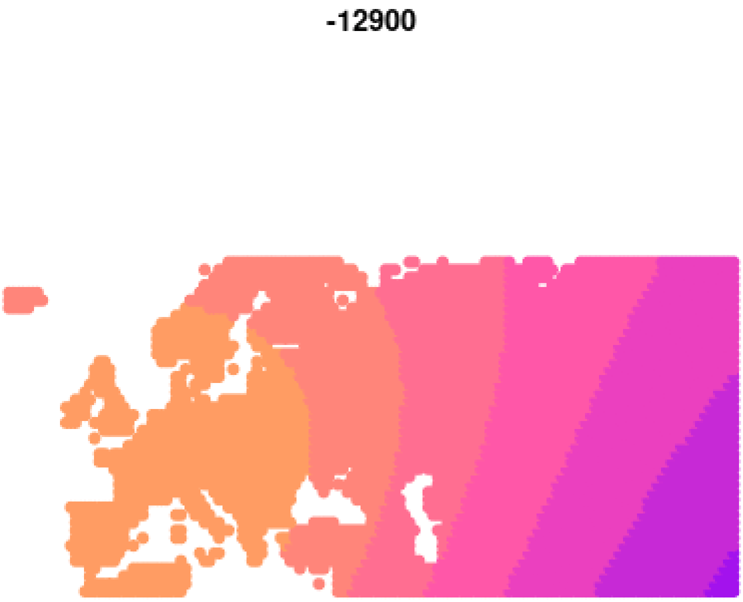

**Animation S3e.7. “MiddleDon_7500BP” kriged ancestry from IBD analysis.**

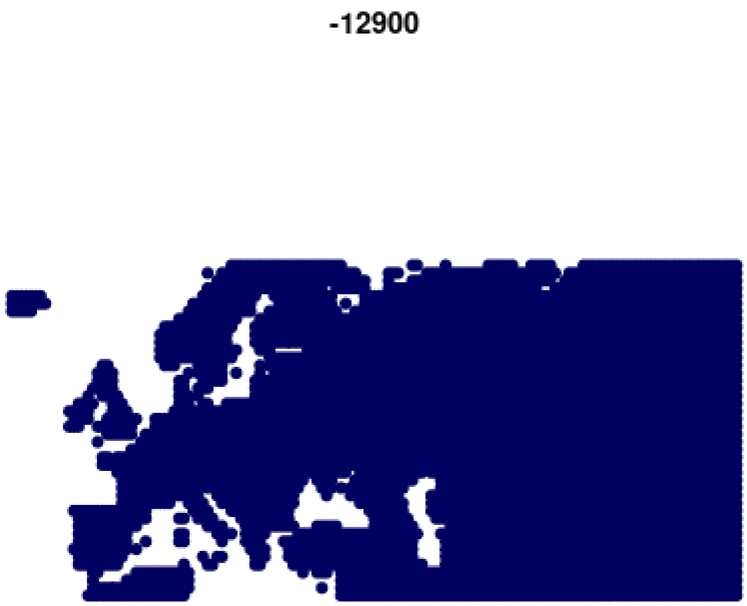


**Animation S3e.8. “Caucasus_13000BP_10000BP” kriged ancestry from IBD analysis.**

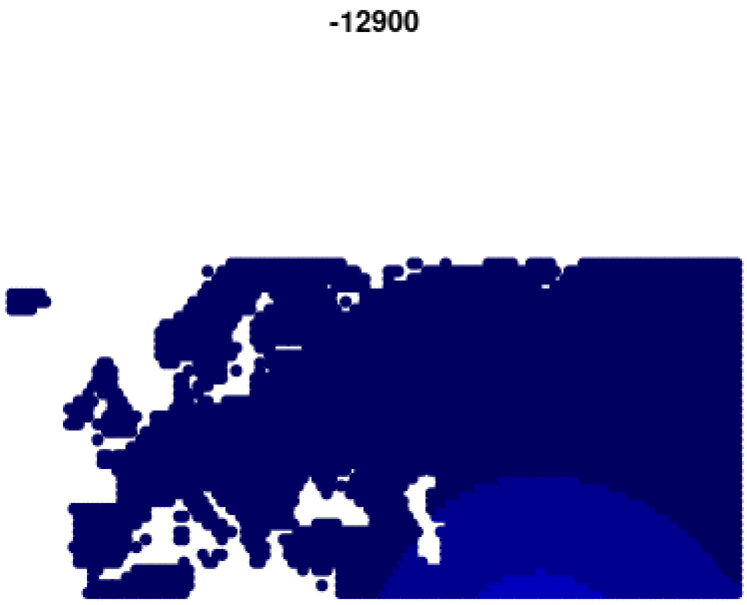

**Animation S3e.9. “SiberiaNE_9800BP” kriged ancestry from IBD analysis.**

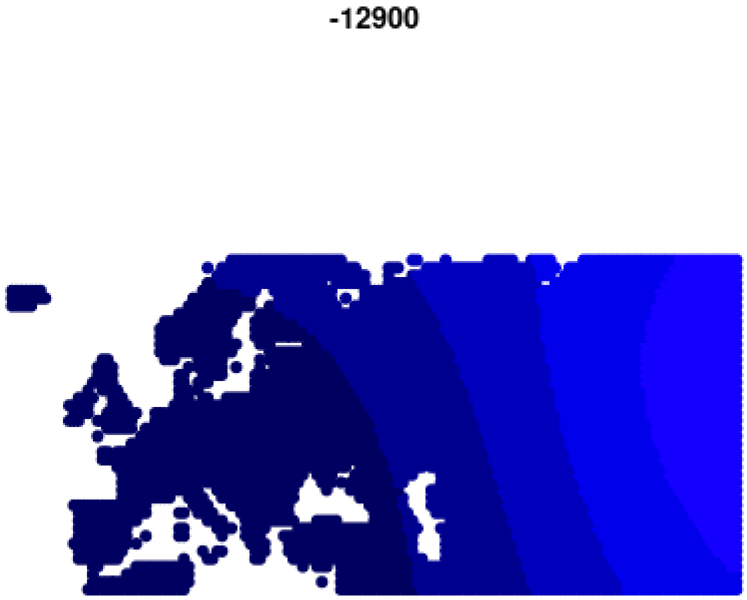


**Animation S3e.10. “SteppeC_8300BP_7000BP” kriged ancestry from IBD analysis.**

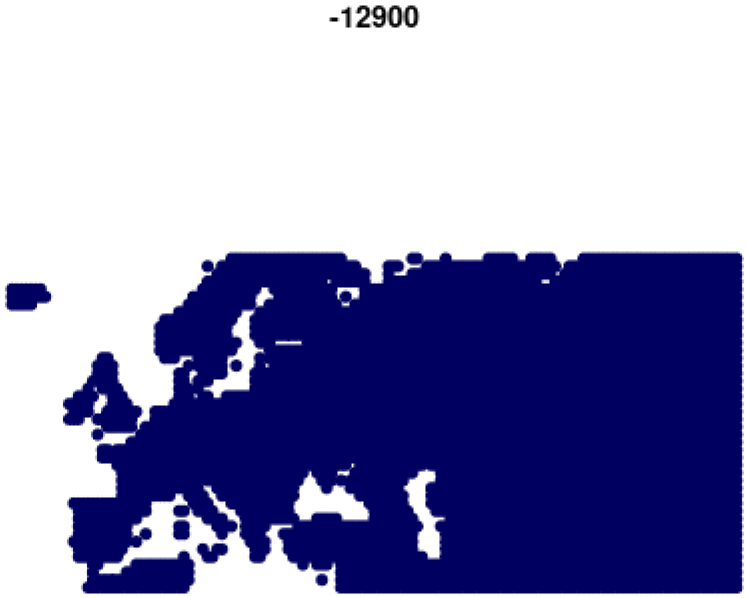

**Animation S3e.11. “Ukraine_10000BP_4000BP” kriged ancestry from IBD analysis.**

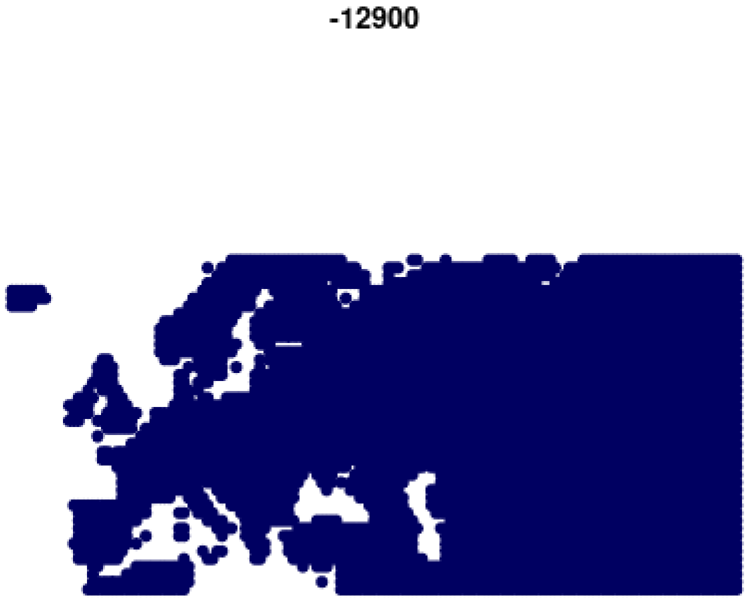

### 3f) HBD/ IBD sharing/ROH/clustering

Martin Sikora[1]
[1]Lundbeck Foundation GeoGenetics Centre, GLOBE Institute, University of Copenhagen, Copenhagen, Denmark

#### Results

##### IBD-based hierarchical graph clustering

We performed hierarchical graph clustering on the 1,492 ancient individuals passing all filters, which were assigned into a final curated set of 122 genetic clusters (Fig. S3f.1). The obtained clusters captured both broad and finer-scale genetic structure, corresponding to shared ancestry within particular spatiotemporal ranges and/or archaeological contexts (Fig. S3f.2). We named these cluster using a “geographic-temporal” nomenclature[11] (e.g. “Denmark_10500BP_6000BP”), in concert with more traditional names for groups of multiple clusters with shared archaeological or subsistence contexts (e.g. “Farmer_Europe_early”) where applicable.

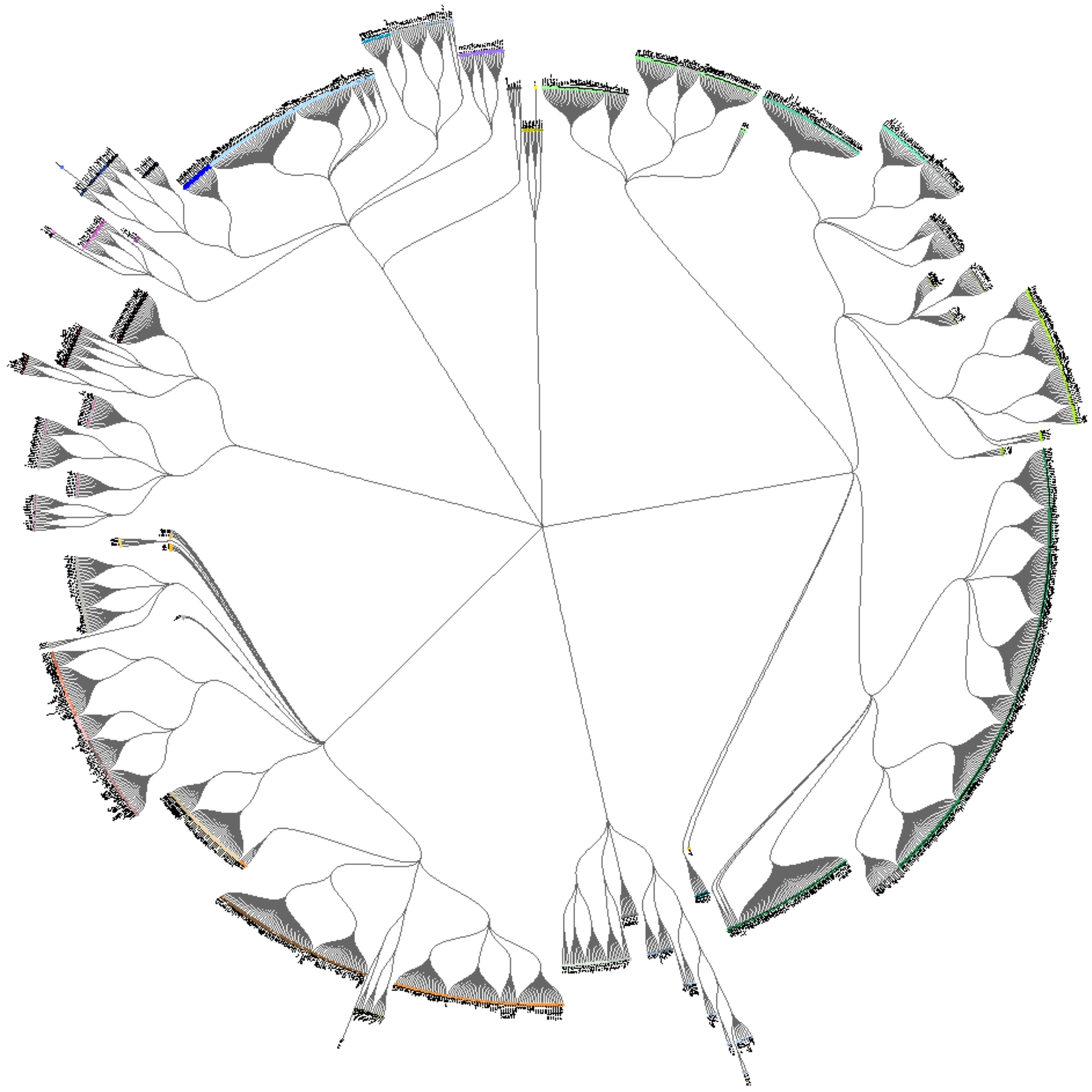

**Fig. S3f.1. Hierarchical graph clustering.** Tree diagram showing final curated hierarchical clustering relationship among the 1,492 imputed ancient genomes passing all filters. Genetic clusters are differentiated using plot symbol colours and shapes.

(hi-res https://www.dropbox.com/s/ozt1vwrm42pr0m3/sfig_ibd_01_cluster_dendro.png?dl=0)

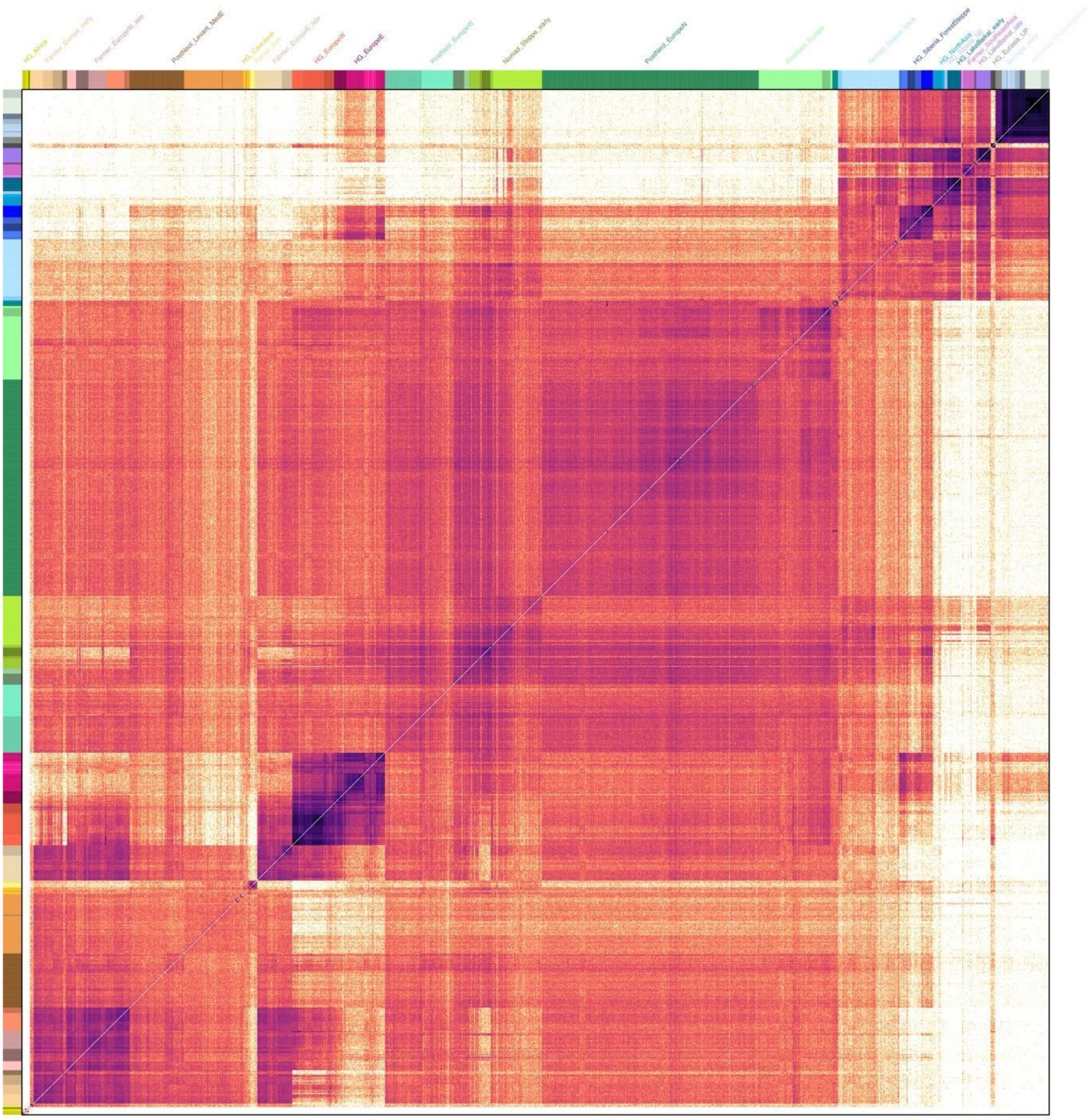


**Fig. S3f.2. IBD sharing similarities.** Heatmap of pairwise IBD-sharing similarities between the 1,492 ancient individuals passing all filters, sorted according to clustering hierarchy. coloured bars indicate cluster membership of individuals. Selected cluster group labels are shown in the top colour bar.
(hi-res https://www.dropbox.com/s/vggfdhb2nb6wzch/sfig_ibd_02_cluster_heatmap.png?dl=0)

At the highest level of the clustering hierarchy, the individuals were partitioned into six global clusters representing broad continent-wide genetic structure.

- Africa_8000BP_400BP
- Europe_15000BP_4000BP

- EuropeWCAsia_25000BP_300BP
- Eurasia_5000BP_200BP
- Asia_45000BP_200BP
- Americas_12000BP_100BP

The following sections provide more detailed descriptions of relevant sub-clusters within the four global clusters from Eurasia.

Europe_15000BP_4000BP

This global cluster includes individuals from western Eurasian Mesolithic and Neolithic contexts with hunter-gatherer ancestry. The individual genetic clusters are partitioned into two cluster groups corresponding to “Eastern hunter-gatherers” and “Western hunter-gatherers” as previously used in the literature (Fig. S3f.3-6):

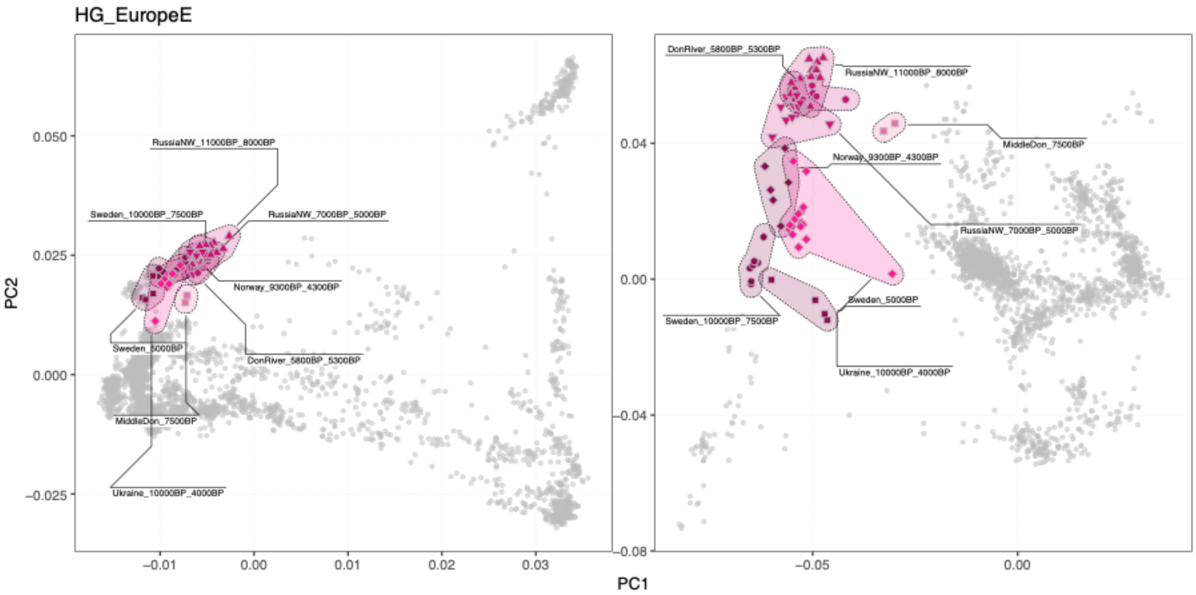


**Fig. S3f.3 PCA for cluster group *HG_EuropeE*.** PCA positions of individuals within specific clusters are highlighted with coloured symbols, and connected with shaded hulls (from PCAs shown in Fig. S3d.7).

(hi-res https://www.dropbox.com/s/a21lvxu9oa7vhkq/sfig_ibd_03_cluster_pca_hgEurE.png?dl=0)

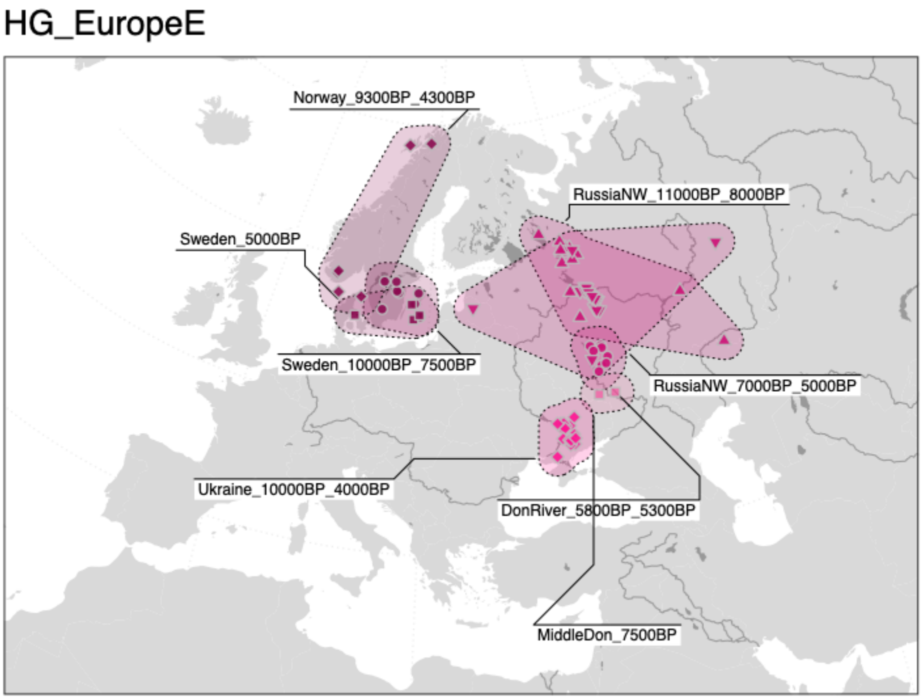


**Fig. S3f.4 Geographic distribution of individuals in cluster group *HG_EuropeE*.** Geographic locations of individuals within specific clusters are highlighted with coloured symbols, and connected with shaded hulls.

(hi-res https://www.dropbox.com/s/590zy1pa1z7tpvd/sfig_ibd_04_cluster_map_hgEurE.png?dl=0)

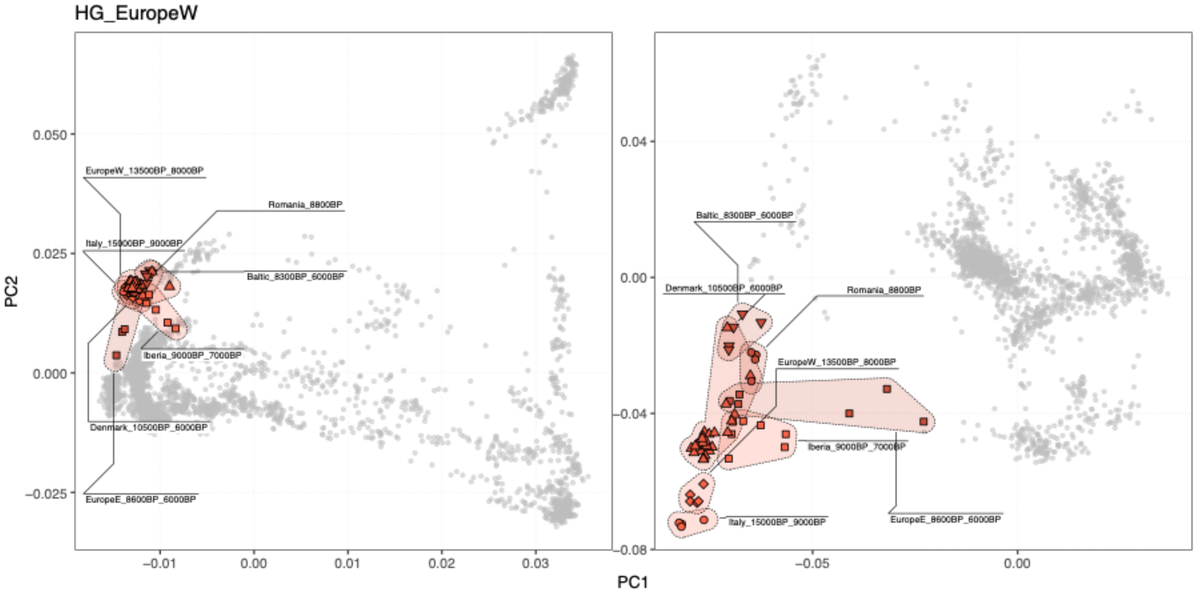


**Fig. S3f.5 PCA for cluster group *HG_EuropeW*.** PCA positions of individuals within specific clusters are highlighted with coloured symbols, and connected with shaded hulls (from PCAs shown in Fig. S3d.7).

(hi-res https://www.dropbox.com/s/q6h8c2b4haqabtp/sfig_ibd_05_cluster_pca_hgEurW.png?dl=0)

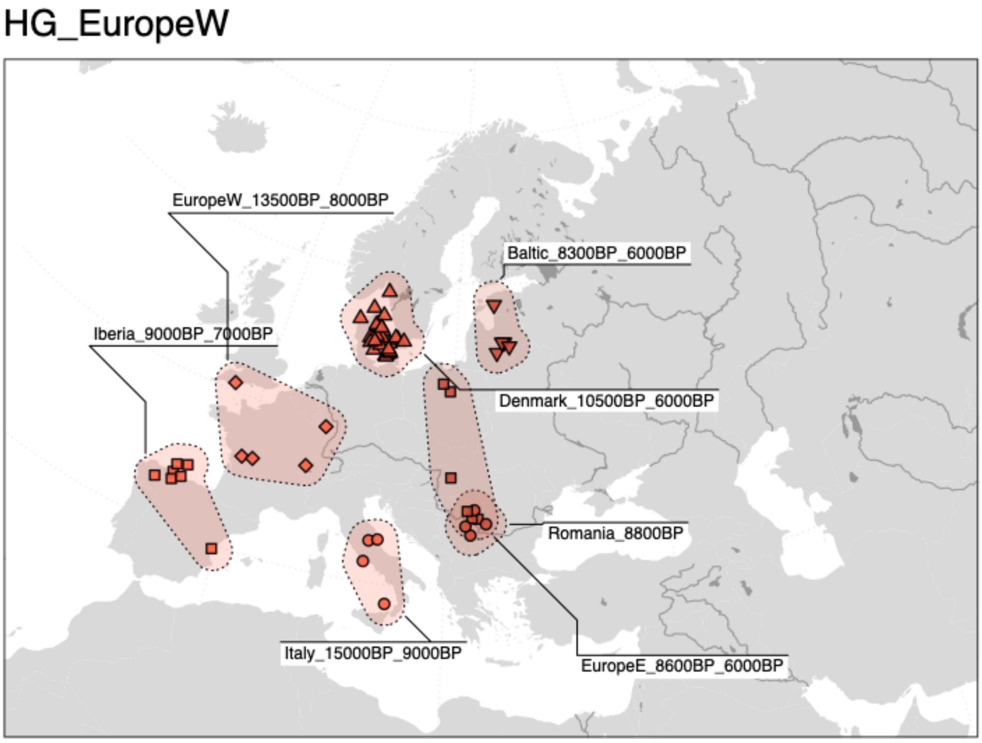


**Fig. S3f.6 Geographic distribution of individuals in cluster group *HG_EuropeW*.** Geographic locations of individuals within specific clusters are highlighted with coloured symbols, and connected with shaded hulls.

(hi-res https://www.dropbox.com/s/g5apww3nismrsxe/sfig_ibd_06_cluster_map_hgEurW.png?dl=0)

EuropeWCAsia_25000BP_300BP

This global cluster includes individuals from western Eurasia with ancestry related to Mesolithic and Neolithic Near Eastern groups. The individual genetic clusters are partitioned into a total of eight cluster groups, including all clusters of “Neolithic farmers” and “Caucasus hunter-gatherers” previously used in the literature (Fig. S3f.7-20).

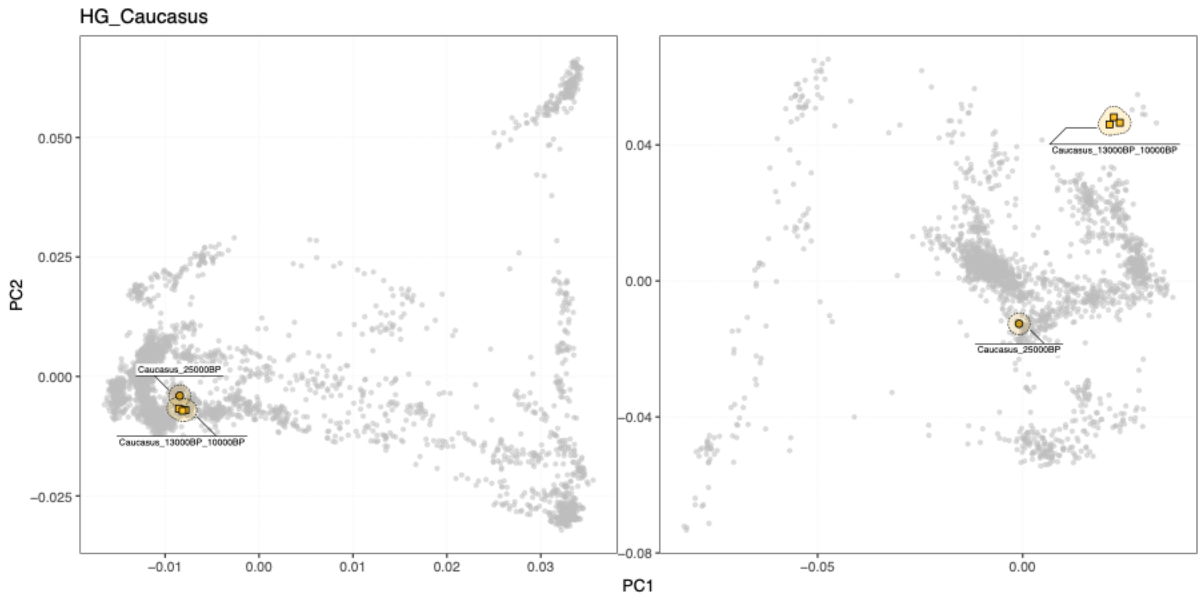


**Fig. S3f.7 PCA for cluster group *HG_Caucasus***. PCA positions of individuals within specific clusters are highlighted with coloured symbols, and connected with shaded hulls (from PCAs shown in Fig. S3d.7).

(hi-res https://www.dropbox.com/s/vqa292d3w1wrs13/sfig_ibd_07_cluster_pca_hgCauc.png?dl=0)

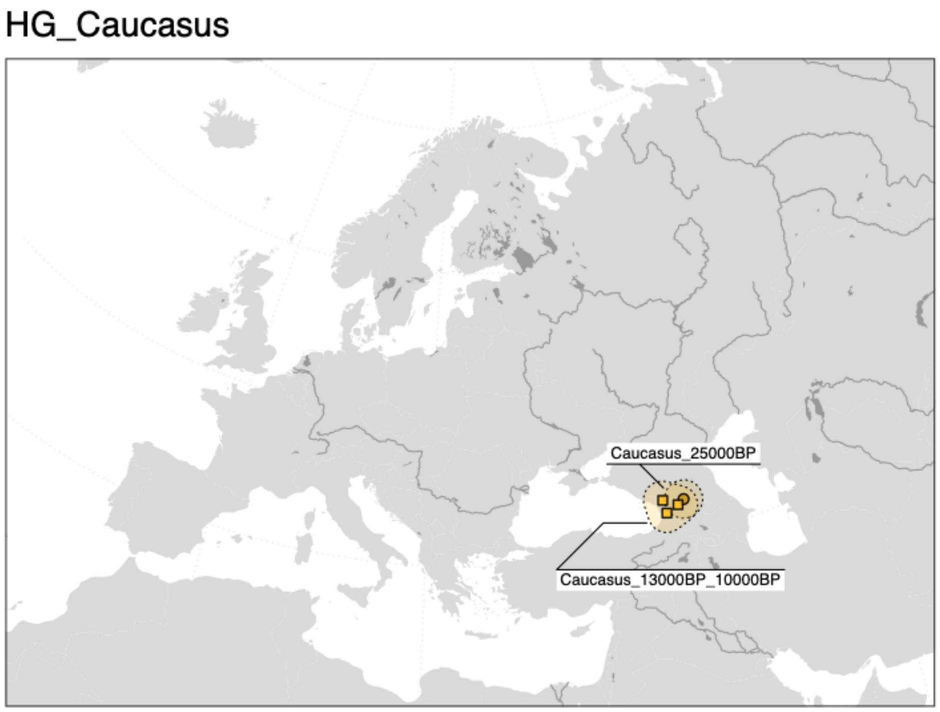


**Fig. S3f.8 Geographic distribution of individuals in cluster group *HG_Caucasus.*** Geographic locations of individuals within specific clusters are highlighted with coloured symbols, and connected with shaded hulls.

(hi-res https://www.dropbox.com/s/540e7ogjp2dyky6/sfig_ibd_08_cluster_map_hgCauc.png?dl=0)

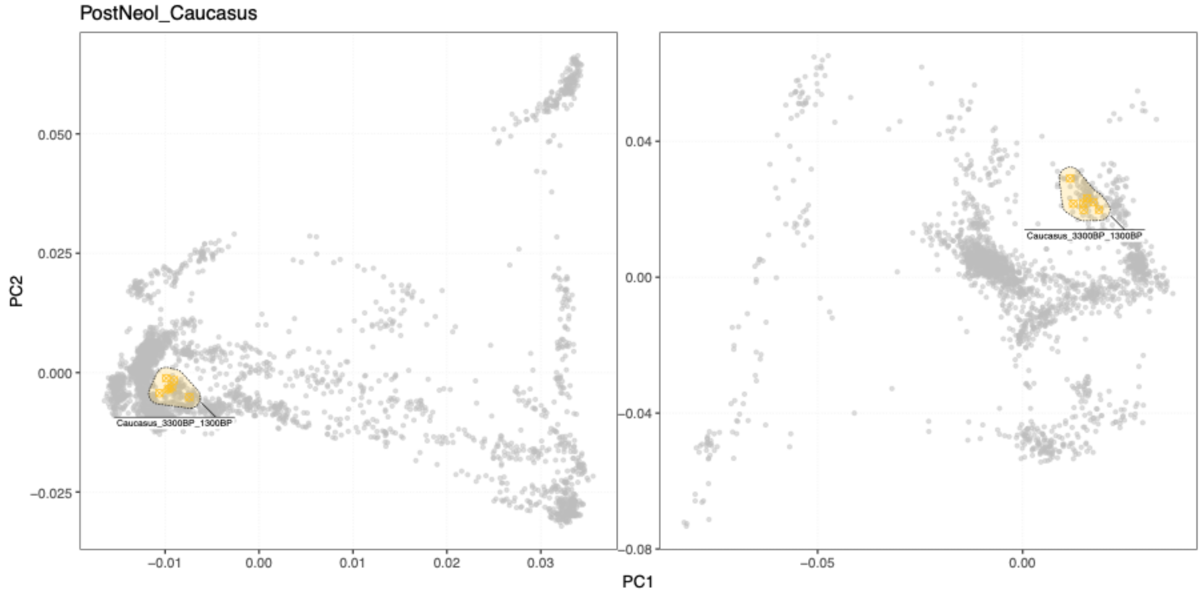


**Fig. S3f.9 PCA for cluster group *PostNeol_Caucasus***. PCA positions of individuals within specific clusters are highlighted with coloured symbols, and connected with shaded hulls (from PCAs shown in Fig. S3d.7).

(hi-res https://www.dropbox.com/s/jg0iwzglo4pkdjs/sfig_ibd_09_cluster_pca_postNeolCauc.png?dl=0)

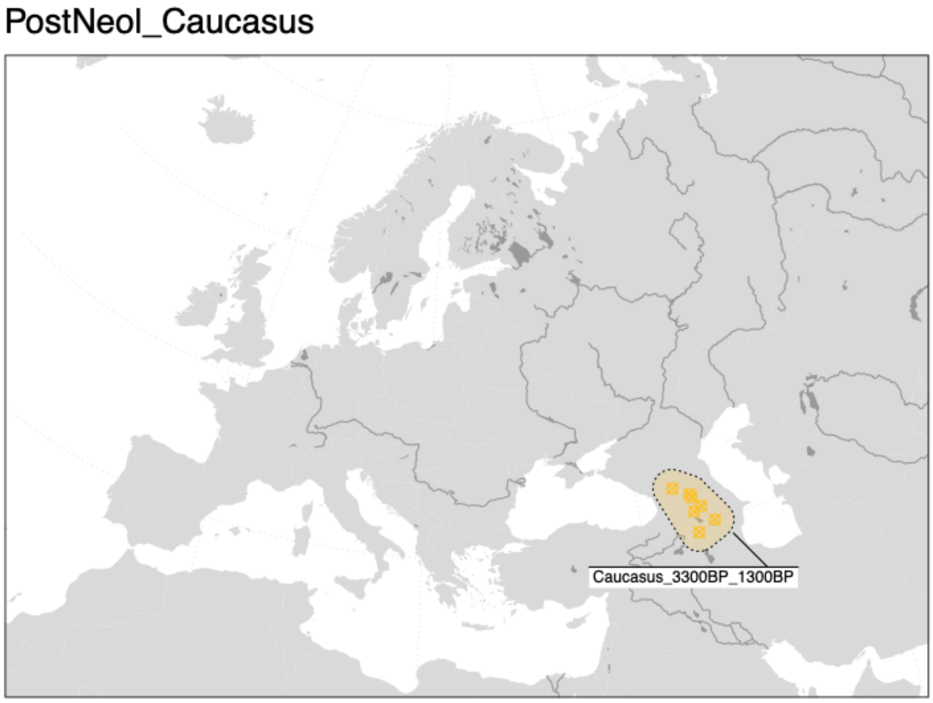

**Fig. S3f.10 Geographic distribution of individuals in cluster group *PostNeol_Caucasus.*** Geographic locations of individuals within specific clusters are highlighted with coloured symbols, and connected with shaded hulls.

(hi-res https://www.dropbox.com/s/9ikx5eci8lz8yg4/sfig_ibd_10_cluster_map_postNeolCauc.png?dl=0)

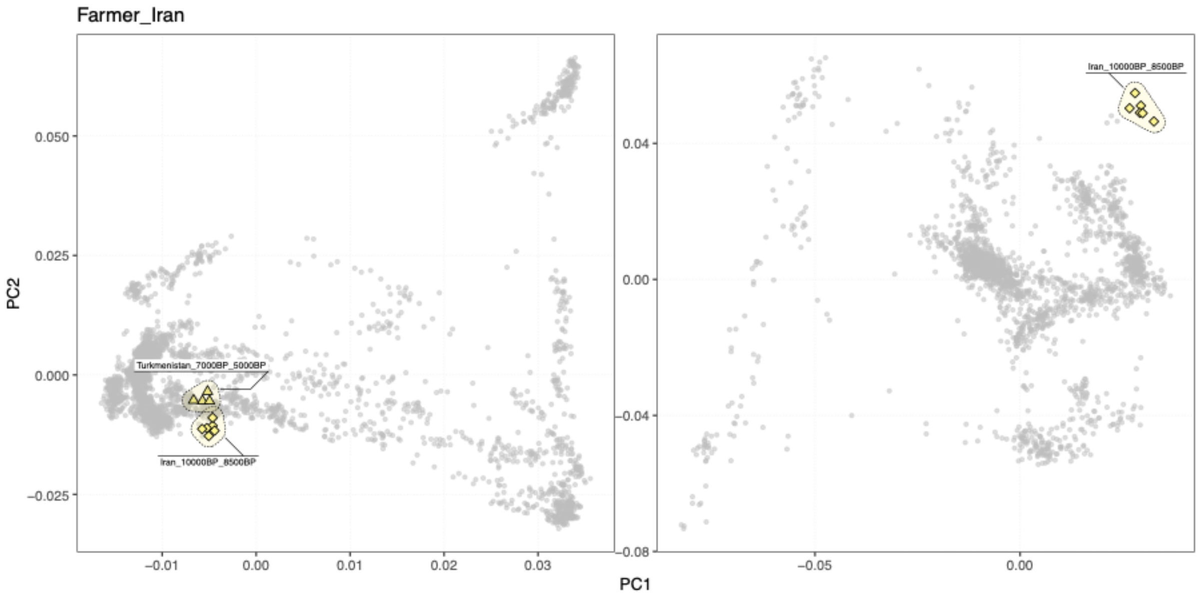


**Fig. S3f.11 PCA for cluster group *Farmer_Iran*.** PCA positions of individuals within specific clusters are highlighted with coloured symbols, and connected with shaded hulls (from PCAs shown in Fig. S3d.7).

(hi-res https://www.dropbox.com/s/koq4nktk63uua10/sfig_ibd_11_cluster_pca_farmerIran.png?dl=0)

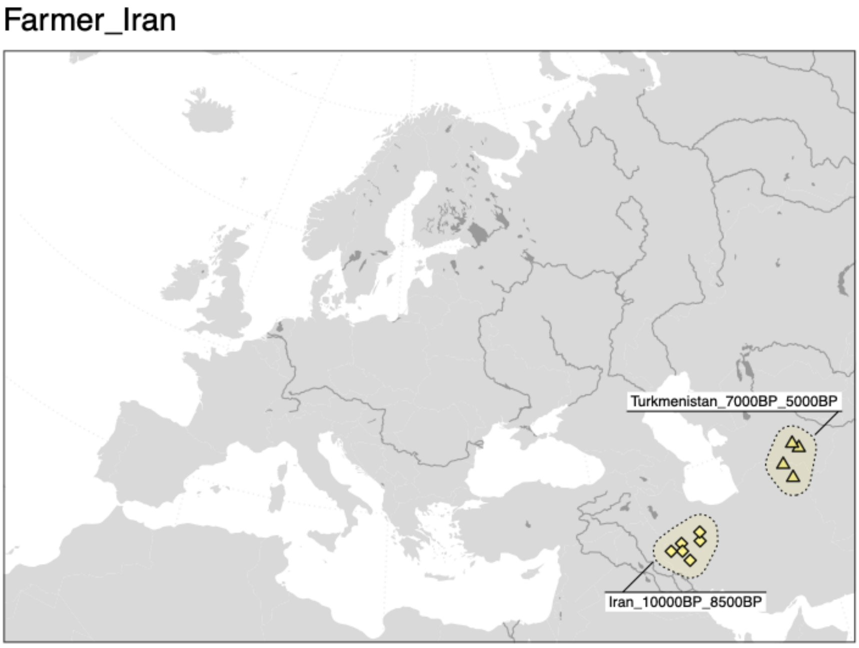

**Fig. S3f.12 Geographic distribution of individuals in cluster group *Farmer_Iran.*** Geographic locations of individuals within specific clusters are highlighted with coloured symbols, and connected with shaded hulls.

(hi-res https://www.dropbox.com/s/0frbab3mhaec2kr/sfig_ibd_12_cluster_map_farmerIran.png?dl=0)

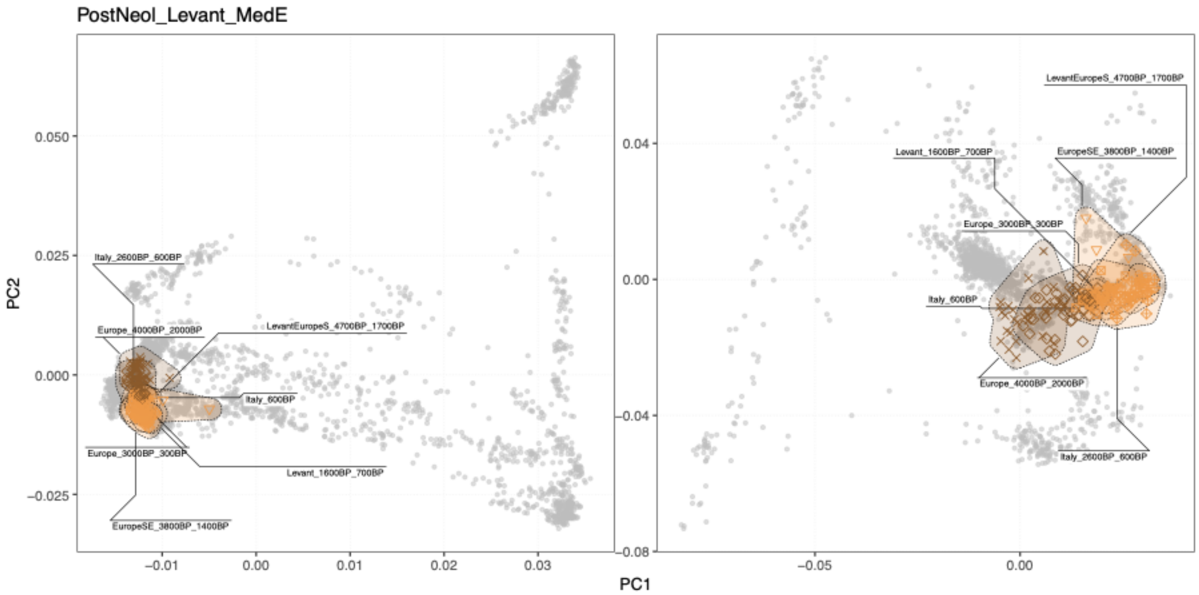


**Fig. S3f.13 PCA for cluster group *PostNeol_Levant_MedE*.** PCA positions of individuals within specific clusters are highlighted with coloured symbols, and connected with shaded hulls (from PCAs shown in Fig. S3d.7).

(hi-res https://www.dropbox.com/s/g213qz6qik5cxfd/sfig_ibd_13_cluster_pca_postNeolLevant.png?dl=0)

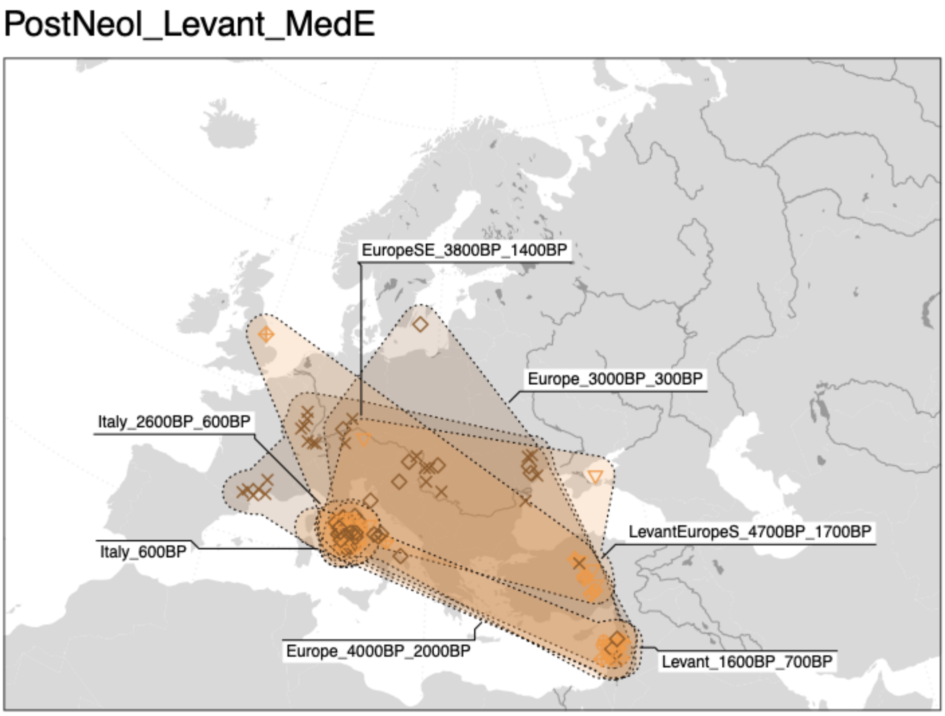


**Fig. S3f.14 Geographic distribution of individuals in cluster group *PostNeol_Levant_MedE.*** Geographic locations of individuals within specific clusters are highlighted with coloured symbols, and connected with shaded hulls.

(hi-res https://www.dropbox.com/s/f1723bwerxthive/sfig_ibd_14_cluster_map_postNeolLevant.png?dl=0)

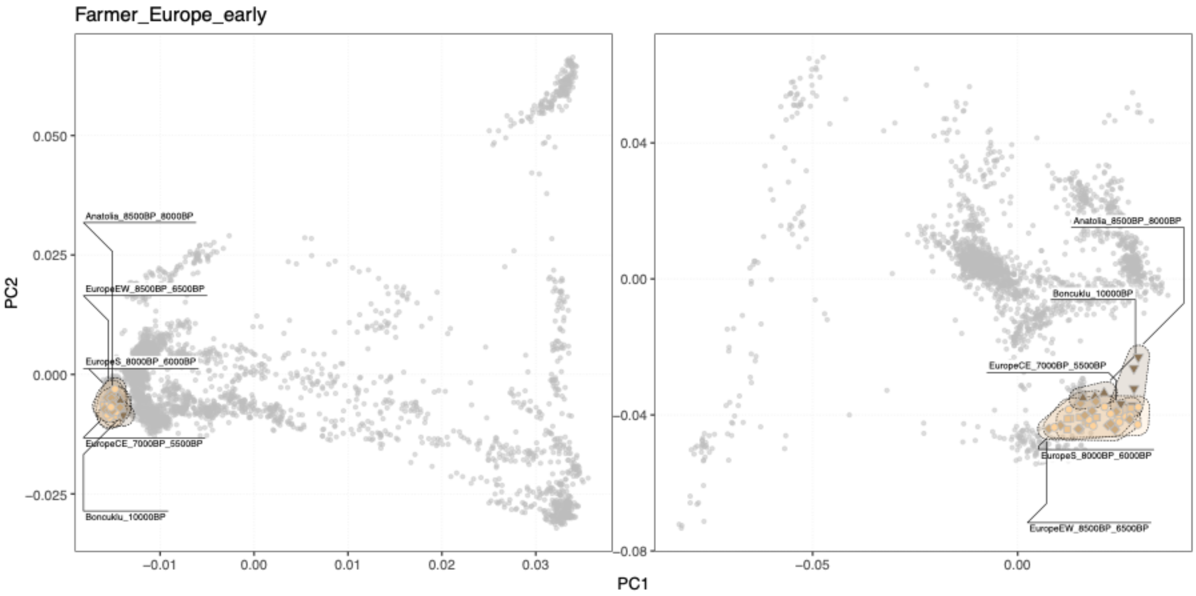


**Fig. S3f.15 PCA for cluster group *Farmer_Europe_early*.** PCA positions of individuals within specific clusters are highlighted with coloured symbols, and connected with shaded hulls (from PCAs shown in Fig. S3d.7).

(hi-res https://www.dropbox.com/s/2uk4xb7bfq1rjw3/sfig_ibd_15_cluster_pca_farmerEurEarly.png?dl=0)

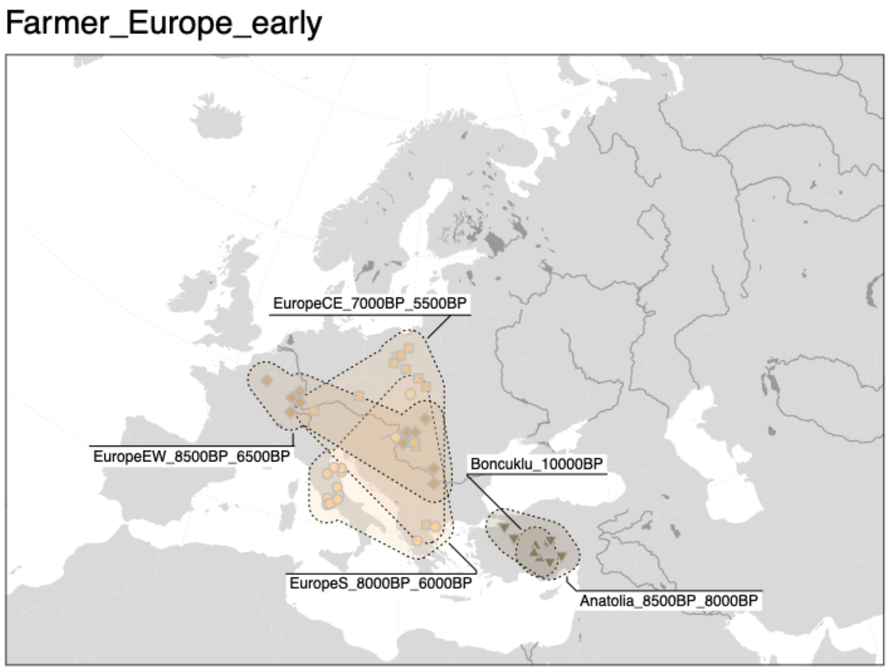


**Fig. S3f.16 Geographic distribution of individuals in cluster group *Farmer_Europe_early.*** Geographic locations of individuals within specific clusters are highlighted with coloured symbols, and connected with shaded hulls.

(hi-res https://www.dropbox.com/s/q5ueuux2nmak9p1/sfig_ibd_16_cluster_map_farmerEurEarly.png?dl=0)

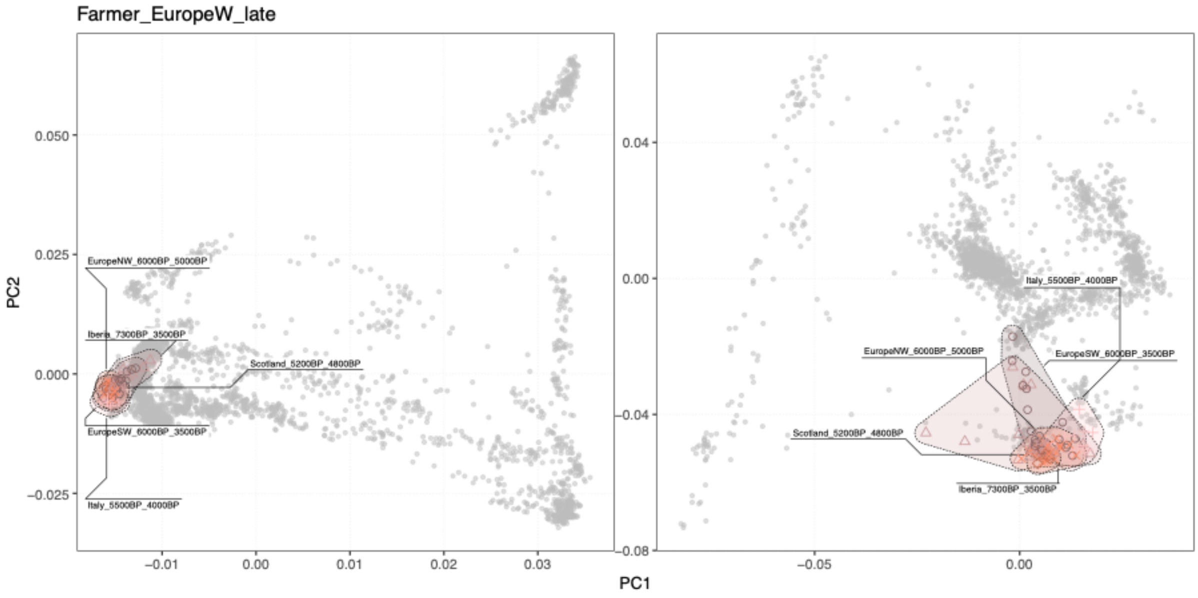


**Fig. S3f.17 PCA for cluster group *Farmer_EuropeW_late*.** PCA positions of individuals within specific clusters are highlighted with coloured symbols, and connected with shaded hulls (from PCAs shown in Fig. S3d.7).

(hi-res https://www.dropbox.com/s/6obhskwydytauqy/sfig_ibd_17_cluster_pca_farmerEurWLate.png?dl=0)

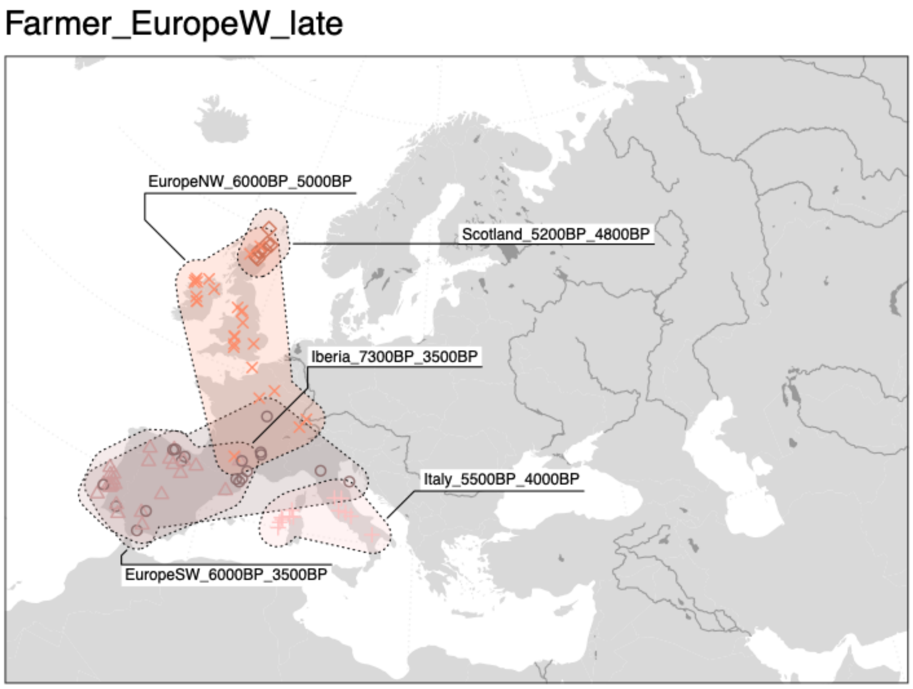


**Fig. S3f.18 Geographic distribution of individuals in cluster group *Farmer_EuropeW_late.*** Geographic locations of individuals within specific clusters are highlighted with coloured symbols, and connected with shaded hulls.

(hi-res https://www.dropbox.com/s/hcdlu7wka0eum0w/sfig_ibd_18_cluster_map_farmerEurWLate.png?dl=0)

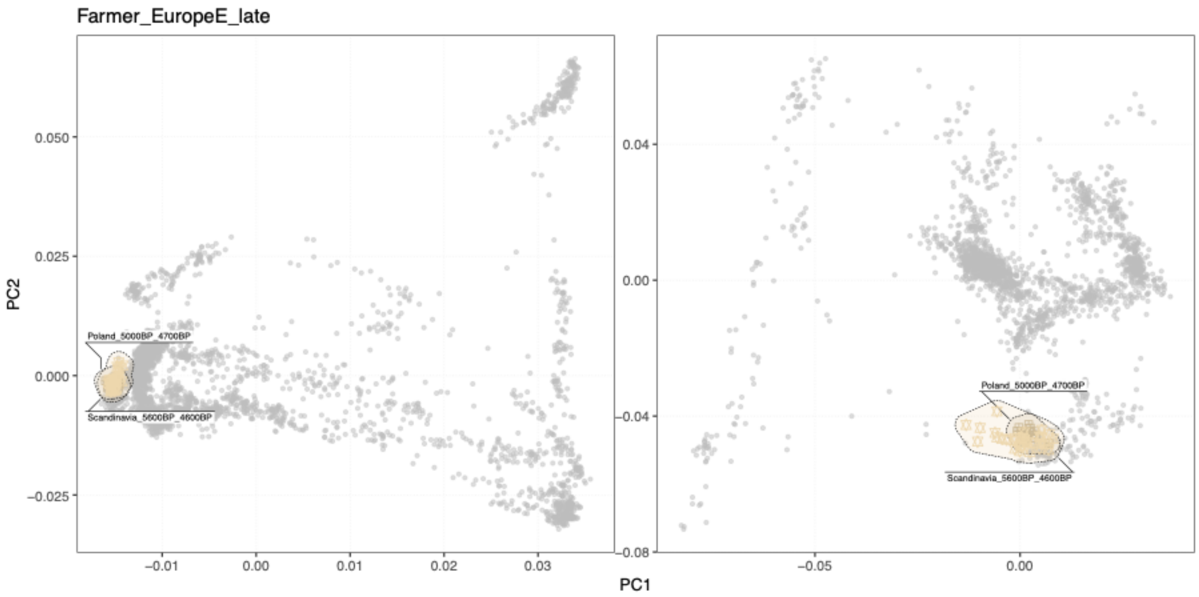


**Fig. S3f.19 PCA for cluster group *Farmer_EuropeE_late***. PCA positions of individuals within specific clusters are highlighted with coloured symbols, and connected with shaded hulls (from PCAs shown in Fig. S3d.7).

(hi-res https://www.dropbox.com/s/lcyek0dnhpzkmw4/sfig_ibd_19_cluster_pca_farmerEurELate.png?dl=0)

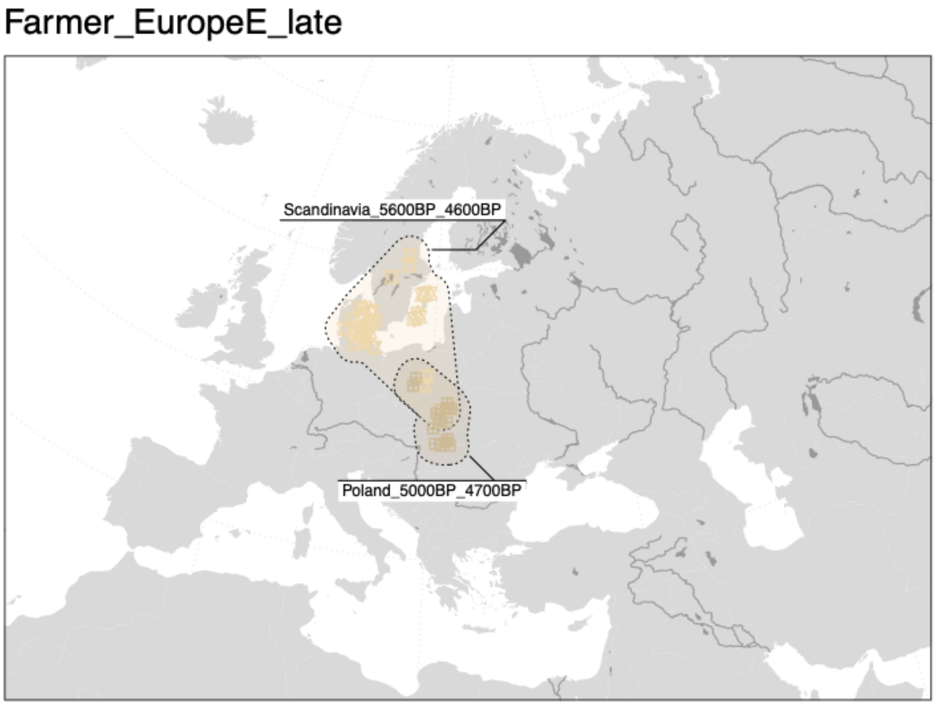


**Fig. S3f.20 Geographic distribution of individuals in cluster group *Farmer_EuropeE_late.*** Geographic locations of individuals within specific clusters are highlighted with coloured symbols, and connected with shaded hulls.
(hi-res https://www.dropbox.com/s/79r3s7lg4zz32gk/sfig_ibd_20_cluster_map_farmerEurELate.png?dl=0)

Eurasia_5000BP_200BP

This global cluster includes individuals from western Eurasia from the Late Neolithic and early Bronze Age onwards. The individual genetic clusters are partitioned into a total of five cluster groups, including all clusters of individuals with "Steppe ancestry" related to Bronze Age Steppe pastoralists previously used in the literature (Fig. S3f.21-28).

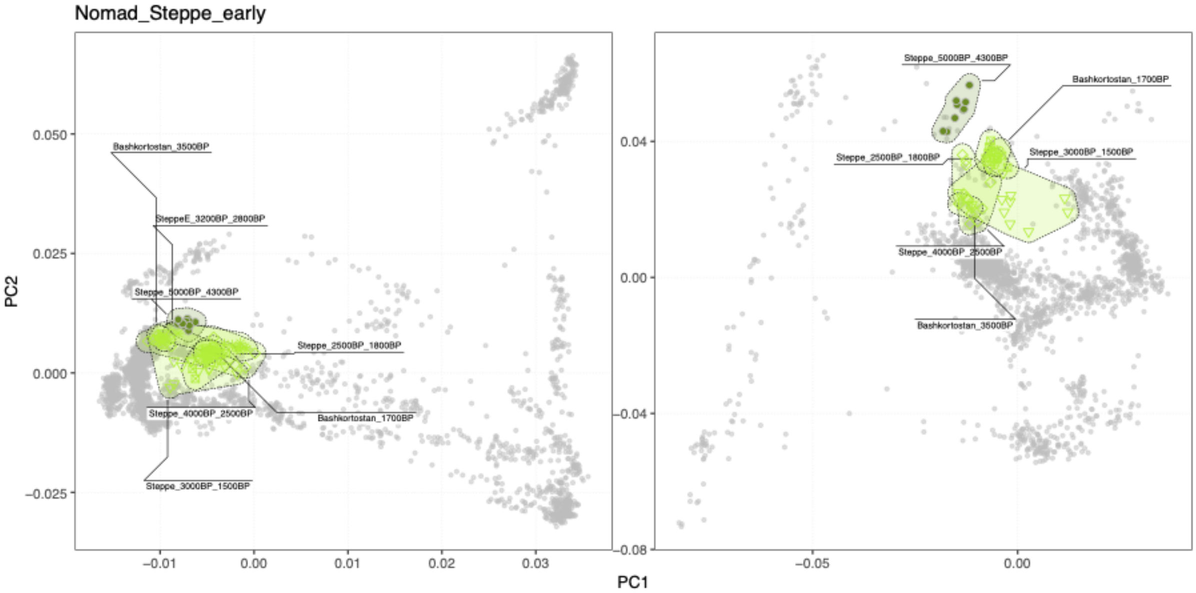


**Fig. S3f.21 PCA for cluster group *Nomad_Steppe_early***. PCA positions of individuals within specific clusters are highlighted with coloured symbols, and connected with shaded hulls (from PCAs shown in Fig. S3d.7).

(hi-res https://www.dropbox.com/s/agh89vha7fis685/sfig_ibd_21_cluster_pca_NomadEarly.png?dl=0)

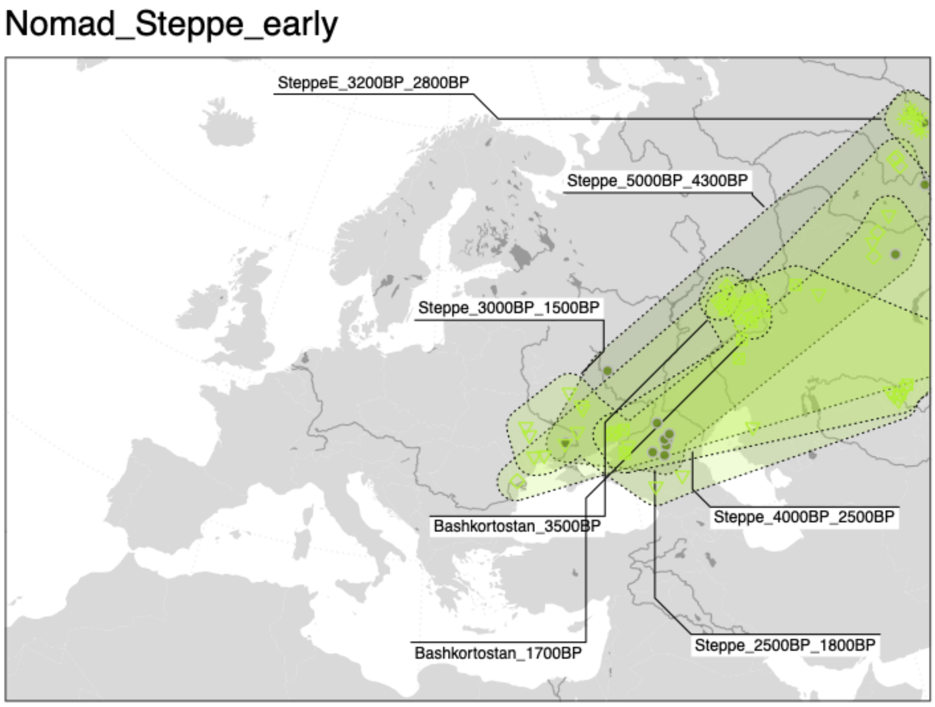

**Fig. S3f.22 Geographic distribution of individuals in cluster group *Nomad_Steppe_early.*** Geographic locations of individuals within specific clusters are highlighted with coloured symbols, and connected with shaded hulls.

(hi-res https://www.dropbox.com/s/wcjeq0sxsslqi6h/sfig_ibd_22_cluster_map_NomadEarly.png?dl=0)

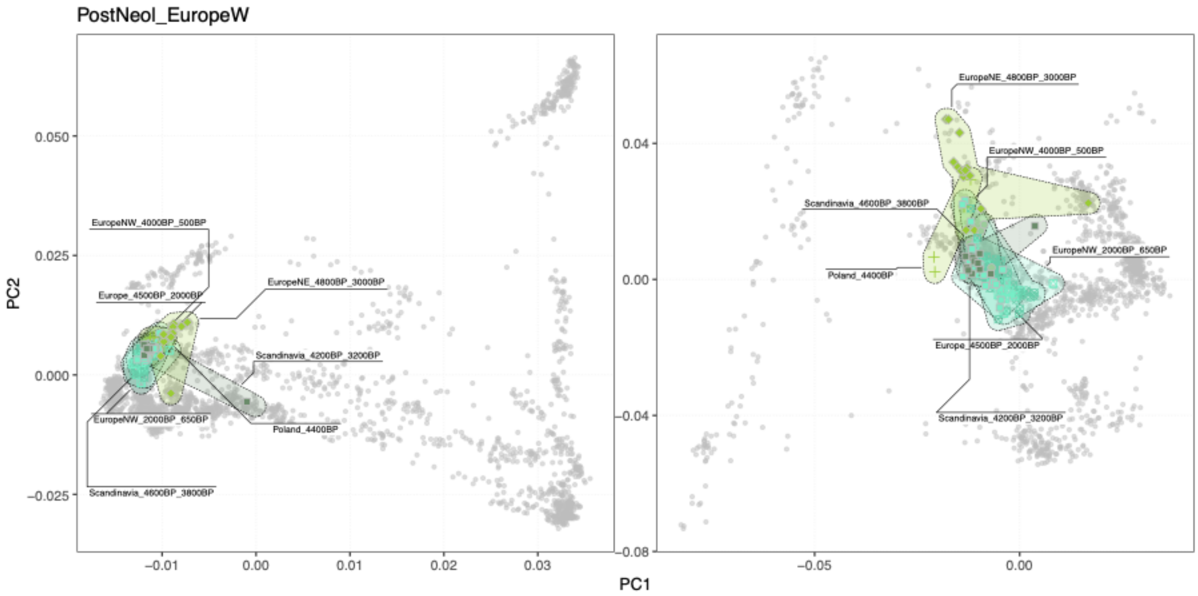


**Fig. S3f.23 PCA for cluster group *PostNeol_EuropeW*.** PCA positions of individuals within specific clusters are highlighted with coloured symbols, and connected with shaded hulls (from PCAs shown in Fig. S3d.7).

(hi-res https://www.dropbox.com/s/3u7el8u48s7fag5/sfig_ibd_23_cluster_pca_postNeolEurW.png?dl=0)

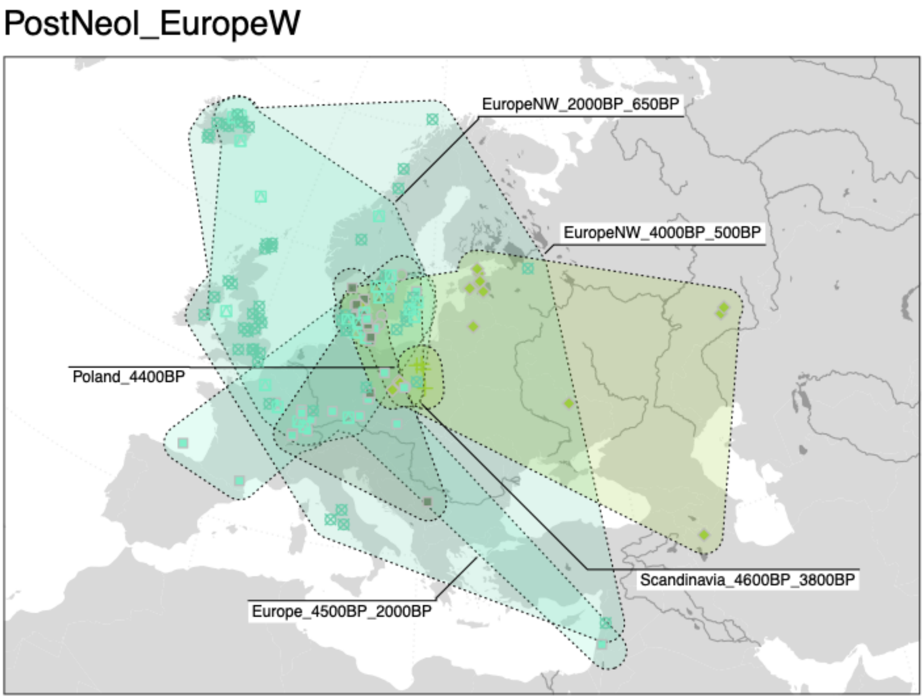


**Fig. S3f.24 Geographic distribution of individuals in cluster group *PostNeol_EuropeW.*** Geographic locations of individuals within specific clusters are highlighted with coloured symbols, and connected with shaded hull.

(hi-res https://www.dropbox.com/s/39gdicoj0lr2xrk/sfig_ibd_24_cluster_map_postNeolEurW.png?dl=0)

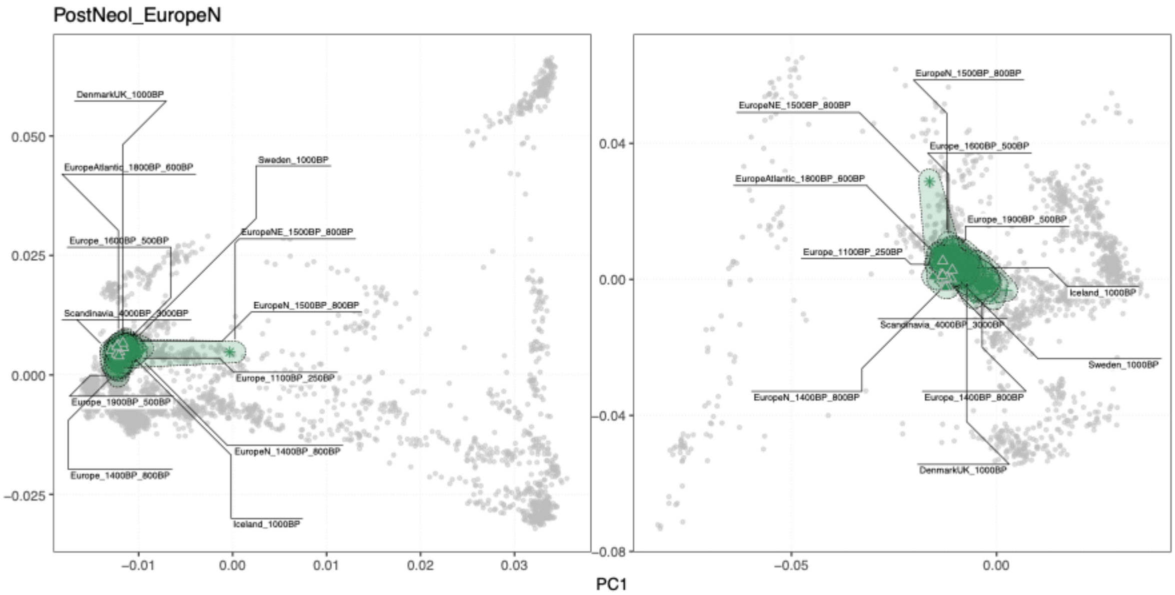


**Fig. S3f.25 PCA for cluster group *PostNeol_EuropeN.*** PCA positions of individuals within specific clusters are highlighted with coloured symbols, and connected with shaded hulls (from PCAs shown in Fig. S3d.7).

(hi-res https://www.dropbox.com/s/fikzciloupbcl9r/sfig_ibd_25_cluster_pca_postNeolEurN.png?dl=0)

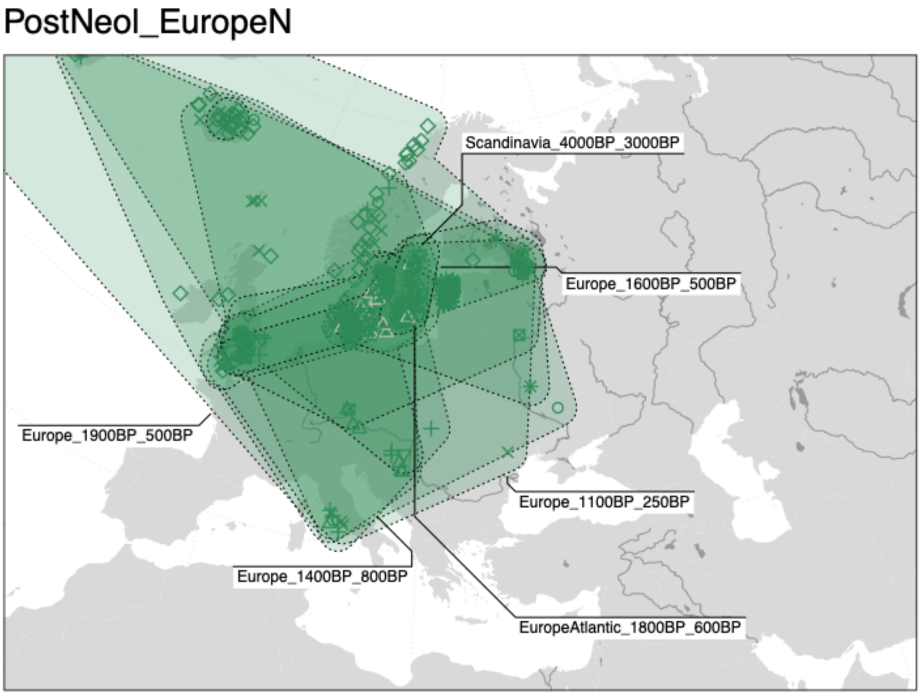


**Fig. S3f.26 Geographic distribution of individuals in cluster group *PostNeol_EuropeN.*** Geographic locations of individuals within specific clusters are highlighted with coloured symbols, and connected with shaded hulls.

(hi-res https://www.dropbox.com/s/5ss7sa6rfof0ljr/sfig_ibd_26_cluster_map_postNeolEurN.png?dl=0)

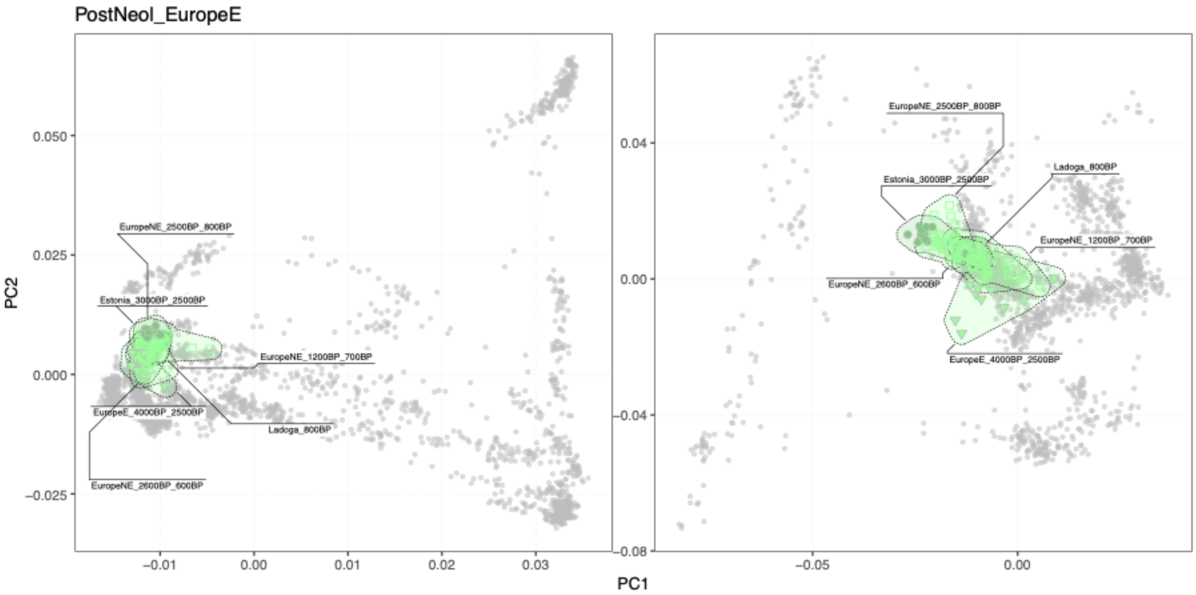

**Fig. S3f.27 PCA for cluster group *PostNeol_EuropeE*.** PCA positions of individuals within specific clusters are highlighted with coloured symbols, and connected with shaded hulls (from PCAs shown in Fig. S3d.7).

(hi-res https://www.dropbox.com/s/pg13ea15u7caccb/sfig_ibd_27_cluster_pca_postNeolEurE.png?dl=0)

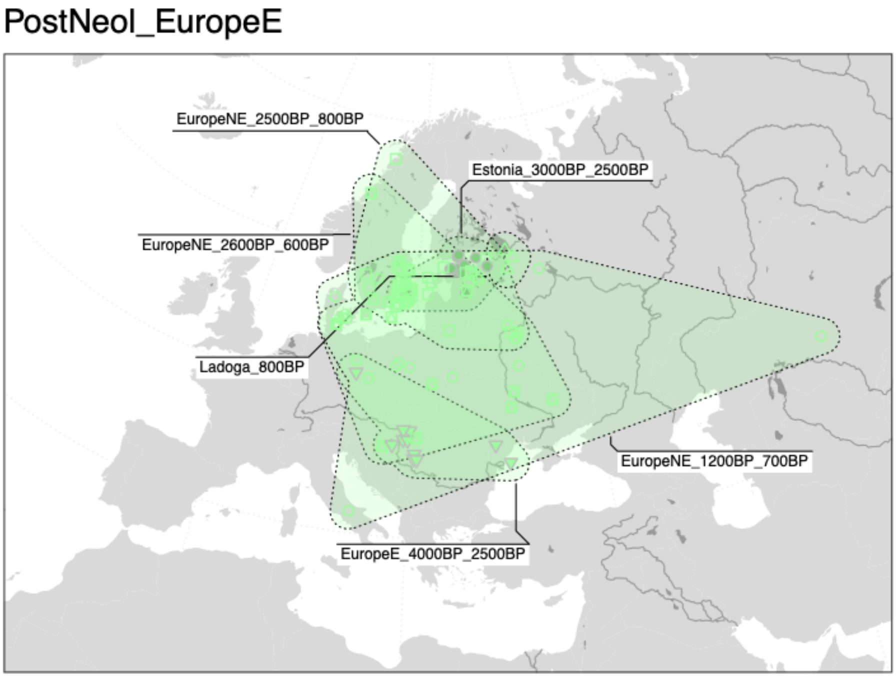


**Fig. S3f.28 Geographic distribution of individuals in cluster group *PostNeol_EuropeE.*** Geographic locations of individuals within specific clusters are highlighted with coloured symbols, and connected with shaded hulls.

(hi-res https://www.dropbox.com/s/3o696sxcv0zkzpm/sfig_ibd_28_cluster_map_postNeolEurE.png?dl=0)

Asia_45000BP_200BP

This global cluster includes diverse sets of clusters of individuals from the Southeast-, East- and North Asia, broadly characterised by “east Eurasian” ancestry. The individual genetic clusters are partitioned into a total of 12 cluster groups (Fig. S3f.29-44).

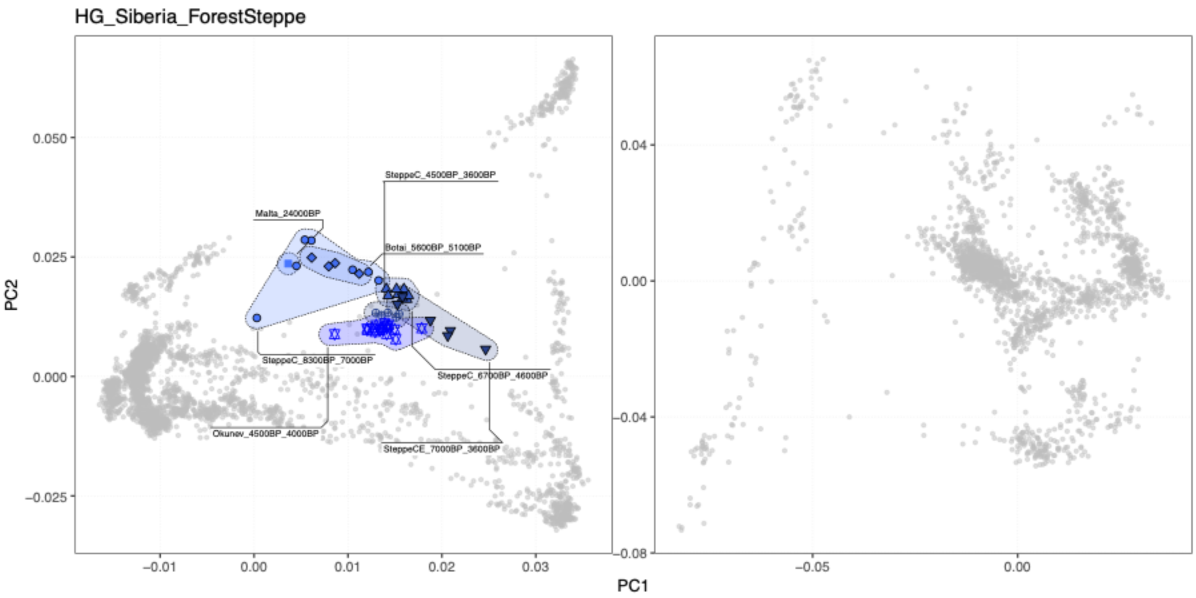


**Fig. S3f.29 PCA for cluster group *HG_Siberia_ForestSteppe***. PCA positions of individuals within specific clusters are highlighted with coloured symbols, and connected with shaded hulls (from PCAs shown in Fig. S3d.7).

(hi-res https://www.dropbox.com/s/57nxlvvuzbgw381/sfig_ibd_29_cluster_pca_hgSibForestSteppe.png?dl=0)

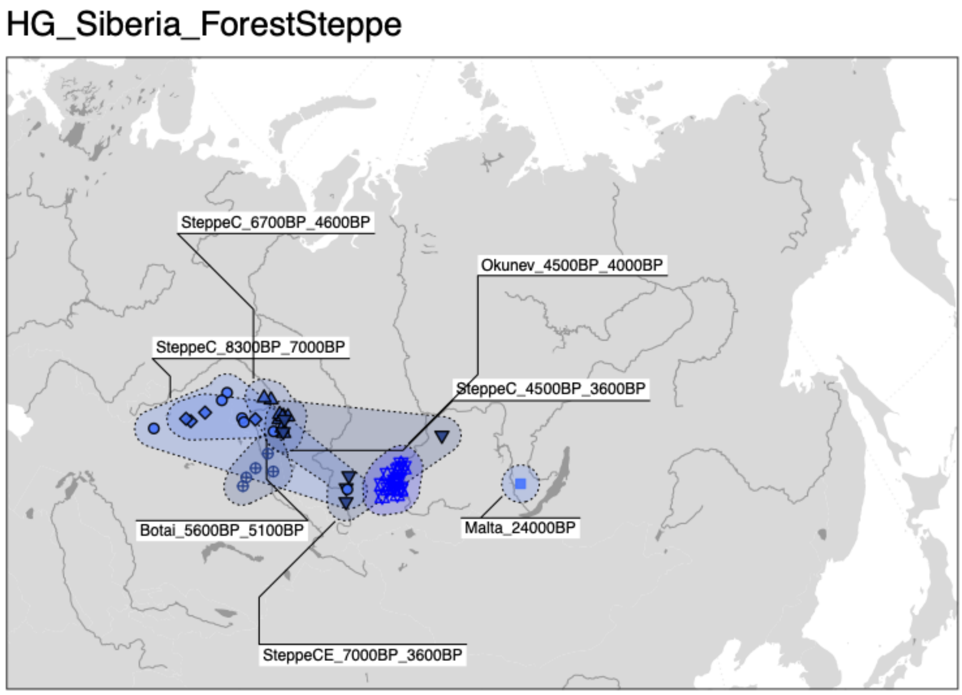


**Fig. S3f.30 Geographic distribution of individuals in cluster group *HG_Siberia_ForestSteppe.*** Geographic locations of individuals within specific clusters are highlighted with coloured symbols, and connected with shaded hulls.

(hi-res https://www.dropbox.com/s/l4332v1vxa2z4dd/sfig_ibd_30_cluster_map_hgSibForestSteppe.png?dl=0)

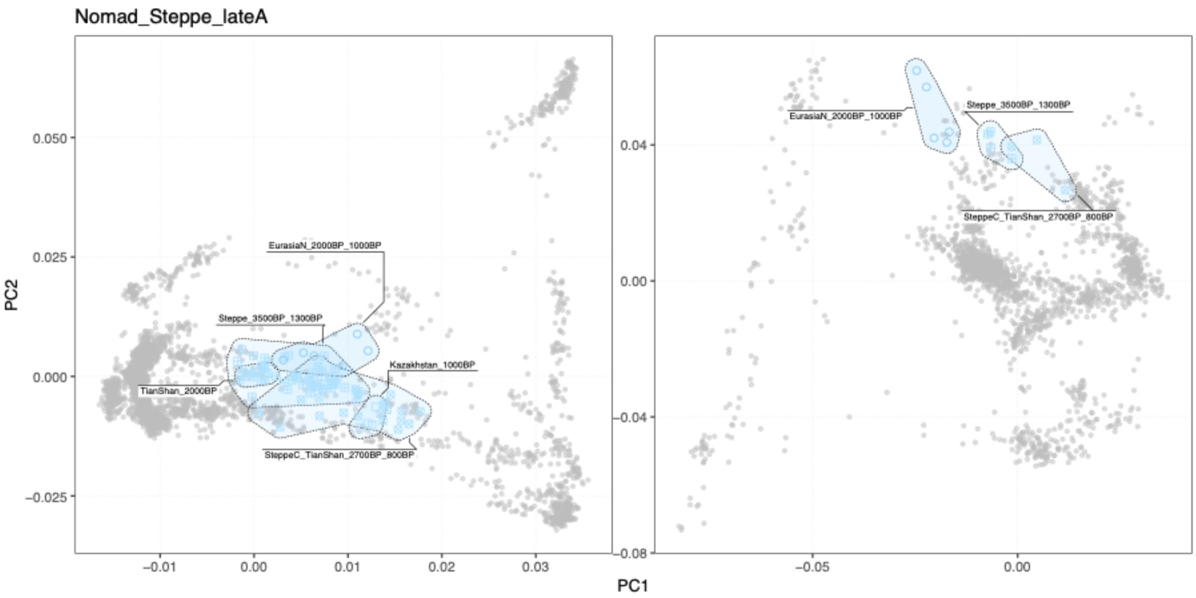


**Fig. S3f.31 PCA for cluster group *Nomad_Steppe_lateA*.** PCA positions of individuals within specific clusters are highlighted with coloured symbols, and connected with shaded hulls (from PCAs shown in Fig. S3d.7).

(hi-res https://www.dropbox.com/s/3ignx2oah5vi60j/sfig_ibd_31_cluster_pca_NomadLateA.png?dl=0)

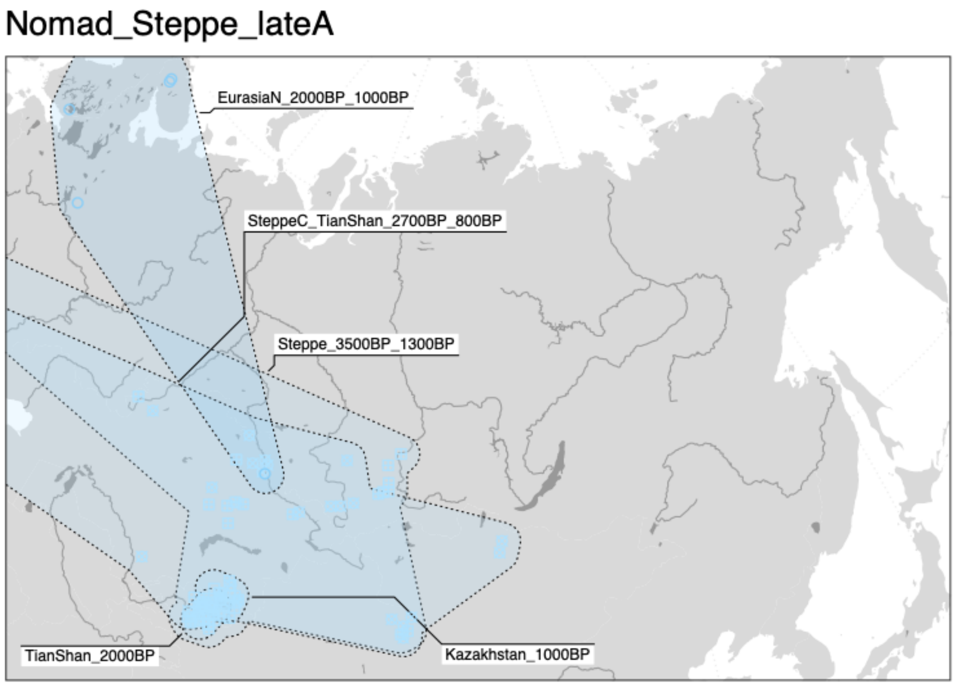


**Fig. S3f.32 Geographic distribution of individuals in cluster group *Nomad_Steppe_lateA.*** Geographic locations of individuals within specific clusters are highlighted with coloured symbols, and connected with shaded hulls.

(hi-res https://www.dropbox.com/s/homoq40xk1d630u/sfig_ibd_32_cluster_map_NomadLateA.png?dl=0)

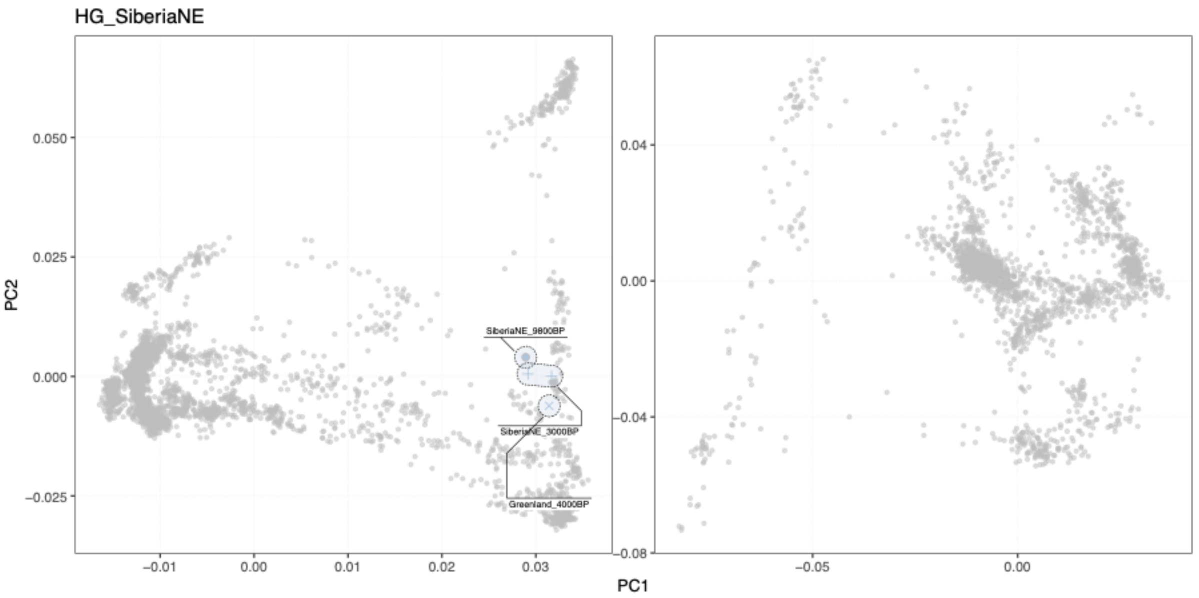


**Fig. S3f.33 PCA for cluster group *HG_SiberiaNE***. PCA positions of individuals within specific clusters are highlighted with coloured symbols, and connected with shaded hulls (from PCAs shown in Fig. S3d.7).

(hi-res https://www.dropbox.com/s/ggpvvldymk7f0g2/sfig_ibd_33_cluster_pca_hgSibNE.png?dl=0)

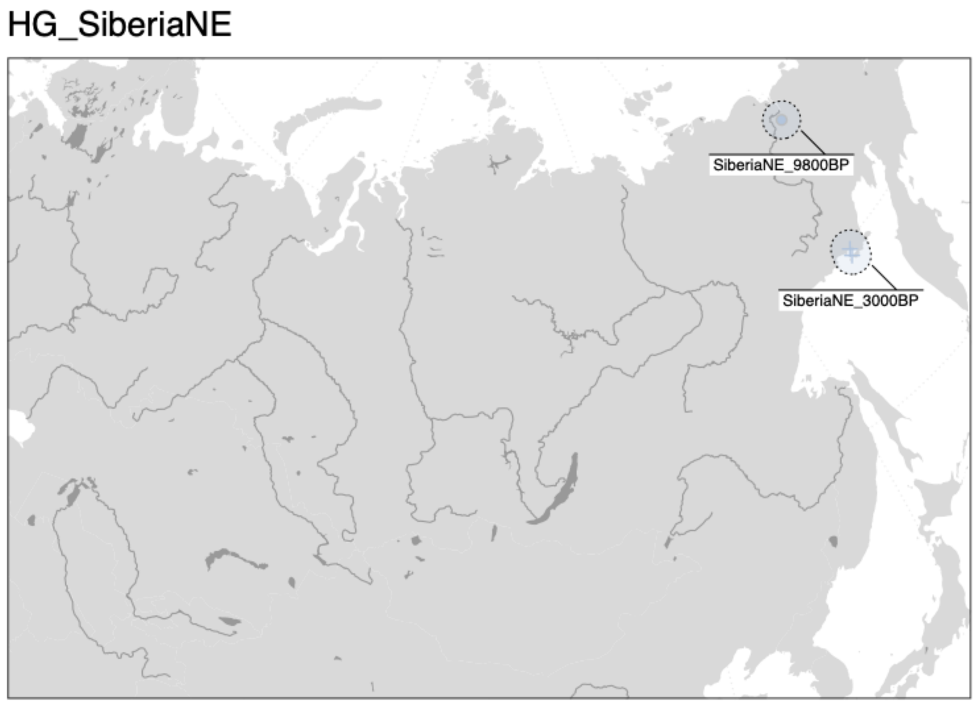


**Fig. S3f.34 Geographic distribution of individuals in cluster group *HG_SiberiaNE.*** Geographic locations of individuals within specific clusters are highlighted with coloured symbols, and connected with shaded hulls.

(hi-res https://www.dropbox.com/s/7o45et220dshbhy/sfig_ibd_34_cluster_map_hgSibNE.png?dl=0)

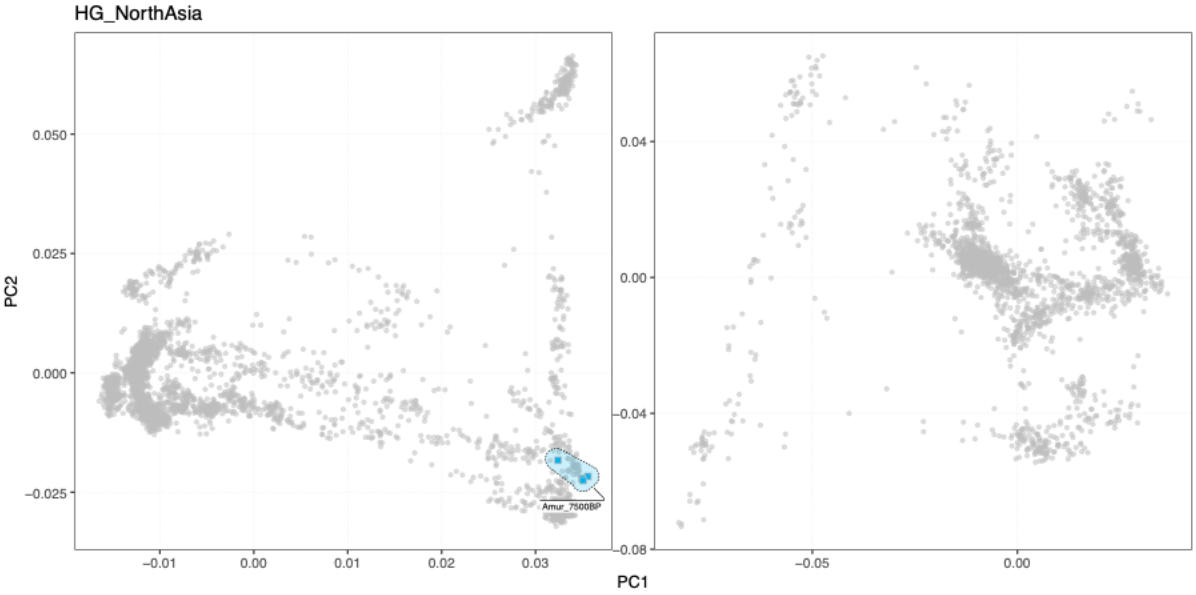


**Fig. S3f.35 PCA for cluster group *HG_NorthAsia***. PCA positions of individuals within specific clusters are highlighted with coloured symbols, and connected with shaded hulls (from PCAs shown in Fig. S3d.7).

(hi-res https://www.dropbox.com/s/k5psams72jz1f5z/sfig_ibd_35_cluster_pca_hgAsiaN.png?dl=0)

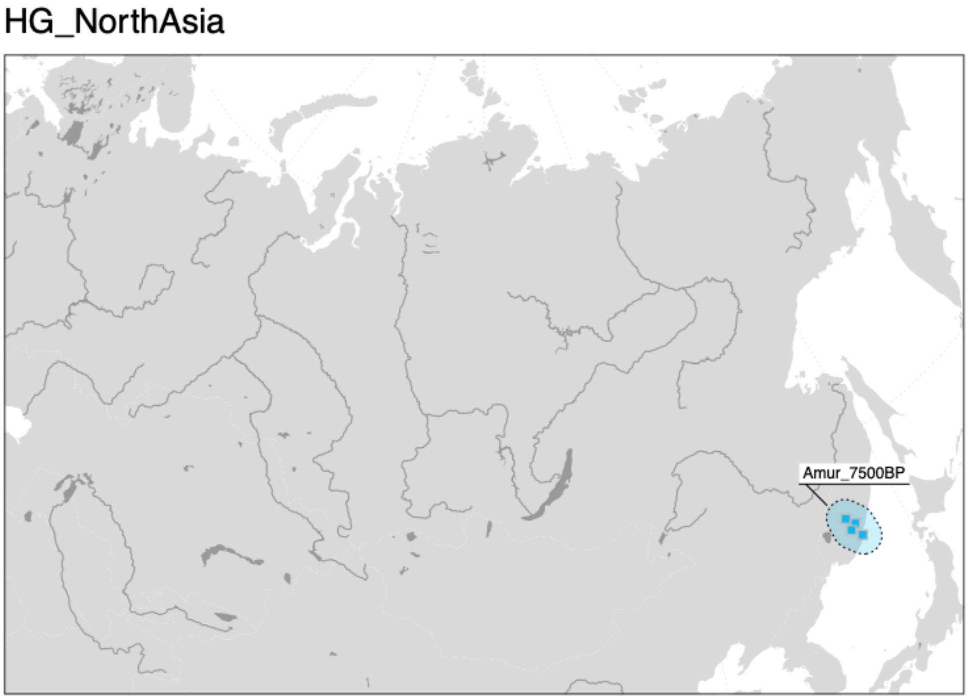

**Fig. S3f.36 Geographic distribution of individuals in cluster group *HG_NorthAsia.*** Geographic locations of individuals within specific clusters are highlighted with coloured symbols, and connected with shaded hulls.

(hi-res [https://www.dropbox.com/s/65nrdlz8yi4hbpl/sfig_ibd_36_cluster_map_hgAsiaN.png?dl=0](https://www.dropbox.com/s/65nrdlz8yi4hbpl/sfig_ibd_36_cluster_map_hgAsiaN.png?dl=0))

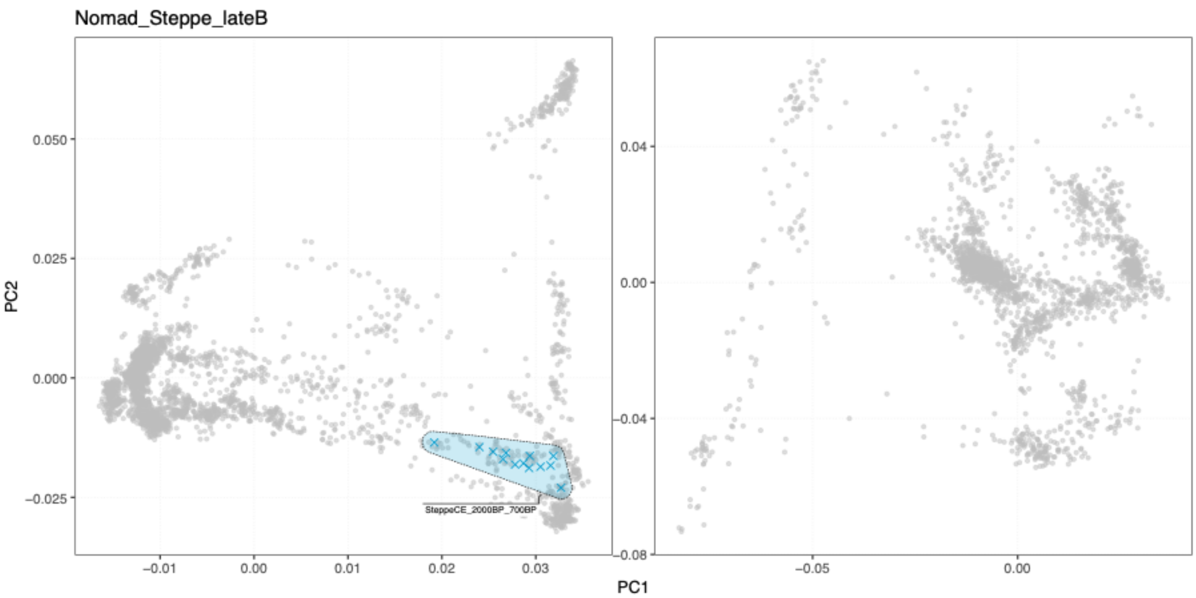


**Fig. S3f.37 PCA for cluster group *Nomad_Steppe_lateB.*** PCA positions of individuals within specific clusters are highlighted with coloured symbols, and connected with shaded hulls (from PCAs shown in Fig. S3d.7).

(hi-res [https://www.dropbox.com/s/bi8nxp89c4v17il/sfig_ibd_37_cluster_pca_NomadLateB.png?dl=0](https://www.dropbox.com/s/bi8nxp89c4v17il/sfig_ibd_37_cluster_pca_NomadLateB.png?dl=0))

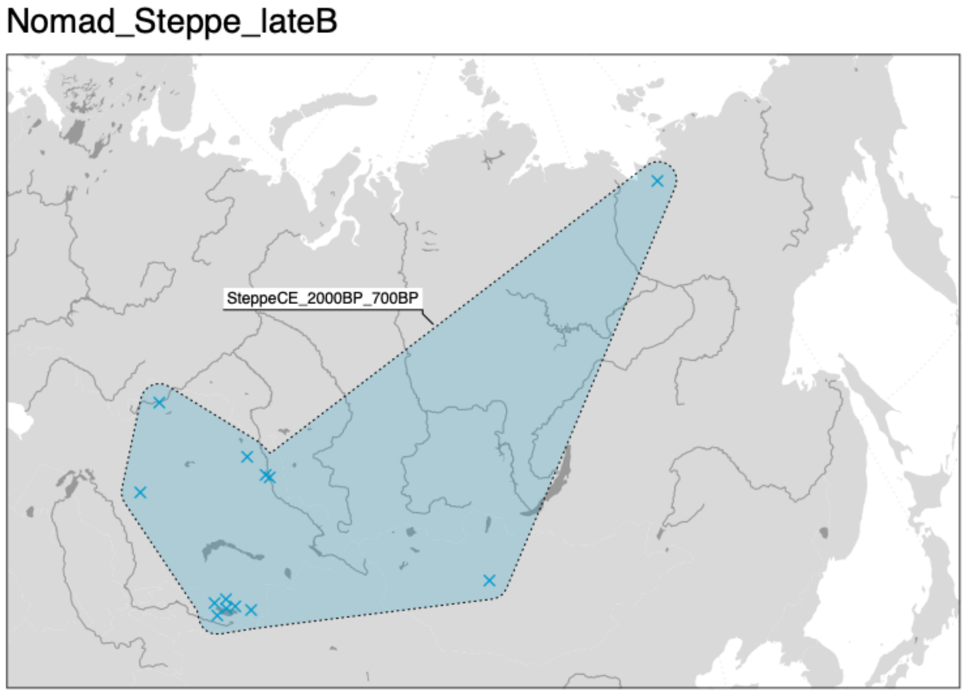

**Fig. S3f.38 Geographic distribution of individuals in cluster group *Nomad_Steppe_lateB.*** Geographic locations of individuals within specific clusters are highlighted with coloured symbols, and connected with shaded hulls.

(hi-res https://www.dropbox.com/s/vvyvnlcyk18d7z5/sfig_ibd_38_cluster_map_NomadLateB.png?dl=0)

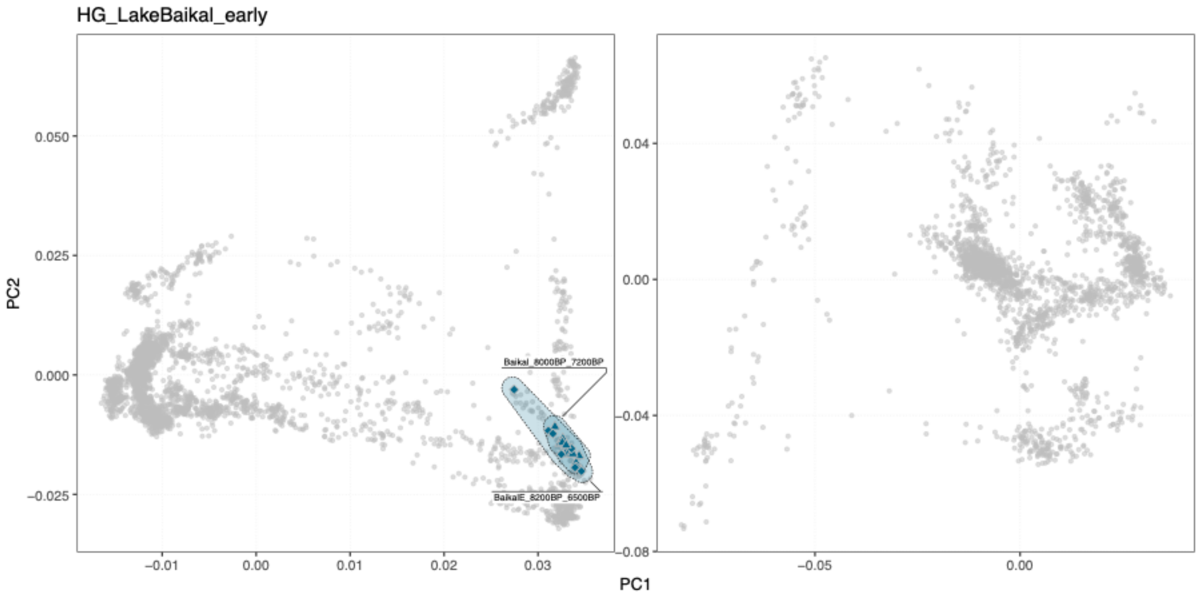


**Fig. S3f.39 PCA for cluster group *HG_LakeBaikal_early*.** PCA positions of individuals within specific clusters are highlighted with coloured symbols, and connected with shaded hulls (from PCAs shown in Fig. S3d.7).

(hi-res https://www.dropbox.com/s/gyo2p4dwd8guq4s/sfig_ibd_39_cluster_pca_hgBaikalEarly.png?dl=0)

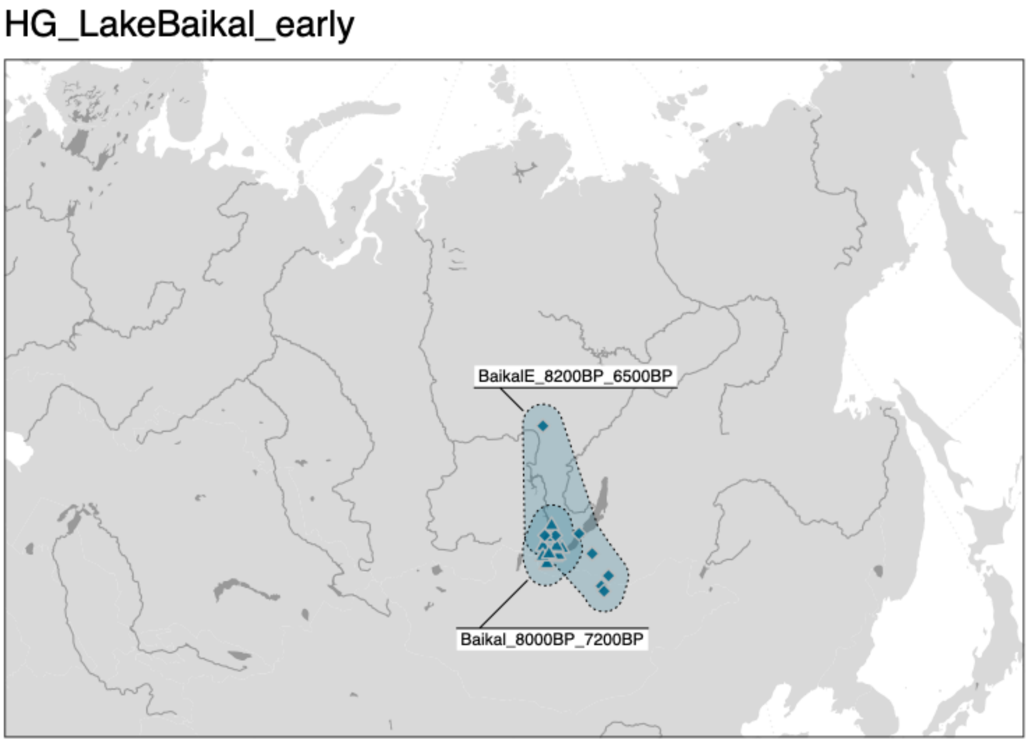

**Fig. S3f.40 Geographic distribution of individuals in cluster group *HG_LakeBaikal_early.*** Geographic locations of individuals within specific clusters are highlighted with coloured symbols, and connected with shaded hulls.

(hi-res https://www.dropbox.com/s/e83ng1wjmu5sbzt/sfig_ibd_40_cluster_map_hgBaikalEarly.png?dl=0)

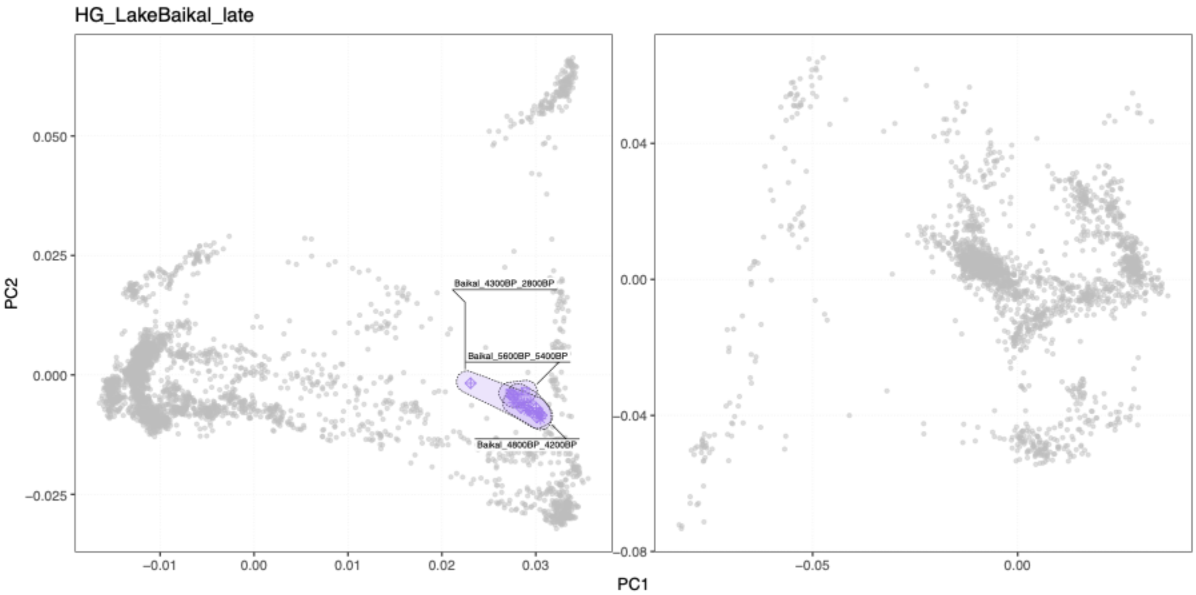


**Fig. S3f.41 PCA for cluster group *HG_LakeBaikal_late*.** PCA positions of individuals within specific clusters are highlighted with coloured symbols, and connected with shaded hulls (from PCAs shown in Fig. S3d.7).

(hi-res https://www.dropbox.com/s/8x68bzwue6qqzpg/sfig_ibd_41_cluster_pca_hgBaikalLate.png?dl=0)

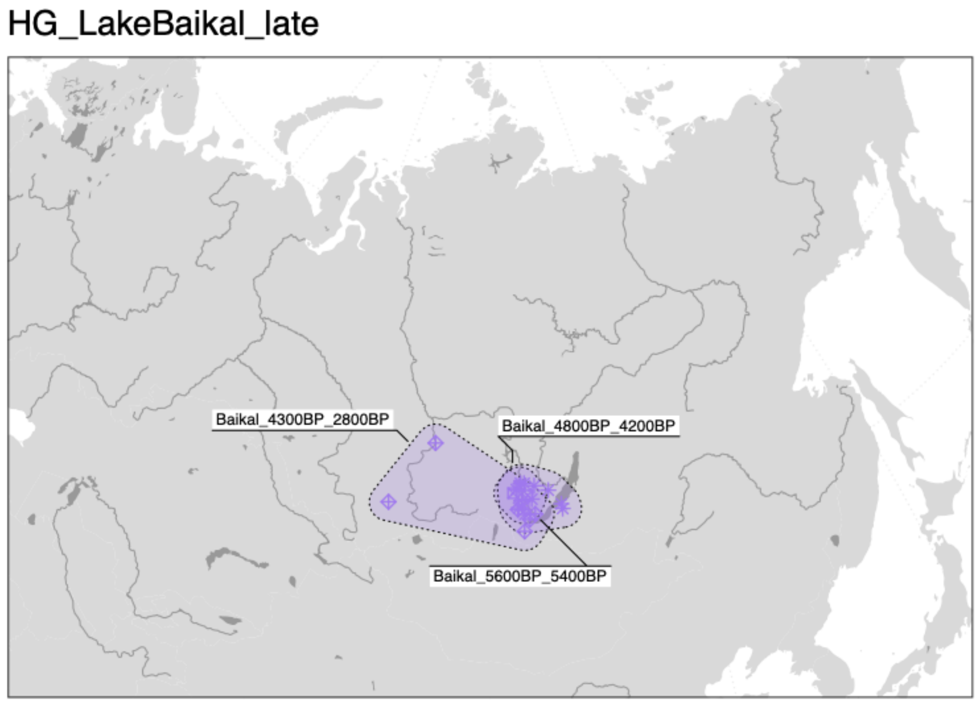


**Fig. S3f.42 Geographic distribution of individuals in cluster group *HG_LakeBaikal_late.*** Geographic locations of individuals within specific clusters are highlighted with coloured symbols, and connected with shaded hulls.

(hi-res [https://www.dropbox.com/s/hokbefzr208wpw2/sfig_ibd_42_cluster_map_hgBaikalLate.png?dl=0](https://www.dropbox.com/s/hokbefzr208wpw2/sfig_ibd_42_cluster_map_hgBaikalLate.png?dl=0))

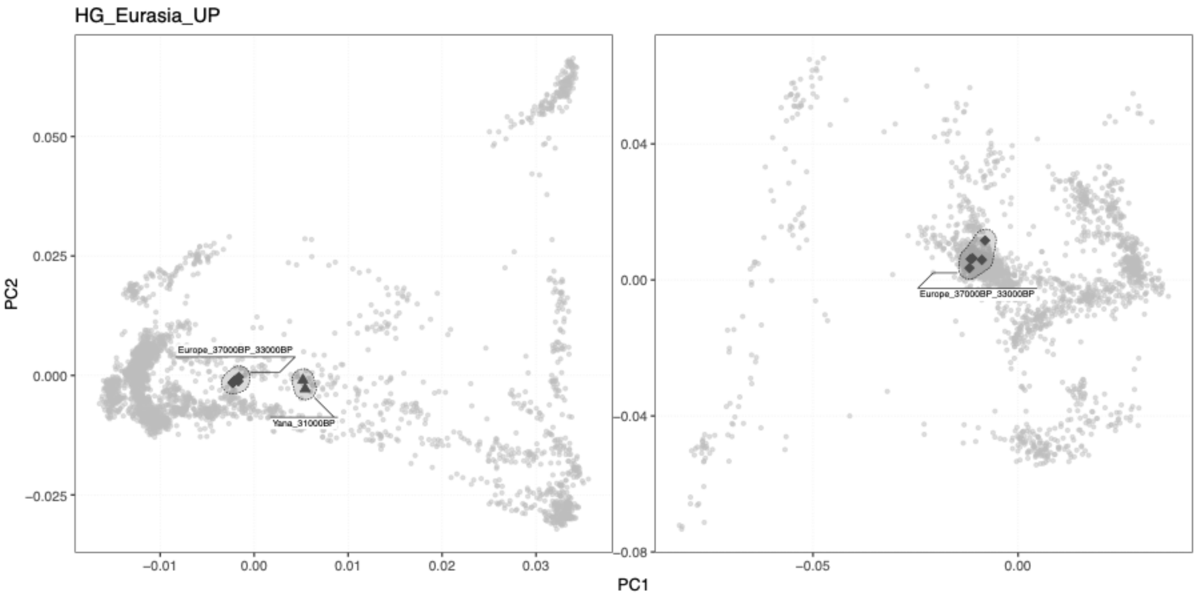


**Fig. S3f.43 PCA for cluster group *HG_Eurasia_UP***. PCA positions of individuals within specific clusters are highlighted with coloured symbols, and connected with shaded hulls (from PCAs shown in Fig. S3d.7).

(hi-res [https://www.dropbox.com/s/rqqd9o458tde2hj/sfig_ibd_43_cluster_pca_hgEurasUP.png?dl=0](https://www.dropbox.com/s/rqqd9o458tde2hj/sfig_ibd_43_cluster_pca_hgEurasUP.png?dl=0))

HG_Eurasia_UP

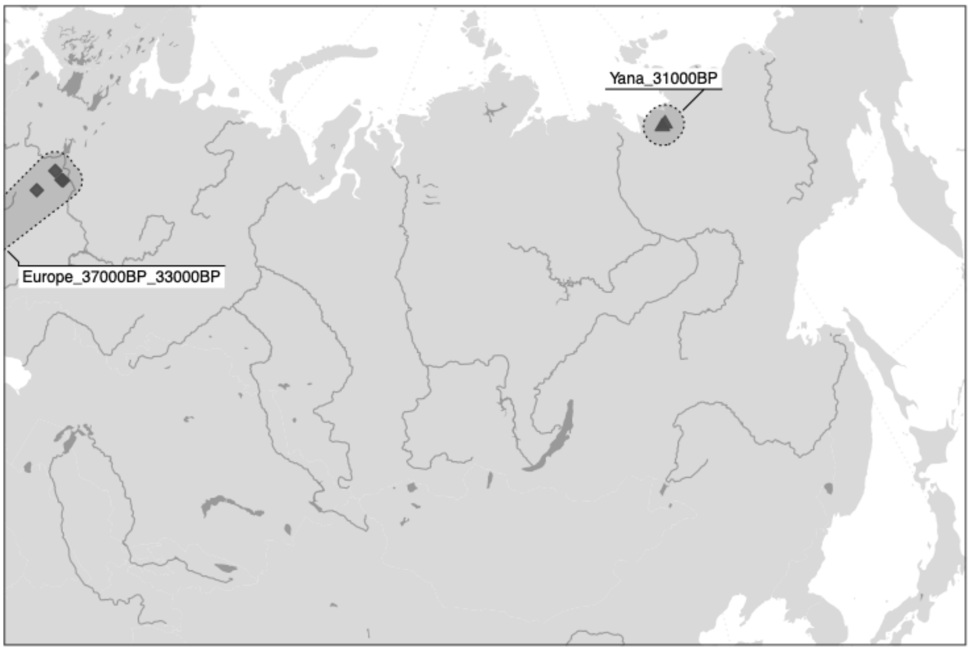


**Fig. S3f.44 Geographic distribution of individuals in cluster group *HG_Eurasia_UP.*** Geographic locations of individuals within specific clusters are highlighted with coloured symbols, and connected with shaded hulls.

(hi-res https://www.dropbox.com/s/xdowuhra8xqnegl/sfig_ibd_44_cluster_map_hgEurasUP.png?dl=0)

#### Runs of homozygosity and IBD sharing within clusters

We quantified genetic relatedness within clusters by investigating runs of homozygosity and pairwise IBD sharing among cluster individuals. The analyses showed broad differences in patterns of genetic relatedness between different cluster groups across Eurasia, associated with both spatiotemporal and subsistence contexts of the individuals. The highest amounts of IBD sharing and ROH were generally found in clusters of individuals from hunter-gatherer contexts. Individuals of comparable age from farming contexts showed lower sharing, consistent with overall higher effective population sizes in farming communities compared to forager groups (Fig. S3f.45-48).

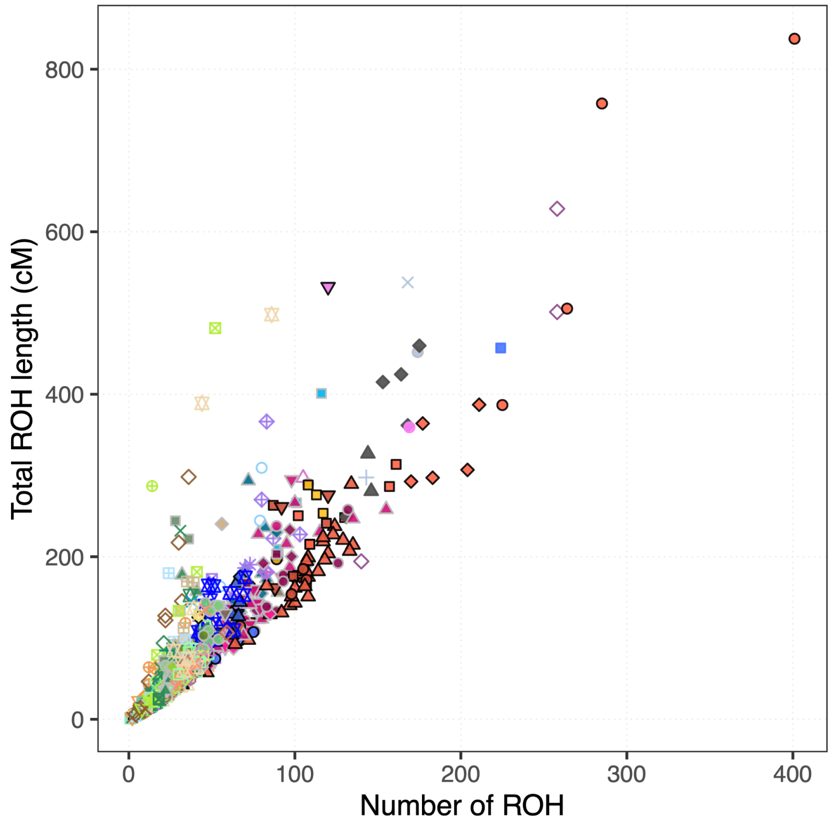


**Fig. S3f.45. Number and total length of ROH segments.** Plot shows the number and total length of ROH segments detected in the respective ancient individual. Symbol colour and shape indicated genetic cluster membership.

(hi-res https://www.dropbox.com/s/ozlivn1bl5esl67/sfig_ibd_45_roh_n_l.png?dl=0)

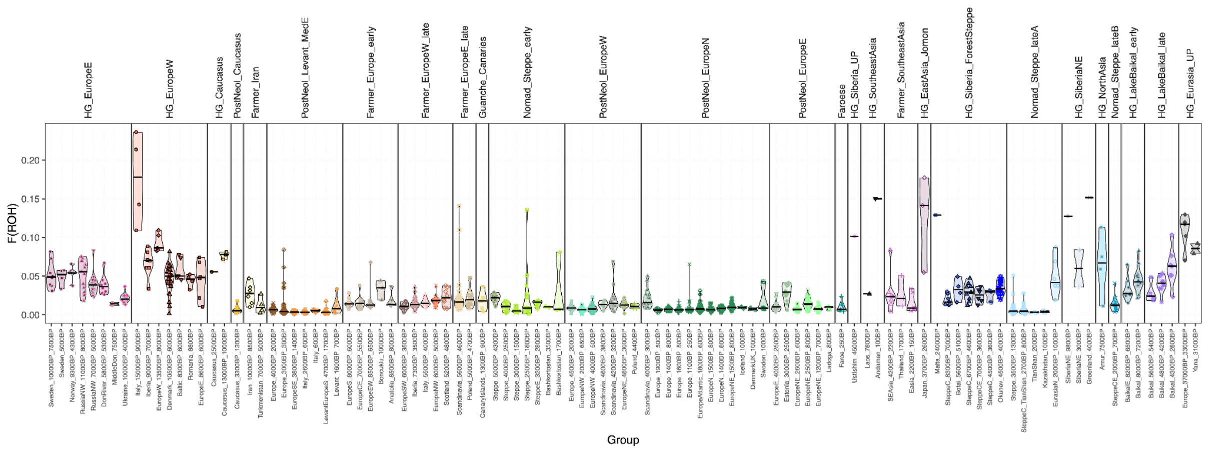


**Fig. S3f.46. Distribution of F(ROH).** Violin plots and symbols showing the distributions of F(ROH) for each individual across genetic clusters.

(hi-res https://www.dropbox.com/s/4skc4iurtigfs2y/sfig_ibd_46_froh_dist.png?dl=0)

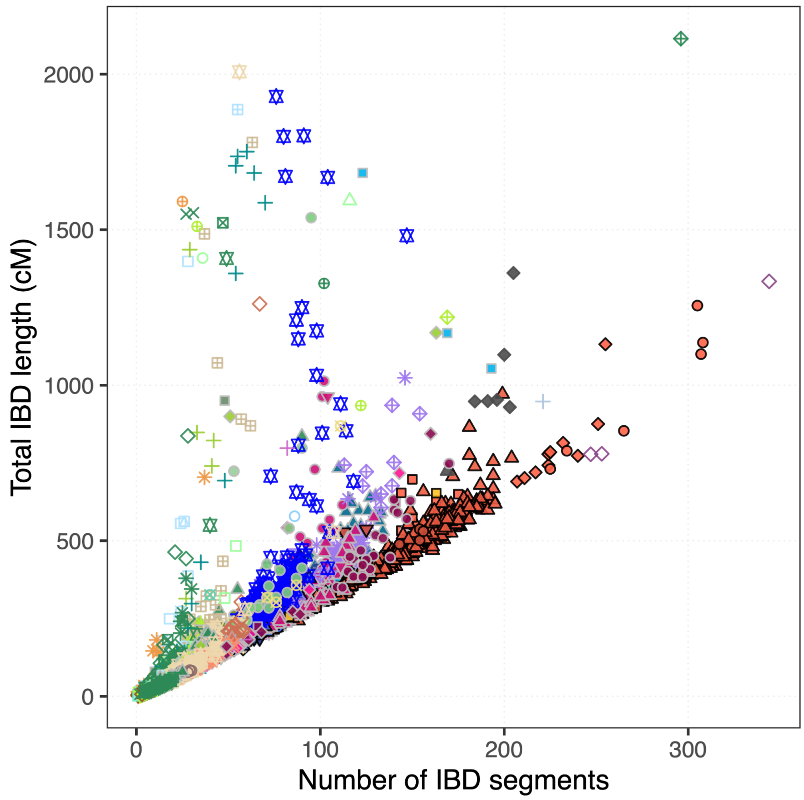


**Fig. S3f.47. Number and total length of IBD segments.** Plot shows the number and total length of IBD segments detected in the respective pair of individuals. Symbol colour and shape indicated genetic cluster membership.

(hi-res https://www.dropbox.com/s/o85v3wlt8cg2thh/sfig_ibd_47_ibd_n_l.png?dl=0)

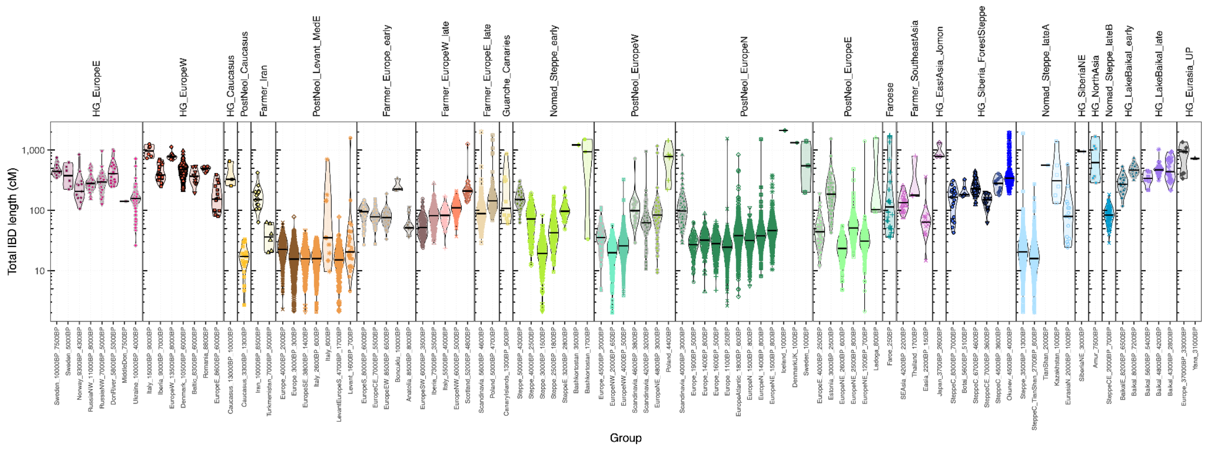


**Fig. S3f.48.** Violin plots and symbols showing the distributions of total length of segments shared IBD between pairs of individuals within the same genetic cluster.

(hi-res https://www.dropbox.com/s/xo4cd09x7y6dmt8/sfig_ibd_48_ibd_dist.png?dl=0)

#### Mixture models

We used IBD "painting profiles" to model sets of target individuals as mixtures of putative source groups. To investigate how these IBD profiles are capturing underlying population structure, we compared their similarities using the "total-variation- distance" (TVD)[12,13] measure. We calculated pairwise TVD values for each pair of individual profiles in the combined ancient and modern dataset, as well as for average profiles aggregated across all individuals within a genetic cluster. Our results show that the painting profiles readily distinguished both broad- and fine-scale genetic differentiation among the individuals and genetic clusters (Extended Data Fig 3e; Fig. S3f.49,50).

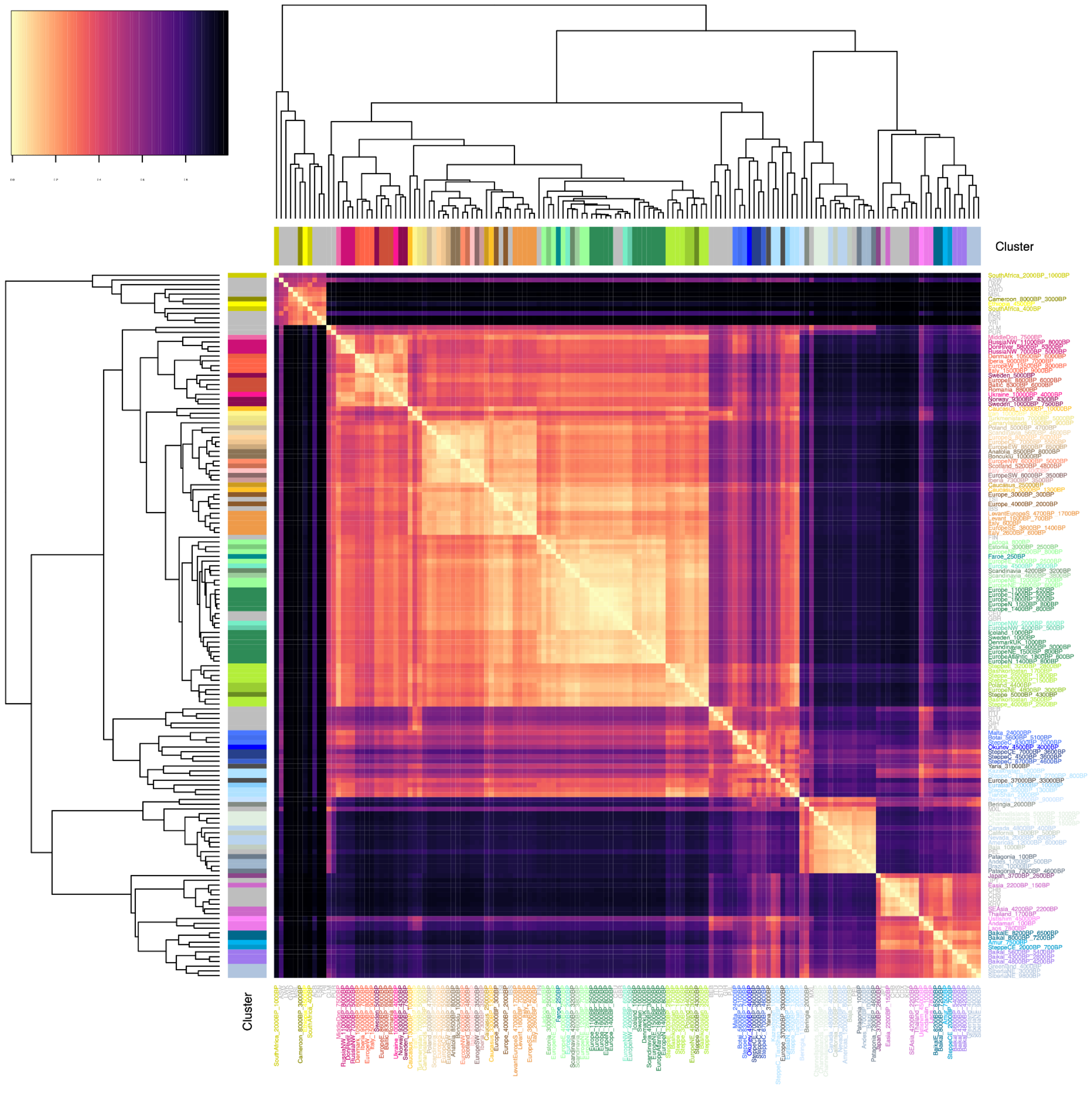


**Fig. S3f.49. Cluster IBD painting profile distances.** Heatmap showing pairwise distance between genetic cluster IBD painting profiles, measured using TVD. Coloured bars indicate cluster membership.

(hi-res https://www.dropbox.com/s/czgngu1fcwh2m5q/sfig_ibd_49_tvd_heatmap.png?dl=0)

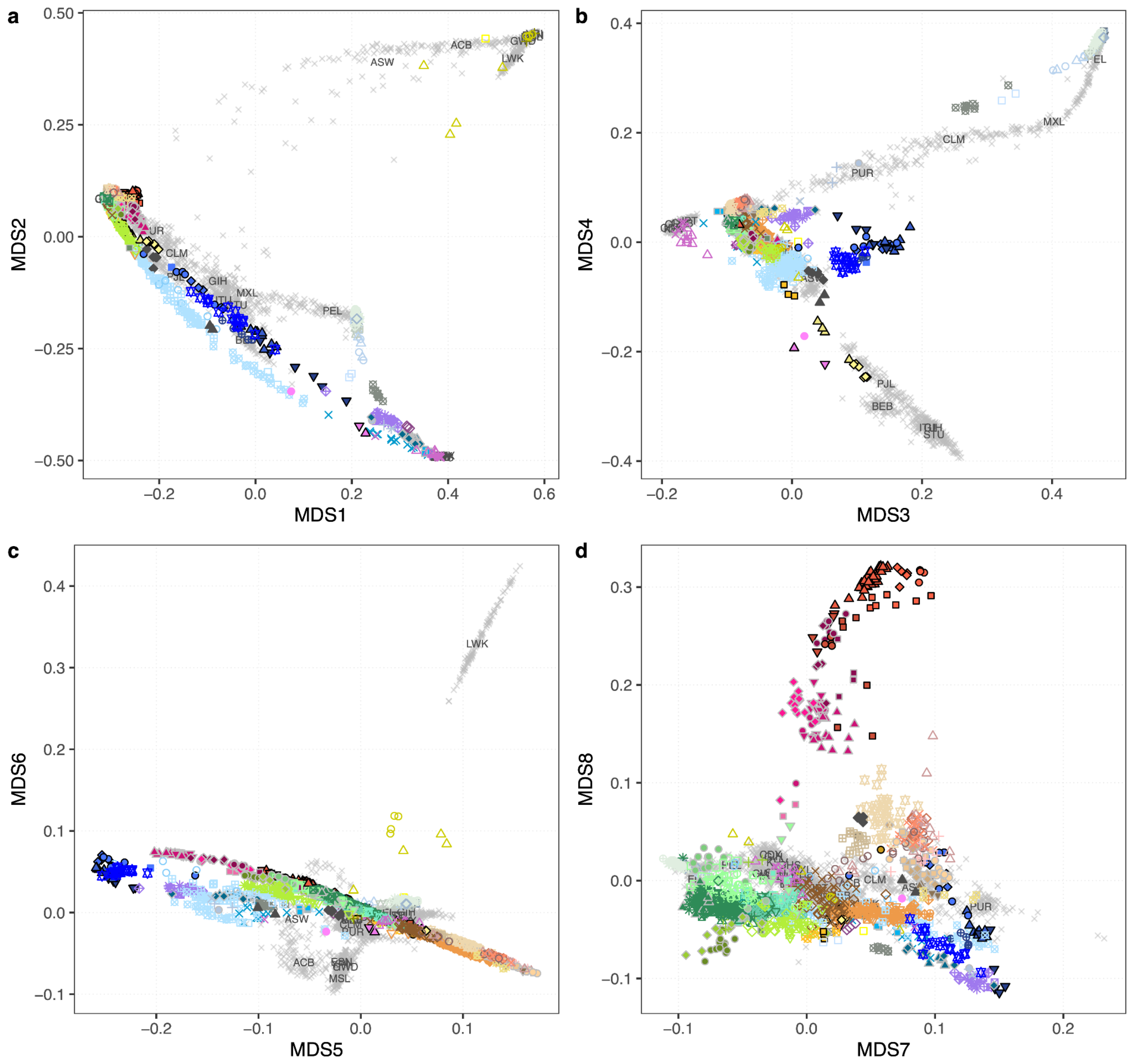


**Fig. S3f.50. Genetic structure inferred from IBD painting profiles.** (a)-(d) Plots show the first 8 dimensions of a multidimensional scaling (MDS) of individual painting profile TVDs across ancient and modern individuals. Genetic cluster membership for ancient individuals is indicated by symbol colour and shape. Present-day individuals are indicated with grey crosses, with labels indicating population median coordinates.

(hi-res https://www.dropbox.com/s/5mvrtwu16ildxf0/sfig_ibd_50_tvd_mds.png?dl=0)

We used these painting profiles in supervised modelling of target individuals as mixtures from different sets of putative source groups (Fig. S3f.51; Supplementary Data VIII). To investigate ancestry compositions across the full set of ancient individuals, we used three sets of source groups reflecting different temporal depths:

- “deep”, a set of groups representing highly differentiated deep ancestry sources
- “postNeol”, using diverse Neolithic and earlier source groups
- “postBA”, using Late Neolithic and Bronze Age source groups

We also performed analyses of more restricted spatiotemporal scope:

- “hgEur”, modelling European hunter-gatherers as mixtures of different sets of early European hunter-gatherers and selected outgroups
- “fEur”, modelling later European farmers as mixtures of earlier farmers and hunter-gatherers

In each source set analysis, we computed source group painting profiles by averaging the profiles of the included individuals within each source group. We then estimated the mixture and proportions of source profiles that best fits the profile observed in the target individuals using non-negative least squares (Supplementary Datas IX-XIII)

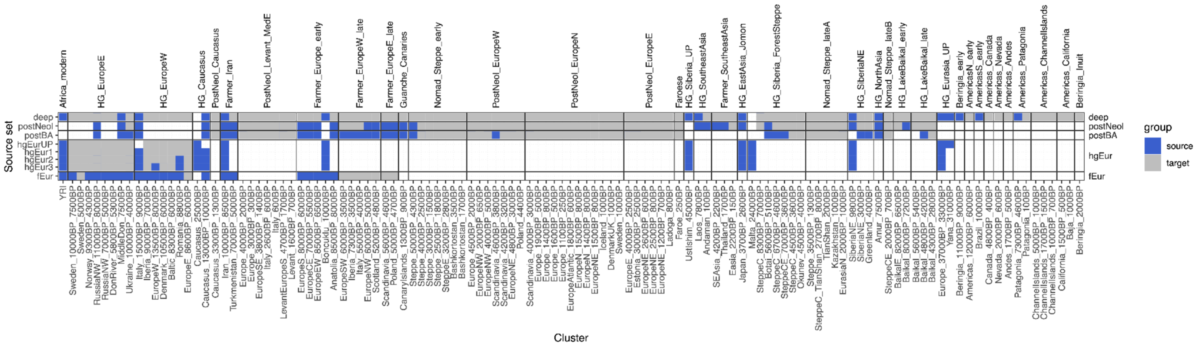


**Fig. S3f.51. Mixture model source and target groups.** Matrix showing the source and target groups used across the five source set analyses.

(hi-res https://www.dropbox.com/s/c839j2ichbziq48/sfig_ibd_51_mixmodel_matrix.png?dl=0)

We carried out a comparison of IBD-based mixture modelling to other commonly used admixture modelling methods using a well-described population model for ancient West eurasian individuals, using four putative source populations (notation as used in the literature):

- Western hunter-gathereres / WHG (Loschbour, Bichon)
- Eastern hunter-gatherers / EHG (Sidelkino, Karelia)
- Caucasus hunter-gatherers / CHG (KK1, SATP, NEO281)

- Anatolian farmers / EEF (Bar8, Bar31)

We estimated ancestry proportions for this model on a set of 185 ancient west Eurasian target individuals using three different methods

- IBD-based mixture modelling as described above
- Supervised ADMIXTURE
- qpAdm

For supervised ADMIXTURE analysis, we used pseudo-haploid genotypes in the “1000G” dataset, LD-pruned for 142,550 transversion SNPs as used in the main unsupervised ADMIXTURE analysis. For qpAdm, we used pseudo-haploid genotypes of all transversion SNPs in the “1000G” dataset, and the “right” populations *Cameroon_8000BP_3000BP, Ethiopia_4500BP, UstIshim_45000BP, Europe_37000BP_33000BP, Malta_24000BP, Caucasus_25000BP, Japan_3700BP_2600BP, Yana_31000BP, Boncuklu_10000BP* and *Iran_10000BP_8500BP*. For each individual, we selected the best-fitting model (tail probability >0.01) with the lowest number of source populations. For individuals without a good model fit (tail probability ≤0.01) we select the best model where adding another source did not improve the fit anymore (nested p-value >0.01).

The results of this analysis are shown in Fig. S3f.52. Using IBD-based mixture models, we recapitulate previously described results of ancestry mixtures in ancient Eurasians, such as

- Norwegian HGs (*Norway_9300BP_4300BP*) modelled as EHG/WHG mixture with EHG as major component[1].
- Iberian HGs (*Iberia_9000BP_7000BP*) predominantly WHG, with an influx of EHG in later individuals[2]. All Iberian individuals also show a small amount of a EEF/CHG mixed component, likely representing a pre-LGM Magdalenian ancestry component previously found in Iberian HGs[3], for which no good proxy source population is included in this model.
- European farmers modelled as predominantly EEF, with WHG admixture in varying amounts[4] (low in early Iberian farmers / Iberia_7300BP_3500BP), higher in later Scandinavian farmers / *Scandinavia_5600BP_4700BP*).
- Steppe pastoralists from the early Bronze Age modelled as predominantly EHG/CHG mixture[5], with a minor component of EEF[6].

- Later European Bronze Age individuals with Steppe-associated EHG/CHG mixture, but increased amount of EEF ancestry[4,7].

We found that results of IBD-based models were consistent with the other two methods, showing only subtle differences. Supervised ADMIXTURE results show overall reduced amounts of CHG ancestry, across all target individuals (Fig. S3f.52, middle). Results for qpAdm are more noisy with some inconsistencies of source populations modelled within target groups (Fig. S3f.52, bottom). This is not entirely surprising as qpAdm is based on sets of *f*-statistics, which can suffer from low statistical power when applied on the level of individuals as performed here. As an example, individuals with lowest coverage in the cluster of Scandinavian farmers (*Scandinavia_5600BP_4700BP*) are modelled as 100% EEF ancestry (e.g. NEO753 - 0.16X; ans005 - 0.13X; ans003 - 0.14X), likely due to lack of power to reject the single source population model with the low number of SNPs used in this analysis. Notably, CHG admixture proportions from qpAdm are comparable to those of IBD-based models, suggesting that supervised ADMIXTURE underestimates CHG ancestry.

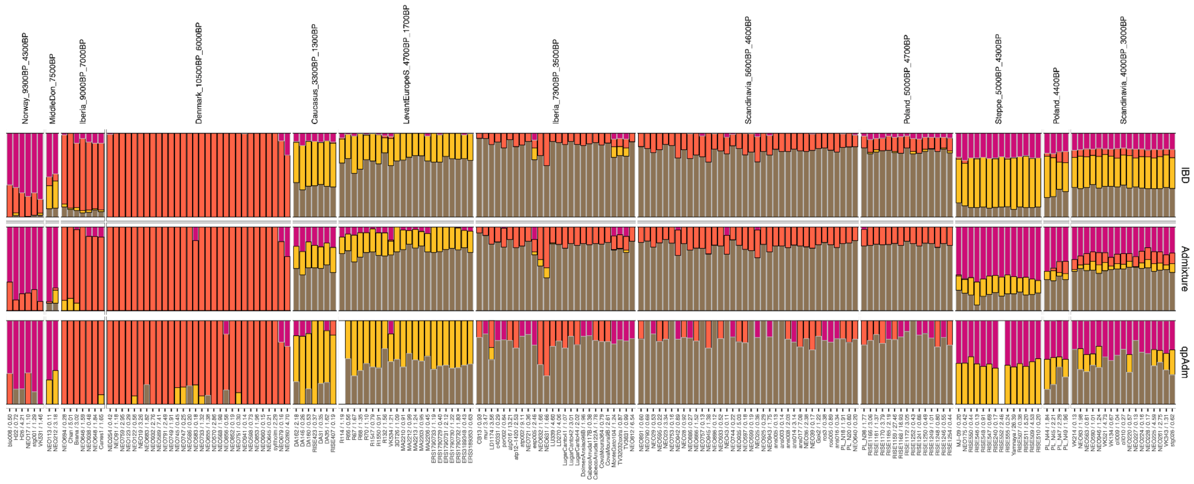


**Fig. S3f.52. Mixture model comparisons.** Barplots showing estimated ancestry proportions for XX ancient west Eurasian individuals using IBD-based mixture models (top), supervised ADMIXTURE (middle) or qpAdm (bottom). Bar colour indicates source population (WHG - orange; EHG - purple; CHG - yellow; EEF - brown). Average genomic coverage of samples is indicated in sample legend.

(hi-res https://www.dropbox.com/scl/fi/25dqnhyno2qxr58a75j4b/sfig_mixmodel_comp.png?rlkey=r0o7oybbxp6p01mgclqfovc37&dl=0)

We note that more subtle biases can occur for individuals at the lower extreme of coverage cutoff in more challenging analysis situations. One such scenario is fine-scale genetic structure modelling in low coverage individuals, e.g. distinguishing contributions from source clusters with very little genetic differentiation. An example of such a case is shown in the modelling distinguishing the early European farmer sources in later farmer individuals (set “fEur”). Our results of this modelling showed that Western European farmers derived their farmer ancestry almost exclusively from a southern European early farmer source (*EuropeS_8000BP_6500BP*, Extended Data Fig. 8e). However, a subset of individuals from Neolithic Britain were modelled as mixtures of both southern and central European early farmer sources (Extended Data Fig. 8a). This pattern occurs predominantly in the lowest coverage individuals (<0.2X), but estimated ancestry proportions are associated with large standard errors and can hence be identified as unreliable (Fig. S3f.53).

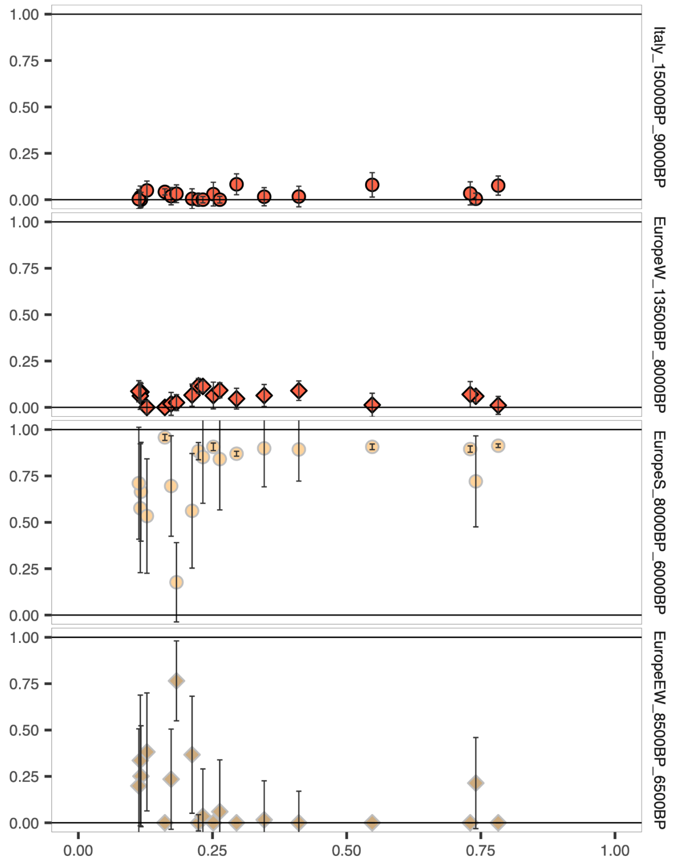


**Fig. S3f.53. Example effect of low coverage in mixture models.** Estimated ancestry proportions and associated standard errors for British Neolithic individuals, as a function of genomic coverage. Symbol colour and shape indicates source population.

(hi-res https://www.dropbox.com/scl/fi/qixlo4z8lmm5dm2yr8c67/sfig_mixmodel_lc.png?rlkey=7xun8xqkdlwpxhgj1q35obqa5&dl=0)

### 4) $^{14}$C chronology and estimates of reservoir effects

Karl-Göran Sjögren[1] and Anders Fischer[1,2]

[1]Department of Historical Studies, University of Gothenburg, 405 30 Gothenburg, Sweden

[2]Sealand Archaeology, Gl. Røsnæsvej 27, 4400 Kalundborg, Denmark

In total, 317 samples were successfully genome sequenced, of which two (NEO901 and NEO902) were only used for imputation due to unexpectedly late dating. In the present project, 272 new $^{14}$C dates were produced. They were measured either on the same samples that were analysed for DNA or on other samples from the same individuals. Most of the dates (274 samples, 32 of which failed) were performed at the $^{14}$CHRONO Centre laboratory at Queen's University, Belfast. Sample pretreatment and laboratory protocols have been described by Reimer et al.[1]. Some additional samples were analysed by the Oxford Radiocarbon Accelerator Unit (ORAU) laboratory (24 samples) and by the Keck-CCAMS Group, Irvine, California, USA (6 samples). Laboratory procedures at these labs are described in Brock et al.[2] and Beaumont et al.[3]. All new samples were also analysed for $\delta^{13}$C and $\delta^{15}$N in order to evaluate diet and possible reservoir effects. In addition, collagen quality values were measured systematically. The great majority of samples were within the range for well preserved collagen (C/N 2.9-3.6). For 32 samples, dating failed due to small sample sizes or low collagen content, however. Ten of these could be re-dated with new samples, but this was not always possible.

We also collected previously produced dates and isotope values for the graves/individuals in question, e.g.[4]. In cases where several radiocarbon measurements have been made on samples from the same individuals, a quality assessment was made of the available dates, avoiding outliers and less credible older radiometric dates. For the remaining cases, combined dates were calculated in OxCal, unless $\chi^2$-testing suggested the samples were not contemporary and should be treated as representing different individuals.

In all, 302 of the 317 samples are directly dated. For the 15 samples, where no direct dating was possible, their absolute ages were estimated on the basis of archaeological context. This includes one case (NEO587, Kongemose) where contamination from preservatives was suspected. Instead, a date from the genetically close relative NEO932; Tudse Hage) was used.

All relevant $^{14}C$ dates are presented in Supplementary Datas II-IV. Supplementary Data II shows radiocarbon ages after evaluation and combination of dates. Dates were calibrated in Oxcal online ver 4.3 using the Intcal20 calibration curve[5]. Supplementary Data III contains the calculation of these corrections, the results of which are inserted also in Table II. For calculation and presentation purposes, midpoints of both the uncorrected and the corrected date ranges were calculated. Supplementary Data IV contains all raw data pertaining to the sampled individuals.

#### Reservoir effects

For samples from coastal regions, a marine reservoir effect (MRE) may be considered. Marine reservoir effects vary over time and space[6,7]. Marine reservoir effects are expressed either as deviation from a known age, R(t), or as ΔR, i.e. deviation from the global marine calibration curve (Marine13,[8]). Corrections can be made in two ways[7,9–11]. The first is subtraction of a diet-weighted fraction of R(t) from the measured BP value, and then calibration by the atmospheric calibration curve. The second method is to use a diet-weighted fraction of the ΔR value and then use the marine curve for calibration. Here we use the first method, as ΔR values are not available for all areas sampled. Regional values for R(t) and/or ΔR are taken from publications (see Table S4.1). There are some sources of uncertainty due to small scale variation and variation over time, which must be kept in mind but cannot be taken into account here.

| **Area** | **Country** | **R(t)** | **1s** | **ΔR** | **1s** | **Comment** | **Source** |
|---|---|---|---|---|---|---|---|
| Kola peninsula | Russia | 392 | | -8 | 53 | modern value | Chrono database (Reimer & Reimer 2001[12]) |
| S Norway | Norway | 380 | 30 | | | | Günther et al. 2018[13] |
| Jutland, Limfjorden | Denmark | 375 | | | | Brackish, variable over time. Mean of 3 values | Philippsen et al. 2013[14] |
| Jutland east coast | Denmark | 350 | | | | | Olsen et al. 2009[15], 2017[16]; Larsen et al. 2018[17] |
| Jutland west coast | Denmark | 400 | | | | modern value | Standard ocean value |
| Nekselø, Zealand | Denmark | 273 | 18 | | | used for E Denmark | Fischer & Olsen 2021[18], cf Philippsen 2018[19] |
| SE Baltic | Latvia | 190 | 43 | | | Brackish, variable over time | Piličiauskas & Heron 2015[20] |
| Atlantic coast | Portugal | 495 | | 95 | 15 | | Monge Soares et al. 2016[21] |
| Cantabria | Spain | 256 | 56 | -105 | 21 | | Monge Soares et al. 2016[21] |
| Bohuslän | Sweden | 259 | 59 | -51 | 48 | modern value | Chrono database (Reimer & Reimer 2001[12]) |
| Scotland | UK | 330 | 48 | -126 | 39 | | Russel et al. 2015[22]; Ascough et al 2017[23] |
| Adriatic | | 378 | 44 | 28 | 45 | without mussels | Faivre et al. 2015[24] |

| | | | | | | | |
|---|---|---|---|---|---|---|---|
| Aegean | | 480 | 72 | 154 | 52 | | Reimer & McCormac 2002[25]; Facorellis & Vardala-Theodorou. 2015[26] |
| W Mediterranean | | 400 | 22 | 40 | 14 | | Reimer & McCormac 2002[25] |
| Mediterranean | | 390 | 15 | 45 | 14 | except Aegean | Siani et al. 2000[27]; Reimer & McCormac 2002[25] |

***Table S4.1. Values used for calculation of reservoir effect corrections and sources.***

Samples from inland sites are assumed not to be influenced by marine reservoir effects. This is a simplification but the issue cannot be resolved without local reference data and detailed knowledge about mobility patterns.

For coastal sites, the degree of marine protein intake is calculated as a linear interpolation between a marine and a terrestrial endpoint. Again, this is a simplification, but modelling of dietary fractions requires detailed knowledge of isotopic values throughout the food chain, and this is only available for some of the study areas, e.g. Denmark. Endpoint values used here are:

- Denmark: $\delta^{13}C$ -21.7 to -10.1‰ were originally measured on coeval local fish and terrestrial fauna [4,cf. 28]. For the sake of stressing uncertainties, we here use the rounded end values of -21‰ and -10‰
- Atlantic coast: $\delta^{13}C$ -21.8 to -12‰ [29] (fauna values offset by +1‰)
- Mediterranean: $\delta^{13}C$ -21 to -11.8‰ [29] (fauna values offset by +1‰).

Reservoir effects in brackish and freshwater environments (FRE) are more difficult to estimate, due to large variation over time, between lakes/estuaries, within lakes/estuaries, and between different fish species within one and the same water system.

The present study includes individuals from the brackish systems of the Limfjord (Denmark) and the Baltic. For the former we use estimates by Philippsen et al.[14]. Reservoir effects in the Baltic Sea have been discussed by several authors, e.g. Hedenström & Possnert[30], Lougheed et al.[31,32], Piličiauskas & Heron[20]. Only two of our individuals (Skateholm I and Sope) are located by the coast of this brackish environment. In these cases there was no need for correction for marine reservoir effect, as direct dating of the former was unsuccessful, whereas the latter had a completely terrestrial $\delta^{13}C$ signal. An individual from Køge Sønakke (NEO759) belongs to a seascape characterised by relatively complex aquatic reservoir effects (Olsen et al. 2017). In this case we have tentatively judged the Nekselø value (Table S4.1) to be an appropriate approximation.

Freshwater reservoir effects have been treated in a number of publications, some of which are relevant to our samples. This for instance applies to samples from Aamose[33], Iron Gates[34], Zvejnieki[35], Dnepr rapids[36,37], Lake Baikal[38,39], Sakhtysh and Minino[40,41]. We have used these estimates where relevant. For other published values, see Kulkova et al.[42],

Svyatko et al.[43], Losey et al.[44], Ramsey et al.[45]. However, for some areas there are no such values available. This concerns mostly sites in Russia. For forest areas where no direct estimates have been published, we use a standard FRE of 500 years.

Another problem is to estimate the dietary contribution of freshwater protein. In some cases, such as Lake Baikal[38], this has been shown to be proportional to $\delta^{15}N$, although other sites show no clear correlation [41]. In the latter case, the relation may be obscured by chronological or other types of variation. In inland regions, freshwater fish consumption is also indicated by low $\delta^{13}C$ values[28,36,46,47]. This is clearly visible in our samples from Ukraine and western Russia.

Isotopic signatures in most of our samples from Scandinavia on the other hand do not suggest very high levels of freshwater fish consumption, as $\delta^{13}C$ is mostly higher than -21‰ and in cases where it is lower than this, $\delta^{15}N$ is also low (<13‰), even for Mesolithic samples. Further, there is a positive correlation between $\delta^{13}C$ and $\delta^{15}N$ values, suggesting two-component mixing between terrestrial and marine sources. Only a few samples deviate from this pattern (Figures S9.1-S9.2).

In this context, we follow regional recommendations, where such are available (eg. Lake Baikal). In most cases, method 2 of Cook et al.[34] (see also Boric & Miracle[48]) is applied. This method does not involve attempts to estimate the proportion of freshwater protein but only groups the $\delta^{15}N$ values into three classes: <11‰, 11-13‰, and >13‰. This translates to 0%, 50% and 100% of the maximum FRE value. This is a coarse estimate, but has the merit of not introducing false precision into the estimates. It can be stressed that large uncertainties and high variability are at hand in the baseline values.

Steppe and arid regions of Russia and Kazakhstan present further problems in distinguishing the effect of freshwater diet from those of environmental factors such as elevation and aridity, which lead to elevated $\delta^{15}N$ in both plants, animals and humans. A further complication arises from the possible consumption of C4 plants, eg. millet, which may elevate $\delta^{13}C$ values. A number of studies[49–5152] have published FRE estimates, which are followed here. These range c. 200-400 years.

### 5) From forager to farmer in western Eurasia: an archaeological overview

Anders Fischer[1,2], T. Douglas Price[3], Karl-Göran Sjögren[1] and Kristian Kristiansen[1]

[1]Department of Historical Studies, University of Gothenburg, 405 30 Gothenburg, Sweden

[2]Sealand Archaeology, Gl. Røsnæsvej 27, 4400 Kalundborg, Denmark

[3]Laboratory for Archaeological Chemistry, University of Wisconsin-Madison, Madison, United States of America

#### Introduction

The present study has drawn on samples across a considerable transect of continental Eurasia, stretching from Lake Baikal to the Atlantic oceanfront (Fig. S5.1), and temporally spanning the conclusion of the Ice Age to the introduction of metallurgy. Human skeletal remains sampled in the course of this research derive from an extensive variety of contexts, including burial mounds, caves, bogs and the sea-floor. Considerable effort has been devoted by archaeologists for more than a century to describe and understand the patterns of cultural and environmental change that have shaped societies in western Eurasia through the later Stone Age (the Mesolithic and Neolithic ages)[1–3]. Here, we provide a synthesis of this research, which serves as a vital basis for the interpretation of novel results presented in this study, as well as an outline of current archaeological understanding.

The Mesolithic comprises a European period of fisher-hunter-gatherer (forager) societies between the end of the last Ice Age (the Pleistocene) and the introduction of agriculture. By convention, the onset of the Mesolithic corresponds to the climatic amelioration from c. 11,700 cal BP[4]. Across western Eurasia, the duration of forager lifestyles was highly variable. In some instances — the Natufian in the Levant, and the Early Neolithic of Northern China[5] — the beginning of agriculture was virtually co-terminus with the end of the Pleistocene. In other regions, outside the influence of the first spread of agriculture — e.g. much of northern Eurasia — a forager way of life continued, in some cases until historical times. In the eastern parts of our focus area, the Neolithic is - according to local research tradition - defined by the introduction of pottery, while agriculture comes later. However, for our purposes, we use the arrival of farming to define the local onset of the Neolithic, unless specified otherwise.

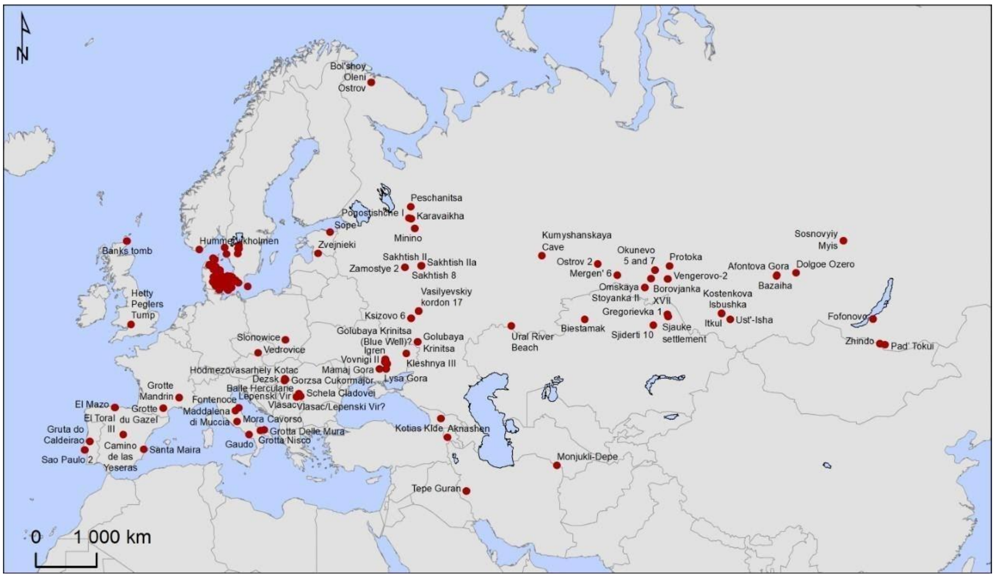


***Figure S5.1. Location of sites sampled in this study.***

The portion of Eurasia that is the focus of this study is presently (and was prehistorically) a highly varied mosaic of landscapes, biomes and cultures. From c. 12,000 cal BP the landscapes themselves underwent dramatic changes in fauna, vegetation, soils and water levels. The marked shift towards warmer climate at the end of the Pleistocene was largely responsible for these changes.

The geographic area of our study can broadly be divided into three large regions, based on archaeological material and trajectories. The first consists of northern, western, central, and southern Europe, the second includes parts of Belarus and Ukraine as well as western Russia, and the third the Urals and western Siberia (geographic division and terminology according to[6]).

#### The cradle of European agriculture - southwestern Asia

Worldwide, the origins and spread of agriculture and a Neolithic way of life was a major turning point in the evolution of human society. Agriculture is a way of obtaining food that involves domesticated plants and animals. However, the transition to farming involved much more than the simple appearance of herding or cultivation: it entailed major, long-term changes in the structure and organisation of societies. With the transition to agriculture, humans began deliberately and systematically to shape their environment far beyond their habitation areas, as well as reshaping their own biology and culture in response (in terms of reproductive rates, social organisation, etc.).

The cultivation of plants and herding of animals, village society, and pottery, did not originate in Europe. Domesticates arrived from southwest Asia, where the Neolithic began some 11,000 years ago and eventually spread into the European subcontinent and beyond[3,7,8]. In the period just preceding domestication, there was intense utilisation of wild plant foods. Between 11,000 and 10,000 cal BP, changes in the size, shape, and structure of several cereals indicate that they were domesticated[9]. The Neolithic, defined by the appearance of domesticated plants, began at that time. Several species of plants were domesticated during the period 11,000–9000 cal BP, including cereals such as emmer wheat, einkorn wheat, and barley; and at least four pulses: lentils, peas, bitter vetch, and chickpeas. In this same time period, animals were domesticated, and herding practices were adopted. Goats may have been the first livestock domesticates[10], soon joined by sheep, pigs, and cattle[11,12].

The rise of agriculture precipitated the first large settled communities, concurrent with major changes in human diet, and in the organisation of societies. The number and the size of communities expanded greatly during early stages of the Neolithic, as populations apparently concentrated in settlements. By 10,000 cal BP, new forms of residential architecture (rectangular houses) appeared, and early public constructions were seen. In southwest Asia, pottery came into use around 9500 cal BP to serve as easily produced containers for holding liquids, cooking, and storage. The establishment of shrines and ritual paraphernalia suggests a formalisation of religious activity. The complete Neolithic package of domesticates, village architecture, and pottery was thus in place shortly before 9000 cal BP, as the Neolithic began to spread to Europe[3].

#### The later Stone Age of western and central Europe

The distribution of human populations across the landscape is central to understanding the population historical dynamics of the Mesolithic and the neolithization period. A key factor contributing to this topic was the rise in sea level that accompanied the warming temperatures of the Holocene (the current geological period since the end of the Pleistocene), inundating huge areas of biologically highly productive land and reaching modern levels around 6000 cal BP (Fig. S5.2). As a consequence, most Mesolithic and many Neolithic coastal sites are under water and very few of these are investigated by marine archaeology[13–15]. At the same time there are indications of population decline in interior parts of western and central Europe during the Late Mesolithic[16]. It can reasonably be inferred that the increasingly dense mixed oak-lime-elm forest of the Atlantic climatic phase would have inhibited Mesolithic people's use of these areas, which became centres of habitation as soon as farming was introduced.

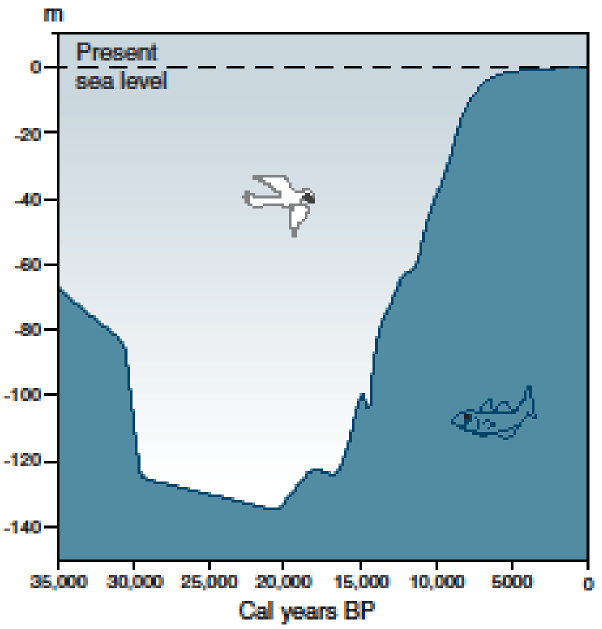


***Figure S5.2*. *Changes in global sea level over the last 35,000 years*.** *Reproduced from Fischer and Jensen[17], based on Lambeck et al.[18].*

Mesolithic settlement locations are spatially clearly associated with the distribution of contemporaneous food resources. Their frequent littoral or riverine positioning reflects the importance of aquatic species in subsistence. Correspondingly, a wide range of fishing gear is known. Likewise, equipment for hunting is richly represented - not least in terms of lithic projectile points, whereas traces of the exploitation of edible plants are generally restricted to sites with exceptional preservation quality[19,20]. Large coresident group size and permanent occupation appear to have characterised Mesolithic settlement in a number of coastal regions of this area. Social interaction among Mesolithic groups across Europe is well-documented in terms of exchange of material culture[17,21].

Immigrant Neolithic farmers gradually replaced Mesolithic foragers across the European subcontinent[22]. This westward spread took place both by land, across the Bosporus from Turkey to the Balkan Peninsula, and by sea from Cyprus and Anatolia, to the archipelago of Greece.

The 8th millennium cal BP witnessed an expansion of farmers out of Greece along two major routes (Fig. S5.3). One was along the north coast of the Mediterranean, probably by seaworthy vessels with intermittent stops that left behind communities of farmers and pastoralists[23,24]. The second arm of the spread went inland, crossing Central Europe almost to the shores of the Atlantic. These Linearbandkeramik (LBK) farmers introduced cultivation and stock rearing, large permanent houses, and a distinctive pottery decorative style to much of Central Europe[25,26]. Many archaeologists have for decades regarded the LBK a result of migration by farmers expanding into new territory[27,28] and genomic research has recently confirmed this inference[29,30].

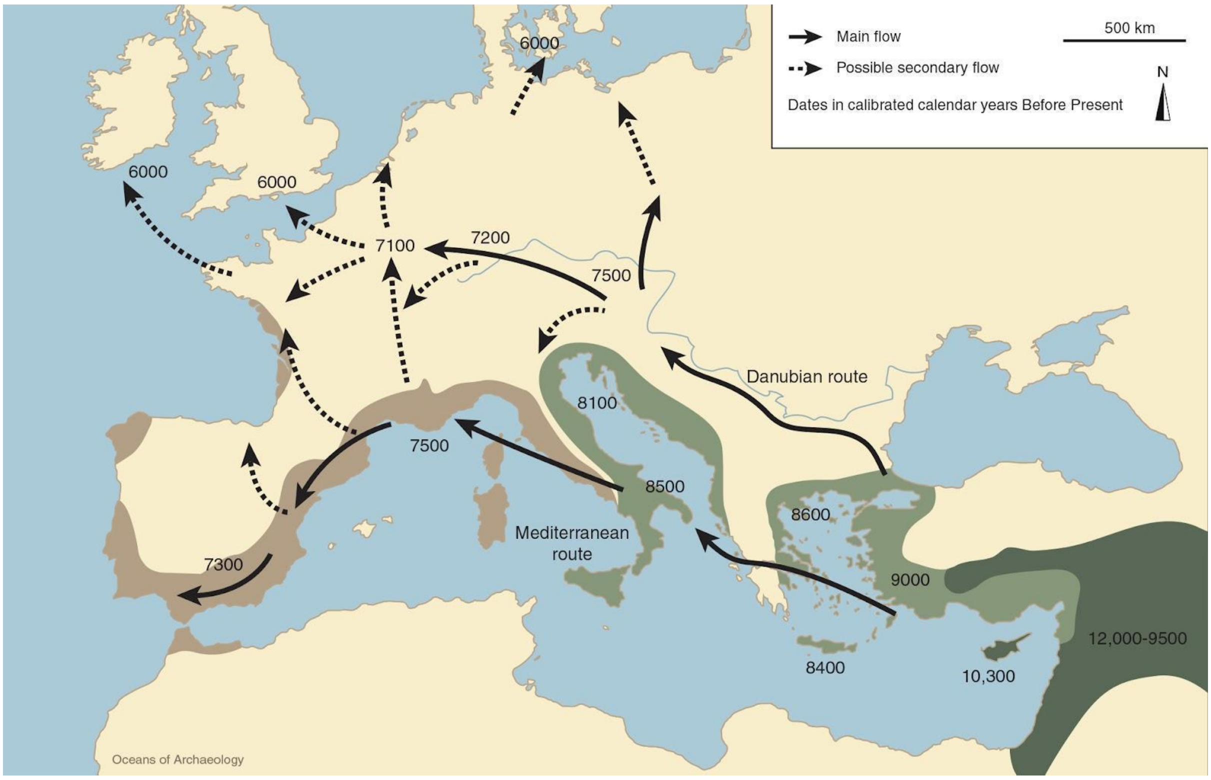


***Figure S5.3. Expansion of agricultural groups through Europe.*** *Reproduced from Fischer[31].*

Based on pottery decoration styles, the Mediterranean branch is usually referred to by names such as Impresso, Cardial and epi-Cardial ware. Groups of Cardial farmers spread inland and probably founded the Neolithic in Iberia and much of France. Genetically, they mixed to a higher degree with local forager (WHG) groups and developed a slightly different genetic profile from the central European branch[32]. In the Netherlands the Swifterbant Culture emerged, apparently with connections to both Cardial farmers and hunter-gatherers from the Ertebølle Culture of Northern Europe[25].

The final stage, expansion into the Alpine regions, the British Isles (Early Neolithic) and Northern Europe (Funnel Beaker Culture), towards the environmental limits for cultivation, took place from c. 6000 cal BP[33]. The entire journey from Greece to Scandinavia and Britain (>2000 km) took about 3000 years. These dispersals from southwest Asia to Europe took place as rapid leaps, followed by long periods of stability and adjustment[34].

Initial farming cultures expanded over broad regions and pottery styles were similar across very large areas[27,35]. Population growth and the development of permanent field systems rapidly resulted in competition and conflict between groups, however, and by 5500 cal BP, the subcontinent was occupied by a large number of culturally diversified populations.

In southeastern Europe developments in the Neolithic were dramatic, witnessing the rise of large settlements often on tells — huge mounds of human refuse and building material, accumulated in the same place over generations. This was a time of major cultural fluorescence among a series of cultures in Serbia, Romania, Bulgaria, and eastern Hungary — population numbers increased, large villages and towns appeared, technological innovations including the first copper production flourished, long distance trade expanded, and social inequality was pronounced[27,36–39].

Later Neolithic settlements across Europe were often located in defensible positions and heavily fortified. At the same time, trade and exchange expanded in scope. A variety of materials and finished goods were moved long distances across Europe[40]. Obtaining raw materials, manufacturing trade items, and transporting finished goods became an important part of Neolithic economic systems. Flint, for example, was mined in Denmark, Belgium, England, and elsewhere and polished into fine axes for trade[33].

#### The later Stone Age of eastern Europe and western Siberia

Important innovations in technology, economy and social organisation took place in eastern Europe and western Siberia prior to the arrival of agriculture. Among the early changes can be emphasised the repopulation of the northern regions after the glacial maximum, the introduction of pressure blade technique, the dispersal of pottery production, and the general tendency for settlement stability, economic intensification, and probably social complexity within late fisher-hunter-gatherer populations.

The earliest of these technological innovations was the development and spread of pressure flaking techniques for the production of sharp-edged lithic blades. This was closely connected with the production of composite hunting equipment such as slotted bone points. The earliest documented instances of pressure flaked blades and microblades are found in eastern Asia and date to at least 20,000 cal BP[41,42]. From this region, the technology seems to have spread both eastwards into America and westwards into Europe. In western Russia and Scandinavia, it is first found in Early Mesolithic assemblages, for instance in the Russian Veretye Culture and in the south Scandinavian late Maglemose and the Kongemose

Cultures[43,44]. Some researchers connect the dispersion of this technology with migrations of people associated with distinctive Eastern Hunter-Gatherer (EHG) genetic ancestry[45].

The emergence of pottery is one of the more significant technological changes in human prehistory. At present, it is widely acknowledged that the oldest pottery appeared in eastern Asia during the Late Palaeolithic. Dates of c. 18,000 cal BP for southern China, c. 16,700 cal BP for Japan and c. 15,900 cal BP for the Russian Far East are suggested[46]. From these very early centres, the technology spread gradually towards the west, reaching the area east of Lake Baikal at c. 14,000 cal BP[46]. Here, however, the dispersal seems to have halted for several thousand years. At present knowledge, pottery appeared c. 8500-8000 cal BP in western Siberia and the Urals[47]. In the region further to the west, i.e. Ukraine and western Russia, pottery was introduced around or shortly after c. 8000 cal BP[48,49], while the Narva pottery and Comb ware in the eastern Baltic zone are again somewhat later, from around 7500 cal BP[50]. This early pottery has been subdivided into a plethora of cultural groups, mostly based on decoration[51,52]. While often richly decorated, vessel shapes are less varied and rather simple, consisting mostly of pointed-based or flat-bottomed, rounded bowls. Analyses of lipids, preserved in the clay matrix, conducted in several geographical regions suggest a variety of foods being processed, with a notable presence of aquatic products, i.e. freshwater fish and molluscs, increasing over time[52–54]. This is consistent with the human isotope data from the present study.

Thus, a general east-west trend is apparent in the dates of early pottery. In local literature pottery is used to define the division between the Mesolithic and the Neolithic ages in these regions, but this innovation apparently did not imply any dramatic change in subsistence economy. Instead, these societies persisted as pottery-using foragers for millennia. Several scenarios have been proposed to account for the emergence and spread of early pottery, such as continuous spread from east to west[55], or independent innovation in various Eurasian foraging communities[46].

Even if no indications of agriculture are associated with the early pottery, important developments seem to occur both in the economic, social and ritual spheres. Thus, an increasing exploitation of seasonally abundant resources, higher degree of settlement stability, emergence of complex settlements (sometimes fortified), large cemeteries and sacrificial sites in the form of ritual mounds, are features of this period[47,56].

In eastern Europe and northwestern Asia, the temporal patterning of the introduction of domesticates is in marked contrast to central and western Europe, with many species appearing later or much later than in more western regions. The delayed introduction of domesticated plants and animals, at least in the form that ultimately derives from southwestern Asia, may perhaps be related to the adaptive success of the local hunter-gatherer societies. It has been suggested that other plants may have been cultivated, such as buckwheat[57], but this remains uncertain at present.

The introduction of agriculture of European type took place via Moldavia and western Ukraine, before 7000 cal BP[58]. Bones of domesticated animals are found in low numbers at the large settlements and cemeteries along the Dnipro rapids, after c. 7000 cal BP. Based on human isotopes, these are thought to play a very minor role in the economy, in favour of the dominance of fish food[59]. A later development is the fully agricultural Trypillia Culture (c. 6100-5000 cal BP), characterised by large, highly structured settlements with up to several thousands inhabitants[60]. These agricultural groups would have coexisted with forager groups along the Dnipro as well as in the forest belt further north. In the forest belt of western Russia (as well as eastern Baltic and Finland), clear indications of agriculture do not appear until well after 5000 cal BP, probably in the context of Corded Ware groups[61,62].

We find throughout much of the region a frontier between farming communities in the west and south, and pottery-using foragers to the northeast, from the Black Sea to the eastern Baltic[63,64 Fig. 8.1]. The latter groups defy in certain ways traditional classifications, and some of them reached high degrees of social organisation and stratification, facilitated by the resource potential of the environment (forest to steppe, intersected with large, productive rivers and lakes, capable of sustaining long-term stable settlements).

Some of this variation has been summarised as a prelude to the formation of the Yamnaya nomadic lifeway from the mid to later 6$^{th}$ millennium cal BP[64,65]. Examples of these groups are found in the Dnipro-Donets Culture, also including Sredni-Stog. In their cemeteries, we find richly furnished graves with symbols of power, most clearly at Khvalynsk at the Volga. These societies were linked together in far-reaching forms of exchange[8,66].

East of the Urals, in western Siberia, we find little or no traces of cultivation and stock keeping prior to c. 5000 cal BP. Only by the mid 5$^{th}$ millennium cal BP do we find indications of agro-pastoralism in northern central Asia, and in the Altai and southern Siberia even later[67,68].

#### References

1. Steenstrup, J. J. S., Forchhammer, J. G. & Worsaae, J. J. A. Geologisk-antiqvariske

Undersøgelser angaaende Landets ældste Natur- og Cultur-Forhold. in *The Neolithisation of Denmark - 150 years of debate* (eds. Fischer, A. & Kristiansen, K.) 34–44 (JR Collis Publications, 2002).

2. Childe, V. G. *The Dawn of European Civilization*. (K. Paul, Trench, Trubner & Company, 1925).
3. Price, T. D. *Europe's First Farmers*. (Cambridge University Press, 2000).
4. Douglas Price, T. The European Mesolithic. *Am. Antiq.* **48**, 761–778 (1983).
5. Lu, H. *et al.* Earliest domestication of common millet (*Panicum miliaceum*) in East Asia extended to 10,000 years ago. *Proc. Natl. Acad. Sci. U. S. A.* **106**, 7367–7372 (2009).
6. Jordan, P. Großgliederung Europas nach kulturräumlichen Kriterien [The large-scale division of Europe according to cultural-spatial criteria]. *Eur. Reg.* **13.2005**, 162–173 (2005).
7. Shennan, S. *The First Farmers of Europe: An Evolutionary Perspective*. (Cambridge University Press, 2018).
8. Chapman, J. *Forging Identities in the Prehistory of Old Europe: Dividuals, Individuals and Communities, 7000-3000 BC*. (Sidestone Press, 2020).
9. Weiss, E. & Zohary, D. The Neolithic Southwest Asian Founder Crops: Their Biology and Archaeobotany. *Curr. Anthropol.* **52**, S237–S254 (2011).
10. Zeder, M. A. & Hesse, B. The initial domestication of goats (Capra hircus) in the Zagros mountains 10,000 years ago. *Science* **287**, 2254–2257 (2000).
11. Zeder, M. A. The Domestication of Animals. *J. Anthropol. Res.* **68**, 161–190 (2012).
12. Frantz, L. A. F., Bradley, D. G., Larson, G. & Orlando, L. Animal domestication in the era of ancient genomics. *Nat. Rev. Genet.* **21**, 449–460 (2020).
13. Galili, E., Horwitz, L. K., Eshed, V. & Rosen, B. Submerged Pottery Neolithic Settlements off the Coast of Israel: Subsistence, Material Culture and the Development of Separate Burial Grounds. in *Under the Sea: Archaeology and Palaeolandscapes of the Continental Shelf* (eds. Bailey, G. N., Harff, J. & Sakellariou, D.) 105–130 (Springer

International Publishing, 2017).

14. Fischer, A. The tip of the iceberg - taking stock of the early prehistoric submarine record for Europe and the eastern Mediterranean. in *Oceans of Archaeology* (eds. Fischer, A. & Pedersen, L.) 150–161 (Jutland Archaeological Society, 2018).
15. Astrup, P. M. *Sea-level Change in Mesolithic Southern Scandinavia: Long- and Short-term Effects on Society and the Environment*. (Aarhus Universitetsforlag, 2018).
16. Jochim, M. The origins of agriculture in south-central Europe. *Europe's first farmers* (2000).
17. Jensen, T. Z. T. & Fischer, A. Gifts from the sea - inland indications of early prehistoric activity at now submerged coasts. in *Oceans of Archaeology* (eds. Fischer, A. & Pedersen, L.) 30–41 (Jutland Archaeological Society, 2018).
18. Lambeck, K., Rouby, H., Purcell, A., Sun, Y. & Sambridge, M. Sea level and global ice volumes from the Last Glacial Maximum to the Holocene. *Proc. Natl. Acad. Sci. U. S. A.* **111**, 15296–15303 (2014).
19. Zvelebil, M. Plant Use in the Mesolithic and its Role in the Transition to Farming. *Proceedings of the Prehistoric Society* **60**, 35–74 (1994).
20. Fischer, A. *et al.* The composition of Mesolithic food – evidence from a submerged settlement on the Argus Bank, Denmark. *Acta Archaeol.* **78**, 163–178 (2007).
21. Peeters, J. *et al. Paradise Lost? Insights into the early prehistory of the Netherlands from development-led archaeology*. (Rijksdienst voor het Cultureel Erfgoed, 2017).
22. Whittle, A., Pollard, J. & Greaney, S. *Ancient DNA and the European Neolithic: Relations and Descent*. (Oxbow Books, 2023).
23. Isern, N., Zilhão, J., Fort, J. & Ammerman, A. J. Modeling the role of voyaging in the coastal spread of the Early Neolithic in the West Mediterranean. *Proc. Natl. Acad. Sci. U. S. A.* **114**, 897–902 (2017).
24. Zeder, M. Out of the Fertile Crescent: The dispersal of domestic livestock through Europe and Africa. in *Human Dispersal and Species Movement* (eds. Boivin, N.,

Crassard, R. & Petraglia, M.) 261–303 (Cambridge University Press, 2017).

25. Constantin, C., Ilett, M. & Burnez-Lanotte, L. La Hoguette, Limburg and the Mesolithic: some questions. in *Pots, farmers and foragers: Poterry traditions and social interaction in the earliest Neolithic of the Lower Rhine Area* 41–48 (Leiden University Press, 2010).
26. Bickle, P. & Whittle, A. *The First Farmers of Central Europe: Diversity in LBK Lifeways*. (Oxbow Books, 2013).
27. Tringham, R. *Hunters, Fishers and Farmers of Eastern Europe, 6000-3000 B.C*. (Hutchinson University Library, 1968).
28. van Andel, T. H. & Runnels, C. M. The Earliest Farmers in Europe. *Antiquity* **69**, 481–500 (1995).
29. Mathieson, I. *et al.* The genomic history of southeastern Europe. *Nature* **555**, 197–203 (2018).
30. Nikitin, A. G. *et al.* Interactions between earliest Linearbandkeramik farmers and central European hunter gatherers at the dawn of European Neolithization. *Sci. Rep.* **9**, 19544 (2019).
31. Fischer, A. The qualities of the submarine Stone Age. in *Oceans of Archaeology* (eds. Fischer, A. & Pedersen, L.) 180–191 (Højbjerg, 2018).
32. Olalde, I. *et al.* A Common Genetic Origin for Early Farmers from Mediterranean Cardial and Central European LBK Cultures. *Mol. Biol. Evol.* **32**, 3132–3142 (2015).
33. Sørensen, L. From Hunter to Farmer in Northern Europe. Migration and adaptation during the Neolithic and Bronze Age. *Acta Archaeologica Supplementa* **85**, (2014).
34. Bocquet-Appel, J.-P., Naji, S., Vander Linden, M. & Kozlowski, J. Understanding the rates of expansion of the farming system in Europe. *J. Archaeol. Sci.* **39**, 531–546 (2012).
35. Midgley, M. *TRB Culture: The First Farmers of the North European Plain*. (Edinburgh University Press, 1992).
36. Higham, T. *et al.* New perspectives on the Varna cemetery (Bulgaria) – AMS dates and

social implications. *Antiquity* **81**, 640–654 (2007).

37. Czekaj-Zastawny, A., Kabaciński, J. & Terberger, T. Long distance exchange in the Central European Neolithic: Hungary to the Baltic. *Antiquity* **85**, 43–58 (2011).

38. Porčić, M. Social complexity and inequality in the Late Neolithic of the Central Balkans: reviewing the evidence. *Doc. Praehist.* **39**, 167–184 (2012).

39. Radivojević, M. & Roberts, B. W. Balkan metallurgy in a Eurasian context. in *The Rise of Metallurgy in Eurasia: Evolution, Organisation and Consumption of Early Metal in the Balkans* (eds. Radivojević, M., Roberts, B., Marić, M., Kuzmanović-Cvetković, J. & Rehren, T.) (Archaeopress Publishing Ltd, 2021).

40. Windler, A. From the Aegean Sea to the Parisian Basin. How Spondylus can rearrange our view on trade and exchange. *Metalla* **20**, 87–115 (2013).

41. Desrosiers, P. M. *The Emergence of Pressure Blade Making: From Origin to Modern Experimentation*. (Springer Science & Business Media, 2012).

47. Piezonka, H. *et al.* The emergence of hunter-gatherer pottery in the Urals and West Siberia: New dating and stable isotope evidence. *J. Archaeol. Sci.* **116**, 105100 (2020).
48. Hartz, S. *et al.* Hunter-Gatherer Pottery and Charred Residue Dating: New Results on Early Ceramics in the North Eurasian Forest Zone. *Radiocarbon* **54**, 1033–1048 (2012).
49. Piezonka, H. *et al.* The Early and Middle Neolithic in NW Russia: radiocarbon chronologies from the Sukhona and Onega regions. *Documenta Praehistorica* **44**, 122–151 (2017).
50. Piezonka, H. *et al.* Stone Age Pottery Chronology in the Northeast European Forest Zone: New AMS and EA-IRMS Results on Foodcrusts. *Radiocarbon* **58**, 267–289 (2016).
51. Jordan, P. & Zvelebil, M. *Ceramics Before Farming: The Dispersal of Pottery Among Prehistoric Eurasian Hunter-Gatherers*. (Left Coast Press, 2009).
52. Piezonka, H. *Jäger, Fischer, Töpfer: Wildbeutergruppen mit früher Keramik in Nordosteuropa im 6. und 5. Jahrtausend v. Chr. (Archäologie in Eurasien)*. (Habelt, 2015).
53. Oras, E. *et al.* The adoption of pottery by north-east European hunter-gatherers: Evidence from lipid residue analysis. *J. Archaeol. Sci.* **78**, 112–119 (2017).
54. Papakosta, V. Early Pottery Use among Hunter-Gatherers around the Baltic Sea. (Department of Archaeology and Classical Studies, Stockholm University, 2020).
55. Jordan, P. *et al.* Modelling the diffusion of pottery technologies across Afro-Eurasia: emerging insights and future research. *Antiquity* **90**, 590–603 (2016).
56. Borzunov, V. A. The neolithic fortified settlements of the Western Siberia and Trans-Urals. *Rossijskaâ arheologiâ* 20–34 (2013).
57. Alenius, T., Mökkönen, T. & Lahelma, A. Early farming in the northern boreal zone: Reassessing the history of land use in southeastern Finland through high-resolution pollen analysis. *Geoarchaeology* **28**, 1–24 (2013).
58. Matuzeviciute, G. M. The adoption of agriculture: archaeobotanical studies and the

earliest evidence for domesticated plants. in *Prehistoric Ukraine: from the first hunters to the first farmers* (eds. Potekhina, I. D. & Lillie, M. C.) 309–326 (Oxbow books, 2020).

59. Budd, C. & Lillie, M. C. The prehistoric populations of Ukraine : stable isotope studies of fisher-hunter-forager and pastoralist-incipient farmer dietary pathways. in *Prehistoric Ukraine: from the first hunters to the first farmers* (eds. Potekhina, I. D. & Lillie, M. C.) 283–307 (Oxbow Books, 2020).
60. Müller, J., Rassmann, K. & Videiko, M. *Trypillia Mega-Sites and European Prehistory: 4100-3400 BCE*. (Routledge, 2016).
61. Heron, C. *et al.* Cooking fish and drinking milk? Patterns in pottery use in the southeastern Baltic, 3300–2400 cal BC. *J. Archaeol. Sci.* **63**, 33–43 (2015).
62. Nordqvist, K. & Heyd, V. The Forgotten Child of the Wider Corded Ware Family: Russian Fatyanovo Culture in Context. *Proceedings of the Prehistoric Society* **86**, 65–93 (2020).
63. Lillie, M. C. & Potekhina, I. D. Introduction. in *Prehistoric Ukraine: from the first hunters to the first farmers* (eds. Potekhina, I. D. & Lillie, M. C.) (Oxbow books, 2020).
64. Anthony, D. W. *The Horse, the Wheel, and Language*. (Princeton University Press, 2010).
65. Parzinger, H. *Die frühen Völker Eurasiens: vom Neolithikum bis zum Mittelalter*. (C.H.Beck, 2006).
66. Chapman, J. Expansion and social change at the time of Varna. *Counterpoint: essays in Archaeology and Heritage Studies in honour of Professor Kristian Kristiansen* 301–308 (2013).
67. Ryabogina, N. E. & Ivanov, S. N. Ancient agriculture in Western Siberia: problems of argumentation, paleoethnobotanic methods, and analysis of data. *Archaeology, Ethnology and Anthropology of Eurasia* **39**, 96–106 (2011).
68. Spengler, R. N., Ryabogina, N., Tarasov, P. E. & Wagner, M. The spread of agriculture into northern Central Asia: Timing, pathways, and environmental feedbacks. *Holocene* **26**, 1527–1540 (2016)

### 6) Catalogue of Danish archaeological sites

Edited by Anders Fischer[1,2] and Per Lysdahl[3]

[1]Department of Historical Studies, University of Gothenburg, 405 30 Gothenburg, Sweden

[2]Sealand Archaeology, Gl. Røsnæsvej 27, 4400 Kalundborg, Denmark

[3]Vendsyssel Historical Museum, Museumsgade 3, DK-9800 Hjørring

#### Introduction

Ancient DNA sampling across Denmark has been undertaken with the explicit aim of analysing a comprehensive assemblage of the total population of human skeletal remains from the Mesolithic and the Early Neolithic, curated in public museums. The genomic data produced should, as far as possible, reflect the full variability of the skeletal population in question with regard to chronology, geography and depositional environments. Additionally, a further direct aim was to maximise representation of social diversity as potentially reflected in burial types, sacrificial depositions and stray human bones from mentioned periods. Skeletons believed to belong to the Middle and Late Neolithic have only been sampled in special cases. Individuals of even later date have merely come into the assemblage by chance. Skeletal remains of 197 humans dating earlier than 3000 cal BP were sampled in the course of this study. Of these, 100 were found to have DNA preservation of acceptable quality. Their find locations are shown in Figures S6.1 and S6.2.

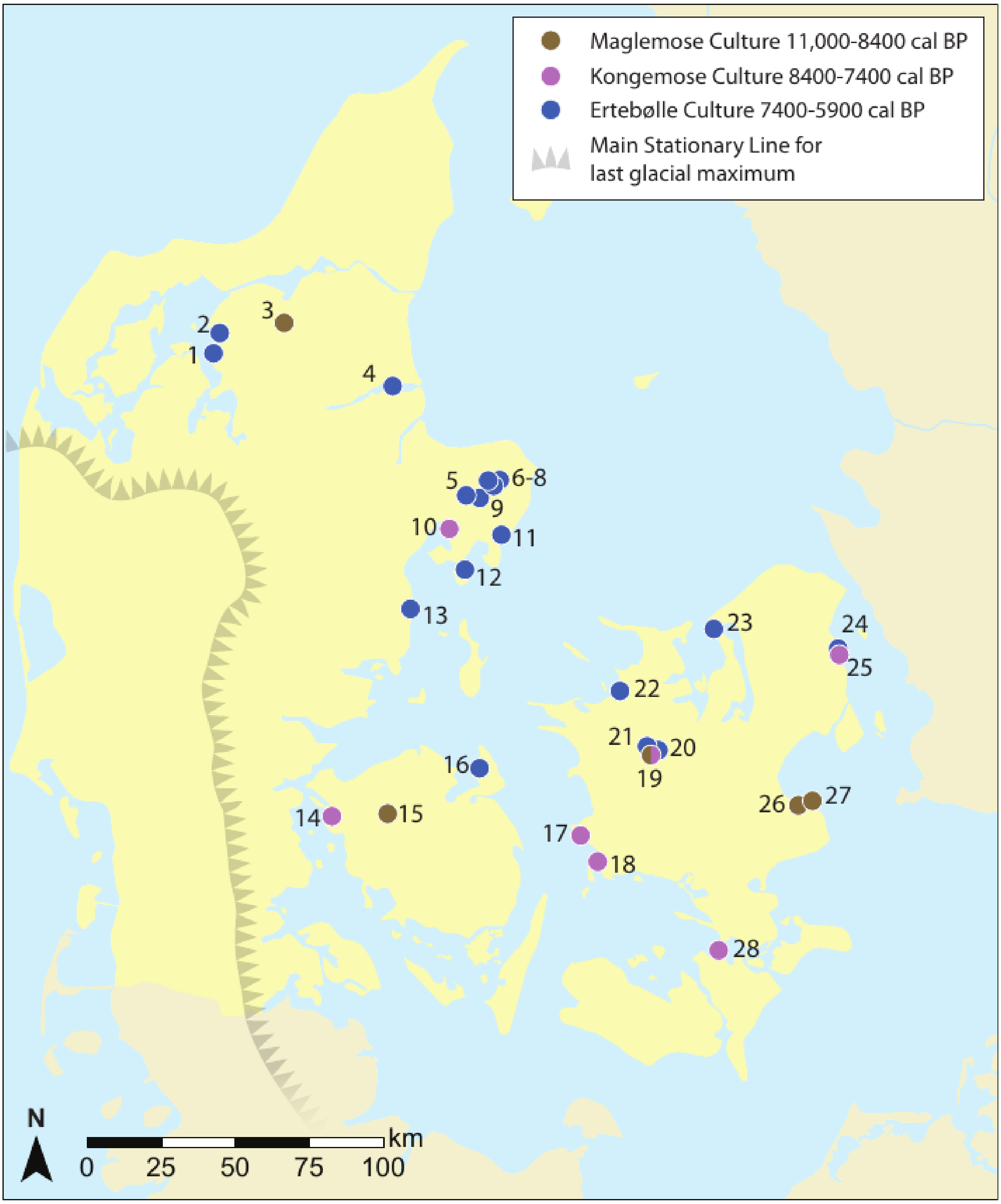


***Figure S6.1. The geographic distribution of the study's 38 Mesolithic individuals from Denmark.*** *All three major chronological groups (Maglemose, Kongemose, Ertebølle) have been sampled to comparable high intensity. The sites in question are: 1 Ertebølle (*locus classicus*), 2 Bjørnsholm, 3 Hedegaard, 4 Havnø, 5 Koed, 6 Nederst, 7-9 Fannerup D, E and F, 10 Rønsten, 11 Holmegård-Djursland, 12 Vængesø II, 13 Norsminde, 14 Tybrind Vig, 15 Koelbjerg, 16 Langø Skaldynge, 17 Korsør Nor, 18 Tudse Hage, 19 Bodal K, 20 Ravnsbjerggård II, 21 Kongemose (*locus classicus*), 22 Dragsholm, 23 Sølager, 24 Henriksholm-Bøgebakken, 25 Vedbæk Boldbaner, 26 Strøby Grøftemark, 27 Køge Sønakke, 28 Orehoved Sejlrende.*

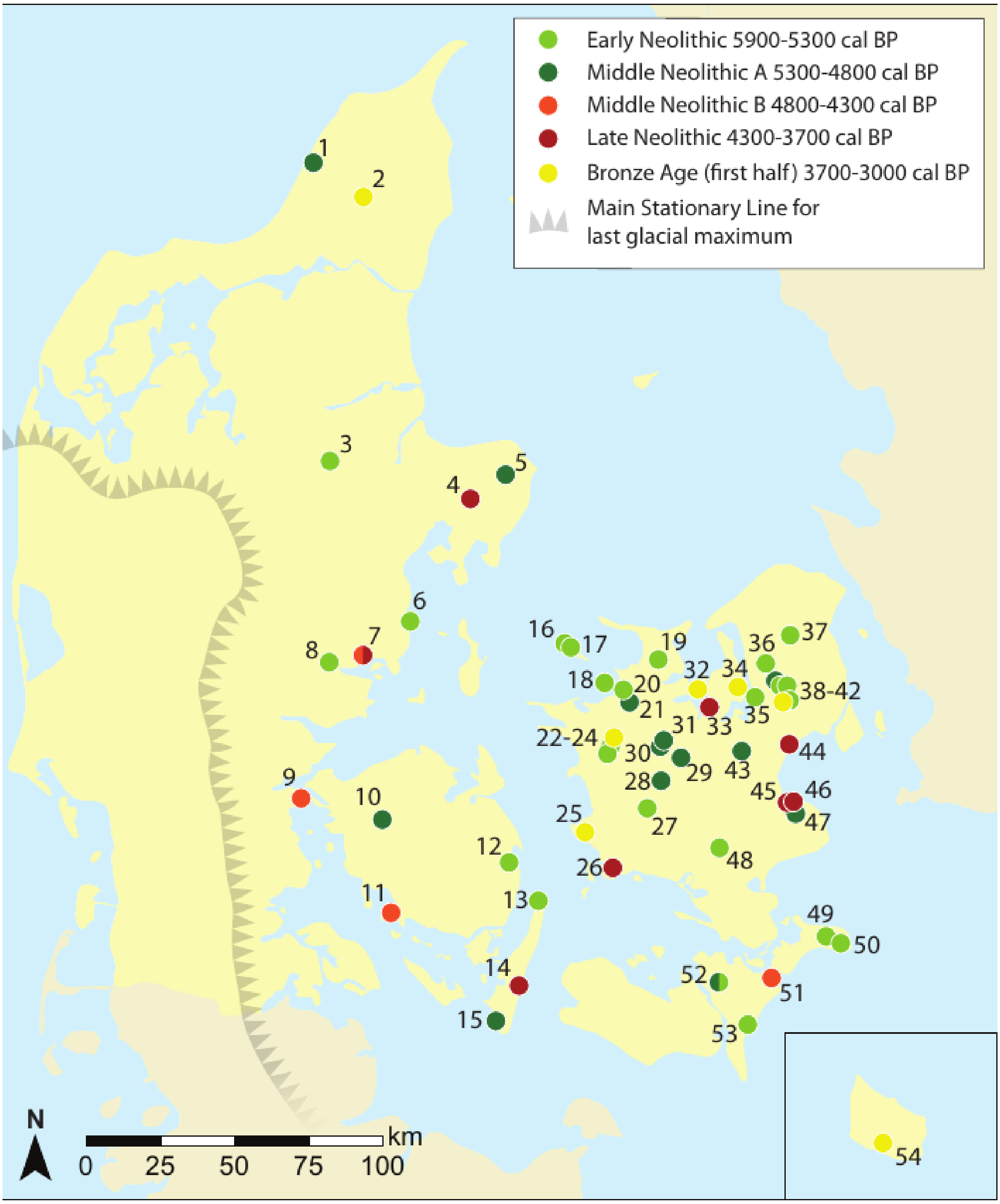


***Figure S6.2. Geographical distribution of the find locations for the study's 55 individuals of Neolithic date and 7 individuals dating to the parts of the Bronze Age earlier than 3000 cal BP.*** *1 Vittrup, 2 Klæstrupholm Mose, 3 Læsten Mose, 4 Kolind, 5 Kainsbakke, 6 Rude, 7 Toftum Mose, 8 Bygholm Nørremark, 9 Stenderup Hage, 10 Neverkær Mose, 11 Klokkehøj, 12 Sludegård Sømose, 13 Lohals, 14 Gammellung, 15 Myrebjerg Mose, 16 Rødhals, 17 Sejerby, 18 Pandebjerg, 19 Vig Femhøve, 20 Dragsholm, 21 Svinninge Vejle, 22 Madesø, 23 Jorløse Mose, 24 Tissøe, 25 Magleø, 26 Borreby, 27 Grøfte, 28 Døjringe, 29 Vanløse Mose, 30 Storelyng Eel Picker, 31 Storelyng Fire Lighter, 32 Bybjerg, 33 Kyndeløse, 34 Lollikhuse, 35 Roskilde Fjord, 36 Jørlundegård, 37*

*Salpetermosen, 38 Sigersdal Mose, 39 Viksø Mose, 40 Sigersdal, 41 Hove Å, 42 Tysmose, 43 Vibygårds Mose, 44 Mosede Mose, 45 Barhøj, 46 Strøby Ladeplads, 47 Avlebjerg, 48 Porsmose, 49 Dalmosegård, 50 Mandemarke, 51 Næs, 52 Lundby-Falster, 53 Elkenøre, 54 Vasagård.*

#### The structure of the catalogue

In the following the sites, from which skeletal material has been sampled, are presented in alphabetical order. Emphasis is put upon topography, chronology, research history and human skeletal remains represented. The headings below begin with the site name – with alternative site names in brackets. Following that comes our project's internal ID number(s) for humans from the site in question, represented in the main text. Then comes the name of the parish in which the site is located and an ID number referring to the national database for archaeological sites, *Fund & Fortidsminder*, housed by the National Agency for Culture and Palaces. Next, reference is made to the major geographic region of the find. Finally comes a brief, generalised culture-historical classification of the site. $^{14}$C dates are given as conventional radiocarbon years Before Present (i.e., uncalibrated $^{14}$C years before 1950 AD). Approaches for the reservoir effect correction of dates are detailed in Supplementary Note 4, while calibrated and reservoir corrected ages of each individual are listed in Supplementary Data III. The full assemblage of dates and isotopic values is presented in Supplementary Data IV. We also provide reference to the specific publications detailing the history of research at each site, listed at the end of each site description under 'literature'.

#### Acknowledgements

During the years-long process of selection and sampling of human skeletal materials for the study, many persons working within archaeology, etc. as researchers, museum employees and citizen scientists have assisted importantly in the acquisition of samples, provenance data, etc. A special thank is given to the following (listed alphabetically), who have been involved in our work with the assemblage dealt with in the present paper: Arne Hedegaard Andersen, Pia Bennike, Sophie Bergerbrant, Kirsten Christensen, Kristian Murphy Gregersen, Vaughan Grimes, Erik Johansen, Ole Thirup Kastholm, Kurt Kjaer, Tage Lotz, Erni Lundberg, Jesper Olsen, Knud Rosenlund, and Hugo Hvid Sørensen.

In addition, we acknowledge those, who have been of assistance in the sampling process or the gathering of provenance data on prehistoric human remains that transpired to have insufficiently well-preserved DNA (although the remains have provided other kinds of interesting data via AMS dating and/or measuring of C, N or Sr isotopes). Among these persons we especially thank the following (site names in brackets): Hans Dahl (Tybrind Vig), Inge Bødker Enghoff (Østenkær), Anne Birgitte Gurlev (Vedbæk Havn), Lars Holten (Aldersro), Niels Nørkjær Johannsen (Stistrup Kær), Lilian Matthes (Knudsgrund/Knudshoved), Klavs Randsborg (Nivå).

Furthermore, we express our sincere thanks to the many members of the research society who have been helpful in evaluating results, etc. This not least applies to Rikke Maring and Marcello Mannino (Aarhus University) and Paula Reimer and Michelle Thompson (Queen's University Belfast).

#### Avlebjerg, NEO961; Strøby 05.06.12-8A, Zealand. Stone cist

*Rune Iversen*

Aulebjerg is situated on a hill above a pronounced river valley. In 1937 human remains were exposed during gravel digging, after which an archaeologist from the National Museum of Denmark arranged a swift excavation of a NE-SW oriented, irregular stone cist. It was 2.6 m long and 0.85 m wide and contained the skeletal remains of two children placed in opposed crouched positions facing south. The burial goods included tooth and amber beads, two bone tubes, a boar tusk and two boar-tusk ornaments. Except for two amber beads, all the burial goods were associated with skeleton I. In 1939 another grave was excavated 12-15 m NE of the previous one. It was covered by a stone heap, measured 2.2 by 0.5 m and contained the skeletal remains of a c. one-metre-long child (individual III) placed with the head toward the E. The grave contained no burial goods.

DNA samples were taken from the individuals I and III. However, sufficient endogenous DNA was only preserved in skeleton I. A radiocarbon date of this individual gave the result 4510±32 $^{14}$C years BP (UBA-40443) corresponding to the early Middle Neolithic TRB (MN I).

Literature: Becker 1951[1]; Iversen 2015[2].

#### Barhøj, NEO92; Strøby 05.06.12-22, Zealand. Stone cist

*Kristoffer Buck Pedersen*

Barhøj belongs to a cluster of burial mounds that were erected about half a kilometre from the present-day and contemporaneous coast. This tumulus was situated 10 m above sea-level on a flat piece of cultivated land. Originally it was surrounded by a circle of large stones. These were removed in 1855-1860, and in 1900 the mound was reported over-ploughed. Anyhow, an excavation in 1932-33 revealed several burials, including one measuring 3.1 by 1.4 m, made of large stone slabs. A tooth from an adult human was sampled for DNA, and proved to have sufficient endogenous DNA content. It dates 3803±31 $^{14}$C BP (UB-37878), which according to the traditional south Scandinavian chronology says Late Neolithic. Fragments of at least two ceramic vessels are typologically dated to the same period.

#### Bjørnsholm, NEO751; Ranum 12.07.10-20, North Jutland. Shell midden with inhumation

*Anders Fischer*

The kitchen midden of Bjørnsholm is one of the largest of its kind known from Denmark. The extent of marine mollusk accumulations along the former sea shore is c. 325 m, while width and thickness are reported to be up to 50 m and 1.2 m, respectively. The shell layers were subject to archaeological investigation in 1931 and 1985-1991 and are well-known in literature, among other reasons, due to their stratigraphically well separated assemblages of flint, pottery and faunal materials dating from the Late Mesolithic and Early Neolithic. It was, however, only apparent through the present project that this site has also provided one of the first intact Mesolithic burials revealed during professional archaeological fieldwork in Denmark. Dated 5792±41 ($\delta^{13}C$ -13.5, $\delta^{15}N$ 14.1, UB-35718) this inhumation without burial goods can be referred to the last half of the Ertebølle Culture [ex. 3].

Literature: e.g. Andersen 1993[4] (with reference to earlier archaeological presentations); Bratlund 1993[5]; Enghoff 1993[6].

#### Bodal K (Knoglebo), NEO814; Stenlille 03.03.09-176, Zealand. Settlement and unassociated human remains

*Anders Fischer*

A rich assemblage of habitation debris derives from ploughed-up wetland sediments over an area >20 m in diameter. It is of transitional Mesolithic-Neolithic typology. Several bones of dog, domesticated cattle and a human tibia are AMS dated to the same narrow time slice. Additionally, there are two unassociated human skeletal remains with a significantly earlier AMS date, even when correcting for marine reservoir effect: a fragmented humerus and a molar (6+). The latter, which has produced a DNA sample of acceptable quality, is dated 6435±44 $^{14}C$ years BP ($\delta^{13}C$ -11.6, $\delta^{15}N$ 14.5, UB-38238).

Literature: Fischer & Gotfredsen 2006[7]; Fischer et al. 2007[8].

#### Borreby, NEO735+737; Magleby 04.04.11-45, Zealand. Passage grave

*Anders Fischer, Morten E. Allentoft and Martin Sikora*

Originally the Borreby passage grave was covered in a tumulus framed by up to 1¼ m high stones that formed a semicircle 16-18 m in diameter. The oval burial chamber, horizontal dimensions c. 5½ by 2 m, was constructed of eleven upright stones and 3 cap stones. The ESE facing, 5½ m long entrance was constructed from 5 pairs of vertical stones and a pair of door jambs, topped by capstones (Fig. S6.3). When excavated in 1859 the chamber and entrance was full to the roof with human skeletal parts, in between which were scattered burial gifts of Middle Neolithic and Late Neolithic date. The excavator judged there were remains of at least 60-70 individuals. The Meso-Neo project sampled three of them, characterised by unusually coarse facial characteristics ('Borreby type' according to previous literature) and suspected to represent foreign ancestry (cf. text on the Madesø individual below). The two of these skulls that produced DNA of acceptable quality were AMS dated to the Late Neolithic period.

Our genetic analyses have indeed documented shared ancestry for these two individuals, and for our third Borreby type individual, from Madesø, since they are all of Y-chromosomal haplotype R1b, falling within the earlier cluster of Scandinavian Late Neolithic and Bronze Age individuals. Interestingly the non-Scandinavian individuals of this cluster are generally from Western Europe, and the R1b haplogroups are also more common there. If this points to migration it would likely be from there (cf. main text's chapter: *Fine-scale structure and multiproxy analysis of Danish transect*).

Literature: Bröste et al. 1956, p. 320[9]; Ebbesen 2008, p. 125[10]; Bennike & Alexandersen 2002, p. 297[11]; Hansen 1993, p. 115[12].

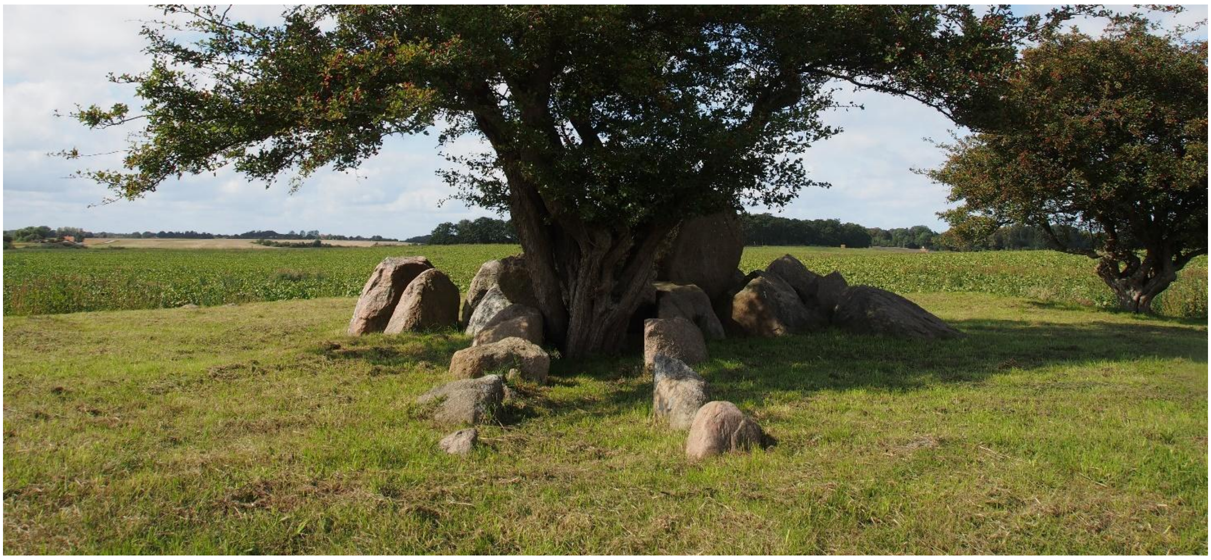

***Figure S6.3. The Borreby passage grave in its present-day appearance, where the surrounding tumulus and most of the cap stones are missing.*** *The passage leading to the burial chamber is seen in the foreground. Photo: Cille Krause, ROMU 2020, courtesy the National Agency for Culture and Palaces.*

#### Bybjerg, NEO563; Orø 03.07.09-34, off Zealand. Mesolithic kitchen midden with Bronze Age burials

*Anders Fischer*

The Bybjerg coastal kitchen midden was recorded while its sediments were queried for field improvement and road fill. It consisted of oyster and snail shells in between which numerous worked flints of Mesolithic character and some stone-set fireplaces were observed. In reports of 1911 to the Danish National Museum the local vicar mentioned two human skeletons interred at the site. The one sampled by the present project, was found below the shell layer, and was framed with a rectangular setting of stones of a size a man could lift. The body had been placed on its back in a stretched out position with the head in WSW. An AMS date relates this skeleton to the Early Bronze Age (3210±32 $^{14}$C years BP, UB-38226). The same general date probably applies to the other burial since its bones were stained by verdigris.

#### Bygholm Nørremark, NEO564; Hatting 17.04.03-128, East Jutland. Earthen long barrow

*Poul Otto Nielsen*

The earthen long barrow of Bygholm Nørremark was excavated 1977-78. It was oriented approximately E-W and measured c. 60 m in length (Fig. S6.4). Based on a series of radiocarbon dates of samples from below and within the mount, it is dated to an early stage of the south Scandinavian Early Neolithic. Grave A contained badly preserved remains of a young male and grave D contained skeletons of four adult individuals (Fig. S6.5), morphologically determined to be one male and three females. At least one of these was killed by force. The long barrow was extended to a length of 75 m in the early Middle

Neolithic, when a megalithic grave chamber was constructed.

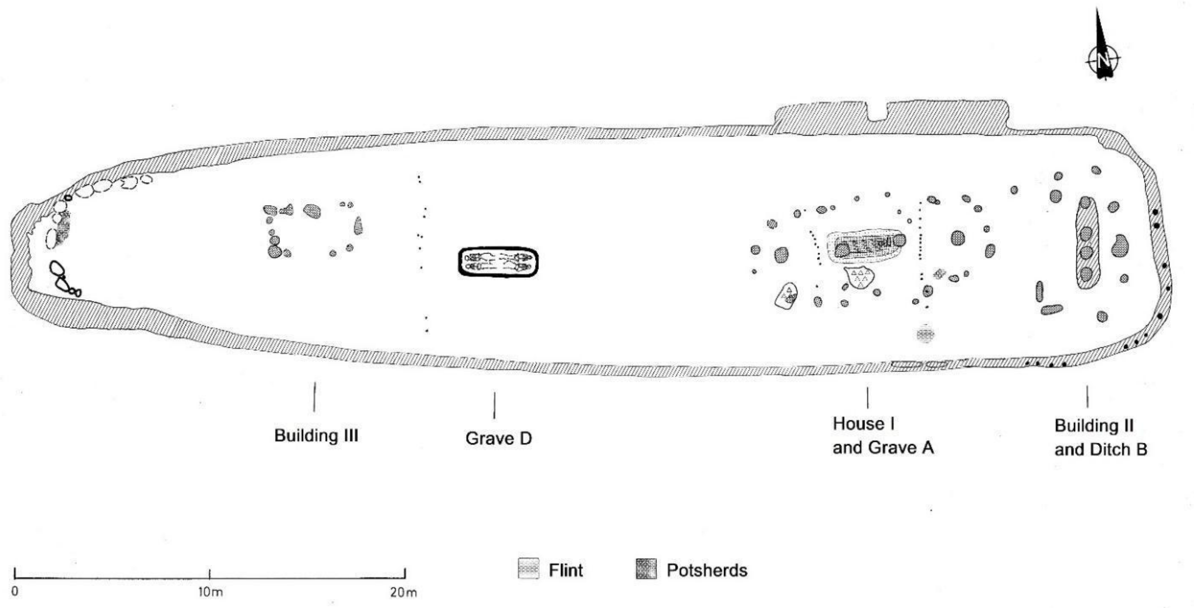


***Figure S6.4. Bygholm Nørremark, Early Neolithic stages.*** *The present genomic study deals with one of the four humans found in grave D, AMS dated 4836±35 $^{14}$C BP.*

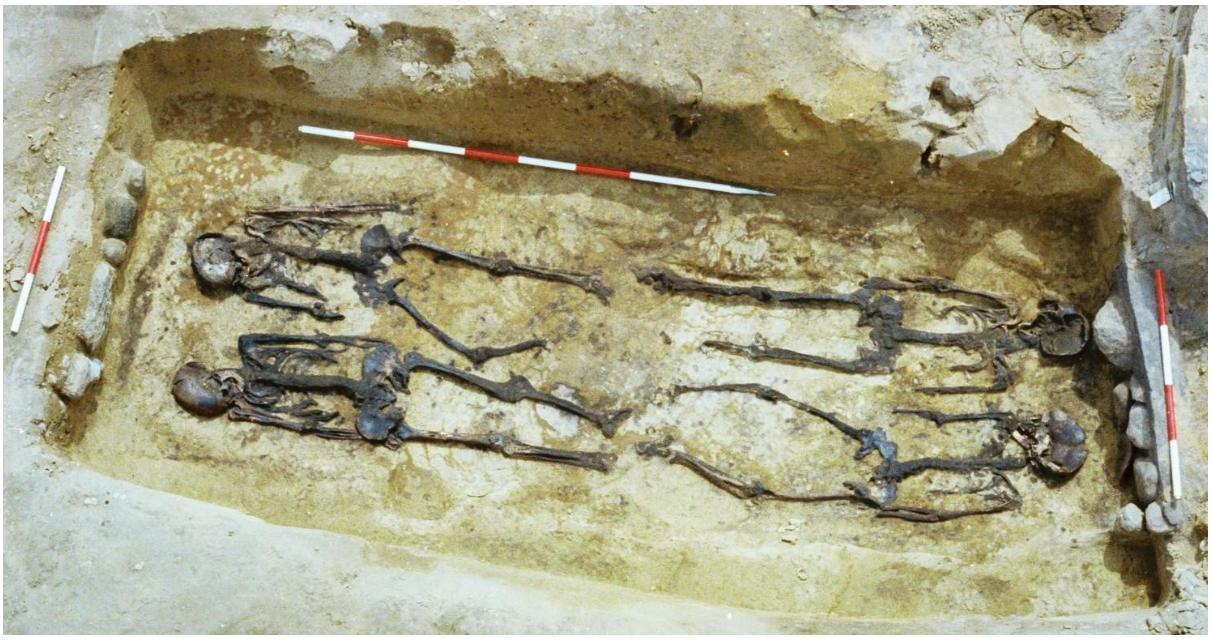

***Figure S6.5. The four humans of Bygholm Nørremark grave D, seen from the north****. It is assumed they were victims of a human sacrifice connected with the primary burial (grave A) elsewhere in the earthen barrow. Photo: Preben Rønne (Danish National Museum) 1978.*

Literature: Rønne 1978[13]; 1979[14]

#### Dalmosegaard, NEO886; Borre 05.05.02-95, Møen. Bog skeleton

*Anders Fischer*

During peat digging in a small wetland, a local farmer in 1941 revealed a highly fragmented human skull. Subsequently, a bone of domestic cattle was also salvaged from the bog deposits. Based on parallels to similar finds with better documentation, the find is inferred as the result of sacrificial deposition. Radiocarbon dates the individual to 4774±52 $^{14}C$ years BP (UB-39141) corresponding to the South Scandinavian Early Neolithic.

#### Dragsholm, NEO732+733+822+962b; Fårevejle 03.04.03-503, Zealand. Dwelling site with a Mesolithic and a Neolithic burial, and a loose human bone

*Erik Brinch Petersen*

The location is a small rise in the innermost part of the Lammefjord on the north western coast of Zealand. The former island and the drained fjord are situated just to the south of the Bay of Sejerø. A rescue excavation by the National Museum was initiated here during the early spring of 1973, and this was followed up by a second investigation in the late summer of the same year. A subsequent excavation by the Odsherred Museum and the University of Madison, USA took place between 2002 and 2004. The first two seasons in 1973 produced a single grave each, first a Late Mesolithic Ertebølle burial with ochre and then an Early Neolithic burial, while the later investigation found a single human lower jaw.

The Mesolithic burial is a double burial with two females, while the Neolithic burial contained a single male individual. The two graves were located next to each other near the top of the rise, and this close connection in space as well as in time, has created quite a stir, as well as the spilling of much ink. Both graves have been dug through a former Ertebølle shell midden, though not very deep. The Neolithic male burial is dated around 5100 $^{14}C$ years BP (AAR-7416, AAR-7416-2). The $\delta^{13}C$ value of -19.6‰ shows a non-marine diet, while the $\delta^{15}N$ measurement around 10‰ indicates a Neolithic diet. His strontium value, 0.709391, is close to that of a local red deer.

In the Mesolithic double burial was a younger individual of eighteen years of age together with an older female between forty and fifty years at death. The younger was found in a crouched position whereas the older one was in a supine position, and much ochre surrounded the two individuals. Both were well dressed, each with a girdle belt of tooth pendants from red deer teeth. Two exotic teeth, Elk and Aurochs, were found among the red deer teeth in the two belts. The younger female also carried a bone spatula decorated with a human figure.

The radiocarbon dating of the two females has been problematic, involving several laboratories. However, we have a date (prior to reservoir correction) around 6000 BP for the red deer bone dagger, the only item not affected by a marine supplement. Thus, there is at least half a millennium between the Neolithic male and the Mesolithic females. The latter ones have high $\delta^{13}C$ values, -10.7‰ for the younger one and -11.7 ‰ for the older one. Their $\delta^{15}N$ signatures are as high as 13.3 and 13.7‰.

Literature: Brinch Petersen 1974[15]; Alexandersen 1988[16], 1989[17]; Brinch Petersen & Egebjerg 2009[18]; Price et al. 2007[19].

#### Døjringe, NEO566; Munke-Bjergby 04.01.08-109, Zealand. Wetland with two bog skeletons

*Anders Fischer*

During peat digging in the year 1942 remains of two human skeletons were unearthed in the vicinity of each other, about 3 m deep in the sediment. Pollen analysis indicated a nearly identical date within the Early or Middle Neolithic. $^{14}C$ analyses later on provided nearly identical ages within the Early Neolithic. Individual I: 4629±31 and 4640±90 $^{14}C$ years BP (UB-40108 and K-3623). Individual II: 4670±90 $^{14}C$ years BP (K-3624). Both skulls have healed traces of trepanation.

Literature: Bröste et al. 1956, pp. 20-23[9]; Bennike 1985, pp. 69-72[20].

#### Elkenøre, NEO888; Idestrup 07.02.05-91, Falster. Bog skeleton

*Anders Fischer*

During peat digging a number of cranial and postcranial human skeletal remains were salvaged. Among these are cranial parts of two individuals, characterised by a dark and a pale patina, respectively. Via AMS dating, the dark one turned out to be from the Early Neolithic (4647±31 $^{14}C$ years BP; UB-40440), whereas the pale one is from the early Iron Age. Based on parallels to better documented sites, the bone assemblage is inferred to be the result of sacrificial depositions.

#### Ertebølle, NEO568+569; Strandby 12.02.12-63, North Jutland. Kitchen midden with inhumations and loose human bones

*Anders Fischer and Søren H. Andersen*

This kitchen midden has given name to the South Scandinavian Late Mesolithic Ertebølle Culture. In the seminal 1900 publication the masses of accumulated shells of marine molluscs were reported to be 1.9 m in height, c. 20 m in width and extending c. 141 metres along the contemporary beach. Excavations 1893-97 and 1979-84 have revealed a rich assemblage of worked flint, pottery, charcoals and bones of fish, birds and mammals. A well-defined stratigraphy (Figure S6.6) is observed, with massive Late Mesolithic (Ertebølle Culture) shell accumulations covered by a less shell-rich culture layer of Early Neolithic date (Funnel Beaker Culture). During the pioneering excavation human skeletal remains were found in five places within the Mesolithic deposits. They represent A) a burial with the skeleton of an adult male; B) skeletal remains of a young child, possibly representing an inhumation that was not noticed during excavation, located right next to the adult individual; C) three loose human bones. Radiocarbon dates of the two skeletons demonstrate them to be contemporary with the Mesolithic midden accumulation.

Literature: Madsen et al. 1900[21]; Andersen & Johansen 1987[22]; Enghoff 1987[23]; Petersen 1987[24]; Müller et al. 2002[25].

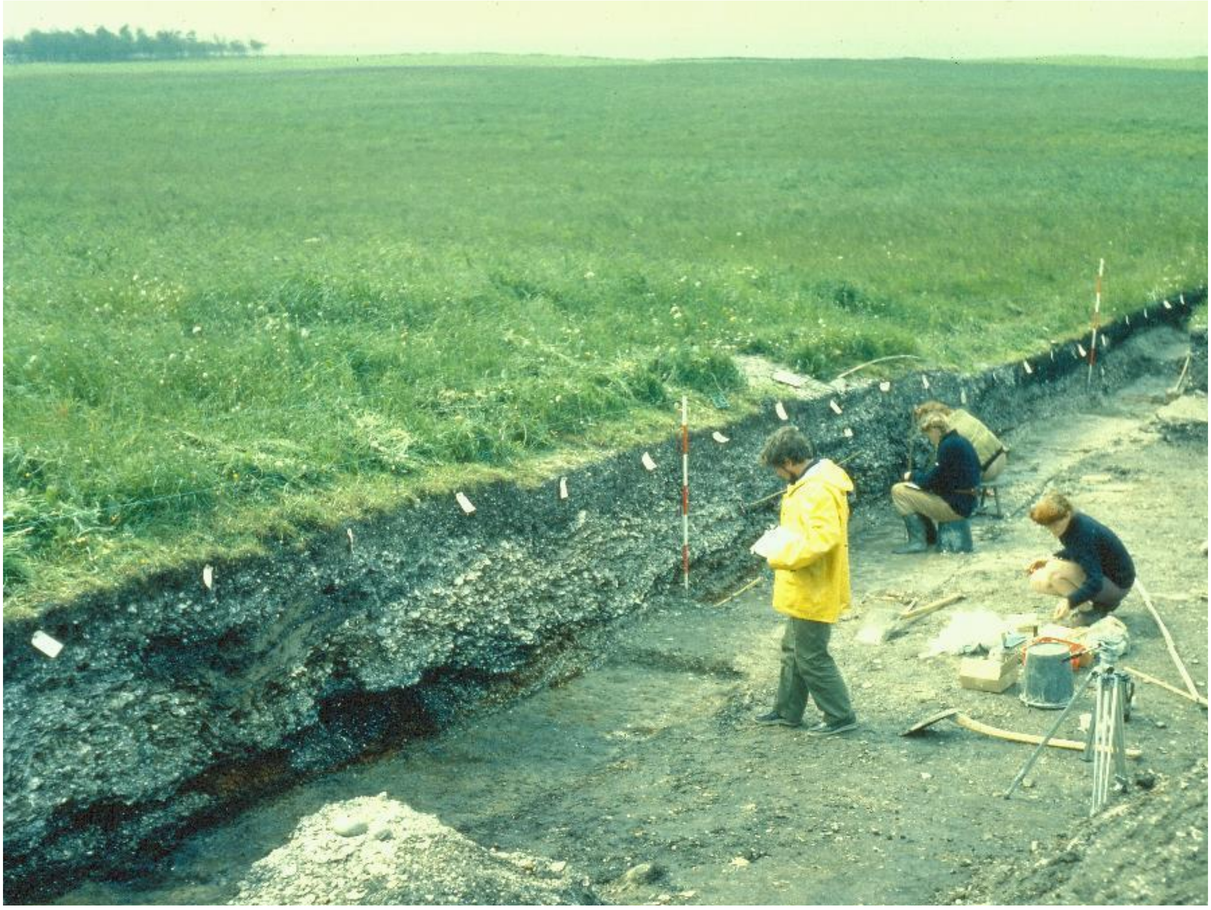

***Figure S6.6. Section through the classical Ertebølle kitchen midden.** The whitish (lower) layer, dating to the Late Mesolithic is dominated by oyster shells, while the black-grey top layer of Early Neolithic date is characterised by shells of cockles. To the left in the section is a large pit with several layers of ash. Photo 1983 Jan Sloth-Carlsen. Courtesy of Søren H. Andersen, Moesgaard Museum.*

#### Fannerup D, NEO855; Ginnerup 14.01.05-71, East Jutland. Kitchen midden with inhumation

*Rikke Maring and Esben Kannegaard*

Ruins of a coastal shell midden, excavated 1992-94. Its deposits of marine mollusc shells were accumulated to a height of as much as 0.7 m and contained animal and fish bones, flint tools, ceramics, fish hooks etc. mainly dating to the latest centuries of the Mesolithic (late Ertebølle Culture). An inhumation burial was revealed in the uppermost shell layers. The site belongs to a cluster of twelve kitchen middens along the shores of the now drained salt water sound Kolindsund.

Literature: Kannegaard 1994[26].

#### Fannerup E, NEO570; Ginnerup 14.01.05-72, East Jutland. Kitchen midden with inhumation

*Anders Fischer*

A Mesolithic coastal kitchen midden, where clearly stratified deposits of shell and sand were accumulated to a height of at least 0.7 m. The shells were dominated by oyster, but also included snails [periwinkle], cockle and common mussel. The width of the midden was about 10 m and its length parallel to the coeval shoreline >31m. Archaeological examination in 1888 led to the revealing of an inhumation burial in the uppermost shell layer. A stone paved hearth covered in sod was observed at a stratigraphically lower level. Via AMS dating (5911±43 $^{14}$C years BP, UB-35705) the interred, 40-50-year-old individual can be referred to the Late Mesolithic Ertebølle Culture.

#### Fannerup F, NEO930; Ginnerup 14.01.05-107, East Jutland. Kitchen midden with inhumation

*Lutz Klassen*

Kitchen midden with a partly disturbed grave dug down into the shell deposits. The deceased was placed in a stretched-out position on its back, and was entirely strewn with red ochre. By means of physical anthropology the well-preserved skeleton is identified as a c. 40-year-old male. It is dated 6377±30 $^{14}$C years BP (AAR-19687). The site is part of a large complex of Mesolithic and Neolithic kitchen middens on a peninsula located at the mouth of the Ørum Å river into the contemporaneous fjord of Kolindsund.

Literature: Rasmussen 1990[27]; Bennike & Alexandersen 1990[28]; Maring & Riede 2019[29].

#### Gammellung (Troldebjerg), NEO934; Lindelse 09.03.04-98, Langeland. Ceremonial bog deposition

*Otto Uldum*

Stray find of a human skull and mandible from a bog extending c. 500 by 300 m. The skull appears to show damages caused by a fatal blow. The find spot is located next to the pronounced hill, Troldebjerg, on which an enclosure site of Middle Neolithic date has been excavated. However, a $^{14}$C date produced via the present project tells that the skull belongs to the south Scandinavian Late Neolithic period: 3573±39 $^{14}$C years BP (UB-39146).

Literature: Skaarup et al. 1985, pp. 71-72[30].

#### Grøfte, NEO571; Kindertofte 04.03.06-3, Zealand. Megalithic monument with two dolmen chambers

*Anders Fischer*

The archaeological site of Grøfte is situated on a low elevation, surrounded by wetlands. Across the small rise in the moraine landscape three long dolmens are located nearly in line. Two small dolmen cists were excavated here in 1946. They were found below an earthen barrow, which measured 9 by at least 80 m and had originally been framed by megalithic stones. At least three individuals were buried in the chambers, furnished with ceramic vessels of Early Neolithic type (Figure S6.7). This date is now confirmed through AMS dating. Expressed in $^{14}C$ years BP, the results were: 4828±35 (individual from cist A, UB-38228) and 4731±32 (individual from cist B, UB-40437).

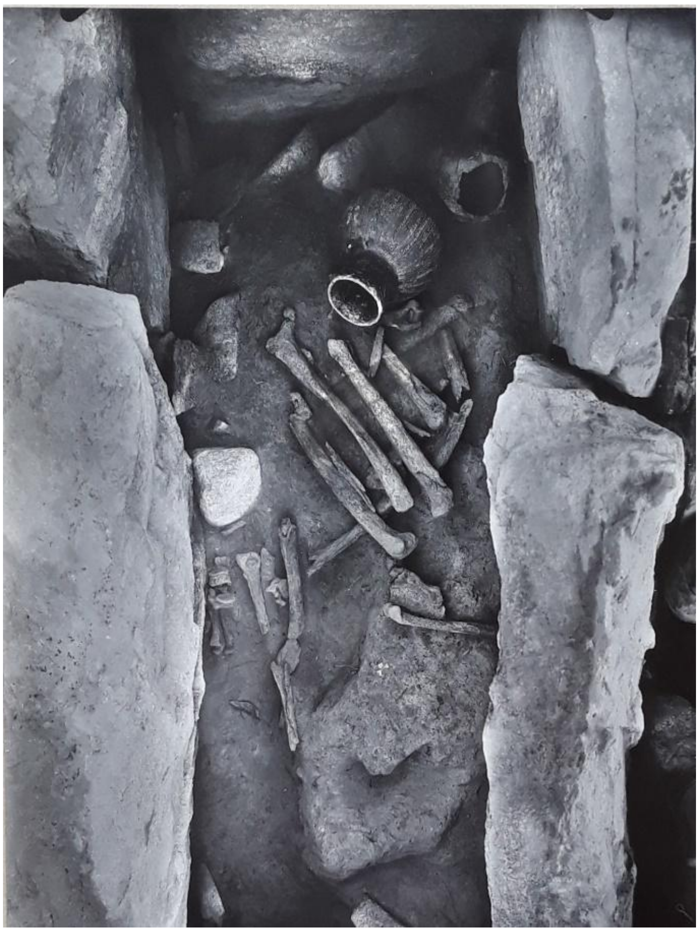

***Figure S6.7. Grøfte dolmen chamber A.*** *An intact and a fragmented lugged flask are seen upper right. Photo: Harald Andersen 1946.*

Literature: Bennike 1990[31]; Ebbesen 1990[32]; Sjögren & Fischer 2023[112].

#### Havnø, NEO941; Visborg 12.04.13-(45?), North Jutland. Human mandible from a kitchen midden

*Anders Fischer*

In 1847 a fragmented human mandible was revealed in a kitchen midden by Havnø. It was handed over to an institution, later on termed the National Museum of Denmark. Via a radiocarbon measurement (5947±33 $^{14}C$ years BP, UB-39153) the find is now dated to the Late Mesolithic Ertebølle Culture.  The site in question is probably the one that was subject to systematic excavation in 1894 and was published under the name of Havnø. The latter excavation recorded Mesolithic shell deposits extending c. 100 m along the coeval coastline and reaching a thickness of up to 0.9 m. In between the shells several human bones were found.

Literature: Madsen et al. 1900, pp. 103-111[21]; Rowley-Conwy 1983[33]; Fischer et al. 2021[34].

#### Hedegaard, NEO13; Bislev 12.05.01-113, North Jutland. Wetland with human skull

*Bjarne Henning Nielsen*

The Hedegaard calvarium is a stray find from a cleanup operation in a stream. The skull has some serious but healed traumas in the back and in the front. For many years it was believed to represent an Early Iron Age bog deposition. However, its archaic characteristics led to AMS dating (AAR-4554): 8680±40 BP, which implies Early Mesolithic and makes it one of the earliest human skeletal remains presently known from Denmark.

Literature: Fischer et al. 2007[8]; Nielsen & Adamsen 2013[35].

#### Henriksholm-Bøgebakken, NEO745+746+747+748+749; Søllerød 02.03.10-157, Zealand. Mesolithic dwelling site with burials

*Erik Brinch Petersen*

Henriksholm-Bøgebakken lies at the fossil fjord of Vedbæk, along which more than forty Mesolithic sites are recorded. The site was first excavated by the National Museum in 1924, but it was only in 1975 that the burials were detected. They include 17 inhumations with 22 individuals. This is not a cemetery, but a settlement with burials. Red deer teeth adorn the richest female, while male persons tend to be equipped with one or two flint blades. Two older individuals were laid to rest on pairs of antlers from red deer, while three males seem to have been buried under parts of dugout canoes.

The individuals dated so far, K- and Ua- dates, span the period c. 7100-6100 $^{14}C$ years BP (not corrected for marine reservoir effect). Their $\delta^{13}C$ and the $\delta^{15}N$ values centre around -15‰ and 15‰, respectively. Five skeletons have been sampled for aDNA, and they were all positive. The selected individuals were adult males from graves 5, 9, 12 and 14 and a female from grave 6/15. The archaeologically most interesting burials were not available for sampling, being presently on exhibition at Gl. Holtegård and the National Museum.

Literature: Albrethsen & Brinch Petersen 1975[36], 1977[37]; Newell et al. 1979[38]; Alexandersen 1988[16], 1989[17]; Brinch Petersen 2015[39].

#### Holmegaard-Djursland, NEO1; Hyllested 14.02.07-24, East Jutland. Kitchen midden with inhumation

*Søren H. Andersen*

A Late Mesolithic (Ertebølle Culture) kitchen midden from which a rich assemblage of worked flint, pottery, charcoal and bones of fish, birds and mammals is available. The deposit of marine shells, originally located just above the beach, measured c. 20-25 m by 15-20 m and was c. 0.5 m thick. Archaeological excavation has revealed various settlement structures, e. g. hearths of two types, large structural stones, layers of fish bones, a flint-working site and a possible dwelling floor. In 1967 an inhumation burial with a very well-preserved skeleton was found deep in the shell layer. Over the skeleton's legs and feet lay two large stones. Due to the excellent circumstances for observation it can be stated with certainty that no burial gifts were present – and no cranium either. According to physical anthropology the interred individual was a male c. 17-20 years of age. Although only a sample of limb bone was at disposal for the project, it proved possible to extract aDNA of acceptable quality. Radiocarbon dates (K-359, OxA-118, OxA-533) refer the skeleton to the middle phase of the Ertebølle Culture.

Literature: Andersen 2018[40]; cf. Newell et al. 1979[38]; Andersen et al. 1986[41]; Fischer et al. 2007[8].

#### Hove Å (Gundsømagle Mose), NEO946; Hvedstrup 02.04.06-29, Zealand. Wetland with bog skeletons

*Anders Fischer*

An ornamented spear head of Early Iron Age type and skeletal remains of two humans were salvaged during peat digging in the 1940s in mire deposits along the Hove River. The individual included in the present study dates to the Bronze Age (2982±44 $^{14}C$ BP, UB-39154), whereas the other one has previously been dated to the Neolithic.

Literature: Frei et al. 2019[42].

#### Jorløse Mose (Jordløse Mose), NEO23; Jorløse 03.06.06-192, Zealand. Wetland with bog skeleton

*Lisbeth Pedersen*

The bog of Jorløse is part of Minor Aamose (in Danish 'Lille Åmose') - a c. 6 $km^2$ mire with an outstanding concentration of Neolithic sacrificial depositions of pottery and skeletons of humans and cattle. The find in question consists of a stray found cranium, revealed during peat cutting in 1943. According to physical anthropological determination it represents a ≥ 40-50-year-old, probably male individual. Via AMS measurements 4877±32 $^{14}C$ years BP (UB-39120) and 4706±40 $^{14}C$ years BP (AAR-11122) it is dated to the South Scandinavian Early Neolithic.

Literature: Fischer & Pedersen 2005[43]; Fischer et al. 2007[8].

#### Jørlundegård, NEO702; Jørlunde 01.03.06-85, Zealand. Wetland with bog skeleton

*Anders Fischer*

During peat cutting, several Neolithic to Wiking Age wetland depositions have been salvaged from a c. 40 ha large mire, located in an undulating moraine landscape. Among these are skeletal remains of several humans. One of these was included in the present study. It represents a 2-6 years-old child, dated to the Early Neolithic (4619±41 $^{14}C$ years BP, UB-35714).

#### Kainsbakke, NEO25; Ginnerup 14.01.05-118, East Jutland. Ritual pit at coastal settlement

*Lutz Klassen*

Kainsbakke is the largest settlement of the Pitted Ware Culture known in Denmark and one of the largest Neolithic settlements of this country. It is located on a low elevation, c. 750 x 500 m in size, that was surrounded by a branch of the fjord of Kolindsund on its southern side and by a 200-400 m wide stretch of bogs on all other sides. A number of large pits were located in a row. The most important of these was pit A47, rectangular in horizontal outline, measuring c. 5.7 by 4.5 m and 1 m in depth (Figure S6.8). It had the character of a single ditch of a causewayed enclosure and was filled with marine shells (predominantly oysters), earth, rocks and several tens of thousands of flint artefacts and debris, remains of c. 530 pottery vessels and several thousand animal bones as well a few isolated bones from two humans. The tooth investigated here is from a maxilla of an older individual, directly dated to 4464±29 $^{14}$C years BP (AAR-21424). Its stable isotope values indicate a modest intake of marine food. The pit was re-opened on several occasions over a period of several hundred years, and the maxilla may be of an earlier date than the contents of Pitted Ware cultural material.

Literature: Rasmussen 1984[44]; 1991[45]; Wincentz et al. 2020[46]; Philippsen et al. 2020[47]; Makarewicz & Pleuger 2020[48].

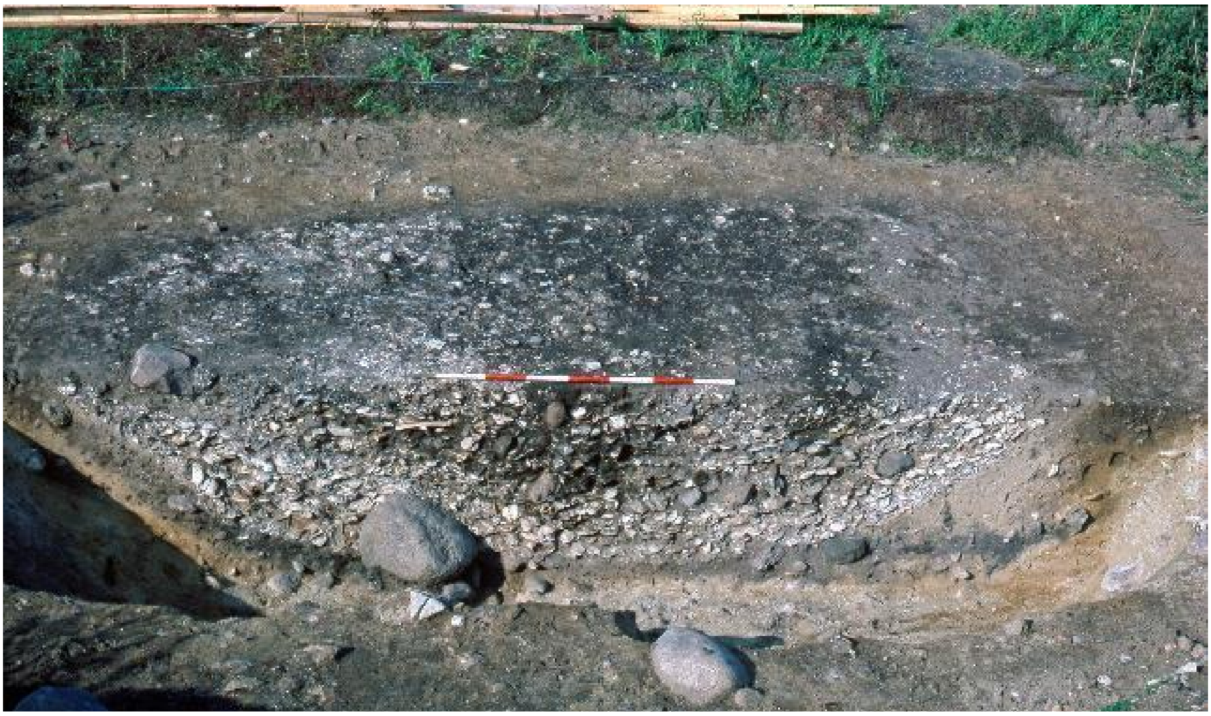

***Figure S6.8. Cross-section through pit A47 at Kainsbakke.*** *The human maxilla included in the present study was found in the massive layer of marine shells. Photo: Lisbeth Wincentz (East Jutland Museum), 1982.*

##### Klokkehøj, NEO580; Horne 09.04.12-2, Funen. Dolmen

*Anders Fischer*

Klokkehøj is a megalithic burial monument with a partly preserved dolmen chamber (Figure S6.9). During restoration work in 1977 a rich assemblage of human bones was retrieved. They represent repeated Neolithic burial activity over a period of c. a thousand years - from 4847±34 $^{14}C$ years BP (UB-37888, individual QØ) to 3883±39 $^{14}C$ years BP (UB-35706, individual RW). Of 17 humans sampled, only one produced aDNA of acceptable preservation quality. That is the individual QK, which is dated 4086±42 $^{14}C$ years BP (UB-35708), and thus belongs to a relatively early part of the epoch of the Single Grave Culture.

Literature: Thorsen 1981; Sjögren & Fischer 2023[112].

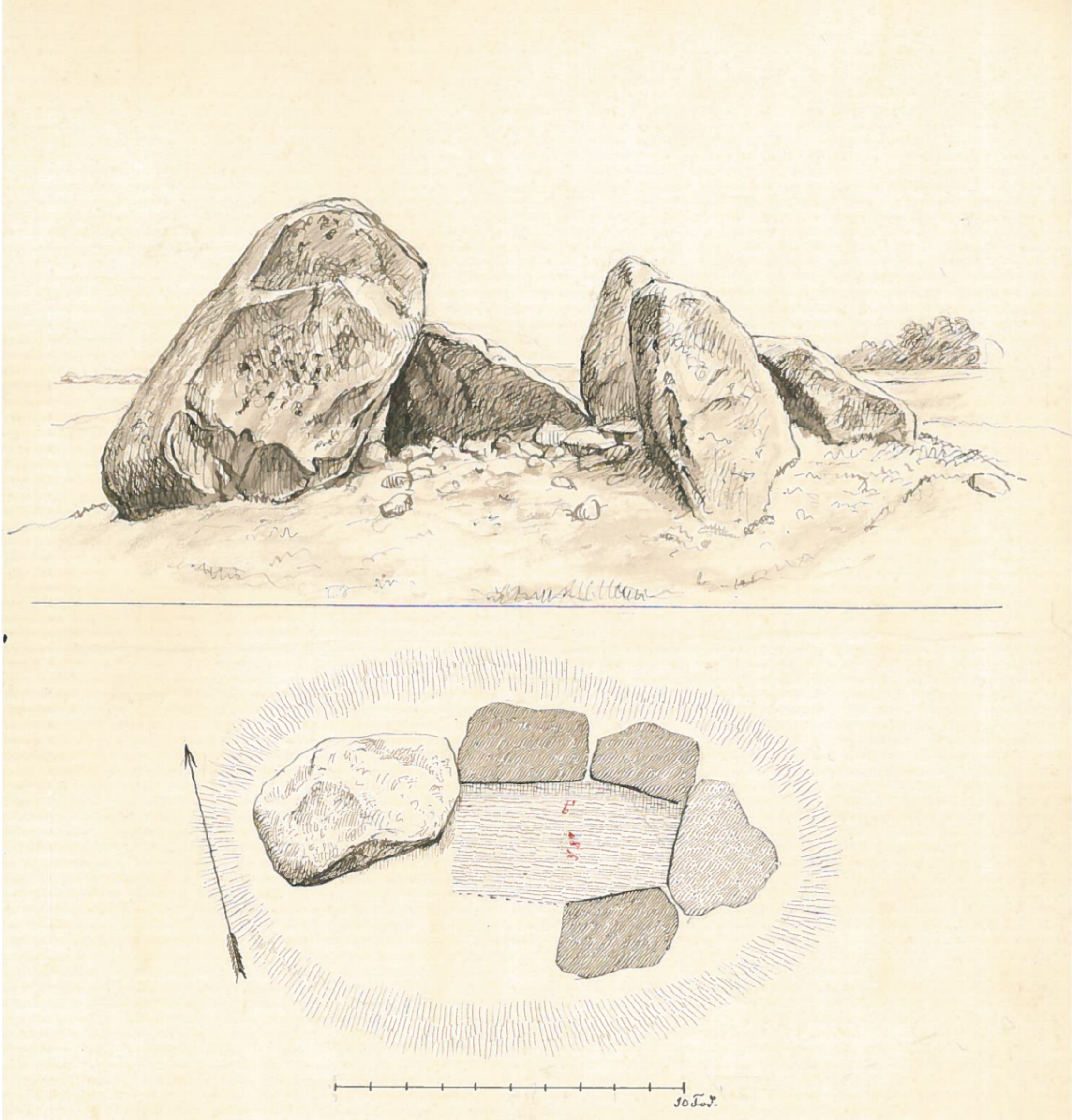

***Figure S6.9. The Klokkehøj dolmen chamber.*** *Although ruined through the activities of a stone mason, a rich assemblage of human skeletal remains, etc. was subsequently salvaged from this monument. It is located on a hill overlooking the sea. Drawing by A. P. Madsen, 1886. 1 Danish foot = 30.5 cm.*

##### Klæstrupholm Mose, NEO951; Jerslev 10.01.06-269, Vendsyssel. Wetland with bog skeleton

*Sidsel Wåhlin*

An almost complete human skeleton, unearthed by manual small-scale peat cutting in an alkaline bog situated in a small ice age river valley. Osteologically, it is determined to be an adult, probably male. There are no other prehistoric finds reported from the bog. Based on the AMS date 3114±54 $^{14}$C years BP (UB-39162) it belongs to the Early Bronze Age, according to the South Scandinavian chronology.

#### Koed, NEO583+586; Koed 14.10.07-37, East Jutland. Kitchen midden and inhumations

*Anders Fischer*

At the Late Mesolithic coastal settlement of Koed, shell-dominated culture layers made up an about one metre thick deposit. According to a report from 1891 the lower parts of the deposits were dominated by common mussel (*Mytilus edulis*) and periwinkle (*Littorina littoria*), while thin layers of oyster (*Ostrea edulis*) were observed higher up. When the site was removed for road construction in the 1940s a number of human skeletons were observed and partly salvaged from the sand below the midden deposits. Red coloration of the soil around one of the skeletons was reported, and present-day inspections have revealed ochre colour on several of the skulls. According to a physical anthropological report of 1944 there are remains of five individuals. The four of them, which have been radiocarbon dated, belong to an early part of the Ertebølle Culture.

Literature: Rasmussen 1990[27].

#### Koelbjerg, NEO254; Vissenbjerg 08.04.15-78, Funen. Wetland with bog skeleton

*Jesper Hansen*

The nearly complete skeleton of a human, estimated in the mid-20s, was found scattered in a bog near Koelbjerg during peat cutting in 1941. Until 2017 it was often referred to as the Koelbjerg Woman but is - due to new physical anthropological studies as well as aDNA analysis - determined as male, approx. 155 cm in height. In the c. 200 by 600 m large bog, two Neolithic flint axes and a Mediaeval sword have also been revealed.

The skeletal remains were preliminary dated to the Mesolithic Maglemosian Culture based on pollen analyses. Subsequently the greater part of the right-side femur was dated to 9250±85 $^{14}$C years BP (K-4063). An AMS determination from the same femur has subsequently given an only slightly older age, 9285±50 $^{14}$C years BP (AAR-8613). At present, Koelbjerg Man is the oldest human skeletal remains known from the Nordic Countries.

Literature: Bröste & Fischer-Møller 1943[49]; Troels-Smith 1943[50]; Tauber 1986[51]; Fischer et al. 2007[8]; Hansen et al. 2017[52].

#### Kolind, NEO738+739; Kolind 14.02.08-13, East Jutland. Settlement deposits with stray human bones

*Anders Fischer*

Settlement excavated during the 1920s and 1930s. Its habitation remains, accumulated to a thickness of as much as 0.9 m, were dominated by Late Mesolithic coastal kitchen midden layers, topped with deposits from the Neolithic. We have sampled a tooth and a *pars petrosa* from two individuals, aged >15 years and <10 years respectively. AMS dates showed both of them to be of Late Neolithic date (3810±40 $^{14}$C BP, AAR-35715; 3691±40 $^{14}$C BP, UB-35716).

Literature: Mathiassen et al. 1942, pp. 44-46, pp. 126-127.

#### Kongemose, NEO587; Stenmagle 04.01.12-232, Zealand. Settlement and stray human skeletal remains

*Anders Fischer*

The Middle Mesolithic epoch in southern Scandinavia is usually called Kongemose Culture after this site in Zealand's largest peat-filled lowland, the Major Aamose (in Danish 'Store Åmose'). This site, once located at an islet in a large lake, is densely littered with cultural remains of Kongemosian age, such as large rhomboid arrow points of flint. Locally it does, however, also show a far less dense scatter of artefacts of Maglemosian age, such as geometric microliths. From ploughed up parts of the site two teeth and a fragment of a human skull have been picked up by an amateur archaeologist. A physical anthropological inspection indicated they may derive from one and the same individual. This, however, does not accord with two AMS dates. If they can be relied upon, these dates refer to somewhat different parts of the Maglemose period: tooth -6, 8060±65 $^{14}$C years BP (AAR-6788) and skull fragment *os parietale*, 8331±49 $^{14}$C years BP (AAR-11350), respectively. The reason could be sample contamination from a product based on fossil oil. It is known for sure that the skull fragment was intensely treated with lacquer. Since such material could not be removed effectively through the standard sample pretreatment procedures applied those days, the AMS date of the skull fragment (and potentially also the tooth) must be suspected to be too old.

Literature: Jørgensen 1956[53].

#### Korsør Nor (Korsør Glasværk), NEO589+791; off Korsør 401433-17, Zealand. Habitation site with inhumations

*Anders Fischer*

During harbour construction in the 1940s rich traces of Mesolithic coastal habitation were revealed 2.5-4.0 m below present sea-level in the inlet of Korsør. Reports from the site indicate the presence of dug-out canoes and fish weirs. Among the material handed over to the National Museum were worked items in flint, bone and antler as well as skeletal remains of at least seven humans. Most important is an extraordinarily well-preserved inhumation burial with a middle-aged male surrounded by a packing of hazel sticks and sheets of bark. The skeletal remains tell of a sturdy, about 168 cm high individual with pronounced muscle attachments. Two healed lesions on the skull seem to result from fighting. The scattered finds of human bones from other parts of the locality may derive from burials that were disintegrated during the submergence of the locality. Well-preserved teeth from three individuals were sampled for DNA and came out with positive results.

Literature: Norling-Christensen & Bröste 1945[54]; Bennike 1997[55]; Schilling 1997[56].

#### Kyndeløse (Møllehøj), NEO878; Kirke Hyllinge 02.06.05-6, Zealand. Passage grave

*Anders Fischer*

Double passage grave in a tumulus, c. 18 m in diameter, 4 m high. The oval chambers share a wall stone. They both measure c. 3½ by 2¼ m in ground plan and have heights of 1¾ m from floor to roof. Their entrance passages are c. 4 m long. When excavated in 1938 the chamber floors were covered by several decimeters of deposits. In these numerous human bones and burial gifts of Middle Neolithic and Late Neolithic date were revealed. A left *pars petrosa* of one of the most well-preserved skulls (PMD XIII, subadult) was sampled for DNA. Additionally, a tooth (5-) from the same skull was sampled for strontium isotope measurement. Through AMS dating, this individual is referred to the Late Neolithic, 3710±35 $^{14}C$ BP (UB-39134).

Literature: Bröste et al. 1956, p. 151[9]; Ebbesen 2008, pp. 51-52[10]; Ebbesen 2009, p. 72-149[57].

#### Køge Sønakke, NEO759; 401379-38, east of Zealand. Human skeleton from the sea floor

*Anders Fischer and Kristoffer B. Pedersen*

From a submerged beach ridge complex c. 8 m below present sea level, a 'green stone' flat hoe and various human skeletal remains have been revealed during industrial extraction of gravel. When accumulated in the Early Mesolithic this beach formation was located at the entrance to a fjord connected with the Baltic Basin, the waters of which around that time were gradually changing from fresh to saline with the raising of the global sea-level. A human skeleton was brought to the surface in 1951. Morphologically it belonged to an adult, most likely a male. Two radiocarbon dates showed it to be Early Mesolithic and stable isotope measurements indicated a protein diet, which to a large extent derived from marine waters. The topographical characteristics could imply that the skeleton represents a burial at a coastal habitation site. Another human skeleton was pumped up with beach ridge materials from the site in 1943 and donated to a public museum. However, an authoritative physical anthropologist considered it of no interest and discarded it.

Literature: Fischer 1997, pp. 16-17[58]; Fischer et al. 2007[8]; Fischer & Petersen 2018[59]; Fischer & Jensen 2018[60].

#### Langø Skaldynge, NEO853; Stubberup 08.01.12-(51?), Funen. Human skull from shell midden

*Anders Fischer*

The find consists of a human skull and mandible with the rather scanty provenance data: "found in a shell midden at Langø, flint items above and below it". Based on biological anthropology, it represents an individual aged >45 years at death. Its morphology indicated a Mesolithic date, which would accord with the described context. A large kitchen midden, named Langø, has been known to scientists since the 1880s. Archaeological excavations during the first half of the 1900s had demonstrated it to date primarily from the Late Mesolithic. A *pars petrosa* was sampled for DNA analysis and AMS dating. The latter confirmed the supposed date. It is in fact one of the latest Ertebølle Culture individuals in our assemblage from Denmark: 5496±57 $^{14}C$ years BP (c. 4100 reservoir corrected years cal BC; UB-37896; $\delta^{13}C$ -10.1, $\delta^{15}N$ 14.5).

Literature: Broholm 1928[61].

#### Lohals, NEO29; Hou 09.02.02-17, Langeland. Shell midden with inhumation

*Otto Uldum*

None-megalithic grave, discovered during construction work in 1910. It contained two human skeletons in a somewhat degraded state. Via its burial goods it was referred to the south Scandinavian Early Neolithic, a date that was confirmed by AMS dates of both individuals, arranged by the present project: 4843±40 $^{14}$C years BP (UB-37900, individual in northern part of the burial) and 4799±43 $^{14}$C years BP (UB-39123, southern skeleton). Physical anthropological determination of gender suggests one to be a female of c. 154 cm body height. The other is of indeterminate gender, possibly male, with a suggested body height of 175 cm. The burial was dug into a shell midden with an original extent of c. 16 x 6 m. The shells were primarily of *Ostrea* and *Cardium*.

Literature: Hansen 1917[62]; Johansen 1917[63]; Skaarup et al. 1985, pp. 324-325[64].

#### Lollikhuse, NEO857; Selsø 01.02.06-77, Zealand. Kitchen midden with stray human remains

*Søren Anker Sørensen*

Coastal kitchen midden, the accumulation of which took place mostly during the Late Mesolithic Ertebølle Culture. The archaeological excavation of shell deposits at the site produced a few finds of settlement material from the Funnel Beaker Culture as well. A stray human tooth from the shell deposits has provided well-preserved DNA (NEO857) and is AMS dated to c. 3400 $^{14}$C years BP (Late Neolithic according to traditional Danish archaeological terminology). It may come from a destroyed burial of that period, as are known from other Danish kitchen middens.

#### Lundby-Falster, NEO865+866; Brarup 07.01.01-28, Falster. Wetland with bog skeletons

*Anders Fischer*

During peat digging in 1941 (a time of intense peat digging for fuel) two human skulls were salvaged and donated to the Anthropological Laboratory in Copenhagen. As to their context the scanty archival data only inform of the observation of animal bones. Morphologically, the skulls represent individuals aged approximately 30-35 and 18-20 years at death, respectively. Via AMS dating they can be referred to the Early Neolithic. The date of the former (individual I) is 4688±31 $^{14}C$ years BP (UB-39128), whereas the latter (individual II) is dated 4743±31 $^{14}C$ years BP (UB-40439).

##### Læsten Mose (Volstrup Mose), NEO945; Ålum 13.12.15-(71?) North Jutland. Wetland with bog skeleton

*Anders Fischer*

Parts of a human skull, found about 1½ m below surface during peat digging, probably in 1943. No further information as to context is available. Via AMS dating, the find can be related to the Early Neolithic: 4674±51 $^{14}C$ years BP (UB-39151).

##### Madesø, NEO752; Jorløse 03.06.06-53, Zealand. Wetland with bog skeleton

*Anders Fischer and Lisbeth Pedersen*

In a peat cutting area in Minor Aamose, next to Lake Madesø, a well-preserved human cranium was found c. 1.75 m below surface. Museums have received several prehistoric items from the same cutting, including two flint daggers of Late Neolithic date. They are all interpreted as the result of sacrificial deposition - cf. text on Jorløse Mose above. According to a physical anthropological inspection of 1945 (K. Bröste unpublished notes, Anthropological Laboratory) the skull represents a male c. 50 years of age. It has healed impact scars on its forehead and is characterised by unusually robust facial characteristics, including an unusually prominent supraorbital ridge (Figure S6.10; cf. comments as to a potential genetic background for these characteristics, mentioned within the above presentation of the Borreby site). Via a radiocarbon date (3523±44 $^{14}C$ years BP, AAR-8302) it can be dated to the South Scandinavian Late Neolithic.

Literature: Stensager 2003[65].

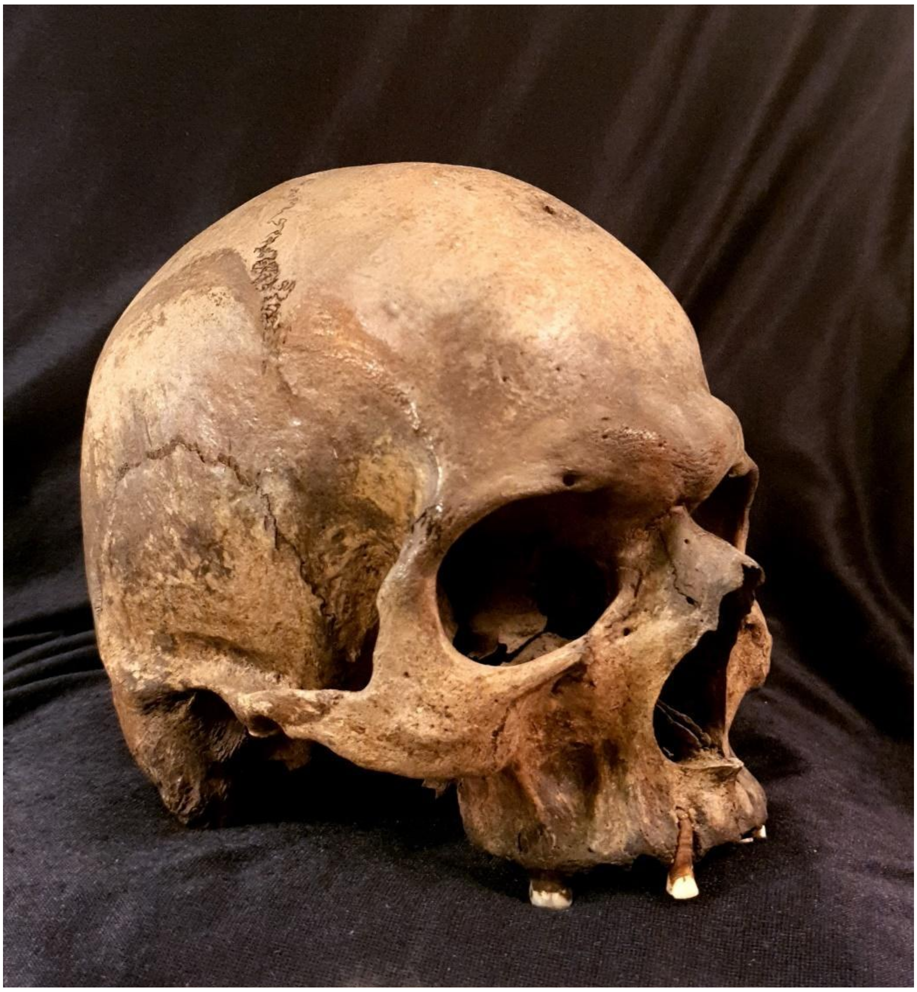

***Figure S6.10. Among the (potentially heritable) archaic traits of the Late Neolithic skull from Madesø are its very pronounced supraorbital ridge.*** *Photo: Marie Louise Jørkov, Anthropological Laboratory.*

#### Magleø, NEO590; off Korsør 401433-6, Zealand. Stray Bronze Age skeleton on Mesolithic habitation site

*Anders Fischer*

Human skeletal remains were found about 1 m below average daily sea-level, in the sea-bed of the inlet of Korsør. Traces of Mesolithic habitation had since long been recorded from the locality, and it was assumed the skeleton represented a burial of coeval age. A radiocarbon measurement, however, dates it to the Early Bronze Age: 3114±30 $^{14}C$ years BP (UB-38231).

Literature: Pedersen 1997[66].

#### Mandemarke, NEO896; Magleby 05.05.07-375, Møen. Wetland with bog skeleton

*Anders Fischer*

During peat digging in 1941 parts of a human skeleton were salvaged and donated to the Zoological Museum in Copenhagen. The donation included bones of sheep and domesticated cattle. No further information is available as to the context of the find. From the well-preserved mandible, a tooth was sampled for DNA analysis and AMS dating. The latter resulted in an Early Neolithic date: 4698±32 $^{14}$C years BP (UB-39554).

#### Mosede Mose (Karlslunde Mose), NEO860+861; Karlslunde 02.05.04-9, Zealand. Wetland with bog skeletons

*Anders Fischer*

In 1943 peat-diggers salvaged skeletal remains of three humans while working about 2 m below surface in a bog named Mosede Mose. There is no further information available on the context for these finds. Most interest had previously been given to the skull of individual I, which shows signs of trauma. The other two were available for DNA sampling and AMS dating, which resulted in Late Neolithic dates: 3568±51 (individual II, UB-37900) and 3756±29 (individual III, UB-39126) $^{14}$C years BP.

#### Myrebjerg Mose, NEO925; Magleby 09.03.06-78, Langeland. Ceremonial wetland deposition

*Otto Uldum and Anders Fischer*

The site consisted of a heap of bones, partly covered by a diffuse stone setting. It was located in a mire extending c. 200 by 100 m. The fragmented bones of at least five humans - children aged about 3 and 4 years, an adult woman and two other young adults - were intermixed with skeletal remains of domestic animals, pottery, worked flint, etc. The site is interpreted as a result of ceremonial deposition. Typologically it is – with some reservation - dated to the Funnel Beaker Culture. This chronological position seems confirmed via the $^{14}$C date 4640±320 BP (K-3711). The radiometric date should, however, be taken with reservations. This is partly due to its large standard error. Moreover, three bone samples from the present project failed to produce an absolute date for the reason of bad bone collagen preservation quality (UB-37913, UB-40132-1 and UB-40441) – a condition that may also have applied to the dated sample. From the mire there are additional finds of a celt and a belt plate of bronze, both dating to the Late Bronze Age.

Literature: Skaarup 1985, pp. 76-77[64].

#### Nederst, NEO856; Albøge 14.02.01-49, East Jutland. Settlement with shell middens and inhumations

*Esben Kannegaard*

During the Late Mesolithic the Nederst site formed an island, measuring a maximum of 400 by 900 m, in a fjord that in the historical period became known as Kolindsund. Archaeological excavations in 1988-92 revealed rich habitation deposits, including three shell middens. In the space between the middens six Early Ertebølle Culture inhumation graves were found. They were characterised by prolific strews of red ochre. Grave 1 was very disturbed by cultivation, and only the back of the skull and part of the jaw were found *in situ*. However, the skeletal remains were in excellent preservation condition, resulting in a tooth sample of acceptable DNA quality for the present project.

Literature: Kannegaard 2016[67].

#### Neverkær Mose, NEO594; Vissenbjerg 08.04.15-24, Funen. Wetland with bog skeleton

*Anders Fischer*

The bog named Neverkær measures c. 1½ by 3 km and is rich in archaeological finds. Several of these belong to the Early Neolithic period like the human skeleton dealt with here. According to biological anthropology it is a male aged 40-50 years at death, stature c. 157 cm. His well-preserved bones were revealed during peat cutting in 1944. There are two AMS dates at hand from this individual: 4548±48 (UB-37886) and 4518±33 (UB-38232) $^{14}$C years BP.

#### Norsminde, NEO852; Malling 15.04.05-71, East Jutland. Kitchen midden with burials and loose human bones

*Søren H. Andersen*

A Late Mesolithic and Early Neolithic coastal kitchen midden below which an inhumation was revealed during archaeological excavation. The grave was oriented NE-SW with the head in SW. Based on physical anthropology it represents a woman 30-40 years of age, live stature c. 153 cm. No grave goods were associated with the burial. Two radiocarbon measurements of the skeleton (K-5199, 5790±95 $^{14}C$ years BP; AAR-8556, 5800±35 $^{14}C$ years BP) indicate it belonged to the younger phase of the Ertebølle Culture. In between the Late Mesolithic and Early Neolithic food and settlement debris of the midden, some stray human bones were found. A second burial that did not produce DNA of acceptable quality is dated to the Late Neolithic.

Literature: Andersen 1991[68]; Fischer et al. 2007[8]; Andersen et al. (in prep)[111].

#### Næs, NEO792; Aastrup 07.02.14-(45?), Falster. Megalithic tomb

*Poul Otto Nielsen*

Megalithic tomb, presumably a passage grave, excavated 1838, containing Funnel Beaker pottery and flint objects from the Middle or Late Neolithic (the National Museum of Denmark, inv. no. 4630-31). Among the bones of several individuals was the skull of a man 30-40 years of age with trepanation (Figure S6.11). $^{14}C$ dated 4020±35 BP (UB-35723), it is one of the very few individuals currently known from the time of the Single Grave Culture in eastern Denmark (MN BII).

Literature: Bröste et al. 1956, pp. 426-27, no. 257[9]; Bennike 1985, p. 84[20].

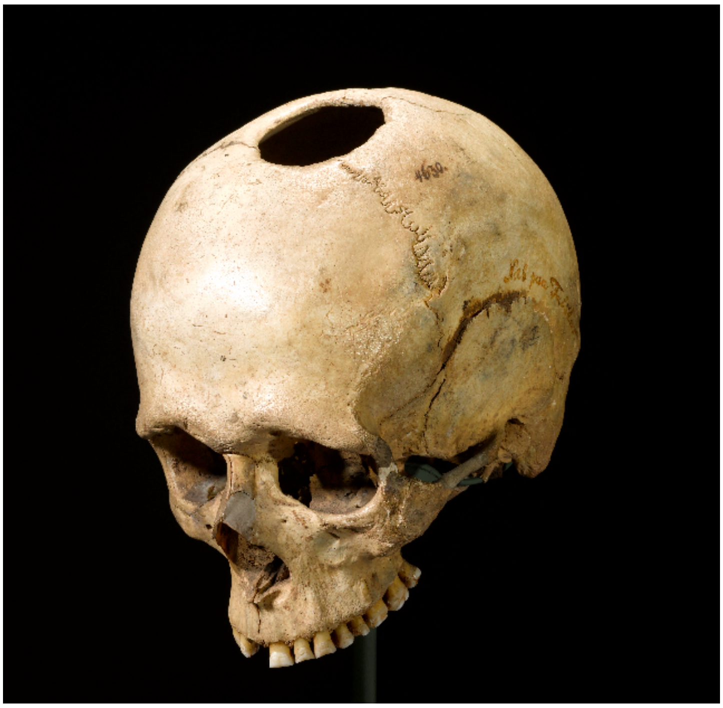

***Figure S6.11. Skull with healed trepanation from Næs, belonging to the period of the Corded Ware Culture influences and genetic impact of steppe ancestry in Denmark.*** *Photo: Roberto Fortuna / Kira Ursem (National Museum of Denmark).*

#### Orehoved Sejlrende, NEO122+123; 401651-33, south of Zealand. Submerged settlement with stray human remains

*Morten Johansen*

The submerged Mesolithic habitation site of Orehoved Sejlrende is located at a water depth of approximately 5 m. In 2015 an area of 94 $m^2$ of coastal settlement and water deposited dump materials from the Kongemose Culture was excavated. Some 21,000 pieces of worked flint and numerous organic materials including finished tools of bone, antler and wood were uncovered. The material includes three stray human teeth from deposits just off the contemporaneous coastline. Two of these have produced DNA of acceptable quality. Their dates (prior to reservoir correction) are 7505±36 (UB-37880) and 7415±53 (UB-38222) $^{14}C$ years BP, confirming their Kongemosian age.

Literature: Johansen & Ravn 2018[69]; Donahue et al. 2019[70].

#### Pandebjerg, NEO595; Føllenslev 03.06.04-76, west of Zealand. Kitchen midden with stray human bones

*Anders Fischer*

The shell midden at Pandebjerg on the small island of Nekselø was partly excavated in 1916. According to its composition of lithic artefacts it was mainly accumulated during the Late Mesolithic. Stray human bones were found in its upper level. They may be the remains of a disturbed burial. Sampling for DNA, AMS, etc. was conducted on a mandible with six extant teeth. Osteologically, it represents an individual 20-25 years of age, probably a male. Via AMS dating it is referred to the Early Neolithic. Its dietary isotopic signature ($\delta^{13}C$ -18.8, $\delta^{15}N$ 11.5) implies a higher intake of marine food than normal for Neolithic individuals from Denmark.

Literature: Brøndsted 1957, p. 167[71]; Fischer et al. 2007[72].

#### Porsmose, NEO795; Toksværd 05.04.10-15, Zealand. Wetland with bog bog skeleton

*Lasse Sørensen*

A nearly complete skeleton of a 35-40-year-old male was revealed in 1946 during excavation of peat within the Porsmose part of a c. 7½ x 2½ km mire, known as Holmegård Bog. His stature is estimated at 166 cm. Two arrows tipped with bone points had struck him, one in the upper jaw and one in the breastbone. The former had penetrated diagonally from above through the right nostril and lodged in the palate (Figure S6.12). The latter would have caused a fatal wound, as the aorta is just behind the breastbone. The skeleton is dated to the Early Neolithic: 4710±90 $^{14}C$ years BP (K-3748). Stable isotopes tell of a terrestrial diet: $\delta^{13}C$ -20.4, $\delta^{15}N$ 8.6. Nearby areas are rich in habitation deposits of Mesolithic and Early Neolithic age.

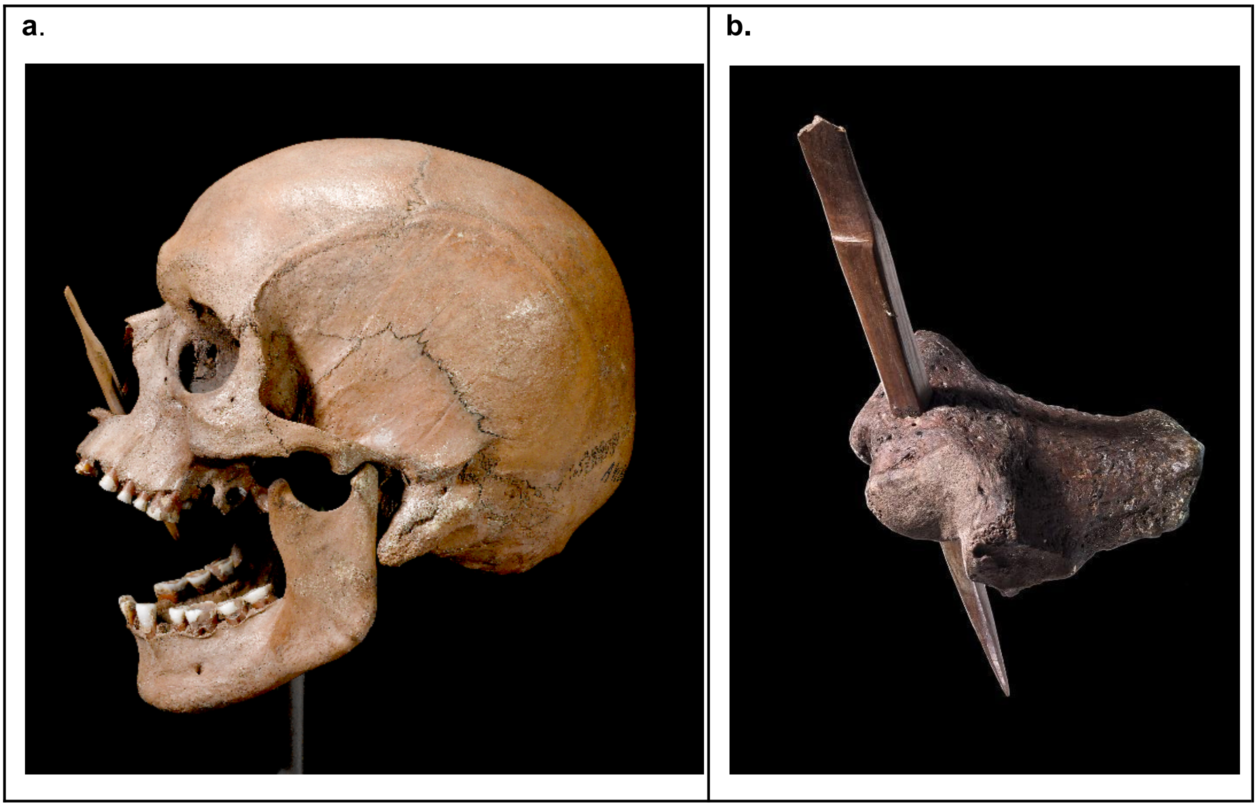


***Figure S6.12. The cause of the death of Porsmose Man****: a 10,6 cm long bone point shot at close range in the face (a) and a 12,7 cm bone point struck the breastbone (b). Photos: Roberto Fortuna and Kira Ursem (a) and Arnold Mikkelsen (b), The National Museum of Denmark.*

Literature: Becker 1952[73]; Bennike & Ebbesen 1987, p. 101[74]; Fischer et al. 2007[72].

#### Ravnsbjerggård II (Rygård), NEO960; Undløse 03.03.18-54, Zealand. Settlement site with bog deposition

*Peter Vang Petersen*

Human skull found by an amateur archaeologist's private excavation in 1955 of a mixed Late Mesolithic/Early Neolithic (late Ertebølle Culture/early Funnel Beaker Culture) settlement situated on the northern part of a small sandy island in the former lake of Major Aamose ('Store Åmose'). The find situation is unclear, but apparently the skull was found in peat layers, probably a dump zone adjacent to the shore of the island. In the peat zone were also unearthed substantial numbers of flint artefacts, axes and awls of antler and bone as well as ceramic sherds of Ertebølle and funnel beaker pottery including an Early Neolithic lugged jar. Polished flint axes and amber beads have also been found on this site.

Literature: Andersen 1983, p. 113 [75]; Koch 1998, p. 352[76].

#### Roskilde Fjord (south of Jyllinge), NEO891; 401256-78 off Zealand. Stray human bone from shell bank

*Anders Fischer*

In 1966 the industrial extraction of fossil oyster shells from the bottom of Roskilde Fjord led to the find of a human mandibula. From the same marine area skeletal remains of red deer and horse were salvaged. Knowing these finds may be of prehistoric age and could be of scientific interest, a local citizen arranged for them to be sent to the Zoological Museum in Copenhagen. Decades later it was realised that sacrificial depositions of precious items and humans frequently took place in Roskilde Fjord during the Neolithic – similarly to what took place in bogs during the same periods. Such ceremonial activity may also be the background of the specimen here dealt with. The present study arranged for its sampling for DNA analysis, AMS dating, etc. This way an Early Neolithic age is established: 4939±45 $^{14}$C years BP (UB-37910). Morphologically, it represents an individual >20 years, possibly a female.

Literature: Davidsen 1983[77]; Fischer & Petersen 2018[59].

#### Rude, NEO41+043; Saksild 15.02.12-6, East Jutland. Long barrow with two stone chambers

*Anders Fischer*

The long barrow of Rude is 8-9 m wide and at least 58 m long. Within it, two dolmen chambers are seen. They are constructed from slabs of split stone. A copper disc of Early Neolithic type derives from the western chamber. Excavation at the eastern end of the mound has demonstrated the remains of a wooden façade and the presence of votive pottery. Scattered human skeletal material is available from both cists. At least the ones from the western chamber potentially represent more than one individual, as is also indicated by differences in time and genetic signatures. According to the earliest AMS dates for DNA samples from the stone chambers these were built at an early stage of the local Early Neolithic: 4901±37 (western cist, UB-37877) and 4838±29 (eastern cist, UB-37876) $^{14}$C years BP.

Literature: Madsen 1980 [78]; Klassen 2000[79]; Sjögren & Fischer 2023[112].

#### Rødhals (Sejerø), NEO8; Sejerø 03.06.07-49, off Zealand. Kitchen midden with a burial and a stray human skull

*Anders Fischer and Anne Birgitte Gotfredsen*

Major parts of the shell midden of Rødhals were accumulated during the last decades of the Mesolithic. The site was located on a tiny island far to sea, and its well-preserved faunal remains reflect a broad-scale exploitation of marine resources including open-sea fowling, deep-water angling and hunting of small whales. For a number of years local amateur archaeologists worked their way through the midden. As a result of this digging activity, faunal material was on several occasions donated to the Zoological Museum in Copenhagen. In 1956 a simple burial was revealed in between the shells. The well-preserved skeleton (Rødhals individual I or 'Rødhals Man') with an intact skull, is included in the present study. It represents an adult male, who dates to the very transition between the Mesolithic and the Neolithic in the region. A recent study has demonstrated the presence of fragments of an additional human skull (Rødhals individual II) among the bones given to the Zoological Museum. The finds from the site also include habitation traces from initial parts of the Early Neolithic, including some of the earliest $^{14}$C dated domestic livestock presently known from Denmark.

Literature: Fischer et al. 2005[80]; Fischer et al. 2021[81]; Fischer et al. (in prep).

#### Rønsten, NEO19; 401275-1, off East Jutland. Settlement with stray human bone

*Anders Fischer*

The Middle Mesolithic coastal settlement of Rønsten is located below about two metres of marine water. Among local divers it has for decades been known as a site rich in lithic items. Occasionally organic materials have also been exposed in the sea-bed. During a period of intense sea-floor erosion the Danish authority for cultural heritage regularly inspected the site and salvaged material of scientific importance. Among these finds was a stray right mandible with three extant teeth. It has proven to have well-preserved DNA and is AMS dated to 7542±42 $^{14}C$ years BP (AAR-11355; $\delta^{13}C$ -15.0, $\delta^{15}N$ 11.8). Consequently the specimen belongs to the Middle Mesolithic epoch of the Kongemose Culture - in accordance with the typological date of the settlement assemblage.

Literature: Fischer 2007[82]; Fischer & Petersen 2018[59]; Fischer & Jensen 2018[60].

#### Salpetermose, NEO28; Hillerød 01.03.01-180A, Zealand. Wetland with bog skeleton

*Thomas Jørgensen*

A nearly complete skeleton of a 25-30-year-old male, was revealed by digging archaeological search trenches in a peat bog. His live height is estimated to be 165 cm. The cause of his death is indicated by an injury in the posterior side of the skull (Figure S6.13). All bones were found in the dug-up soil. An arrowhead of bone was found with the skeleton. No impact marks from the arrowhead were found on the human bones. Two samples from the skeleton have been AMS dated: 4789±25 (AAR-21343) and 4752±29 (AAR-21344) $^{14}C$ years BP. Nearby areas are rich in habitation deposits of Mesolithic and Early Neolithic age.

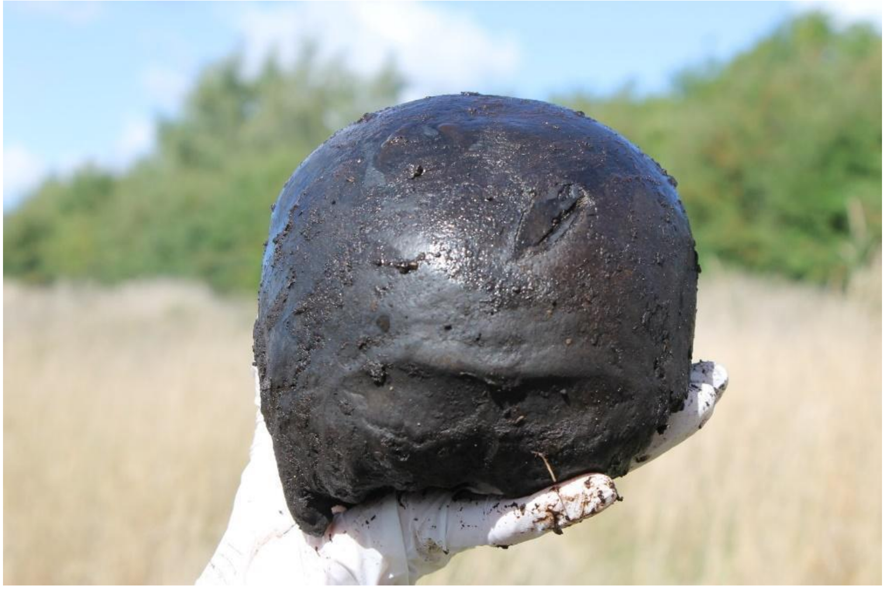

***Figure S6.13. The possible cause of death of Salpetermose Man**: a 4 cm long impact fracture, probably from a blow with a stone axe. Photo: Esben Aarsleff (Museum Nordsjælland) 2014.*

Literature: Jørgensen & Hagedorn 2015[83].

#### Sejerby, NEO757; Sejerø 03.06.07-36B, off Zealand. Burial

*Anders Fischer*

In 1927 the skeleton of an adult human was salvaged right below the bottom of the plough layer, located on even ground at the island of Sejerø. It had been buried on a 2 by ¾ m large paving of small beach-rounded stones, framed with somewhat larger stones. Lacking any kind of burial gift, the grave was – those days - considered not datable. Although of moderate bone preservation, the present study has extracted DNA of acceptable quality and has arranged for an AMS date based on a tooth. This way an Early Neolithic date is established: 4746±45 $^{14}$C years BP (UB-35721). Morphologically, the skeleton is determined juvenile or young adult, possibly female, stature c. 1.51 m.

#### Sigersdal, NEO7; Stenløse 01.06.05-110, Zealand. Wetland with bog skeletons

*Anders Fischer*

The bog of Sigersdal is located in an undulating moraine landscape where the Neolithic Funnel Beaker Culture erected monumental burials on high ground and sacrificed ceramic vessels, flint axes, domestic animals and human bodies in its watery bottom. Outstanding examples of the latter phenomena are the skeletons of two young individuals, revealed during peat digging in 1949. In literature they are termed Sigersdal individuals A and B, and have called special attention due to the fact that a cord was found around the neck of one of them (the genetically male individual A). This observation was seen as a clear indication that this 18–20-year-old did not meet death in the bog voluntarily. Both skeletons were sampled for DNA, and individual A produced data of acceptable quality. It is dated 4650±140 $^{14}$C years BP (K-3744), whereas individual B is dated 4680±75 $^{14}$C years BP (K-3745).

Literature: Bennike & Ebbesen 1987[74]; Koch 1998, find site 37[76]; Sparrevohn 2009[84].

##### Sigersdal Mose, NEO753; Stenløse 01.06.05-99, Zealand. Wetland with bog skeleton

*Anders Fischer*

The rather fragmentary remains of a human skeleton were found in the peat-infilled tunnel valley some 300 m east of the aforementioned site. They were donated to the Danish National Museum in 1947. As part of the present project an AMS analysis of the individual has been conducted, resulting in an Early Neolithic date: 4853±29 $^{14}$C years BP (UB-39125). According to biological anthropology this person was <18 years at death.

Literature: Sparrevohn 2009, Appendix 2, site 31[84].

##### Sludegaard Sømose, NEO933; Frørup 09.06.06-34A, Funen. Wetland sacrificial site with bog skeletons

*Jesper Hansen*

Prehistoric depositions of pottery, flints, wooden objects and bones  were revealed during peat cutting in a mire. The site is located in a small river valley in an undulating moraine landscape and extends c. 600 by 200 m. A pile of at least ten mandibles from wild boar has previously been dated to 5220±90 $^{14}$C years BP (K-4632). The present study includes a nearly complete human skeleton of a child app. 2 years of age. It dates to 4688±56 $^{14}$C years BP (UBA-39145). Additionally, the assemblage includes two human crania, one of which is morphologically determined male, maturus.

Literature: Albrectsen 1954[85]; Tauber 1987[86]; Noe-Nygaard & Richter 1990[87]; Ebbesen 1996[88].

#### Stenderup Hage, NEO943; South Jutland 401513c-1. Submerged settlement and stray human bones

*Anders Fischer*

Since 1946 this site has given rise to finds of prehistoric date from the beach to about 4 m below sea-level. A considerable amount of flint material is of Late Mesolithic date. The same probably applies to faunal remains, nut shells, hazel rods, etc. some of which are observed *in situ* in gyttja formations. In 1971 two fragments of a human skull were picked up from the sea-bed in about 1½ m of water. According to physical anthropology they represent a single individual of around 50-60 years, probably male. The present study has arranged for a radiocarbon analysis, 4072±61 $^{14}$C years BP (UB-39152), which places this individual at the close of the Funnel Beaker Culture or/and the beginning of the Single Grave Culture.

#### Storelyng Eel Picker (Øgaarde boat III), NEO597; Undløse 03.03.18-331, Zealand. Wetland with bog skeleton

*Anders Fischer*

The site belongs to a cluster of sacrificial Neolithic depositions in the bog of Major Aamose. It came to the light of day in connection with peat extraction for fuel in 1943. An archaeological investigation demonstrated the presence of a 7 m long log boat of alder wood, fixed to the spot with vertical rods of hazel. Right next to the vessel lay a human skeleton, inferred by the excavator to have originally been buried in the boat – an interpretation that do not conflict with radiocarbon dates of the skeleton and the boat to 4570±60 (K-3746) and 4590±120 (K-1165) $^{14}$C years BP, respectively. Morphologically, the individual is determined to be a mature male. Depositions of a second boat of alder wood, a ceramic vessel, a bone knife and a flint item within a distance of 10 m probably also represent sacrificial activity – although not necessarily absolutely contemporaneous.

Literature: Koch 1998, 311-312[76]; Richards & Koch 2001[89]; Pedersen & Fischer 2005, Fig. 5[90]; Fischer et al. 2007[72].

#### Storelyng Fire Lighter (Østrup homo II), NEO602; Undløse 03.03.18-26B, Zealand. Wetland with bog skeleton

*Anders Fischer*

Thanks to the presence of a permanently staffed archaeological field station nearby, the present skeleton is one of the exceptions, where scientists were given the chance of investigating a bog skeleton *in situ*. This took place in 1950 in relation to extraction of peat for fuel. Next to the hip of the skeleton lay what was seen as the remains of a long-gone purse holding a 'strike-a-light' consisting of pyrite, tinder fungus (*Fomes fomentarius*) and flint. Additionally, this small cluster of items included a further five flint flakes and a small awl made of bird bone. Morphologically, the individual was a male around 30 years old at death. The site is located in Major Aamose, one of the richest areas for Early Neolithic votive deposition in Northern Europe. A death in the context of sacrificial activity may also have been the fate of this individual. Its Early Neolithic date is established via radiocarbon dating: 4530±90 (K-5741) and 4523±37 (AAR-10248) $^{14}C$ years BP.

Literature: Koch 1998, pp. 350 & 353[76]; Richards & Koch 2001[89]; Fischer et al. 2007, Table 1 and p. 2143[72].

#### Strøby Grøftemark, NEO91; Strøby 05.06.12-54, Zealand. Wetland with bog skeleton

*Anders Fischer and Kristoffer B. Pedersen*

In a mire, extending c. 200 by 100 m in a gently undulating moraine landscape, parts of a human skeleton were revealed during peat cutting in 1943. Morphologically the skeletal remains were judged male and 25-35 years at death. Stratigraphic information indicated a relatively early date, which has been confirmed by two radiocarbon determinations of nearly identical results: 8215±55 (AAR-11311) and 8211±40 (UB-39958) $^{14}C$ years BP. There are no further observations of the presence of Mesolithic humans reported from the site nor from its vicinity within a distance of several hundred metres.

#### Strøby Ladeplads, NEO93; Strøby 050612-31, Zealand. Stone cist

*Kristoffer B. Pedersen*

Originally in a large mound measuring at least 25 metres in diameter and 2.5 metres in height, a stone cist was excavated. It is one of several mounds situated close to the present-day and contemporaneous coast. A small excavation in the cist was conducted by professional archaeologists in 1969 to document the circumstances of a previous private excavation.

From the cist, skeletal parts of at least 7 adults were salvaged. At least one of these was a man. Additionally, there were tibiae from a newborn child. Only one of the individuals produced DNA of acceptable quality. An AMS date of the tooth in question says 3512±33 $^{14}$C years BP (UB-37879), which implies Late Neolithic according to traditional South Scandinavian terminology.

##### Svinninge Vejle, NEO898; Svinninge 03.07.12-91, Zealand. Reclaimed fjord with stray human bone

*Anders Fischer, Lisbeth Pedersen and Anne Birgitte Gotfredsen*

The low-land of Svinninge Vejle was transgressed by the sea during an early part of the Holocene. Around 1850 a decades-long process of land reclaim began. It implied considerable efforts in digging drainage canals etc., a process that attracted the National Geological Survey as well as a locally practising doctor, who enthusiastically collected prehistoric artefacts and ancient skeletal remains to the benefit of museum collections in the nearby town of Kalundborg as well as the Zoological Museum in Copenhagen. In 1924 digging work in the former fjord-bed led to the find of a fragment of a human skull in sediments of presumed prehistoric date. This bone remained undated until the present study arranged an AMS analysis of its *pars petrosa*. It proved to be of Neolithic age: 4539±72 $^{14}$C years BP (UB-37912). According to a biological anthropological inspection the individual was an adult, probably a male.

Literature: Fischer et al. (submitted).

##### Sølager, NEO598; Torup 01.05.09-7, Zealand. Kitchen midden with stray human bones

*Anders Fischer*

Around 1850 a scientific examination of shell deposits at the Sølager site was conducted with the aim of clarifying if shell deposits observed by scientists in several locations along ancient coastlines of Denmark were the controversially early products of mankind. A human origin was confirmed. Resumed excavations in the early 1900s demonstrated a 1.7 m deep stratigraphy with unpolished flint axes and a coarse style of pottery at its lower levels, and polished flint axes and finer ceramics further up – now known as Late Mesolithic Ertebølle Culture and Early Neolithic Funnel Beaker Culture, respectively. A human mandible from a young adult with an extant tooth was available from the lowermost level. Via an AMS date arranged by the present study, its Late Mesolithic age is demonstrated: 5553±40 $^{14}C$ years BP (UB-37887). Relatively high $\delta^{13}C$ and $\delta^{15}N$ values (-10.5, 14.6) tell that marine food took up a considerable proportion of this individual's protein diet, and consequently that a comparably high reservoir effect applies to the radiocarbon date.

Literature: Skaarup 1973[91]; Newell et al. 1979, p. 70-71[38]; Fischer & Kristiansen 2002[92].

#### Tissøe, NEO942; Buerup 03.02.02-219, Zealand. Lake deposition (?)

*Anders Fischer and Lisbeth Pedersen*

Human skull from an adult female. It is provenanced 'Tissøe', which is an old-time spelling for Lake Tissø, the name of which refers to the Nordic pagan god of war, Tir. The present-day extent and depth of the lake is 12 $km^2$ and 13.5 m respectively. It is located in the Aamose-Tissø Valley, the watery landscapes of which served for numerous sacrificial depositions from the Mesolithic and well into the Christian era. Based on the archaeological record the votive activities at Lake Tissø peaked three times: during the Neolithic, the Pre-Roman Iron Age and the Viking Age to Early Medieval epochs. An AMS date 4846±53 $^{14}C$ years BP (UB-39159) showed the skull to be from the South Scandinavian Early Neolithic.

Literature: Pedersen 2004[93]; Fischer 2004[94].

#### Toftum Mose, NEO870+872+875+876; Søvind 16.05.08-17, East Jutland. Wetland with bog skeletons

*Anders Fischer and Anne Birgitte Gotfredsen*

The stores of the Anthropological Laboratory hold skeletal remains of several humans from Toftum Bog, which is a roundish peat filled basin not much more than 100 m in diameter. It is located right at the foot of a hill archaeologically known from a Funnel Beaker Culture causewayed enclosure dating to the transition from the Early to the Middle Neolithic. Our study has produced well-preserved DNA from four of the skeletons. Based on radiocarbon dates two of them can be referred to the close of the Single Grave Culture, while the other two belong to the Late Neolithic. Recordings of finds of skeletal remains from the wetland site date back to 1873. They also inform on the observation of a boat and numerous animal bones, representing wild as well as domesticated species. In addition, the archival materials tell of the finds of two tongued wedges – a type associated with the Single Grave Culture in Denmark and Corded Ware groups in Central and Eastern Europe. A pollen sample from an Aurochs skull is dated to Zone VIII. Its high frequency of *Fraxinus* in combination with presence of *Plantago lanceolata* indicates it is coeval with or later than the arrival of the Single Grave Culture (J. Troels-Smith letter of 1961).

Literature: Degerbøl & Fredskild 1970, pp. 20, 60, 199[95]; Madsen 1978[96]; 2020[97].

##### Tudse Hage, NEO932; 401441-11, west of Zealand. Submarine settlement site with stray human remains

*Per Lotz*

Stray human skeletal remains were salvaged during an archaeological sea bottom survey of an area rich in cultural material from Mesolithic coastal habitation. Besides a large collection of flint artefacts, the assemblage consists of a variety of weapons, tools and waste products in plant material, wood, bone and antler, dating to the Kongemose and Ertebølle periods. The present genetic study deals with an os temporale with a well-preserved *pars petrosa*. Samples from different parts of this bone are AMS-dated 6794±49 (UB-40929) and 6608±47 (UB-38242) $^{14}C$ years BP, respectively. The reason for the notable discrepancy in age is most likely differences in degree of sample contamination from a stabilising lacquer ('pioloform'). Since this varnish will have been based on fossil oil it is probable that the younger of the two dates (UB-38242) comes closest to the true age.

Literature: Lotz 2000[98], 2018[99]; Fischer & Pedersen 2018, pp. 78, 139, 140, 180[59].

##### Tybrind Vig, NEO683; 401521-6, west of Funen. Submerged settlement with burials

*Otto Uldum and Anders Fischer*

In an area 2-3 m below present sea level, rich remains from Mesolithic coastal habitation have been revealed. Among the archaeologically excavated deposits are two burials, both with skeletal remains of two individuals. DNA samples were taken from three of these skeletons. Only one produced genetic material of acceptable quality. It derives from an inhumation grave excavated in 2008 in about 3 m of water. Its two individuals were placed on their backs, side by side in a communal pit, without any grave goods. Physical anthropological analysis suggests the larger skeleton to be an adult female and the smaller one to be a child aged 9-12 years. Radiocarbon dates of the two individuals confirm their Mesolithic age: 6820±55 (AAR-9341) and 6905±55 (AAR-9342) $^{14}$C years BP.

Literature: Fischer et al. 2007[72]; Uldum 2011[100]; Andersen 2013[101].

##### Tysmose, NEO790; Ledøje 02.02.09-65, Zealand. Wetland with bog skeletons

*Anders Fischer and Anne Birgitte Gotfredsen*

Two complete human skeletons were revealed in a dense scatter deep in bog sediments of Tysmose. According to biological anthropology, they were c. 10 and c. 8 years at death, respectively. Pollen analysis indicated they belonged to an early stage of the Neolithic. This is now confirmed via AMS analyses arranged by the present study. Expressed in $^{14}$C years BP, the former (individual I) dates 4871±32 (UB-39159), whereas the latter (individual II) dates 4959±43 (UB-35722). In the possession of the Zoological Museum is a nearly intact skeleton of a red deer that was also salvaged from this bog in 1940.

##### Vanløse Mose, NEO599; Stenmagle 04.01.12-418, Zealand. Wetland with bog skeletons

*Anders Fischer*

Skeletal remains of two humans were found side by side in 1943 in a peat-filled basin extending c. 2.5 by 1 km along the upper reaches of the Aamose River. No other archaeological finds are recorded within a distance of 100 m. A biological anthropological inspection determined the skeletons most likely to be a female and a male. The latter, which was estimated to be <40 years at death, has been analysed for DNA with a positive outcome. According to an AMS date, this individual is Neolithic and can be attributed to the Funnel Beaker Culture. The date is 4485±41 $^{14}$C years BP (AAR-10994).

##### Vasagård, NEO815; Åker 06.02.05-58, Bornholm. Passage grave

*Poul Otto Nielsen*

Passage grave in a long barrow, whose grave chamber was excavated in 1938, resulting in finds from both the Middle Neolithic and Bronze Age (the National Museum of Denmark inv. no. A38837-40, B13208-10). The passage to the chamber (Figure S6.14) was excavated in 2008 in connection with restoration (National Museum file no. 8333/03-35). In the passage well-preserved bones of an unusually tall (male?) person were found. According to the AMS date: 3248±32 $^{14}$C years BP (UB-38239), the individual belongs to the South Scandinavian Early Bronze Age, whereby the passage to the chamber had been modified and re-used as a grave cist.

Literature: Ebbesen 1985, p. 207[102]; Hansen 2014, pp. 48-56[103].

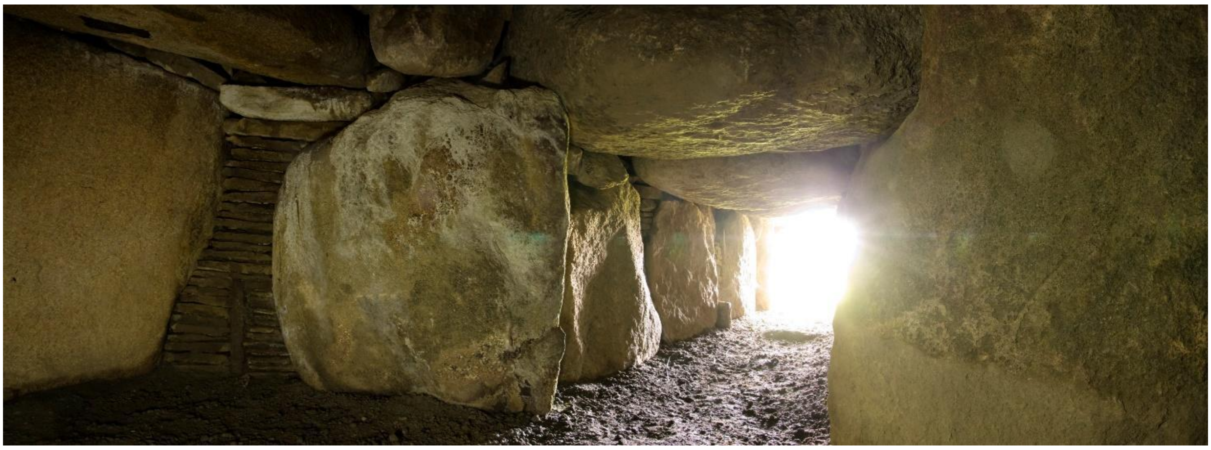

***Figure S6.14. The entrance to the Middle Neolithic Vasagård passage grave, in which an Early Bronze Age individual with well-preserved DNA was revealed.*** *Photo: Martin Stoltze, 2008.*

#### Vedbæk Boldbaner, NEO600; Søllerød 02.03.10-135, Zealand. Settlement with burials and stray human bones

*Anders Fischer*

An islet in the former Vedbæk Fjord was inhabited during several hundred years of the Kongemose period. Archaeological excavation took place in the 1940s in connection with construction works. As a result, a large assemblage of flint artefacts was retained, as were faunal remains and items in bone, antler and wood. Among the material discharged in shallow water off the habitation area were some human bones. Two proper burials were found in the sand below the culture layer: an inhumation grave and a cremation grave. The former was characterised by a 20-25 cm deep pit, horizontally extending 1.8 by 0.7 m. In this the skeleton of a mature male lay extended on its back. Based on a radiocarbon date, this individual is firmly established as of Kongemosian age: 7117±55 (Ua-23792). The present study sampled a tooth (-3), which proved to have well-preserved DNA.

Literature: Mathiassen 1946[104]; Bröste et al. 1956, p. 13[9]; Petersen 1977[105]; Newell et al. 1979, pp. 77-79[38]; Brinch Petersen 2015, p. 195[39].

##### Vibygårds Mose, NEO935; Syv 02.01.10-17, Zealand. Wetland with bog skeleton

*Per Lotz*

In a mire of c. 1700 by 300 m in a gently undulating moraine landscape, peat cutting has revealed skeletal remains of two humans and a Bronze Age celt. The present study deals with a cranium of a 35-50-year-old individual, morphologically of male character. AMS dated 4573±33 $^{14}C$ years BP (UB-39147) it can be referred to the Early Neolithic period. It has marks of a healed trepanation, possibly the earliest of its kind known from Denmark.

Literature: Nielsen 1978[106]; Bennike 1985, pp. 90-91[20].

##### Vig Femhøve, NEO744; Vig 03.04.12-150, Zealand. Megalithic monument with a dolmen chamber

*Anders Fischer*

The rectangular monument termed Femhøve ('Five mounds') (Figure S6.15) is surrounded by a frame of megalithic stones and has a dolmen cist approximately in its centre. At the opening of the monument in 1909 the floor of the 1.7 x 0.9 m large chamber was packed with disorderly located skeletal remains of as many as six individuals. Additionally, two ceramic vessels of Early Neolithic type were found in this level. Higher up, and separated by a layer of small stones, the bones of yet another individual were revealed together with burial gifts of Bronze Age date. Teeth from three individuals from the lower layer were chosen for DNA analysis and AMS dating. The latter confirmed the expectation of their Early Neolithic data. Expressed in $^{14}C$ years BP the results were: 4657±36 (individual 1, UB-37892), 4833±40 (individual 2, UB-40810) and 4709±52 (individual 3, UB-37893).

Literature: Müller 1911, pp. 282-285[107]; Sjögren & Fischer 2023[112].

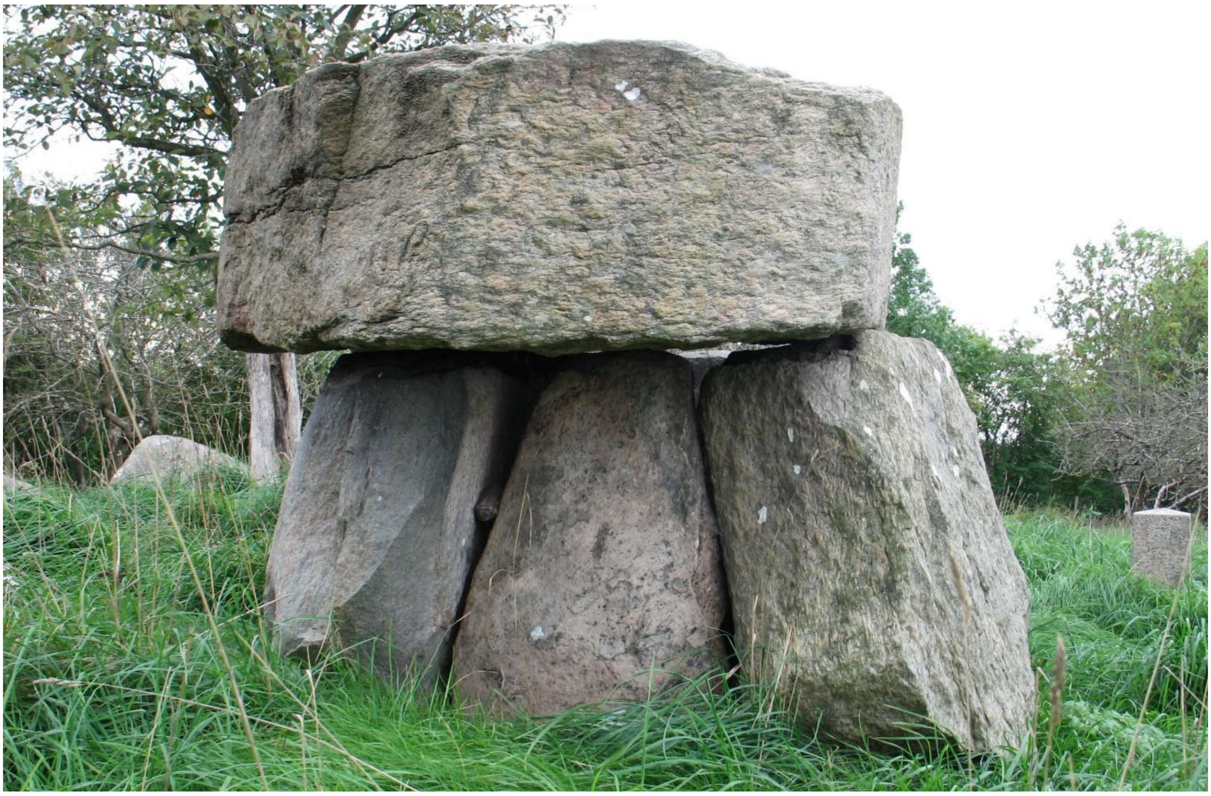

***Figure S6.15. The 1½ m high burial chamber of Vig Femhøve, in which a dense deposit of human bones was found.*** *Prior to its excavation in 1909 this 'Urdolmen' was completely covered in a 3.5 m high mound of crushed flint, a stone packing and a topping of earth. Photo: Anders Fischer 2018.*

#### Viksø Mose (Rolandsgårdens Mose), NEO601; Veksø 01.06.06-42, Zealand. Wetland with bog skeleton

*Anders Fischer*

A nearly intact human skeleton was found during peat-digging for fuel in 1946. The location is in the swampy bottom of a tunnel valley, rich in Neolithic votive depositions mainly from the Funnel Beaker Culture. Within a radius of 350 m there are recordings of a stone paving, a ceramic vessel, several flint axes, fragments of a human skull and several skulls of domestic cattle in the bog sediments. The location of the abovementioned bog skeleton of Sigersdal Mose lies round 1½ km further down-stream. The present individual represents an adult/mature female with gracile bones. Her estimated stature was c. 1.52 m. An AMS date on the tooth (3-) sampled for DNA gave the result 5050±15 $^{14}$C years BP (UCIA-232706; $\delta^{13}$C -20.9, $\delta^{15}$N 9.5), and four other radiocarbon dates cluster around that value. According to the dietary isotope values this individual had a protein diet typical of Early Neolithic farmers from Denmark. She is currently the earliest radiocarbon dated human with an Anatolian genetic ancestry known from this country.

Literature: Koch 1998, p. 277[76]; Fischer et al. 2007[72]; Sparrevohn 2009[84].

##### Vittrup, NEO33; Børglum 10.01.02-56, Vendsyssel. Wetland with bog skeleton

*Anders Fischer, Per Lysdahl and Sidsel Wåhlin*

Deposits of bone, wood and pottery were discovered in 1915 during peat cutting in the valley of a small stream. The present study deals with a skull and a mandible morphologically male and 30-40 years of biological age. The cause of death is indicated by several fractures to the skull, resulting from repeated strikes from a blunt instrument. A wooden club was found within a distance of half a metre. Additionally, a richly decorated Neolithic pottery vessel and skeletal remains of aurochs and domestic cattle were salvaged from the site. Via AMS analysis the human remains are dated to an early part of the South Scandinavian Middle Neolithic (MN A).

Literature: Friis & Lysdahl 1999 [108]; Fischer et al. (submitted).

##### Vængesø II, NEO3; Helgenæs 14.05.06-39, East Jutland. Kitchen midden with burials and loose human bones

*Søren H. Andersen*

A Late Mesolithic and Early Neolithic coastal kitchen midden in which two inhumation graves and a number of scattered human bones have been revealed. Although only a sample of limp bone was at disposal for the project, it proved possible to extract aDNA of acceptable quality from one of these individuals (skeleton BMY). Based on physical anthropology it represents a man 20-35 years of age, stature c. 1.65 m. On the left side of his frontal bone there was a healed lesion, and on the front of the eleventh thoracic vertebra there was a round lesion, probably the mark of an arrow or spear that caused death. Two large stones had been placed over the hip regions. No burial goods were associated with the grave. However, a radiocarbon measurement (5540±65 $^{14}C$ years BP, K-3921), corrected for a significant quantity of marine protein, dates him to the very latest part of the Late Mesolithic Ertebølle Culture, close to the time of the introduction of domesticated cattle and funnel beaker pottery in the region.

Literature: Andersen 2018[40]; cf. Andersen et al. 1986[41]; Meiklejohn et al. 1998[109]; Fischer et al. 2007[72].

(Aarhus University Press, Aarhus, 2020).

47. Philippsen, B., Iversen, R. & Klassen, L. The Pitted Ware culture chronology on Djursland. New evidence from Kainsbakke and other sites. in *The Pitted Ware Culture on Djursland. Supra-regional significance and contacts in the Middle Neolithic of southern Scandinavia* (ed. Klassen, L.) 257–277 (Aarhus University Press, Aarhus, 2020).
48. Makarewicz, C. & Pleuger, S. Herder-hunter-fishers and agricultural contacts. Zooarchaeological perspectives on pitted ware animal exploitation strategies from Djursland. in *The Pitted Ware Culture on Djursland. Supra-regional significance and contacts in the Middle Neolithic of southern Scandinavia* (ed. Klassen, L.) 279–339 (Aarhus University Press, Aarhus, 2020).
49. Bröste, K., Fischer-Møller, K. & Skelettet, K. Et Fund fra tidlig Maglemosetid [Le squelette de Koelbjerg. *Aarbøger for Nordisk Oldkyndighed og Historie* 211–231 – (1943).
50. Troels-Smith, J. Geologisk Datering af Koelbjerg-Skelettet [Datation géologique du squelette de Koelbjerg. *Aarbøger for Nordisk Oldkyndighed og Historie* 232–238 – (1943).
51. Tauber, H. Koelbjerg-kvindens alder og ernæring. *Fynske Minder* 28–33 (1986).
52. Hansen, J. *et al.* The Maglemosian skeleton from Koelbjerg revisited: Identifying sex and provenance. *Danish Journal of Archaeology* **6**, 55–66 (2017).
53. Jørgensen, S. Kongemosen: Endnu en Aamose-Boplads fra Ældre Stenalder. *Kuml* 23–40 (1956).
54. Norling-Christensen, H. & Bröste, K. Skeletgraven fra Korsør Nor. Et menneskefund fra ældre stenalder. *Fra Nationalmuseets Arbejdsmark* 5–17 (1945).
55. Bennike, P. *Death in the Mesolithic. Two old men from Korsør Nor.* 99–105 (A/S Storebælt Fixed Link, 1997).
56. Schilling, H. *The Korsør Nor site. The permanent dwelling place of a hunting and fishing*

*people in life and death*. 93–98 (A/S Storebælt Fixed Link, 1997).

57. Ebbesen, K. *Danske Jættestuer*. (Attica, 2009).
58. Fischer, A. People and the sea – settlement and fishing along the Mesolithic coasts. in *The Danish Storebælt Since the Ice Age – Man, Sea and Forest* (eds. Pedersen, L., Fischer, A. & Aaby, B.) 63–77 (A/S Storebælt Fixed Link, 1997).
59. Fischer, A. & Petersen, P. V. A sea of archaeological plenty. in *Oceans of Archaeology* (eds. Fischer, A. & Pedersen, L.) 68–83 (Højbjerg, 2018).
60. Fischer, A. & Jensen, T. Z. T. *Radiocarbon dates for submarine and maritime finds from early prehistory*. 202–221 (Jutland Archaeological Society, Højbjerg, 2018).
61. Broholm, H. C. Langøfundet. En boplads fra ældre Stenalder paa Fyn. *Aarbøger for Nordisk Oldkyndighed og Historie* – (1928).
62. Hansen, F. Några enmandsgraver från stenåldern. *Fornvännen* 67–89 (1917).
63. Johansen, K. F. Jordgrave fra dyssetid. *Aarbøger for Nordisk Oldkyndighed og Historie* 131–147 (1917).
64. Skaarup, J., Nyegaard, G., Aaris-Sørensen, K., Petersen, K. S. & Bennike, P. *Yngre stenalder på øerne syd for Fyn*. (Langelands Museum, 1985).
65. Stensager, A. O. Menneskeofre i Jorløse og Verup Moser, in Årets gang 2002. Beretning for Kalundborg og Omegns Museum. 50–53 (2003).
66. Pedersen, L. *When the world view turned upside down*. 122–123 (A/S Storebælt Fixed Link, 1997).
67. Kannegaard, E. Late Mesolithic ochre graves at Nederst, Denmark: ochre rituals and customs of personal adornment, in Mesolithic burials – rites, symbols and social organization of early postglacial communities. in *Landesamt für Denkmalplege und Archäologie Sachsen-Anhalt/Landesmuseum für Vorgeschichte* (eds. Grünberg, J. M., Gramsch, B., Larsson, L., Orschiedt, J. & Meller, H.) 81–93 (2016).
68. Andersen, S. H. Norsminde. *J. Dan. Archaeol.* **8**, 13–40 (1991).
69. Johansen, M. & Ravn, M. Orehoved – high-tech hand excavation under water. in

*Oceans of Archaeology* (eds. Fischer, A. & Pedersen, L.) 120–121 (Oxbow Books, 2018).

70. Donahue, R. E., Fischer, A., Burroni, D. B., Malm, T. & Johansen, M. Microwear analysis aiding excavation prioritization at the submerged Mesolithic settlement of Orehoved. *Journal of Archaeological Science: Reports* **23**, 540–548 (2019).
71. Brøndsted, J. *Danmarks oldtid: Stenalderen*. vol. 1 (Gyldendal, 1957).
72. Fischer, A. *et al.* Coast-inland mobility and diet in the Danish Mesolithic and Neolithic: evidence from stable isotope values of humans and dogs. *J. Archaeol. Sci.* **34**, 2125–2150 (2007).
73. Becker, C. J. Skeletfundet fra Porsmose ved Næstved. *Nationalmuseets Arbejdsmark* 25–30 (1952).
74. Bennike, P. & Ebbesen, K. The bog find from Sigersdal. Human sacrifice in the Early Neolithic. *Journal of danish Archaeology* **5**, 85–111 (1987).
75. Andersen, K. *Stenalderbebyggelsen i Den vestsjællandske Åmose*. (Fredningsstyrelsen, 1983).
76. Koch, E. *Neolithic Bog Pots from Zealand*. vol. 16 (The Royal Society of Northern Antiquaries, 1998).
77. Davidsen, K. Stenalderfund fra østersbanker ved Kølholm i Roskilde Fjord (English summary: Neolithic underwater votive offerings. *Antikvariske Studier* **6**, 127–136 (1983).
78. Madsen, T. En tidligneolitisk langhøj ved Rude i Østjylland. *KUML* 79–108 (1980).
79. Klassen, L. *Frühes Kupfer im Norden: Untersuchungen zu Chronologie, Herkunft und Bedeutung der Kupferfunde der Nordgruppe der Trichterbecherkultur*. vol. 36 (Aarhus Universitetsforlag, 2000).
80. Fischer, A., Bennike, B. & Heinemeier, J. Særling fra Sejerø. in *Årets gang 2004. Beretning og regnskab for Kalundborg og Omegns Museum* (ed. Pedersen, L.) 64–68 (Kalundborg og Omegns Museum, 2005).
81. Fischer, A., Gotfredsen, A. B., Meadows, J., Pedersen, L. & Stafford, M. The Rødhals

kitchen midden – marine adaptations at the end of the Mesolithic world. *Journal of Archaeological Science: Reports* **39**, 103102 (2021).

82. Fischer, A. Coastal fishing in Stone Age Denmark--evidence from below and above the present sea level and from human bones. *Shell Middens in Atlantic Europe* **30**, 54–69 (2007).
83. Jørgensen, T. & Hagedorn, L. Salpetermoseliget in Alle tiders Nordsjælland, Museum Nordsjællands Årbog 2015. 118–122 Preprint at (2015).
84. Sparrevohn, L. R. Omkring en å. Tragtbægerkulturens særlige steder. in *Plads og rum i tragtbægerkulturen. Nordiske Fortidsminder. Serie C* (ed. Schülke, A.) vol. 6 45–65 (The Royal Society of Northern Antiquaries, 2009).
85. Albrectsen, E. Et offerfund i Sludegårds mose. *Fynske Minder* 4–14 (1954).
86. Tauber, H. Danish radiocarbon dates of archaeological samples. *Arkæologiske udgravninger I Danmark 1986* 182–199 (1987).
87. Noe-Nygaard, N. & Richter, J. Seventeen Wild Boar Mandibles from Sludegårds Sømose – Offal or Sacrifice? in *Experimentation and Reconstruction in Environmental Archaeology. Symposia of the Association for Environmental Archaeology No. 9, Roskilde 23-26 September 1988* (ed. Robinson, D. E.) (Oxbow Books, 1990).
88. Ebbesen, K. Mesolitisk måltidsoffer i Sludegårds Sømose. *SDA-nyt* **12**, (1996).
89. Richards, M. P. & Koch, E. Neolitisk kost. Analyser af kvælstof-isotopen 15N i mennesker fra yngre stenalder. *Aarbøger for Nordisk Oldkyndighed og Historie* 7–17 (2001).
90. Pedersen, L. & Fischer, A. Kultur- og naturmiljøer i naturpark Åmosen-Tissø – to sider af samme sag. in *Kulturarv i Naturpark Åmosen-Tissø* (eds. Fischer, A. & Pedersen, L.) 5–16 (Naturpark Åmosen-Tissø, Kalundborg og Omegns Museum & Nationalmuseet, 2005).
91. Skaarup, J. Hesselø – Sølager. in *Jagdstationen der südskandinavischen Trichterbecherkultur* (Akademisk Forlag, 1973).

92. *The Neolithisation of Denmark: 150 years of debate*. (JR Collis Publications, 2002).
93. kulturhistorie og natur. *Kalundborg* 9–26 (2004).
94. Fischer, A. Tissø og Åmoserne som trafikforbindelse og kultsted i stenalderen. in *Tissø og Åmoserne – kulturhistorie og natur. Historisk Samfund for Holbæk Amt 2003* (ed. Pedersen, L.) 27–44 (2004).
95. Degerbøl, M. & Fredskild, B. The Urus (Bos primigenius Bojanus) and Neolithic domesticated cattle (Bos Taurus domesticus Linné) in Denmark. The Royal Danish Academy of Sciences and Letters. *Biologiske Skrifter* **17,1**, (1970).
96. Madsen, T. Toftum ved Horsens. Et 'befæstet' anlæg tilhørende tragtbægerkulturen. *Kuml* 161–184 (1978).
97. Madsen, T. Fra grænselandet mellem to kulturer: Tragtbægerkultur og enkeltgravskultur i det centrale Østjylland. *Kuml* **69**, 9–84 (2020).
98. Lotz, P. Tudsehage – an underwater settlement on Sjælland. *Maritime Archaeology Newsletter from Roskilde Denmark* **14**, 8–13 (2000).
99. Lotz, P. Stenalderredskaber af ben og hjortetak – samt status fra Tudse Hage. *Fund & Fortid –Arkæologi for alle* **4**, 4–8 (2018).
100. Uldum, O. The excavation of a Mesolithic double burial from Tybrind Vig. in *Submerged Prehistory (Oxbow* (ed. Benjamin, J.) 15–21 (2011).
101. Andersen, S. H. *Tybrind Vig: Submerged Mesolithic Settlements in Denmark*. (Jutland Archaeological Society, Højbjerg, 2013).
102. Ebbesen, K. Bornholms dysser og jættestuer. Bornholmske Samlinger II. *Rk* **18**, 175–211 (1985).
103. Hansen, S. I. Bornholms jættestuer. in *Solstensøen. På sporet af Bornholms bondestenalder (Bornholms Museum, Nationalmuseet & Wormianum* (eds. Nielsen, P. O., Nielsen, F. O. S. & Adamsen, C.) 47–80 (2014).
104. Mathiassen, T. En Boplads fra ældre Stenalder ved Vedbæk Boldbaner. *Søllerødbogen* 19–35 (1946).

### 7) Catalogue of non-Danish archaeological sites

Edited by Fabrice Demeter[1,2], Karl-Göran Sjögren[3] and Ruairidh Macleod[1,4]

[1]Lundbeck Foundation GeoGenetics Centre, Globe Institute, University of Copenhagen.

[2]National Museum of Natural History, Ecoanthropology and Ethnobiology, Musée de l'Homme, Paris.

[3]Department of Historical Studies, University of Gothenburg, 405 30 Gothenburg, Sweden

[4]GeoGenetics Group, Department of Zoology, University of Cambridge, Cambridge, UK.

The format of the following section is similar to the above catalogue of Danish archaeological sites; in each instance a description of the nature of the site and samples is provided, followed by a short list of publications on the site, providing a bibliography of relevant research relating to samples analysed here. The distribution of sampling is extensively cosmopolitan across western Eurasia (outside of the sampling focus on Denmark, see Fig. S7.1), and predominantly represent the Mesolithic and Neolithic periods, though with individual samples from the Upper Palaeolithic (Kotias Klde, Georgia) and Medieval (Shauke 1, Kazakhstan) periods, at the extremes of the temporal distribution. The former (NEO283), is directly dated to 26,052 - 25,323 cal. BP (95%) and is reported alongside a Mesolithic individual (NEO821) from the same site, while the latter appears to have been intrusive and is reported alongside genomes from a Bronze Age and an Iron Age individual from the same site. Commensurate with the geographical area covered by these countries, the largest proportion of samples outside of Denmark originate from Russia and Ukraine (Fig. S7.2). In Russia, human remains from 28 distinct archaeological localities were sampled, including the Neolithic site of Sakhtysh II (within the Sakhtysh locality), for which 13 unique ancient genomes were sequenced, the single site with most intense aDNA sampling across the whole dataset. The occurrence of relatively large prehistoric hunter-gatherer cemeteries is particularly characteristic of the Neolithic in Eastern Europe and Siberia, with notable examples comprising up to hundreds of graves at cemeteries at Lake Baikal (of the Early Neolithic *Kitoi* or *Kitoy* mortuary tradition). In Ukraine, archaeological sites where human remains have been sampled are dominated by an association with the Dnipro (alternatively Dnepr or Dneiper) river valley (in the case of seven out of eight localities). Many of these are associated with the Dnipro-Donets culture, a significant Mesolithic to Neolithic cultural complex from ca. 7000 to 6200 BP, and also known for large collective hunter-gatherer cemeteries. In western Europe, further sampling of Swedish archaeological sites contribute to the focus of ‘filling in the gap’ of understanding of the Mesolithic to Neolithic transition in Scandinavia, including novel genomes from the notable Mesolithic cemetery at Skateholm and a number of individuals at a Late Neolithic/Early Bronze Age gallery grave at Fredriksberg, Västergötland. Across the numerous other localities sampled, these represent a considerable diversity of archaeological contexts and taphonomic environments, in many cases reliant on limited documentation of excavations from the mid or early 20th Century from which to document these remains.

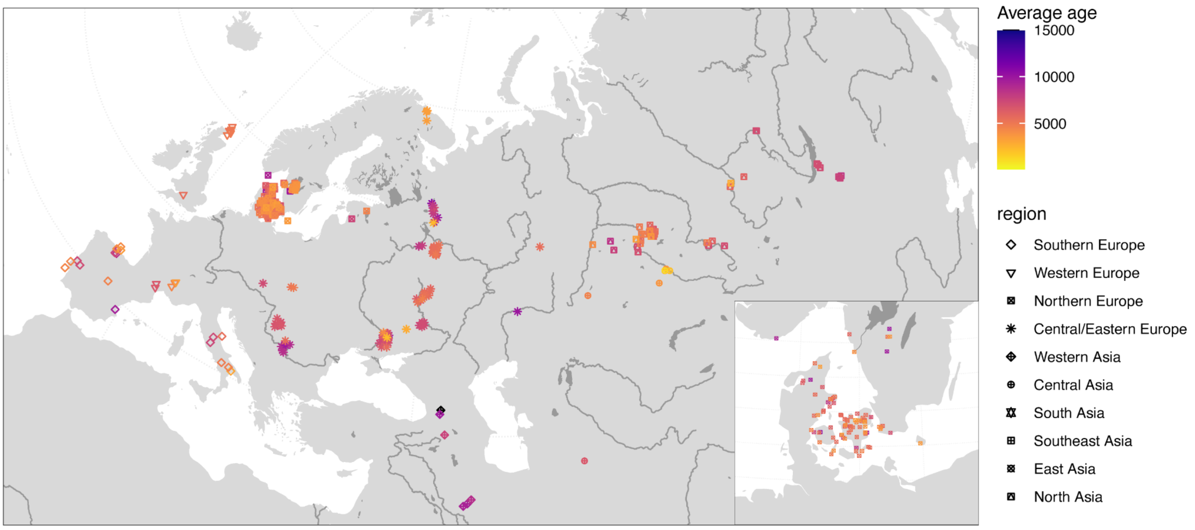


**Figure S7.1. The total distribution of sampled ancient human remains in this study.** Adapted from Fig. 1A.

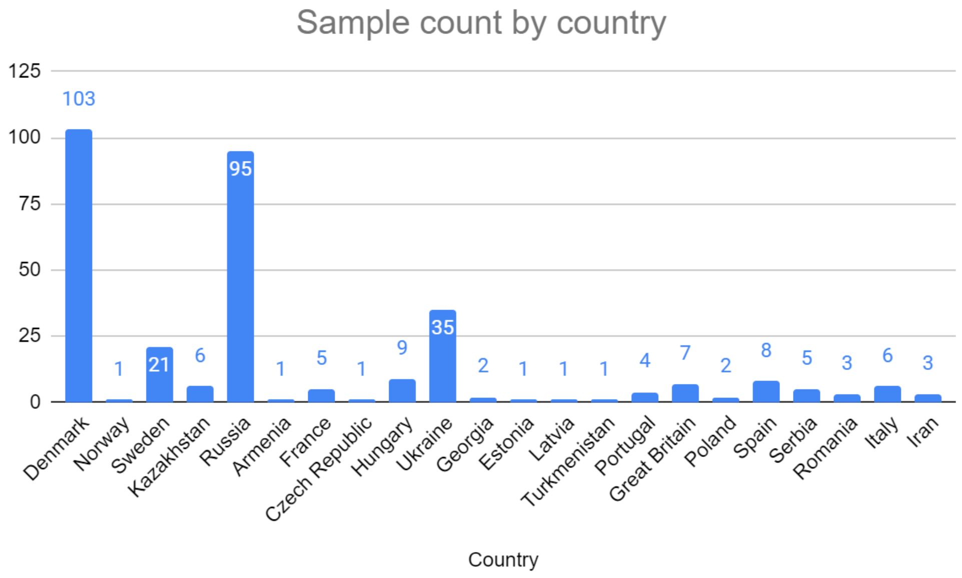


**Figure 7.2. Histogram showing number of sampled ancient human remains for which suitable ancient genome data was obtained, by country.**

### Armenia

#### Aknashen, Armavir, Armenia. Settlement and burials.

*Ruben Badalyan & Levon Yepiskoposyan*

The site of Aknashen-Khatunarkh is located in the Ararat Plain, in the basin of the Sev jur (Metsamor), at an altitude of 838 m, in the province (marz) of Armavir (6 km south of Vagharshapat, on the northeast periphery of the village of Aknashen). The site is an artificial hill (blur), circular in plan, 100 m in diameter (a surface area of about 0.8 hectares), with a flat top rising about 3.5 m above the plain. The excavations of the site have been conducted since 2004 by the Armenian-French joint expedition.

The site belongs to the Late Neolithic "Aratashen-Shulaveri-Shomutepe" culture and dates back to the first half of the VI millennium BC. Human remains occur both from a cultural layer of the synchronous deliberately committed burial (Tr.6, UF 11, F.15) and of household waste (unburied remains of a newborn, Tr.1, UF 8/NEO110), and intrusive burial of the end of Early Bronze - the beginning of the Middle Bronze Age (second half of the III millennium BC; Tr.7, UF 5, F2).

Literature: Badalyan et al. 2010[1]; Chataigner et al. 2014[2]

### Czech Republic

#### Vedrovice, Moravia, Czech Republic. Cemeteries and settlement.

*Václav Smrčka*

At least three different cemeteries have been found at this Linear Pottery site. Between 1961 and 1974 V. Ondruš investigated 11 graves within the settlement in the  tract of land “Široká u lesa“ [3]. Funerals H1/1963- H11/1974 and H2/1985 have been found in an area enclosed by a ditch (Erdwerk). North from this densely populated place there is a regular burial ground where the dead had been continually buried since the phase Ib1 of the Linear pottery culture [4]. A total of around 110 burials have been excavated at the site [5].

Besides classical contracted funerals into separate gravel pits (H4,H7 and H10), a settlement pit (H1) or burial in a construction pit (H5), and also cases of discarding (H9) and anthropophagy (H11) were recorded.

One sample was analysed for aDNA: NEO128, grave 10/74. The skeleton of a 40-50 year old male was found in this grave. Orientation of the grave pit was E-W (with head positioned towards the East, feet to the West). The male was positioned lying on his right side (right lateral recumbent). Position of the skull was right temple, unnaturally bent backwards, the rear of the skull touching the spine. View was to NE.

Left humerus was along the body, forming a right angle with forearm and palm in front of mandible, thumb and index in extended position and other fingers flexed. Right hand was stretched along the body and the palm touched the left knee joint. Positioning of the lower extremities was “semi-crouched”: the left femur was perpendicular to the body axis and the right knee joint was in the NW corner of the grave pit. Tibiae were crossed with the left tibia behind, right foot covering the left foot.

There was a hoof-shaped wedge in the area of the chest, 2 pcs of products of flaking, pottery fragment under the mandible, and a lump of graphite under the skull.

The male in grave 10/1974 exhibited a high ratio of 87Sr/86Sr in the femur compared with the tooth enamel and with the shell. He may have travelled between distinct environmental localities during his lifetime [6].

Literature: Ondruš V. 1961-1974[3]; Podborský V. (ed.), 2002[4]; Smrčka V. et al. 2005[6]; Richards et al. 2008[7]; Bickle and Whittle 2013[5]; Zvelebil & Pettitt 2013[8].

#### England

##### Hetty Pegler’s Tump (Uley Barrow), Gloucestershire, England. Long barrow.

*Alan K. Outram and Catriona J. McKenzie*

Hetty Pegler’s Tump, otherwise known as Uley Barrow, is a Neolithic Long Barrow about 8km SW of the modern town of Stroud, Gloucestershire, which was first excavated by Fry in 1821 [9]. It is an early Neolithic chambered tomb of the Cotswold-Severn type [10]. The long barrow is approximately 37 m long with a maximum breadth of 26 m. The entrance passage is at the SE end and originally led to five stone burial chambers, three of which now survive [9]. The early excavations recovered human skeletal material and the teeth and jaws of wild boar from the chambers [9], but in the upper levels there was also a human skeleton buried with Roman coins of the house of Constantine [9,11], showing later re-use of this funerary monument.

The fragmented skeletal material analysed in this study was, according to Gloucester museum records, discovered by a workman digging a fence line around the monument. Given the re-use of the monument and the nature of the discovery, direct dating of the remains was essential and has yielded a determination of 3641-3384 cal BC (95.4%, UBA-40120) confirming Neolithic date. The surviving remains from this discovery comprise four bones: an almost complete mandible (16 tooth sockets with 12 teeth present), an incomplete maxilla (8 tooth sockets with 5 teeth present), a right temporal (part of the squamous portion) and a left temporal (including petrous portion, jugular fossa and the internal auditory meatus). There is no duplication of skeletal elements and so it is possible that these bones may all have come from the same individual.

In terms of sex estimation, the only morphological traits available for analysis are the presence of a slight mental protuberance on the mandible and a pointed rather than squared symphyseal region of the mandible [12]. These two traits indicate that the remains may have come from a female individual. In terms of age at death, three third molars are fully erupted (the fourth was not present for analysis) indicating that this individual was older than 23.5 years of age at time of death [13]. Assessment of dental wear for age estimation following Brothwell [14] suggests that the left mandibular and maxillary molars are from an individual aged between 35-45 years of age, while the right mandibular molars are indicative of an individual older than 45 years of age. Overall, it is most likely that the remains came from an adult woman older than 35 years of age at time of death.

Literature: Brothwell 1981, Williams 1988. [9–14]; Clifford 1966[9]; Thomas 1988[10]; Williams 1998[11]; Brickley 2004[12]; AlQahtani et al. 2010[13].

#### Estonia

##### Sope

*Sergey V Vasilyev and S. B. Borutskaya*

The Corded Ware burial site at Sope is situated on a sandy knoll on the bank of the Sope. The majority of the skeletons were unearthed by farmers at the end of 19th – beginning of 20th century. The available information distinguishes two burial grounds within the larger burial area – Sope A and Sope B.

Sope A was discovered at the end of the 19th century. During the excavation of a cellar a human skeleton in a “sitting position” was unearthed, and in 1924 three accumulations of human remains were visible inside the cellar walls. Thus, this burial ground included at least four inhumations. Sope B was discovered in 1908 when the field was ploughed for the first time and allegedly seven skeletons with some bone and one amber artefact were found). Reportedly, one of the individuals lay on its side, with the lower limbs flexed at the knee. All the recovered bones were reburied nearby. Two more individuals were recovered from the field during archaeological excavations in 1926 [15] and 1933 [16]. Additionally, one skeleton has been found somewhere in the area, but the specific location is unknown.

Excavations revealed sandy soil that contained fragments and spots of charcoal, a small pottery shard and a few animal bones [15]. In the middle of the excavation plot, approximately 35 cm below the present surface in an area of 100×30 cm an accumulation of human and animal bones was found. No evidence of grave structures or above-ground markings were discovered. The body position of the individual was originally interpreted as tightly flexed and it was suggested that the corpse might have been bound prior to burial.

Previously, DNA results from Sope were published by Allentoft et al. [17]. This individual (RISE00, Sope I) was an adult female, probably in a flexed position. In the present project the grave Sope II was sampled, an adult woman buried in contracted position (NEO306). The same individual was analysed by Saag et al. [18]. Both individuals were dated to the Corded Ware period.

Literature: Moora 1926[15]; Indreko 1935[16]; Kriiska et al. 2007[19]; Allentoft et al. 2015[17]; Varul et al. 2019[20]; Saag et al. 2021[18].

#### France

##### Grotte Gazel, Aude, France. Double burial.

*Patrice Courtaud*

This cave (Sallèles-Cabardès, Aude, France) was occupied from the Upper Paleolithic until the historic times with several levels from Mesolithic to late Neolithic. This cave is mainly known by the remains from the Early Neolithic excavated by J. Guilaine between 1963 and 1971. A single burial, Gazel 1 (ERL 12288) 5733 ± 50 BP - 4692-4462 BC (94.5 % ) belongs to the Epicardial or early Chasséen [21,22]. The grave sampled here, Gazel 2, was a double grave including a child (around 5 y.) and an adult female (Figure S7.3). The two skeletons are very closely related. The archaeo-thanatological study conducted by H. Duday [23] showed that the two individuals were deposited simultaneously. This discovery is important as there is little funerary evidence from this period in this region of Languedoc and more generally in southern France. One of the questions is the relationship between the two deceased. The age estimation of the female based on the auricular surface of the hip bone [24] suggests a death occurring after the age of 40 years. The adult female in Gazel 2 was dated to (UBA-40111) 5740 ± 32 BP- 4686-4503 BC (95,4%), supporting the contemporaneity of the two burials.

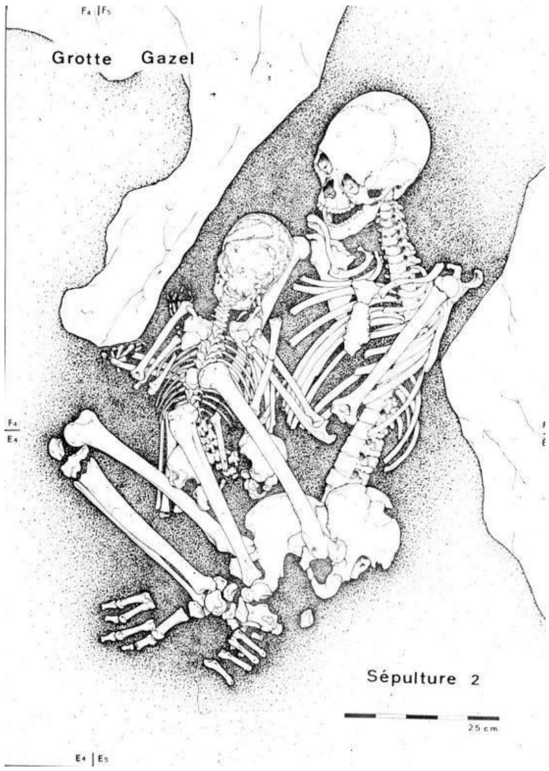


***Figure S7.3. The double grave at Grotte Gazel.*** *From Duday and Guilaine* 1980[23]

Literature: Duday and Guilaine 1980[23]; Guilaine & Manen 1995[21]; 2007[22]; Goude and Fontugne 2016[25].

#### Grotte Mandrin, Drôme, Rhône-Alpes, France. Rock shelter.

*Ludovic Slimak & Laure Metz*

Grotte Mandrin is a vaulted rockshelter located in the Middle Rhône valley, in Mediterranean France. The site has been excavated since 1990 and records a long Pleistocene sequence recording a dozen of archaeological layers, from layers J to B, concerning Neanderthal occupations and Early Anatomically modern Humans [26,27]. Recently, a tooth from a modern human was dated to ca 54000 BP, so far the oldest evidence for modern humans in western Europe[298]. The main fossil records of this very rich archaeological sequence concern occupations from MIS 5 to 3. This sequence of Pleistocene successions ends with a single Holocene unit that records several phases of human cremations from the late Neolithic and Bronze age. This Holocene layer layer A was mainly excavated in the 1960's by Gaston Etienne. The archaeological material from layer A is mainly composed of human remains, commonly burned, associated with a rare archaeological material composed of few foliate arrowheads, some pottery fragments and rare Mediterranean pierced shells. Grotte Mandrin was then only used during this Holocene phase for body treatments. This layer A was recently excavated in the entrance of the rockshelter in 2014 and 2015. The material analysed in this study come from both the recent 2015 excavations (human samples Man 15 A 1249 and Man 15 A 101) and from the Gaston Etienne 1960 waste that was screened in 2015 (Man A Déblais Gaston).

Literature: Slimak 2008[26]; 2019[27], Slimak et al. 2022[298].

#### Georgia

##### Kotias Klde, Chiatura, Georgia. Cave site.

*David Lordkipanidze*

The Kotias Klde cave is located in west Georgia, Chiatura region, near the village Sveri on the limestone Mandaeti plateau, in the Sadzalikhevi Gorge, 716 metres above sea level. While at the entrance it is quite spacious (23 metres wide and 8 metres high), most of it comprises a narrow and deep tunnel with the archaeological remains confined to the entrance area. The cave was discovered in 1968 by the Kvirila basin archaeological expedition headed by David Tushabramishvili, the Georgian State Museum, and assigned to the Neolithic – Early Bronze periods. It was tested in 2003 as part of the international (Georgian-USA-Israeli) project led by Tengiz Meshveliani, the Georgian National Museum. Systematic archaeological excavations were conducted in 2004-2006 and 2010 revealing a sequence of Upper Paleolithic (Layer C), Mesolithic (Layer B) and Neolithic (Layer A) occupations.

A grave was discovered in 2004 containing a complete skeleton of an adult male which was dug from layer A into layer B, almost reaching the stone pavement of sublayer B2. Stones crushing the skull, the knees and lower limbs of the skeleton were placed at the time of the burial. Radiocarbon dating of the tibia provided a date of 9,690-9,540 years cal BP. DNA from this individual was published by Jones et al. [28] and used to define the genetic group of Caucasian hunter-gatherers.

In the present project, six tooth samples were analysed (NEO279-NEO284), four of which yielded usable DNA. NEO281, found in layer B (basket #62, layer B, sq. G9, depth 335-340), NEO282, found in layer B (basket #153, layer B, sq. Igb, depth 335-350) and NEO284, also found in layer B, were found to belong to one and the same individual, which is also the same as the articulated Mesolithic skeleton published by Jones et al.[28]. These three samples were merged into NEO281. A new 14C date was obtained for this individual: UBA-40023, 8735±44 uncal BP, 9894-9552 cal BP 95.4%, slightly older than the previous date.

The tooth sampled as NEO283 was found deeper in the stratigraphy (layer B3/C, 234/9-10:242, sq. G9c, depth: 390-395). The date obtained for this sample was UBA-40025, 21412±194 uncal BP, 26009-25261 cal BP. This corresponds to the Paleolithic finds in layer C.

Literature: Tushabramishvili 1971[29]; Tushabramishvili and Nebieridze 1974[30]; Meshveliani et al. 2006[31]; 2007[32]; 2010[33]; 2013[34]; Bar-Oz et al. 2009[35]; Jones et al. 2015[28].

#### Hungary

##### Deszk – Olajkút

*Václav Smrčka & Olivér Gábor*

This Körös culture site was excavated by Otto Trogmayer [36]. In grave 6 was a gracile skeleton of a 30-35 year old female with a height of 155 cm and with torus palatinus on the palatal plates. An M2 tooth was analysed for aDNA and was dated to 6950±48 uncal BP (UBA-39965).

Literature: Farkas & Marcsik 1988[37]; Lipták 1974[38]; Ottó 1968[39]; Ottó 1984[40]; Whittle et al. 2002[41].

##### Hödmezövasarhely Kotac

*Václav Smrčka & Olivér Gábor*

The Körös culture site of Hődmezőváráshely-Kotac is not at this point archaeologically published. The skeletal remains are housed in the University of Szeged. One sample was analysed: NEO137, grave 25/2, no. 132. It was dated to 6728±42 uncal BP (UBA-39966).

Literature: [42,43]

#### Gorzsa Cukormajor, region Csongrád, Hungary

*Václav Smrčka & Olivér Gábor*

Hódmezővásárhely-Gorzsa is a settlement of the Tisza culture. The site lies in the environs of Hódmezővásárhely, at the confluence of the Tisza and Maros rivers. However, the archaeological site is not identical with the detached farmstead of Gorzsa shown on road maps; it lies some 10 km to its southwest. The site is marked as Főldvár (earthwork fort) on the 18th and 19th century maps. This  name has been replaced by names preserving the memory of the former owners (Cukor major). The name Főldvár probably coincides with the Mediaeval name of this area given by Slavic settlers. Goruša, the Old Slavic form of Gorzsa, is mound elevation. It is used to denote the entrance to a hillfort in modern Russian. According to an old description the remains of "fortified castle built in olden days“ being surrounded by rivers and marshes on all sides closed to strangers [44].

A total of 50 burials have so far been uncovered. These burials form discrete grave clusters; the preliminary serological analyses indicate that these burials contained the deceased of four successive generations of one genetic unit. These grave clusters appear, moreover, to have been organised according to sex.

The graves were uniformly southeast – northwest oriented. The deceased were buried in contracted  positions, with females laid on their left, and males on their right side. About one third of the graves contained vessels that had been deposited beside the skull. Female burials also yielded bone pins, and one male burial contained a marble macehead.

Seven individuals were analysed successfully for aDNA (Table S7.1). They were all dated to a short span of time at 5800-6100 BP uncal.

| **Sample** | **Description** |
|---|---|

| | |
|---|---|
| NEO143, 8999, grave 2 | Young person (18-20/22 years old). Deficient and fragmented postcranial skeleton with fragmented mandibula. |
| NEO147, 9005, grave 10 | Adult female with skull in good condition and postcranial skeleton. The neurocranium is medium length, narrow and high mesokran, hypsikran, akrokran, prot. occ. externa 0. The outline of the skull in norma verticalis is pentagonoid. The forehead is narrow and aurymetop. Glabella: 2. graduate. Due to the capacity of skull: aristenkephal. The face is narrow and of medium height., low, leptoprosop mesén, orthognath. Fossa canina 1. graduate. Orbita hypsikonch. Nose: camaerrhin, spina nasalis anterior 2. graduate. Palate brachystaphylin, alveolaris prognathia 1. graduate. On lumbal vertebrae: print of Schmorl's node.<br><br>Stature: 162 cm. Taxon: gracilis mediterrán—x (m—x) (2. fig.). Literature: MFMÉ 1987 |
| NEO148, 9014, grave 24 | Adult with skull in good condition and postcranial skeleton. The neurocranium long, narrow, dolichokran, chamaekran, tapeinokran. The outline of the skull in norma verticalis is pentagonoid, occipitalis externa O. The forehead is narrow and metriometop, glabella 2. grad. Due to the capacity of skull: euenkephal. The face is short orthognath. Fossa canina: 3. graduate. orbita: mesokonch. Nose: hyperchamaerrhin, alveolaris prognathia 3. graduate. Palate: mesostaphylin. In lambrasutures: ossa Wormiana. The first segment of ascrum was not grown together with the following one. On the right humerus: foramen entepicondyloideum.<br><br>Stature: 157 cm. Taxon: gracilis mediterranean – x(m-x) |
| NEO140, 9706, grave 27 | Banner János: Hódmezővásárhely, Gorzsa-Czukor major. Dolgozatok 1933-34.<br>Bálint Alajos: Hódemezőváráshely, Gorzsa-Czukor major. Dolgozatok 1937.<br>Gazdapusztai Gyula: Jelentés a Hódmezővásárhely-Gorzsa |

| | |
|---|---|
| | Cukor-majornál végzett ásatásról. RégFüz I/9 (1958)<br>Horváth F: Hódmezővásárhely Gorzsa. ArchÉrt 109 (1982) |
| NEO142, 9707, grave 28 | nd |
| NEO149, grave 49 | nd |
| NEO145, grave 60 | nd |

***Table S7.1. Details of samples from Gorzsa included in the present study.***

Literature: Banner 1934[43]; Bálint 1937[45]; Gazdapusztai 1958[46]; Horváth 1982[44]

#### Iran

##### Tepe Guran, Hulailan Valley, Central Zagros, Iran, grave/pit find

*Peder Mortensen & Pernille Bangsgaard*

The site was excavated in 1963 by the Danish Archaeological Expedition to Iran. It is a classic tell site with remains from the Iron Age, Bronze Age and the Neolithic Periods, with the latter including 18 layers of unbroken habitation (dated to 6700 – 5500 BC. BC). The three samples all come from these Neolithic layers. Two samples originate in area GI. Sample 71 is from level Q, where a tightly flexed female skeleton was found, her age app. 25 years. Sample 75 is from Level V and inside a pit four individuals were found in a disarticulated state, a likely reburial. Sample 75 is the partial skull of an adult around 20-30 years old. From area GIII a pit contained a male skeleton in a tightly flexed position, aged around 35-40 years.

Literature: Meldgaard et al. 1964[47]; Alexandersen et al. 1981[48]; Thrane 2001[49]; Mortensen 2014[50]

#### Italy

##### Fontenoce, Recanati, Italy

*Gabriele Scorrano, Cristina Martinez-Labarga, Mario Federico Rolfo, Mara Silvestrini, Francesca Radina, Paola Aurino, Mauro Calattini, Lucia Sarti, Enrico Cappellini & Olga Rickards*

The archaeological site of Fontenoce di Recanati is located at few kilometres away from the Adriatic coast in Central Italy. The Chalcolithic settlement was excavated under the supervision of Dr. Mara Silvestrini starting in 1984, when an isolated “grotticella” grave was brought to the light. In the subsequent years, several graves associated to Rinaldone culture and trans-Adriatic influence were discovered. All the radiometric dates set Fontenoce di Recanati in the Early Copper Age (3600-3300 cal BC). The sample analysed is a female dated to 4661 ± 30 BP (rib).

The human remains appeared to be in a good level of preservation, and most of the inhumations present complete skeletons, with the bones in anatomical association. There were single and multiple burials, with the inhumations usually lying in a flexed position on the right side, with great variability of the limbs. The exceptional human bone preservation allowed several interdisciplinary studies, such as the occupational markers analysis showing an intense physical activity, such as hunting, while the stable isotope analysis indicated a diet with remarkably high but not exclusive consumption of animal proteins.

Litterature: Silvestrini et al. 1992[51]; 2005[52]; 2011[53]; Cencetti et al. 2005[54]; Dolfini et al. 2010[55]; De Angelis et al. 2019[56].

##### Gaudo, Paestum, Italy

*Gabriele Scorrano, Cristina Martinez-Labarga, Mario Federico Rolfo, Mara Silvestrini, Francesca Radina, Paola Aurino, Mauro Calattini, Lucia Sarti, Enrico Cappellini & Olga Rickards*

The excavation and discovery of the Eneolithic necropolis of Gaudo di Paestum, in the locality of Gaudo between the Fiumarello and Contrada Seliano, took place in three important moments:

1. In the autumn of 1943, strictly connected to an exceptional and contingent event, represented by the Second World War, during the works for the construction of a military airport.

In this first stage the excavation took place with mechanical vehicles and the tombs were unfortunately completely damaged; about 50 vases, flint daggers and a copper dagger were recovered.

In June 1944, during the manual extraction of other quantities of tuff, tomb 10 was intercepted leading to the discovery of the culture of Gaudo. The work done by both Lieutenant Morris, who was in charge of the work at the time, and by Lieutenant Brinson, director of the B.P. Mobile Archaeological Unit, in anticipation of any archaeological finds was fundamental.

Lieutenant Brinson carried out the survey of about 9 tombs previously identified by the Allied army and he excavated tomb 10, today known as the Brinson tomb, with great stratigraphic rigor.

The disposition of the human remains indicated that the burials had taken place in succession for a long period of time, and that the ceramics and skeletons had been pushed to the bottom each time and piled on top of each other to make room for the new burials. Furthermore, since the skeletons had the legs collapsed laterally in a squatting position, on the basis of the detached position of the skull, it was possible to assume that the bodies were originally buried curled up on their side or in a sitting position. However, it could not be excluded that these were secondary burials, meaning that the bodies may have been previously exposed elsewhere and the skeletons brought later to their last "home".

2. In 1945, Pellegrino Claudio Sestieri inspected the necropolis of Gaudo and collected materials. Then, two excavation campaigns were carried out in 1945, others between 1946 and 1947 bringing to light a total of 19 tomb contexts whose outfits flowed into the Paestum Museum, which was inaugurated in 1952.

3. The last excavations were conducted by the "Soprintendenza di Salerno" on several occasions since 1962 by Giuseppe Voza who discovered 14 new tombs. Finally, in August 1969 the emergency intervention of the staff of the Paestum museum.

The superimposition of Brinson's planimetric sketch with Voza's planimetry allowed Antonio Salerno and Gianni Bailo Modesti to understand that the tomb 10 excavated by Brinson actually constituted only one of the cells of a bicellular complex digged by Voell in 1962. This second cell was defined as "grave e".

The first cell contained the disconnected remains of about 12 individuals: 10 adults, 1 adolescent and 1 child, while in the second cell were discovered the remains pertinent to about 7 individuals: 5 adults, and 2 sub-adults, still preserved in the Museum of Paestum.

There are some radiocarbon dates from two individuals: individual 5 from tomb IXA: (LTL 13376A) 4593 ± 45 BP, 3520-3310 BC (67.2%, 2σ), 3300-3260 BC (1.5%, 2σ), 3240-3100 BC (26.7%, 2σ); and from individual 13 of the IXB tomb of Paestum: (LTL 3338A) 4478 ± 45 BP, 3360-3010 BC (95.4%, 2σ). The chronology between the middle of the IV millennium and the middle of the III millennium, remains substantially unpublished and mainly derives from the presence in Paestum of tombs with multiple depositions in which there are up to 70 individuals in a single cell.

Literature: Aurino 2011[57], 2013[58].

#### Grotta delle Mura, Monopoli, Bari, Italy

*Gabriele Scorrano, Cristina Martinez-Labarga, Mario Federico Rolfo, Mara Silvestrini, Francesca Radina, Paola Aurino, Mauro Calattini, Lucia Sarti, Enrico Cappellini & Olga Rickards*

The archaeological site of Grotta delle Mura is located in the municipality of Monopoli (Apulia), a few metres away from the Adriatic coast in southern Italy. It was discovered by Prof. Anelli in 1952. The excavation has been under the supervision of “Dipartimento di Scienze storiche e dei Beni culturali” of Siena University since 1985.

On the basis of the different artefacts found and some radiocarbon dating performed on charcoal and bone remains, the settlements span from Middle Palaeolithic to the Ancient Neolithic. Several stratigraphic units have been identified, with the most ancient being associated with the Mesolithic and the Final Epigravettian (from SU 125 to SU 130). From SU 130 dated to 11420 ± 100 BP, comes one of the most important Italian Epipaleolithic graves, a child. The sample analysed in the present paper (NEO806) came from SU 123, which is associated with the Ancient Neolithic level. However, the sample was directly dated to the Bronze Age (3014±26 uncal BP, UBA-40109).

Literature: Calattini 2002[59]; Calattini & Tessaro 2016[60]; Calattini et al. 2003[61], 2017[62].

#### Grotta Nisco, Cassano Murge, Bari, Italy, Cave

*Gabriele Scorrano, Cristina Martinez-Labarga, Mario Federico Rolfo, Mara Silvestrini, Francesca Radina, Paola Aurino, Mauro Calattini, Lucia Sarti, Enrico Cappellini & Olga Rickards*

The Eneolithic site of Grotta Nisco is a karst cave located in the territory of Cassano Murge (Bari) 380 metres above the sea level, studied by the "Soprintendenza per i Beni Archeologici della Puglia". It was a collective tomb dated between 2,500 and 1,800 BCE.

The very narrow entrance leads, after a drop of more than 1m in a vestibule leading to the central hall around which several irregular rooms are placed on different levels.

In room 1, skeletal remains of 3 disarticulated adults were found, skeletons A and B, both adult males, had a metal stab with nails, flint tools and 3 arrow cusps, the adult female skeleton C, had a clay spindle. In the environment there were other fragments of various small sizes. Large bovid bones were also found, some with clear signs of slaughter.

In room 5, located in the rear part of room 1, the skeletal remains of an individual in a secondary position were recovered, with a kit consisting of a dagger with ogival contour and triangular blade, along with a mug with geometric enterprise decoration.

Overall, a total of 18631 bone fragments were recovered, 4807 of which were properly classified as human (1709, 35.5%) and faunal (3098). The minimum number of individuals was 20: 14 adults, 2 juveniles, 3 infants and, probably a full-term foetus should be added.

The mean stature was about 159 cm, and the males probably were subjected to a greater biomechanical load on the lower limbs, with evidence of running on rough substrates, climbing on rocks, throwing objects and crouching. The general health status was good, only a few cases of malnutrition are evident in the early stages of childhood, but in adulthood, nutrition had to be good based on animal husbandry products and low carbohydrate intake. The frequency of caries is very low, even the loss of ante-mortem teeth and there are no signs attributable to anemias. Based on the analysis of the faunal remains (with 60% of remains attributable to sacrificed *Ovis* or *Capra* and the presence of dog), a pastoral economy based on sheep and goat breeding may be hypothesised.

Literature: Venturo et al. 2011[63]; Radina and Savino 2018[64].

#### Maddalena di Muccia A, Province of Macerata, Marche Region, Italy. Settlement.

*Alfredo Coppa*

The Maddalena di Muccia site was discovered and partially investigated in the 1960s by Delia Lollini [65] who had found structures, ceramics and lithic industry dated to the Ancient Neolithic. Since 2001, as part of a larger program, which involved the University of Rome La Sapienza, the area has been systematically investigated, with extensive excavations and regular research campaigns carried out in the years 2001-2003, 2005, 2006 and 2008 [66].

The sample examined (NEO695) comes from the excavations of the 1960s in which only one burial was found (A) of which the skull with mandible, without the splanchnocranium, and various parts of the postcranial skeleton are preserved. The skeleton belongs to an adult female individual, which was determined by the morphology of the skull and hip bone [67,68 a,69 b]. The skeleton was found in the deposits of the ancient Neolithic ceramic impressed with an uncalibrated dating of 6580 ± 75 BP which, calibrated at 2σ, chronologically places the site between 5640 and 5460 BC [70]. To these dating results are added those coming from the new excavations (US114- 6638 ± 59, 6637 ± 83, 6440 ± 50 BP) which confirm the chronology of the site in the 6th millennium BC.

The dental morphological traits of this sample have been employed, together with those of many other Neolithic ruins, in a study as direct indicators of biological affinities among the populations that inhabited the Italian peninsula from the Upper Palaeolithic-Mesolithic to Mediaeval times. The results from the principal component analysis, maximum likelihood, mean measure of divergence, and multidimensional scaling do indicate a net separation of the Paleo-Mesolithic sample from the other groups that is not related to dental reduction. This suggests that the shift in dental morphology was the product of Neolithic populations migrating into the peninsula from other areas. Nonetheless, the Paleo-Mesolithic populations share several discriminative traits with the Neolithic group. The biological relevance of such evidence suggests that, to some minor extent, the spread of agriculture did not occur by total population replacement [71].

Literature: Lollini 1965[65]; Corrain and Capitanio 1968[67]; Alessio et al. 1970[70]; Lollini 1991[68]; 1991[69]; Coppa et al. 2007[71]; Manfredini 2014[66].

##### Mora Cavorso cave, Jenne, Rome, Italy. Cave.

*Gabriele Scorrano, Cristina Martinez-Labarga, Mario Federico Rolfo, Mara Silvestrini, Francesca Radina, Paola Aurino, Mauro Calattini, Lucia Sarti, Enrico Cappellini & Olga Rickards*

Grotta Mora Cavorso is a cave with a stratified deposit, located in the inner Apennines in Central Italy. The archaeological deposit spans from Upper Palaeolithic to historical age. The focus of the deposit is the Neolithic level, which holds one of the largest Neolithic burial deposits of central Italy. Multidisciplinary investigations carried out between 2006 and 2018 identified approximately 600 highly fragmented human bones in two little inner rooms (UP and LR) of the carsick complex.

The minimum number of individuals amounts to 28, and the estimation of the age at death indicates: 11 foetus, perinatal and infant (I-II), 1 juvenile, 5 young adults and 11 adults. The majority of human bones were found in one of the two rooms (LR), chaotically piled with no skeletal connection. A smaller number of human bones, still partially articulated, were found in the contiguous room (UR). In this context the human bones were found lying directly on the crust, without any apparent delimitation of the funerary space. Radiometric dating of the Neolithic context are between 6874 ± 45 BP (jaw of a domestic caprine) and 6000±45 BP (charcoal of hearth), and set the Neolithic level in the early phase of the Neolithic period of the Italian peninsula. The archaeological finds in the burial level are scarce, as eleven shell and stone beads, possibly ornaments worn by the dead, were found, together with pottery, a stone polished axe and five flint bladelets as grave goods. Rare and almost exclusively domestic mammals have been identified.

Here we report results from an adult individual (NEO834), dated to 6499±41 uncal BP, UBA-40801.

Literature: Scorrano et al. 2015[72], 2019[73], Rolfo et al. 2016[74], 2017[75], Silvestri et al. 2020[76].

#### Kazakhstan

##### Gregorievka 1, Pavlodar, Kazakhstan. Cemetery.

*V. K. Merz, translated from Russian by Aija Macane*

The Gregorievka 1 flat earth burial ground, grave No. 2, is located 100 m south-east of the grave No. 1, in the village of the same name, on the first terrace above the floodplain on the right bank of the river Irtysh. The grave was discovered on a riverbank bluff, where a dark filling of the grave pit and small bones were recorded. At a depth of 0.6 m, the grave had a square shape 1.5 x 1.6 m in size, oriented to the north with a slight inclination to the east, and with a collapsed southern edge. By the northern and eastern sides of the grave, small pits were recorded. The depth of the grave reached 1 m, and it has been plundered in the past. The red-black filling of the grave contained fragments of charcoal, scattered human bones, as well as bones of cattle and sheep, fragments of ceramics decorated with “stepping” comb stamp impressions (made by rolling), a bone object resembling an awl, and fragments of ochre. At the bottom of the pit, near the eastern wall a spot of ochre was documented [77 b,78].

The postcranial skeleton was studied by the senior researcher of the Physical Anthropology Sector of the Tyumen Scientific Centre of the Siberian Branch of the Russian Academy of Sciences, Candidate of Historical Sciences K.N. Solodovnikov. It belongs to a woman about 18 years old (NEO899). The skull belongs to the dolichocephalic Europioid type with a narrow face, similar to the type of the male population of the Yelunin culture in the Upper Ob region. The skeleton was dated to 3696±20 uncal BP, OxA-39283, similar to the date from burial 1.

Based on peculiarities of the funeral practices and grave inventory, this burial belongs to the Yelunin culture of the early animal herders in the interfluve of the Ob and Irtysh. Despite the fact that the grave had been looted and small material was obtained from it, the burial can still be attributed to the Yelunin culture and reflects the particularity of the development of the Yelunin population in the southern part of the middle reaches of the Irtysh River [79].

A 14C date, UBA-32665, was previously obtained from grave 1, indicating the interval of its construction between 2037–2285 BC [80]. Earlier, before any results of radiocarbon analysis were obtained, this burial was attributed to the Yamnaya cultural type [77,78,81]. However, the obtained radiocarbon date forces us to abandon the original interpretation linking this burial to Yamnaya, confirmed also by the significant difference between the anthropological type of the woman in the grave 1 and the populations of the Yamnaya and Afanasievo cultures [81].

Thus, based on these facts, both burials should be considered within the “Klinskaya” sub-group of the burials of Yelunin culture, and reflect the specificity of the burial practices of the Yelunin population in the southern part of the middle reaches of the Irtysh River ([79]). However, it should be noted that the anthropological type of women buried in burial 1 and 2 is very different from each other. That suggests the multicomponent character of the population of Yelunin culture, which has been repeatedly noted by anthropologists ([82]). The Irtysh valley was a contact zone between the populations of the steppe, forest-steppe and sub-taiga zones.

Literature: Tur & Solodovnikov 2003[82]; Merz, V. K. 2007[78]; 2007[77], Merz & Merz 2010[81]; Khokhlov et al. 2016[83]; Merz, I. V. 2017[79]; Damgaard 2018[80].

#### Shauke settlement, Pavlodarsk, Kazakhstan. Settlement and burial

*V. K. Merz, translated from Russian by Aija Macane*

The Shauke settlement is located on the first terrace above the floodplain on the right bank of the River Irtysh. It is situated 0.5 km north of the village of the same name, on the site of an old sandpit. The settlement was discovered in 1992 by V.K. Merz. Repeated collection of archaeological material from the Bronze – Early Iron Age (III–I millennium BC) was carried out at the site in 1996, 2001, 2006 and 2011.

NEO901 Shauke settlement, Burial 2: The burial No. 2 was discovered in 1996 by V.K. Merz while surveying parts of settlement which had been subject to wind erosion. The burial was completely preserved, the skeleton lay on the left side with its head to the north. No inventory was documented. Direct dating showed the skeleton to be from the Mediaeval period.

The postcranial skeleton was studied in 2012 by the senior researcher of the Physical Anthropology Sector of the Tyumen Scientific Centre of the Siberian Branch of the Russian Academy of Sciences, Candidate of Historical Sciences K.N. Solodovnikov. It belongs to a woman, 30-40 years old. The skull is gracile, sharply dolichocephal with elongated proportions of the cranial and facial parts of the skull, an Europeoid with sharply wedge-shaped face and slightly protruding nose. No analogous examples can be found among the Neolithic-Bronze Age populations of the interfleuve of Ob and Irtysh rivers.

Literature: Merz 2011[84]; Svyatko et al. 2015[85]; Merz & Svyatko 2016[86].

#### Shauke 1, Pavlodarsk, Kazakhstan. Cemetery.

*V. K. Merz, translated from Russian by Aija Macane*

The burial ground Shauke 1 is located on the first terrace above the floodplain on the right bank of the river Irtysh, 1.2 km north of the village of the same name, on the site of a sandpit. The burial ground was discovered in 2010 by V.K. Merz, who discovered accumulations of human bones, ceramics and other objects on the eastern edge of the quarry. The finds were concentrated in several heaps, which received provisional designation as graves in the order of their discovery No. 1–3. While surveying the edges of the quarry, 140 m to the west of the location of the artefacts, the remains of a destroyed grave was discovered. At a depth of 1.7 m from the modern-day surface, a rectangular spot of red and black soil concretion, 1.08x0.3 m in size, was documented. A grave layer or filling, 15 cm thick, was recorded on top of this spot. On this basis it was concluded that the three discovered burials had been transferred by a loader from this very place. The excavations carried out at this site on May 27, 2012, revealed that the spot consists of the north-eastern edge of a rectangular grave pit. No finds were found in the filling (Merz 2011: Fig. 2). Nearby, from the soil fallen down from the quarry walls and from an area newly opened by a bulldozer, fragments of ceramics of the Yelunin culture were collected and grave No. 4 was discovered. Postcranial skeletons were studied by senior researcher of the Physical Anthropology Sector of the Tyumen Scientific Centre of the Siberian Branch of the Russian Academy of Sciences, Candidate of Historical Sciences K.N. Solodovnikov.

To study the reservoir effect in the region, paired samples were taken from burials No. 1–4 (1 human bone, 1 animal bone) in 2014. The following dates were obtained from grave No. 2:

UBA-268929, human bone: 3772±33 uncal BP, 2293–2047 cal BC

UBA-268939, sheep bone: 3706±36 uncal BP, 2202–1980 cal BC

The following dates were obtained from grave No. 3:

UBA-268949, human bone: 3782±33 uncal BP, 2334–2050 cal BC

UBA-268959, sheep bone: 3761±40 uncal BP, 2292–2036 cal BC

Archaeological material accompanying the bone remains indicates that the burials of the Shauke 1 burial ground belong to the Yelunin archaeological culture, based on four AMS dates. Burial No. 2 is dated to the XXIII–XX centuries BC, burial No. 3 to the XXIII–XXI centuries BC [85,86].

All in all, the radiocarbon and isotopic studies carried out on objects of the Early Bronze Age in the southern part of the Middle Irtysh region revealed the existence of a freshwater reservoir effect in the region. That means that 14C-dates, especially of human bones, turn out to be much older. For the middle reaches of the Irtysh, within the Pavlodar region, the reservoir effect is ≈ 157–224 years ([85]).

NEO900 Shauke 1 Burial 2

Burial No. 2 contained an open, jar-shaped vessel, decorated with a “stepping” comb stamp (made by rolling). A piece of a stone crucible and a tanged arrowhead, and bones of sheep and calf were found nearby. Skeletal remains belonged to a man, 40–50 years old. The skull is dolichocephalic, Europeoid with a narrow face, similar to the type of male population of the Yelunin culture of the Upper Ob region. The sample was dated to 3831±20 uncal BP, OxA-39284.

NEO902 Shauke 1 Burial 3

Burial No. 3 contained two wall sherds of a clay vessel, one of which had a raised cordon decorated with comb stamp impressions. A fragment of an astragalus of a small ruminant covered with green oxides from non-ferrous metal, a fragment of a metal object and 37 animal astragali (cattle – 2 pcs, sheep – 35 pcs), as well as horse and sheep bones were found nearby. Bone remains belonged to an adolescent, presumably male, 14–16 years old. The skull is dolichocephalic, probably Europeoid with somewhat softened Europeoid features. Possibly similar to the type of population of the Yelunin and Krotovo cultures in the interfluve of the Ob and Irtysh. The NEO902 sample was dated to the Iron Age, 2337±18 uncal BP, OxA-39286. This sample therefore most likely comes from another individual, and the bone assemblage may be considered as mixed, probably in connection with the gravel extraction.

Literature: Merz 2011[84]; Svyatko et al. 2015[85]; Merz & Svyatko 2016[86].

#### Sjiderty 10, Pavlodarsk, Kazakhstan. Barrows.

*V. K. Merz, translated from Russian by Aija Macane*

The Sjiderty 10 burial ground is located 5.5 km to the south of the Sjiderty station, in a valley formed by the high right bank of the Sjiderty river and the hills. Two burial mounds (at a distance of 5 m from each other) are situated at the lowest part of the valley, near the confluence of a temporary watercourse and the river. The burial ground was discovered in 2001 by V.K. Merz. In 2001 V.K. Merz investigated the mound No. 1, and in 2002 mound No. 2. Both of them have been plundered earlier. The mounds were stone-and-earth-mixed barrows in the form of an irregular circle, with a diameter of 8x9 and 9x7.6 m, and a height of 0.45 and 0.15 m.

Square settings built of large stones were discovered under the barrows. In the middle of them, earth graves with oval pits contained collective burials of adults and children, placed in crouched positions on their backs, with their heads oriented to the north-west. The walls of the graves were burnt and charcoal was noted in the fill, while concretions of burnt soil and ochre were present in the graves.

The burial inventory is represented by fragments of pottery, decorated with a “stepping” comb stamp (made by rolling), mollusc shells (Unio sp.), flint chips and flakes, fragments of arrowheads and a bronze knife and a turquoise bead. Also bones of horse, cattle and small ruminants were discovered with the burials, interpreted as remains of sacrificial food.  A shrine was discovered in the north-eastern sector of the mound No.1, and contained, in addition to the above-mentioned categories of objects, a grinding stone and a bone awl.

The postcranial skeletons were studied by the head of the anthropological department of the Altai State University, Candidate of Historical Sciences S.S. Tur, and the senior researcher of the Physical Anthropology Sector of the Tyumen Scientific Centre of the Siberian Branch of the Russian Academy of Sciences, Candidate of Historical Sciences K.N. Solodovnikov.

A 14C date (COAH – 4860, 3835 ± 90 uncal BP) was obtained from a charcoal sample from the grave in mound No. 1, and suggests that the object was constructed in the interval between 2563–2031 BC [79,86 Fig. 2,87,88].

The burial ground belongs to the Sjiderian variant of the Yelunin culture, and illustrates the specific development of the Yelunin population in the north-eastern Saryarka (Merz, 2017: 20). Particular aspects of the funerary practice and inventory suggest cultural influences from eastern Europe (Merz, 2018: 138).

Mound No. 2 had an irregularly shaped flat stone-earth barrow 9x7.6 m in size and 0.15 m high. A square setting of large stones was discovered underneath it. Horse bones and a stone ball were found in the filling of the mound. In the centre, there was a heavily destroyed oval pit, 2.4x1.6 m in size, dug 1.39 m deep into the ground, and oriented in WNW-ESE. An accumulation of charcoal was found by the eastern wall, at a depth of 0.8 m, and at the bottom of the pit a concretion of soil with a diameter of 0.33 m. Remains of three people were found, sprinkled with ochre: 1) a woman of mature (?) age; 2) a neonatus or a foetus in perinatal period; 3) a child about 5 ± 1 years old (NEO904). Fragments of horse, cattle and sheep bones were found among the human bones. A turquoise bead was found in the grave filling (Merz, 2003). The radiocarbon date UBA-39933 was obtained from the teeth of the five year old child.

Literature: Merz, V. K. 2002[87], 2003[88], Merz & Svyatko 2016[86]; Merz, I. V. 2017[I.V. 79], 2018[I.V. 89].

#### Bestamak, Kazakhstan. Cemetery.

*I. V. Shevnina & A. V. Logvin*

The burial ground Bestamak is located on the right bank of the river Buruktal (Kostanay region, Northern Kazakhstan) The site was investigated by the Turgai archaeological expedition in 1991-2013. The necropolis functioned in antiquity for a long period of time: the later Stone Age - the Bronze Age - the Early Iron Age - the Middle Ages. Most of the burials belong to the Sintashta period of the Bronze Age. Pit No. 117 is rounded in shape, 1.15 m in diameter, 0.9 m deep. In the burial, a skeleton of a woman of 50-60 years old was found in a sitting position, facing the pit wall. The whole skeleton is abundantly sprinkled with ocher. Behind the buried back was a stone ball with a diameter of 15 mm and a flake without retouching (Logvin, Shevnina, 2009, p. 104-110; Logvin, Shevnina, 2009, p. 142-151; Logvin, Shevnina, 2013, p. 231-244).

Literature: Logvin & Shevnina 2009[90]; 2013[91 2013]; Logvin et al. 2009[92].

#### Latvia

##### Zvejnieki, Latvia. Cemetery and settlement.

*Elizaveta V. Veselovskaya & Sergey Vasilyev*

Zvejnieki is a large Mesolithic and Neolithic burial ground and settlement, located in Northern Latvia at the Ruja River head by lake Burtnieks. Excavations of the 1964-1971 expeditions of the Institute of History of Latvia Academy of Sciences were conducted under the guidance of F.A. Zagorskis. 302 Mesolithic and Neolithic burials were discovered. The cemetery has a long period of use, from ca 7500 to 2600 cal BC. According to the inventory, it belongs to the Kunda Mesolithic culture, which was distributed in the territory of Latvia and Estonia from the 8th to 5th millennium BC [93]. The Comb Ware and Corded Ware cultures are also represented. Most burials are single burials in supine position, with variable orientation and covered with ochre. Several graves were excavated by Lars Larsson [e.g. 94]. The total number of burials is at least 348.

Two anthropological variants are represented [95]. One is Caucasoid, sharply dolichocranial with a medium-wide, high, significantly profiled face and protruding nose. The other is dolicho-mesocranial with a wider and flattened face. There are sculptural reconstructions made on the skulls of representatives of both types by G.V. Lebedinsky.

Dietary isotopes were analysed by Eriksson [96]. Previous aDNA analyses were published by Jones et al. [97] and Mathieson et al. [98], suggesting the presence of Western Hunter-Gatherer ancestry.

In the present project one individual is reported (NEO307), dated to the Mesolithic. A further sampled individual (NEO308) was dated to the Mediaeval period and was not included in further analysis. The two skulls were sent to Moscow for facial reconstruction by Lebedinsky [99].

NEO307 is a male adult, 30-40 years old, from burial 2. The skeleton was buried in a supine position, sprayed with ochre but without artefacts. It was dated to the Middle Mesolithic, 7313±49 uncal BP, UBA-40033. A previously published date, 6900±65 uncal BP, Ua-3638, is considered less reliable.

Literature: Denisova 1973[95]; Zagorskis & Zagorska 1973[93]; Eriksson 2003[96]; Zagorskis 2004[99];Jones et al. 2017[97]; Larsson et al. 2017[94]; Mathieson et al. 2018[98].

#### Norway

##### Hummervikholmen, Søgne, Agder, Norway. Submarine find.

*Per Persson*

In 1994 the landowner dredged the small harbour on the island Hummervikholmen. He then found a human skull that later was radiocarbon dated to the Mesolithic. The find spot was at c. 1 m depth in the sea. In the following years, divers from the Norwegian Maritime Museum documented the site and collected more human bones [100]. In 2013 the remaining part of the undisturbed sediment in the harbour was excavated in a joint project between the Norwegian Maritime Museum and the Museum of Cultural History [101]. In total about 20 human bones were collected. There are skulls/-fragments from at least three individuals. No artefacts were found at the site. Trial pits have been excavated on the island with negative results. There are in total 11 radiocarbon dates on human bones from the site, except for one outlier they are in the range c. 8400-8850 uncal BP. An investigation on diet from stable isotopes shows an extreme marine diet; average δC13 was -13.7 and δN15 was 19.8 [102]. From the stratigraphy there are 9 radiocarbon dates on natural wood both above and below the skeletal remains, dating the bones to c. 7500-7000 cal BC. These datings in combination with nearby dating of shoreline displacement, show that the find spot was dry land between c. 8000 and 7000 cal BC and that it was up to c. 5 m above the contemporary sea level. Most likely these individuals have been buried on dry land and the graves destroyed by the sea when the site was transgressed and covered by marine sediments. Two other individuals from Hummervikholmen were investigated for ancient DNA by Günther et al. [103].

One sample was successfully analysed for DNA (NEO017). This is a tooth (X82) found close to a cranium (X90) of a male aged 33-45 years.

Literature: Sellevold & Skar 1999[100]; Eggen & Nymoen 2014[101]; Skar et al. 2016[102]; Günther et al. 2018[103]

#### Poland

##### Słonowice, Kazimierza Wielka district, site 5, Poland. Cemetery (long barrows)

*Krzysztof Tunia & Piotr Włodarczak*

A cemetery of the Funnel Beaker culture was excavated in 1979-2012 by Krzysztof Tunia (Institute of Archaeology and Ethnology, Polish Academy of Sciences). The research included the lower part of the southern slope of the upland gently descending to the Małoszówka river. There were discovered 10 tombs (megaxylons, known also as tombs of Niedźwiedź type, corresponding to earthen long barrows in west European terms), as well as a ceremonial square. The singular and (occasionally) double burials were recorded within and outside wooden constructions of tombs. Based on singular radiocarbon dates and typochronological studies, the cemetery is dated to c. 3600-3300 BC. At present, Słonowice is the largest cemetery from the middle Eneolithic period in south-eastern Poland. Two samples were analysed from this site.

NEO640, Feature (grave) No. 12/II. A 2.2 x 0.8 m rectangular grave pit, with a longer axis located in the W-E direction. 1.45 m deep (from the modern surface of the earth). A feature spatially connected with megaxylon No. I - dug into the ditch, from which the soil was taken for the tomb embankment, thus chronologically younger than it. At the bottom of the grave was a skeleton located along the W-E axis. The deceased has been situated in an upright position with arms placed along the torso, head to the west.

NEO641, Feature (grave) No. 10/XXIV. A 2.7 x 1.3 m rectangular grave pit, with a longer axis located in the W-E direction. 1.05 m deep (from the modern surface of the earth). A feature spatially connected with megaxylon No. IV- located about 1 m to the N from its northern edge. Grave equipment - 1 flint flake from the Volhynian flint. At the bottom of the grave was a skeleton located along the W-E axis. The deceased has been situated in an upright position, head to the west. Skeleton partially damaged.

Literature: Tunia 2006[104]; Przybyła & Tunia 2013[105].

#### Portugal

##### Gruta do Caldeirão, Tomar, Portugal. Cave site.

*João Zilhão*

The samples analysed come from Horizon NA2 of Gruta do Caldeirão. This archaeological horizon comprises non-articulated human remains, sheep bones, Cardial-decorated ceramics and other impressed  wares intruded into Upper Magdalenian layer Eb, which, due to a post-Pleistocene sedimentation hiatus, formed the cave floor at the time of Early Neolithic funerary activity. Coupled with the analysis of spatial distributions, the physical anthropological study concluded that the set of human remains assigned to NA2 represented four adults and a child, with the dental remains suggesting the possibility of a fifth adult being present. Available dating results indicate that this NA2 ensemble includes individuals buried in at least two different moments, c. 5200-5300 cal BC, and c. 5350-5450 cal BC; the direct dating of the samples analysed here shows that the right temporal R11-8 belongs to the earlier phase and the left temporal Q12-186 to the later phase. Stable isotope analysis of these remains shows that they had a terrestrial diet with no measurable input of aquatic foods, in marked contrast with coeval Late Mesolithic individuals buried in the Muge shell-middens of the Tagus valley, ~60 km downstream. Previous analysis of the mitochondrial DNA preserved in twelve dental samples assigned Gruta do Caldeirão's Early Neolithic humans to haplogroups H (six), H/U (three), U* (two) and V (one), and suggested genetic discontinuity with the region's Mesolithic hunter-gatherer populations (Chandler et al 2005).

Literature: Zilhão 1992; Jackes and Lubell 1992; Zilhão 1993, 2000; Lubell et al. 1994; Chandler et al. 2005; Isern et al. 2017 [106–111].

##### São Paulo II, Almada, Lisbon, Portugal. Hypogeum.

*Ana Maria Silva*

In 1989 during repairs in the churchyard of the Old Dominican Convent of São Paulo at Almada (Lisboa, Portugal), a rock cut cave tomb hewn into the limestone was discovered. This hypogeum structure, named São Paulo 2, was used as a collective burial place from the end of the Neolithic until the beginning of the Bronze Age according to the recovered archaeological artefacts. Radiocarbon dating of two human bones indicated a Late Neolithic chronology: 2905–1950 cal BC (UBAR-629) and 2553–2137 cal BC (UBAR-630) with a 95% confidence level [112,113]. The uncovered human remains corresponded to a minimum of 254 individuals (both sexes), 131 adults (>15 years) and 123 non-adults [112,113]. Apart from one non-adult skeleton, all other elements were found disarticulated, due to the burial practice and post-mortem activities that took place inside the cave. Among the more relevant data are the detection of developmental abnormalities in tarsal bones [114] and three adult skulls with evidence of complete trepanation. These were performed by scraping method, and two of them, with signs of remodelling [115]. The detailed osteological study of this sample has been published by Silva [112,113].

Literature: Silva 2002, 2011, 2012 [112,113,115]

#### Romania

##### Schela Cladovei, Danube Gorges, Romania. Settlement with burials.

*Dušan Borić*

Schela Cladovei, in Romania, is a large, open-air site on an Early Holocene terrace adjacent to the River Danube, c. 67 km downriver from Vlasac. It is situated 7 km below the Iron Gates I dam of the Danube Gorges/Iron Gates area. Discovered in 1964, the first excavations were undertaken by the Romanian archaeologist Vasile Boroneanţ. An area of 127 m long strip was excavated along the riverbank up to 1989. From 1992 onward, a joint Romanian-British research project was co-directed by V. Boroneanţ and C. Bonsall. More recently, A. Boroneanţ has joined as the Romanian project partner. The site covers the Late Mesolithic and Early Neolithic occupation mainly. A large series of AMS 14C dates on animal and human remains places the Late Mesolithic occupation between c. 7200 and 6300 cal BCE, and the Early Neolithic occupation between 6000 and 5600 cal BCE. At least 75 burials, containing the remains of over a hundred individuals, have been excavated from the Schela Cladovei, most of them dating to the Late Mesolithic.

We report genetic data from two individuals with no further stratigraphic details.

Literature: Boroneanţ, V. et al. 1999 [116]; Bonsall 2008[117]; Boroneanţ, A. et al. 2014[118].

##### Băile Herculane – Peştera Hoţilar, Banat, Romania. Cave.

*Dušan Borić*

Băile Herculane (The Bath of Hercules), also known as Peştera Hoţilar (The Cave of Thieves), is situated deeper in the hinterland of the Danube Gorges area on the Romanian side, in the Banat region. The site was excavated in 1954–1955, and more intensively in 1960–1961, 1965, 1968–1970, and 1972. It is situated on the cliffs above the Cerna River, one of the Danube tributaries. The deposits of this cave encompass a sequence with Mousterian, Upper Palaeolithic, and Epipalaeolithic/Mesolithic levels (Dinan 1996; Nicolâescu et al. 1957; Mogoşanu 1978). Lithic typology is similar to the site of Cuina

Turcului I-II found on the banks of the Danube in the Danube Gorges, with a small assemblage of only c. 100 artefacts. Among the raw materials used were grey radiolarite, which dominates, followed by red radiolarite, with a presence of several pieces of the Balkan (yellow-spotted) flint. The absence of quartz is notable and among formal tools there are only several microliths (backed bladelet, trapeze, burin). There are numerous fish and mollusc remains, and mammal remains include bear, red deer, and beaver. A cooler environment has been modelled for the Early Holocene occupation phase in the cave on the basis of the presence of rodents and some floral remains. One charcoal measurement (Gr-16978) from Early Holocene levels provides a Mesolithic date: 11,490±75 BP.

We report genetic data from one individual with no further stratigraphic detail, dated to the Neolithic.

Literature: Nicolăescu-Plopșor et al. 1957[119]; Mogoşanu 1978[120]; Dinan 1996[121]; Boroneanţ 2011[122].

#### Russia

##### Afontova Gora, Krasnoyarsk, Southern Siberia, Russia. Flat grave.

*Mikhail Sablin*

Afontova Gora is a complex of archaeological sites located on the left bank of the Yenisei River near the city of Krasnoyarsk, Russia. The complex was first excavated in 1884 by the Russian archaeologist I.T. Savenkov. There are many burial sites from different epochs. Dating of the human bone SR-8482 around 5.3 kya uncal BP, corresponding to the Kuznetsk-Altai culture, Late Neolithic in the Siberian chronology. The sample was personally found by I. T. Savenkov in 1884. Previous aDNA analyses were published by Raghavan et al. [123] and Allentoft et al. ([17], RISE553 and RISE554).

One new sample was analysed: NEO102, SR-8482, dated to 5280±30 uncal BP (UCIAMS-14767).

Literature: Raghavan et al. 2014[123]; Allentoft et al. 2015[17].

##### Bazaiha village, Krasnoyarsk district, Southern Siberia, Russia. Cemetery.

*Mikhail Sablin*

Bazaiha Late Bronze Age cemetery is located on the right bank of the Yenisei River near the village of Bazaiha, Russia. As early as 1883-1885 Russian archaeologist I.T. Savenkov excavated a few flat graves at that cemetery. In one of them, a bronze Celt was discovered. Dating of the human tooth SR-8461 around 2.9 kya BP, corresponding to the Karasuk culture, Late Bronze Age in the Siberian chronology. The sample was personally found by I.T. Savenkov. The material has not been published before.

One sample was analysed successfully, NEO070, SR-8461, individual 4, dated to the Late Bronze Age.

##### Borovyanka XVII, Omsk region, Russia. Cemetery.

*Olga Poshekhonova*

The Borovyanka XVII burial ground is located on the bank of Batakovo Lake on the left side of the Irtysh River near the Borovyanka village, Bol'sherechinsk district, Omsk region. The landscape zone is Northern forest-steppe. The site was excavated by L.I. Pogodin in 1999-2001. While description of several burials was published no dedicated publication has been made so far. The site consists of burials dated from the Eneolithic to the Middle Ages. The most numerous are Late Bronze burials which are attributed to Chernozerye culture type [124,125]. About 20 burials which are supposed to be a single Eneolithic complex were dated by the middle of 3rd millennium BC [126]. Most of the burials are single but a few double and collective (up to 8 skeletons) ones were also excavated. The skeletons lay stretched on their back with bent legs and their heads oriented to the North or more rarely to the North-East.

Burials of non-complete skeletons or single skulls as well as a few cases of partly burned skeletons were reported. It was argued that high variability of burial traditions in Borovyanka XVII resulted from multi-ethnicity of Eneolithic society related to the. Based on characteristic artifacts found in graves including the ceramics similar to those of the Ekaterininskaya culture, the Keltiminar fish-like and petal-like pendants, and arrow points the closest analogies can be found in the Khutor Bor IV and Okunevo V-VII burial grounds from the Irtysh region and the burial ground on the Bol'shoy Andreevsky Lake from Tobol region. Three samples are reported here:

NEO080 – grave 83, skeleton E, museum number IPDN 78-19. Morphologically female of 55+ years old. The skeleton was found in a collective grave which consists of 8 or 9 individuals. Only the skull, femurs, left tibia and partially the bones of the foot have survived. Under the skull, pelvis and ankle bones ochre layers up to 3 cm were recorded. The skeleton lay stretched on its back with its head to northeast. An Eneolithic dating was argued and is supported by the 14C date of the skeleton.

NEO081 - grave 69, museum number IPDN 78-52. Morphologically male of 55+ years old. The skeleton was found in a single disturbed burial. The postcranial bones and mandible lay in the southwest part of the grave while the skull lay separately in the northeast part. A fragment of non-ferrous metal plate was found near the skull and fragments of the Chernozerye and Ekateriniskaya ceramics were found in the grave filling. The grave was dated to the Bronze Age and attributed to Chernozerye culture [126]. The direct dating of the sample (3588±29, UBA-39953) supports a Bronze Age date.

NEO083 - grave 36, Skeleton B, museum number IPDN 78-29. Morphologically male 55+ years old. The burial was found beneath the burial of a child of 7-9 years old (skeleton A). Both skeletons lay stretched on their backs with their heads oriented to the northeast. The long bones and the skull of the child and the adult were partly burned. Also in the child burial the fragmented and partly burned bones of an adult (possibly female) were found. The similarity in funeral ritual of the burials and shell beads found in the graves suggests that all burials belong to the same Eneolithic population and were done in a short period of time [126]. The 14C date of the adult male was UBA-39955, 5433±37 uncal BP, supporting a Neolithic date.

Literature: Khvostov 2001[126]; Polevodov & Shestobitova 2006[124]; Shestobitova 2007[125]

#### Dolgoe Ozero (Long lake), Krasnoyarsk, Russia. Settlement and graves.

*Veselovskaya Elizaveta V. and Vasilyev Sergey*

The ancient settlement near the Long Lake is located on the western outskirts of the Kansk city, Krasnoyarsk Territory. From the end of the XIX century and until the beginning of the 1960's, the sand dune in which the burial ground was discovered served as a quarry for the extraction of sand, as a result of which the archaeological site was not completely preserved.

The first archaeological surveys at the Long Lake were carried out in 1909-1910 by the archaeologists of Kazan University, E.A. Popov and A.E. Ermolaev, who collected material here and dated the site to the Early Bronze Age - Iron Age.

In September 1958, in the settlement near the Long Lake, stationary excavations were carried out by the Leningrad archaeologist G.A. Maksimenkov [127]. Two single graves and one double were discovered in the northwestern part of the quarry, which Maksimenkov dated to the early Neolithic, approximately the Isakovo-Serovo period. The graves are located in parallel, forming one continuous row. The skeletons are oriented with their feet towards the lake, not corresponding to the cardinal points. Judging by the surviving remains of the buried, the burial ground has a generic nature, rather than a family one. A piece of ochre was found by the foot bones of one of the skeletons; a flake was inserted into the hand of another buried. No other archaeological finds were found in the graves.

The main archaeological material from the locality was obtained as a result of collections on its surface, mainly in the northern part of the site:

• 1. Up to eleven types of stone arrowheads.

• 2. Scrapers on flakes of random shapes.

• 3. Two micro chisels.

The nature of archaeological material finds analogies with Western regions, largely repeating the finds in the areas of the Krasnoyarsk district.

In anthropological terms, the population that left the burial ground was interpreted as characterised by a combination of Caucasoid and Mongoloid traits. The only whole skull (sampled here as NEO 292) is massive, with a very high and wide flattened face. Zygomatic bones are very large, canine fosses are very weakly expressed. The lower jaw is massive with expanded rami. The skull was determined to belong to an adult male, ca 40-45 years old. It was dated to 5448±41 uncal BP, UBA-40027.

Literature: Maksimenkov 1964[127]

#### Fofonovo, Baikal region, Russia. Cemetery.

*Vasilyev Sergey & S. B. Borutskaya*

For the first time, skulls from the Fofonovsky burial ground were published by G.F. Debets [128]. At his disposal were only 4 skulls from the excavations of M.M. Gerasimov of 1934 and 1936, which discovered over 40 burials of the Eneolithic and developed bronze. Studies of the Fofonovsky burial ground were continued by A.P. Okladnikov, who headed the Buryat-Mongolian expedition in 1948 - 1952. In 1959, M.M. Gerasimov, as part of the Irkutsk expedition under the leadership of M.P. Gryaznov, the excavations of the Fofonovsky burial ground were resumed.

The continuous excavation in the eastern part of the burial ground made it possible to open 41 burials belonging to three stages or cultural complexes. These excavations made it possible to make valuable stratigraphic observations, which contributed to the fact that all burials, regardless of the availability of accompanying dating equipment, could be divided into three stratigraphic and chronological groups - Kitoysky, Glazkovsky and, possibly, Shiversky.

The most interesting materials were given by the burials of Kitoyan association - jewellery made of shells and teeth of wild boar and maral, bone daggers with silicone inserts, various stone and bone tools found in the grave usually in a bundle or in heaps. Burials are characterised by abundant backfilling with ochre. In 1988, at the Fofonovsky burial ground, excavations by archaeologist V.P. Konev. For one of the burials, a radiocarbon age of 6600 ± 100 years is given. Already in our time, the excavations of EA Zhambaltarova in 2007 - 2008 provided paleoanthropological material from several other Kitoyan burials.

Three individuals are reported here, all dated to the Early Neolithic, ca 6800-7000 uncal BP (osteological determinations are not available):

NEO199, burial 10

NEO200, burial 2(2)

NEO201, burial 7(5)

Literature: Alekseev and Gokhman 1984[129]; Gerasimova 1992[130], Vasilyev et al. 2010[131].

#### Golubaya Krinitsa, Middle Don, Russia. Cemetery.

*A.M. Skorobogatov*

The site was discovered in 2011 by Valery Berezutsky [132]. The burial ground is located on the right bank of the Black Kalitva River (a tributary of the Don River), near its mouth. Excavations were carried out in 2015-2016 under the leadership of Andrey Skorobogatov. A total of 18 burials were studied (single, paired and collective).

The burials were in rectangular pits, characterised by orientation to the south - southeast and southeast. The position of the buried is stretched out on the back, with arms located along the body. The bones are sprinkled with red ochre.

The burials were accompanied by inventory: fossil sea shells, Unio shells and products from their wings, bone decorations (wild boar fangs, beaver teeth and groundhogs), bone tools, a copper product, flint tips, flint knives, and ceramics.

The complex finds analogies in the Mariupol-type burial grounds widespread in the territory of modern Ukraine (Mariupol, Nikolsky, Lysogorsky, Yasinovatsky burial grounds), and can date back to the 6th millennium BC.

Six samples were analysed, with datings ranging ca 6400-6700 uncal BP, corresponding to c. 5000-5400 cal BC:

NEO113, kurgan 10, burial 10

NEO204, burial 4

NEO207, burial 7 skeleton 2

NEO209, burial 7 skeleton 4

NEO210, burial 8

NEO212, burial 10

Literature: Berezutsky et al. 2011[132].

##### Itkul (Bol’shoy Mys), Altai region, Russia. Cemetery.

*Marina Rykun & Vyacheslav Moiseyev*

The Itkul’ burial ground is located in the Altai lowland area between Pleshkovo and Vershinino villages, Zonalnoe district, Altai region. The site is situated on the top of the Bol’shoy Mys (Big Cape) of Itkul’ Lake. In the Neolithic-Eneolithic times the cape would be separated by water from the mainland.

16 burials arranged in 3 rows were excavated by B.Kh. Kadikov in 1962. Earlier one or four skeletons were excavated by a local amateur. All buried lay stretched on their back, heads oriented to North-East. The burial inventory is scarce and found mostly in the male burials. Few pendants made of animal teeth were found in both male and female burials. No ceramics were found. At present time the burial ground is attributed to Bol’shoy Mys Eneolithic culture [133].

NEO063 - burial 2, morphologically male of 30-35 years old [134]. Burial inventory included a stone axe, flake and polished bar. Several deer and bear teeth were found in the neck zone of the skeleton. It was dated to 6656±34 uncal BP, UBA-39940.

NEO064 - burial 8 (doubled with burial 7), morphologically female of 25-35 years old [134]. Burial inventory included stone rod of the fishing hook, several beaver and marmot teeth and pendants made of deer and roe deer, two gypsum «roses». The skeleton was dated to 5744±41 cal BP, UBA-39941.

Literature: Dremov 1980[134]; Kiryushin et al. 2000[135]; Kiryushin 2002[133].

#### Karavaikha, Vologda region, Russia. Cemetery.

*Alexandra Buzhilova*

The site of Karavaikha was located on the right bank of the Eloma River, which flows into the Lake Vozhe from the west (Kirillovsky District, Vologda Region). Since 1938, during seven field seasons, this site was excavated by Bryusov [136]. A total of 38 burials were opened. Now bone remains from the excavations of 1938 and 1939 are housed in the Museum of Anthropology of Moscow State University (22 burials).

The burials discovered in 1938 were mostly destroyed. Nevertheless, it was possible to establish that there were remains of adult individuals only, most of them buried on their backs with the head oriented to the south. In one burial (No. 7, #8623), the remains were in a sitting position. All burials were single, except for grave No. 2, in which there were remains of two individuals. Part of the skeletons was covered by ochre (No. 2-7, 9-11).

Excavations of 1939 gave a greater number of burial options. So, in addition to human remains on the back (No. 12, 15-18), there was a person lying on the right side (No. 21) and one individuals was in a sitting position (No. 13, #8761). The bodies of the buried were oriented with their heads to different parts of the world (to the south, north and west). In addition to adults, a teenager (No. 17) and two children (No. 15, 18) were buried. Some of the skeletons were covered by ochre (No. 12, 14, 15, 18 and 22).

Bryusov stressed that burials 1 (#8624), 3 and 11 (#8625) could be the earliest in this part of the excavation; and the rest of the burials were from later time. In the early years of the excavation, Bryusov established the chronological difference in the burial complex of Karavaikha, dating it from the late Mesolithic to the Eneolithic [136]. This was partially confirmed by attempts of radiocarbon analysis (#8624 burial 1 - 8200 ± 50 GIN-7173, #8622 burial 6 - 4420 ± 50 GIN-7172 and #8623 burial No. 7 - 6880 ± 90 GIN-7176) [137]. In the present project, the datings were concentrated to around 7700-7500 uncal BP. This applies also to burial 6, which was redated (Supplementary Data IV).

Anthropological studies have shown the heterogeneity of the series, which could be associated with use of the burial complex for a long time; researchers also note a mixture of craniological complexes [138,139].

Sampled individuals:

NEO559: 8622 – burial 6, M, Adultus

NEO556: 8623 – burial 7, M?, Adultus2-Maturus, healed skull trauma

NEO555: 8624 – burial 1, M, Maturus2-Senilis

NEO557: 8625 – burial 11, M, Maturus2-Senilis,  healed skull trauma

NEO558: 8761 – burial 13, M, Adultus 1

NEO560: 8762 – burial 20, M?, Maturus 2, healed skull trauma

NEO561: 8763 – burial ?, F?, Maturus 2- Senilis

Literature: Akimova 1953[138]; Gerasimov 1955[139]; Bryusov 1961[136]; Utkin & Kostyleva 2001[137].

#### Kostenkova Izbushka, Altai region, Russia. Cemetery.

*Marina Rykun & Vyacheslav Moiseyev*

The Kostenkova Izbushka burial ground is located on the opposite shore to the Itkul' site of Itkul' Lake. The site was excavated by Y.F. Kiryushin in 1981. Two of three excavated burials were attributed to the Eneolithic Bol'shoy Mys culture.

NEO065 - burial 5, morphologically male of 30-40 years old. The individual was killed by an arrow shot from a close distance. The bone arrow found at the head base destroyed several frontal teeth, went through the larynx and stuck at the cervical vertebrae. The burial inventory included fragments of ceramic vessels and two beaver maxillas. The skeleton was dated to 4516±33 uncal BP, UBA-39942.

Literature: Kiryushin et al. 2000[135].

#### Kumyshanskaya cave, Ural, Russia

*Pavel Kosintsev and Yu. Serikov*

This karst cave is located on the bank of the Kumysh River in the Middle Urals (57.58 N 58.20 E). The entrance to the cave measuring 3 x 4 m is located at an altitude of 4 m from the river level. The length of the cave is 9 m, the width is 2 m, and the height is up to 5 m. Excavations were carried out in 1998-2000. The excavation area is 17 m2.

The cave deposits consist of 4 layers (from top to bottom): a layer of humus, a layer of black sandy loam, a layer of brown loam, a layer of brown clay. In layers 2 and 3, artefacts from the Middle Ages, Iron Age, Bronze Age, Eneolithic, Neolithic and Mesolithic were found. Upper Palaeolithic artefacts have been found in the upper horizon of layer 4. In layers 1, 2 and 3, bones of a mountain hare (*Lepus timidus*), squirrel (*Sciurus vulgaris*), beaver (*Castor fiber*), wolf (*Canis lupus*), marten (*Martes martes*), brown bear (*Ursus arctos*), elk (*Alces alces*), reindeer (*Rangifer tarandus*) (about 2000 specimens) were found. In layer 4, bones of mountain hare (*Lepus timidus*), bobak (*Marmota bobak*), wolf (*Canis lupus*), arctic fox (*Vulpes lagopus*), mammoth (*Mammuthus primigenius*), horse (*Equus ferus*), woolly rhinoceros (*Coelodonta antiquitatis*), reindeer (*Rangifer tarandus*), bison (*Bison priscus*), saiga (*Saiga tatarica*)(more than 500 specimens) were found.

A radiocarbon date of 12,430±260, SOAN-4846, was obtained from the mammoth bone from the upper horizon of layer 4, and a radiocarbon date of 33,670±300, OxA-10929 was obtained from the lower horizon of layer 4 from the woolly rhinoceros bone.

In the part of the cave farthest from the entrance, at a depth of -0.3 -0.4 m, there was a cluster of human bones (more than 100 specimens). It had an oval shape measuring 0.3 – 0.6 m x 2.0 m. The bones were covered with ochre. Bones of 2 foetuses 7 – 9 months old, 2 newborn individuals (skull bones, pelvis, ribs, tibia, femur, humerus and radius bones) and bones of 3 adult individuals (skull fragments, ribs, metapodia, phalanges), probably women: 2 adults (over 30 years old) and one young (20 – 25 years old) were found. A radiocarbon date of 4 635±85 uncal BP, SOAN-5155, was obtained from the bones of adult individuals.

In this project, one sample was analysed for DNA: NEO687, 910/2456, Kumyshanskaya 2. This is a cranial fragment from a small child (inf I). It was dated to 4922±47 uncal BP, UBA-40107.

Literature: Serikov 2004[140], 2013[141].

#### Ksizovo 6, Lipetsk, Russia. Settlement and burial ground.

*R.V. Smolyaninov & S.B. Borutskaya*

The archaeological complex of Ksizovo 6 was investigated by R.V. Smolyaninov in 2004-2005, two excavations with a total area of 253 sqm [142]. It combines the features of a settlement and a burial ground of the Neolithic-Bronze Age periods. Located near the village of Ksizovo, Zadonsky district, Lipetsk region at the confluence of the river Don with its right tributary river Snova. The site is located on a flat surface with a height of about 5.5-6.0 m above the river edge at the foot of a steep ledge of the root bank of the Don River with a height of 20-25 m.

As a result of the excavations, a ceramic collection was obtained from several hundred vessels of the Neolithic-Bronze Age epochs dating from the second half of the VI - II millennium BC. The site has cultural strata varying from 1 to 2 metres thick. In both excavations, burials were revealed. In total, the remains of 17 individuals from 15 burials were studied from them: 6 men, 5 women, and 6 children. Anthropological definitions made by S.V. Vasilyev and S.B. Borutskaya [143].

Based on the analysis of the studied burials, we believe that the flat burials were left by the population of three archaeological cultures. Elongated burials with a northern and northeastern orientation date back to the Neolithic era and to the first quarter of the VI millennium cal BC. The Srednestogovskoy culture of the Eneolithic era includes elongated and shortened burials, which date to the 1st half of the 5th millennium BC (cal. BC). To the Catacomb culture of the Bronze Age, middle of III millennium BC, one crouched burial 3 from excavation site 2 belongs.

Four samples from three burials were analysed:

NEO172, burial 4. Neolithic, dated to 6181±47 uncal BP, UBA-39984.

NEO173, burial 2 skeleton 1. Sredny culture, dated to 5789±34 uncal BP, UBA-39985.

NEO174, burial 2 skeleton 2. Sredny culture, dated to 4865±33 uncal BP, UBA-39986.

NEO175, burial 3. Catacomb culture, dated to 4032±34 uncal BP, UBA-39987.

Literature: Lavrushin et al. 2009[142]; Vasilyev et al. 2018[143].

##### Mergen’ 6, Tumen region, Russia. Settlement

*Olga Poshekhonova & Svetlana Skochina & Dmitri Enshin*

The Mergen’ 6 multilayer settlement is located on the bank of Mergen’ Ishim district, Tumen region, Russia. The landscape zone is defined as the forest-steppe of West Siberia. The site was excavated by V.A.Zakh in 1990; in 2002, 2004 and 2013 by S.N. Skochina and D.N. Enshin. The Neolithic settlement consists of 13 buildings arranged in a semi-circular complex adjusted to the cape form [144]. Based on dating of different materials including (animal horn and bone, human bone, ceramics of Boborykino and Koshkino types, and ceramic soot) the dating of the settlement was defined as the last quarter of the 7 millennium BC (Piezonka et al, 2020) which corresponds to the Early Neolithic period.

NEO072 (museum number IPDN 116-1) - morphologically female of 30-45 years old. A fragment of the female skull was found in hut 15 near a hearth together with a tool for making bast shoes, and several crushed animal bones. On the opposite side of the hearth a skull of a dog was found. This finding was identified as the evidence of ‘building sacrifice’, which is characteristic to Koshkino and Boborikino cultural traditions [144]. The earlier obtained dating OxA-33489 7355±40 BP, 6361-6086 cal BCE (δ15N+15.3‰, δ13C -20.15‰) is close to the dating of based on animal bones and ceramic soot found in the hut; (Poz-94074 7060±50 BP and Poz-98334 7270 ± 50 BP) [145].

NEO073 (museum number IPDN 116-2) - a child up to 1 year old (genetically female). The burial of the child was found in hut 21. The pit grave (0,55×0,25×0,1 m) was dug into the pit floor of the house by 0.1 m. The skeleton of good preservation lay on its stomach with the arms stretched along the body oriented from North to South. No artefacts were found in the grave. The sample was dated to 7317±62 uncal BP.

Both samples have been genetically studied earlier [146].

Literature: Enshin 2012[147]; 2014[144]; Piezonka et al. 2020[145]; Narasimhan et al. 2019[146].

#### Minino I and II, Vologda region, Russia. Cemetery.

*Alexandra Buzhilova*

Minino I and Minino II are two Mesolithic-Neolithic burial complexes, which were excavated on the bank of Kubenskoe Lake during a few seasons from 1996 by an archaeological expedition of the Institute of Archaeology, RAS. The Kubenskoe Lake is a large water-pool of glacier origin situated to the south from latitude 60° North in European Russia (Vologda region). Around 170 archaeological sites have been registered in the area (from the Mesolithic to Early Mediaeval time). There were not too many convenient places to settle near the marshy sides of the lake; therefore, those suiting habitation were settled repeatedly. The Mesolithic materials originate from 10 sites and 4 of them are from the archaeological complex Minino, which was represented both by artefacts and 2 of them represented by human remains [148].

The burials are located on the elevated edge of the modern floodplain terrace. The distance between the excavations of these two sites is 230 m. The orientation of the graves prevails along the edge of the terrace, along the southeast-northwest line, parallel to the shore of a lake.

Minino I and Minino II burials were revealed in the Mesolithic cultural deposit. These are several groups of asynchronous burials (among them 22 single burials, 5 paired burials and 2 triple ones) [149]. Some burial goods point toward parallels to the local Veret'e and Butovo archaeological cultures. However, the majority of objects and a radiocarbon analysis suggest a wide chronological span from the Mesolithic until the Early Neolithic [150,151].

The anthropological materials (39 individuals), which were studied in bioarchaeological context, provided the basis for a detailed reconstruction of lifestyles. The indicators of physical activity demonstrate the general picture of occupational stress matched to known cases of hunter-gatherers. A series of male skulls from Minino indicates undoubted analogies with the synchronous population of the nearest territories of northern-eastern Europe and partly represented anthropological complexes of Late Palaeolithic and Early Mesolithic hunters. In the early stages of the development of the region, the population demonstrates high life expectancy; and the mean age at death decreases in the final chronological period [152]. Isotopic analysis of human bones gives the data of diet changing over time [153]. These data could well mirror some biological and cultural changes over time.

Four samples were analysed, three of which dated to the Mesolithic and one to the Late Bronze Age:

NEO536, Minino II, burial 2 skeleton 2. Adult female?, dated to the Mesolithic

NEO537, Minino I, burial 3. Adult male, dated to the Mesolithic

NEO538, Minino I, burial 16. Adult male, dated to the late Bronze Age

NEO539, Minino I, burial 20, Adult male, dated to the Mesolithic

Literature: Makarov 2001[148]; Suvorov & Buzhilova 2004[149]; Wood 2006[153]; Suvorov 2007[150]; Wood et al. 2013[154]; Buzhilova 2016[152].

#### Okunevo 5 and 7, Omsk region, Russia. Cemetery.

*Olga Poshekhonova*

The Okunevo 5 and 7 burial grounds are located on a narrow cape formed by the Tara and Irtysh rivers; Murmantsevo district, Omsk region, Russia. In terms of the landscape, the territory belongs to the subtaiga zone of Western Siberia. Only ground graves and ritual places but no settlements were found in the vicinity.

The multi-layer site of Okunevo 5 consists of a sacrificial place and a cemetery. The site was found by V.A. Mogilnikov and was excavated by B.A. Konnikov, V.A. Mogilnikov and V.I. Matyushchenko in 1976, 1981 and 1987-1989 respectively. The cultural layers date from the Neolithic to the Middle Ages.

The Okunevo 7 burial ground directly adjoins the Okunevo 5 site. It was excavated by A.I.Petrov, V.I. Matyushchenko and V.A. Mogil'nikov in 1980's and 1990's. Burials dating from Neolithic to the Middle Ages were studied at the site.

For genetic analysis, Eneolithic-Early Iron Age samples (the end of the 3rd – first part of the 2nd millennium BP) from both sites have been taken. The close location of these graves suggests that all of them belong to a single cemetery which was artificially divided by its excavators in two sites. The graves were arranged in rows divided by at least six deep ditches of uncertain dating. All samples belong to the first (earliest) group of burials but their cultural attribution was not fully defined. At least a part of the graves of this group of burials belongs to the Ekaterininskaya culture [155,156]. Because most artefacts were found between graves they can obviously be defined as funeral offerings.

Three samples were analysed, all dated to the Eneolithic, ca 4500-4700 uncal BP:

NEO068 - burial 62, morphologically male 20-21 years old. Grave size is 170x75 cm, oriented from North to South. The artefacts found in the grave include 40 horn beads, a bronze pendant and a piercer, and a ceramic tetrahedral flat-bottomed vessel, decorated by comb ornament [155].

NEO077 - burial 66, morphologically male, 40-45 years old. Grave size 160x80 cm, the skeleton oriented from North to South lay stretched. The skull lay on a sand "pillow" 8-10 cm high. Ochre spots were found near the skull and between the femurs. In the North-East part of the grave a fragment of a wooden cylindrical item with a diameter of 5 mm and a length of 3 cm was found [155].

NEO079 - burial 69, morphologically female, 18-20 years old. The grave pit was traced along a spot of humus soil. It had a rectangular shape, 220x90 cm and a depth of 80 cm. The skeleton was covered with wooden chopped blocks. Their thickness varied from 25 cm to 10-15 cm, the width was 10-20 cm. A thin layer of ocher was poured over the entire area occupied by the bones. A poorly preserved skeleton of a newborn child lay between the tibias of the adult. A large quartzite scraper was found near the right arm of the adult, a scraper-knife of the same material was on the left. A ceramic ball and two fragments of ceramics are also found here [155].

Literature: Matyushchenko & Polevodov 1994 [155]; Petrov 2014[156].

#### Omskaya Stoyanka (Omsk settlement), Russia. Settlements and cemeteries.

*Olga Poshekhonova*

The complex of archaeological sites usually referred to as the 'Omsk settlement' is located in the city of Omsk on the left bank of the Irtysh River, Russia. In the Neolithic the site was on a small island. From the landscape point of view this is the forest-steppe zone of Western Siberia. The site includes several settlements and burial grounds. The burial complexes are dated to the Neolithic and Bronze Age.

The site was discovered in 1918 by C.A. Kovler. Since then a number of excavations have been done [157]. In 1998 B.A. Konnikov excavated five burials. Three of them were severely destroyed while two were not disturbed. These grave pits partly overlapped each other.

NEO075 - (grave pit1, skeleton 2) morphologically female of 20-25 years old. Few artefacts including two wolverine drilled canines, two small stone knives, and a flake were found in the grave. Direct carbon dating has been done earlier - 6656±50 BP (UBA-23603). A new date was almost identical, 6654±40 uncal BP, UBA-39947.

NEO078 - (pit grave 2, skeleton 3) morphologically male of 25-30 years old. The pit grave was 10 cm deeper than grave 1 and partly overlapped it. No artefacts were found in the grave. Direct dating was 6560±49 uncal BP, UBA-39950, i.e. slightly younger than NEO075.

Both skeletons lay stretched in anatomical order with their heads oriented to the Irtysh River from South to North or North-East. Archaeologically the burials were dated to the 4th millennium BC [158]. Albeit no cultural attribution has been proved for these burials, some similarity with Sopka 2 was nevertheless declared [159].

Literature: Metel' 2016[157], Konnikov 1998[158], 2013[159].

#### Ostrov 2, Tumen region, Russia. Cemetery.

*Olga Poshekhonova & Svetlana Skochina*

The Ostrov 2 Eneolithic sanctuary is located in the Iset' River basin (Tobol tributary); Yalutorovo district, Tumen region, Russia. The site was excavated by A.V. Matveev in 1995.

The sanctuary consists of several clusters of pits. In two pits human remains were found (pits 3 and 4). The site is attributed to the Eneolithic comb ceramics of Shapkul' culture of local South Ural origin [160].

NEO076 – child, 7-8 years old. The child skeleton was found in an oval pit 3 (3x0,98 m) which is one of the central pits of the sanctuary. Close to the child remains a mandible fragment of a female, 18-20 years old, and fragments of femur and tibia which possibly belong to the same individual were also excavated. The only artefacts found in the grave are a scraper made of pink flint and a jasper blade [160]. The sample was dated to 4225±29 uncal BP, UBA-39948.

Literature: Matveev et al. 2015[160].

#### Pad' Tokui, Krasnochikoysky region, Russia. Cemetery.

*S. V. Vasilyev & S. B. Borutskaya*

The Neolithic burial grounds of Zjindo and Pad' Tokui are located on the territory of the Krasnochikoysky region of Western Transbaikalia, in the Chikoi river basin (the eastern part of the Selenga river basin and Lake Baikal). Now mountain taiga biotopes are common there. Excavations were carried out mainly in 2004-2005 by the Chikoy archaeological expedition of the Trans-Baikal State University under the direction of M.V. Konstantinov [161].

Pad' Tokui burial ground is located in the Daur Highlands on the left bank of the Menza River, 7 km south-west of the village. Menza on a gentle slope of a hill, on the site of an old abandoned arable land. This place rises more than 900 m above sea level. The monument is a flat burial ground. In 2005, two burials were discovered and excavated.

One sample was analysed, from burial 2 (Zhindo 6/NEO116). It was dated to 7378±72 uncal BP, UBA-33757.

Literature: Konstantinov et al. 2005[161]; Vasilyev et al. 2018[143].

#### Peschanitsa, Archangelsk region, Russia. Burial(s).

*S. V. Vasilyev & S. B. Borutskaya*

In 1986, during archaeological research of the Mesolithic site in the Peschanitsa area, located 800 m from the modern shore of Lake Lacha (Kargopolsky district of the Arkhangelsk region), S.V. Oshibkina discovered the skeletal remains of an ancient person. In a quarry that destroyed the site, a skull was discovered (object 1). Near this location, in the excavation, bones of the thoracic region, pelvis and fragments of bones of the upper extremities were found. At a distance of 1.5-3 m, at a depth of 60 cm from the modern-day surface, a pit with only leg bones was found (object 2). The bones lay on a thin layer of white clay and were abundantly covered with red ochre. They were accompanied by fragmented animal bones and three flint flakes. Small coals are marked in the clay layer.

All this, according to the author of the excavation, suggests that the burial of the legs was not accidental. The skull and the other bones may possibly belong to one and the same person. The appearance of the material remains of the Peschanitsa site and the features of the funeral rite make it possible to attribute it to the Mesolithic culture of Veretye, as well as the Popovo burial ground, and several settlements located near Lacha and Vozhe lakes.

The skull, sampled here as NEO202, was previously dated to 10728±50 BP uncal, UBA-41633, and the leg bones to 9890±120, GIN-4858 [18]. A new dating of the skull resulted in the date 10030±56 uncal BP, UBA-40011. This makes it more likely that the skull and leg bones are contemporary.

Literature: Oshibkina 1997[162]; 1998[163]; 2017[164]; Mamonova 1995[165]; Saag et al. 2021[18].

#### Pogostishche I, Vologda region, Russia. Cemetery.

*Alexandra Buzhilova*

The site was discovered by A. Ya. Bryusov in 1938. A scientific paper was published in 1951, in which the author mentions that the site of Pogostishche 1 is located opposite the confluence of the Ukhtomka River and the Modlona River near the village of Pogostishche, Charozersky district, Vologda region [166]. Only one skull of a juvenil woman is stored in the Museum of Anthropology, Moscow State University.

Modern researchers note that the site is located on the floodplain terrace – a relatively high and dry place. There are very few such places in the basin of Lake Vozhe, which explains the high density of archaeological sites in the area. Nowadays, almost two dozen archaeological sites are known. They were discovered and partially explored in different years by A. Ya. Bryusov, S. V. Oshibkina, N. A. Makarov, and N. V. Kosorukova. Since 2002, the later Stone Age sites have been systematically studied by the archaeological expedition of the Cherepovets State University [167].

Information about the site of Pogostishche 1, presented by Bryusov, is extremely fragmentary. There is no information about the methodology of excavation; only a schematic plan of the location of the excavation is presented, subsequently published by Oshibkina [168]. Bryusov notes that the artefacts were in the sand. Oshibkina, who checked the stratigraphy of the site, writes about the same. Bryusov reports that the findings were not numerous: in the excavation there were found tools made of bone and stone, including 6 scrapers from large flakes, a fragment of a leaf-like flint tip of a dart, a fragment of a knife-like plate, 2 flint retouchers, 4 nucleuses, half of the stone disc-shaped top of the club with a hole in the middle, a bone awl 16.5 cm long, a fragment of the second awl, a fragment of a bone harpoon, a fragmented horn clutch of an axe, a bone arrowhead with a biconical head [166]. Also animal bones, flakes and fragments of flint, slate and quartz were found.

Bryusov suggested that the site of Pogostishche 1 should be dated no later than the end of the 4th millennium BC. According to Oshibkina, the site of the Pogostishche I can be compared with the site of Lower Veretie I (to the conditions of the cultural layer, as well as by the characteristics of the finds), and should be dated to the Boreal period [169]. According to Makarov, the site does not represent a pure Mesolithic complex, since pit-comb ceramics and smooth-wall ceramics of the Iron Age were found in the pits in the upper layer [170].

A sample from the single skull (NEO554) was analysed here. The skull was determined as belonging to a young female individual. It was dated to 7472±42 uncal BP, UBA-40069.

Literature: Bryusov 1951[166]; Oshibkina 1983[169]; Makarov et al. 2001[148]; Oshibkina 2006[168]; Kosorukova 2012[167].

#### Protoka, Novosibirsk region, Russia. Cemetery.

*Veselovskaya Elizaveta V. & Vasilyev Sergey*

The Protoka burial ground is located in the Kyshtovsky district of the Novosibirsk region. Under one of the 9 mounds from the Early Iron Age, 14 Neolithic burials were revealed, in which 26 people were buried.

According to the state of the bones in the Neolithic part of the burial ground, scientists conclude that the burials are secondary. Apparently, the dead were first kept in an open place, possibly in the winter. This may explain the absence of some bones that animals could pull apart. Then the dead were reburied in groups of up to 8 people, laying everyone with their heads to the northwest.

N.V. Polosmak, the excavator of the Protoka burial ground, attributed its Neolithic part to the Middle Irtysh culture. Later V.I. Molodin attributed it to Verchne-Obskaya Neolithic culture.

The study of craniological material has been interpreted as indicating the presence of Mongoloid complex features.

One individual was sampled (NEO309, burial n19).It was dated to 5575±36 uncal BP, UBA-40035. Data for morphological age and sex is lacking from this individual.

Literature: Troitskaya & Novikov 2004[171]

#### Sakhtysh, Ivanovo region, Russia. Settlements and cemeteries.

*Veselovskaya Elizaveta, Vasilyev Sergey & Kostyleva Elena*

Sakhtysh, a unique complex of 15 archaeological settlement and cemetery sites, is located in the centre of the Russian Plain in the Teykovsky District of the Ivanovo Region at the source of the Koyki (Kiyki) River from the Sakhtysh Paleolake. Chronological range of the multilayer settlements: early Mesolithic - Middle Ages; burial grounds: Neolithic - Eneolithic.

Burials of that time were found at the Sakhtysh I, II, IIa, VII and VIII sites. They were inhumated into the ground.

Neolithic burials belong to the Lyalovo archaeological culture (Middle Neolithic). People who came from Northern Europe (Karelia, Finland) took part in the formation of this culture. This is confirmed by specific features of material culture and anthropological type of sculls (Kostyleva, Utkin. 2019). Burials of the Eneolithic epoch belong to two phases of the Volosovo culture. Burials of the early phase possess some links with Eastern Baltic region, such as numerous amber decorations from the area of Lake Lubanos (now in Latvia) in the male burials, as well as the anthropological type of people characteristic of the Baltic zone. In the burials belonging to the late phase of the Volosovo culture, the eastern direction of cultural connections may be traced, in particular, the presence of serpentine stone ornaments in female burials, deposits of which are known near the South Urals. Perhaps this is due to exogamous marriages [172–174].

The dating of the Ljalovo burials is about 5000 years BC, the earliest Volosovo is c. 4000 BC, and the latest c. 3500 years BC [175,176].

##### Sakhtysh II

The archaeological site Sakhtysh II was discovered by Prof. D.A. Krainov in 1962. Excavations of the Sakhtysh II site were carried out during 14 field seasons (1962-1964, 1966, 1978-1982, 1984-1987, 2001). During this period, 23 burials were excavated, located on an area of 1500 square metres. The burials are clearly subdivided into two cultural and chronological groups: Lyalovo (4 burials) and late Volosovo ones (19 burials).

The Lyalovo burials have a North-South orientation with slight deviations. The burial ground of the Volosovo culture does not have a clear structure; the orientation of the burials is different. Single burials prevail, but there are also some group burials: in two graves there were 4 people each and in another one - 18. In a number of burials, traces of violent death were noticed.

Most of the ornaments found in the Volosovo burials at the Sakhtysh II site were made of bone. The second largest type of ornaments is pendants made from animal teeth. They were mostly from dog teeth. Amber ornaments are scarce.  Basically, these are pendants, several buttons and rings. There are also stone pendants in burials.

In addition to human burials, three burials of dogs were discovered. Burials of dogs are chronologically linked to the Volosovo layer. There is reason to consider dog burials as a kind of ritual.

Three samples were analysed from this site:

NEO158, burial 4; Burial 4 (excavation of 1963) was collective: it contained three males (skeletons # 2, 3, 4) and one female (skeleton # 1). The male skeletons had no skulls, while the female skeleton had a skull. Thus, a tooth for analysis could only be taken from a woman's skull. The woman was determined to be 16-18 years old[172 Table 2]. Determination of sex and age was carried by Prof. G.V. Lebedinskaya, who personally participated in the excavations. The burial belongs to the late Volosovo culture. However, it was genetically determined as a male (SI table I).

NEO178, burial 12. Adult male (late phase of Volosovo culture)

NEO192, burial 19. Adult male (Lyalovo culture).

NEO192 was dated to ca 6200 BP uncal. while the three others were c. one millennium later.

**Sakhtysh IIA**

The archaeological site Sakhtysh IIA was opened by D.A. Krajnov and E. Kostyleva in 1986. Excavations at the site continued intermittently for 11 years (1987 - 1994, 1999, 2004, 2015), six of which D.A. Krainov personally supervised [177].

As a result of excavations on an area of 824 m2, 72 burial sites were discovered: 57 classed as Volosovo and 15 classed as Lyalovo [172].

An analysis of the planigraphy of burials revealed the structure of both burial grounds and the sequence of burials. Representatives of Lyalovo culture buried their dead in the north-south direction, parallel to the river in a position stretched out on their backs, less often on their stomachs.  Representatives of Volosovo culture oriented the dead with their heads toward the river (to the south-west), in a position stretched out on their backs. A study of the stratigraphy of the Volosovo graves and the decorations made it possible to divide the burials into early and late. Early phase male burials were accompanied by amber ornamentation from the Eastern Baltic region (Lake Lubanos area) and lay at a greater depth (30-50 cm from the present surface). Late burials were located less deep (15-30 cm from the present surface) and were accompanied by decorations made from animal teeth, bone and stone. Serpentine stone decorations were found in female burials of this phase. This allows us to propose the eastern direction of communications - up to the Ural Mountains.

Anthropological studies of the materials were carried out by G.V. Lebedinskaya, T.I. Alekseeva, V.N. Fedosova (Philbert), M.V. Dobrovolskaya (Kozlovskaya). Sex-age determinations and craniological measurements were made, the chemical composition of bones was studied, and nutritional preferences were revealed. G.V. Lebedinskaya performed a number of graphic and sculptural reconstructions of the external appearance of the bearers of the Ljalovo and Volosovo cultures, which clearly demonstrated their differences [178,179].

It is worth paying attention to burial # 11. The date of burial makes it possible to reconsider its cultural affiliation. It clearly "drops out" from the dating of the Lyalovo burials from Sakhtysh IIa (burials # 40 and 42) and Sakhtysh II (burial # 19). The funeral ritual is also unusual: a young woman is buried in a strongly crouched position on her left side, with her head facing east (with a slight deviation). Its attribution to the Lyalovo culture was based only on the position of the skeleton below the level of the cultural layer in the natural soil ortsands [172]. The funeral ritual noted in burial 11 is characteristic of the cultures of the Bronze Age, including the Fatyanovo culture, which replaced Volosovo on the territory of this culture. Women of this culture were buried on their left side with their heads pointing east [180].  The dates of the Fatyanovo culture available in modern literature determine the time of its existence between  ~2880 (2750) and 2500 (2300) CalBC, pre-Fatyanovo refers to ~3000–2550 CalBC [181,182]. However the absence of any burial inventory in the grave gives the ability to accurately determine its cultural identity. It is obvious that this burial does not belong to the Lyalovo culture.

14 samples were analysed successfully, detailed in Table S7.2.

| Sample no | Context | Material | Labno | BP uncal | 1s | Sex | Age |
|---|---|---|---|---|---|---|---|

| | | | | | | | |
|---|---|---|---|---|---|---|---|
| NEO179 | Grave 13 skel 2 | tooth | UBA-39991 | 4919 | 30 | female | 50-60 |
| NEO180 | Grave 36 (lower) | tooth | UBA-39992 | 5314 | 34 | male | 20-25 |
| NEO181 | Grave 58 | tooth | UBA-39993 | 5328 | 39 | female | 40-45 |
| NEO182 | Grave 46 | tooth | UBA-39994 | 4767 | 35 | female | 20-25 |
| NEO183 | Grave 33 | tooth | UBA-39995 | 5011 | 35 | male | 50-55 |
| NEO185 | Grave 40 | tooth | UBA-39997 | 6393 | 39 | male | 50-60 |
| NEO186 | Grave 42 | tooth | UBA-39998 | 6317 | 91 | male | 20-25 |
| NEO187 | Grave 11 | tooth | UBA-39999 | 4616 | 38 | female | 20-25 |
| NEO188 | Grave 9 | tooth | UBA-40000 | 4916 | 35 | male | 50-55 |
| NEO189 | Grave 39 | tooth | UBA-40001 | 5157 | 35 | male | 30-35 |
| NEO193 | Grave32 | tooth | UBA-40004 | 4981 | 37 | male | 40-45 |
| NEO194 | Grave 34 | tooth | UBA-40005 | 5143 | 34 | male | 50-55 |
| NEO195 | Grave 35 | tooth | UBA-40006 | 5118 | 59 | male | 35-40 |
| NEO197 | Grave 36 (upper) | tooth | UBA-40007 | 4827 | 34 | male | 40-45 |

***Table S7.2. Details of samples from Sakhtysh analysed in this study.***

The dates of burials No. 40 and 42 of the Lyalovo culture to a certain extent agree with the dates obtained earlier in the laboratories of Aarhus and the Geological Institute of the Russian Academy of Sciences [175].

NEO179, burial 13 skeleton 2. This was originally recorded as belonging to Skakhtysh II, but burial # 13 from the Sakhtysh II site (excavation of 1978) was a solitary one and belonged to a teenager [172 Table 2]. Double (paired burial) # 13 was at the Sakhtysh IIa site (excavation of

1988). Skeleton 2 belonged to a 50-60-year-old man identified by G.V. Lebedinskaya, T.I. Alekseeva and M.V. Kozlovskaya (Dobrovolskaya) by stratigraphy and artefacts from the late Volosovo culture.

**Sakhtysh VIII**

The archaeological site Sakhtysh VIII was opened by D.A. Krainov in 1964.
Excavations at the Sakhtysh VIII site were carried out for 11 seasons (1965, 1970, 1971, 1973-1978, 1995, 2021). The excavation area is 1448 sq.m. 39 burials were identified. One burial belonged to the Lyalovo culture, 38 to the Volosovo culture. Most of them were destroyed.
The burial of the Lyalovo culture was oriented with its head to the north. The grave pit was located parallel to the river. The buried of the Volosovo culture, most often, lay stretched out on their backs with their heads toward the river (to the east). Many ornaments were found in the burials: pendants, beads, rings and buttons-plaques made of stone, bone and amber. Serpentines and slate were the main materials for making stone ornaments. They were found in burials of the late phase of the Volosovo culture. Burials of the early phase contained amber ornaments.
One sample is reported here: NEO184, from the burial of the Volosovo culture, a tooth of an adult female 40-45 years old. It was dated to 5014±36 BP uncal (UBA-39996).

Literature: Krajnov 1972[180]; Krajnov et al. 1994[177]; Alekseeva et al. 1997[178]; Kostyleva & Utkin 2010[172]; Piezonka et al. 2013[175]; Engovatova et al. 2015[179]; Kostyleva & Macane 2018[173]; Kostyleva et al. 2018[174]; Krenke 2019[181]; Macane et al. 2019[176]; Utkin & Kostyleva 2019[183]; Nordqvist & Heyd 2020[182].

#### Sosnovyi Mys, Angara valley, Russia. Settlement and cemetery.

*Nikolai A. Saveliev, Aleksey A. Timoshenko & Andrey Gromov*

The settlement and burial ground of Sosnovyi Mys are located in the Angara valley, Northern Angara region, on the lower end of the Sosnovyi Island, between the Kata and Yodarma River mouths.

The site was discovered by Nikolay I. Drozdov in 1974. Large-scale archaeological rescue work was performed in 2011-2012. The burial ground, where 8 burials were found, is located in the eastern part of the Sosnovyi Island, on the periphery of the settlement.

NEO841, Burial 4 was discovered on a base of the cultural horizon 3 which is attributed to the Neolithic period. The skeleton of a child of 5-7 years old was placed between large stones put in a rectangular construction. The burial was severely destroyed by a tree root system. Only hands, spinal column, and several ribs were found in an anatomical order while fragmented skull bones lay in separate compact clusters.

The grave inventory consists of two pendants made of split wild boar fangs with the holes at the ends. They lay around the neck vertebrae in the form of a circle with a diameter of 14 cm. Fragments of the upper and lower jaw of the child, and separately lying teeth are found inside of this circle. The skeleton was dated to 6781±35 uncal BP, UBA-40116.

NEO843, Burial 7 is a double burial. It was discovered in the cultural horizon 3. The burial pit was not traced, the grave constructions are absent. Skeleton 1 (sampled as NEO843): A male 20-30 years old lay on his back (along the East-West line), his head tilted to his left shoulder, facing South-West. The skull is crushed. This skeleton was dated to 6678±37 uncal BP, UBA-40117.

A bone tip with a bevelled base is found in the grave's upper part, at the level of the skull. Two musk deer fangs were placed at the east side of the skull. At the southeastern side of the skull 8 musk deer fangs and 2 pear-shaped pendants made from maral (Caspian red deer) teeth were found.

A comparative analysis of the burial tradition and inventory allows us to attribute the Sosnovyi Mys site to the Kitoy culture. At present time the site is the most northwestern point of the Kitoy people dispersion.

Literature: Okladnikov 1976[184]; Drozdov & Privalikhin 2003[185]; Saveliev et al. 2011[186]; Timoshchenko 2012[187].

#### Ural River, Orenburg region, Cis-Urals, Russia. River beach find.

*Mikhail Sablin*

A partly destroyed human skull was personally found on the Ural River beach in the Orenburg region by the Russian palaeontologist N.K. Vereshchagin in 1956. Dating of the human bone NEO100, SR-8457 around 9.1 kya 14C BP corresponds to the Romanov-Il'murzin culture, Mesolithic in the Cis-Urals chronology. The material has not been published before.

#### Ust'-Isha, Altai region, Russia. Cemetery.

*Marina Rykun & Vyacheslav Moiseyev*

The Ust'-Isha Neolithic burial ground is located near Ust'-Isha village (Krasnogorsk district, Altai region), close to Isha River mouth in the zone of low mountains of Altai. The site was excavated by B.Kh. Kadikov in 1961. 11 Neolithic burials were found below an Iron Age cultural layer dating 500-100 BC [135]. Several burials were damaged by Iron Age storage pits. Archaeologically the burial ground is attributed to Kuznetsk-Altai Neolithic culture.

The male burials inventory included stone and bone daggers, stone knives, arrows and darts heads, bone harpoon and others while female burials had few artefacts.

NEO067 - burial 7, morphologically female of 20-30 years old [134]. Several pendants made of elk and deer teeth were found in the neck zone of the skeleton. The sample was dated to 6640±55 uncal BP, UBA-39943.

Literature: Dremov 1980[134]; Kiryushin et al. 2000[135].

#### Vasilyevsky Cordon 17, Lipetsk, Russia. Settlement and burial ground

*R. V. Smolyaninov*

The archaeological complex Vasilyevsky Cordon 17 was investigated by R. V. Smolyaninov and A. A. Sviridov in 2012-2015 with a total area of 348 sq.m. It combines the features of a settlement and a burial ground of the Neolithic-Eneolithic eras. Located near the village Preobrazhenovka Dobrovsky district of the Lipetsk region in the floodplain of the river Voronezh (left tributary of the Don river).

As a result of the excavations, six foundation pits were investigated, a large number of bones, stones, and ceramic collection tools were obtained from several hundred vessels, mainly from the Srednestogovskaya culture of the Eneolithic era, dating from the first half of the 4th millennium BC.

The excavation revealed 28 burials. They are divided into three groups: wrinkled on the back, wrinkled on the side and elongated on the back. Based on the analysis of the studied burials, we believe that this soil repository was abandoned mainly by the population of the Srednestogovskaya culture of the Eneolithic era in the first quarter of the 4th millennium BC.

Nine human samples are reported here (see SI sample table).

Literature: Sviridonov 2013[188]; Smolyaninov 2013[189].

#### Vengerovo-2A, Novosibirsk Region, Russia

*V.I. Molodin, D. V. Pozdnyakov, L. N. Mylnikova, M. S. Nesterova, translated from Russian by Aija Macane*

The Vengerovo-2A burial ground is located on the edge of the second terrace above the floodplain on the left bank of the river Tartas (Vengerovsky District, Novosibirsk Region, Western Siberia).

Two collective burials of the late Neolithic period have been investigated. The mounds consisted of complex architectural structures, including a central burial pit and a ditch surrounding it, which in ancient times were covered with a barrow. The minimum number of individuals in the central grave consisted of 8 and 19 individuals, respectively. Various methods of the body disposal were recorded (inhumation, cremation, secondary and partial burials) and the dead were buried in different positions.

Analysis of the accompanying inventory demonstrates a pronounced north-west vector of parallels. In addition to those artefacts, with extremely wide territorial and chronological distribution (polished axes, adzes, teardrop-shaped pendants from bone and stone, tubular beads, artefacts made from incisors and canines of animals, shells), the rest of the items are, in one way or another, associated with the Neolithic materials from the taiga zone of the Western Siberia, the forested Trans-Urals, the Baltic states and Karelia.

Two burials from the complex No. 1 provided radiocarbon dates (skeleton 1: COAH-8738, 6250 ± 70 uncal BP and skeleton 2: COAH-8739, 6220 ± 80 uncal BP). According to them the usage of the site dates back to the end of the 7th millennium BC [190]. The new datings on these skeletons gave somewhat different results, see Supplementary Data IV. From complex 2, two individuals were dated: MAMS-29411 - 6322 ± 23 (Complex 2, sk. 17) and NSKA-02199 - 6368 ± 89 (Complex 2, sk. 18).

Cranially, Vengerovo people display the Northern Eurasian trait combination. This fact along, with skeletal and paleogenetic findings, places them within what can be described as the Uralian and Western Siberian Neolithic community.

First of all, it can be stated that all currently known craniological material from the Barabinsk forest-steppe belongs to the typological structure of the second level of racial differentiation - the north Eurasian anthropological formation. We can also talk about two components that formed the basis of the Vengerovo 2A paleopopulation. One of them is of the autochthonous origin and is close to people buried at the Sopka-2/1 and Protoka sites. The second component has its origins in the Volga-Ural interfluve. This component had the greatest impact on the male part of the population, while the female part of the population retained a certain local identity [191].

Eleven samples were analysed from this site, of which one was dated to the Mesolithic, eight to the Neolithic, one to the Chalcolithic and one to the Bronze Age. Details of the samples are found in the SI sample table. While most of these dates are comparable to the earlier datings, two individuals (NEO910 and NEO921) gave datings later than expected. We have no explanation for this discrepancy.

Literature: Chikisheva et al. 2015[191]; Molodin et al. 2016[190].

#### Zamostje 2, Sergiev-Posad district, Moscow region, Russia. Settlement.

*Olga V. Lozovskaya & Vyacheslav Moiseyev*

Zamostje 2 is a multilayer Late Mesolithic – Early and Middle Neolithic wetland settlement located at the Volga-Oka region, Sergiev-Posad district, Moscow region, Russia. The site is cut by the Dubna River. The archaeological layers are below modern water level.

According to archaeological data the groups of hunter-fisher-gatherers appeared on the shore of a vast post glacial lake along with the retreat of the water. Fluctuations in water levels continued for all 7th mil. cal BC [192]. The economy of Zamostje 2 people included hunting for elk and beaver, wetland and waterfowl, active and passive fishing (pike, perch, cyprinids), gathering berries, roots, and seeds [193,194].

No burials were found on the site. A number of human bones were probably brought to the site by its inhabitants while deciduous teeth might be lost by children naturally. Archaeological layers dates are: LM LL c.6600–6400 cal BC, LM UL (c. 6300–5900 cal BC), Final Mesolithic (c. 5900–5750 cal BC), Early Neolithic (c.5700–5300 cal BC) and Middle Neolithic (c. 4900–4300 cal BC). The ceramics were found in the Neolithic layers with no evidence of agriculture.

The results of the analysis of the metric characteristics of the permanent molars from Zamostje 2 suggest that the inhabitants of the site might have had common origins with the Onega culture population from Yuzhny Oleny Island [195]. More generally, the studied dental sample shows similarities to a wide range of Eastern European populations collectively referred to in paleogenetics as Eastern hunter-gatherers.

NEO087, sample 8, is a molar taken from the fragmented mandible of a young female of 18-20 years old. The sample was excavated in 1990 by Vladimir Lozovski in the dark grey sandy sapropel (gyttja) among hundreds of other artefacts made of bone, stone, and wood without burial context in the Late Mesolithic Upper layer (LM UL, square B3). The 14C dating of the sample is 7663±44 BP (KIA-51435) (6600–6440 cal BC) and can be slightly older than the surrounding layer.

NEO088, sample 9, is an incisor taken from the right half of the maxilla of an adult 20–30 years old. The sample together with few archaeological finds was found in 1998 by Vladimir Lozovski without burial context in a drainage trench in dark grey sapropel (gyttja) with shells between Upper and Lower Late Mesolithic layers (LM UL/LL). A 14C dating of the sample is 7581±41 BP (KIA-53296) (6500–6380 cal BC). A new dating gave a similar result, 7533±38 uncal BP, UBA-39957.

A detailed discussion of dating and results of stable isotope analysis of samples 8 and 9 is published by Meadows et al. [196]. The morphological description of dentition is in Zubova et al. [195].

Literature: Lozovski et al. (eds.) 2013[193]; Lozovski et al. 2014[192]; Lozovskaya (ed.) 2018[194]; Zubova et al. 2019[195]; Meadows et al. 2020[196].

#### Zjindo, Krasnochikoysky region, Russia. Cemetery.

*S. V. Vasilyev and S. B. Borutskaya*

The Neolithic burial ground of Zjindo is located on the territory of the Krasnochikoysky region of Western Transbaikalia, in the Chikoi river basin (the eastern part of the Selenga river basin and Lake Baikal). Now mountain taiga biotopes are common there. Excavations were carried out mainly in 2004-2005 by the Chikoy archaeological expedition of the Trans-Baikal State University under the direction of M.V. Konstantinov [161].

The Zjindo burial ground is located on the right bank of the river Chikoi on the lower part of the elevated slope, which is part of the system of spurs of the Malkhan Range. 2 km downstream of the river, the left bank of Chikoya already belongs to Mongolia. The elevation is about 680 m. The height above the river is 10-25 m. An artificial erosion ditch located across the slope surface has developed into a powerful ravine. In the ravine zone, from its top to the mouth, within 600 m, there is a burial ground consisting of a series of soil burials. It has been studied since 2005. 7 burials of varying degrees of conservation have been identified.

Two individuals were sampled:

NEO115, Zhindo # 7, burial N2, skeleton 1a. Dated to 6943±51 uncal BP, UBA-33756.

NEO117, Zhindo # 9, burial no.2, skeleton 2. Dated to 7964±63 uncal BP, UBA-33758.

Literature: Konstantinov et al. 2005[161]; Vasilyev et al. 2018[143].

#### Scotland

##### Banks Chambered Tomb, South Ronaldsay, Orkney, Scotland

*Nick Card, with contributions from Dan Lee*

Two phases of partial excavation were conducted by the University of the Highlands and Islands in 2010 and 2011 on a newly discovered chambered tomb. The area was disturbed during development work, but the heart of the monument was largely intact. The tomb consists of a linear central chamber, aligned E–W and c5m long, leading to five burial cells sealed by large capstones. The original ridge-like mound would have been up to 80m long, 20m wide and 2.5m high, but was truncated to the level of the capstones. The central area is partly subterranean and constructed in a quarried area into bedrock. The entrance passage is to the N. There are two larger cells at the W and E end of the central chamber, a single cell to the N and two cells to the S.

The excavation in phase one give a rare insight into the closing up of a tomb, with the slab layer and 'closing' deposits of human bone deliberately laid on top. This was then left open for a while (otter spraint was found on top of the slab layer) before final back filling. Otters were present right from the very first phase of burial in the west cell, right up to the closing layers, with spraint evident throughout, showing that the tomb was open (at least some of the time) and even shared with animals. There were numerous animal bones including sheep among the human remains (yet to be analysed fully) but no pottery was recovered from the ORCA excavations (although some were recovered by informal excavation in the passage subsequently). The human remains were disarticulated.

The N and E cells were partly excavated during Phase 1 and were found to contain human bones that formed closing deposits. These had been placed upon a layer of slabs, which is presumed to seal the lower unexcavated burial deposits. Six fragments of bone, predominantly cranium, were found in each cell, and a whole cranium had been placed in the E cell as if in a final gesture. The remains of the upper chamber backfill deposits were excavated during Phase 2. Further deposits of cranium and long bone were found within the SE cell above the slab layer.

In Phase 2 the damaged W cell was fully excavated. Above the floor were several layers of disarticulated human bone within soft semi-waterlogged silts. The evidence indicates that there were several phases of use within the cell, with distinct concentrations of bones. All the samples submitted came from the W cell. The other cells remain unexcavated and are preserved in situ.

Literature: Lee 2011a[197], 2011b[198].

#### Serbia

##### Vlasac, Danube Gorges, Serbia. Settlement with burials.

*Dušan Borić*

The site of Vlasac contains Early and Late Mesolithic and Early Neolithic deposits. Vlasac is situated in the Lady's Whirlpool Gorge of the Danube Gorges/Iron Gates area, on the right (Serbian) bank of the river 3 km downstream from the site of Lepenski Vir. The site was investigated in 1970–1971 by D. Srejović and Z. Letica examined an area of 640 m2 along the riverbank below 70 m a.s.l., which was subsequently submerged beneath the reservoir created by the Iron Gates I dam. More recent excavations were undertaken between 2006–9 by D. Borić, who examined a further 326 m2 upslope of the area excavated in 1970–1. The two excavation periods produced over a hundred burials of primarily or exclusively Mesolithic date, comprising primary inhumations and secondary inhumations and cremations [199–203]. A total of 68 AMS 14C dates on human and animal remains and charcoal from Vlasac range (after fresh water reservoir correction) between c. 9300–6000 cal BCE.

We report genetic data from three individuals, all of which were found in the period 1970–1971:

NEO655, burial 32 represents a fully articulated extended supine inhumation found in the vicinity of dwelling 2, immediately beneath burial 31 that was directly AMS dated by AA-57777 to 8196±69 BP (7756±82 BP after the correction for the reservoir effect was applied), with this date representing a terminus ante quem for burial 32.

NEO677, burial 41 represents a disarticulated skull found at the bottom of the stratigraphic sequence, on bedrock in quad. D/I.

NEO657, burial 83a represents a disarticulated human mandible found on the shoulder of an articulated supine burial marked as 83, which was directly AMS-dated by OxA-5826 to 7760 ±100 BP (7756±82 BP after the correction for the reservoir effect was applied).

Literature: Srejović & Letica 1978[199]; Borić et al. 2008[200]; 2009[201]; 2014[202]; Borić and Griffiths 2015[203].

#### Lepenski Vir, Danube Gorges, Serbia. Settlement with burials.

*Dušan Borić*

Lepenski Vir is one of the best-known archaeological sites in Europe, covering the Mesolithic and Early Neolithic periods. The site was discovered in the 1960s during archaeological surveys in advance of the construction of the Iron Gates I dam. Excavations between 1965–70 led by Dragoslav Srejović examined an area of c. 2500 m2 where an unprecedented array of archaeological features and artefacts relating to repeated use of the site over thousands of years was found [204–207]. These included the remains of around 70 buildings with trapezoidal bases and (often) furnished with lime plaster floors and stone-lined hearths, over 200 burial contexts [204], and exceptional numbers of stone and bone artworks and body ornaments. A revised chronological framework recognizes three main phases of site occupation of the site: Early–Middle Mesolithic, c. 9500–7300 cal BCE (‘Proto-Lepenski Vir’), Mesolithic-Neolithic Transition, c. 6150–5950 cal BCE (‘Lepenski Vir I–II’), Early Neolithic, 5950–5500 cal BCE (‘Lepenski Vir III’) (Borić 2019; Borić et al. 2018). Based on a total of 108 AMS 14C dates that cover Mesolithic and Early Neolithic contexts, no evidence of a Late Mesolithic (7300–6200 cal BCE) occupation has been identified at Lepenski Vir.

We report genetic data from two individuals:

NEO669, burial 126. Burial 126 is a disarticulated mandible, possibly of an Early/Middle Mesolithic date, found beneath a large trapezoidal base building structure of the Mesolithic/Neolithic transition phase I-II. Previously obtained genomic information on this individual has been reported in Mathieson et al. 2018.

NEO658, Burial 88: This sample was probably mislabeled as coming from Vlasac, but since no burial 88 exists at Vlasac, it more likely is from Lepenski Vir. If this is correct, it is the grave of an older adult, probably female. Burial 88 at Lepenski Vir was previously dated to 7130±90 uncal BP (OxA-5831, Boric 2019).

Literature: Srejovic 1972[207]; Radovanovic 1996[206]; Borić 2016[204]; Borić et al. 2018[208]; Mathieson et al. 2018[98]; Borić 2019[205].

#### Spain

##### Camino de las Yeseras, San Fernando de Henares, Madrid, Spain. Chalcolithic Tomb, Área 15, EI-08-I

*Corina Liesau, Patricia Ríos, Concepción Blasco, Jorge Vega & Roberto Menduiña*

Camino de Las Yeseras is a large, third millennium BC ditched enclosure covering more than 22 ha, strategically located on a terrace near the confluence of the Henares and Jarama rivers. More than 8500 structures were documented after the area was mechanically cleared during the 2006/2007 campaign. Circa 1200 features (pits, hut structures, tombs, structured depositions) were excavated by Argea Consultores, S.L. archaeological company in collaboration with the research team of Camino de las Yeseras from the Autonomous University of Madrid.

The analysed sample (individual 2) comes from a skeleton of an 18-20-year-old female from a tomb excavated in 2006 by Argea S.L. company. The tomb is a pit with dimensions of 1.60 m in diameter and 0.86 m in depth. In the bottom of the pit, 4 primary inhumations have been located, two infantile buried nearby, but not in physical contact, covered partially by two adult females in a crouched position, a 26-31-year-old female (individual 3) and at least by a younger female (individual 2) superposing the latter one, the body of individual 3. Several grave goods have been recovered in the centre of the pit, a granite mill, a plain bowl upside down, and a necklace bead. Intermingled with the human remains some cranial and appendicular bones of a lamb or kid have been recovered.

This tomb located in the south area of the site beneath the enclosure 4, is until yet one of the oldest ones documented in Camino de las Yeseras. The 14C dating of the individual number 2 is 4021 ± 30 BP, 2620-2471 cal BC 2σ (Ua-39308). The 14C dating of the female individual 4 gave the result 3990 ± 40 BP, 2621-2350 cal BC 2σ (Ua- 35016), evidencing contemporaneity, as well as the close contact of the skeletons, indicates a multiple burial in a filled space. The structure was half-filled with sediment and sealed by a compact clay layer. Remarkable, and until now as a unique case in this site, is the fact that several centuries later, another female (individual 1) was buried in a crouched position in an upper level of the tomb. The C14 dating of this adult female (individual 1), less than 27-year-old gave the result 3590 ± 30BP; 2028-1884 cal BC 2σ (Ua-35015). This led us to propose that probably this burial exhibit during a long period, some visual external markers to be located and respected.

The tomb is in close contact with enclosure 4, well documented by surface characterization up to 60m (of which 42m of them have been excavated). The enclosure is generally U-shaped, 1.35–3m in width, and with depths ranging between 0.45m and 1.40m. Another interesting aspect to highlight is that the tomb was so close to the enclosure 4, that when the circular trajectory of the enclosure was designed, it was deviated outside to avoid the destruction of the tomb. Once excavated the enclosure and still in use, some erosional processes took place, probably torrential rains, that affected the wall of the enclosure near the tomb. To avoid the destruction of the tomb the inhabitants of Camino de las Yeseras intentionally protected this zone around the tomb with an inner buttress of a compact earth step inside the enclosure.

Literature: Liesau et al. 2008[209]; Vega et al. 2009[210]; Blasco et al 2011[211]; Ríos 2011[212]; Gómez et al. 2011[213]; Ríos et al. 2014[214]; Liesau 2017[215]; Arteaga et al. 2017[216].

##### El Toral III, Andrín, Asturias, Spain. Rock shelter.

*Igor Gutiérrez-Zugasti*

The rockshelter of El Toral III was almost completely dug by María Noval and a team of the Institute of Prehistory (IIIPC – University of Cantabria) in 2009 as part of a rescue excavation. The site contained a huge Mesolithic shell midden that was heavily eroded by environmental and anthropogenic causes throughout the Holocene, although some intact stratigraphic units were still preserved at the time of excavation. Radiocarbon dates placed the Mesolithic occupation of the site between ~9500 and 7300 cal BP. Shell midden units were mainly composed of shells, lithics, and fish and mammal bone remains. Three post-holes were identified in unit 8. A partially disarticulated skeleton was found inside the mesolithic shell midden, in units 21 and 22. Apart from the Mesolithic evidence, the presence of pottery and human remains from disturbed units evidenced that the site was also visited by human populations between the Neolithic and the Bronze Age. Radiocarbon dates placed those visits in two different periods: 5600-5400 cal BP and 4300-3600 cal BP. Despite the heavy erosion that affected the site, an empty funerary cist was found. Anthropological and isotopic analysis of the human remains are still in progress as part of the PhD of Borja González-Rabanal.

Literature: Noval 2013[217]; Bello-Alonso et al. 2015[218]; Rigaud & Gutiérrez-Zugasti 2016[219].

#### El Mazo, Andrín, Asturias, Spain. Rock shelter.

*Igor Gutiérrez-Zugasti, David Cuenca Solana, Manuel González-Morales*

The inner test pit at the rockshelter of El Mazo was excavated in 2009-2012 by a team of the Institute of Prehistory (IIIPC – University of Cantabria). This is one of the most well-documented shell midden sites in northern Iberia. Although significant parts of the site have suffered from heavy erosion throughout the Holocene, part of the deposit was well preserved in the inner part of the rockshelter, close to the walls. Excavation of the inner test pit produced 27 mesolithic stratigraphic units dated between ~9000 and 7300 cal BP, providing an almost continuous record of occupations at the site. Microstratigraphic investigations have demonstrated the existence of several subunits inside some of the units identified during the excavation. The shell midden units were mainly composed of shells, with lesser presence of lithics and fish and mammal bone remains. Several in-situ hearths were also identified. Four human teeth were found across the stratigraphic sequence. The excavation of three additional test pits in other parts of the site have allowed to establish the formation and erosion processes at the site, as well as the presence of stratigraphic units belonging to the Bronze Age. Some isolated human remains (three teeth and a bone) were found in the outer test pit. Anthropological and isotopic analysis of the human remains are still in progress as part of the PhD of Borja González-Rabanal.

Literature: Gutiérrez Zugasti & González Morales 2013[220]; Gutiérrez-Zugasti et al. 2013[221]; 2014[222]; García-Escárzaga et al. 2015[223]; Rigaud and Gutiérrez-Zugasti 2016[219]; Gutiérrez-Zugasti et al. 2016[224]; Gutiérrez Zugasti et al. 2018[225]; García-Escárzaga et al. 2019[226].

#### Coves de Santa Maira, Castell de Castells, Alacant, Spain. Cave.

*Carles Lalueza-Fox, J. Emili Aura Tortosa, Domingo C. Salazar-García*

Coves de Santa Maira is a site situated c. 600 m.a.s.l., in a Mediterranean middle-mountain environment. Archaeological excavations were performed in two sectors: West (= SM-W) and East (= SM-CG). Archaeological materials used for this study were recovered from the site in 2009, during excavations carried out by the Universitat de València [227].

The dental piece analysed in this study comes from SM-CG, where short human occupations have been reported during the Late Glacial Maximum (23 – 19 kyr cal BP), the Tardiglacial (11.5 kyr BP) and the Neolithic (5,7 kyr BP). The tooth appeared in the upper levels of the

Pleistocene package, where several non-articulated human remains were recovered over a surface of c. 1 m2 under a pit. Two AMS radiocarbon dates were performed on this assemblage: on a human bone (Beta-261220: 8540 ± 50) and on a *Quercus* charcoal (Beta-313425: 8540 ± 50) [228]. The dental piece used for this study appeared only 30 cms away from the directly dated human bone.

The above-mentioned assemblage is intrusive, appears in Late Upper Palaeolithic levels but comes from a Holocene Mesolithic context. In SM-W there is evidence of Mesolithic layers with a similar chronology. From this other sector come human remains from 3 individuals with evidence of anthropic processing (cutmarks, bite marks, fractures and thermoalterations) [229]. Isotopic data show a wide variety of resource exploitation and suggest a connection with the coast, 30 km away [230].

Literature: Aura Tortosa 2014[227]; Salazar-García et al. 2014[230]; Verdasco Cebrián 2016[231]; Morales-Pérez et al. 2017[229]; Carrión Marco et al. 2018[228], Aura Tortosa et al., 2020a[299], Aura Tortosa et al., 2020b[300]

#### Sweden

##### Bredgården, RAÄ 113 Marbäck sn, Västergötland, Sweden. Wetland find. SHM 33241.

*Maria Vretemark*

The remains of a nearly complete human skeleton were found in 1994 when some trenches were dug in gyttja layers in a former lake. A total of 18 well preserved bones from the skeleton were found at a depth of 1.2 metres, dislocated over a relatively large area. The body is supposed to have been deposited or buried in a small lake. The bones had been disarticulated during decomposition in the bottom sediments. According to C14 dating, the human skeleton is Mesolithic, around 7800 BC. Observations on the bones reveal that this was an adult male with stature around 174 cm. He had lost some teeth ante mortem and the still present ones were heavily worn. There was also clear evidence of inflammatory processes in the jaws due to apical lesions. The age at death was estimated to be 45+ years. A tooth was sampled for DNA (NEO027).

Literature: Borrman et al. 1996[232]; Jonsson and Gerdin 1997[233].

#### Dösemarken, Scania, Sweden. Grave and settlement.

*Yvonne Magnusson*

The area (Limhamn 155:355) is located in the south-west of Malmö, directly east of the limestone quarry in Limhamn and was excavated in 2006 by Malmö Kulturmiljö.

This was a coastal area made up of arable land and the eastern part consisted of a former wetland, Hyllie mosse. The marshland has been fairly extensive but has since long been dried out and cultivated.

The area is part of what is known as Dösemarken, and there is information about the remains of a stone chamber grave nearby, possibly a dolmen (Hyllie sn., RAA no. 25:1), but its exact location is unknown. The existence of one or more stone chamber graves in the surroundings is indicated by the place names of the fields, Stora and Lilla Döse.

In Area A, a well-preserved skeleton grave was found (grave 1671, MK 321), along with a large number of settlement remains in the form of houses, pits, pit systems, hearths, a well, an oven, and postholes.

The tomb was rectangular, 1.9 x 1.2 m and 0.5 m deep, placed in a north-south direction and surrounded by a frame with large, evenly laid stones. The main end was marked especially with horizontally laid longer stones and with a circular shaped stone in between each long stone. In addition to the nearly intact skeleton, fragments of ceramics, flint flakes, and a bone awl was found at the foot end.

The body had been placed on the left side with its lower extremities bent, the upper arms along the body and forearms bent to the torso, with hands originally by the head. The position of the skeleton was in a north-south direction with its face directed to the east. The skeleton was very brittle and fell apart when touched.

This was the remains of a young woman who died at the age of 15 - 18. The size of the leg bones shows that the woman was relatively short, about 155 cm. Enamel hypoplasia occurs on several teeth, which have been formed at the age of 5–8 years and show that during this period the individual was subjected to physiological stress in the form of, for example, illness or malnutrition. The shape of the femur and tibia indicates that they have been subjected to some stress and can be seen as an indication that the buried woman has been relatively physically active during her short life.

Based on 14C analysis of a tooth, the grave can be dated to 2470 - 2230 BC. The design and location of the grave show similarities to graves from Battle Axe Culture and pieces of ceramic which were found confirm this hypothesis.

The orientation of some bones indicates that the grave was disturbed relatively shortly after the burial, about 2 or 3 years. The lack of grave goods, apart from a bone awl, could also be a sign that the grave has been looted.

The closeness of Neolithic house remains suggests that this can be interpreted as part of a burial ground belonging to one of the houses. A total of twelve house remains were found. Together with other remains of buildings, this indicates an intensive and varied settlement activity. The buildings have an assumed continuity from the Late Early Neolithic to the Late Neolithic/Early Bronze Age. One tooth was sampled from A1671, NEO044.

Literature: Magnell 2006[234]; Ifverson 2007[235].

#### Evensås, Skaftö 85, Bohuslän, Sweden. Flat graves.

*Karl-Göran Sjögren*

Human bones were found in 1930 during gravel digging in a late glacial shell bank. The find was investigated by Johan Alin later the same year. According to Alin the remains were from a single individual buried in supine position with the head towards the west. No artefacts accompanied the skeleton, but flint finds indicated a settlement at the site. Later osteological determinations by Gejvall and Ahlström showed the presence of two individuals. Individual A was a juvenile and individual B was a male, c. 25-30 years old. It is unclear if they had been buried in the same grave or not.

Several datings and isotopic analyses have been made, showing a very high intake of marine protein in both individuals. Datings range ca 5000-5200 BP, i.e. late Mesolithic, but after reservoir correction both must be regarded as early Neolithic in date, although no indication of Neolithic diet is seen in the isotope values. An upper right first molar from the adult male was sampled for DNA, NEO260.

Literature: Niklasson 1932[236]; Johansson 1974 nr 3[237]; Nordqvist 2000[238]; Liden et al. 2004[239]; Ahlström & Sjögren 2007[240]; Sjögren et al. 2009[241].

#### Fredriksberg, Falköping stad 5, Västergötland, Sweden. Gallery grave.

*Malou Blank*

The gallery grave at Fredriksberg in Falbygden, western Sweden, was excavated and restored by Västergötlands museum in 1973. The slightly trapezoid chamber measured

5.3×2 m and was oriented NNE-SSW. It was constructed of limestone slabs and consisted of a chamber and an ante-chamber covered by a collapsed roof. During the excavation a pottery sherd, a flint dagger (Lomborg type IIB), flint flakes, amber pendants, a slate whetter, a bone bead, bone needles, a bone awl, a bone cylinder, a few animal bones and commingled human remains were recovered. The number of buried individuals are estimated to be at least 28 to 30, both males and females, adults and children. Dating of the skeletons range from c. 2200 to 1650 BC, corresponding to the Late Neolithic and the beginning of the Early Bronze Age period I in the Scandinavian chronology. In addition to the aDNA data presented in this study, other scientific analyses were conducted, including Sr isotopes and stable isotopes. Most of the sampled individuals (12/21) exhibit childhood Sr isotope ratios which can be expected outside the local area of Falbygden. The stable isotopes indicate a terrestrial diet with a rather high intake of plant foods.

16 samples were analysed from this site, eight of which yielded usable DNA, presented in Table S7.3.

| **Sample id** | **Context** | **Material** | **Description** | **Sex** | **Ageclass** | **Age** |
|---|---|---|---|---|---|---|
| NEO220 | F115II:2 | tooth | PM1 dxt | M | juv/adult | 15-20 |
| NEO221 | F83III:1 | tooth | canine sin | M? | adult | >30 |
| NEO223 | F98III:2 | tooth | dpm1 dxt | | inf I | 3 |
| NEO224 | F121 | tooth | PM2 sin | F | adult | >35 |
| NEO225 | F123 | tooth | PM2 dxt | F? | adult | >40 |
| NEO226 | F90 | tooth | dc dxt | | inf I | 3-4 |
| NEO227 | F122 | tooth | canine sin | M? | mat | 40-50 |
| NEO228 | F108IV:3 | tooth | PM1 dxt | M? | mat | 45-55 |

***Table S7.3. Details of samples from Fredriksberg in the present study.***

Literature: Weiler 1977[242]; Blank et al. 2018[243], 2020[244].

#### Frälsegården, Gökhem 94, Västergötland, Sweden. Passage grave.

*Karl-Göran Sjögren*

The passage grave at Frälsegården in Falbygden, western Sweden, was excavated in 1999-2001 by Gothenburg University. In spite of damage and ploughing, this constitutes the most well-documented bone material from a Scandinavian megalithic tomb. The grave had been a ca 10 m long and 2 m wide chamber of limestone slabs, with a passage towards the east, also ca 10m long. The presence of a number of whole or partially articulated skeletons was one of the most significant results of the excavation. These range from almost complete skeletons to partial articulations. In addition, there is a mass of disarticulated bones but also some bones that seem to have been treated differently, such as a skull group and a couple of bone packages. The number of buried individuals are estimated to be at least 51, but more likely ca 78-80. Datings of the skeletons range mainly ca 3100-2900 BC, corresponding to the late Funnel Beaker Culture, period MN A in the Scandinavian chronology. A large number of scientific analyses have been made, including aDNA, dietary isotopes, Sr isotopes. Two individuals tested positive for plague infection, so far the oldest identified cases. Sr isotopes suggest that some 25% of the individuals were born outside the local area.

One sample was analysed in this project, NEO259 (ID120033, bone ID3644, Tooth ID 318), a lower left PM4 tooth.

Literature: Ahlström 2009[245], Skoglund et al. 2014[246]; Sjögren 2008[247], 2015[248], 2017[249]; Sjögren et al. 2009[241]; Rascovan et al. 2019[250].

#### Hanaskede, N Ving sn, Västergötland, Sweden. Wetland find. VGM 106100.

*Maria Vretemark*

In autumn 1990 a single human cranium was found during drainage of a small kettle hole near lake Hornborgasjön in Västergötland. No other parts of the skeleton were found despite thorough search around the find spot, nor any artefacts. The cranium was C14 dated to the Mesolithic period, around 8000 BC. At that time the lake Hornborgasjön was larger than today and there were lots of Mesolithic settlements on the shores. When it decomposed the separate skeletal parts were scattered around and eventually embedded in the bottom sediments of the kettle hole. The cranium was from a male with an age at death estimated to 40+ years. Some of his teeth had been lost in life due to severe dental wear. An apical infection was noted in the upper right lateral incisor. He also had a scar from a healed injury in his frontal bone, just above the right eye. Dietary isotopes from two teeth and a cranial sample were published by Eriksson [96].

One tooth sample was analysed, NEO018.

Literature: Vretemark 1996[251]; Eriksson 2003[96]

##### Hindby mosse, MHM 1505, Scania, Sweden. Central place.

*Yvonne Magnusson*

The archaeological excavations of Hindby Mosse were carried out by Malmö Museum between 1967 and 1974. Today, the site is beneath a motorway, Inre Ringvägen, which transverses the site and it is located about 600 m southwest of Hindby junction in the southern part of Malmö.

In prehistoric time the site was located on a fairly flat isthmus between two shallow bays in a lake which today is Hindby mosse. Just south of the settlement, a stream debouched into the lake. In the vicinity there are several neolithic settlements and a long dolmen.

Hindby mosse is an unusually large settlement and is an example of a central place dating to the Middle Neolithic period and the Funnel Beaker culture. In the area several remains of buildings have been found. The almost circular cultural layer covers 1.5 hectares and in the middle is an 80 x 60 m large featureless area surrounded by two semi-circular activity areas, one in the north and one in the south. The total site has not been investigated, but the area was divided into four larger trenches and several smaller trenches and test pits.

The artefacts consisted mainly of flint tools such as burnt and unburnt axes, arrowheads, scrapers and flakes, but ceramics are also well represented on the site. Most vessel fragments are from storage vessels, but also beakers and bowls of finer ware and richly decorated are represented.

A large number of animal and human bones were also found. Bones and horns from slaughtered cattle and from hunting were used as raw material for making tools.

Crucial for interpreting the place as a central place for the local population with elements of ritual activities is its circular structure, how it is situated on an isthmus, the number of fragments from human skulls and the presence of burnt axes.

Seven samples were taken, three of which were successful:

Cranium 1, NEO036. This skull was found centrally in context 4 and with the "face" down. The back of the head was crushed and was otherwise relatively compressed and partially burnt. About 10 cm west of the skull was a skull fragment. It is unclear if it is related to the skull. Next to the skull's northern side was found a scraper. Near the skull was also a tusk from a wild boar. No traces of the rest of the skeleton were discovered. No marked burial or other marking in the skull area was observed. In osteological analysis, it was assessed that the skull belonged to an individual aged 20 - 25 years.

Cranium from square 15/71, NEO038. In osteological analysis, it was judged that the skull belonged to a young individual. Other documentation is missing.

Cranium 2, NEO039. The second skull was found on the outskirts of context 4. The skull was compressed so that the skull top was depressed in the lower part. Adjacent to the skull was a piece of a jaw with teeth. Immediately adjacent to this jaw was a mandible from a larger animal (pig or dog) and nearby a femur, possibly human, was also found. As in the case with the first skull, there is no marked burial or other marking. In osteological analysis, it was estimated that the skull belonged to a young individual, about 7 - 9 years old.

Literature: Archival material, unpublished, S 03:12-14: Malmö Museer; Salomonsson 1971[252]; Jeppsson 1976[253]; Svensson 1986[254]; Sandén et al. 2010[255]; Nilsson 2020[256].

#### Kastanjegården, Scania, Sweden. Cemetery.

*Yvonne Magnusson*

During 1972 - 1974, Malmö Museums conducted excavations at Kastanjegården for future development of the area. The excavation area is located in the southern part of Malmö at Kastanjegården and south of Fosie church. Here, settlement remains from the younger Stone Age, older Iron Age and the transition between the Viking Age and the Middle Ages emerged. A flat earth grave field dated to Middle Neolithic B (Battle Axe Culture) was found. The burial ground was oriented in a north-south direction and covered an area of about 17 x 10 m.

Grave 105 was located in a north-south direction and the north-west part of the grave was partially destroyed by sand extraction. The grave consisted of a stone-framed structure measuring 2.5 x 1.5 m with the remains of a decomposed wooden coffin. In the lower part of the stone structure was found a stone with two cup marks. In the upper part of the structure lay a grindstone.

Skeletal remains that were severely weathered and partially crushed were found in the grave. Only the skull and parts of the hands were preserved, the rest was decomposed. The skeleton turned out to be a woman aged 25 - 30, lying on her right side with her head to the south and facing east. Two children were buried in the same grave, both about 5 - 7 years old.

The child in the southern part was placed next to the woman, either in front of or behind her and with its head in the south. The child in the northern part of the grave had probably been lying at the woman's feet with its head to the north facing east.

The grave goods consisted of two earthenware vessels, seven amber beads, a flint axe, four flint chips, two scrapers and five flint flakes. The two vessels were of typical Battle Axe Culture type, belonging to pottery group J.

Sometime during her lifetime, the woman had been hit severely to the left side of her head. The blows had been so violent that a tooth in the upper jaw and one in the lower jaw had been badly damaged and inflamed. Whether this is the result of abuse or an accident could not be decided, but the woman during the later part of her life suffered persistent pain in the jaw and severe toothache. The teeth in the other half of the jaw were also more worn than the injured part, suggesting that she was in pain from chewing the food.

A tooth from the adult woman in grave 105 was sampled, NEO051, MHM 4555.

Literature: Winge 1974 [257]; 1976[258]; Rosborn 1999[259].

#### Sillvik, Torslanda 43, Bohuslän, Sweden. Flat grave.

*Karl-Göran Sjögren*

Remains of three individuals were found in 1929 during gravel digging in a late glacial shell bank. Further bones were reported, but have not been recovered. Finds of two flint daggers close by indicate a late Neolithic date. Individual A is a male >40 years old, individual B is a probable male, 20-30 years old, and individual C is a juvenile. Individual B was 14C-dated to the Late Neolithic. Despite the coastal location, his δ13C value indicates only marginal intake of marine protein.

The sample, NEO261, is a lower left PM from individual B.

Literature: Niklasson 1956[260]; Sjögren et al. 2009[241].

#### Skateholm I, Scania, Sweden. Cemetery.

*Lars Larsson*

The sample was taken from one of 65 burials excavated from a combined settlement site and cemetery at Skateholm I in the southernmost part of Sweden. Skateholm I is one of two cemeteries situated on islands in a former lagoon dated to 5200–4800 BC. The material culture belongs to the Ertebølle Culture.  Both sites were submerged due to transgression during the late Mesolithic.

The sample, NEO679, is a tooth from a mature female (ca 55 years old) buried in feature 52, grave 26.

Literature: Larsson 2004[261]; 2016[262].

#### Vattenledningen, Scania, Sweden. Cemetery and settlement.

*Yvonne Magnusson*

Due to construction of a water supply line in 2005 between Hyllie water tower in Malmö and Vellinge, Malmö Kulturmiljö conducted an archaeological excavation.

The area which was surveyed consisted of long narrow excavation trenches in north-south direction through the northern part of Vellinge municipality. The area is located just east of Hököpinge and just over a km west of Hököpinge church village, and north of Pilebäcken which flows in an easterly direction towards the Öresund. The area is on a ridge, which slopes southwards towards Pilebäcken.

The excavation area turned out to be an area of relatively intense prehistoric activity. The remains that emerged consisted of settlements, graves and fire-related remains which dated from the end of the Early Neolithic to the Roman Iron Age, a span of about 3000 years.

In the southern part of Area 3, a burial ground with six flat earth graves, hearths, pits and scattered post holes appeared. Five of the six flat graves were dated to the transition between the Late Neolithic Age and the Bronze Age and one to the later part of the Early Bronze Age.

The grave A 430 has been interpreted as a skeleton grave, possibly a double grave where the bodies are buried in a row. It appeared as an approximately 3.50 m long coloured staining in east-west direction with scattered skeletal parts in the surface. Skeletal parts, which were highly fragmented, from two people were found in two concentrations, one in the western and one in the eastern part of the grave.

In the western part of the grave there was a crushed skull and different bone remnants. The body had been placed with its head to the west facing north. In the eastern part of the grave, there were a few additional cranial parts and bone remains. Traces of soot were found around the skull. According to the osteological analysis, the bones were from two adult males, who died at the age of 20 and 25-30 years. One skeletal sample was 14Cdated with the result 1943–1748 BC, corresponding to the Late Neolithic in Scandinavian chronology. In the grave filling, scattered artefacts were found in the form of two flint flakes and a fossil sea urchin.

One sample, NEO052, from A430 was analysed. The sample was a tooth from the individual found in the grave N° 430.

Literature: Arcini 2007[263]; Gidlöf & Gruber 2009[264].

#### Ängdala, Scania, Sweden. Grave and flint mines.

*Yvonne Magnusson*

The area around Ängdala farm at Kvarnby and Södra Sallerup on the eastern outskirts of Malmö contains the only known Early Neolithic flint mines discovered in Sweden. They were discovered in the early 1900s when digging in the area for mining chalk, and archaeological excavations were carried out at several occasions in the area, mainly during the 1970s until the 1990s.

The mines are different in size, but usually 2 - 7 metres deep. There are also many open quarries in the area, which are characterised by pits with chalk on the surface. They are a maximum of two metres deep. In some places it has been possible to pick flint nodules in the upper soil layer without digging.

The mining activity is dated to the Early Neolithic (ca 4000 - 3300 BC). Samples show that the mines were re-used during the later part of the Neolithic era, during the Bronze Age and even during the Iron Age. C14 dating shows that the flint quarrying was most intense in the area between 4000 BC and 3650 BC.

When excavating a flint mine (A2408 C) in 1991, a skeletal burial (Grave A2408, MHM 6902) was found in the soil of the depression formed by the mine. On a stone layer and with larger stones under the head, a young woman about 18 - 20 years old had been buried. Only a few skeletal parts were found, but most probably she was buried stretched out on her back. Above the dead had been laid soil and in the next layer there was a sooty layer of stone with some bird bones. With some reservation, this layer can be interpreted as an applied layer in connection with ceremonies at the burial. A couple of flints have been interpreted as grave goods. The skeleton was C14 dated to the Early Neolithic I.

A tooth sample from this individual was analysed, NEO046.

Literature: Rudebeck 1986[265]; Frejd & Rudebeck 2013[266]; Berggren 2018[267]; Malmö Museum archive.

#### Turkmenistan

##### Monjukly Depe. Settlement.

*Elizaveta V. Veselovskaya and Vasilyev Sergey*

An ancient Neolithic settlement on the left bank of the Tejen River (South Turkmenistan) was discovered in 1960 by archaeologist A.A. Marushchenko. Excavations were also carried out 1961 by Berdiev and 2010-2014 by Bernbeck & Pollock. Datings range ca 6000-5650 cal BC and 4800-4350 cal BC millennium BC which corresponds to the Jeyton and Anau I A cultures, i.e. Neolithic and Eneolithic [268]. The Eneolithic settlement arose on the ruins of the village of the Jeytun culture [269]. Ceramics of the late stage of the Jeytun culture, silicon drills and scrapers were found, although microplates predominate, and there are also many copper products.

The location is important for establishing regional chronology because here the Chalcolithic layers follow the Neolithic layers. However, in 2010 subsequent excavations revealed a long break in settlement between the end of the Neolithic settlement (layers XV, 6200-5600 BC) and the Chalcolithic migration (layers IV-I, 4650-4340 BC). Based on this, the "Mean horizon" was defined here, which, apparently, is limited to the Kaka region in Turkmenistan and precedes the phase IA of the Anau culture. Layers IV-I were excavated over a large area. They contain standard houses with a square layout and columns in the middle of the rooms. In the upper two layers, a closed open space was discovered, in which, judging by the animal bones found here, banquets took place. In the lowest layer IV, a house was found with a wall painting depicting two people, also with some abstract patterns. The inhabitants of the Monzhukli depots lived by cattle breeding and agriculture. Among herd animals, sheep and goats predominated. Cattle, as well as their skulls, played an important role at feasts. As for wild animals, the remains of gazelles and onagers were found. Barley and wheat played an important role in arable farming, and analyses potentially indicated simple irrigation. Very little pottery was produced in the Chalcolithic Monzhukli Depe. On a general level, there are ceramic parallels to the Sialk II/Ceshme Ali horizon of the Iranian highlands.

A burial ground was found with seven buried, whose bones are partially covered with ocher. Two of them are head-oriented to the north-west, the rest to the north and northeast. The anthropological type of the buried is characterised as the eastern Central Asian Mediterranean [270]. In the studied population, two anthropological types are distinguished: one Proto-European with a low and wide face, and the other having southern signs (small protrusion of the nasal bones, pronounced prognathism), which brings it closer to the South Indian anthropological type.

Literature: Korobkova 1969[269]; Ginzburg & Trofimova 1972[270]; Bernbeck & Pollock 2016[268].

#### Ukraine

##### Igren 8. Cemetery.

*Alexandra Buzhilova & Natalia Berezina*

Multi-layered site Igren 8 was located on the banks of the Samara River, a tributary of the Dnepr River. It was discovered in 1945 by an expedition of the Institute of Archeology of the Academy of Sciences of Ukraine [271].

Archaeologists date the site to the Neolithic – Eneolithic. The early Eneolithic age in the area is associated with local Sredniy Stog II archaeological culture [272].

Preserved bone remains reflect both the Neolithic period and later time. The bone remains are stored in the Museum of Anthropology, MSU.

According to anthropologists, the skulls are dolichocranial with a large cerebral part and with average height and width of the face; the nose is protruding [273]. According to Potekhina, two anthropological types are represented in the Eneolithic of Ukraine: 1) massive hypermorphic variant and 2) gracile mesomorphic one. Both types are presented in the craniological series of Igren 8. The researcher notes that the gracile craniological complex in the population of this culture may be the result of mixing with representatives of the earlier Tripolie culture [274].

Literature: Dobrovolsky 1949[271]; Konduktorova 1973[273]; Telegin 1973[272]; Potekhina 1999[274].

#### Kleshnya III, the Seversky Donets River, Ukraine. Burial in a settlement.

*Inna Potekhina*

The site Kleshnya III (Claw III) was excavated by V. Man'ko and S. Telizhenko in 1998 in the Kreminsky district of the Lugansk region, Ukraine. The site is located on the cape of the floodplain terrace of the lake Kleshnya 2, on the right bank of the Seversky Donets River. Excavations revealed the settlement and burial of the Early Neolithic era, as well as the cultural horizons of the Late Eneolithic and Late Bronze Age.

For DNA analysis, bone samples were taken from the Early Neolithic and Late Eneolithic burials.

Burial No. 1 (Early Neolithic) dates to the last third of the 7th millennium BC, contained the skeleton of a woman (25-35 y.o.) buried in a stretched position, sprinkled with ocher. Above her was a child’s skull. Unfortunately, their samples did not yield results in genetic analysis.

In contrast, the burial of the Late Eneolithic era proved to be suitable for DNA analysis. Burial No. 2, NEO278 belonged to an elderly man (maturus-senilis) who suffered from a disease of the knee joint. The burial has been attributed to the Repin culture; for it, radiocarbon dates were obtained in the range from 4170 ± 60 BP to 4050 ± 60 BP (Ki-7848, Ki-7849). However, a direct date on the sample resulted in 2860±25 (UBA-40022), i.e. Bronze Age.

Literature:  Manko & Telizhenko 2000[275]; 2002[276];

#### Lysa Gora, Lower Dnepr, Ukraine. Burial ground.

*Inna Potekhina*

Lysa Gora (Bald Mountain) burial ground is one of the most ancient in the group of Mariupol-type Mesolithic-Neolithic cemeteries in the Dnieper basin. The cemetery was excavated in 1959 by A. V. Bodyansky. It is situated 5 km to the west from the village of Vasylivka, Zaporizhzhya region, in an outcrop of a loessic terrace at the Kakhovka reservoir. The main part of the cemetery was covered with red ocher, this area consisted of a lens up to 60-70cm thick. There were five burial pits of round or elongated oval form, the total number of skeletons recovered exceeded 50. These included six-seven skeletons without burial pits (probably the earliest in the cemetery), eight child skeletons in pits I-III, 26 partially burnt skeletons in pits IV and V which probably comprised a single ossuary, and about six additional cremation burials. In the elongated portion were skeletons in the extended position while in the annular pit there were more than twenty human skulls. The skeletons in the grave pits were mainly in a ruined state.

The grave goods from Lysa Gora were both varied and numerous. Within the area of red ochre and the grave pits there were found sherds from 80 vessels broken during the burial rites. The pots were flat bottomed with biconical shapes and collared rims. Ornamentation usually covered the entire surface of the vessel, including its base. Stroked ornament predominated with incised and linear ornament less frequent. More than twenty flint artefacts, including large knives and knifelike blades and scrapers were found. There was a variety of ornaments among which were seven plates of boar tusk enamel of the Mariupol-type, fish teeth, deer tooth pendants and numerous annular beads of various sizes, ranging from 0.6 to 3.5cm. in diameter. They were fashioned from shale, gagate, bone and shell.

The cemetery is associated with the Neolithic habitation sites near the Dnieper rapids. The presence of a large amount of pottery in this cemetery convinced that the Mariupol-type cemeteries in the Dnieper valley and in the adjacent steppe territories were related to the Nadporizhzhya-Azov group of the Dnieper-Donets ethno-cultural community.

Two burials from Lysa Gora cemetery were sampled for DNA analysis.
Burial #3, NEO262 belongs to a 45-50 years old male. A sample of the left *Pars petrosa* was taken for DNA analysis. It was dated to 6175±35 uncal BP (UBA-40018).
Burial kv.8, NEO265 belongs to a 35-50 years old male (?). The 1st lower molar, left side, was sampled for DNA analysis. It was dated to 5276±38 uncal BP (UBA-40019).

Literature: Telegin & Potekhina 1987, pp.11-15, 110, 111[277].

#### Mamai Gora, Lower Dnieper, Ukraine. Burial ground.

*Inna Potekhina*

The Neolithic cemetery Mamai Gora is on Mamai Hill near the city of Kamyanka-Dniprovska in the Zaporizhzhya region, Ukraine. It was investigated by G.N. Toshchev in 1989-2001. 26 graves were unearthed in this burial ground. Those buried lay on their backs in a stretched position, with their heads to the east and southeast. Ocher was recorded in 13 burials, but the custom of using ocher, apparently, has not yet fully settled. The ocher was poured on the bottom before laying the person buried in the pit, it was used when filling the pit, and it covered individual bones. Most of the inventory is represented by ornaments which were obviously sewn on hats and clothes in the chest, shoulders, belt and feet. These are pendants from underdeveloped teeth of a deer, beads from mother-of-pearl shells, bone and stone. Fragments of wide flint plates and high trapezes with retouching were also found in the graves.

The Neolithic burial ground Mamai Gora is a typical representative of a large group of Mariupol-type cemeteries of the Dnieper-Donets cultural-historical region in the Northern Black Sea region. The radiocarbon dates obtained from the bones of the three burials indicate that the burial site existed in the period of about 5200-5000 (5950-5750) BC.

For genomic analysis, samples of teeth and *Pars Petrosa* bones from two burials (No. 6 and No. 9) were taken.

Burial # 6, NEO268 belonged to a 30-35 years old male. The skeleton was at a depth of 1.2m. The bones of the legs were separated by a crack and lay 0.3m below the level of the upper half of the skeleton. The man was buried in a stretched position on his back, his head to the ESE. The left arm is laid along the body, only the humerus is preserved from the right arm. At the humerus of the right hand and to the right of the skull, ring-shaped flat beads from shells (35 units) and 2 deer pendants were found. A radiocarbon date of 7055±70 BP (Ki-9193) was obtained for this skeleton.

Burial # 9, NEO270 belonged to a 45-50 years old male. The skeleton lay in an extended position on his back, his head to the SEE. The bones of the arms are stretched along the body, the face is turned to the left. On the frontal part of the skull were 37 pendants made of the deer teeth and 31 ring-shaped beads from shells, five of which with a corrugated surface. There were 14 pendants above the elbow joint of the right hand, 8 pendants above the wrist of this hand. Above the elbow and at the wrist of the left hand, 3 pendants were revealed. In the area of the "abdomen" two "belts of deer teeth" were traced. The first “belt” consisted of 14, and the second – of 43 pendants. Under the skeleton ocher is fixed. The burial was dated twice, to 6960±70 BP (Ki-8184) and to 6823±38 uncal BP (UBA-40021).

Literature: Tuboltsev 1992[278]; Toshchev 2005 [279]; Andrukh & Toshchev 2009, P.13-34[280].

#### Vasilevka I, Dnepr river valley, Ukraine. Cemetery.

*Alexandra Buzhilova & Natalia Berezina*

Burial ground Vasilevka I was discovered in 1953 by A.V. Bodyansky on the left bank of the Dnepr River near the village of Vasilevka in the area of the modern town Dnepr, Ukraine. The excavation of the Vasilevka was done later by A.D. Stolyar (1957). On the area of about 46 square meters, 26 burials were opened. The bone remains are stored in the Museum of Anthropology, MSU.

Burials, as a rule, were single; in three cases, paired burials were recorded (Nos. 2, 14, 18). The body orientation of the head was mainly to the east. The crouched body position on the right side was most often. Some skeletons were covered partly by ochre (Nos. 2, 4, 5, 6, 7, 9, 10, 14, 14a, 17, 18a).

The river shells were found in three burials (Nos. 2, 9, 18a), stones with traces of heat treatment lay in two graves (Nos. 14, 19), and microlithic flint artifacts were found in three other graves (Nos. 10, 17, 18). Based on these findings, as well as parts of other flint tools from the cultural layer, Stolyar dated the site to the Mesolithic.

A study of the skulls showed that in the series there may be at least two anthropological components, which are reflected in two craniological complexes. The first variant demonstrates high and narrow-faced orthognathic skulls with high orbits; the second one presents the broad-faced skulls with low orbits. The first craniological complex was found also in the anthropological series of the Voloshsky burial ground − a synchronous site, which was located just 16 km from Vasilyevka 1. According to Konduktorova [273,281], the presence of several craniological variants in the Mesolithic clearly shows that the population of this territory had a different geographical origin, and the most common craniological complex could reflect the Late Palaeolithic anthropological background.

Literature: Konduktorova 1957[281]; Stolyar 1957[282]; Konduktorova 1973[273]; Grünberg 2000[283].

#### Vovnigi I and Vovnigi II, Dnepr river valley, Ukraine. Cemeteries.

*Alexandra Buzhilova & Natalia Berezina*

The burial ground of Vovnigi I (Left-bank Vovnigi) was discovered by M. Ya. Rudinsky in 1949. The site was excavated on the left bank of the Dnepr River in front of the village of Vovnigi, Solonyansky district. Numerous burials have been discovered in the mainland loess.

They were densely stacked in a row, one burial was next to another, and often one was above the other. In total, 31 burials were opened. The bone remains are stored in the Museum of Anthropology, MSU.

The orientation of the buried was along the north-south line, and body orientation of the head was to the southeast. The burials, located in the central part, were richly covered by ochre; and no ochre was found in peripheral burials. Studying the positions of the bodies, the researcher suggested that the bodies were strongly pulled together in several places or swaddled [284]. The burial ground was used over a long time. Graves formed at close range and at about the same depth gradually formed one large pit. According to Rudinsky, such a cluster can be considered as a kind of ossuary. Samples N° 9481, 9482, 9483, 9484, 9487.

The burial ground of Vovnigi II (Right-bank Vovnigi) was located on the right bank of the Dnepr River in the centre of the village of Vovnigi, Solonyansky district. Rudyansky excavated about 100 square metres and opened 130 burials in 1952. Part of the bone remains is stored in the Museum of Anthropology, MSU, Moscow (63 burials) and another part is in the Museum of Anthropology and Ethnography, RAS (Kunstkamera), St. Petersburg (37 burials).

The burials located in the central part were covered by ochre; and, the same as in the site of Vovnigi I, no ochre was found in the peripheral burials [284]. The researcher identified three groups of burials: 1) in the northwestern sector, the burials lay along the north-south line, and partially some burials overlapped the others; 2) in the northeastern sector, the burials did not overlap and were located more freely one from another; 3) in the central part of the site there is a massive multi-layer accumulation of burials. The orientation of the bodies of those buried was the same as in the site of Vovnigi I. Samples N° 9840, 9844, 9874, 9876, 9879, 9892, 9864.

Rudyansky [284] notes that there are no significant archaeological finds to give a fairly narrow chronological interval for both monuments. In his opinion, both sites were formed during the Neolithic. Based on some features of burials, the author stressed that these may be late Neolithic burials.

A partial sample of Vovnigi II, housed in St Petersburg, has been dated to 5,470–4,783 cal BC (OxA-5938, OxA-5939, and OxA-5940) [285].

Anthropologists did not reveal any differences between the anthropological sequences of the two burial grounds. In many ways, the skulls are very close to the Late Palaeolithic skulls of Europe [286,287]. Previous aDNA analyses were published by Jones et al. [97].

Literature: Rudinsky 1955[284]; Konduktorova 1956[286]; Gokhman 1966[287]; Telegin et al. 2003[288]; Lillie & Budd 2011[285]; Jones et al. 2017[97].

#### Voloshskoe, Dnepr river valley, Ukraine. Cemetery.

*Alexandra Buzhilova & Natalia Berezina*

The burial ground of Voloshskoe was discovered in 1952 during archaeological explorations by A.V. Bodyansky on the bank of the Dnepr River near the village of Voloshskoe in the area of the modern town Dnepr, Ukraine. During the work Bodyansky excavated nine graves and later V.N. Danilenko finished the excavation of the site [289]. In total there were 18 burials (19 individuals). The skeletal remains are stored in the Museum of Anthropology, MSU.

Most burials were single, with grave No. 8 a paired burial (man and woman). The most common burial rite was a variant of the crouched body position (11 individuals), with the head oriented to the southeast. Moreover, 10 remains were buried lying on the right side (burials No. 1, 2 and 4 - 10), and one person laid on the left side (No. 3). The second group includes graves with remains lying on the back and oriented with their heads in the southeast.

According to Danilenko [289], the burial ground was located on the second loess terrace of the Dnepr River, that is, at the same level as the final Palaeolithic sites of the region. Archaeological analysis allowed the researcher to confirm that most of the burials belong to the Epipalaeolithic period. Both archaeologists and anthropologists note the heterogeneity of the group, and one of the reasons could be that the burial place was used for a long time [273,289–291]. The sex and age structure of the group indicates a clear male prevalence. In addition to a small number of women, only one immature individual was noted in the sample. The group does not differ from other Late Palaeolithic and Mesolithic samples of its structure. This period is characterised by the burial of mostly men, and very rarely women or immature individuals. The average age at death refers to the maximum values of this indicator, according to palaeodemography of the Mesolithic groups [292]).

Palaeopathological analysis divides this group between two patterns of trauma. The first includes two cases with healed skull injuries (Nos. 6 and 17) and persons without any traumas. The second sample includes individuals buried with evidence of postmortem body manipulations and/or perimortem traumas of the skull that could have caused death (Nos. 3, 13, 14, 15 and 16). This group demonstrates episodes of aggressive intrusions and skirmishes in the area [293]. Sample 9881

Literature: Danilenko 1955[289]; Debetz 1955[290]; Telegin 1957[291]; Alexeev 1972[292]; Konduktorova 1973[273]; Buzhilova & Berezina 2016[293].

#### Volnensky, Dnepr river valley, Ukraine. Cemetery.

Veselovskaya Elizaveta V. & Vasilyev Sergey

The burial ground is located on the left bank of the Dnieper River above the currently flooded Volny rapids, near the city Zaporozhye. Excavations were carried out in 1956 by an expedition of the Institute of Archeology of the Ukrainian SSR [291]. The site belongs to the Dnieper-Donetsk culture, which existed on the territory of the Dnieper Nadporozhye in the Neolithic era from the fifth to the beginning of the third millennium BC. The burial ground is a tribal cemetery, in which burials were carried out over many generations. Stratigraphy and ritual features make it possible to distinguish three groups of burials of different times [294] single burials in elongated rounded “nest” pits; 2) collective burials in square graves; 3) single burials in small shuttle-visible (in the form of a boat) pits. The buried were laid on their backs in an extended position, covered with ocher. Funeral equipment is extremely poorly preserved.

Despite the differences in the funeral ritual in anthropological terms, the series from the Volnensky burial ground turned out to be homogeneous. The skulls are exceptionally large, massive, with a strongly pronounced relief, with a long and wide cerebral part. The forehead is wide, sloping. The face is very tall and wide, orthognathic, strongly profiled in the horizontal plane, and canine fossae are deep. Orbits are low. The nose is high, strongly protruding. The lower jaw is very large and massive [295]. The unique skull features are estimated as proto-European type, having no analogies in the Neolithic of Western Europe and Western Asia.

Literature: Telegin 1957[291]; 1968[294]; Surnina 1961[295].

**Acknowledgments**

G.S. was supported by a Marie Skłodowska-Curie Individual Fellowship “PALAEO-ENEO”, a project funded by the European Union EU Framework Programme for Research and Innovation Horizon 2020 (Grant Agreement number 751349).

Long term research in Grotte Mandrin was made possible with the support of the Auvergne Rhône-Alpes Service Régional de l’Archéologie and the city of Malataverne (Drôme, France).

Research on the Ural River site (M.S.) conducted with the participation of ZIN RAS (grant № 075-15-2021-1069).

75. Rolfo, M. F., Achino, K. F. & Silvestri, L. Mora Cavorso Cave: a collective underground burial in Neolithic central Italy. in *Proceedings of the XVII World UISPP Congress* (eds.

T., T., M., D.-Z. B., A.m., S., C., C. & R, B.) 33–40 (Archaeopress Publishing Ltd, 2017).

76. Silvestri, L., Achino, K. F., Gatta, M., Rolfo, M. F. & Salari, L. Grotta Mora Cavorso: Physical, material and symbolic boundaries of life and death practices in a Neolithic cave of central Italy. *Quat. Int.* **539**, 29–38 (2020).

77. Merts, V. K. Pogrebeniye yamnogo tipa na Irtyshe (A Yamnaya-type burial on the Irtysh); in Altaye-Sayanskaya gornaya strana i sosedniye territorii v drevnosti (Altai-Sayan mountainous country and neighboring territories in antiquity. in *Altaye-Sayanskaya gornaya strana i sosedniye territorii v drevnosti (Altai-Sayan mountainous country and neighboring territories in antiquity)* 23–26 (Izdatel'stvo Instituta arkheologii i etnografii SO RAN (Publishing House of the Institute of Archeology and Ethnography SB RAS), 2007).

78. Merts, V. K. K probleme migratsiy v epokhu rannego metalla (o pogrebenii yamnogo tipa na Irtyshe) (To the problem of migrations in the era of the early metal (about the burial of the Yamnaya type on the Irtysh). in *Problemy arheologii: Ural i Zapadnaya Sibir' (k 70-letiyu T. M. Potemkinoy) (Problems of archeology: the Urals and Western Siberia (to the 70th anniversary of T.M. Potemkina))* (ed. Vokhmeytsev, M. P.) 71–75 (Izdatelstvo Kurganskogo universiteta, 2007b).

79. Merts, I. V. *Kul'tura naseleniya Vostochnogo Kazakhstana v epokhu ranney bronzy: avtoref. dis. … kan. ist. nauk: 07.00.06. (Culture of the population of East Kazakhstan in the Early Bronze Age: snyopsis of dissertation for candidate of historical sciences: 07.00.06.)*. 26 (Altay State University, 2017).

80. Damgaard, P. de B. The first horse herders and the impact of early Bronze Age steppe expansions into Asia. vol. 360 1422– Preprint at (2018).

81. Merts, V. K. & Merts, I. V. Pogrebeniya «yamnogo» tipa Vostochnogo i Severo-vostochnogo Kazakhstana (k postanovke problemy) (Burials of the 'Yamnaya' type of East and North-East Kazakhstan [to the problem statement]). in *Afanas'yevskiy sbornik (Afanasyevsky collection)* 134–144 (Izdatelstvo Azbuka, 2010).

82. Tur, S. S. & Solodovnikov, K. N. Kraniologicheskiye materialy yeluninskoy kul’tury epokhi ranney bronzy Verkhnego Priob'ya (Craniological materials of the Yelunin culture of the Early Bronze Age of the Upper Ob region). in *Pogrebal’nyy obryad naseleniya epokhi ranney bronzy Verkhnego Priob'ya (po materialam gruntovogo mogil'nika Teleutskiy Vzvoz-I) (Funeral practices of the population of the Early Bronze Age of the Upper Ob region [based on the materials of the Teleutsky Vzvoz-I burial ground])* (eds. Kiryushin, Y. F., Grushin, S. P. & Tishkin, A. A.) 142–176 (Altay State University, 2003).
83. Khokhlov, A. A., Solodovnikov, K. N., Rykun, M. P., Kravchenko, G. G. & Kitov, Y. P. Kraniologicheskiye dannyye k probleme svyazi populyatsiy yamnoy i afanas’yevskoy kul'tur Yevrazii nachal'nogo etapa bronzovogo veka (Craniological data on the problem of the relationship between the populations of the Yamnaya and Afanasievsk cultures of Eurasia in the initial stage of the Bronze Age). *Vestnik arkheologii, antropologii i etnografii (Bulletin of archeology, anthropology and ethnography)* **3**, 86–106 (2016).
84. Merts, V. K. Arkheologicheskoye izucheniye Severo-Vostochnogo Kazakhstana za 20 let nezavisimosti (Archaeological studies of North-East Kazakhstan during 20 years of independence). in *Svideteli tysyachiletiy: arkheologicheskaya nauka Kazakhstana za 20 let (1991-2011): sbornik nauchnykh statey (Witnesses of millennia: archaeological science of Kazakhstan for 20 years (1991-2011): collection of scientific articles)* 386–407 (2011).
85. Svyatko, S. V., Mertz, I. V. & Reimer, P. J. Reservoir Effect on Re-Dating of Eurasian Steppe Cultures: First Results for Eneolithic and Early Bronze Age North-East Kazakhstan. *Radiocarbon* **57**, 625–644 (2015).
86. Merts, I. V. & Svyatko, S. V. Radiouglerodnaya khronologiya pamyatnikov rannego bronzovogo veka Severo-Vostochnogo i Vostochnogo Kazakhstana. Pervyy opyt (Radiocarbon chronology of the sites of the Early Bronze Age in North-Eastern and Eastern Kazakhstan. *First experience); in Teoriya i praktika arkheologicheskikh issledovaniy (Theory and practice of archaeological research* **1**, 126–150 (2016).

87. Merts, K. V. K izucheniyu pamyatnikov epokhi ranney bronzy Kazakhstana // (The study of the monuments of the Early Bronze Age in Kazakhstan.). in *Arkheologicheskiye issledovaniya v Kazakhstane (Archaeological research in Kazakhstan)* 34–41 (2002).
88. Merts, V. K. O novykh pamyatnikakh epokhi ranney bronzy Kazakhstana (About new monuments of the Early Bronze Age in Kazakhstan). in *Istoricheskiy opyt khozyaystvennogo i kul'turnogo osvoyeniya Zapadnoy Sibiri: Chetvertyye nauch. chteniya pamyati prof. A. P. Borodavkina (Historical experience of economic and cultural development of Western Siberia: Fourth scientific readings in the memory of prof. A.P. Borodavkin)* 132–141 (Altay State University, 2003).
89. Merts, I. V. Vliyaniye vostochnoyevropeyskikh kul'turnykh obrazovaniy na formirovaniye naseleniya epokhi ranney bronzy Vostochnogo Kazakhstana (Influences of eastern European cultural entities on the formation of the population of the Early Bronze Age in East Kazakhstan); in XXI Ural'skoye arkheologicheskoye soveshchaniye, posvyashchennoye 85-letiyu so dnya rozhdeniya G.I. Matveyevoy i 70-letiyu so dnya rozhdeniya I. in *B. Vasil'yeva. Materialy Vserossiyskoy nauchnoy konferentsii s mezhdunarodnym uchastiyem (XXI Ural archaeological meeting dedicated to the 85th anniversary of G.I. Matveeva and the 70th anniversary of the birth of I.B. Vasiliev. Materials of the All-Russian scientific conference with international participation* 138–140 (SGSPU, 2018).
90. Logvin, A. V. & Shevnina, I. V. Stone-age burials of the Bestamak burial ground. in *Problems of studying the cultures of the early Bronze Age of the steppe zone of Eastern Europe* 142–151 (2009).
91. Logvin, A. V. & Ševnina, I. V. Die nekropole von Bestamak. in *Unbekannes Kasachstan Archaologie im Herzen Asiens* 231–244 (2013).
92. Logvin, A. V., Shevnina, I. V. & Kolbina, A. V. About burials of the Eneolithic in the 'sitting' position from the Bestamak burial ground. in *Problems of the archaeological study of the Southern Urals* 104–110 (2009).

93. Zagorskis, F. & Zagorska, I. Raskopki mezoliticheskoy stoyanki Zveyniyeki II (Excavations of the Mesolithic site Zvejnieki II). *Arkheologicheskiye otkrytiya 1972g (Archaeological discoveries 1972)* (1973).
94. Larsson, L., Stutz, L. N., Zagorska, I., Berzins, V. & Cerina, A. New aspects of the Mesolithic-Neolithic cemeteries and settlement at Zvejnieki, northern Latvia. *Acta Archaeol.* **88**, 57–93 (2017).
95. Denisova, R. Y. Antropologicheskiy sostav i genezis mezoliticheskogo naseleniya Latvii (Anthropological composition and genesis of the Mesolithic population of Latvia). *Sovetskaya etnografiya (Soviet Ethnography)* **1**, (1973).
96. Eriksson, G. *Norm and difference. Stone Age dietary practice in the Baltic region*. (Archaeological Research Laboratory, Stockholm University, 2003).
97. Jones, E. R. *et al.* The Neolithic Transition in the Baltic Was Not Driven by Admixture with Early European Farmers. *Curr. Biol.* **27**, 576–582 (2017).
98. Mathieson, I. *et al.* The genomic history of southeastern Europe. *Nature* **555**, 197–203 (2018).
99. Zagorskis, F. *Zvejnieki (Northern Latvia) Stone Age Cemetery (Translated by V. Bērziņś)*. vol. 1292 (Archaeopress, 2004).
100. Sellevold, B. J. & Skar, B. The First Lady of Norway. in *NIKU 1994-1999 Kulturminneforskningens Mangfold, NIKU Temahefte* (eds. Gundhus, G., Seip, E. & Ulriksen, E.) 6–11 (1999).
101. Eggen, I. & Nymoen, P. *Funnsted for mesolittiske skjeletter Hummervikholmen av Hallandvik, 32/69, Søgne kommune, Vest-Agder*. (Kulturhistorisk Museum, 2014).
102. Skar, B., Lidén, K., Eriksson, G. & Sellevold, B. J. A Submerged Mesolithic Grave Site Reveals Remains of the First Norwegian Seal Hunters. in *Marine Ventures: Archaeological Perspectives on Human-Sea Relations* (eds. Bjerck, H. B. et al.) 225–239 (Equinox Publishing, 2016).
103. Günther, T. *et al.* Population genomics of Mesolithic Scandinavia: Investigating early

postglacial migration routes and high-latitude adaptation. *PLoS Biol.* **16**, e2003703 (2018).

104. Tunia, K. *Temenos" kultury pucharów lejkowatych w Słonowicach, pow. Kazimierza Wielka. Badania 1979-2002. Trzecie sprawozdanie*. 335–340 (Instytut Archeologii UMCS, Instytut Archeologii i Etnologii PAN, 2006).
105. Przybyła, M. M. & Tunia, K. Investigations in 2012 of the southern part of the Funnel Beaker culture temenos at Słonowice near the Małoszówka river. Fourth report. *Environment and subsistence--forty years after Janusz Kruk's "Settlement studies* 139–162 (2013).
106. Jackes, M. & Lubell, D. The Early Neolithic human remains from Gruta do Caldeirão. Gruta do Caldeirão. in *Instituto Português do Património Arquitectónico e Arqueológico* (ed. Lisboa, O. N. A. J. Z.) 259–295 (1992).
107. Zilhão, J. *Gruta do Caldeirão. O Neolítico Antigo*. (Instituto Português do Património Arquitectónico e Arqueológico, 1992).
108. Zilhão, J. The Spread of Agro-Pastoral Economies across Mediterranean Europe: A View from the Far West. *Journal of Mediterranean Archaeology* **6**, 5–63 (1993).
109. Lubell, D., Jackes, M., Schwarcz, H., Knyf, M. & Meiklejohn, C. The Mesolithic-Neolithic Transition in Portugal: Isotopic and Dental Evidence of Diet. *J. Archaeol. Sci.* **21**, 201–216 (1994).
110. Chandler, H., Sykes, B. & Zilhão, J. Using ancient DNA to examine genetic continuity at the Mesolithic-Neolithic transition in Portugal. in *Actas del III Congreso del Neolítico en la Península Ibérica* (eds. P. Arias, R. O. & García-Moncó., C.) 781–786 (2005).
111. Isern, N., Zilhão, J., Fort, J. & Ammerman, A. J. Modeling the role of voyaging in the coastal spread of the Early Neolithic in the West Mediterranean. *Proc. Natl. Acad. Sci. U. S. A.* **114**, 897–902 (2017).
112. Silva, A. M. Antropologia funerária e Paleobiologia das populações Portuguesas (Litorais) do Neolítico final/Calcolítico. Dissertação de Doutoramento em Antropologia

Biológica. (FCTUC. Universidade de Coimbra, 2002).

113. Silva, A. M. Antropologia Funerária e Paleobiologia das Populações Portuguesas (litorais) do Neolítico final/Calcolítico. in *Textos Universitários de Ciências Sociais e Humanas. Lisboa, Fundação Calouste Gulbenkian, Fundação para a Ciência e Tecnologia* (2012).

120. Mogoşanu, F. *Paleoliticul din Banat*. (Editura Academiei Republicii Socialiste Romania, 1978).

127. Maksimenkov, G. A. Mogil'nik u Dogogo ozera v g. Kanske (Burial ground at Long Lake in Kansk). *Voprosy antropologii (Questions of anthropology)* (1964).

128. Debets, G. *Paleoantropologiya SSSR*. (1948).

129. Alekseev, V. P. & Gokhman, I. I. Anthropology of the Asian part of the USSR. Preprint at (1984).

130. Gerasimova, M. M. Skulls of the Fofonovsky burial ground. *Antiquities of Lake Baikal. Irkutsk* (1992).

131. Vasilyev, S. V., Gerasimova, M. M., Borutskaya, S. B., Frisen, S. Y. & Zhambaltarova, E. D. Anthropological study of the Fofonovsky burial ground (Transbaikalia) of the Neolithic and Early Bronze Age. *Herald of Anthropology* **18**, 113–127 (2010).

132. Berezutsky, V. D., Kileinikov, V. V. & Skorobogatov, A. M. *Mariupol-type burial of the Middle Don. Archaeological site of Eastern Europe: interuniversity collection of scientific papers*. 76–88 (Voronezh State Pedagogical University, 2011).

133. Kiryushin, Y. F. *Eneolithic and Early Bronze Age of the southern part of West Siberia. (Eneolit i rannyaya bronza yuga Zapadnoy Sibiri)*. (Barnaul University, 2002).

134. Dremov, V. A. The anthropological materials from the Ust'-Isha and Itkul' burial grounds.

in *Paleoantropologia Sibiri (Pslrosnthropology of Siberia* (eds. Okladnikov, A. P. & Alexeev, V. P.) 19–46 (Nauka, 1980).

135. Kiryushin, Y. F., Kungurova, N. Y. & Kadikov, B. K. The Ancient burial places of the Northern Altai foothills (Drevneyshie mogil'niki severnykh predgoriy Altaia). Preprint at (2000).
136. Bryusov, A. Y. The Karavaikha site. in *The archeology of the Vologda region* (ed. Bryusov, A. Y.) 72–95 (1961).
137. Utkin, A. V. & Kostyleva, E. L. Burials in the site of Karavaikha. *Russian archaeology (Russkaya arkheologiya)* 55–66 (2001).
138. Akimova, M. S. New paleoanthropological finds of the Neolithic era in the forest zone of the European part of the USSR. *Brief Communications of the Institute of Ethnography of the USSR (Kratkiye soobshcheniya Instituta Etnografii SSSR)* **18**, 55–65 (1953).
139. Gerasimov, M. M. Facial restoration through the skull: a modern and fossilized person. *Proceedings of the Institute of Ethnography (new series). Trudy Instituta Etnografii (novaya seriya)* **28**, 558 (1955).
140. Serikov, Y. B. *Cave sanctuaries of the Chusovaya River*. (Nizhnetagilskaya gosudarstvennaya socialno-pedagogicheskaya akademiya, 2004).
141. Serikov, Y. B. An unusual collective burial of the epoch Neolithic in the Kumyshan cave (Middle Urals). *Vestnik of Archeology, Anthropology and Ethnography* **2**, 4–10 (2013).
142. Lavrushin, Y., Spiridonova, E. A., Bessudnov, A. N. & R.v. *Smolyaninov Natural disasters in the Holocene of the Upper Don basin*. (GEOS Publishing House, 2009).
143. Vasilyev, S. V., Smolyaninov, R. V., Borutskaya, S. B. & Bessudnov, A. N. Population of the Neolithic-Eneolithic of the Upper Don and its funeral rituals (based on the materials of the Ksizovo 6 burial ground). Late prehistory of Eurasia: social models and cult practice. *Stratum* 167–195 (2018).
144. En'shin, D. N. Neoliticheskie zhilishcha poselenii ozera Mergen (The Neolithic dwellings of the settlement of the Lake Mergen). *Vestnik arkheologii, antropologii i etnografi*

*(Bulletin of Archaeology, Anthropology and Ethnography)* **1**, 14–23 (2014).

145. Piezonka, H. *et al.* The emergence of hunter-gatherer pottery in the Urals and West Siberia: New dating and stable isotope evidence. *J. Archaeol. Sci.* **116**, 105100 (2020).
146. Narasimhan, V. M. *et al.* The formation of human populations in South and Central Asia. *Science* **365**, (2019).
147. En'shin, D. N., Skochina, S. N. & Zakh, V. A. K voprosu o poselencheskoi obryadnosti v neolite Nizhnego Priishim'ya (po materialam poseleniya Mergen' 6) [On the issue of settlement rituals in the Neolithic of the Lower Ishim River region (based on the materials of the settlement of Mergen 6)]. *Vestnik arkheologii, antropologii i etnografii (Bulletin of Archaeology, Anthropology and Ethnography)* **4**, 43–52 (2012).
148. Makarov, N. A. Six years of excavation of the Minino archaeological complex. in *Kubenskoye Lake: a look through millennia* (ed. Makarov, N. A.) 3–6 (2001).
149. Suvorov, A. v. & Buzhilova, A. p. Unusual Stone Age funerary complexes of the site Minino on Lake Kubenskoe. *OPUS: Interdisciplinary research in archaeology* 41–54 (2004).
150. Suvorov, A. v. Methods of handling the bodies of the dead in the Stone Age burials of the sites of Minino I and II on Lake Kubenskoe. in *Russian Culture of the New Century: Problems of Studying, Preserving and Using the Historical and Cultural Heritage* (ed. Sudakov, G. V.) 21–26 (Book Heritage, 2007).
151. Wood, R. E. *et al.* Freshwater radiocarbon reservoir effects at the burial ground of Minino, Northwest Russia. *Radiocarbon* **55**, 163–177 (2013).
152. Buzhilova, A. P. A reconstruction of the lifestyle of early humans by natural-science methods. *Her. Russ. Acad. Sci.* **86**, 297–304 (2016).
153. Wood, R. *Chronometric and paleodietary studies at the Mesolithic and Neolithic burial ground of Minino, NW Russia: Dissertation for the MSc in archaeological Science*. (Oxford University, 2006).
154. Wood, R. E. *et al.* Freshwater radiocarbon reservoir effects at the burial ground of

Minino, northwest Russia. *Radiocarbon* **55**, 163–177 (2013).

155. Matyushchenko, V. I. & Polevodov, A. V. *The complex of the archaeological sites at the Tatar Uval near the Okunevo Village (Komplex arkheologicheskih pamyatnikov na Tatarskom uvale u derevni Okunevo)*. (Nauka, 1994).
156. Petrov, A. I. *The Neolithic and Early Bronze Age in the Middle Irtysh region (Epokha pozdnego neolita i ranney bronzy v Srednem Priirtysh'e)*. (Omsk University, 2014).
157. Metel', I. V. The history of the study of the complex of the archaeological sites the 'Omsk settlement'. *'Magistra Vitae': an electronic journal on historical sciences and archeology* 134–143 (2016).
158. Konnikov, B. A. *The complex of sites 'Omsk settlements': discoveries and findings. (Komplex pamyatnikov 'Omskaya stoyanka': otkrytiya i nakhotki)*. (Omsk State Pedagogical University, 1998).
159. *The Omsk settlement. The Album (Omskaya stoyanka. Al'bom)*. (2013).
160. Matveev, A. V., Matveeva, N. P., Serikov, Y. B. & Skochina, S. N. *The ritual Eolithic sites (Kul'tovye pamyatniki epokhi eneolita)*. (Tumen University, 2015).
161. Konstantinov, M. V., Ekimova, L. V. & Gantimurova, M. I. Neolithic burials in the vicinity of Jindo. Sources, formation and development of Eurasian multiculturalism. Cultures and Societies of North Asia in the Historical Past and Present. in *Materials of the I (XLV) Russian with international participation of the Archaeological and Ethnographic Conference of Students and Young Scientists (RAESK-XLV): April 12-16 2005* 129 (Publishing House of the Russian Orthodox Church 'Radian', 2005).
162. Oshibkina, S. v. Spree I. Settlement of the Mesolithic in the North of Eastern Europe. Preprint at (1997).
163. Oshibkina, S. V. Archaeological characteristics of the Mesolithic burial grounds of the European North of Russia. *VKA* (1998).
164. Oshibkina, S. V. *Art from Mesolithic period based on materials from Veretje culture*. (2017).

165. Mamonova, N. N. Paleoanthropological materials of the Mesolithic of the northern regions of Russia (preliminary report. *VA* (1995).
166. Bryusov, A. Y. Pile settlement on the Modlona River and other sites in the Charozersky district of the Vologda region. in *Materials of the Institute of Archaeology* (ed. Bryusov, A. Y.) vol. 20 7–76 (1951).
167. Kosorukova, N. v. Peat Mesolithic site Pogostishche XIV in the basin of Lake Vozhe (based on research 2005, 2008, 2009. in *History and Archeology of the Russian North* 58–63 (2012).
168. Oshibkina, S. v. *Mesolithic of the region of Eastern Onega*. 322 (2006).
169. Oshibkina, S. v. *Mesolithic of the region of Sukhona basin and Eastern Onega*. 296 (1983).
170. Makarov, N. a., Zakharov, S. d. & Buzhilova, A. p. *Medieval settlement in Beloozero region*. 496 (Languages of Russian culture, 2001).
171. Troitskaya, T. N. & Novikov, A. V. *Arkheologiya Zapadno-Sibirskoy ravniny: Uchebnoye posobiye (Archeology of the West Siberian Plain: Textbook)*. 136 (2004).
172. Kostyleva, E. L. & Utkin, A. V. *Neo-Eneolithic burial grounds of the Upper Volga and Volga-Oka interfluves: Planographic and chronological structures*. 300 (Publishing house TAUS, 2010).
173. Kostyleva, E. L. & Macane, A. Ural and Volga-Oka interfluves in the Eneolithic: connecting threads (Based on the materials of the monuments of the Sakhtysh paleo lake). in *XXI Ural Archaeological Conference. Materials of the All-Russian scientific conference with international participation* (Publishing House of SGSPU, 2018).
174. Kostyleva, E. L., Utkin, A. V. & Macane, A. ‘Eastern traces’ in the Eneolithic burials of the Sakhtysh burial grounds. *Tver archaeological collection* 565–577 (2018).
175. Piezonka, H., Kostyleva, E., Zhilin, M. G., Dobrovolskaya, M. & Terberger, T. Flesh or fish? First results of archaeometric research of prehistoric burials from Sakhtysh IIa, Upper Volga region, Russia. *Documenta Praehistorica* **40**, 57–73 (2013).

176. Macāne, A., Nordqvist, K. & Kostyleva, E. Marmot incisors and bear tooth pendants in Volosovo hunter-gatherer burials. New radiocarbon and stable isotope data from the Sakhtysh complex, Upper-Volga region. *Journal of Archaeological Science: Reports* **26**, 101908 (2020).
177. Krajnov, D. A., Kostyleva, E. L. & Utkin, A. V. Graveyard and 'sanctuary' on the settlement Sakhtysh IIA. *Rossiyskaya arkheologiya* 118–130 (1994).
178. Alekseeva, T. I. *et al.* Neolit lesnoy polosy Vostochnoy Yevropy (Antropologiya Sakhtyshskikh stoyanok) [Neolithic forest belt of East Europe (Anthropology of the Sakhtysh sites)]. *Nauchnyy mir* 191 (1997).
179. Engovatova, A. V., Dobrovol'skaya, M. V. & Kostyleva, E. L. Izotopnye harakteristiki individov iz pogrebenij neoliticheskogo mogil'nika Sahtysh-2A: dannye k rekonstrukcii pitaniya. in *Arheologiya Podmoskov'ya. Materialy nauchnogo seminara* (ed. Engovatova, A. V.) 138–145 (2015).
180. Крайнов, Д. А. *Древнеишая история Волго-Окского междуречия: Фатяновская культура II тысячилетия до н. е*. (Наука, 1972).
181. Krenke, N. A. Radiouglerodnaya hronologiya fat'yanovskoj kul'tury. *Rossijskaya arheologiya* 110–116 (2019).
182. Nordqvist, K. & Heyd, V. The Forgotten Child of the Wider Corded Ware Family: Russian Fatyanovo Culture in Context. *Proceedings of the Prehistoric Society* **86**, 65–93 (2020).
183. Utkin, A. V. & Kostyleva, E. L. Eshchyo raz o proiskhozhdenii l'yalovskoj kul'tury. *Verhnedonskoj arheologicheskij sbornik* **11**, 314–322 (2019).
184. Okladnikov, A. P. *Neliticheskie pamyatniki Nizhney Angary (Neolithic Sites of Lower Angara)*. 327 (Nauka Publ, 1976).
185. Drozdov, N. I. & Privalikhin, V. I. Arkheologicheskie razvedki na Sredney Angare (Archaeological Survey on the Middle Angara Region). in *Arkheologicheskie otkrytiya 1974 goda (Archaeological Discoveries of 1974)* 80–94 (2003).

186. Saveliev, N. A., Timoshchenko, A. A. & Badmaev, D. A. Spasatel'nye raboty Pyatogo Ust'-Ilimskogo otryada na pamyatnike Sosnovyi Mys v 2011 godu (Rescue archaeological work of the Fifth Ust-Ilim Detachment at the Sosnovy Mys site in 2011). in *Problems of Archaeology, Ethnography, Anthropology of Siberia and Neighboring Territories* vol. XVII 457–463 (IAET SB RAS Publ T., 2011).

187. Timoshchenko, A. A. Spasatel'nye raboty na stoyanke, mogil'nike Sosnovyi Mys v zone zatopleniya Boguchanskoy GES (Rescue archaeological work on the Sosnovyi Mys burial ground in the flood zone of the Boguchany Dam). in *Arkheologiya, etnologiya I antropologiya Evrazii: issledovaniya i gipotezy (materialy dokladov) (Archeology, ethnology and anthropology of Eurasia: research and hypotheses (reports))* 110–11 (2012).

188. Sviridov, A. A. *Report on the excavations of the settlement Vasilyevsky Kordon 17 in the Dobrovsky district of the Lipetsk region in 2012*. (2013).

189. Smolyaninov, R. V. *Report on the excavations of the settlement Vasilyevsky Kordon 17 in the Dobrovsky district of the Lipetsk region in 2013, 2014, 2015*. (2016).

190. Molodin, V. I., Mylnikova, L. N. & Nesterova, M. S. The Vengerovo-2A Neolithic Cemetery, Southwestern Siberia: Results of a Multidisciplinary Study. *Archaeology, Ethnology & Anthropology of Eurasia* **44**, 30–46 (2016).

191. Chikisheva, T. A., Pozdnyakov, D. V. & Zubova, A. V. Kraniologicheskiye osobennosti paleopopulyatsii neoliticheskogo mogilnika Vengerovo-2A v Barabinskoi lesostepi (Craniological features of the paleopopulation of the Neolithic burial ground Vengerovo-2A in the Barabinsk forest-steppe). *Teoriya i praktika arkheologicheskih issledovaniy (Theory and practice of archaeological research)* **2**, 144–162 (2015).

192. Lozovski, V., Lozovskaya, O., Mazurkevich, A., Hookk, D. & Kolosova, M. Late Mesolithic–Early Neolithic human adaptation to environmental changes at an ancient lake shore: The multi-layer Zamostje 2 site, Dubna River floodplain, Central Russia. *Quat. Int.* **324**, 146–161 (2014).

193. *Zamostje 2. Lake Settlement of the Mesolithic and Neolithic Fisherman in Upper Volga Region*. (IHMC RAS, 2013).

194. Site Zamostje 2 and landscape evolution in the Volga-Oka region during the Holocene. *IHMC RAS* (2018) doi:10.31600/978-5-9909872-8-9-2018.

195. Zubova, A. V., Moiseev, V. G. & Kulkov, A. M. Mesolithic Human Teeth from Zamostye-2, Moscow Region. *Archaeology, Ethnology & Anthropology of Eurasia* **47**, 120–127 (2019).

196. Meadows, J., Lozovskaya, O., Bondetti, M., Drucker, D. G. & Moiseyev, V. Human palaeodiet at Zamostje 2, central Russia: Results of radiocarbon and stable isotope analyses. *Quat. Int.* **541**, 89–103 (2020).

197. Lee, D. Banks Chambered Tomb, South Ronaldsay, Orkney, Discovery and Excavation in Scotland. *New* **12**, (2011).

198. Lee, D. *Banks Chambered tomb, South Ronaldsay*. (Internal Data Structure Report, 2011).

199. Srejović, D. & Letica, Z. *Vlasac: Mezolitsko naselje u Djerdapu: arheologija, tom 1*. (Srpska akademija nauka i umetnosti, 1978).

200. Borić, D., French, C. A. I. & Dimitrijević, V. Vlasac Revisited: Formation Processes, Stratigraphy and Dating. *Documenta Praehistorica* **35**, 293–320 (2008).

201. Borić, D., Raičević, J. & Stefanović, S. Mesolithic Cremations as Clements of Secondary Mortuary Rites at Vlasac (Serbia. *Documenta Praehistorica* **36**, 247–282 (2009).

202. Borić, D. *et al.* Late Mesolithic lifeways and deathways at Vlasac (Serbia). *J. Field Archaeol.* **39**, 4–31 (2014).

203. Borić, D. & Griffiths, S. The Living and the Dead, Memory and Transition: Bayesian Modelling of Mesolithic and Neolithic Deposits from Vlasac, the Danube Gorges. *Oxford Journal of Archaeology* **34**, 343–364 (2015).

204. Borić, D. *Deathways at Lepenski Vir: Patterns in Mortuary Practice*. (Serbian

Archaeological Society, 2016).

205. Borić, D. Lepenski Vir chronology and stratigraphy revisited. *Starinar* 9–60 (2019).

206. Radovanovic, I. *The iron gates mesolithic*. (International Monographs in Prehistory Ann Arbor (Michigan), 1996).

207. Srejovic, D. *Europe's First Monumental Sculpture: New Discoveries At Lepenski Vir*. (Thames and Hudson, 1972).

208. Borić, D. *et al.* High-Resolution AMS Dating of Architecture, Boulder Artworks and the Transition to Farming at Lepenski Vir. *Sci. Rep.* **8**, 14221 (2018).

209. Liesau, C. *et al.* Un espacio compartido por vivos y muertos: El poblado calcolítico de fosos de Camino de las Yeseras (San Fernando de Henares, Madrid)". *Complutum* **19**, 97–120 (2008).

210. Vega, J. *et al.* El recinto de fosos de Camino de las Yeseras (San Fernando de Henares, Madrid). Actas de las cuartas Jornadas de Patrimonio arqueológico en la Comunidad de Madrid. *Dirección General de Patrimonio Histórico* **Madrid**, 251–261 (2009).

211. Blasco, C., Rios, P. & Liesau, C. *Yacimientos calcolíticos con campaniforme de la región de Madrid: nuevos estudios*. (Universidad Autónoma de Madrid Madrid, 2011).

212. Ríos, P. Territorio y sociedad en la Región de Madrid durante el III milenio a.C. El referente del yacimiento de Camino de las Yeseras. *Patrimonio Arqueológico de Madrid* **7**, (2011).

213. Gómez, J. L. *et al.* Los protagonistas". *Nuevos estudios. Patrimonio Arqueológico de Madrid* **6**, 101–132 (2011).

214. Ríos, P., Liesau, C. & Blasco, C. Funerary practices in the ditched enclosures of Camino de las Yeseras: Ritual, Temporal and Spatial Diversity". *Recent Prehistoric Enclosures and Funerary Practices in Europe: Proceedings of the International Meeting held at the Gulbenkian Foundation* **2676**, 139–14 (2014).

215. Liesau, C. Fauna in Living and Funerary Contexts of the 3rd Millennium BC in Central

Iberia. in *Key resources and sociocultural developments in the Iberian Chalcolithic* (eds. Bartelheim, M., Bueno, P. & Kunst, M.) (2017).

216. Arteaga, C. *et al.* The ditched enclosure of Camino de las Yeseras (Madrid): a sedimentological approach to the study of some singular structures". *Espacio, Tiempo y Forma. Serie I Prehistoria y Arqueología* **10**, 77–94 (2017).
217. Noval Fonseca, M. Excavación arqueológica en la cueva de El Toral III (Andrín, Llanes. in *Excavaciones Arqueológicas en Asturias 2007-2012, Consejería de Cultura y Deporte del Gobierno del Principado de Asturias* 381–384 (2013).
218. Bello Alonso, P., Ozkorta Escribano, L. & Gutiérrez Zugasti, I. Un acercamiento al aprovechamiento de los recursos litorales durante el Mesolítico: los invertebrados marinos del abrigo de El Toral III. in *La Investigación Arqueomalacológica en la Península Ibérica: Nuevas Aportaciones* (eds. Llanes, A. I., Gutiérrez Zugasti, I., Cuenca Solana, D. & González Morales, M. R.) 91–99 (Nadir Ediciones, 2015).
219. Rigaud, S. & Gutiérrez-Zugasti, I. Symbolism among the last hunter-fisher-gatherers in northern Iberia: Personal ornaments from El Mazo and El Toral III Mesolithic shell midden sites. *Quat. Int.* **407**, 131–144 (2016).
220. Gutiérrez Zugasti, I. & González Morales, M. R. Intervención arqueológica en la cueva de El Mazo (Llanes, Asturias): campañas de 2009, 2010 y 2012. in *Excavaciones Arqueológicas en Asturias 2007-2012, Consejería de Cultura y Deporte del Gobierno del Principado de Asturias* 159–167 (2013).
221. Gutiérrez-Zugasti, I. *et al.* Back to the Asturian: first results from the Mesolithic shell midden site of El Mazo (Asturias, Northern Spain. in *Anciens peuplements littoraux et relations Homme/Milieu sur les côtes de l'Europe Atlantique / Ancient Maritime Communities and the Relationship between People and Environment along the European Atlantic Coasts, BAR International Series 2570* (eds. Daire, M. Y. et al.) 483–490 (2013).
222. Gutiérrez-Zugasti, I. *et al.* La ocupación de la costa durante el Mesolítico en el Oriente

de Asturias: primeros resultados de las excavaciones en la cueva de El Mazo (Andrín, Llanes. *Archaeofauna* **23**, 25–38 (2014).

223. García-Escárzaga, A., Gutiérrez-Zugasti, I. & González-Morales, M. R. Análisis arqueomalacológico de la unidad estratigráfica 108 del conchero mesolítico de El Mazo (Llanes, Asturias): conclusiones socio-económicas y metodológicas. in *La Investigación Arqueomalacológica en la Península Ibérica: Nuevas Aportaciones Nadir Ediciones* (eds. Gutiérrez-Zugasti, I., Cuenca-Solana, D. & González-Morales, M. R.) 77–89 (2015).
224. Gutiérrez-Zugasti, I. *et al.* Collection and consumption of echinoderms and crustaceans at the Mesolithic shell midden site of El Mazo (northern Iberia): Opportunistic behaviour or social strategy? *Quat. Int.* **407, Part B**, 118–130 (2016).
225. Gutiérrez Zugasti, I. *et al.* Intervención arqueológica en la cueva de El Mazo (Andrín, Llanes). Campañas de 2013, 2014, 2015 y 2016. in *Excavaciones Arqueológicas en Asturias 2013-2016* 133–142 (Consejería de Educación y Cultura Principado de Asturias, 2018).
226. García-Escárzaga, A. *et al.* Stable oxygen isotope analysis of Phorcus lineatus (da Costa, 1778) as a proxy for foraging seasonality during the Mesolithic in northern Iberia. *Archaeol. Anthropol. Sci.* **11**, 5631–5644 (2019).
227. Aura Tortosa, J. E. Coves de Santa Maira. in *Pleistocene and Holocene Hunter-Gatherers in Iberia and the Gibraltar Strait* (eds. Carbonell, E., Castro, J. M. & Arsuaga, J. L.) 353–356 (Fundación Atapuerca - Universidad de Burgos, 2014).
228. Carrión Marco, Y., Verdasco Cebrián, C. C., Morales Pérez, J. V. & Aura Tortosa, J. E. Au-delà du radiocarbone : analyse de taxons et contexte combinés pour la détection de problèmes taphonomiques. Un exemple dans les Grottes de Santa Maira (Alicante, Espagne. *ArcheoSciences, revue d'archéométrie* **42**, 35–43 (2018).
229. Morales-Pérez, J. V. *et al.* Funerary practices or food delicatessen? Human remains with anthropic marks from the Western Mediterranean Mesolithic. *Journal of*

*Anthropological Archaeology* **45**, 115–130 (2017).

230. Salazar-García, D. C. *et al.* Isotope evidence for the use of marine resources in the Eastern Iberian Mesolithic. *J. Archaeol. Sci.* **42**, 231–240 (2014).
231. Verdasco Cebrián, C. C. Estudio microsedimentológico de niveles arqueosedimentarios depositados en cuevas y abrigos en el País Valenciano durante el Pleistoceno-Holoceno (11.000-5.000 BP. (Universitat de València, 2016).
232. Borrman, H. *et al.* Dental conditions and temporomandibular joints in an early mesolithic bog man. *Swed. Dent. J.* **20**, 1–14 (1996).
233. Jonsson, L. & Gerdin, A.-L. Bredgårdsmannen. Rapport över arkeologisk undersökning på Bredgården 1:1, Marbäck sn, Västergötland. *UV Väst rapport* **1997**, (1997).
234. Magnell, O. *Osteologisk rapport över skelett från grav (A 1671), Dösemarken, Hyllie sn, förundersökning 2006*. (2006).
235. Ifverson, P. Dösemarken Fastigheten Limhamn 155:355 Hyllie socken i Malmö stad Skåne län, Arkeologisk förundersökning 2006. *Malmö Kulturmiljö Enheten för Arkeologi, Rapport* **2007**, (2007).
236. Niklasson, N. Ett bidrag till kännedomen om begravningsskicket under stenåldern. in *Bidrag till kännedomen om Göteborgs och Bohusläns fornminnen och historia* 211–223 (1932).
237. Johansson, E. Stenåldersbebyggelsen på Skaftö. *Bohusläns hembygdsförbunds årsskrift* 49–60 (1974).
238. Nordqvist, B. Coastal Adaptations in the Mesolithic. A study of coastal sites with organic remains from the Boreal and Atlantic periods in Western Sweden. *GOTARC Series B* (2000).
239. Lidén, K., Eriksson, G., Nordqvist, B., Götherström, A. & Bendixen, E. 'The wet and the wild followed by the dry and the tame' – or did they occur at the same time? Diet in Mesolithic – Neolithic southern Sweden. *Antiquity* **78**, 23–33 (2004).
240. Ahlström, T. & Sjögren, K.-G. Kvinnan från Österöd--ett tidigmesolitiskt skelett från

Bohuslän. *In Situ Archaeologica* **7**, (2007).

241. Sjögren, K.-G., Price, T. D. & Ahlström, T. Megaliths and mobility in south-western Sweden. Investigating relations between a local society and its neighbours using strontium isotopes. *Journal of Anthropological Archaeology* **28**, 85–101 (2009).
242. Weiler, E. Fornlämning 5, hällkista. *Raä och SHM Rapport UV* **1977**, (1977).
243. Blank, M., Tornberg, A. & Knipper, C. New Perspectives on the Late Neolithic of southwestern Sweden. An interdisciplinary investigation of the Gallery Grave Falköping stad 5. *Open Archaeology* **4**, 1–35 (2018).
244. Blank, M., Sjögren, K.-G. & Storå, J. Old bones or early graves? Megalithic burial sequences in southern Sweden based on 14C datings. *Archaeol. Anthropol. Sci.* **12**, (2020).
245. Ahlström, T. Underjordiska dödsriken – humanosteologiska undersökningar av neolitiska kollektivgravar. *Coast to coast books* (2009).
246. Skoglund, P. *et al.* Genomic diversity and admixture differs for Stone-Age Scandinavian foragers and farmers. *Science* **344**, 747–750 (2014).
247. Sjögren, K.-G. Fragment av ordning. Undersökning av överplöjda megalitgravar vid Frälsegården, Gökhems socken, Västergötland, 1999-2001. *GOTARC Serie* **D**, (2008).
248. Sjögren, K.-G. News from Frälsegården. Aspects on Neolithic burial practices. *Neolithic Diversities. Perspectives from a conference in Lund, Sweden, s 200-210. Acta Archaeologica Lundensia, Series in 8o* (2015).
249. Sjögren, K.-G. Modeling Middle Neolithic Funnel Beaker diet on Falbygden, Sweden. *Journal of Archaeological Science: Reports* **12**, 295–306 (2017).
250. Rascovan, N. *et al.* Emergence and Spread of Basal Lineages of Yersinia pestis during the Neolithic Decline. *Cell* **176**, 295–305.e10 (2019).
251. Vretemark, M. Kraniet från Hanaskede – vår äldste skaraborgare. *Västergötlands fornminnesförenings tidskrift* **1996**, 212–214 (1996).
252. Salomonsson, B. Hindby mosse-boplatsen. Malmö Stads historia första delen. Preprint

at (1971).

253. Jeppsson, E. Rapport arkeologisk undersökning av förhistoriskt boplatslager å Hindby mosse, stadsäga Malmö kommun, Skåne, Malmö Museers arkiv. Preprint at (1976).
254. Svensson, M. Trattbägarboplatsen ‘Hindby Mosse’ – aspekter på dess struktur och Funktion. *Elbogen. Malmö fornminnesförenings tidskrift. Årg* **16**, (1986).
255. Sandén, U., Brink, K., Högberg, A., Nilsson, L. & Skoglund, P. Fester och festande vid Hyllie – nya tolkningar av mellanneolitiska platser och bebyggelsemönster i sydvästra Skåne. *Fornvännen* **105**, (2010).
256. Nilsson, L. Hindbymosseboplatsen – En mångfacetterad rituell plats från yngre stenålder. *Osteologisk rapport* (2020).
257. Winge, G. Rapport över arkeologisk undersökning av kulturlager, gropar, härdar, grophus, stolphål och gravar vid Kastanjegården i Fosie och Lockarp. *Malmö* (1974).
258. Winge, G. Gravfältet vid Kastanjegården. vol. Malmöfynd 3 Preprint at (1976).
259. Rosborn, S. *Den skånska historien före skriftspråket*. (Malmö, 1999).
260. Niklasson, N. Torslanda i forntiden. in *Torslanda, en sockenskildring* 37–60 (1956).
261. Larsson, L. Skateholm. in *Ancient Europe 8000 B.C. to A.D. 1000. Encyclopedia of the Babarian World 1* (eds. Boguski, P. & Crabtree, P.) (2004).
262. Larsson, L. Some aspects of mortuary practices at the Late Mesolithic cemeteries at Skateholm, southernmost Sweden. *International Conference Halle (Saale), Germany, 18th–21st September 2013. Tagungen des Landesmuseums für Vorgeschichte Halle Band* **13**, 175–184 (2016).
263. Arcini, C. *Osteologisk undersökning av gravmaterialet från vattenledningen Malmö*. (Arkivmaterial, Malmö Museer, 2007).
264. Gidlöf, K. & Gruber, A. Arkeologisk slutundersökning 2005, Vattenledningen. Vattenöverföringsledning mellan Hyllie vattentorn och vattenverket i Vellinge. Hököpinge socken i Vellinge kommun, Skåne län. *Malmö Museer, Arkeologienheten, Rapport* **2009**, (2009).

265. Rudebeck, E. Ängdala. Flintgruvor från yngre stenåldern, S. Preprint at (1986).

266. Frejd, J. & Rudebeck, E. Pilbladet 1 (f.d. del av Södra Sallerup 180:36), Arkeologisk förundersökning 2012. *Rapport* **2013**, (2013).

267. Berggren, Å. Pilbladet 1, Tidigneolitiska gruvor. *Rapport* **2018**, (2018).

268. Bernbeck, R. & Pollock, S. Scalar differences: temporal rhythms and spatial patterns at Monjukli Depe, southern Turkmenistan. *Antiquity* **90**, 64–80 (2016).

269. Korobkova, G. F. Orudiya truda i khozyaystvo neoliticheskikh plemen Sredney Azii (Tools and economy of the Neolithic tribes of Central Asia). *Materialy i issledovaniya po arkheologii SSSR (Materialy i issledovaniya po arkheologii SSSR)* (1969).

270. Ginzburg, V. V. & Trofimova, T. A. *Paleoantropologiya Sredney Azii (Paleoanthropology of Central Asia)*. (1972).

271. Dobrovolsky, A. V. Eighth Igrenian Neolithic site. *Archaeology of the Dnepr* 243–252 (1949).

272. Telegin, D. Y. Middle Stog culture of the Copper Age. 168 Preprint at (1973).

273. Konduktorova, T. S. *Anthropology of the Ukrainian population of the Mesolithic, Neolithic and Bronze Ages*. 128 (Nauka, 1973).

274. Потехина, И. Д. Население Украины в эпохи неолита и раннего энеолита по антропологическим данным. *Киев: ИА НАНУ* (1999).

275. Manko, V. A. & Telizhenko, S. A. Early Neolithic site Kleshnya III on the Middle Donets (Ранненеолитическая стоянка Клешня 3 на Среднем Донце). in *Antiquities of the Seversky Donets (Древности Северского Донца)* vol. 4 5–34 (Shlyakh, 2000).

276. Manko, V. A. & Telizhenko, S. A. *Mesolithic, Neolithic, Eneolithic of the Donets Basin: catalogue of radiocarbon dates (Мезолит, неолит, энеолит Подонечья: каталог радиокарбонных дат)*. (Shlyakh, 2002).

277. Telegin, D. Y. & Potekhina, I. *Neolithic cemeteries and populations in the Dnieper Basin*. (British Archaeological Report S383, 1987).

278. Tuboltsev, O. V. Neolithic burials of the Mamaj Mountains. *Archaeological Bulletin* **3**,

(1992).

279. Toshchev, G. N. Die neolithische Nekropole Mamaj-Gora im unteren Dneprgebiet. **XXXIV**, (2005).

280. Andrukh, S. I. & Toshchev, G. N. *Могильник Мамай-Гора, книга 4 (Mamai-Gora burial ground, book 4)*. (Zaporizhzhya National University, 2009).

281. Konduktorova, T. S. Paleoanthropological materials from the Mesolithic cemetery Vasilyevka I. *Soviet anthropology (Sovetskaya antropologiya)* 189–210 (1957).

282. Stolyar, A. D. The Mesolithic burial ground of Vasilyevka on the Dnieper. *Soviet anthropology (Sovetskaya antropologiya)* 179 – 189 (1957).

283. Grünberg, J. M. *Mesolithische Bestattungen in Europa, Katalog*. (Leidorf (Rahden/Westf.), 2000).

284. Rudinsky, M. Y. Late Neolithic burial grounds of Vovnigi (on the issue of the design of Mariupol type burial grounds. **4**, 147–151 (1955).

285. Lillie, M. & Budd, C. The Mesolithic-neolithic transition in eastern Europe: Integrating stable isotope studies of diet with palaeopathology to identify subsistence strategies and economy. in *Human Bioarchaeology of the Transition to Agriculture* 43–62 (John Wiley & Sons, Ltd, 2011).

286. Konduktorova, T. S. Skulls from the Vovnigi Late Neolithic burial grounds. *Brief Communications of the Institute of Archeology of the Academy of Sciences of the Ukrainian SSR(Kratkiye soobshcheniya Instituta arkheologii AN Ukrainskoi SSR)* **6**, 68–71 (1956).

287. Gokhman, I. I. The population of Ukraine during the Mesolithic and Neolithic (anthropological essay). 195 Preprint at (1966).

288. Telegin, D. Y. & Potekhina, I. D. *Neolithization in Ukraine. British Archaeological Reports*. (Archaeopress, 2003).

294. Telegin, D. Y. *Dnieper-Donetsk culture*. (1968).

295. Surnina, T. S. Paleoanthropological material from the Volnensky Neolithic burial ground. *Trudy Instituta etnografii. Novaya seriya (Proceedings of the Institute of Ethnography. New series)* **71**, (1961).
